# Distinct cellular DNA methylation mechanisms underlie common and rare genetic risk for brain disorders

**DOI:** 10.64898/2026.03.09.710649

**Authors:** Jiyun Zhou, Cuining Liu, Xiaoming Liu, Yuan Zhang, Yu Wei, Joo Heon Shin, Brady Maher, Chunyu Liu, Chongyuan Luo, Kai Wang, Daniel R. Weinberger, Shizhong Han

## Abstract

Noncoding genetic variation contributes to brain disorder risk, but the mechanisms through which it acts in specific brain cell types remain unclear. DNA methylation (DNAm), a highly cell type-specific regulatory layer in the brain, may mediate noncoding genetic risk, yet whether methylation at CG (mCG) and neuron-enriched non-CG (mCH) dinucleotides contribute differently to that risk remains unknown. Here we develop a deep learning framework that predicts DNAm from DNA sequence and estimates variant effects across 186 brain cell subtypes in both mCG and mCH, leveraging single-nucleus DNAm profiles from 46 brain regions. The models reveal distinct transcription factor (TF) programs underlying the two methylation contexts, with mCH-associated TFs showing stronger evolutionary constraint. Predicted variant effects agree closely with cell type-matched mQTLs in both direction and magnitude. Common variants predicted to affect mCG, particularly in excitatory neurons, show substantially greater heritability enrichment for brain-related traits than variants affecting mCH. By contrast, noncoding *de novo* mutations in autism preferentially perturb mCH, but not mCG, at conserved neuronal regulatory regions. This pattern is replicated across two independent cohorts totaling 5,782 probands and 4,053 unaffected siblings. Together, these findings indicate that common and rare noncoding variants contribute to brain disorders through distinct DNA methylation mechanisms.

## Introduction

The noncoding genome harbors substantial genetic risk for brain disorders, yet the regulatory mechanisms linking noncoding variation to disease in specific brain cell types remain poorly understood^1–3^. DNA methylation (DNAm) is a highly cell type-specific epigenetic mark that regulates gene expression and is essential for brain development and function^4–10^, making it a strong candidate mediator of noncoding genetic risk. Crucially, genetic studies have identified widespread DNAm quantitative trait loci (mQTLs) across the genome, linking noncoding variation to inter-individual differences in DNAm^11–15^. However, most mQTL studies in the human brain have relied on bulk tissue, obscuring cell type-specific effects. Moreover, these studies have focused almost exclusively on methylation at CG dinucleotides (mCG), overlooking non-CG methylation (mCH), which accumulates in neurons during postnatal development and plays critical roles in neuronal maturation^6,16–19^.

Single-cell technologies enable DNAm profiling in individual cells^9,20,21^ and, in principle, allow mQTL detection at the cell-type level. However, the high cost of these approaches limits the sample sizes needed for robust mQTL discovery, particularly for rare variants. Deep learning models trained on DNA sequence offer a complementary strategy, as they can predict variant effects without requiring large population-based samples. Several models^22–25^, and most recently AlphaGenome^26^, have demonstrated remarkable performance in predicting variant effects on chromatin accessibility, gene expression, and splicing directly from DNA sequence. However, none of these models explicitly model DNAm, leaving this critical epigenetic layer largely unexplored. To address this gap, we previously developed INTERACT, a transformer-based deep learning framework that predicts variant effects on DNAm levels in bulk brain tissue^27^ and across 13 brain cell types from the dorsolateral prefrontal cortex^28^. However, INTERACT was restricted to the CG context and a limited number of cell types from a single brain region, leaving the vast cellular and methylation context diversity of the human brain unexplored.

Here, we develop a multi-task deep learning framework, trained on single-nucleus DNAm profiles from 46 brain regions^29^, to predict DNAm from DNA sequence and estimate variant effects across 186 brain cell subtypes in both mCG and mCH contexts. The models uncover distinct transcription factor (TF) programs underlying the two methylation contexts, with mCH-associated regulators showing stronger evolutionary constraint. We demonstrate that common variants associated with brain-related traits preferentially influence mCG in excitatory neurons, whereas noncoding *de novo* mutations in autism spectrum disorder (ASD) preferentially perturb mCH at conserved neuronal regulatory regions, a pattern replicated across the Simons Foundation Powering Autism Research for Knowledge (SPARK)^30^ and Simons Simplex Collection (SSC)^31^ cohorts. Together, these results indicate distinct cellular DNAm mechanisms through which common and rare noncoding variants contribute to brain disorders.

## Results

### A multi-task deep learning framework for predicting DNAm across brain cell types

We developed a multi-task deep learning framework that predicts DNAm levels at individual cytosines across multiple cell types from local DNA sequence (2 kb) (**Fig. 1**). The framework uses a two-stage hierarchical training strategy to jointly learn sequence features associated with DNAm across related cell types. In the first stage, we trained three independent multi-task models for broad cell classes—excitatory neurons (telencephalic excitatory), inhibitory neurons (inhibitory/non-telencephalic), and non-neuronal cells—each jointly predicting DNAm levels across cell types within that class. In the second stage, each broad-class model was fine-tuned for individual cell types to predict DNAm levels in their refined subtypes. To capture context-specific regulatory programs, we trained separate models for CG and the four most prevalent CH contexts in neurons: CAC, CAG, CAT and CTC.

**Figure 1.**
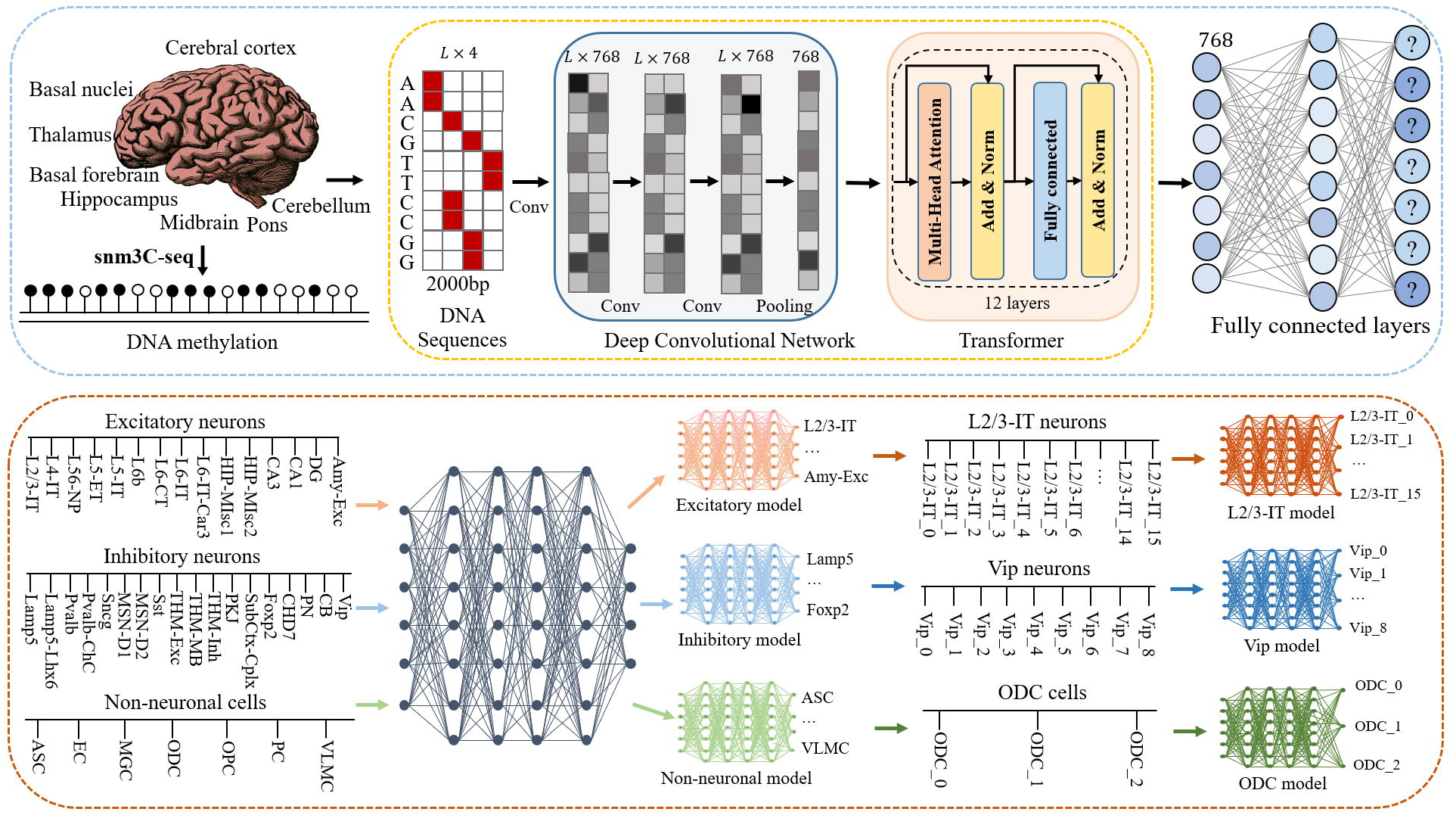
Schematic of the model architecture and two-step training strategy. Upper panel, model architecture. The model takes as input a one-hot-encoded 2-kb DNA sequence centered on the target cytosine and processes it through convolutional layers to extract sequence features, transformer encoder layers to capture long-range dependencies, and fully connected layers to predict DNAm levels at the central cytosine across multiple cell types. Separate models were trained for each methylation context (CG, CAC, CAG, CAT and CTC). **Lower panel**, two-step hierarchical training strategy consisting of pretraining and fine-tuning. **(i) Pretraining**, models were trained separately for broad cell classes—telencephalic excitatory neurons, inhibitory/non-telencephalic neurons, and non-neuronal cells (CG only). Each pretrained model jointly predicts DNAm levels across cell types within its class; for example, the excitatory model predicts across L2/3-IT, L4-IT, L5-IT, and other excitatory neuron cell types. **(ii) Fine-tuning**, each pretrained model was fine-tuned to jointly predict DNAm levels across cell subtypes within each cell type. Examples shown include fine-tuning the excitatory model on L2/3-IT subtypes, the inhibitory model on Vip subtypes, and the non-neuronal model on oligodendrocyte (ODC) subtypes.

We evaluated performance on held-out cytosines from chromosome 22 (**Supplementary Table 1**). Prediction accuracy varied across cell subtypes and methylation contexts, as assessed by mean squared error (MSE) (**Fig. 2A**). The CG context showed the highest average MSE across subtypes (mean MSE = 0.040), followed by CAC (0.031), CTC (0.013), CAG (0.011), and CAT (0.010). Within the CG context, excitatory neuronal subtypes had higher average MSE (0.047) than non-neuronal (0.037) and inhibitory neuronal subtypes (0.033). Variation in observed DNAm levels across held-out cytosines closely mirrored this pattern (**Fig. 2B**) and was strongly correlated with MSE across cell subtypes and contexts (Pearson’s r = 0.96; **Fig. 2C**).

**Figure 2.**
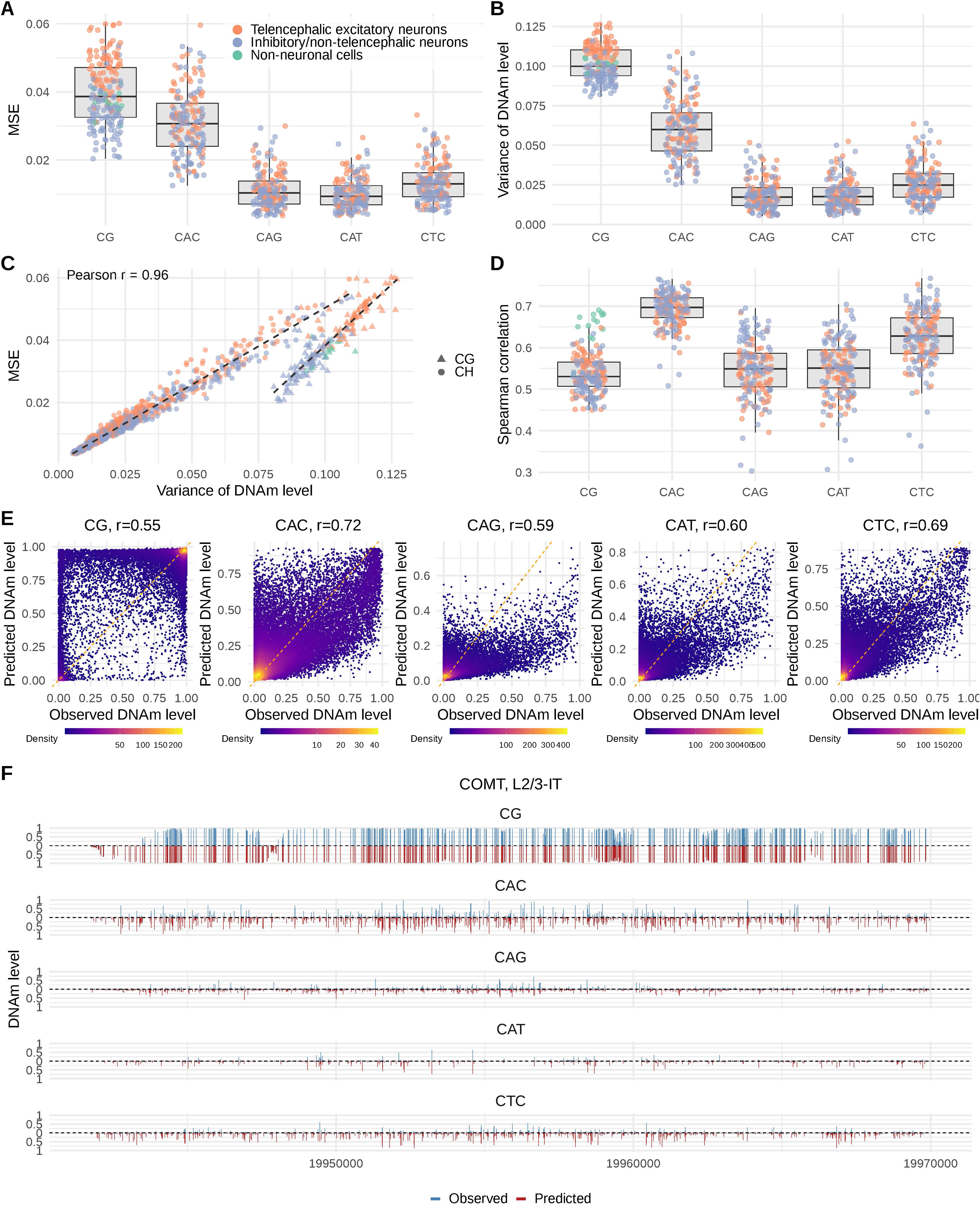
Model prediction performance. **(A)** Prediction accuracy across cell subtypes, measured by mean squared error (MSE) on held-out cytosines from chromosome 22. Results are shown for each methylation context (CG, CAC, CAG, CAT, CTC) and colored by broad cell class (excitatory, inhibitory, and non-neuronal cells). Each point represents one cell subtype. **(B)** Variance in observed DNAm levels across held-out cytosines from chromosome 22 for each cell subtype and methylation context. **(C)** Correlation between DNAm variance and prediction error (MSE) across all models. Each point represents one cell subtype–methylation context pair. Points are colored by broad cell class and shaped by methylation context (CG versus CH). **(D)** Prediction accuracy for each cell subtype, measured by the Spearman correlation between predicted and observed DNAm levels across held-out cytosines. **(E)** Example scatter plots comparing observed and predicted DNAm levels for 30,000 randomly selected cytosines from chromosome 22 in one L2/3-IT subtype. Each panel shows one methylation context (CG, CAC, CAG, CAT or CTC), with the Spearman correlation shown in each panel. **(F)** Example of observed (top) and predicted (bottom) DNAm levels at individual cytosine positions across the five methylation contexts at the *COMT* locus on chromosome 22 in the same L2/3-IT subtype shown in **E**.

We further assessed performance using the Spearman correlation between predicted and observed DNAm levels across held-out cytosines (**Fig. 2D**). CAC showed the highest average correlation across cell subtypes (mean Spearman’s r = 0.69), followed by CTC (r = 0.63), CAG (r = 0.55), CAT (r = 0.55), and CG (r = 0.54). As illustrative examples, **Figure 2E** shows the correspondence between observed and predicted DNAm levels across methylation contexts for one L2/3-IT neuronal subtype, and **Figure 2F** shows predictions at individual cytosine positions at the *COMT* locus.

### Model learns distinct TF programs across cell types and DNAm contexts

To gain biological insights from the trained models, we examined first-layer convolutional filters of models fine-tuned for 40 cell types to identify learned DNA motifs and mapped them to known TF binding motifs. The number of inferred TFs varied across cell types and methylation contexts (**Fig. 3A and Supplementary Table 2**). In the CG context, inhibitory neuron cell types showed the highest average number of TFs (N=57), followed by non-neuronal (N=52) and excitatory neuron cell types (N=41). By contrast, excitatory neuron cell types showed more TFs across CH contexts (CAC, N=74; CAG, N=85; CAT, N=110; CTC, N=80) than inhibitory neuron cell types (CAC, N=45; CAG, N=50; CAT, N=38; CTC, N=52), suggesting broader CH-associated regulatory programs in excitatory neurons. Hierarchical clustering of cell types based on learned TF profiles across CG and CH contexts largely recapitulated known cellular relationships, with excitatory neurons, inhibitory neurons, and non-neuronal cells forming distinct clusters (**Fig. 3B**), indicating that the models capture cell type-specific regulatory programs encoded in DNAm patterns.

**Figure 3.**
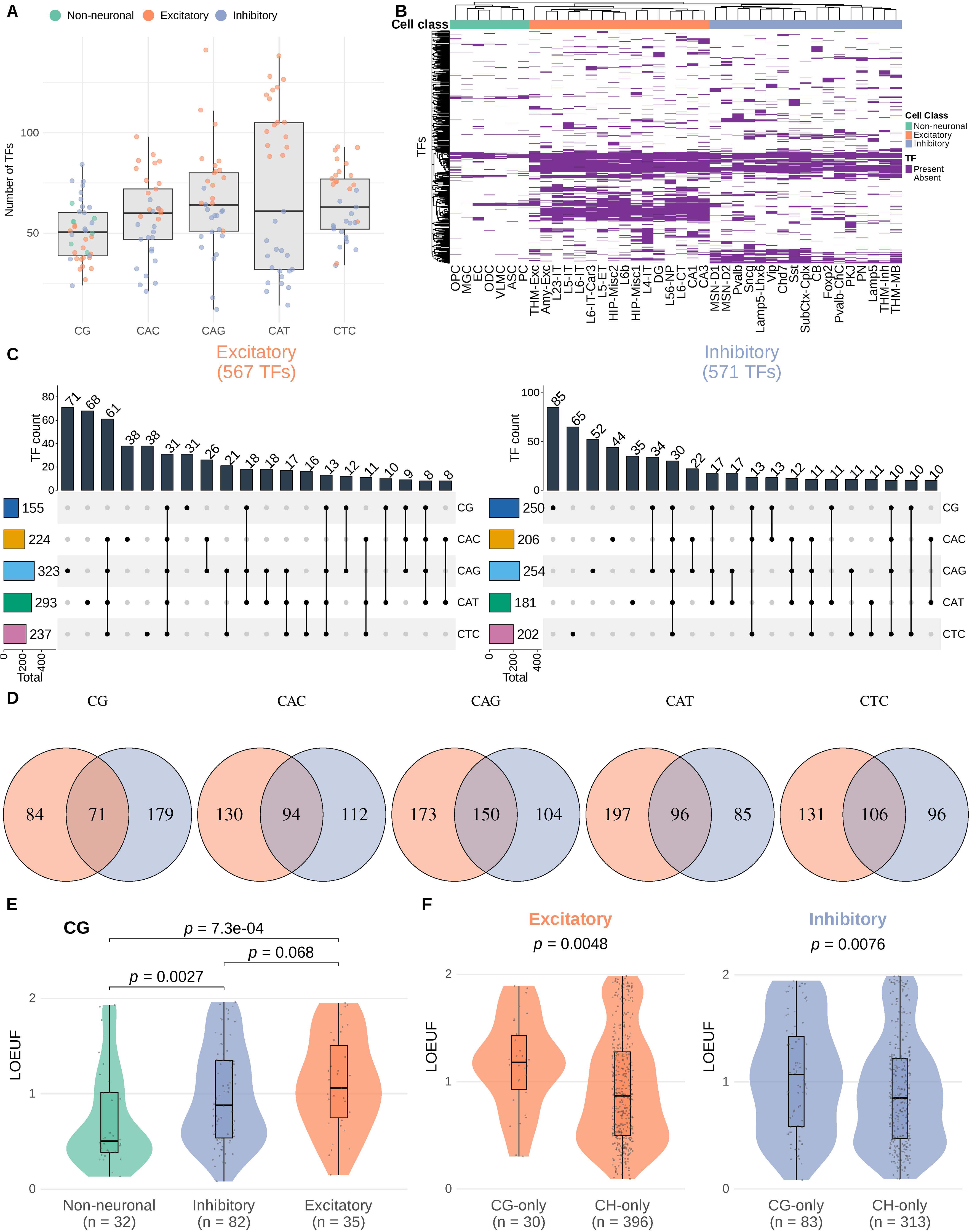
Transcription factor programs differ across methylation contexts and brain cell types. **(A)** Boxplot showing the number of transcription factor (TFs) detected per cell type in each methylation context (CG, CAC, CAG, CAT, CTC). Each dot represents one cell type, colored by broad cell class (Non-neuronal, Excitatory, Inhibitory). **(B)** Binary presence/absence heatmap of all TFs (rows) across cell types (columns), aggregated across all five methylation contexts. Rows and columns are hierarchically clustered using Jaccard distance (complete linkage). **(C)** UpSet plots showing the overlap of detected TFs across the five methylation contexts, separately for excitatory (left) and inhibitory (right) neurons. Top bar plots indicate the number of TFs in each intersection; left bar plots show the total number of TFs per context. Intersections are ordered by size (decreasing). **(D)** Venn diagrams comparing excitatory versus inhibitory TF sets within each methylation context. **(E)** Violin plots comparing LOEUF scores of CG-exclusive TFs across the three broad cell classes (Non-neuronal, Inhibitory, Excitatory). A TF is classified as exclusive to a class if it is detected in that class only (not in either of the other two) in the CG context. P-values are from one-sided Wilcoxon rank-sum tests. **(F)** Violin plots comparing LOEUF scores of CG-only versus CH-only TFs, separately for excitatory (left) and inhibitory (right) neuronal class. CG-only TFs are detected in CG but not in any CH context (CAC, CAG, CAT, CTC); CH-only TFs are detected in at least one CH context but not in CG. P-values are from one-sided Wilcoxon rank-sum tests (CH-only < CG-only).

We next examined how inferred TFs were distributed across methylation contexts within each broad neuronal class (**Fig. 3C**). In excitatory neurons (567 unique TFs), the largest context-specific groups were CAG-only (N=71) and CAT-only (N=68), whereas 61 TFs were shared across all four CH contexts but absent from CG, and only 31 TFs were shared across all five contexts. In inhibitory neurons (571 unique TFs), the largest context-specific group was CG-only (N=85), followed by CTC-only (N=65) and CAG-only (N=52), with 30 TFs shared across all five contexts. Comparison between neuronal classes within each DNAm context revealed both shared and class-specific TFs (**Fig. 3D**). CG preferentially captured inhibitory-associated TFs, whereas CH contexts preferentially captured excitatory-associated TFs. For example, NEUROD2, a canonical excitatory neuron TF, was detected across all CH contexts in excitatory neurons but was absent in inhibitory neurons, whereas the inhibitory-associated TF LMX1B was detected predominantly in inhibitory neurons in the CG context.

We compared evolutionary constraint of inferred TFs across cell types and DNAm contexts using LOEUF (loss-of-function observed/expected upper bound fraction)^32^. Within the CG context, excitatory neuron cell types generally showed higher median LOEUF values than inhibitory and non-neuronal cell types (**Supplementary Figure 1**). Formal testing showed that TFs specific to non-neuronal cells had significantly lower LOEUF values than those specific to excitatory (p = 7.3 × 10^⁻^□) or inhibitory neurons (p = 0.0027), with a trend toward lower values in inhibitory than excitatory neuron TFs (p = 0.068) (**Fig. 3E**). This pattern corroborates our previous observation that inhibitory neuron-specific differentially methylated regions show greater sequence conservation than excitatory neuron-specific regions^9^.

Critically, TFs detected specifically in CH contexts had lower LOEUF values than those detected in CG contexts in both excitatory and inhibitory neurons (**Fig. 3F**), indicating that mCH-associated TF programs are under stronger evolutionary constraint than mCG-associated programs. This result is consistent with evidence that mCH is evolutionarily conserved across vertebrates^17^ and that disruption of mCH regulation through loss of MECP2 or DNMT3A causes severe neurodevelopmental phenotypes^19,33–36^.

### *In silico* mutagenesis predicts cell type-specific variant effects on DNAm

We performed *in silico* mutagenesis to predict the effects of common variants (minor allele frequency > 0.01 in CEU samples from the 1000 Genomes Project) on DNAm across methylation contexts and cell subtypes. To assess prediction accuracy, we leveraged mQTL data in the CG context from four broad cell classes (excitatory neurons, inhibitory neurons, astrocytes, and oligodendrocytes) derived from a single-cell DNAm dataset with matched genotype information^37^. SNP-CG pairs with higher-ranked predicted effects showed, on average, greater directional concordance with mQTL effects and larger mQTL z-scores (**Fig. 4A**). Notably, concordance was highest when predictions were compared with mQTLs from the matched cell class, indicating that the model captures cell type specificity. For example, among the top 0.01% of ranked SNP-CG pairs, directional concordance for excitatory neuron subtypes evaluated against excitatory neuron mQTLs reached an average of 89.5%, compared with 78.5% for inhibitory neuron subtypes, 75.9% for astrocyte subtypes and 70.8% for oligodendrocyte subtypes. Similarly, astrocyte subtypes evaluated against astrocyte mQTLs reached 93.9% concordance, compared with 82.6% for oligodendrocyte subtypes, 71.1% for inhibitory neuron subtypes and 61.4% for excitatory neuron subtypes.

**Figure 4.**
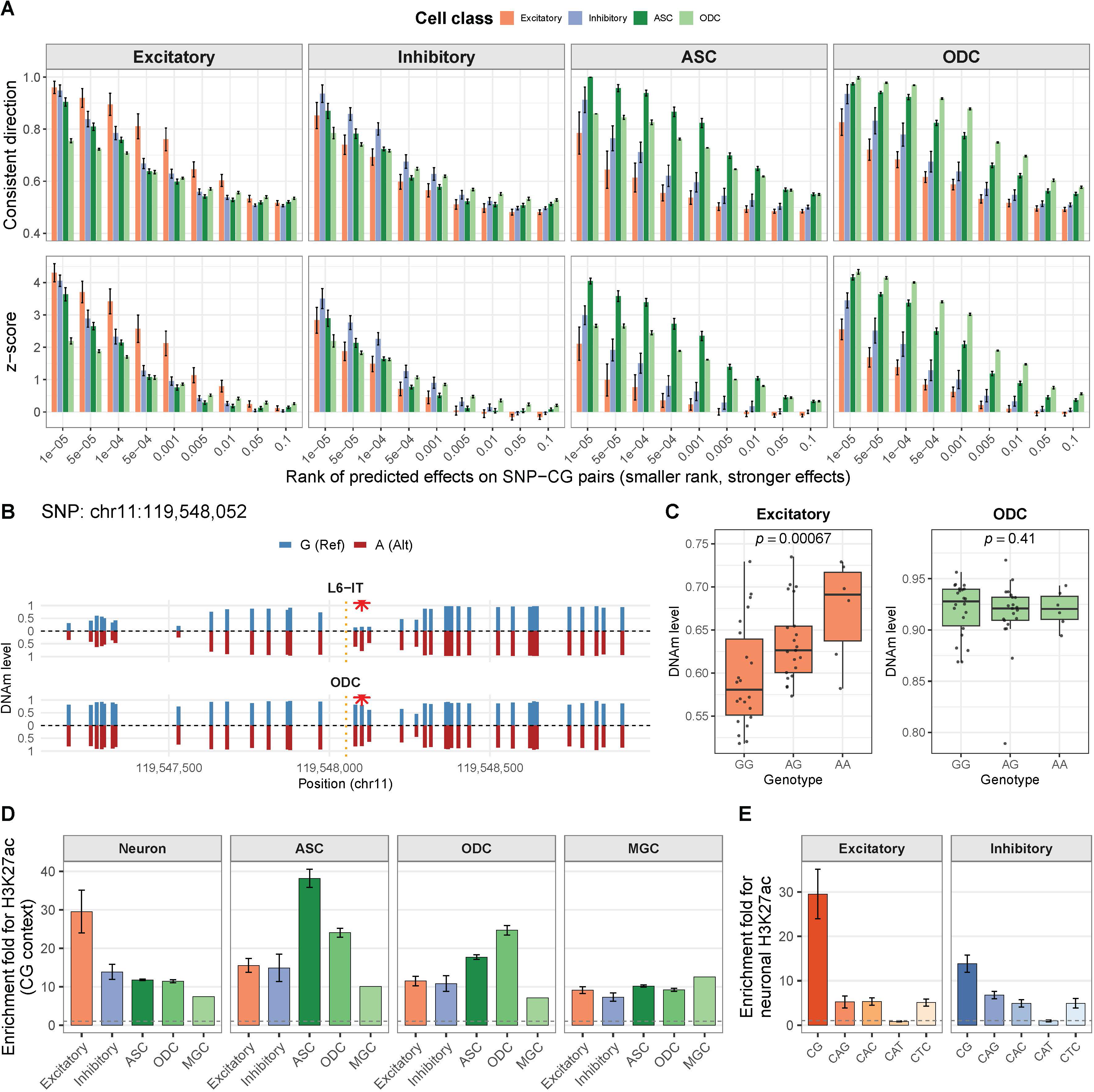
Validation of predicted common variant effects on DNAm using mQTL and active chromatin marks from broad cell classes. **(A)** Validation of predicted variant effects on mCG using mQTL data from four broad cell classes: excitatory neurons, inhibitory neurons, astrocytes (ASC) and oligodendrocytes (ODC). Each column represents the cell class of mQTL used for evaluation. Colored bars indicate the cell class of the predicted subtypes from our models (61 excitatory and 36 inhibitory subtypes from cortical regions, selected to match the cortical origin of the mQTL data, together with 6 ASC and 3 ODC subtypes). **Top row**: For each cell subtype, directional consistency was defined as the proportion of SNP-CG pairs for which the predicted effect direction matched the observed mQTL effect direction. Bars show the mean of this proportion across cell subtypes within each predicted cell class. **Bottom row**: For each cell subtype, the mQTL z-score measures the strength of association between genotype and methylation for SNP–CG pairs, weighted by the direction of predicted effect. Bars show the mean weighted z-score across cell subtypes within each predicted cell class, with larger values indicating stronger association signals with concordant directions. The x-axis denotes the rank threshold of predicted effects, with smaller values corresponding to stronger predicted effects. Error bars represent standard deviation across cell subtypes within each predicted cell class. (**B**) Predicted allele-specific DNAm levels at CG sites surrounding SNP chr11:119,548,052 in an L6-IT subtype (top) and ODC (bottom). Vertical bars represent predicted DNAm levels for the reference allele (blue, upward) and alternative allele (red, downward) at each CG site. The orange dotted line marks the SNP position, and the red asterisk indicates the CG position (chr11:119,548,082) with the largest predicted allelic difference. (**C**) Observed DNAm levels at CG (chr11:119,548,082) stratified by genotype at SNP chr11:119,548,052 in excitatory neurons (left) and ODC (right). Consistent with the model’s predictions, a significant genotype–DNAm association was detected in excitatory neurons (p = 6.7 × 10^⁻^□) but not in ODC (p = 0.41). (**D**) H3K27ac enrichment of top-ranked variants predicted to affect mCG across cell classes. For each cell subtype, variants were ranked by predicted CG impact score, and the top 10% were tested for enrichment of H3K27ac signal from four broad cell classes (Neuron, ASC, ODC, MGC), relative to the bottom 10%. Bars show the mean enrichment fold across cell subtypes within five broad cell classes in our models (Excitatory, Inhibitory, ASC, ODC, MGC). The dashed line at fold = 1 indicates no enrichment. Error bars represent standard deviation across cell subtypes. (**E**) Neuronal H3K27ac enrichment across methylation contexts. For each neuronal subtype, variants were ranked by predicted impact score within each methylation context (CG, CAG, CAC, CAT, CTC). Fold enrichment of neuronal H3K27ac signal was computed for the top 10% ranked variants relative to the bottom 10% ranked variants for each neuronal subtype. Bars show the mean fold enrichment across subtypes within excitatory (left) and inhibitory (right) neuron classes. The dashed line at fold = 1 indicates no enrichment. Error bars represent standard deviation across subtypes.

As an illustrative example, the model predicted a strong allelic effect of a SNP at chr11:119,548,052 on a nearby CG site (chr11:119,548,082) in an L6-IT excitatory neuron subtype, but minimal effect in oligodendrocytes (**Fig. 4B**). Consistent with this prediction, mQTL analysis of this SNP–CG pair showed a significant association in excitatory neurons (p = 6.7 × 10^−4^) in the same direction but not in oligodendrocytes (p = 0.41) (**Fig. 4C**).

### Common variants impacting mCG preferentially reside in active chromatin regions

To summarize the overall impact of each variant on nearby cytosines, we defined a variant impact score as the aggregated relative DNAm change within ±1 kb of the variant, taken in the dominant direction (**Methods**). To assess the regulatory potential of variants with large impact scores, we ranked variants within each cell subtype and computed the enrichment of each rank interval for H3K27ac peaks from four broad cell classes (neurons, astrocytes, oligodendrocytes, and microglia)^38^, using the bottom 10% of variants as the baseline.

In the CG context, variants with larger impact scores showed progressively stronger enrichment for H3K27ac across all cell subtypes, with the strongest signals observed in matched or closely related cell subtypes (**Supplementary Figure 2**). **Figure 4D** summarizes the average enrichment for the top 10% of ranked variants across cell subtypes within five broad cell classes. For example, the top 10% of variants predicted to affect mCG in excitatory neuron subtypes showed approximately 30-fold enrichment for neuronal H3K27ac, with weaker enrichment for top variants from inhibitory neuron, astrocyte, oligodendrocyte, and microglial subtypes. We next asked whether variants predicted to affect mCH show comparable enrichment. In both excitatory and inhibitory neuron subtypes, variants impacting mCH showed substantially weaker enrichment for neuronal H3K27ac across the CAC, CAG, and CTC contexts, and no enrichment in the CAT context (**Fig. 4E and Supplementary Figure 2**).

To validate these patterns at higher cellular resolution, we leveraged single-nucleus ATAC–seq data from postmortem human brain^39^ to test enrichment of top-ranked variants in accessible chromatin regions in cell subtypes. This analysis recapitulated the H3K27ac results: variants impacting mCG showed robust enrichment that generally peaked in matching or closely related subtypes (**Supplementary Figure 3**), whereas variants impacting mCH exhibited markedly weaker enrichment across CH contexts (**Supplementary Figures 4-7**). Together, these results indicate that common variants impacting mCG preferentially localize to active regulatory regions, whereas variants affecting mCH show limited enrichment in active chromatin.

### Common variants impacting mCG contribute more to heritability of brain-related traits

We next tested whether variants with large predicted impact scores are enriched for heritability across 20 brain-related traits, with height included as a negative control, using stratified linkage disequilibrium score regression (S-LDSC) (**Supplementary Table 3**). For each cell subtype, we constructed annotations from the top 10% of common variants ranked by predicted impact on mCG or mCH. Variants predicted to affect mCG showed strong heritability enrichment across most neuronal cell subtypes for nearly all brain-related traits (FDR < 0.05; except for Alzheimer’s disease), whereas enrichment was substantially weaker in non-neuronal cell types and absent for height (**Fig. 5A**). Notably, heritability enrichment was significantly higher in excitatory than inhibitory neuronal subtypes for 16 of 20 traits (two-sided Wilcoxon rank-sum test, FDR < 0.05; **Fig. 5B**), mirroring the weaker evolutionary constraint of excitatory neuronal regulatory programs observed above.

**Figure 5.**
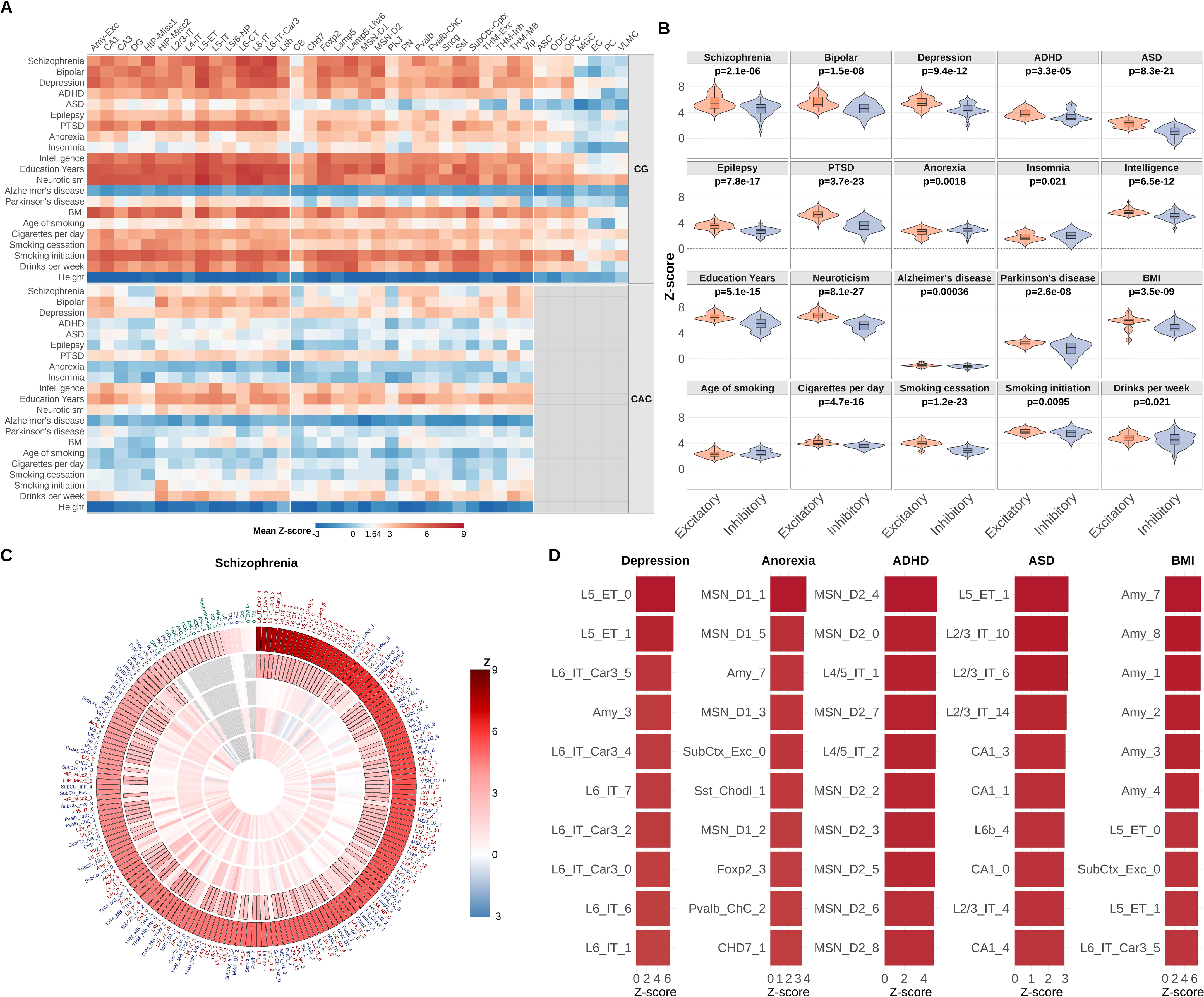
Common variants impacting mCG are enriched for the heritability of brain-related traits in neuronal cell types. **(A)** Heatmap of heritability enrichment across cell types and methylation contexts. Each tile shows the mean S-LDSC coefficient z-score, averaged across subtypes within each cell type, for 21 traits (20 brain-related plus height as negative control). Results are shown for mCG and one mCH context (CAC). Grey tiles indicate missing data for non-neuronal cell types in the CAC context, as mCH models were trained only for neuronal cell types. The dashed line at z = 1.64 corresponds to one-sided p = 0.05. **(B)** Comparison of heritability enrichment between excitatory and inhibitory neurons for brain-related traits in the CG context. Violin plots show the distribution of z-scores across subtypes. Two-sided p-values from Wilcoxon rank-sum tests are shown for traits with significant differences (FDR < 0.05). **(C)** Polar heatmap of heritability enrichment for schizophrenia across all 186 cell subtypes and five methylation contexts. Each concentric ring represents a methylation context (outer to inner: CG, CAC, CAG, CAT, CTC), and each radial position represents a cell subtype, ordered by decreasing z-score in the CG context. Tile color indicates the z-score; tiles with black borders denote significant enrichment (FDR < 0.05). Grey tiles indicate missing data as mCH model was not trained for non-neuronal cell types. Cell subtype labels are colored by cell class. **(D)** Top 10 cell subtypes ranked by z-score in the CG context for five representative traits.

By contrast, variants predicted to impact mCH showed markedly weaker enrichment across brain-related traits (**Fig. 5A and Supplementary Figure 8**), consistent with the stronger evolutionary constraint of mCH-associated TF programs. Together, these convergent lines of evidence—from evolutionary constraint, active chromatin enrichment and heritability enrichment—indicate that common genetic variation influencing brain traits preferentially acts through mCG-linked regulatory mechanisms, whereas mCH-associated effects make a comparatively limited contribution.

To explore how predicted variant effects implicate disease-relevant cell types, we ranked all 186 cell subtypes by heritability enrichment for each trait (**Supplementary Figure 9**). **Figure 5C** shows the ranking for schizophrenia: although many neuronal subtypes were enriched for schizophrenia risk, the highest-ranked cell subtypes were deep-layer excitatory neurons, consistent with preferential transcriptional dysregulation of deep-layer cortical projection neurons in schizophrenia^40^. We also highlight the top ten ranked cell subtypes for five additional traits (**Fig. 5D**). Depression similarly showed strong enrichment in deep-layer excitatory neurons, which showed preferential transcriptional alterations in this disorder^41^. Inhibitory subtypes were most enriched for anorexia and ADHD, driven primarily by medium spiny neurons (MSNs), with D1-MSNs enriched for anorexia and D2-MSNs for ADHD. This pattern is consistent with the functional opposition of direct and indirect striatal pathways in behavioral control^42^. Superficial-layer and hippocampal excitatory neurons were most enriched for ASD, in line with studies implicating upper-layer excitatory neurons^43^ and hippocampal dysfunction^44^ to ASD. Amygdala excitatory neurons showed the strongest enrichment for BMI, aligning with studies that implicate amygdala in the regulation of body mass and obesity^45–47^.

### Noncoding *de novo* mutations in ASD preferentially impact mCH, but not mCG

The weaker heritability enrichment of common variants predicted to impact mCH, together with the stronger evolutionary constraint of mCH-associated TF programs, suggests that variants disrupting mCH-linked regulation are subject to stronger purifying selection and therefore rarely reach high allele frequencies. We thus hypothesized that mCH-mediated genetic risk is carried predominantly by rare variants of large effect. To test this, we used our models to predict the impact of noncoding de novo mutations (DNMs) on DNAm across methylation contexts and neuronal subtypes, using DNMs identified in 3,508 ASD probands and 2,218 unaffected siblings from the SPARK cohort (302,603 DNMs in probands and 196,898 in siblings), and sought replication in the independent SSC cohort, comprising 2,274 probands and 1,835 unaffected siblings (209,396 DNMs in probands and 171,258 in siblings).

Across all DNMs, differences in predicted impact scores between probands and siblings were minimal across 172 neuronal cell types in SPARK (**Fig. 6**), consistent with the expectation that most DNMs are functionally neutral. We therefore restricted the analysis to DNMs overlapping functionally annotated regions, including evolutionarily conserved elements, active neuronal chromatin regions (H3K27ac peaks), and their intersection. Neither conservation nor H3K27ac filtering alone yielded significant proband–sibling differences after multiple-testing correction. By contrast, DNMs located within the intersection of neuronal H3K27ac peaks and conserved regions showed significantly larger predicted impact scores in probands than in siblings across many neuronal cell subtypes (FDR < 0.05), with stronger effects at more stringent conservation thresholds. Under the most stringent filter (H3K27ac peaks and phastCons > 0.8), the signal was strongest for mCAC (163 of 172 cell subtypes at FDR < 0.05), followed by mCTC (108 of 172) and mCAG (98 of 172), whereas mCAT showed a weaker trend that did not survive correction. By contrast, mCG showed no proband–sibling differences under any functional filter (FDR > 0.05). In the SSC cohort, under the same stringent filter, most SPARK discoveries were replicated at nominal significance (p < 0.05): 91% for mCTC (98 of 108), 71% for mCAC (116 of 163), and 54% for mCAG (53 of 98), whereas mCG showed no signal (**Fig. 6**). These results support a model in which noncoding DNMs in ASD preferentially perturb mCH, but not mCG.

**Figure 6.**
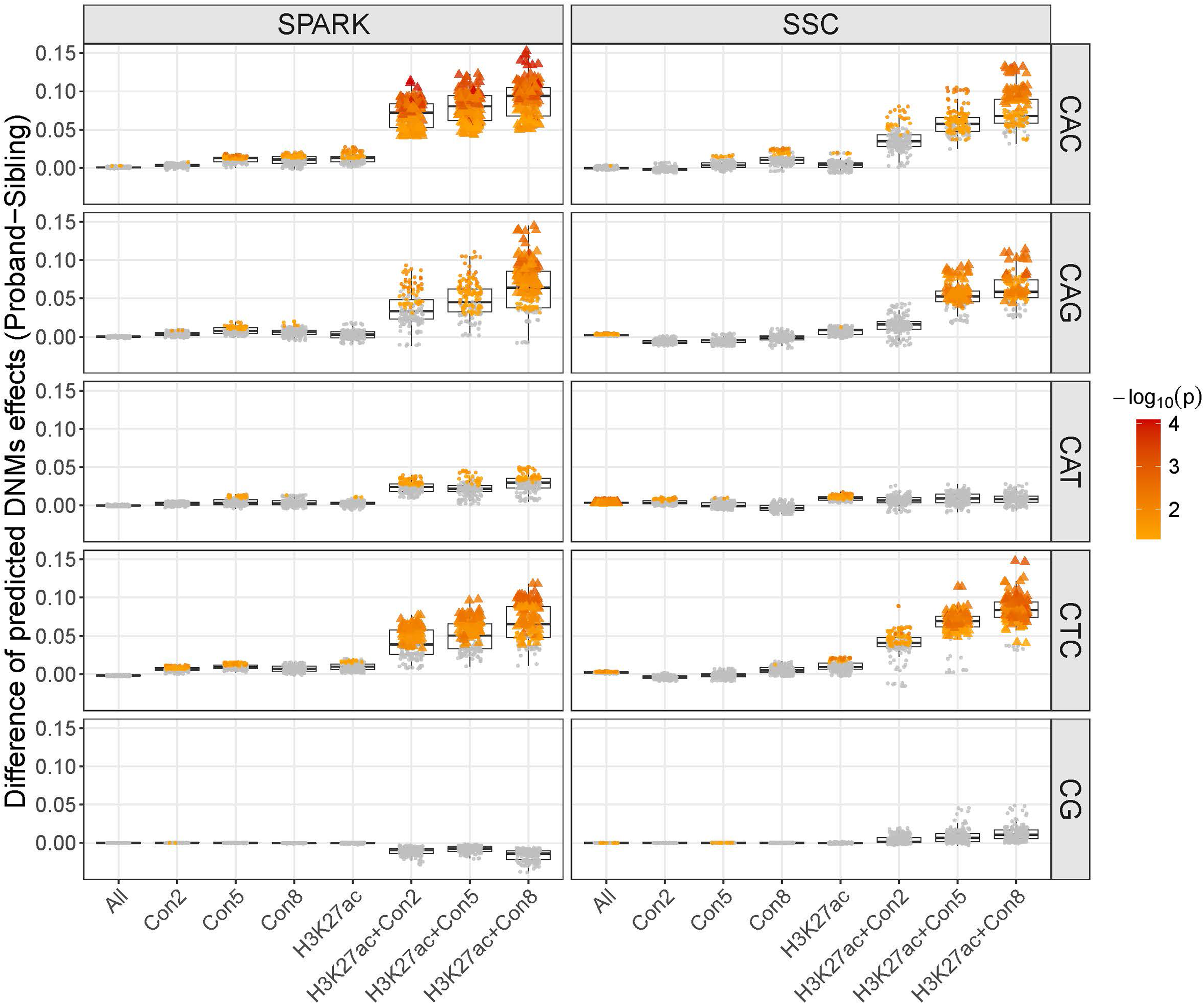
Noncoding *de novo* mutations in conserved and neuronally active chromatin regions show larger impacts on mCH in ASD probands than in unaffected siblings. Comparison of predicted DNAm impact score between probands and unaffected siblings across 172 neuronal subtypes. Each point in the boxplot represents one neuronal subtype and shows the Hodges–Lehmann estimate of the difference in predicted impact score (proband minus sibling; y-axis) across all DNMs, or across subsets of DNMs stratified by functional annotation within each methylation context. Functional annotations include evolutionarily conserved regions at different thresholds (Con2: phastCons > 0.2; Con5: phastCons > 0.5; Con8: phastCons > 0.8), neuronal H3K27ac peaks (H3K27ac), and their intersections. Colored points indicate nominal significance (*P* < 0.05) from one-sided Wilcoxon rank-sum test, and triangles denote points with FDR < 0.05.

## Discussion

Our results indicate that common and rare noncoding variants influence brain-related traits through distinct cellular DNAm mechanisms in the human brain. Common variants preferentially act through mCG, particularly in excitatory neurons, whereas noncoding DNMs in ASD preferentially perturb mCH at conserved neuronal regulatory regions. This contrast is mirrored by the evolutionary signatures of the TF programs learned by our models: mCH-associated TFs are more evolutionarily constrained than those linked to mCG, suggesting that mCH-linked regulatory programs are less tolerant of genetic perturbation. Variants disrupting these constrained programs are therefore expected to remain rare or arise *de novo*, whereas variation affecting the more permissive mCG-associated regulatory landscape can persist at higher frequencies and contribute to polygenic risk.

The stronger constraint on mCH-associated TF programs is consistent with the biology of a specialized neuronal methylation system that emerged in vertebrates and became deeply conserved^17^. Neuronal mCH is largely restricted to vertebrates, accumulates postnatally in neurons, and marks a conserved set of developmentally regulated genes. Its core machinery also appears unusually sensitive to perturbation. Pathogenic mutations in DNMT3A, a principal writer of mCH, cause Tatton-Brown-Rahman syndrome, a neurodevelopmental disorder characterized by intellectual disability and often accompanied by autistic features^33,34^, whereas mutations in MeCP2, a major reader of mCAC, cause Rett syndrome^36^. Moreover, selective loss of MeCP2 recognition of mCAC is sufficient to produce severe Rett-like phenotypes in mice^19^. Together, these observations support a model in which the vertebrate neuronal mCH system is especially intolerant of perturbation relative to the broader mCG regulatory landscape. Our finding that noncoding DNMs in ASD preferentially perturb mCH at conserved neuronal regulatory regions extends this model by suggesting that mCH-linked regulation can be disrupted not only by mutations in core trans-acting mCH machinery, but also by cis-regulatory sequence changes that alter DNAm at specific loci.

Our heritability analyses further show that, within the CG context, common-variant enrichment is generally stronger in excitatory than inhibitory neurons across brain-related traits, paralleling the weaker constraint of mCG-associated TF programs in excitatory neurons learned by our models. This pattern is consistent with comparative epigenomic and transcriptomic evidence that inhibitory neurons are more deeply conserved than excitatory neurons. In our prior single-cell methylome study, regulatory DNA elements showed stronger cross-species conservation in inhibitory than in excitatory neurons^9^, and cross-species single-cell transcriptomic analyses similarly showed that major GABAergic interneuron classes are deeply conserved, whereas glutamatergic neurons exhibit greater divergence^48–52^. Greater conservation of inhibitory neuronal regulatory programs may therefore limit the accumulation of common functional variants in these cells, biasing polygenic risk towards the less constrained regulatory architecture of excitatory neurons. Notably, within the CG context, the strongest evolutionary constraint was observed among TFs specific to non-neuronal cells, suggesting that glial and other non-neuronal regulatory programs—despite showing the weakest heritability enrichment based on common variants—may also be important targets of purifying selection and could contribute to disease risk through rare variants that remain to be identified.

This work has several limitations. First, our models were trained on DNAm profiles from three adult brains and use reference-genome sequence as input, limiting their ability to capture inter-individual genetic variation and developmental dynamics. Training on larger cohorts with matched sequence data, and extending these analyses to developmental brain samples, should help address both limitations. Second, the association between rare noncoding variation and mCH was evaluated only in ASD; whether similar mCH-linked rare-variant mechanisms operate in other brain disorders remains unknown. Third, our framework is designed to identify risk variants acting through DNAm and does not capture variants that influence disease through regulatory mechanisms independent of DNAm.

Despite these limitations, our findings support a model in which common and rare noncoding variants engage distinct DNA methylation systems in the human brain. More broadly, they provide a framework for interpreting how noncoding variation contributes to brain disorders in a cell-type- and methylation-context-resolved manner.

## Materials and Methods

### Training dataset

We trained models on a single-nucleus DNAm dataset from the human brain spanning 46 regions from three adult male donors. Details about data generation and processing were described in a previous study^29^. Briefly, DNAm profiles were generated by snmC-seq3, together with snm3C-seq for jointly profiling of DNAm and chromatin conformation from 17 regions. After quality control, the dataset comprised approximately 517,000 nuclei (399,000 neurons and 118,000 non-neuronal cells), clustered into 40 major cell types (16 excitatory, 17 inhibitory and 7 non-neuronal), which were further subdivided into 188 subtypes.

For each major cell type and subtype defined in the original study^29^, we quantified DNAm in the mCG and mCH (mCAC, mCAG, mCAT, mCTC) contexts as the fraction of methylated reads in pseudo-bulk aggregates. Coverage thresholds were tailored to each cell type to balance data quality and training sample size: 50× for most major cell types, relaxed to 30× for L5-ET, THM-Exc, CA1, EC, VLMC, and PC, and to 20× for CA3, THM-Inh, HIP-Misc1, and PKJ due to smaller cell numbers. Across the 40 major cell types, training sample sizes ranged from approximately 2.1 to 105.1 million cytosine sites depending on methylation context, including 2.1–26.5 million in mCG and 2.1–77.3 million (mCAC), 2.4–105.1 million (mCAG), 2.5–94.6 million (mCAT) and 2.2–87.2 million (mCTC) in mCH contexts (**Supplementary Table 4**).

For cell subtypes, we applied a minimum coverage threshold of 10×, excluding two subtypes (L5_ET_2 and SubCtx_Exc_2) because of insufficient cell numbers (95 and 53 cells, respectively), yielding 186 subtypes for downstream analysis. Across subtypes, training sample sizes ranged from 8.6 to 26.3 million cytosine sites in mCG and from 2.5 to 77.1 million (mCAC), 3.1 to 105.1 million (mCAG), 3.1 to 94.6 million (mCAT) and 2.7 to 86.9 million (mCTC) in mCH contexts (**Supplementary Table 5**).

For both pretraining and fine-tuning, mCG and mCH sites were partitioned into training, validation, and test sets by chromosome. Sites on chromosomes 1–20 were used for training, chromosome 21 for validation, and chromosome 22 as an independent test set for evaluating prediction performance.

### Model architecture

Our model comprises three main modules: a convolutional neural network (CNN), a Transformer encoder, and a fully connected network (**Fig. 1**). The model takes a one-hot-encoded 2-kb DNA sequence as input and outputs the DNAm level at the central CG or CH site across multiple cell types. We chose a 2-kb window based on prior benchmarking of sequence lengths (1, 2, 3, and 4 kb)^27^. Full architectural details are provided in **Supplementary Table 6**.

The CNN module consists of three convolutional layers with 11-bp kernels and either 768 filters for CG models or 512 filters for CH models. Each layer is followed by rectified linear unit (ReLU) activation and batch normalization. A max-pooling layer (kernel size = stride = 20) aggregates features across non-overlapping 20-bp segments, followed by batch normalization and dropout (rate = 0.5). The pooled features are then passed to a Transformer encoder with 12 layers and 12 attention heads in CG models, or 8 layers and 8 attention heads in CH models. Each encoder layer contains multi-head self-attention and position-wise feed-forward blocks with residual connections, normalization, and dropout (rate = 0.1). The fully connected network contains a single hidden layer with 512 units and dropout (rate = 0.1), followed by a sigmoid-activated output layer with one unit for each cell type.

### Pretraining and fine-tuning

Our CG models contain more than 121 million parameters and our CH models more than 43 million parameters. To address limited training data, particularly for cell subtypes with fewer cells, we implemented a two-step hierarchical training strategy. First, we pretrained models in a multi-task learning framework by grouping cell types into broad functional classes. For mCG, we pretrained three models for excitatory neurons (16 cell types), inhibitory neurons (17 cell types), and non-neuronal cells (7 cell types). For mCH, we pretrained separate excitatory and inhibitory models for each CH context (CAC, CAG, CAT and CTC), yielding eight pretraining models in total. This strategy enables learning of features shared across related cell types while reducing computational burden.

In the second step, we fine-tuned the pretrained models in a multi-task learning framework using subtype-specific training samples to capture features unique to individual cell subtypes. Specifically, excitatory models were fine-tuned on subtypes within each of the 16 excitatory major cell types across CG and all four CH contexts; inhibitory models were fine-tuned on subtypes within each of the 17 inhibitory cell types across CG and all four CH contexts; and non-neuronal models were fine-tuned on subtypes within each of the 7 non-neuronal cell types in the CG context only.

### DNA motif analysis

We examined filters in the first convolutional layer of each fine-tuned model to identify DNA motifs associated with DNAm levels in each cell type and methylation context. Following a previously described method^53^, we identified DNA motifs for each filter by selecting sequence subsets that produced activation values exceeding half of the maximum activation observed across all scanned sequences. These sequences were aligned to generate a position weight matrix, which was then matched to annotated transcription factor binding motifs in the *Homo sapiens* CIS-BP database using Tomtom v4.10.1^54^. Matches with FDR < 0.05 were considered significant.

### *In silico* mutagenesis

We performed *in silico* mutagenesis to estimate variant effects on DNAm levels in each cell type and methylation context. For each variant *v*, and for each cytosine *c* within ±1,000 bp of *v*, we built two 2-kb sequences centered on *c*: a reference sequence and a mutant sequence carrying the alternate allele of *v*. For each cell type *t* and methylation context *k* ∊ {*CG, CAC, CAG, CAT, CTC*}, the fine-tuned model was used to predict DNAm at *c* from each sequence. Let *m_ref_* (*c*; *t*,*k*), *m_mut_* (*c*; *t*,*k*) ∊ [0, 1] denote predicted DNAm for the reference and mutant sequences, respectively. We defined the relative change of DNAm at cytosine *c* as:

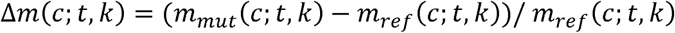

We then aggregated increased and decreased DNAm levels separately over the neighborhood *N*(*v*), defined as all cytosines *c* within ±1,000 bp of *v*:

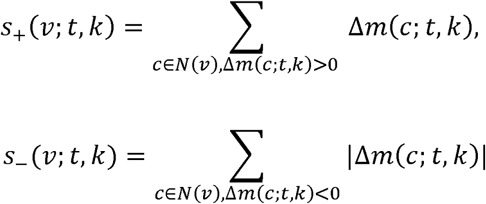

The overall impact score was defined as the aggregated effects in the dominant direction:

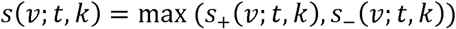

We performed this *in silico* mutagenesis for 9,042,066 SNPs (minor allele frequency > 0.01) observed in the CEU samples from the 1000 Genomes Project. SNPs were ranked in descending order of their overall impact score for downstream functional evaluation.

To evaluate the regulatory potential of variants with large impact scores, we tested their enrichment for active chromatin marks (H3K27ac) using profiles generated previously for four broad brain cell types—neurons, microglia, astrocytes, and oligodendrocytes^38^. To evaluate functional relevance at finer cellular resolution, we further leveraged single-nucleus ATAC-seq data from postmortem human brain^39^, which provide open chromatin profiles for refined subtypes. We then assessed the enrichment of variants ranked by impact score within open chromatin regions across these refined cell types.

### mQTL evidence from broad cell classes in the human brain

We previously generated whole-genome sequencing data from bulk tissue and snmCT-seq (paired single-nucleus methylation and transcriptome) profiles from postmortem cortex tissue (DLPFC) for 49 donors in an ASD case-control study^37^. Using per-nucleus whole-genome methylation profiles together with prior cell type annotations, we generated donor-by-cell type pseudobulks for four broad cell classes (excitatory neurons, inhibitory neurons, astrocytes, and oligodendrocytes) to obtain more robust estimates of DNAm levels at each CG site while preserving a reasonable degree of cell type specificity for validating predicted variant effects on DNAm. Methylation fractions were tabulated at each CG site, smoothed using the BSmooth algorithm in bsseq^55^, and quantile-normalized across donors before mQTL analysis. Donors were genotyped using the DRAGEN pipeline with developer recommendations^56^. Genotypes were harmonized with the 1000 Genomes reference panel, retaining variants with ≤5% missingness and present in the reference panel, before phasing and imputation using BEAGLE v5.4^57^.

Variants with MAF > 0.10 and r² > 0.70 (if imputed) and all CG sites were retained for local mQTL testing within a ±1 kb window. Smoothed, quantile-normalized methylation fractions at each CG were regressed on variant dosage using QTLTools^58^, adjusting for donor diagnosis (ASD or neurotypical control), age (years and log-years), two ancestry principal components, and *k* methylation principal components. Methylation PCA was computed on logit-transformed methylation fractions for cytosines with between-donor interquartile range ≥ 0.10; *k* was determined algorithmically for each cell type, excluding principal components highly correlated with known covariates (univariate Pearson |r| ≥ 0.50 with any covariate, or variance explained r² ≥ 0.50 by the set of all known covariates).

To evaluate the accuracy of *in silico* mutagenesis, we restricted analyses to SNP–CG pairs with mQTL association *P* < 0.01 to reduce inclusion of variants unlikely to influence DNAm. For each cell subtype, SNP-CG pairs were ranked by predicted effect size from the corresponding fine-tuned model. Within ranked intervals, we quantified the proportion of SNP–CG pairs showing directional concordance between predicted variant effects and observed mQTL effects. We further assessed prediction performance by comparing mQTL signal strength across ranked intervals using the mean association *z*-score weighted by the predicted direction of effect. Specifically, *z*-score was multiplied by +1 when the predicted effect direction was positive and by −1 when it was negative. This signed *z*-score captures agreement in effect direction while preserving effect magnitude, such that a larger mean signed *z*-score indicates stronger association signals with concordant directions.

### Stratified LDSC

We performed S-LDSC^59^ to evaluate the enrichment of heritability of 20 brain-related traits for top 10% ranked variants by their impact scores in each cell subtype within each methylation context. We also included human height as a negative control to examine whether our findings are specific to brain-related traits. We downloaded GWAS summary statistics of each trait from the sources listed in **Supplementary Table 7**. S-LDSC was run for each list of variants with the baseline LD model v2.2 that included 97 annotations to control for the LD between variants with other functional annotations in the genome. We used HapMap Project phase 3 SNPs as regression SNPs, and 1000 Genomes SNPs of European ancestry samples as reference SNPs, which were all downloaded from the LDSC resource website. To evaluate the unique contribution of predicted regulatory variants to trait heritability, we used the *z* score of per-SNP heritability from S-LDSC. This metric allows us to discern the unique contributions of candidate annotations while accounting for contributions from other functional annotations in the baseline model. The *P* values are derived from the *z*-score assuming a normal distribution with one-tailed test, and FDR was computed from the *p* values based on Benjamini and Hochberg procedure.

### *De novo* mutation analysis for ASD

We analyzed DNMs in ASD probands and unaffected siblings from two independent cohorts, SPARK and SSC, which were called using our custom bioinformatic pipeline as described previously^60^. Briefly, candidate DNMs were identified from family-level VCFs generated by DeepVariant^61^. We then applied stringent quality filters using BCFtools^62^ and BEDTools (v2.31.0)^63^ to remove likely false positives based on read depth, mapping quality, and allele balance, among other quality metrics. The SPARK discovery cohort (March 2023 release) comprises 3,508 affected probands and 2,218 unaffected siblings from 3,357 families. The SSC replication cohort comprises 2,274 affected probands and 1,835 unaffected siblings from 2,274 families. We annotated all candidate DNMs using ANNOVAR^64^. Our analysis focused on noncoding DNMs, comprising 302,603 (in probands) and 196,898 (in siblings) in SPARK, and 209,396 (in probands) and 171,258 (in siblings) in SSC.

We predicted the impact scores of DNMs on DNAm for each cell subtype and methylation context using the same *in silico* mutagenesis approach described above. We annotated DNMs for evolutionary conservation using phastCons^65^ and phyloP scores^66^ from 46-way vertebrate alignments downloaded from the UCSC Genome Browser and lifted over to hg38 genome coordinates using CrossMap software^67^. We further annotated DNMs for overlap with neuronal active chromatin regions using H3K27ac peaks obtained from a previous study^38^.

We compared predicted DNM impact scores between probands and unaffected siblings using one-sided Wilcoxon rank-sum tests, testing for larger effects in probands. Effect sizes were estimated as the Hodges-Lehmann estimator, defined as the median of all pairwise differences between proband and sibling scores, which is the natural location shift estimator associated with the Wilcoxon test. Tests were performed separately for each neuronal cell type within each methylation context, and p-values were adjusted for multiple testing using the Benjamini-Hochberg procedure across cell types within each context.

## Acknowledgments

We acknowledge the Psychiatric Genomics Consortium, UK BioBank, GSCAN (the GWAS & Sequencing Consortium of Alcohol and Nicotine), the International Parkinson’s Disease Genomics Consortium, and the CTGlab (the Complex Trait Genetics lab at the VU University Amsterdam the Amsterdam University Medical Centre) for making GWAS results publicly available.

## Funding

National Institutes of Health grant R01MH121394 (SH)

National Institutes of Health grant R01MH112751 (SH)

National Institutes of Health grant HG013359 (KW)

Lieber Institute for Brain Development

## Author contributions

Conceptualization: JZ, DRW, SH

Methodology: JZ, SH

Investigation: JZ, CL1, XL, YZ, YW, JS, BM, CL2, CL3, KW, SH

Visualization: JZ, SH

Supervision: DRW, SH

Writing—original draft: JZ, DRW, SH

Writing—review & editing: JZ, CL1, XL, YZ, YW, JS, BM, CL2, CL3, KW, DRW, SH

## Data and materials availability

The single nucleus DNAm dataset used for training the deep learning models was downloaded from the GEO database with accession code GSE215353. All other data needed to evaluate the conclusions in the paper are present in the paper and/or the Supplementary Materials.

Deep learning model code and usage are available from GitHub: https://github.com/LieberInstitute/Brain_INTERACT

## Supplementary Figure Legends

**Supplementary Figure 1. LOEUF constraint scores of TFs across cell types by methylation context.** Violin plots showing the distribution of LOEUF (loss-of-function observed/expected upper bound fraction) scores for transcription factors (TFs) detected as significant (FDR < 0.05) in each cell type, shown separately for each methylation context: CG, CAC, CAG, CAT, and CTC (top to bottom). Cell types are ordered by median LOEUF along the x-axis, with the number of TFs in parentheses. Fill colors indicate broad cell class: non-neuronal (green), Inhibitory (blue), and Excitatory (red). Lower LOEUF values indicate greater intolerance to loss-of-function variants, reflecting stronger evolutionary constraint. LOEUF scores were obtained from gnomAD v4.1 constraint metrics.

**Supplementary Figure 2. Predicted DNAm regulatory variants in the CG context, and to a lesser extent in CH contexts, are enriched for H3K27ac peaks in matching cell class.**

**(A)** Enrichment of CG-context DNAm regulatory variants for cell class-specific H3K27ac peaks. Each panel displays variants from models for each cell type (rows), ranked by predicted impact score (columns), and their enrichment for H3K27ac peaks in each broad cell class (astrocyte, microglia, oligodendrocyte [Oligo], and neuron). For each cell type, enrichment was summarized as the mean fold-enrichment across all cell subtypes within that cell type. Tile color indicates the mean proportion of variants overlapping H3K27ac peaks across cell subtypes for a given rank interval and broad cell class; numbers indicate fold-enrichment relative to the bottom 10% of variants (rank interval “0.9–1.0”). Rank interval “0.0–0.1” represents the top 10% of variants with the largest predicted impact scores. **(B)** Comparison of enrichment for neuronal H3K27ac peaks across variants from different methylation contexts. Each panel shows variants from models for neuronal cell types (rows), ranked by predicted impact score (columns) in each methylation context. Enrichment was summarized as the mean fold-enrichment across cell subtypes within each neuronal cell type. Top-ranked variants (top 10%) in the CG context show substantially stronger enrichment than those in CH contexts.

**Supplementary Figure 3. Predicted mCG regulatory variants show enrichment for accessible chromatin regions in matching or closely related cell subtypes.** Enrichment of mCG regulatory variants for accessible chromatin peaks across refined neuronal and non-neuronal subtypes in human brain. Each panel displays variants from models for cell types (rows), ranked by predicted impact score (columns), and their enrichment for snATAC-seq peaks in each refined cell subtype. For each cell type, enrichment was summarized as the mean fold-enrichment across all subtypes within that cell type. Rank interval “0.0–0.1” represents the top 10% of variants with the largest predicted impact scores. Tile color indicates the mean proportion of variants overlapping accessible chromatin peaks across cell subtypes for a given rank interval and refined cell subtype; numbers indicate fold-enrichment relative to the bottom 10% of variants (rank interval “0.9–1.0”). Enrichment generally peaks in matching or closely related cell subtypes in the snATAC-seq data, recapitulating the H3K27ac enrichment patterns observed in Fig. 4D and validating these findings at higher cellular resolution.

**Supplementary Figure 4. Predicted mCAC regulatory variants show weaker enrichment for accessible chromatin regions than mCG variants.** Enrichment of mCAC regulatory variants for single-nucleus ATAC-seq (snATAC-seq) accessible chromatin peaks across refined neuronal and non-neuronal cell subtypes in human brain. Each panel displays variants from models for cell types (rows), ranked by predicted impact score (columns), and their enrichment for snATAC-seq peaks in each refined cell subtype. For each cell type, enrichment was summarized as the mean fold-enrichment across all subtypes within that cell type. Rank interval “0.0–0.1” represents the top 10% of variants with the largest predicted impact scores. Tile color indicates the mean proportion of variants overlapping accessible chromatin peaks across cell subtypes for a given rank interval and refined cell subtype; numbers indicate fold-enrichment relative to the bottom 10% of variants (rank interval “0.9–1.0”). Enrichment is markedly weaker than the mCG-context pattern observed in Supplementary Fig. 3, consistent with the reduced H3K27ac enrichment for mCH variants shown in Fig. 4E.

**Supplementary Figure 5. Predicted mCAG regulatory variants show weaker enrichment for accessible chromatin regions than mCG variants.** As in Supplementary Fig. 4, but for the CAG context, enrichment is markedly weaker than the mCG-context pattern observed in Supplementary Figure 3, consistent with the reduced H3K27ac enrichment for mCH variants shown in Fig. 4E.

**Supplementary Figure 6. Predicted mCAT regulatory variants show weaker enrichment for accessible chromatin regions than mCG variants.** As in Supplementary Fig. 4, but for the CAT context, enrichment is markedly weaker than the mCG-context pattern observed in Supplementary Figure 3, consistent with the reduced H3K27ac enrichment for mCH variants shown in Fig. 4E.

**Supplementary Figure 7. Predicted mCTC regulatory variants show weaker enrichment for accessible chromatin regions than mCG variants.** As in Supplementary Fig. 4, but for the CTC context, enrichment is markedly weaker than the mCG-context pattern observed in Supplementary Figure 3, consistent with the reduced H3K27ac enrichment for mCH variants shown in Fig. 4E.

**Supplementary Figure 8. Heatmap of heritability enrichment across cell types and methylation contexts.** Each tile shows the mean S-LDSC coefficient Z-score averaged across subtypes within each cell type (column), for 21 traits (20 brain-related plus height as negative control). Results are shown for mCG and four mCH context (CAC, CAG, CAT and CTC). Grey tiles indicate missing data for non-neuronal cell types in mCH contexts, as mCH models were trained only for neuronal cell types. The dashed line at z = 1.64 corresponds to one-sided p = 0.05.

**Supplementary Figure 9. Common variants affecting DNAm in the CG context contribute more to heritability of brain-related traits and disorders than those in the CH context.** Results are shown for 20 brain-related traits and disorders, with height included as a negative control. The color scale denotes the z-score of per-SNP heritability enrichment. The outermost ring represents the CG context, followed by CAC, CAG, CAT, and CTC in the innermost ring. Cell type labels around the ring are sorted by z-score from the CG context; label colors indicate the three broad cell classes (excitatory neurons, inhibitory neurons, non-neuronal cells). Tiles outlined in black indicate z-scores with FDR < 0.05.

