## Supplementary Figure 3 for "Distinct cellular DNA methylation mechanisms underlie common and rare genetic risk for brain disorders"

### ACBGM−Bergmann glia

| | 0.0−0.1 | 0.1−0.2 | 0.2−0.3 | 0.3−0.4 | 0.4−0.5 | 0.5−0.6 | 0.6−0.7 | 0.7−0.8 | 0.8−0.9 | 0.9−1.0 |
|---|---|---|---|---|---|---|---|---|---|---|
| L6b | 98.17 | 13.82 | 7.90 | 5.45 | 4.03 | 2.92 | 2.26 | 1.70 | 1.25 | 1.00 |
| L6−IT−Car3 | 86.14 | 12.02 | 7.32 | 5.08 | 3.57 | 2.85 | 2.21 | 1.75 | 1.40 | 1.00 |
| L6−IT | 96.83 | 13.49 | 8.15 | 5.70 | 4.31 | 3.26 | 2.48 | 1.84 | 1.41 | 1.00 |
| L6−CT | 112.18 | 16.16 | 9.25 | 6.46 | 4.77 | 3.47 | 2.72 | 1.96 | 1.42 | 1.00 |
| L56−NP | 66.68 | 8.50 | 5.41 | 4.03 | 3.18 | 2.65 | 2.12 | 1.77 | 1.35 | 1.00 |
| L5−IT | 86.86 | 11.87 | 7.17 | 4.87 | 3.67 | 2.77 | 2.33 | 1.81 | 1.40 | 1.00 |
| L5−ET | 99.71 | 13.27 | 7.45 | 5.41 | 3.93 | 3.10 | 2.31 | 1.74 | 1.37 | 1.00 |
| L4−IT | 72.67 | 10.00 | 5.68 | 4.05 | 3.03 | 2.33 | 2.00 | 1.57 | 1.34 | 1.00 |
| L23−IT | 87.57 | 12.52 | 7.11 | 4.99 | 3.74 | 2.81 | 2.21 | 1.65 | 1.37 | 1.00 |
| HIP−Misc2 | 72.41 | 9.71 | 6.02 | 4.11 | 2.95 | 2.52 | 2.01 | 1.56 | 1.24 | 1.00 |
| HIP−Misc1 | 97.64 | 13.42 | 7.66 | 5.20 | 3.99 | 3.07 | 2.46 | 2.08 | 1.51 | 1.00 |
| DG | 109.59 | 17.04 | 9.96 | 6.37 | 4.55 | 3.20 | 2.34 | 1.80 | 1.41 | 1.00 |
| CA3 | 124.89 | 18.49 | 10.17 | 7.08 | 4.99 | 3.54 | 2.65 | 2.01 | 1.40 | 1.00 |
| CA1 | 104.87 | 14.89 | 9.43 | 6.13 | 4.67 | 3.29 | 2.58 | 1.98 | 1.45 | 1.00 |
| Amy−Exc | 80.51 | 12.18 | 7.52 | 5.07 | 3.68 | 2.78 | 2.11 | 1.80 | 1.30 | 1.00 |
| Vip | 100.61 | 11.80 | 6.27 | 4.02 | 2.87 | 2.22 | 1.81 | 1.47 | 1.08 | 1.00 |
| THM−MB | 67.79 | 8.25 | 5.28 | 3.94 | 3.14 | 2.77 | 2.21 | 1.67 | 1.33 | 1.00 |
| THM−Inh | 96.93 | 12.71 | 7.41 | 4.95 | 3.69 | 2.98 | 2.48 | 1.93 | 1.69 | 1.00 |
| THM−Exc | 113.50 | 16.21 | 9.45 | 6.29 | 4.55 | 3.45 | 2.35 | 2.01 | 1.50 | 1.00 |
| SubCtx−Cplx | 110.68 | 14.94 | 7.95 | 5.26 | 3.76 | 2.74 | 2.06 | 1.61 | 1.26 | 1.00 |
| Sst | 74.74 | 9.34 | 5.38 | 3.86 | 3.06 | 2.66 | 2.10 | 1.64 | 1.38 | 1.00 |
| Sncg | 109.00 | 13.28 | 7.08 | 4.57 | 3.38 | 2.45 | 1.94 | 1.62 | 1.38 | 1.00 |
| Pvalb−ChC | 91.49 | 11.21 | 6.46 | 4.44 | 3.35 | 2.63 | 2.14 | 1.91 | 1.30 | 1.00 |
| Pvalb | 74.00 | 8.92 | 5.19 | 3.69 | 2.58 | 2.20 | 1.87 | 1.60 | 1.27 | 1.00 |
| PN | 111.65 | 14.09 | 7.19 | 4.85 | 3.16 | 2.19 | 1.75 | 1.51 | 1.06 | 1.00 |
| MSN−D2 | 63.63 | 9.01 | 5.68 | 4.10 | 3.13 | 2.61 | 2.25 | 1.75 | 1.55 | 1.00 |
| MSN−D1 | 60.46 | 8.02 | 5.07 | 3.80 | 3.07 | 2.53 | 1.97 | 1.74 | 1.54 | 1.00 |
| Lamp5−Lhx6 | 124.93 | 15.16 | 7.71 | 4.85 | 3.24 | 2.48 | 1.89 | 1.62 | 1.25 | 1.00 |
| Lamp5 | 122.56 | 14.66 | 7.78 | 4.83 | 3.40 | 2.49 | 1.82 | 1.67 | 1.25 | 1.00 |
| Foxp2 | 78.22 | 10.35 | 6.32 | 4.47 | 3.39 | 2.66 | 2.19 | 1.75 | 1.29 | 1.00 |
| Chd7 | 118.72 | 14.75 | 7.77 | 4.82 | 3.39 | 2.65 | 2.03 | 1.58 | 1.45 | 1.00 |
| CB | 130.50 | 17.80 | 9.73 | 5.85 | 3.87 | 3.08 | 2.47 | 1.81 | 1.54 | 1.00 |
| VLMC | 106.34 | 13.81 | 7.45 | 5.12 | 3.90 | 2.99 | 2.39 | 1.76 | 1.39 | 1.00 |
| PC | 97.16 | 12.91 | 7.60 | 5.07 | 3.90 | 3.13 | 2.43 | 1.85 | 1.46 | 1.00 |
| OPC | 248.45 | 27.35 | 12.20 | 7.30 | 5.19 | 3.11 | 2.80 | 1.76 | 1.41 | 1.00 |
| ODC | 184.77 | 22.24 | 10.87 | 6.80 | 4.60 | 3.35 | 2.26 | 1.92 | 1.38 | 1.00 |
| MGC | 54.36 | 7.06 | 4.73 | 3.63 | 3.12 | 2.39 | 2.25 | 1.67 | 1.26 | 1.00 |
| EC | 66.42 | 8.62 | 5.55 | 4.40 | 3.25 | 2.80 | 2.32 | 1.71 | 1.36 | 1.00 |
| ASC | 229.32 | 23.21 | 9.60 | 5.50 | 3.55 | 2.72 | 2.08 | 1.73 | 1.43 | 1.00 |

Percent: 0.005 0.010 0.015 0.020

### ASCNT−Non−telencephalon astrocytes

| | 0.0−0.1 | 0.1−0.2 | 0.2−0.3 | 0.3−0.4 | 0.4−0.5 | 0.5−0.6 | 0.6−0.7 | 0.7−0.8 | 0.8−0.9 | 0.9−1.0 |
|---|---|---|---|---|---|---|---|---|---|---|
| L6b | 48.04 | 11.03 | 6.96 | 4.89 | 3.72 | 2.81 | 2.20 | 1.75 | 1.43 | 1.00 |
| L6−IT−Car3 | 46.51 | 10.93 | 6.89 | 5.09 | 3.81 | 2.97 | 2.45 | 2.00 | 1.56 | 1.00 |
| L6−IT | 46.97 | 10.98 | 7.07 | 5.16 | 3.90 | 2.99 | 2.42 | 1.93 | 1.47 | 1.00 |
| L6−CT | 48.69 | 11.29 | 7.11 | 5.10 | 3.91 | 3.03 | 2.38 | 1.89 | 1.41 | 1.00 |
| L56−NP | 33.07 | 7.14 | 4.87 | 3.69 | 2.95 | 2.41 | 2.08 | 1.73 | 1.38 | 1.00 |
| L5−IT | 42.89 | 9.63 | 6.13 | 4.30 | 3.39 | 2.68 | 2.22 | 1.81 | 1.46 | 1.00 |
| L5−ET | 44.94 | 9.86 | 6.17 | 4.47 | 3.39 | 2.70 | 2.16 | 1.71 | 1.34 | 1.00 |
| L4−IT | 38.48 | 8.77 | 5.52 | 3.96 | 3.06 | 2.55 | 2.04 | 1.70 | 1.37 | 1.00 |
| L23−IT | 44.69 | 10.64 | 6.65 | 4.65 | 3.63 | 2.91 | 2.30 | 1.81 | 1.47 | 1.00 |
| HIP−Misc2 | 40.80 | 9.39 | 5.97 | 4.28 | 3.30 | 2.60 | 2.12 | 1.78 | 1.49 | 1.00 |
| HIP−Misc1 | 44.66 | 10.20 | 6.28 | 4.40 | 3.43 | 2.66 | 2.02 | 1.75 | 1.39 | 1.00 |
| DG | 56.64 | 14.27 | 8.73 | 6.13 | 4.44 | 3.26 | 2.54 | 2.01 | 1.54 | 1.00 |
| CA3 | 58.91 | 14.28 | 8.89 | 6.08 | 4.37 | 3.37 | 2.53 | 2.03 | 1.48 | 1.00 |
| CA1 | 48.82 | 11.81 | 7.69 | 5.44 | 3.83 | 3.04 | 2.36 | 1.94 | 1.43 | 1.00 |
| Amy−Exc | 41.58 | 10.05 | 6.73 | 4.84 | 3.70 | 2.86 | 2.24 | 1.81 | 1.43 | 1.00 |
| Vip | 47.71 | 9.79 | 5.43 | 3.66 | 2.71 | 2.19 | 1.76 | 1.52 | 1.21 | 1.00 |
| THM−MB | 33.66 | 7.14 | 4.90 | 3.79 | 3.03 | 2.59 | 2.21 | 1.77 | 1.42 | 1.00 |
| THM−Inh | 41.38 | 9.01 | 5.76 | 4.10 | 3.27 | 2.46 | 2.12 | 1.78 | 1.45 | 1.00 |
| THM−Exc | 61.00 | 14.53 | 8.81 | 6.20 | 4.48 | 3.45 | 2.52 | 2.16 | 1.60 | 1.00 |
| SubCtx−Cplx | 55.80 | 12.33 | 7.28 | 4.93 | 3.64 | 2.79 | 2.21 | 1.74 | 1.38 | 1.00 |
| Sst | 37.21 | 7.75 | 5.02 | 3.74 | 2.98 | 2.43 | 2.02 | 1.64 | 1.35 | 1.00 |
| Sncg | 51.03 | 10.45 | 5.95 | 4.22 | 3.20 | 2.56 | 2.04 | 1.71 | 1.41 | 1.00 |
| Pvalb−ChC | 46.06 | 9.52 | 5.95 | 4.16 | 3.33 | 2.67 | 2.24 | 1.73 | 1.46 | 1.00 |
| Pvalb | 37.93 | 7.72 | 4.82 | 3.50 | 2.74 | 2.29 | 1.90 | 1.60 | 1.27 | 1.00 |
| PN | 58.01 | 12.63 | 6.78 | 4.71 | 3.43 | 2.66 | 2.03 | 1.66 | 1.37 | 1.00 |
| MSN−D2 | 29.56 | 6.88 | 4.67 | 3.57 | 2.88 | 2.42 | 2.03 | 1.70 | 1.45 | 1.00 |
| MSN−D1 | 28.78 | 6.61 | 4.48 | 3.40 | 2.77 | 2.27 | 1.96 | 1.61 | 1.33 | 1.00 |
| Lamp5−Lhx6 | 53.12 | 10.91 | 6.04 | 4.10 | 3.07 | 2.34 | 1.80 | 1.46 | 1.16 | 1.00 |
| Lamp5 | 57.11 | 11.62 | 6.61 | 4.40 | 3.24 | 2.56 | 1.97 | 1.57 | 1.33 | 1.00 |
| Foxp2 | 38.09 | 8.58 | 5.49 | 3.87 | 2.96 | 2.46 | 2.01 | 1.65 | 1.33 | 1.00 |
| Chd7 | 53.98 | 11.19 | 6.41 | 4.26 | 3.13 | 2.52 | 2.00 | 1.67 | 1.32 | 1.00 |
| CB | 60.31 | 13.40 | 7.66 | 5.22 | 3.93 | 2.87 | 2.47 | 1.93 | 1.44 | 1.00 |
| VLMC | 51.26 | 11.04 | 6.67 | 4.68 | 3.68 | 2.94 | 2.31 | 1.80 | 1.37 | 1.00 |
| PC | 44.84 | 9.65 | 6.23 | 4.42 | 3.46 | 2.72 | 2.38 | 1.83 | 1.36 | 1.00 |
| OPC | 116.23 | 21.07 | 10.06 | 6.22 | 4.31 | 3.09 | 2.50 | 1.98 | 1.49 | 1.00 |
| ODC | 92.11 | 18.06 | 9.63 | 5.97 | 4.27 | 3.24 | 2.41 | 1.92 | 1.32 | 1.00 |
| MGC | 27.25 | 6.14 | 4.18 | 3.40 | 2.87 | 2.40 | 2.02 | 1.68 | 1.36 | 1.00 |
| EC | 33.85 | 7.20 | 5.04 | 3.88 | 3.18 | 2.61 | 2.20 | 1.84 | 1.39 | 1.00 |
| ASC | 104.02 | 16.93 | 7.97 | 4.97 | 3.45 | 2.62 | 2.02 | 1.56 | 1.29 | 1.00 |

Percent: 0.01 0.02 0.03 0.04 0.05

### ASCT−Telencephalon astrocytes

| | 0.0−0.1 | 0.1−0.2 | 0.2−0.3 | 0.3−0.4 | 0.4−0.5 | 0.5−0.6 | 0.6−0.7 | 0.7−0.8 | 0.8−0.9 | 0.9−1.0 |
|---|---|---|---|---|---|---|---|---|---|---|
| L6b | 37.35 | 11.10 | 7.07 | 5.04 | 3.80 | 2.91 | 2.32 | 1.80 | 1.38 | 1.00 |
| L6−IT−Car3 | 33.79 | 10.10 | 6.60 | 4.82 | 3.72 | 2.96 | 2.38 | 1.91 | 1.45 | 1.00 |
| L6−IT | 35.44 | 10.71 | 6.99 | 5.07 | 3.86 | 3.01 | 2.43 | 1.89 | 1.46 | 1.00 |
| L6−CT | 36.87 | 10.83 | 7.03 | 5.15 | 3.87 | 3.01 | 2.36 | 1.81 | 1.42 | 1.00 |
| L56−NP | 24.68 | 6.91 | 4.75 | 3.64 | 2.99 | 2.44 | 2.07 | 1.74 | 1.42 | 1.00 |
| L5−IT | 30.81 | 8.98 | 5.74 | 4.19 | 3.27 | 2.59 | 2.12 | 1.72 | 1.38 | 1.00 |
| L5−ET | 36.36 | 10.32 | 6.53 | 4.66 | 3.58 | 2.83 | 2.30 | 1.82 | 1.41 | 1.00 |
| L4−IT | 28.77 | 8.41 | 5.41 | 3.96 | 3.12 | 2.58 | 2.16 | 1.78 | 1.39 | 1.00 |
| L23−IT | 34.52 | 10.28 | 6.72 | 4.80 | 3.78 | 2.99 | 2.40 | 1.91 | 1.49 | 1.00 |
| HIP−Misc2 | 28.84 | 8.50 | 5.43 | 4.03 | 3.14 | 2.55 | 2.06 | 1.70 | 1.41 | 1.00 |
| HIP−Misc1 | 32.57 | 9.47 | 6.15 | 4.35 | 3.30 | 2.61 | 2.10 | 1.74 | 1.37 | 1.00 |
| DG | 40.27 | 12.85 | 8.20 | 5.76 | 4.31 | 3.27 | 2.48 | 2.02 | 1.47 | 1.00 |
| CA3 | 42.92 | 13.22 | 8.38 | 5.84 | 4.43 | 3.32 | 2.54 | 1.96 | 1.47 | 1.00 |
| CA1 | 38.96 | 11.98 | 7.88 | 5.64 | 4.21 | 3.21 | 2.53 | 2.01 | 1.51 | 1.00 |
| Amy−Exc | 32.10 | 10.08 | 6.79 | 4.95 | 3.80 | 2.95 | 2.37 | 1.90 | 1.43 | 1.00 |
| Vip | 33.32 | 8.86 | 5.20 | 3.63 | 2.71 | 2.23 | 1.85 | 1.54 | 1.28 | 1.00 |
| THM−MB | 24.83 | 6.79 | 4.68 | 3.68 | 3.00 | 2.50 | 2.14 | 1.79 | 1.46 | 1.00 |
| THM−Inh | 26.48 | 7.55 | 4.89 | 3.66 | 2.89 | 2.33 | 1.94 | 1.59 | 1.28 | 1.00 |
| THM−Exc | 42.45 | 12.93 | 8.26 | 5.85 | 4.25 | 3.38 | 2.62 | 2.09 | 1.59 | 1.00 |
| SubCtx−Cplx | 39.29 | 11.25 | 6.89 | 4.71 | 3.51 | 2.72 | 2.15 | 1.71 | 1.36 | 1.00 |
| Sst | 26.18 | 7.04 | 4.59 | 3.44 | 2.78 | 2.31 | 1.97 | 1.62 | 1.33 | 1.00 |
| Sncg | 34.21 | 8.96 | 5.50 | 3.91 | 3.03 | 2.44 | 2.02 | 1.67 | 1.34 | 1.00 |
| Pvalb−ChC | 31.52 | 8.56 | 5.31 | 3.91 | 3.15 | 2.53 | 2.08 | 1.69 | 1.39 | 1.00 |
| Pvalb | 27.52 | 7.28 | 4.61 | 3.37 | 2.67 | 2.28 | 1.89 | 1.61 | 1.35 | 1.00 |
| PN | 40.97 | 11.45 | 6.63 | 4.58 | 3.36 | 2.60 | 2.01 | 1.66 | 1.34 | 1.00 |
| MSN−D2 | 21.57 | 6.41 | 4.47 | 3.42 | 2.79 | 2.34 | 1.97 | 1.66 | 1.40 | 1.00 |
| MSN−D1 | 20.15 | 5.96 | 4.07 | 3.18 | 2.58 | 2.14 | 1.84 | 1.53 | 1.28 | 1.00 |
| Lamp5−Lhx6 | 39.31 | 10.44 | 6.04 | 4.13 | 3.08 | 2.44 | 1.97 | 1.68 | 1.33 | 1.00 |
| Lamp5 | 39.42 | 10.30 | 6.12 | 4.17 | 3.08 | 2.54 | 1.96 | 1.63 | 1.30 | 1.00 |
| Foxp2 | 27.66 | 7.84 | 5.09 | 3.73 | 2.95 | 2.37 | 2.00 | 1.65 | 1.34 | 1.00 |
| Chd7 | 36.71 | 9.80 | 5.80 | 4.01 | 3.04 | 2.41 | 1.97 | 1.59 | 1.33 | 1.00 |
| CB | 41.23 | 11.88 | 7.04 | 4.82 | 3.69 | 2.89 | 2.42 | 1.86 | 1.44 | 1.00 |
| VLMC | 34.87 | 9.58 | 6.06 | 4.51 | 3.52 | 2.84 | 2.36 | 1.80 | 1.42 | 1.00 |
| PC | 30.93 | 8.61 | 5.81 | 4.31 | 3.44 | 2.79 | 2.33 | 1.84 | 1.46 | 1.00 |
| OPC | 67.35 | 16.29 | 8.50 | 5.62 | 3.93 | 3.02 | 2.38 | 1.78 | 1.38 | 1.00 |
| ODC | 52.75 | 13.74 | 7.80 | 5.15 | 3.82 | 2.87 | 2.31 | 1.80 | 1.36 | 1.00 |
| MGC | 18.35 | 5.31 | 3.80 | 3.06 | 2.72 | 2.30 | 1.98 | 1.66 | 1.38 | 1.00 |
| EC | 22.28 | 6.18 | 4.38 | 3.55 | 2.96 | 2.48 | 2.14 | 1.79 | 1.42 | 1.00 |
| ASC | 64.38 | 14.37 | 7.28 | 4.65 | 3.34 | 2.60 | 1.99 | 1.61 | 1.29 | 1.00 |

Percent: 0.02 0.04 0.06 0.08

### Amy−Exc−Glutamatergic neurons from amygdala

| | 0.0−0.1 | 0.1−0.2 | 0.2−0.3 | 0.3−0.4 | 0.4−0.5 | 0.5−0.6 | 0.6−0.7 | 0.7−0.8 | 0.8−0.9 | 0.9−1.0 |
|---|---|---|---|---|---|---|---|---|---|---|
| L6b | 36.62 | 8.90 | 5.29 | 3.57 | 2.65 | 2.04 | 1.66 | 1.28 | 1.10 | 1.00 |
| L6−IT−Car3 | 42.21 | 9.47 | 5.43 | 3.63 | 2.62 | 2.09 | 1.66 | 1.40 | 1.15 | 1.00 |
| L6−IT | 54.91 | 13.10 | 7.15 | 4.49 | 3.21 | 2.33 | 1.81 | 1.40 | 1.16 | 1.00 |
| L6−CT | 50.40 | 11.58 | 6.41 | 4.14 | 3.00 | 2.25 | 1.75 | 1.38 | 1.19 | 1.00 |
| L56−NP | 23.72 | 5.43 | 3.46 | 2.56 | 2.06 | 1.74 | 1.50 | 1.27 | 1.14 | 1.00 |
| L5−IT | 39.37 | 9.14 | 5.16 | 3.45 | 2.54 | 1.94 | 1.61 | 1.30 | 1.13 | 1.00 |
| L5−ET | 54.49 | 11.74 | 6.35 | 4.11 | 2.94 | 2.26 | 1.79 | 1.41 | 1.18 | 1.00 |
| L4−IT | 31.09 | 7.51 | 4.48 | 3.12 | 2.38 | 1.93 | 1.60 | 1.34 | 1.15 | 1.00 |
| L23−IT | 49.15 | 11.81 | 6.36 | 4.05 | 2.89 | 2.16 | 1.64 | 1.29 | 1.09 | 1.00 |
| HIP−Misc2 | 30.75 | 7.06 | 4.02 | 2.91 | 2.21 | 1.79 | 1.48 | 1.26 | 1.11 | 1.00 |
| HIP−Misc1 | 38.81 | 8.83 | 5.12 | 3.42 | 2.57 | 1.92 | 1.62 | 1.35 | 1.21 | 1.00 |
| DG | 38.47 | 10.33 | 6.29 | 4.40 | 3.09 | 2.37 | 1.79 | 1.49 | 1.14 | 1.00 |
| CA3 | 37.95 | 9.74 | 5.94 | 3.88 | 2.82 | 2.15 | 1.69 | 1.36 | 1.17 | 1.00 |
| CA1 | 53.99 | 13.53 | 7.55 | 4.87 | 3.36 | 2.34 | 1.77 | 1.40 | 1.12 | 1.00 |
| Amy−Exc | 62.67 | 16.28 | 9.15 | 5.54 | 3.74 | 2.67 | 1.98 | 1.53 | 1.22 | 1.00 |
| Vip | 16.58 | 3.89 | 2.48 | 1.89 | 1.54 | 1.33 | 1.17 | 1.10 | 1.01 | 1.00 |
| THM−MB | 16.82 | 3.74 | 2.56 | 2.07 | 1.75 | 1.52 | 1.37 | 1.26 | 1.15 | 1.00 |
| THM−Inh | 15.81 | 3.72 | 2.60 | 2.06 | 1.76 | 1.49 | 1.35 | 1.18 | 1.09 | 1.00 |
| THM−Exc | 20.99 | 5.27 | 3.56 | 2.66 | 2.07 | 1.75 | 1.50 | 1.26 | 1.15 | 1.00 |
| SubCtx−Cplx | 23.88 | 5.76 | 3.68 | 2.69 | 2.08 | 1.70 | 1.43 | 1.25 | 1.09 | 1.00 |
| Sst | 20.85 | 4.58 | 2.88 | 2.18 | 1.76 | 1.53 | 1.34 | 1.22 | 1.08 | 1.00 |
| Sncg | 17.88 | 3.97 | 2.68 | 2.05 | 1.71 | 1.48 | 1.35 | 1.21 | 1.05 | 1.00 |
| Pvalb−ChC | 21.08 | 4.68 | 3.13 | 2.35 | 1.94 | 1.64 | 1.45 | 1.30 | 1.18 | 1.00 |
| Pvalb | 19.95 | 4.33 | 2.72 | 2.10 | 1.71 | 1.48 | 1.33 | 1.19 | 1.10 | 1.00 |
| PN | 22.15 | 5.36 | 3.48 | 2.47 | 1.93 | 1.67 | 1.41 | 1.27 | 1.08 | 1.00 |
| MSN−D2 | 28.96 | 6.94 | 4.12 | 2.90 | 2.20 | 1.73 | 1.42 | 1.21 | 1.07 | 1.00 |
| MSN−D1 | 25.97 | 6.28 | 3.79 | 2.71 | 2.07 | 1.63 | 1.43 | 1.21 | 1.06 | 1.00 |
| Lamp5−Lhx6 | 21.72 | 4.87 | 3.00 | 2.21 | 1.74 | 1.49 | 1.31 | 1.19 | 1.11 | 1.00 |
| Lamp5 | 19.20 | 4.28 | 2.73 | 2.03 | 1.67 | 1.42 | 1.20 | 1.09 | 1.01 | 1.00 |
| Foxp2 | 28.01 | 6.35 | 3.60 | 2.55 | 1.94 | 1.58 | 1.31 | 1.13 | 1.07 | 1.00 |
| Chd7 | 18.43 | 4.17 | 2.74 | 2.11 | 1.69 | 1.46 | 1.29 | 1.15 | 1.04 | 1.00 |
| CB | 17.93 | 4.65 | 3.17 | 2.43 | 2.01 | 1.74 | 1.50 | 1.35 | 1.18 | 1.00 |
| VLMC | 13.57 | 3.30 | 2.37 | 1.94 | 1.74 | 1.54 | 1.35 | 1.20 | 1.12 | 1.00 |
| PC | 14.03 | 3.45 | 2.54 | 2.08 | 1.78 | 1.56 | 1.42 | 1.26 | 1.11 | 1.00 |
| OPC | 16.53 | 3.92 | 2.62 | 2.05 | 1.75 | 1.46 | 1.31 | 1.19 | 1.08 | 1.00 |
| ODC | 16.04 | 3.89 | 2.68 | 2.11 | 1.80 | 1.53 | 1.38 | 1.20 | 1.08 | 1.00 |
| MGC | 9.05 | 2.07 | 1.68 | 1.45 | 1.37 | 1.26 | 1.18 | 1.15 | 1.06 | 1.00 |
| EC | 10.10 | 2.32 | 1.85 | 1.64 | 1.41 | 1.35 | 1.23 | 1.16 | 1.09 | 1.00 |
| ASC | 14.77 | 3.39 | 2.30 | 1.81 | 1.55 | 1.40 | 1.28 | 1.16 | 1.04 | 1.00 |

Percent 0.01 0.02 0.03 0.04 0.05

### BFEXA−GABAergic neurons from basal forebrain and amygdala

| | 0.0−0.1 | 0.1−0.2 | 0.2−0.3 | 0.3−0.4 | 0.4−0.5 | 0.5−0.6 | 0.6−0.7 | 0.7−0.8 | 0.8−0.9 | 0.9−1.0 |
|---|---|---|---|---|---|---|---|---|---|---|
| L6b | 34.55 | 6.24 | 3.88 | 2.63 | 2.08 | 1.62 | 1.36 | 1.20 | 1.04 | 1.00 |
| L6−IT−Car3 | 32.73 | 5.60 | 3.40 | 2.50 | 1.92 | 1.68 | 1.46 | 1.25 | 1.12 | 1.00 |
| L6−IT | 35.62 | 6.36 | 3.85 | 2.89 | 2.20 | 1.81 | 1.54 | 1.33 | 1.08 | 1.00 |
| L6−CT | 38.10 | 6.43 | 3.79 | 2.74 | 2.13 | 1.71 | 1.40 | 1.21 | 1.06 | 1.00 |
| L56−NP | 26.72 | 4.16 | 2.67 | 1.98 | 1.67 | 1.44 | 1.28 | 1.19 | 1.07 | 1.00 |
| L5−IT | 31.58 | 5.30 | 3.39 | 2.35 | 1.87 | 1.56 | 1.35 | 1.24 | 1.16 | 1.00 |
| L5−ET | 43.98 | 7.28 | 4.05 | 2.86 | 2.34 | 1.77 | 1.51 | 1.25 | 1.09 | 1.00 |
| L4−IT | 24.71 | 4.25 | 2.71 | 2.02 | 1.71 | 1.39 | 1.31 | 1.16 | 1.06 | 1.00 |
| L23−IT | 35.87 | 6.61 | 3.83 | 2.76 | 2.12 | 1.81 | 1.43 | 1.19 | 1.09 | 1.00 |
| HIP−Misc2 | 25.77 | 4.28 | 2.67 | 2.10 | 1.68 | 1.48 | 1.23 | 1.17 | 1.10 | 1.00 |
| HIP−Misc1 | 28.26 | 4.97 | 3.13 | 2.23 | 1.80 | 1.46 | 1.29 | 1.13 | 1.04 | 1.00 |
| DG | 25.98 | 5.03 | 3.26 | 2.41 | 2.01 | 1.55 | 1.32 | 1.20 | 1.12 | 1.00 |
| CA3 | 27.76 | 5.24 | 3.47 | 2.48 | 1.96 | 1.54 | 1.33 | 1.19 | 1.05 | 1.00 |
| CA1 | 40.59 | 7.63 | 4.44 | 3.12 | 2.29 | 2.01 | 1.54 | 1.30 | 1.12 | 1.00 |
| Amy−Exc | 39.98 | 7.56 | 4.70 | 3.25 | 2.50 | 2.03 | 1.66 | 1.36 | 1.09 | 1.00 |
| Vip | 17.46 | 2.77 | 1.80 | 1.43 | 1.23 | 1.08 | 0.99 | 0.92 | 0.91 | 1.00 |
| THM−MB | 28.73 | 4.40 | 2.88 | 2.06 | 1.68 | 1.38 | 1.29 | 1.15 | 1.04 | 1.00 |
| THM−Inh | 22.84 | 3.78 | 2.51 | 1.99 | 1.70 | 1.47 | 1.28 | 1.23 | 1.06 | 1.00 |
| THM−Exc | 26.32 | 4.63 | 3.02 | 2.32 | 1.76 | 1.53 | 1.26 | 1.15 | 1.04 | 1.00 |
| SubCtx−Cplx | 30.11 | 5.19 | 3.11 | 2.18 | 1.71 | 1.39 | 1.19 | 1.02 | 0.93 | 1.00 |
| Sst | 25.13 | 3.83 | 2.40 | 1.78 | 1.39 | 1.20 | 1.09 | 1.03 | 0.95 | 1.00 |
| Sncg | 21.26 | 3.37 | 2.26 | 1.81 | 1.51 | 1.33 | 1.20 | 1.10 | 1.01 | 1.00 |
| Pvalb−ChC | 22.95 | 3.69 | 2.33 | 1.81 | 1.51 | 1.31 | 1.20 | 1.10 | 1.04 | 1.00 |
| Pvalb | 20.81 | 3.16 | 2.08 | 1.60 | 1.28 | 1.17 | 1.06 | 0.98 | 0.94 | 1.00 |
| PN | 20.31 | 3.40 | 2.36 | 1.80 | 1.46 | 1.30 | 1.16 | 1.04 | 1.04 | 1.00 |
| MSN−D2 | 47.31 | 7.84 | 4.03 | 2.42 | 1.77 | 1.35 | 1.17 | 0.99 | 0.93 | 1.00 |
| MSN−D1 | 41.73 | 6.90 | 3.76 | 2.51 | 1.81 | 1.44 | 1.18 | 1.01 | 0.95 | 1.00 |
| Lamp5−Lhx6 | 24.17 | 3.89 | 2.41 | 1.80 | 1.45 | 1.27 | 1.08 | 0.99 | 0.90 | 1.00 |
| Lamp5 | 23.19 | 3.76 | 2.31 | 1.77 | 1.46 | 1.25 | 1.04 | 1.03 | 0.94 | 1.00 |
| Foxp2 | 39.28 | 6.33 | 3.26 | 2.11 | 1.59 | 1.26 | 1.05 | 0.94 | 0.86 | 1.00 |
| Chd7 | 20.38 | 3.24 | 2.14 | 1.63 | 1.32 | 1.17 | 1.07 | 1.01 | 0.98 | 1.00 |
| CB | 17.40 | 3.05 | 2.15 | 1.76 | 1.55 | 1.35 | 1.23 | 1.16 | 1.06 | 1.00 |
| VLMC | 15.21 | 2.52 | 1.83 | 1.54 | 1.40 | 1.26 | 1.16 | 1.10 | 1.08 | 1.00 |
| PC | 15.69 | 2.60 | 1.95 | 1.67 | 1.37 | 1.27 | 1.23 | 1.13 | 1.08 | 1.00 |
| OPC | 20.41 | 3.44 | 2.25 | 1.89 | 1.57 | 1.35 | 1.26 | 1.18 | 1.06 | 1.00 |
| ODC | 18.63 | 3.29 | 2.15 | 1.76 | 1.49 | 1.33 | 1.15 | 1.10 | 1.01 | 1.00 |
| MGC | 11.45 | 1.69 | 1.33 | 1.24 | 1.13 | 1.06 | 1.08 | 1.01 | 0.94 | 1.00 |
| EC | 12.44 | 1.88 | 1.52 | 1.41 | 1.23 | 1.19 | 1.13 | 1.06 | 1.07 | 1.00 |
| ASC | 19.29 | 3.22 | 2.08 | 1.67 | 1.46 | 1.29 | 1.17 | 1.13 | 1.05 | 1.00 |

Percent 0.005 0.010 0.015 0.020 0.025

### BNGA−GABAergic neurons from basal nuclei

| | 0.0−0.1 | 0.1−0.2 | 0.2−0.3 | 0.3−0.4 | 0.4−0.5 | 0.5−0.6 | 0.6−0.7 | 0.7−0.8 | 0.8−0.9 | 0.9−1.0 |
|---|---|---|---|---|---|---|---|---|---|---|
| L6b | 43.53 | 6.78 | 4.66 | 3.23 | 2.60 | 2.18 | 1.82 | 1.55 | 1.23 | 1.00 |
| L6−IT−Car3 | 58.77 | 8.99 | 5.53 | 3.90 | 3.11 | 2.43 | 2.12 | 1.58 | 1.28 | 1.00 |
| L6−IT | 71.05 | 11.61 | 7.02 | 4.95 | 3.76 | 3.07 | 2.56 | 1.95 | 1.64 | 1.00 |
| L6−CT | 55.51 | 8.49 | 5.34 | 3.79 | 2.88 | 2.37 | 2.02 | 1.65 | 1.28 | 1.00 |
| L56−NP | 35.72 | 5.04 | 3.32 | 2.65 | 2.09 | 1.90 | 1.55 | 1.34 | 1.14 | 1.00 |
| L5−IT | 48.13 | 7.19 | 4.60 | 3.30 | 2.53 | 2.14 | 1.85 | 1.50 | 1.19 | 1.00 |
| L5−ET | 62.70 | 9.39 | 5.66 | 3.72 | 2.96 | 2.38 | 1.71 | 1.44 | 1.24 | 1.00 |
| L4−IT | 43.46 | 6.56 | 4.49 | 3.27 | 2.61 | 2.17 | 1.90 | 1.63 | 1.37 | 1.00 |
| L23−IT | 55.41 | 9.21 | 5.50 | 3.71 | 2.80 | 2.25 | 1.87 | 1.54 | 1.33 | 1.00 |
| HIP−Misc2 | 40.44 | 6.29 | 4.00 | 3.08 | 2.26 | 2.00 | 1.65 | 1.44 | 1.24 | 1.00 |
| HIP−Misc1 | 43.43 | 6.67 | 4.32 | 3.20 | 2.37 | 2.13 | 1.62 | 1.33 | 1.24 | 1.00 |
| DG | 48.35 | 8.18 | 5.57 | 3.98 | 2.95 | 2.42 | 1.84 | 1.47 | 1.29 | 1.00 |
| CA3 | 53.07 | 9.26 | 5.94 | 4.15 | 3.12 | 2.55 | 2.10 | 1.80 | 1.52 | 1.00 |
| CA1 | 64.57 | 10.83 | 6.69 | 4.16 | 3.22 | 2.65 | 2.01 | 1.53 | 1.27 | 1.00 |
| Amy−Exc | 68.84 | 12.09 | 7.23 | 4.87 | 3.78 | 2.80 | 2.15 | 1.72 | 1.23 | 1.00 |
| Vip | 28.83 | 4.31 | 2.85 | 2.17 | 1.74 | 1.51 | 1.30 | 1.23 | 1.14 | 1.00 |
| THM−MB | 39.20 | 5.27 | 3.69 | 2.97 | 2.38 | 1.93 | 1.65 | 1.44 | 1.28 | 1.00 |
| THM−Inh | 33.87 | 5.11 | 3.33 | 2.51 | 2.05 | 1.78 | 1.68 | 1.45 | 1.09 | 1.00 |
| THM−Exc | 41.20 | 6.75 | 4.43 | 3.19 | 2.63 | 1.85 | 1.82 | 1.33 | 1.32 | 1.00 |
| SubCtx−Cplx | 49.99 | 7.73 | 4.84 | 3.42 | 2.61 | 2.10 | 1.63 | 1.39 | 1.16 | 1.00 |
| Sst | 38.82 | 5.42 | 3.48 | 2.55 | 2.09 | 1.73 | 1.57 | 1.35 | 1.20 | 1.00 |
| Sncg | 27.83 | 3.80 | 2.60 | 2.05 | 1.81 | 1.56 | 1.42 | 1.29 | 1.16 | 1.00 |
| Pvalb−ChC | 38.03 | 5.40 | 3.53 | 2.82 | 2.27 | 1.96 | 1.62 | 1.57 | 1.33 | 1.00 |
| Pvalb | 35.44 | 4.92 | 3.08 | 2.30 | 2.01 | 1.70 | 1.45 | 1.36 | 1.20 | 1.00 |
| PN | 35.92 | 5.35 | 3.46 | 2.61 | 1.85 | 1.72 | 1.45 | 1.25 | 1.22 | 1.00 |
| MSN−D2 | 122.32 | 18.34 | 8.36 | 4.87 | 3.24 | 2.36 | 1.76 | 1.45 | 1.17 | 1.00 |
| MSN−D1 | 85.50 | 12.70 | 6.33 | 4.10 | 2.90 | 2.21 | 1.64 | 1.30 | 1.20 | 1.00 |
| Lamp5−Lhx6 | 31.36 | 4.36 | 3.05 | 2.25 | 1.84 | 1.65 | 1.47 | 1.25 | 1.07 | 1.00 |
| Lamp5 | 27.63 | 4.02 | 2.70 | 2.01 | 1.57 | 1.46 | 1.26 | 1.24 | 1.02 | 1.00 |
| Foxp2 | 84.07 | 11.95 | 5.69 | 3.46 | 2.34 | 1.78 | 1.47 | 1.21 | 1.06 | 1.00 |
| Chd7 | 31.76 | 4.52 | 3.03 | 2.35 | 1.93 | 1.66 | 1.45 | 1.24 | 1.16 | 1.00 |
| CB | 29.79 | 4.92 | 3.42 | 2.75 | 2.20 | 1.86 | 1.74 | 1.48 | 1.26 | 1.00 |
| VLMC | 23.11 | 3.53 | 2.54 | 2.08 | 1.73 | 1.68 | 1.48 | 1.35 | 1.26 | 1.00 |
| PC | 20.57 | 3.01 | 2.18 | 1.93 | 1.62 | 1.55 | 1.35 | 1.19 | 1.10 | 1.00 |
| OPC | 27.88 | 4.16 | 2.88 | 2.24 | 2.01 | 1.62 | 1.42 | 1.32 | 1.24 | 1.00 |
| ODC | 26.40 | 4.11 | 2.75 | 2.19 | 1.86 | 1.61 | 1.36 | 1.33 | 1.19 | 1.00 |
| MGC | 18.67 | 2.51 | 2.09 | 1.86 | 1.67 | 1.63 | 1.39 | 1.36 | 1.17 | 1.00 |
| EC | 17.54 | 2.48 | 1.93 | 1.76 | 1.54 | 1.49 | 1.34 | 1.24 | 1.06 | 1.00 |
| ASC | 26.61 | 3.87 | 2.76 | 2.13 | 1.87 | 1.63 | 1.54 | 1.38 | 1.22 | 1.00 |

Percent 0.003 0.006 0.009 0.012

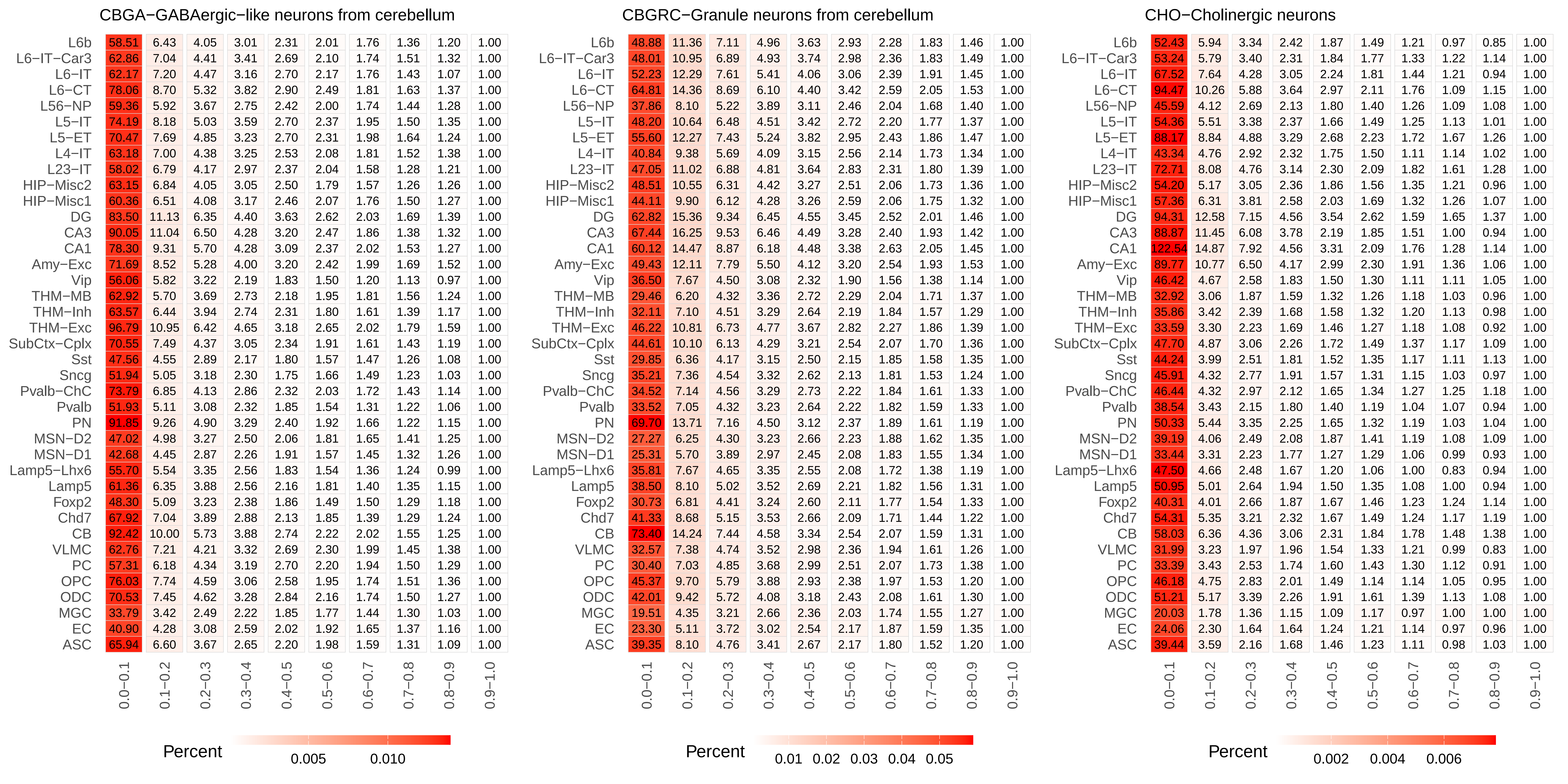
CBGA−GABAergic−like neurons from cerebellum
L6b 58.51 6.43 4.05 3.01 2.31 2.01 1.76 1.36 1.20 1.00
L6−IT−Car3 62.86 7.04 4.41 3.41 2.69 2.10 1.74 1.51 1.32 1.00
L6−IT 62.17 7.20 4.47 3.16 2.70 2.17 1.76 1.43 1.07 1.00
L6−CT 78.06 8.70 5.32 3.82 2.90 2.49 1.81 1.63 1.37 1.00
L56−NP 59.36 5.92 3.67 2.75 2.42 2.00 1.74 1.44 1.28 1.00
L5−IT 74.19 8.18 5.03 3.59 2.70 2.37 1.95 1.50 1.35 1.00
L5−ET 70.47 7.69 4.85 3.23 2.70 2.31 1.98 1.64 1.24 1.00
L4−IT 63.18 7.00 4.38 3.25 2.53 2.08 1.81 1.52 1.38 1.00
L23−IT 58.02 6.79 4.17 2.97 2.37 2.04 1.58 1.28 1.21 1.00
HIP−Misc2 63.15 6.84 4.05 3.05 2.50 1.79 1.57 1.26 1.26 1.00
HIP−Misc1 60.36 6.51 4.08 3.17 2.46 2.07 1.76 1.50 1.27 1.00
DG 83.50 11.13 6.35 4.40 3.63 2.62 2.03 1.69 1.39 1.00
CA3 90.05 11.04 6.50 4.28 3.20 2.47 1.86 1.38 1.32 1.00
CA1 78.30 9.31 5.70 4.28 3.09 2.37 2.02 1.53 1.27 1.00
Amy−Exc 71.69 8.52 5.28 4.00 3.20 2.42 1.99 1.69 1.52 1.00
Vip 56.06 5.82 3.22 2.19 1.83 1.50 1.20 1.13 0.97 1.00
THM−MB 62.92 5.70 3.69 2.73 2.18 1.95 1.81 1.56 1.24 1.00
THM−Inh 63.57 6.44 3.94 2.74 2.31 1.80 1.61 1.39 1.17 1.00
THM−Exc 96.79 10.95 6.42 4.65 3.18 2.65 2.02 1.79 1.59 1.00
SubCtx−Cplx 70.55 7.49 4.37 3.05 2.34 1.91 1.61 1.43 1.19 1.00
Sst 47.56 4.55 2.89 2.17 1.80 1.57 1.47 1.26 1.08 1.00
Sncg 51.94 5.05 3.18 2.30 1.75 1.66 1.49 1.23 1.03 1.00
Pvalb−ChC 73.79 6.85 4.13 2.86 2.32 2.03 1.72 1.43 1.14 1.00
Pvalb 51.93 5.11 3.08 2.32 1.85 1.54 1.31 1.22 1.06 1.00
PN 91.85 9.26 4.90 3.29 2.40 1.92 1.66 1.22 1.15 1.00
MSN−D2 47.02 4.98 3.27 2.50 2.06 1.81 1.65 1.41 1.25 1.00
MSN−D1 42.68 4.45 2.87 2.26 1.91 1.57 1.45 1.32 1.26 1.00
Lamp5−Lhx6 55.70 5.54 3.35 2.56 1.83 1.54 1.36 1.24 0.99 1.00
Lamp5 61.36 6.35 3.88 2.56 2.16 1.81 1.40 1.35 1.15 1.00
Foxp2 48.30 5.09 3.23 2.38 1.86 1.49 1.50 1.29 1.18 1.00
Chd7 67.92 7.04 3.89 2.88 2.13 1.85 1.39 1.29 1.24 1.00
CB 92.42 10.00 5.73 3.88 2.74 2.22 2.02 1.55 1.25 1.00
VLMC 62.76 7.21 4.21 3.32 2.69 2.30 1.99 1.45 1.38 1.00
PC 57.31 6.18 4.34 3.19 2.70 2.20 1.94 1.50 1.29 1.00
OPC 76.03 7.74 4.59 3.06 2.58 1.95 1.74 1.51 1.36 1.00
ODC 70.53 7.45 4.62 3.28 2.84 2.16 1.74 1.50 1.27 1.00
MGC 33.79 3.42 2.49 2.22 1.85 1.77 1.44 1.30 1.03 1.00
EC 40.90 4.28 3.08 2.59 2.02 1.92 1.65 1.37 1.16 1.00
ASC 65.94 6.60 3.67 2.65 2.20 1.98 1.59 1.31 1.09 1.00
0.0−0.1 0.1−0.2 0.2−0.3 0.3−0.4 0.4−0.5 0.5−0.6 0.6−0.7 0.7−0.8 0.8−0.9 0.9−1.0
Percent 0.005 0.010
CBGRC−Granule neurons from cerebellum
L6b 48.88 11.36 7.11 4.96 3.63 2.93 2.28 1.83 1.46 1.00
L6−IT−Car3 48.01 10.95 6.89 4.93 3.74 2.98 2.36 1.83 1.49 1.00
L6−IT 52.23 12.29 7.61 5.41 4.06 3.06 2.39 1.91 1.45 1.00
L6−CT 64.81 14.36 8.69 6.10 4.40 3.42 2.59 2.05 1.53 1.00
L56−NP 37.86 8.10 5.22 3.89 3.11 2.46 2.04 1.68 1.40 1.00
L5−IT 48.20 10.64 6.48 4.51 3.42 2.72 2.20 1.77 1.37 1.00
L5−ET 55.60 12.27 7.43 5.24 3.82 2.95 2.43 1.86 1.47 1.00
L4−IT 40.84 9.38 5.69 4.09 3.15 2.56 2.14 1.73 1.34 1.00
L23−IT 47.05 11.02 6.88 4.81 3.64 2.83 2.31 1.80 1.39 1.00
HIP−Misc2 48.51 10.55 6.31 4.42 3.27 2.51 2.06 1.73 1.36 1.00
HIP−Misc1 44.11 9.90 6.12 4.28 3.26 2.59 2.06 1.75 1.32 1.00
DG 62.82 15.36 9.34 6.45 4.55 3.45 2.52 2.01 1.46 1.00
CA3 67.44 16.25 9.53 6.46 4.49 3.28 2.40 1.93 1.42 1.00
CA1 60.12 14.47 8.87 6.18 4.48 3.38 2.63 2.05 1.45 1.00
Amy−Exc 49.43 12.11 7.79 5.50 4.12 3.20 2.54 1.93 1.53 1.00
Vip 36.50 7.67 4.50 3.08 2.32 1.90 1.56 1.38 1.14 1.00
THM−MB 29.46 6.20 4.32 3.36 2.72 2.29 2.04 1.71 1.37 1.00
THM−Inh 32.11 7.10 4.51 3.29 2.64 2.19 1.84 1.57 1.29 1.00
THM−Exc 46.22 10.81 6.73 4.77 3.67 2.82 2.27 1.86 1.39 1.00
SubCtx−Cplx 44.61 10.10 6.13 4.29 3.21 2.54 2.07 1.70 1.36 1.00
Sst 29.85 6.36 4.17 3.15 2.50 2.15 1.85 1.58 1.35 1.00
Sncg 35.21 7.36 4.54 3.32 2.62 2.13 1.81 1.53 1.24 1.00
Pvalb−ChC 34.52 7.14 4.56 3.29 2.73 2.22 1.84 1.61 1.33 1.00
Pvalb 33.52 7.05 4.32 3.23 2.64 2.22 1.82 1.59 1.33 1.00
PN 69.70 13.71 7.16 4.50 3.12 2.37 1.89 1.61 1.19 1.00
MSN−D2 27.27 6.25 4.30 3.23 2.66 2.23 1.88 1.62 1.35 1.00
MSN−D1 25.31 5.70 3.89 2.97 2.45 2.08 1.83 1.55 1.34 1.00
Lamp5−Lhx6 35.81 7.67 4.65 3.35 2.55 2.08 1.72 1.38 1.19 1.00
Lamp5 38.50 8.10 5.02 3.52 2.69 2.21 1.82 1.56 1.31 1.00
Foxp2 30.73 6.81 4.41 3.24 2.60 2.11 1.77 1.54 1.33 1.00
Chd7 41.33 8.68 5.15 3.53 2.66 2.09 1.71 1.44 1.22 1.00
CB 73.40 14.24 7.44 4.58 3.34 2.54 2.07 1.59 1.31 1.00
VLMC 32.57 7.38 4.74 3.52 2.98 2.36 1.94 1.61 1.26 1.00
PC 30.40 7.03 4.85 3.68 2.99 2.51 2.07 1.73 1.38 1.00
OPC 45.37 9.70 5.79 3.88 2.93 2.38 1.97 1.53 1.20 1.00
ODC 42.01 9.42 5.72 4.08 3.18 2.43 2.08 1.61 1.30 1.00
MGC 19.51 4.35 3.21 2.66 2.36 2.03 1.74 1.55 1.27 1.00
EC 23.30 5.11 3.72 3.02 2.54 2.17 1.87 1.59 1.35 1.00
ASC 39.35 8.10 4.76 3.41 2.67 2.17 1.80 1.52 1.20 1.00
0.0−0.1 0.1−0.2 0.2−0.3 0.3−0.4 0.4−0.5 0.5−0.6 0.6−0.7 0.7−0.8 0.8−0.9 0.9−1.0
Percent 0.01 0.02 0.03 0.04 0.05
CHO−Cholinergic neurons
L6b 52.43 5.94 3.34 2.42 1.87 1.49 1.21 0.97 0.85 1.00
L6−IT−Car3 53.24 5.79 3.40 2.31 1.84 1.77 1.33 1.22 1.14 1.00
L6−IT 67.52 7.64 4.28 3.05 2.24 1.81 1.44 1.21 0.94 1.00
L6−CT 94.47 10.26 5.88 3.64 2.97 2.11 1.76 1.09 1.15 1.00
L56−NP 45.59 4.12 2.69 2.13 1.80 1.40 1.26 1.09 1.08 1.00
L5−IT 54.36 5.51 3.38 2.37 1.66 1.49 1.25 1.13 1.01 1.00
L5−ET 88.17 8.84 4.88 3.29 2.68 2.23 1.72 1.67 1.26 1.00
L4−IT 43.34 4.76 2.92 2.32 1.75 1.50 1.11 1.14 1.02 1.00
L23−IT 72.71 8.08 4.76 3.14 2.30 2.09 1.82 1.61 1.28 1.00
HIP−Misc2 54.20 5.17 3.05 2.36 1.86 1.56 1.35 1.21 0.96 1.00
HIP−Misc1 57.36 6.31 3.81 2.58 2.03 1.69 1.32 1.26 1.07 1.00
DG 94.31 12.58 7.15 4.56 3.54 2.62 1.59 1.65 1.37 1.00
CA3 88.87 11.45 6.08 3.78 2.19 1.85 1.51 1.00 0.94 1.00
CA1 122.54 14.87 7.92 4.56 3.31 2.09 1.76 1.28 1.14 1.00
Amy−Exc 89.77 10.77 6.50 4.17 2.99 2.30 1.91 1.36 1.06 1.00
Vip 46.42 4.67 2.58 1.83 1.50 1.30 1.11 1.11 1.05 1.00
THM−MB 32.92 3.06 1.87 1.59 1.32 1.26 1.18 1.03 0.96 1.00
THM−Inh 35.86 3.42 2.39 1.68 1.58 1.32 1.20 1.13 0.98 1.00
THM−Exc 33.59 3.30 2.23 1.69 1.46 1.27 1.18 1.08 0.92 1.00
SubCtx−Cplx 47.70 4.87 3.06 2.26 1.72 1.49 1.37 1.17 1.09 1.00
Sst 44.24 3.99 2.51 1.81 1.52 1.35 1.17 1.11 1.13 1.00
Sncg 45.91 4.32 2.77 1.91 1.57 1.31 1.15 1.03 0.97 1.00
Pvalb−ChC 46.44 4.32 2.97 2.12 1.65 1.34 1.27 1.25 1.18 1.00
Pvalb 38.54 3.43 2.15 1.80 1.40 1.19 1.04 1.07 0.94 1.00
PN 50.33 5.44 3.35 2.25 1.65 1.32 1.19 1.03 1.04 1.00
MSN−D2 39.19 4.06 2.49 2.08 1.87 1.41 1.19 1.08 1.09 1.00
MSN−D1 33.44 3.31 2.23 1.77 1.27 1.29 1.06 0.99 0.93 1.00
Lamp5−Lhx6 47.50 4.66 2.48 1.67 1.20 1.06 1.00 0.83 0.94 1.00
Lamp5 50.95 5.01 2.64 1.94 1.50 1.35 1.08 1.00 0.94 1.00
Foxp2 40.31 4.01 2.66 1.87 1.67 1.46 1.23 1.24 1.14 1.00
Chd7 54.31 5.35 3.21 2.32 1.67 1.49 1.24 1.17 1.19 1.00
CB 58.03 6.36 4.36 3.06 2.31 1.84 1.78 1.48 1.38 1.00
VLMC 31.99 3.23 1.97 1.96 1.54 1.33 1.21 0.99 0.83 1.00
PC 33.39 3.43 2.53 1.74 1.60 1.43 1.30 1.12 0.91 1.00
OPC 46.18 4.75 2.83 2.01 1.49 1.14 1.14 1.05 0.95 1.00
ODC 51.21 5.17 3.39 2.26 1.91 1.61 1.39 1.13 1.08 1.00
MGC 20.03 1.78 1.36 1.15 1.09 1.17 0.97 1.00 1.00 1.00
EC 24.06 2.30 1.64 1.64 1.24 1.21 1.14 0.97 0.96 1.00
ASC 39.44 3.59 2.16 1.68 1.46 1.23 1.11 0.98 1.03 1.00
0.0−0.1 0.1−0.2 0.2−0.3 0.3−0.4 0.4−0.5 0.5−0.6 0.6−0.7 0.7−0.8 0.8−0.9 0.9−1.0
Percent 0.002 0.004 0.006

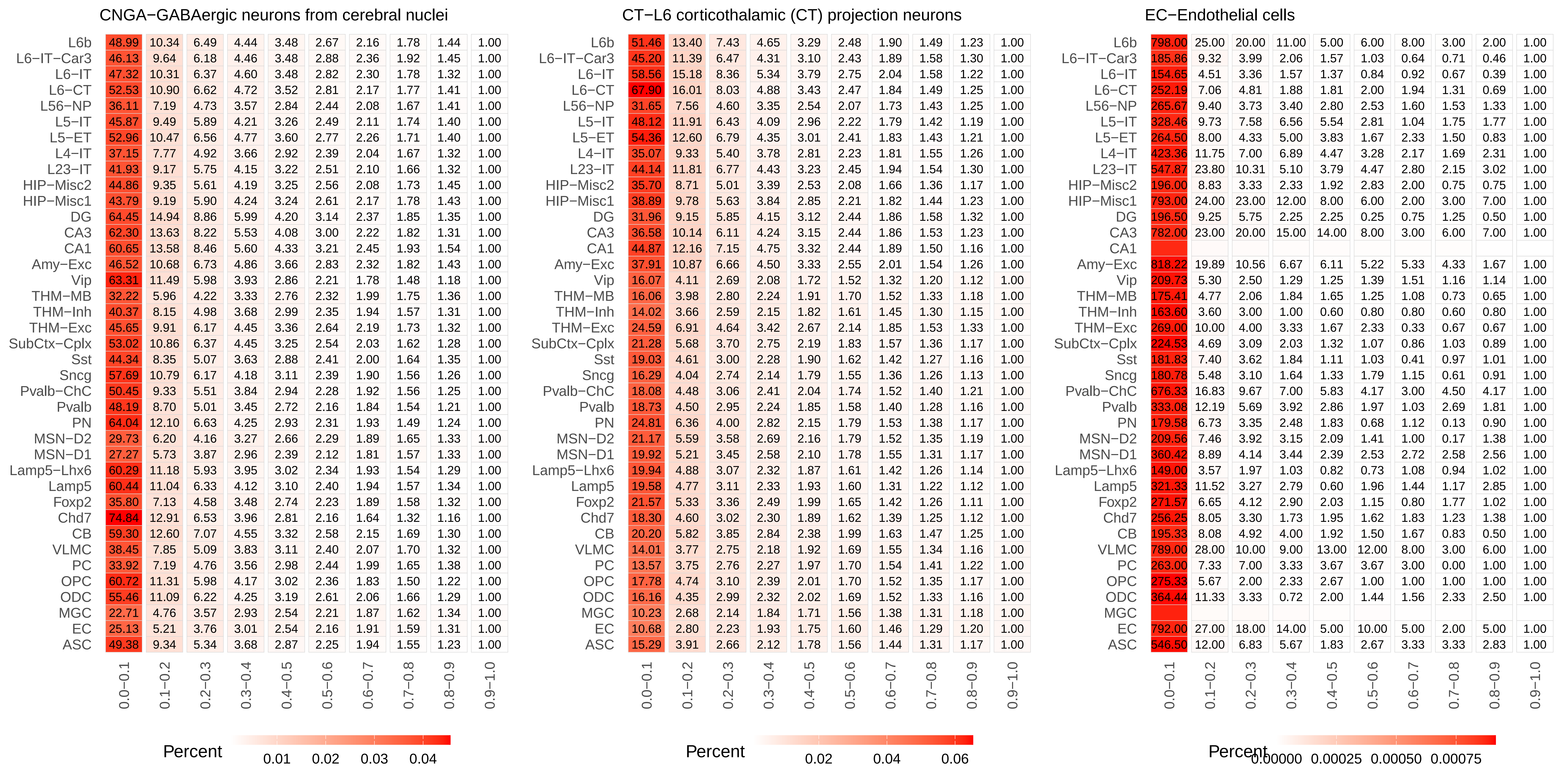
CNGA−GABAergic neurons from cerebral nuclei
CT−L6 corticothalamic (CT) projection neurons
EC−Endothelial cells
L6b
L6−IT−Car3
L6−IT
L6−CT
L56−NP
L5−IT
L5−ET
L4−IT
L23−IT
HIP−Misc2
HIP−Misc1
DG
CA3
CA1
Amy−Exc
Vip
THM−MB
THM−Inh
THM−Exc
SubCtx−Cplx
Sst
Sncg
Pvalb−ChC
Pvalb
PN
MSN−D2
MSN−D1
Lamp5−Lhx6
Lamp5
Foxp2
Chd7
CB
VLMC
PC
OPC
ODC
MGC
EC
ASC
0.0−0.1
0.1−0.2
0.2−0.3
0.3−0.4
0.4−0.5
0.5−0.6
0.6−0.7
0.7−0.8
0.8−0.9
0.9−1.0
Percent
0.01 0.02 0.03 0.04
Percent
0.02 0.04 0.06
Percent
0.00000 0.00025 0.00050 0.00075

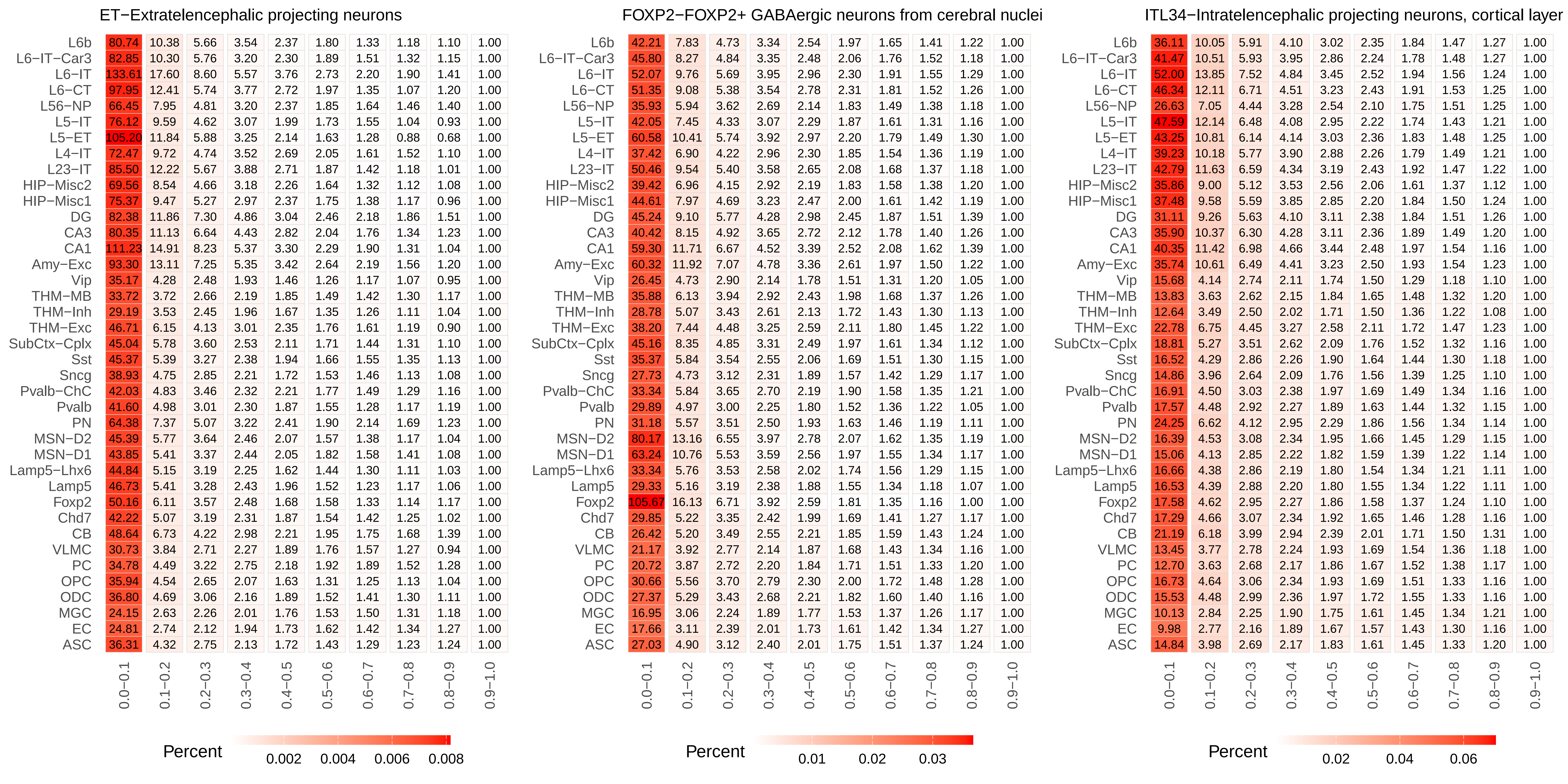

ET−Extratelencephalic projecting neurons
FOXP2−FOXP2+ GABAergic neurons from cerebral nuclei
ITL34−Intratelencephalic projecting neurons, cortical layer
L6b
L6−IT−Car3
L6−IT
L6−CT
L56−NP
L5−IT
L5−ET
L4−IT
L23−IT
HIP−Misc2
HIP−Misc1
DG
CA3
CA1
Amy−Exc
Vip
THM−MB
THM−Inh
THM−Exc
SubCtx−Cplx
Sst
Sncg
Pvalb−ChC
Pvalb
PN
MSN−D2
MSN−D1
Lamp5−Lhx6
Lamp5
Foxp2
Chd7
CB
VLMC
PC
OPC
ODC
MGC
EC
ASC
0.0−0.1
0.1−0.2
0.2−0.3
0.3−0.4
0.4−0.5
0.5−0.6
0.6−0.7
0.7−0.8
0.8−0.9
0.9−1.0
Percent
0.002
0.004
0.006
0.008
Percent
0.01
0.02
0.03
Percent
0.02
0.04
0.06

ITL45−Intratelencephalic projecting neurons, cortical layer 4/5 like

| | 0.0−0.1 | 0.1−0.2 | 0.2−0.3 | 0.3−0.4 | 0.4−0.5 | 0.5−0.6 | 0.6−0.7 | 0.7−0.8 | 0.8−0.9 | 0.9−1.0 |
|---|---|---|---|---|---|---|---|---|---|---|
| L6b | 36.94 | 9.47 | 5.51 | 3.78 | 2.74 | 2.11 | 1.70 | 1.39 | 1.20 | 1.00 |
| L6−IT−Car3 | 46.73 | 11.03 | 5.98 | 3.81 | 2.69 | 2.15 | 1.69 | 1.42 | 1.19 | 1.00 |
| L6−IT | 55.89 | 13.46 | 7.23 | 4.51 | 3.14 | 2.33 | 1.72 | 1.40 | 1.11 | 1.00 |
| L6−CT | 55.61 | 12.91 | 6.90 | 4.40 | 3.12 | 2.30 | 1.81 | 1.42 | 1.20 | 1.00 |
| L56−NP | 30.83 | 7.31 | 4.55 | 3.23 | 2.47 | 2.02 | 1.67 | 1.40 | 1.25 | 1.00 |
| L5−IT | 53.98 | 12.29 | 6.41 | 3.93 | 2.74 | 2.02 | 1.60 | 1.33 | 1.13 | 1.00 |
| L5−ET | 46.23 | 10.51 | 5.95 | 4.02 | 2.81 | 2.28 | 1.74 | 1.42 | 1.23 | 1.00 |
| L4−IT | 39.36 | 9.68 | 5.42 | 3.66 | 2.61 | 2.05 | 1.60 | 1.37 | 1.19 | 1.00 |
| L23−IT | 45.98 | 11.59 | 6.57 | 4.23 | 3.05 | 2.34 | 1.80 | 1.43 | 1.17 | 1.00 |
| HIP−Misc2 | 38.68 | 8.92 | 4.94 | 3.32 | 2.49 | 1.94 | 1.52 | 1.33 | 1.10 | 1.00 |
| HIP−Misc1 | 38.44 | 9.29 | 5.36 | 3.69 | 2.65 | 2.07 | 1.69 | 1.38 | 1.23 | 1.00 |
| DG | 30.62 | 8.41 | 5.25 | 3.80 | 2.75 | 2.17 | 1.73 | 1.44 | 1.16 | 1.00 |
| CA3 | 36.58 | 9.75 | 5.80 | 3.83 | 2.88 | 2.23 | 1.65 | 1.39 | 1.09 | 1.00 |
| CA1 | 45.38 | 11.69 | 6.88 | 4.63 | 3.13 | 2.34 | 1.89 | 1.47 | 1.13 | 1.00 |
| Amy−Exc | 39.66 | 10.90 | 6.58 | 4.43 | 3.18 | 2.39 | 1.88 | 1.49 | 1.24 | 1.00 |
| Vip | 15.51 | 3.74 | 2.52 | 1.97 | 1.64 | 1.43 | 1.23 | 1.16 | 1.09 | 1.00 |
| THM−MB | 14.81 | 3.53 | 2.49 | 2.09 | 1.80 | 1.56 | 1.42 | 1.27 | 1.18 | 1.00 |
| THM−Inh | 13.27 | 3.32 | 2.40 | 1.95 | 1.64 | 1.44 | 1.33 | 1.20 | 1.06 | 1.00 |
| THM−Exc | 23.50 | 6.23 | 4.20 | 3.11 | 2.48 | 1.94 | 1.73 | 1.41 | 1.18 | 1.00 |
| SubCtx−Cplx | 18.69 | 4.74 | 3.22 | 2.38 | 1.93 | 1.63 | 1.39 | 1.24 | 1.11 | 1.00 |
| Sst | 17.63 | 4.13 | 2.71 | 2.09 | 1.75 | 1.53 | 1.36 | 1.23 | 1.14 | 1.00 |
| Sncg | 15.10 | 3.62 | 2.51 | 2.02 | 1.71 | 1.48 | 1.36 | 1.22 | 1.11 | 1.00 |
| Pvalb−ChC | 17.41 | 4.06 | 2.79 | 2.23 | 1.82 | 1.56 | 1.38 | 1.25 | 1.13 | 1.00 |
| Pvalb | 18.36 | 4.28 | 2.77 | 2.12 | 1.74 | 1.51 | 1.34 | 1.22 | 1.11 | 1.00 |
| PN | 23.50 | 5.89 | 3.73 | 2.74 | 2.05 | 1.74 | 1.49 | 1.32 | 1.14 | 1.00 |
| MSN−D2 | 18.51 | 4.61 | 3.04 | 2.32 | 1.89 | 1.61 | 1.40 | 1.23 | 1.09 | 1.00 |
| MSN−D1 | 16.97 | 4.27 | 2.88 | 2.22 | 1.80 | 1.54 | 1.37 | 1.18 | 1.10 | 1.00 |
| Lamp5−Lhx6 | 16.96 | 4.01 | 2.68 | 2.04 | 1.68 | 1.49 | 1.31 | 1.21 | 1.10 | 1.00 |
| Lamp5 | 16.49 | 3.95 | 2.64 | 2.04 | 1.71 | 1.48 | 1.26 | 1.16 | 1.09 | 1.00 |
| Foxp2 | 18.85 | 4.55 | 2.86 | 2.15 | 1.71 | 1.48 | 1.30 | 1.17 | 1.09 | 1.00 |
| Chd7 | 16.48 | 4.00 | 2.69 | 2.09 | 1.72 | 1.48 | 1.32 | 1.16 | 1.08 | 1.00 |
| CB | 20.11 | 5.45 | 3.62 | 2.72 | 2.24 | 1.87 | 1.63 | 1.46 | 1.26 | 1.00 |
| VLMC | 13.73 | 3.46 | 2.56 | 2.08 | 1.81 | 1.60 | 1.48 | 1.33 | 1.18 | 1.00 |
| PC | 13.21 | 3.42 | 2.49 | 2.08 | 1.81 | 1.62 | 1.44 | 1.32 | 1.15 | 1.00 |
| OPC | 16.23 | 4.09 | 2.77 | 2.14 | 1.76 | 1.53 | 1.38 | 1.29 | 1.14 | 1.00 |
| ODC | 15.88 | 4.10 | 2.85 | 2.25 | 1.87 | 1.60 | 1.50 | 1.32 | 1.16 | 1.00 |
| MGC | 10.63 | 2.70 | 2.13 | 1.83 | 1.66 | 1.54 | 1.37 | 1.31 | 1.19 | 1.00 |
| EC | 10.40 | 2.63 | 2.05 | 1.79 | 1.59 | 1.50 | 1.38 | 1.26 | 1.16 | 1.00 |
| ASC | 14.03 | 3.41 | 2.36 | 1.91 | 1.63 | 1.47 | 1.35 | 1.24 | 1.15 | 1.00 |

Percent 0.01 0.02 0.03 0.04 0.05

ITL5−Intratelencephalic projecting neurons, cortical layer 5

| | 0.0−0.1 | 0.1−0.2 | 0.2−0.3 | 0.3−0.4 | 0.4−0.5 | 0.5−0.6 | 0.6−0.7 | 0.7−0.8 | 0.8−0.9 | 0.9−1.0 |
|---|---|---|---|---|---|---|---|---|---|---|
| L6b | 28.19 | 8.97 | 5.35 | 3.71 | 2.76 | 2.17 | 1.71 | 1.41 | 1.19 | 1.00 |
| L6−IT−Car3 | 25.94 | 7.95 | 4.81 | 3.39 | 2.57 | 2.05 | 1.65 | 1.42 | 1.19 | 1.00 |
| L6−IT | 35.27 | 11.04 | 6.42 | 4.26 | 3.13 | 2.35 | 1.82 | 1.47 | 1.20 | 1.00 |
| L6−CT | 34.92 | 10.35 | 5.98 | 4.05 | 2.92 | 2.22 | 1.77 | 1.39 | 1.23 | 1.00 |
| L56−NP | 19.45 | 5.90 | 3.82 | 2.86 | 2.24 | 1.85 | 1.57 | 1.35 | 1.17 | 1.00 |
| L5−IT | 33.16 | 9.92 | 5.56 | 3.64 | 2.66 | 2.02 | 1.62 | 1.31 | 1.11 | 1.00 |
| L5−ET | 32.87 | 9.46 | 5.44 | 3.69 | 2.76 | 2.18 | 1.74 | 1.42 | 1.19 | 1.00 |
| L4−IT | 23.73 | 7.47 | 4.55 | 3.28 | 2.48 | 1.98 | 1.64 | 1.40 | 1.20 | 1.00 |
| L23−IT | 29.11 | 9.32 | 5.55 | 3.73 | 2.78 | 2.20 | 1.73 | 1.38 | 1.19 | 1.00 |
| HIP−Misc2 | 24.44 | 7.23 | 4.38 | 3.09 | 2.36 | 1.92 | 1.54 | 1.31 | 1.15 | 1.00 |
| HIP−Misc1 | 24.71 | 7.49 | 4.53 | 3.29 | 2.47 | 2.00 | 1.63 | 1.36 | 1.17 | 1.00 |
| DG | 20.17 | 7.02 | 4.65 | 3.50 | 2.72 | 2.13 | 1.72 | 1.49 | 1.27 | 1.00 |
| CA3 | 22.60 | 7.68 | 4.90 | 3.43 | 2.66 | 2.06 | 1.65 | 1.42 | 1.16 | 1.00 |
| CA1 | 28.65 | 9.46 | 5.86 | 4.05 | 2.95 | 2.29 | 1.77 | 1.42 | 1.16 | 1.00 |
| Amy−Exc | 25.83 | 8.90 | 5.66 | 3.93 | 2.97 | 2.36 | 1.86 | 1.48 | 1.23 | 1.00 |
| Vip | 10.82 | 3.33 | 2.30 | 1.79 | 1.53 | 1.36 | 1.19 | 1.12 | 1.06 | 1.00 |
| THM−MB | 10.50 | 3.21 | 2.32 | 1.92 | 1.65 | 1.46 | 1.34 | 1.22 | 1.12 | 1.00 |
| THM−Inh | 9.24 | 2.96 | 2.19 | 1.81 | 1.59 | 1.39 | 1.27 | 1.16 | 1.08 | 1.00 |
| THM−Exc | 15.88 | 5.45 | 3.74 | 2.81 | 2.26 | 1.85 | 1.58 | 1.39 | 1.20 | 1.00 |
| SubCtx−Cplx | 13.46 | 4.36 | 3.03 | 2.33 | 1.88 | 1.60 | 1.39 | 1.22 | 1.11 | 1.00 |
| Sst | 12.60 | 3.77 | 2.55 | 2.01 | 1.68 | 1.48 | 1.32 | 1.22 | 1.11 | 1.00 |
| Sncg | 11.09 | 3.42 | 2.36 | 1.87 | 1.60 | 1.41 | 1.30 | 1.18 | 1.07 | 1.00 |
| Pvalb−ChC | 12.21 | 3.69 | 2.56 | 2.01 | 1.69 | 1.49 | 1.34 | 1.23 | 1.11 | 1.00 |
| Pvalb | 12.61 | 3.75 | 2.50 | 1.94 | 1.65 | 1.45 | 1.30 | 1.22 | 1.10 | 1.00 |
| PN | 15.63 | 4.99 | 3.27 | 2.46 | 1.96 | 1.64 | 1.39 | 1.27 | 1.14 | 1.00 |
| MSN−D2 | 12.48 | 4.00 | 2.75 | 2.11 | 1.74 | 1.51 | 1.35 | 1.18 | 1.09 | 1.00 |
| MSN−D1 | 11.49 | 3.65 | 2.56 | 1.98 | 1.66 | 1.45 | 1.29 | 1.14 | 1.06 | 1.00 |
| Lamp5−Lhx6 | 12.96 | 3.89 | 2.61 | 1.97 | 1.64 | 1.44 | 1.28 | 1.17 | 1.09 | 1.00 |
| Lamp5 | 12.76 | 3.83 | 2.59 | 2.01 | 1.67 | 1.41 | 1.24 | 1.14 | 1.07 | 1.00 |
| Foxp2 | 13.26 | 4.04 | 2.68 | 2.05 | 1.71 | 1.47 | 1.29 | 1.17 | 1.08 | 1.00 |
| Chd7 | 11.93 | 3.66 | 2.51 | 1.97 | 1.67 | 1.45 | 1.29 | 1.17 | 1.09 | 1.00 |
| CB | 13.16 | 4.52 | 3.15 | 2.41 | 2.01 | 1.76 | 1.53 | 1.38 | 1.23 | 1.00 |
| VLMC | 9.17 | 2.98 | 2.20 | 1.88 | 1.65 | 1.47 | 1.38 | 1.24 | 1.14 | 1.00 |
| PC | 9.13 | 2.96 | 2.33 | 1.91 | 1.66 | 1.50 | 1.38 | 1.26 | 1.15 | 1.00 |
| OPC | 11.28 | 3.61 | 2.47 | 1.98 | 1.67 | 1.47 | 1.33 | 1.20 | 1.11 | 1.00 |
| ODC | 10.72 | 3.49 | 2.51 | 2.01 | 1.71 | 1.50 | 1.37 | 1.23 | 1.12 | 1.00 |
| MGC | 6.67 | 2.16 | 1.75 | 1.54 | 1.43 | 1.35 | 1.26 | 1.20 | 1.13 | 1.00 |
| EC | 7.07 | 2.25 | 1.81 | 1.61 | 1.48 | 1.39 | 1.30 | 1.20 | 1.10 | 1.00 |
| ASC | 9.97 | 3.13 | 2.23 | 1.82 | 1.56 | 1.41 | 1.29 | 1.22 | 1.12 | 1.00 |

Percent 0.02 0.04 0.06 0.08

ITL6_1−Intratelencephalic projecting neurons, cortical laye

| | 0.0−0.1 | 0.1−0.2 | 0.2−0.3 | 0.3−0.4 | 0.4−0.5 | 0.5−0.6 | 0.6−0.7 | 0.7−0.8 | 0.8−0.9 | 0.9−1.0 |
|---|---|---|---|---|---|---|---|---|---|---|
| L6b | 39.21 | 11.36 | 6.49 | 4.33 | 3.09 | 2.34 | 1.83 | 1.46 | 1.17 | 1.00 |
| L6−IT−Car3 | 43.52 | 11.53 | 6.35 | 4.05 | 2.86 | 2.21 | 1.72 | 1.44 | 1.20 | 1.00 |
| L6−IT | 57.20 | 15.78 | 8.20 | 5.07 | 3.46 | 2.46 | 1.88 | 1.47 | 1.16 | 1.00 |
| L6−CT | 47.70 | 12.88 | 6.97 | 4.47 | 3.12 | 2.32 | 1.79 | 1.42 | 1.20 | 1.00 |
| L56−NP | 23.58 | 6.65 | 4.25 | 3.16 | 2.47 | 2.04 | 1.73 | 1.44 | 1.23 | 1.00 |
| L5−IT | 42.10 | 11.53 | 6.25 | 3.99 | 2.81 | 2.10 | 1.70 | 1.35 | 1.11 | 1.00 |
| L5−ET | 46.51 | 12.20 | 6.62 | 4.26 | 3.03 | 2.32 | 1.84 | 1.43 | 1.21 | 1.00 |
| L4−IT | 32.78 | 9.23 | 5.41 | 3.70 | 2.72 | 2.14 | 1.74 | 1.45 | 1.25 | 1.00 |
| L23−IT | 44.43 | 12.80 | 7.10 | 4.60 | 3.25 | 2.43 | 1.88 | 1.51 | 1.24 | 1.00 |
| HIP−Misc2 | 29.79 | 8.29 | 4.86 | 3.39 | 2.52 | 1.96 | 1.66 | 1.38 | 1.15 | 1.00 |
| HIP−Misc1 | 34.84 | 9.41 | 5.36 | 3.67 | 2.71 | 2.04 | 1.72 | 1.40 | 1.20 | 1.00 |
| DG | 26.74 | 8.49 | 5.39 | 3.91 | 2.87 | 2.23 | 1.73 | 1.53 | 1.19 | 1.00 |
| CA3 | 29.96 | 9.34 | 5.71 | 3.90 | 2.95 | 2.31 | 1.78 | 1.48 | 1.23 | 1.00 |
| CA1 | 38.13 | 11.54 | 6.96 | 4.58 | 3.24 | 2.39 | 1.85 | 1.47 | 1.15 | 1.00 |
| Amy−Exc | 37.88 | 12.01 | 7.26 | 4.76 | 3.44 | 2.59 | 2.00 | 1.53 | 1.26 | 1.00 |
| Vip | 13.71 | 3.91 | 2.62 | 2.04 | 1.70 | 1.46 | 1.29 | 1.18 | 1.09 | 1.00 |
| THM−MB | 13.00 | 3.68 | 2.63 | 2.15 | 1.84 | 1.62 | 1.43 | 1.28 | 1.18 | 1.00 |
| THM−Inh | 12.41 | 3.68 | 2.61 | 2.15 | 1.84 | 1.56 | 1.45 | 1.31 | 1.16 | 1.00 |
| THM−Exc | 20.41 | 6.36 | 4.34 | 3.22 | 2.56 | 2.06 | 1.76 | 1.47 | 1.23 | 1.00 |
| SubCtx−Cplx | 18.24 | 5.45 | 3.63 | 2.71 | 2.14 | 1.81 | 1.55 | 1.33 | 1.17 | 1.00 |
| Sst | 16.67 | 4.58 | 3.02 | 2.33 | 1.94 | 1.65 | 1.45 | 1.30 | 1.17 | 1.00 |
| Sncg | 14.11 | 3.94 | 2.69 | 2.12 | 1.77 | 1.53 | 1.40 | 1.25 | 1.12 | 1.00 |
| Pvalb−ChC | 15.63 | 4.35 | 2.97 | 2.35 | 1.97 | 1.68 | 1.50 | 1.34 | 1.16 | 1.00 |
| Pvalb | 16.57 | 4.53 | 2.97 | 2.24 | 1.87 | 1.62 | 1.41 | 1.30 | 1.15 | 1.00 |
| PN | 19.30 | 5.66 | 3.64 | 2.69 | 2.06 | 1.76 | 1.49 | 1.31 | 1.16 | 1.00 |
| MSN−D2 | 18.47 | 5.41 | 3.55 | 2.63 | 2.11 | 1.76 | 1.49 | 1.29 | 1.12 | 1.00 |
| MSN−D1 | 16.65 | 4.93 | 3.26 | 2.45 | 1.95 | 1.66 | 1.43 | 1.23 | 1.07 | 1.00 |
| Lamp5−Lhx6 | 16.92 | 4.65 | 3.04 | 2.29 | 1.82 | 1.57 | 1.36 | 1.22 | 1.10 | 1.00 |
| Lamp5 | 16.27 | 4.52 | 2.93 | 2.25 | 1.83 | 1.56 | 1.32 | 1.21 | 1.10 | 1.00 |
| Foxp2 | 18.61 | 5.17 | 3.22 | 2.38 | 1.90 | 1.60 | 1.37 | 1.22 | 1.08 | 1.00 |
| Chd7 | 15.08 | 4.25 | 2.85 | 2.19 | 1.82 | 1.55 | 1.37 | 1.24 | 1.10 | 1.00 |
| CB | 16.26 | 5.19 | 3.52 | 2.63 | 2.20 | 1.89 | 1.60 | 1.46 | 1.25 | 1.00 |
| VLMC | 11.78 | 3.57 | 2.57 | 2.12 | 1.87 | 1.66 | 1.52 | 1.29 | 1.18 | 1.00 |
| PC | 11.51 | 3.52 | 2.63 | 2.17 | 1.88 | 1.64 | 1.50 | 1.34 | 1.15 | 1.00 |
| OPC | 14.98 | 4.43 | 2.99 | 2.31 | 1.97 | 1.67 | 1.50 | 1.36 | 1.18 | 1.00 |
| ODC | 13.76 | 4.12 | 2.91 | 2.28 | 1.97 | 1.66 | 1.50 | 1.31 | 1.18 | 1.00 |
| MGC | 8.46 | 2.51 | 2.02 | 1.74 | 1.61 | 1.47 | 1.35 | 1.25 | 1.14 | 1.00 |
| EC | 8.84 | 2.61 | 2.04 | 1.81 | 1.64 | 1.51 | 1.40 | 1.29 | 1.13 | 1.00 |
| ASC | 12.92 | 3.71 | 2.57 | 2.07 | 1.75 | 1.54 | 1.41 | 1.30 | 1.15 | 1.00 |

Percent 0.02 0.04 0.06

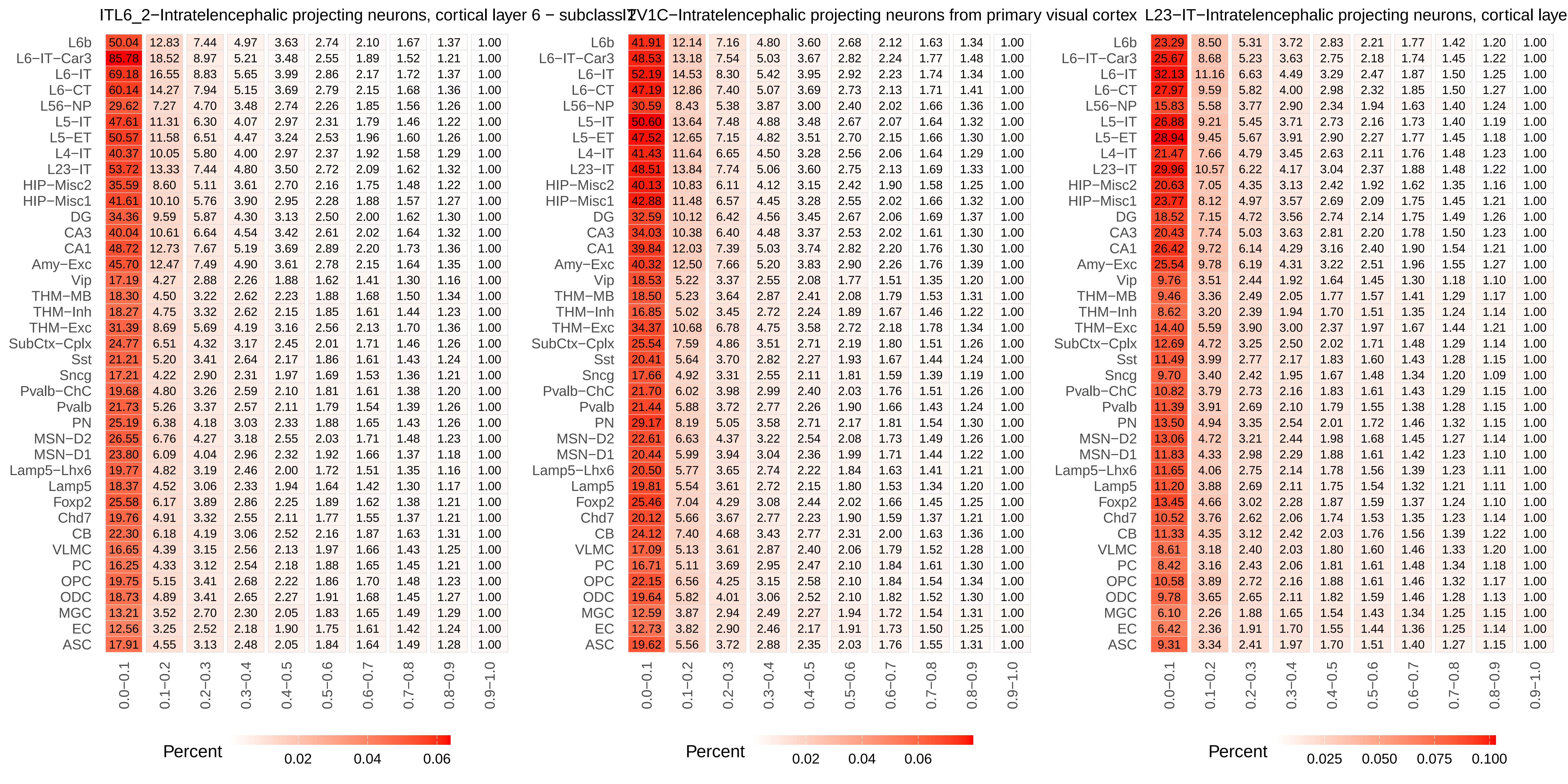

ITL6_2−Intratelencephalic projecting neurons, cortical layer 6 − subclass 2
IT V1C−Intratelencephalic projecting neurons from primary visual cortex
L23−IT−Intratelencephalic projecting neurons, cortical laye
L6b
L6−IT−Car3
L6−IT
L6−CT
L56−NP
L5−IT
L5−ET
L4−IT
L23−IT
HIP−Misc2
HIP−Misc1
DG
CA3
CA1
Amy−Exc
Vip
THM−MB
THM−Inh
THM−Exc
SubCtx−Cplx
Sst
Sncg
Pvalb−ChC
Pvalb
PN
MSN−D2
MSN−D1
Lamp5−Lhx6
Lamp5
Foxp2
Chd7
CB
VLMC
PC
OPC
ODC
MGC
EC
ASC
0.0−0.1
0.1−0.2
0.2−0.3
0.3−0.4
0.4−0.5
0.5−0.6
0.6−0.7
0.7−0.8
0.8−0.9
0.9−1.0
Percent
0.02
0.04
0.06
Percent
0.02
0.04
0.06
Percent
0.025
0.050
0.075
0.100

### L4−IT−Intratelencephalic projecting neurons, cortical layer 4

| | 0.0−0.1 | 0.1−0.2 | 0.2−0.3 | 0.3−0.4 | 0.4−0.5 | 0.5−0.6 | 0.6−0.7 | 0.7−0.8 | 0.8−0.9 | 0.9−1.0 |
|---|---|---|---|---|---|---|---|---|---|---|
| L6b | 38.21 | 10.40 | 6.13 | 4.16 | 3.11 | 2.41 | 1.90 | 1.53 | 1.28 | 1.00 |
| L6−IT−Car3 | 44.82 | 11.21 | 6.24 | 4.08 | 2.99 | 2.34 | 1.81 | 1.53 | 1.26 | 1.00 |
| L6−IT | 53.87 | 14.01 | 7.63 | 4.83 | 3.47 | 2.60 | 1.93 | 1.55 | 1.24 | 1.00 |
| L6−CT | 48.29 | 12.10 | 6.77 | 4.41 | 3.21 | 2.40 | 1.88 | 1.51 | 1.28 | 1.00 |
| L56−NP | 29.88 | 7.51 | 4.64 | 3.33 | 2.60 | 2.11 | 1.75 | 1.45 | 1.23 | 1.00 |
| L5−IT | 53.12 | 12.89 | 6.92 | 4.34 | 3.04 | 2.32 | 1.75 | 1.42 | 1.17 | 1.00 |
| L5−ET | 45.51 | 11.33 | 6.35 | 4.29 | 3.10 | 2.44 | 1.95 | 1.52 | 1.27 | 1.00 |
| L4−IT | 44.13 | 11.11 | 6.12 | 4.17 | 2.95 | 2.31 | 1.79 | 1.51 | 1.24 | 1.00 |
| L23−IT | 48.25 | 12.74 | 7.16 | 4.66 | 3.35 | 2.56 | 1.96 | 1.53 | 1.25 | 1.00 |
| HIP−Misc2 | 40.24 | 9.83 | 5.56 | 3.66 | 2.72 | 2.10 | 1.75 | 1.39 | 1.19 | 1.00 |
| HIP−Misc1 | 40.39 | 9.97 | 5.62 | 3.82 | 2.81 | 2.18 | 1.83 | 1.47 | 1.24 | 1.00 |
| DG | 31.29 | 9.12 | 5.66 | 4.07 | 3.06 | 2.45 | 1.93 | 1.57 | 1.30 | 1.00 |
| CA3 | 34.00 | 9.61 | 6.04 | 4.12 | 3.13 | 2.38 | 1.89 | 1.53 | 1.23 | 1.00 |
| CA1 | 40.48 | 11.32 | 6.80 | 4.71 | 3.35 | 2.50 | 1.99 | 1.59 | 1.23 | 1.00 |
| Amy−Exc | 40.61 | 11.80 | 7.16 | 4.83 | 3.55 | 2.66 | 2.10 | 1.65 | 1.33 | 1.00 |
| Vip | 15.54 | 4.02 | 2.70 | 2.08 | 1.72 | 1.51 | 1.31 | 1.21 | 1.11 | 1.00 |
| THM−MB | 15.43 | 3.94 | 2.81 | 2.26 | 1.93 | 1.72 | 1.51 | 1.33 | 1.20 | 1.00 |
| THM−Inh | 13.92 | 3.71 | 2.68 | 2.13 | 1.79 | 1.59 | 1.41 | 1.29 | 1.14 | 1.00 |
| THM−Exc | 26.34 | 7.56 | 4.97 | 3.56 | 2.73 | 2.17 | 1.82 | 1.51 | 1.25 | 1.00 |
| SubCtx−Cplx | 20.30 | 5.53 | 3.62 | 2.72 | 2.14 | 1.81 | 1.55 | 1.35 | 1.16 | 1.00 |
| Sst | 18.16 | 4.57 | 3.01 | 2.31 | 1.92 | 1.66 | 1.45 | 1.31 | 1.19 | 1.00 |
| Sncg | 14.81 | 3.82 | 2.60 | 2.07 | 1.76 | 1.55 | 1.38 | 1.24 | 1.12 | 1.00 |
| Pvalb−ChC | 17.61 | 4.47 | 2.99 | 2.33 | 1.92 | 1.65 | 1.44 | 1.30 | 1.16 | 1.00 |
| Pvalb | 18.82 | 4.69 | 2.97 | 2.26 | 1.91 | 1.62 | 1.44 | 1.32 | 1.17 | 1.00 |
| PN | 24.06 | 6.39 | 4.03 | 2.90 | 2.22 | 1.89 | 1.55 | 1.37 | 1.20 | 1.00 |
| MSN−D2 | 20.20 | 5.34 | 3.49 | 2.65 | 2.09 | 1.76 | 1.50 | 1.31 | 1.18 | 1.00 |
| MSN−D1 | 18.75 | 5.01 | 3.31 | 2.50 | 2.01 | 1.70 | 1.51 | 1.30 | 1.16 | 1.00 |
| Lamp5−Lhx6 | 16.84 | 4.26 | 2.85 | 2.15 | 1.80 | 1.52 | 1.37 | 1.24 | 1.13 | 1.00 |
| Lamp5 | 16.21 | 4.18 | 2.77 | 2.17 | 1.78 | 1.56 | 1.32 | 1.22 | 1.11 | 1.00 |
| Foxp2 | 21.16 | 5.35 | 3.31 | 2.42 | 1.94 | 1.65 | 1.39 | 1.27 | 1.13 | 1.00 |
| Chd7 | 17.05 | 4.43 | 2.95 | 2.24 | 1.90 | 1.63 | 1.43 | 1.29 | 1.18 | 1.00 |
| CB | 20.06 | 5.75 | 3.82 | 2.84 | 2.36 | 2.01 | 1.70 | 1.47 | 1.29 | 1.00 |
| VLMC | 14.30 | 3.88 | 2.87 | 2.30 | 1.98 | 1.74 | 1.53 | 1.38 | 1.22 | 1.00 |
| PC | 13.59 | 3.80 | 2.77 | 2.25 | 1.92 | 1.71 | 1.54 | 1.37 | 1.17 | 1.00 |
| OPC | 17.13 | 4.69 | 3.09 | 2.38 | 1.99 | 1.73 | 1.56 | 1.38 | 1.21 | 1.00 |
| ODC | 15.90 | 4.42 | 3.06 | 2.42 | 2.01 | 1.73 | 1.59 | 1.33 | 1.20 | 1.00 |
| MGC | 11.26 | 3.09 | 2.39 | 2.00 | 1.82 | 1.64 | 1.47 | 1.35 | 1.20 | 1.00 |
| EC | 11.08 | 3.00 | 2.32 | 1.98 | 1.80 | 1.61 | 1.47 | 1.35 | 1.17 | 1.00 |
| ASC | 15.20 | 3.99 | 2.76 | 2.21 | 1.86 | 1.65 | 1.47 | 1.34 | 1.21 | 1.00 |

Percent 0.02 0.04 0.06

### L6B−Intratelencephalic projecting neurons, cortical layer 6B

| | 0.0−0.1 | 0.1−0.2 | 0.2−0.3 | 0.3−0.4 | 0.4−0.5 | 0.5−0.6 | 0.6−0.7 | 0.7−0.8 | 0.8−0.9 | 0.9−1.0 |
|---|---|---|---|---|---|---|---|---|---|---|
| L6b | 44.72 | 11.98 | 6.48 | 4.08 | 2.81 | 2.14 | 1.65 | 1.37 | 1.14 | 1.00 |
| L6−IT−Car3 | 29.94 | 8.01 | 4.88 | 3.36 | 2.47 | 1.98 | 1.58 | 1.36 | 1.19 | 1.00 |
| L6−IT | 37.70 | 10.28 | 5.97 | 4.10 | 2.98 | 2.29 | 1.80 | 1.48 | 1.18 | 1.00 |
| L6−CT | 40.08 | 10.09 | 5.63 | 3.83 | 2.77 | 2.20 | 1.71 | 1.41 | 1.18 | 1.00 |
| L56−NP | 21.65 | 5.45 | 3.53 | 2.66 | 2.14 | 1.79 | 1.55 | 1.30 | 1.12 | 1.00 |
| L5−IT | 31.82 | 8.44 | 4.87 | 3.28 | 2.43 | 1.95 | 1.60 | 1.35 | 1.14 | 1.00 |
| L5−ET | 38.64 | 9.57 | 5.41 | 3.56 | 2.66 | 2.12 | 1.68 | 1.38 | 1.16 | 1.00 |
| L4−IT | 23.73 | 6.59 | 4.10 | 3.04 | 2.35 | 1.89 | 1.58 | 1.37 | 1.15 | 1.00 |
| L23−IT | 29.39 | 8.17 | 4.92 | 3.33 | 2.49 | 2.00 | 1.61 | 1.32 | 1.18 | 1.00 |
| HIP−Misc2 | 24.53 | 6.33 | 3.85 | 2.77 | 2.19 | 1.79 | 1.50 | 1.29 | 1.14 | 1.00 |
| HIP−Misc1 | 28.18 | 7.37 | 4.48 | 3.12 | 2.32 | 1.86 | 1.56 | 1.32 | 1.15 | 1.00 |
| DG | 21.04 | 6.30 | 4.25 | 3.20 | 2.47 | 2.06 | 1.69 | 1.46 | 1.25 | 1.00 |
| CA3 | 23.37 | 6.82 | 4.43 | 3.17 | 2.46 | 2.01 | 1.64 | 1.44 | 1.17 | 1.00 |
| CA1 | 27.45 | 7.83 | 4.91 | 3.46 | 2.64 | 2.06 | 1.66 | 1.35 | 1.13 | 1.00 |
| Amy−Exc | 24.89 | 7.37 | 4.75 | 3.37 | 2.61 | 2.10 | 1.73 | 1.40 | 1.20 | 1.00 |
| Vip | 12.53 | 3.32 | 2.26 | 1.77 | 1.51 | 1.35 | 1.19 | 1.12 | 1.05 | 1.00 |
| THM−MB | 13.16 | 3.38 | 2.42 | 1.98 | 1.72 | 1.57 | 1.38 | 1.27 | 1.15 | 1.00 |
| THM−Inh | 11.18 | 3.07 | 2.22 | 1.89 | 1.64 | 1.44 | 1.35 | 1.21 | 1.12 | 1.00 |
| THM−Exc | 18.83 | 5.53 | 3.79 | 2.80 | 2.31 | 1.90 | 1.62 | 1.36 | 1.16 | 1.00 |
| SubCtx−Cplx | 16.84 | 4.71 | 3.15 | 2.37 | 1.91 | 1.65 | 1.44 | 1.25 | 1.09 | 1.00 |
| Sst | 14.81 | 3.74 | 2.51 | 1.97 | 1.66 | 1.46 | 1.31 | 1.20 | 1.09 | 1.00 |
| Sncg | 14.64 | 3.76 | 2.59 | 2.04 | 1.73 | 1.49 | 1.36 | 1.25 | 1.08 | 1.00 |
| Pvalb−ChC | 13.81 | 3.63 | 2.57 | 2.02 | 1.73 | 1.54 | 1.36 | 1.24 | 1.13 | 1.00 |
| Pvalb | 14.56 | 3.65 | 2.46 | 1.95 | 1.67 | 1.46 | 1.33 | 1.21 | 1.12 | 1.00 |
| PN | 17.99 | 4.86 | 3.27 | 2.39 | 1.95 | 1.61 | 1.43 | 1.27 | 1.16 | 1.00 |
| MSN−D2 | 15.06 | 4.14 | 2.80 | 2.16 | 1.82 | 1.58 | 1.36 | 1.22 | 1.12 | 1.00 |
| MSN−D1 | 14.22 | 3.89 | 2.64 | 2.05 | 1.72 | 1.51 | 1.36 | 1.18 | 1.08 | 1.00 |
| Lamp5−Lhx6 | 17.09 | 4.35 | 2.79 | 2.09 | 1.68 | 1.45 | 1.30 | 1.17 | 1.07 | 1.00 |
| Lamp5 | 17.61 | 4.44 | 2.91 | 2.20 | 1.78 | 1.49 | 1.23 | 1.14 | 1.04 | 1.00 |
| Foxp2 | 16.10 | 4.17 | 2.74 | 2.10 | 1.73 | 1.45 | 1.28 | 1.19 | 1.09 | 1.00 |
| Chd7 | 14.33 | 3.73 | 2.55 | 1.97 | 1.68 | 1.45 | 1.29 | 1.16 | 1.08 | 1.00 |
| CB | 15.35 | 4.57 | 3.17 | 2.39 | 2.02 | 1.78 | 1.55 | 1.37 | 1.18 | 1.00 |
| VLMC | 11.88 | 3.29 | 2.44 | 2.02 | 1.73 | 1.57 | 1.42 | 1.31 | 1.11 | 1.00 |
| PC | 11.61 | 3.25 | 2.45 | 2.04 | 1.80 | 1.56 | 1.46 | 1.29 | 1.14 | 1.00 |
| OPC | 14.89 | 4.02 | 2.76 | 2.10 | 1.82 | 1.54 | 1.41 | 1.30 | 1.12 | 1.00 |
| ODC | 13.84 | 3.86 | 2.72 | 2.14 | 1.83 | 1.61 | 1.43 | 1.30 | 1.15 | 1.00 |
| MGC | 8.48 | 2.26 | 1.87 | 1.60 | 1.51 | 1.44 | 1.28 | 1.23 | 1.14 | 1.00 |
| EC | 9.32 | 2.56 | 1.98 | 1.77 | 1.59 | 1.48 | 1.40 | 1.28 | 1.17 | 1.00 |
| ASC | 13.41 | 3.62 | 2.47 | 1.97 | 1.67 | 1.49 | 1.38 | 1.26 | 1.11 | 1.00 |

Percent 0.01 0.02 0.03 0.04 0.05

### LAMP5_1−LAMP5+ GABAergic neurons

| | 0.0−0.1 | 0.1−0.2 | 0.2−0.3 | 0.3−0.4 | 0.4−0.5 | 0.5−0.6 | 0.6−0.7 | 0.7−0.8 | 0.8−0.9 | 0.9−1.0 |
|---|---|---|---|---|---|---|---|---|---|---|
| L6b | 42.64 | 10.01 | 5.92 | 4.15 | 3.04 | 2.42 | 1.86 | 1.57 | 1.28 | 1.00 |
| L6−IT−Car3 | 30.46 | 6.84 | 4.42 | 3.29 | 2.55 | 2.16 | 1.81 | 1.56 | 1.30 | 1.00 |
| L6−IT | 38.94 | 9.20 | 5.72 | 4.07 | 3.23 | 2.58 | 2.03 | 1.62 | 1.30 | 1.00 |
| L6−CT | 38.73 | 8.65 | 5.20 | 3.77 | 2.90 | 2.33 | 1.91 | 1.53 | 1.27 | 1.00 |
| L56−NP | 24.46 | 5.12 | 3.46 | 2.67 | 2.18 | 1.91 | 1.66 | 1.45 | 1.23 | 1.00 |
| L5−IT | 32.86 | 7.32 | 4.60 | 3.29 | 2.57 | 2.03 | 1.77 | 1.50 | 1.25 | 1.00 |
| L5−ET | 47.21 | 10.17 | 6.06 | 4.06 | 3.16 | 2.45 | 1.98 | 1.54 | 1.29 | 1.00 |
| L4−IT | 24.84 | 5.68 | 3.64 | 2.89 | 2.32 | 1.95 | 1.63 | 1.41 | 1.17 | 1.00 |
| L23−IT | 33.62 | 7.96 | 5.00 | 3.57 | 2.83 | 2.29 | 1.90 | 1.55 | 1.28 | 1.00 |
| HIP−Misc2 | 26.32 | 5.85 | 3.78 | 2.75 | 2.18 | 1.86 | 1.59 | 1.43 | 1.22 | 1.00 |
| HIP−Misc1 | 33.27 | 7.59 | 4.83 | 3.56 | 2.67 | 2.07 | 1.84 | 1.53 | 1.29 | 1.00 |
| DG | 38.70 | 10.00 | 6.30 | 4.31 | 3.35 | 2.55 | 2.02 | 1.66 | 1.28 | 1.00 |
| CA3 | 35.22 | 8.52 | 5.42 | 3.77 | 2.87 | 2.26 | 1.82 | 1.56 | 1.23 | 1.00 |
| CA1 | 45.00 | 10.70 | 6.63 | 4.65 | 3.32 | 2.58 | 2.09 | 1.70 | 1.37 | 1.00 |
| Amy−Exc | 33.95 | 8.35 | 5.40 | 3.87 | 3.01 | 2.40 | 1.96 | 1.59 | 1.28 | 1.00 |
| Vip | 33.26 | 7.01 | 4.00 | 2.78 | 2.08 | 1.72 | 1.39 | 1.21 | 1.06 | 1.00 |
| THM−MB | 22.37 | 4.53 | 3.08 | 2.38 | 1.99 | 1.74 | 1.57 | 1.37 | 1.21 | 1.00 |
| THM−Inh | 23.69 | 5.24 | 3.42 | 2.62 | 2.24 | 1.83 | 1.57 | 1.35 | 1.20 | 1.00 |
| THM−Exc | 29.65 | 6.96 | 4.54 | 3.36 | 2.63 | 2.19 | 1.86 | 1.58 | 1.25 | 1.00 |
| SubCtx−Cplx | 32.22 | 7.23 | 4.49 | 3.20 | 2.46 | 1.97 | 1.64 | 1.35 | 1.16 | 1.00 |
| Sst | 30.23 | 6.02 | 3.79 | 2.76 | 2.23 | 1.86 | 1.60 | 1.35 | 1.16 | 1.00 |
| Sncg | 44.59 | 9.06 | 4.96 | 3.28 | 2.43 | 1.97 | 1.62 | 1.33 | 1.15 | 1.00 |
| Pvalb−ChC | 34.14 | 6.90 | 4.13 | 2.98 | 2.32 | 1.91 | 1.61 | 1.44 | 1.15 | 1.00 |
| Pvalb | 28.19 | 5.49 | 3.36 | 2.37 | 1.92 | 1.64 | 1.42 | 1.26 | 1.13 | 1.00 |
| PN | 31.38 | 6.95 | 4.28 | 2.98 | 2.23 | 1.95 | 1.59 | 1.35 | 1.19 | 1.00 |
| MSN−D2 | 24.09 | 5.54 | 3.64 | 2.76 | 2.27 | 1.90 | 1.66 | 1.41 | 1.21 | 1.00 |
| MSN−D1 | 21.96 | 4.96 | 3.33 | 2.58 | 2.08 | 1.77 | 1.53 | 1.37 | 1.19 | 1.00 |
| Lamp5−Lhx6 | 59.65 | 10.76 | 5.05 | 3.01 | 2.03 | 1.59 | 1.29 | 1.00 | 0.91 | 1.00 |
| Lamp5 | 66.23 | 12.12 | 5.74 | 3.43 | 2.38 | 1.80 | 1.41 | 1.18 | 1.07 | 1.00 |
| Foxp2 | 27.76 | 6.05 | 3.83 | 2.82 | 2.25 | 1.87 | 1.61 | 1.39 | 1.27 | 1.00 |
| Chd7 | 37.53 | 7.78 | 4.46 | 3.10 | 2.31 | 1.84 | 1.55 | 1.30 | 1.18 | 1.00 |
| CB | 29.42 | 7.17 | 4.54 | 3.41 | 2.67 | 2.21 | 1.95 | 1.62 | 1.31 | 1.00 |
| VLMC | 20.55 | 4.66 | 3.14 | 2.51 | 2.10 | 1.87 | 1.62 | 1.35 | 1.16 | 1.00 |
| PC | 22.11 | 5.08 | 3.59 | 2.88 | 2.33 | 2.01 | 1.80 | 1.49 | 1.35 | 1.00 |
| OPC | 33.16 | 7.07 | 4.19 | 2.95 | 2.41 | 1.90 | 1.57 | 1.37 | 1.15 | 1.00 |
| ODC | 32.12 | 7.32 | 4.46 | 3.18 | 2.50 | 2.10 | 1.77 | 1.40 | 1.15 | 1.00 |
| MGC | 13.41 | 2.95 | 2.27 | 1.96 | 1.81 | 1.64 | 1.46 | 1.34 | 1.18 | 1.00 |
| EC | 15.53 | 3.41 | 2.60 | 2.16 | 1.93 | 1.73 | 1.53 | 1.40 | 1.21 | 1.00 |
| ASC | 30.18 | 6.45 | 3.86 | 2.82 | 2.19 | 1.80 | 1.55 | 1.35 | 1.17 | 1.00 |

Percent 0.01 0.02 0.03 0.04 0.05

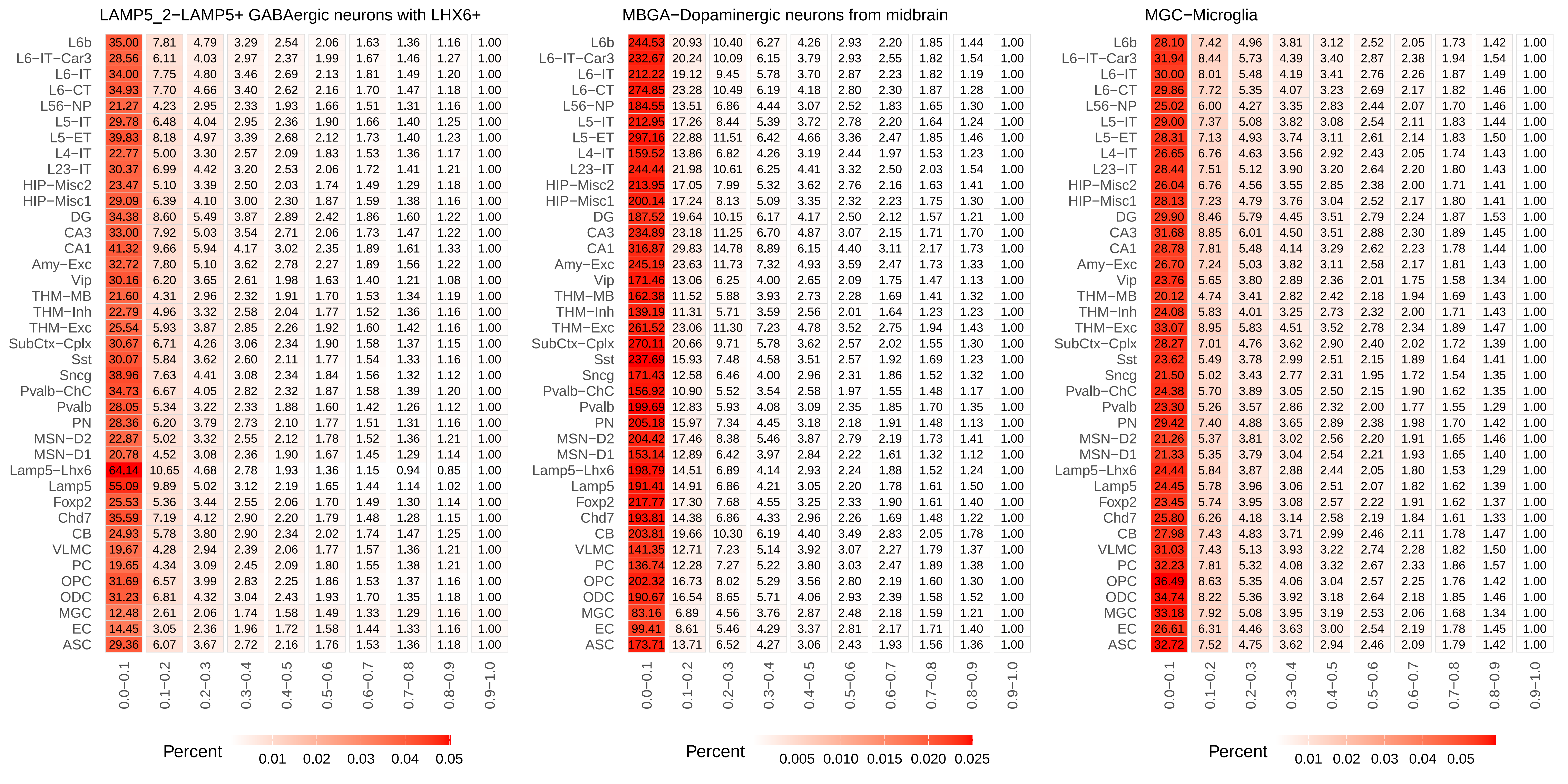
LAMP5_2−LAMP5+ GABAergic neurons with LHX6+
L6b 35.00 7.81 4.79 3.29 2.54 2.06 1.63 1.36 1.16 1.00
L6−IT−Car3 28.56 6.11 4.03 2.97 2.37 1.99 1.67 1.46 1.27 1.00
L6−IT 34.00 7.75 4.80 3.46 2.69 2.13 1.81 1.49 1.20 1.00
L6−CT 34.93 7.70 4.66 3.40 2.62 2.16 1.70 1.47 1.18 1.00
L56−NP 21.27 4.23 2.95 2.33 1.93 1.66 1.51 1.31 1.16 1.00
L5−IT 29.78 6.48 4.04 2.95 2.36 1.90 1.66 1.40 1.25 1.00
L5−ET 39.83 8.18 4.97 3.39 2.68 2.12 1.73 1.40 1.23 1.00
L4−IT 22.77 5.00 3.30 2.57 2.09 1.83 1.53 1.36 1.17 1.00
L23−IT 30.37 6.99 4.42 3.20 2.53 2.06 1.72 1.41 1.21 1.00
HIP−Misc2 23.47 5.10 3.39 2.50 2.03 1.74 1.49 1.29 1.18 1.00
HIP−Misc1 29.09 6.39 4.10 3.00 2.30 1.87 1.59 1.38 1.16 1.00
DG 34.38 8.60 5.49 3.87 2.89 2.42 1.86 1.60 1.22 1.00
CA3 33.00 7.92 5.03 3.54 2.71 2.06 1.73 1.47 1.22 1.00
CA1 41.32 9.66 5.94 4.17 3.02 2.35 1.89 1.61 1.33 1.00
Amy−Exc 32.72 7.80 5.10 3.62 2.78 2.27 1.89 1.56 1.22 1.00
Vip 30.16 6.20 3.65 2.61 1.98 1.63 1.40 1.21 1.08 1.00
THM−MB 21.60 4.31 2.96 2.32 1.91 1.70 1.53 1.34 1.19 1.00
THM−Inh 22.79 4.96 3.32 2.58 2.04 1.77 1.52 1.36 1.16 1.00
THM−Exc 25.54 5.93 3.87 2.85 2.26 1.92 1.60 1.42 1.16 1.00
SubCtx−Cplx 30.67 6.71 4.26 3.06 2.34 1.90 1.58 1.37 1.15 1.00
Sst 30.07 5.84 3.62 2.60 2.11 1.77 1.54 1.33 1.16 1.00
Sncg 38.96 7.63 4.41 3.08 2.34 1.84 1.56 1.32 1.12 1.00
Pvalb−ChC 34.73 6.67 4.05 2.82 2.32 1.87 1.58 1.39 1.20 1.00
Pvalb 28.05 5.34 3.22 2.33 1.88 1.60 1.42 1.26 1.12 1.00
PN 28.36 6.20 3.79 2.73 2.10 1.77 1.51 1.31 1.16 1.00
MSN−D2 22.87 5.02 3.32 2.55 2.12 1.78 1.52 1.36 1.21 1.00
MSN−D1 20.78 4.52 3.08 2.36 1.90 1.67 1.45 1.29 1.14 1.00
Lamp5−Lhx6 64.14 10.65 4.68 2.78 1.93 1.36 1.15 0.94 0.85 1.00
Lamp5 55.09 9.89 5.02 3.12 2.19 1.65 1.44 1.14 1.02 1.00
Foxp2 25.53 5.36 3.44 2.55 2.06 1.70 1.49 1.30 1.14 1.00
Chd7 35.59 7.19 4.12 2.90 2.20 1.79 1.48 1.28 1.15 1.00
CB 24.93 5.78 3.80 2.90 2.34 2.02 1.74 1.47 1.25 1.00
VLMC 19.67 4.28 2.94 2.39 2.06 1.77 1.57 1.36 1.21 1.00
PC 19.65 4.34 3.09 2.45 2.09 1.80 1.55 1.38 1.21 1.00
OPC 31.69 6.57 3.99 2.83 2.25 1.86 1.53 1.37 1.16 1.00
ODC 31.23 6.81 4.32 3.04 2.43 1.93 1.70 1.35 1.18 1.00
MGC 12.48 2.61 2.06 1.74 1.58 1.49 1.33 1.29 1.16 1.00
EC 14.45 3.05 2.36 1.96 1.72 1.58 1.44 1.33 1.16 1.00
ASC 29.36 6.07 3.67 2.72 2.16 1.76 1.53 1.36 1.18 1.00
0.0−0.1 0.1−0.2 0.2−0.3 0.3−0.4 0.4−0.5 0.5−0.6 0.6−0.7 0.7−0.8 0.8−0.9 0.9−1.0
Percent
0.01 0.02 0.03 0.04 0.05
MBGA−Dopaminergic neurons from midbrain
L6b 244.53 20.93 10.40 6.27 4.26 2.93 2.20 1.85 1.44 1.00
L6−IT−Car3 232.67 20.24 10.09 6.15 3.79 2.93 2.55 1.82 1.54 1.00
L6−IT 212.22 19.12 9.45 5.78 3.70 2.87 2.23 1.82 1.19 1.00
L6−CT 274.85 23.28 10.49 6.19 4.18 2.80 2.30 1.87 1.28 1.00
L56−NP 184.55 13.51 6.86 4.44 3.07 2.52 1.83 1.65 1.30 1.00
L5−IT 212.95 17.26 8.44 5.39 3.72 2.78 2.20 1.64 1.24 1.00
L5−ET 297.16 22.88 11.51 6.42 4.66 3.36 2.47 1.85 1.46 1.00
L4−IT 159.52 13.86 6.82 4.26 3.19 2.44 1.97 1.53 1.23 1.00
L23−IT 244.44 21.98 10.61 6.25 4.41 3.32 2.50 2.03 1.54 1.00
HIP−Misc2 213.95 17.05 7.99 5.32 3.62 2.76 2.16 1.63 1.41 1.00
HIP−Misc1 200.14 17.24 8.13 5.09 3.35 2.32 2.23 1.75 1.30 1.00
DG 187.52 19.64 10.15 6.17 4.17 2.50 2.12 1.57 1.21 1.00
CA3 234.89 23.18 11.25 6.70 4.87 3.07 2.15 1.71 1.70 1.00
CA1 316.87 29.83 14.78 8.89 6.15 4.40 3.11 2.17 1.73 1.00
Amy−Exc 245.19 23.63 11.73 7.32 4.93 3.59 2.47 1.73 1.33 1.00
Vip 171.46 13.06 6.25 4.00 2.65 2.09 1.75 1.47 1.13 1.00
THM−MB 162.38 11.52 5.88 3.93 2.73 2.28 1.69 1.41 1.32 1.00
THM−Inh 139.19 11.31 5.71 3.59 2.56 2.01 1.64 1.23 1.23 1.00
THM−Exc 261.52 23.06 11.30 7.23 4.78 3.52 2.75 1.94 1.43 1.00
SubCtx−Cplx 270.11 20.66 9.71 5.78 3.62 2.57 2.02 1.55 1.30 1.00
Sst 237.69 15.93 7.48 4.58 3.51 2.57 1.92 1.69 1.23 1.00
Sncg 171.43 12.58 6.46 4.00 2.96 2.31 1.86 1.52 1.32 1.00
Pvalb−ChC 156.92 10.90 5.52 3.54 2.58 1.97 1.55 1.48 1.17 1.00
Pvalb 199.69 12.83 5.93 4.08 3.09 2.35 1.85 1.70 1.35 1.00
PN 205.18 15.97 7.34 4.45 3.18 2.18 1.91 1.48 1.13 1.00
MSN−D2 204.42 17.46 8.38 5.46 3.87 2.79 2.19 1.73 1.41 1.00
MSN−D1 153.14 12.89 6.42 3.97 2.84 2.22 1.61 1.32 1.12 1.00
Lamp5−Lhx6 198.79 14.51 6.89 4.14 2.93 2.24 1.88 1.52 1.24 1.00
Lamp5 191.41 14.91 6.86 4.21 3.05 2.20 1.78 1.61 1.50 1.00
Foxp2 217.77 17.30 7.68 4.55 3.25 2.33 1.90 1.61 1.40 1.00
Chd7 193.81 14.38 6.86 4.33 2.96 2.26 1.69 1.48 1.22 1.00
CB 203.81 19.66 10.30 6.19 4.40 3.49 2.83 2.05 1.78 1.00
VLMC 141.35 12.71 7.23 5.14 3.92 3.07 2.27 1.79 1.37 1.00
PC 136.74 12.28 7.27 5.22 3.80 3.03 2.47 1.89 1.38 1.00
OPC 202.32 16.73 8.02 5.29 3.56 2.80 2.19 1.60 1.30 1.00
ODC 190.67 16.54 8.65 5.71 4.06 2.93 2.39 1.58 1.52 1.00
MGC 83.16 6.89 4.56 3.76 2.87 2.48 2.18 1.59 1.21 1.00
EC 99.41 8.61 5.46 4.29 3.37 2.81 2.17 1.71 1.40 1.00
ASC 173.71 13.71 6.52 4.27 3.06 2.43 1.93 1.56 1.36 1.00
0.0−0.1 0.1−0.2 0.2−0.3 0.3−0.4 0.4−0.5 0.5−0.6 0.6−0.7 0.7−0.8 0.8−0.9 0.9−1.0
Percent
0.005 0.010 0.015 0.020 0.025
MGC−Microglia
L6b 28.10 7.42 4.96 3.81 3.12 2.52 2.05 1.73 1.42 1.00
L6−IT−Car3 31.94 8.44 5.73 4.39 3.40 2.87 2.38 1.94 1.54 1.00
L6−IT 30.00 8.01 5.48 4.19 3.41 2.76 2.26 1.87 1.49 1.00
L6−CT 29.86 7.72 5.35 4.07 3.23 2.69 2.17 1.82 1.46 1.00
L56−NP 25.02 6.00 4.27 3.35 2.83 2.44 2.07 1.70 1.46 1.00
L5−IT 29.00 7.37 5.08 3.82 3.08 2.54 2.11 1.83 1.44 1.00
L5−ET 28.31 7.13 4.93 3.74 3.11 2.61 2.14 1.83 1.50 1.00
L4−IT 26.65 6.76 4.63 3.56 2.92 2.43 2.05 1.74 1.43 1.00
L23−IT 28.44 7.51 5.12 3.90 3.20 2.64 2.20 1.80 1.43 1.00
HIP−Misc2 26.04 6.76 4.56 3.55 2.85 2.38 2.00 1.71 1.41 1.00
HIP−Misc1 28.13 7.23 4.79 3.76 3.04 2.52 2.17 1.80 1.41 1.00
DG 29.90 8.46 5.79 4.45 3.51 2.79 2.24 1.87 1.53 1.00
CA3 31.68 8.85 6.01 4.50 3.51 2.88 2.30 1.89 1.45 1.00
CA1 28.78 7.81 5.48 4.14 3.29 2.62 2.23 1.78 1.44 1.00
Amy−Exc 26.70 7.24 5.03 3.82 3.11 2.58 2.17 1.81 1.43 1.00
Vip 23.76 5.65 3.80 2.89 2.36 2.01 1.75 1.58 1.34 1.00
THM−MB 20.12 4.74 3.41 2.82 2.42 2.18 1.94 1.69 1.43 1.00
THM−Inh 24.08 5.83 4.01 3.25 2.73 2.32 2.00 1.71 1.43 1.00
THM−Exc 33.07 8.95 5.83 4.51 3.52 2.78 2.34 1.89 1.47 1.00
SubCtx−Cplx 28.27 7.01 4.76 3.62 2.90 2.40 2.02 1.72 1.39 1.00
Sst 23.62 5.49 3.78 2.99 2.51 2.15 1.89 1.64 1.41 1.00
Sncg 21.50 5.02 3.43 2.77 2.31 1.95 1.72 1.54 1.35 1.00
Pvalb−ChC 24.38 5.70 3.89 3.05 2.50 2.15 1.90 1.62 1.35 1.00
Pvalb 23.30 5.26 3.57 2.86 2.32 2.00 1.77 1.55 1.29 1.00
PN 29.42 7.40 4.88 3.65 2.89 2.38 1.98 1.70 1.42 1.00
MSN−D2 21.26 5.37 3.81 3.02 2.56 2.20 1.91 1.65 1.46 1.00
MSN−D1 21.33 5.35 3.79 3.04 2.54 2.21 1.93 1.65 1.40 1.00
Lamp5−Lhx6 24.44 5.84 3.87 2.88 2.44 2.05 1.80 1.53 1.29 1.00
Lamp5 24.45 5.78 3.96 3.06 2.51 2.07 1.82 1.62 1.39 1.00
Foxp2 23.45 5.74 3.95 3.08 2.57 2.22 1.91 1.62 1.37 1.00
Chd7 25.80 6.26 4.18 3.14 2.58 2.19 1.84 1.61 1.33 1.00
CB 27.98 7.43 4.83 3.71 2.99 2.46 2.11 1.78 1.47 1.00
VLMC 31.03 7.43 5.13 3.93 3.22 2.74 2.28 1.82 1.50 1.00
PC 32.23 7.81 5.32 4.08 3.32 2.67 2.33 1.86 1.57 1.00
OPC 36.49 8.63 5.35 4.06 3.04 2.57 2.25 1.76 1.42 1.00
ODC 34.74 8.22 5.36 3.92 3.18 2.64 2.18 1.85 1.46 1.00
MGC 33.18 7.92 5.08 3.95 3.19 2.53 2.06 1.68 1.34 1.00
EC 26.61 6.31 4.46 3.63 3.00 2.54 2.19 1.78 1.45 1.00
ASC 32.72 7.52 4.75 3.62 2.94 2.46 2.09 1.79 1.42 1.00
0.0−0.1 0.1−0.2 0.2−0.3 0.3−0.4 0.4−0.5 0.5−0.6 0.6−0.7 0.7−0.8 0.8−0.9 0.9−1.0
Percent
0.01 0.02 0.03 0.04 0.05

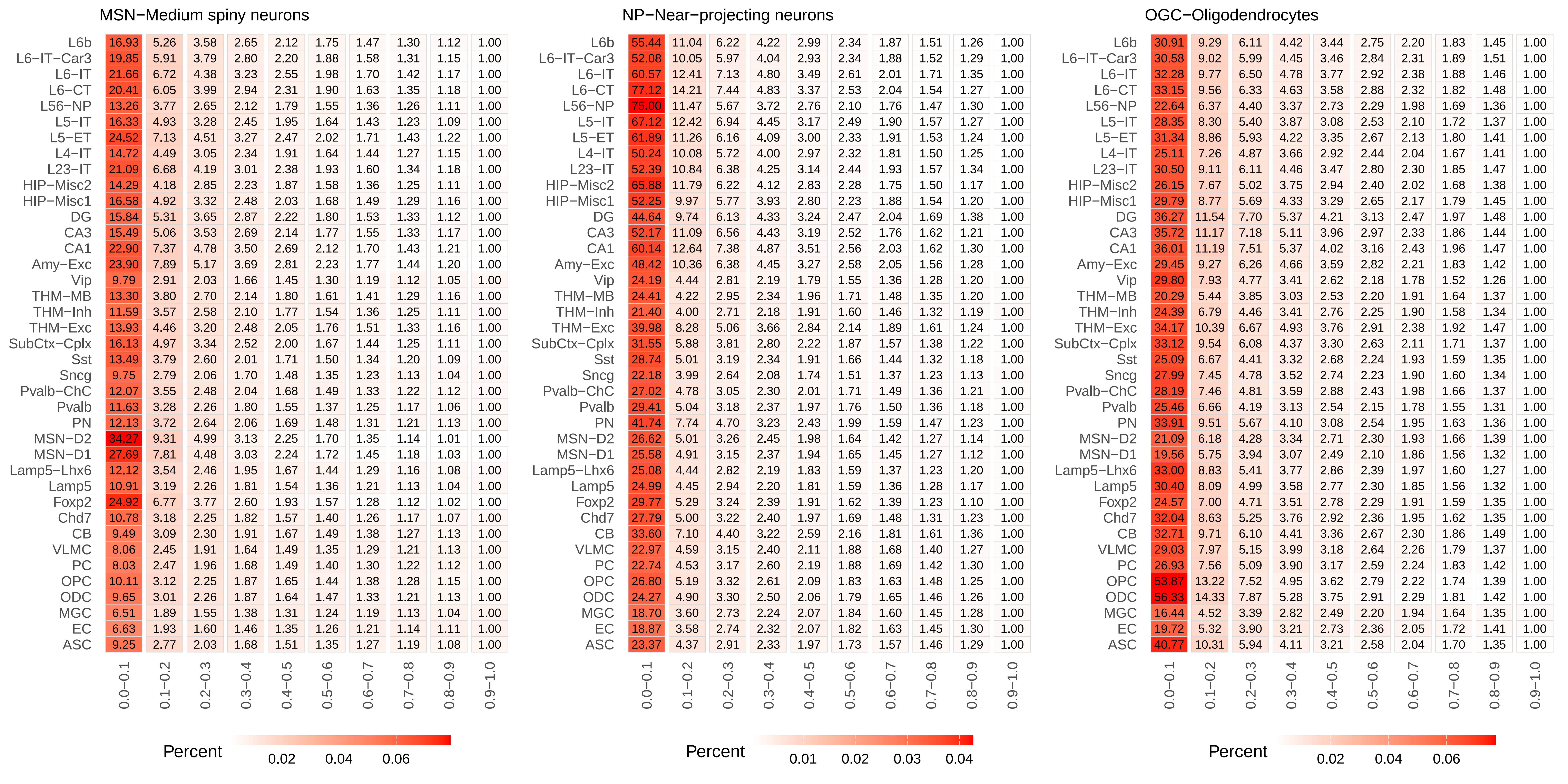
MSN−Medium spiny neurons
L6b 16.93 5.26 3.58 2.65 2.12 1.75 1.47 1.30 1.12 1.00
L6−IT−Car3 19.85 5.91 3.79 2.80 2.20 1.88 1.58 1.31 1.15 1.00
L6−IT 21.66 6.72 4.38 3.23 2.55 1.98 1.70 1.42 1.17 1.00
L6−CT 20.41 6.05 3.99 2.94 2.31 1.90 1.63 1.35 1.18 1.00
L56−NP 13.26 3.77 2.65 2.12 1.79 1.55 1.36 1.26 1.11 1.00
L5−IT 16.33 4.93 3.28 2.45 1.95 1.64 1.43 1.23 1.09 1.00
L5−ET 24.52 7.13 4.51 3.27 2.47 2.02 1.71 1.43 1.22 1.00
L4−IT 14.72 4.49 3.05 2.34 1.91 1.64 1.44 1.27 1.15 1.00
L23−IT 21.09 6.68 4.19 3.01 2.38 1.93 1.60 1.34 1.18 1.00
HIP−Misc2 14.29 4.18 2.85 2.23 1.87 1.58 1.36 1.25 1.11 1.00
HIP−Misc1 16.58 4.92 3.32 2.48 2.03 1.68 1.49 1.29 1.16 1.00
DG 15.84 5.31 3.65 2.87 2.22 1.80 1.53 1.33 1.12 1.00
CA3 15.49 5.06 3.53 2.69 2.14 1.77 1.55 1.33 1.17 1.00
CA1 22.90 7.37 4.78 3.50 2.69 2.12 1.70 1.43 1.21 1.00
Amy−Exc 23.90 7.89 5.17 3.69 2.81 2.23 1.77 1.44 1.20 1.00
Vip 9.79 2.91 2.03 1.66 1.45 1.30 1.19 1.12 1.05 1.00
THM−MB 13.30 3.80 2.70 2.14 1.80 1.61 1.41 1.29 1.16 1.00
THM−Inh 11.59 3.57 2.58 2.10 1.77 1.54 1.36 1.25 1.11 1.00
THM−Exc 13.93 4.46 3.20 2.48 2.05 1.76 1.51 1.33 1.16 1.00
SubCtx−Cplx 16.13 4.97 3.34 2.52 2.00 1.67 1.44 1.25 1.11 1.00
Sst 13.49 3.79 2.60 2.01 1.71 1.50 1.34 1.20 1.09 1.00
Sncg 9.75 2.79 2.06 1.70 1.48 1.35 1.23 1.13 1.04 1.00
Pvalb−ChC 12.07 3.55 2.48 2.04 1.68 1.49 1.33 1.22 1.12 1.00
Pvalb 11.63 3.28 2.26 1.80 1.55 1.37 1.25 1.17 1.06 1.00
PN 12.13 3.72 2.64 2.06 1.69 1.48 1.31 1.21 1.13 1.00
MSN−D2 34.27 9.31 4.99 3.13 2.25 1.70 1.35 1.14 1.01 1.00
MSN−D1 27.69 7.81 4.48 3.03 2.24 1.72 1.45 1.18 1.03 1.00
Lamp5−Lhx6 12.12 3.54 2.46 1.95 1.67 1.44 1.29 1.16 1.08 1.00
Lamp5 10.91 3.19 2.26 1.81 1.54 1.36 1.21 1.13 1.04 1.00
Foxp2 24.92 6.77 3.77 2.60 1.93 1.57 1.28 1.12 1.02 1.00
Chd7 10.78 3.18 2.25 1.82 1.57 1.40 1.26 1.17 1.07 1.00
CB 9.49 3.09 2.30 1.91 1.67 1.49 1.38 1.27 1.13 1.00
VLMC 8.06 2.45 1.91 1.64 1.49 1.35 1.29 1.21 1.13 1.00
PC 8.03 2.47 1.96 1.68 1.49 1.40 1.30 1.22 1.12 1.00
OPC 10.11 3.12 2.25 1.87 1.65 1.44 1.38 1.28 1.15 1.00
ODC 9.65 3.01 2.26 1.87 1.64 1.47 1.33 1.21 1.13 1.00
MGC 6.51 1.89 1.55 1.38 1.31 1.24 1.19 1.13 1.04 1.00
EC 6.63 1.93 1.60 1.46 1.35 1.26 1.21 1.14 1.11 1.00
ASC 9.25 2.77 2.03 1.68 1.51 1.35 1.27 1.19 1.08 1.00
0.0−0.1 0.1−0.2 0.2−0.3 0.3−0.4 0.4−0.5 0.5−0.6 0.6−0.7 0.7−0.8 0.8−0.9 0.9−1.0
Percent
0.02 0.04 0.06
NP−Near−projecting neurons
L6b 55.44 11.04 6.22 4.22 2.99 2.34 1.87 1.51 1.26 1.00
L6−IT−Car3 52.08 10.05 5.97 4.04 2.93 2.34 1.88 1.52 1.29 1.00
L6−IT 60.57 12.41 7.13 4.80 3.49 2.61 2.01 1.71 1.35 1.00
L6−CT 77.12 14.21 7.44 4.83 3.37 2.53 2.04 1.54 1.27 1.00
L56−NP 75.00 11.47 5.67 3.72 2.76 2.10 1.76 1.47 1.30 1.00
L5−IT 67.12 12.42 6.94 4.45 3.17 2.49 1.90 1.57 1.27 1.00
L5−ET 61.89 11.26 6.16 4.09 3.00 2.33 1.91 1.53 1.24 1.00
L4−IT 50.24 10.08 5.72 4.00 2.97 2.32 1.81 1.50 1.25 1.00
L23−IT 52.39 10.84 6.38 4.25 3.14 2.44 1.93 1.57 1.34 1.00
HIP−Misc2 65.88 11.79 6.22 4.12 2.83 2.28 1.75 1.50 1.17 1.00
HIP−Misc1 52.25 9.97 5.77 3.93 2.80 2.23 1.88 1.54 1.20 1.00
DG 44.64 9.74 6.13 4.33 3.24 2.47 2.04 1.69 1.38 1.00
CA3 52.17 11.09 6.56 4.43 3.19 2.52 1.76 1.62 1.21 1.00
CA1 60.14 12.64 7.38 4.87 3.51 2.56 2.03 1.62 1.30 1.00
Amy−Exc 48.42 10.36 6.38 4.45 3.27 2.58 2.05 1.56 1.28 1.00
Vip 24.19 4.44 2.81 2.19 1.79 1.55 1.36 1.28 1.20 1.00
THM−MB 24.41 4.22 2.95 2.34 1.96 1.71 1.48 1.35 1.20 1.00
THM−Inh 21.40 4.00 2.71 2.18 1.91 1.60 1.46 1.32 1.19 1.00
THM−Exc 39.98 8.28 5.06 3.66 2.84 2.14 1.89 1.61 1.24 1.00
SubCtx−Cplx 31.55 5.88 3.81 2.80 2.22 1.87 1.57 1.38 1.22 1.00
Sst 28.74 5.01 3.19 2.34 1.91 1.66 1.44 1.32 1.18 1.00
Sncg 22.18 3.99 2.64 2.08 1.74 1.51 1.37 1.23 1.13 1.00
Pvalb−ChC 27.02 4.78 3.05 2.30 2.01 1.71 1.49 1.36 1.21 1.00
Pvalb 29.41 5.04 3.18 2.37 1.97 1.76 1.50 1.36 1.18 1.00
PN 41.74 7.74 4.70 3.23 2.43 1.99 1.59 1.47 1.23 1.00
MSN−D2 26.62 5.01 3.26 2.45 1.98 1.64 1.42 1.27 1.14 1.00
MSN−D1 25.58 4.91 3.15 2.37 1.94 1.65 1.45 1.27 1.12 1.00
Lamp5−Lhx6 25.08 4.44 2.82 2.19 1.83 1.59 1.37 1.23 1.20 1.00
Lamp5 24.99 4.45 2.94 2.20 1.81 1.59 1.36 1.28 1.17 1.00
Foxp2 29.77 5.29 3.24 2.39 1.91 1.62 1.39 1.23 1.10 1.00
Chd7 27.79 5.00 3.22 2.40 1.97 1.69 1.48 1.31 1.23 1.00
CB 33.60 7.10 4.40 3.22 2.59 2.16 1.81 1.61 1.36 1.00
VLMC 22.97 4.59 3.15 2.40 2.11 1.88 1.68 1.40 1.27 1.00
PC 22.74 4.53 3.17 2.60 2.19 1.88 1.69 1.42 1.30 1.00
OPC 26.80 5.19 3.32 2.61 2.09 1.83 1.63 1.48 1.25 1.00
ODC 24.27 4.90 3.30 2.50 2.06 1.79 1.65 1.46 1.26 1.00
MGC 18.70 3.60 2.73 2.24 2.07 1.84 1.60 1.45 1.28 1.00
EC 18.87 3.58 2.74 2.32 2.07 1.82 1.63 1.45 1.30 1.00
ASC 23.37 4.37 2.91 2.33 1.97 1.73 1.57 1.46 1.29 1.00
0.0−0.1 0.1−0.2 0.2−0.3 0.3−0.4 0.4−0.5 0.5−0.6 0.6−0.7 0.7−0.8 0.8−0.9 0.9−1.0
Percent
0.01 0.02 0.03 0.04
OGC−Oligodendrocytes
L6b 30.91 9.29 6.11 4.42 3.44 2.75 2.20 1.83 1.45 1.00
L6−IT−Car3 30.58 9.02 5.99 4.45 3.46 2.84 2.31 1.89 1.51 1.00
L6−IT 32.28 9.77 6.50 4.78 3.77 2.92 2.38 1.88 1.46 1.00
L6−CT 33.15 9.56 6.33 4.63 3.58 2.88 2.32 1.82 1.48 1.00
L56−NP 22.64 6.37 4.40 3.37 2.73 2.29 1.98 1.69 1.36 1.00
L5−IT 28.35 8.30 5.40 3.87 3.08 2.53 2.10 1.72 1.37 1.00
L5−ET 31.34 8.86 5.93 4.22 3.35 2.67 2.13 1.80 1.41 1.00
L4−IT 25.11 7.26 4.87 3.66 2.92 2.44 2.04 1.67 1.41 1.00
L23−IT 30.50 9.11 6.11 4.46 3.47 2.80 2.30 1.85 1.47 1.00
HIP−Misc2 26.15 7.67 5.02 3.75 2.94 2.40 2.02 1.68 1.38 1.00
HIP−Misc1 29.79 8.77 5.69 4.33 3.29 2.65 2.17 1.79 1.45 1.00
DG 36.27 11.54 7.70 5.37 4.21 3.13 2.47 1.97 1.48 1.00
CA3 35.72 11.17 7.18 5.11 3.96 2.97 2.33 1.86 1.44 1.00
CA1 36.01 11.19 7.51 5.37 4.02 3.16 2.43 1.96 1.47 1.00
Amy−Exc 29.45 9.27 6.26 4.66 3.59 2.82 2.21 1.83 1.42 1.00
Vip 29.80 7.93 4.77 3.41 2.62 2.18 1.78 1.52 1.26 1.00
THM−MB 20.29 5.44 3.85 3.03 2.53 2.20 1.91 1.64 1.37 1.00
THM−Inh 24.39 6.79 4.46 3.41 2.76 2.25 1.90 1.58 1.34 1.00
THM−Exc 34.17 10.39 6.67 4.93 3.76 2.91 2.38 1.92 1.47 1.00
SubCtx−Cplx 33.12 9.54 6.08 4.37 3.30 2.63 2.11 1.71 1.37 1.00
Sst 25.09 6.67 4.41 3.32 2.68 2.24 1.93 1.59 1.35 1.00
Sncg 27.99 7.45 4.78 3.52 2.74 2.23 1.90 1.60 1.34 1.00
Pvalb−ChC 28.19 7.46 4.81 3.59 2.88 2.43 1.98 1.66 1.37 1.00
Pvalb 25.46 6.66 4.19 3.13 2.54 2.15 1.78 1.55 1.31 1.00
PN 33.91 9.51 5.67 4.10 3.08 2.54 1.95 1.63 1.36 1.00
MSN−D2 21.09 6.18 4.28 3.34 2.71 2.30 1.93 1.66 1.39 1.00
MSN−D1 19.56 5.75 3.94 3.07 2.49 2.10 1.86 1.56 1.32 1.00
Lamp5−Lhx6 33.00 8.83 5.41 3.77 2.86 2.39 1.97 1.60 1.27 1.00
Lamp5 30.40 8.09 4.99 3.58 2.77 2.30 1.85 1.56 1.32 1.00
Foxp2 24.57 7.00 4.71 3.51 2.78 2.29 1.91 1.59 1.35 1.00
Chd7 32.04 8.63 5.25 3.76 2.92 2.36 1.95 1.62 1.35 1.00
CB 32.71 9.71 6.10 4.41 3.36 2.67 2.30 1.86 1.49 1.00
VLMC 29.03 7.97 5.15 3.99 3.18 2.64 2.26 1.79 1.37 1.00
PC 26.93 7.56 5.09 3.90 3.17 2.59 2.24 1.83 1.42 1.00
OPC 53.87 13.22 7.52 4.95 3.62 2.79 2.22 1.74 1.39 1.00
ODC 56.33 14.33 7.87 5.28 3.75 2.91 2.29 1.81 1.42 1.00
MGC 16.44 4.52 3.39 2.82 2.49 2.20 1.94 1.64 1.35 1.00
EC 19.72 5.32 3.90 3.21 2.73 2.36 2.05 1.72 1.41 1.00
ASC 40.77 10.31 5.94 4.11 3.21 2.58 2.04 1.70 1.35 1.00
0.0−0.1 0.1−0.2 0.2−0.3 0.3−0.4 0.4−0.5 0.5−0.6 0.6−0.7 0.7−0.8 0.8−0.9 0.9−1.0
Percent
0.02 0.04 0.06

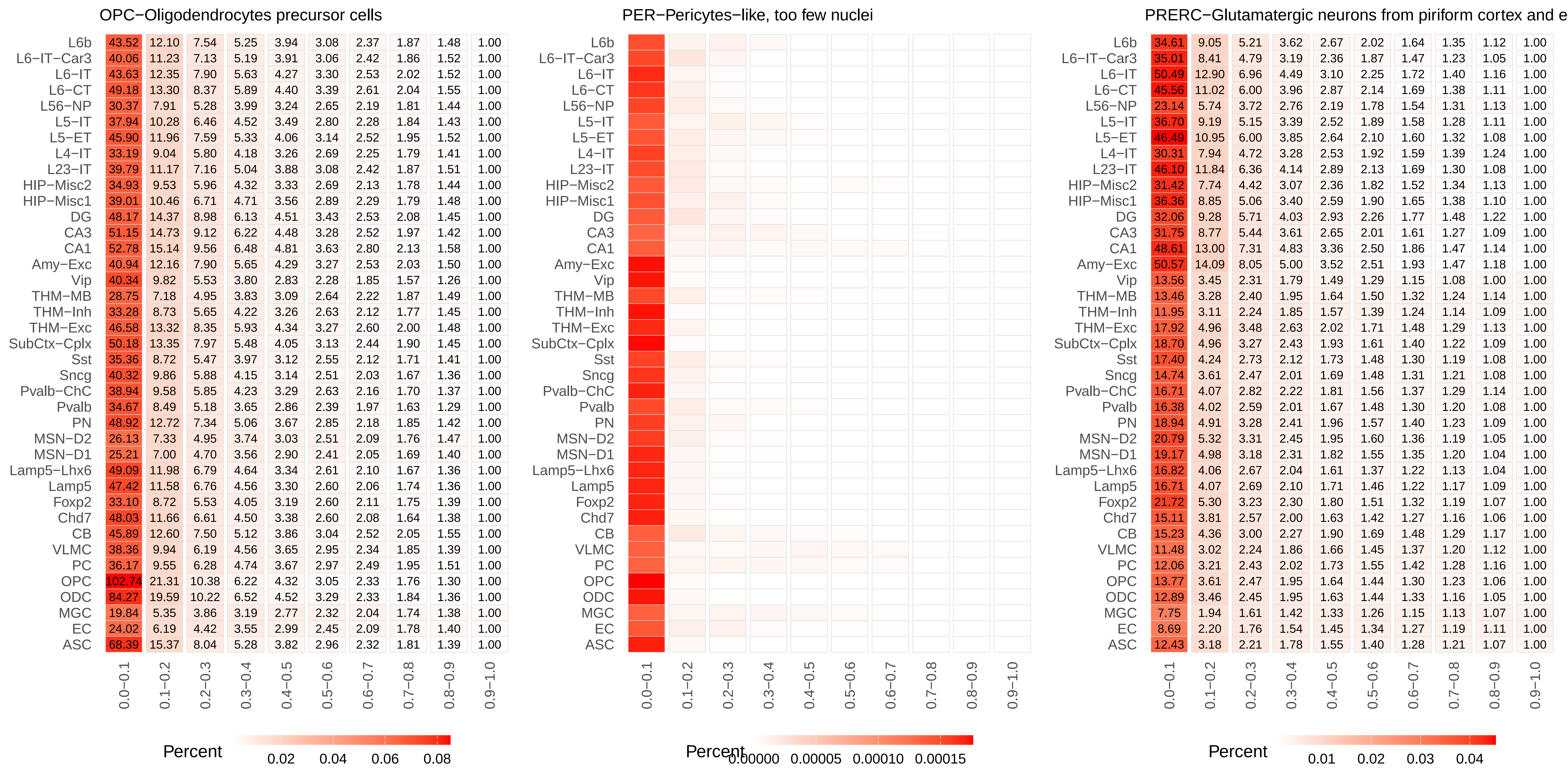
OPC−Oligodendrocytes precursor cells
L6b 43.52 12.10 7.54 5.25 3.94 3.08 2.37 1.87 1.48 1.00
L6−IT−Car3 40.06 11.23 7.13 5.19 3.91 3.06 2.42 1.86 1.52 1.00
L6−IT 43.63 12.35 7.90 5.63 4.27 3.30 2.53 2.02 1.52 1.00
L6−CT 49.18 13.30 8.37 5.89 4.40 3.39 2.61 2.04 1.55 1.00
L56−NP 30.37 7.91 5.28 3.99 3.24 2.65 2.19 1.81 1.44 1.00
L5−IT 37.94 10.28 6.46 4.52 3.49 2.80 2.28 1.84 1.43 1.00
L5−ET 45.90 11.96 7.59 5.33 4.06 3.14 2.52 1.95 1.52 1.00
L4−IT 33.19 9.04 5.80 4.18 3.26 2.69 2.25 1.79 1.41 1.00
L23−IT 39.79 11.17 7.16 5.04 3.88 3.08 2.42 1.87 1.51 1.00
HIP−Misc2 34.93 9.53 5.96 4.32 3.33 2.69 2.13 1.78 1.44 1.00
HIP−Misc1 39.01 10.46 6.71 4.71 3.56 2.89 2.29 1.79 1.48 1.00
DG 48.17 14.37 8.98 6.13 4.51 3.43 2.53 2.08 1.45 1.00
CA3 51.15 14.73 9.12 6.22 4.48 3.28 2.52 1.97 1.42 1.00
CA1 52.78 15.14 9.56 6.48 4.81 3.63 2.80 2.13 1.58 1.00
Amy−Exc 40.94 12.16 7.90 5.65 4.29 3.27 2.53 2.03 1.50 1.00
Vip 40.34 9.82 5.53 3.80 2.83 2.28 1.85 1.57 1.26 1.00
THM−MB 28.75 7.18 4.95 3.83 3.09 2.64 2.22 1.87 1.49 1.00
THM−Inh 33.28 8.73 5.65 4.22 3.26 2.63 2.12 1.77 1.45 1.00
THM−Exc 46.58 13.32 8.35 5.93 4.34 3.27 2.60 2.00 1.48 1.00
SubCtx−Cplx 50.18 13.35 7.97 5.48 4.05 3.13 2.44 1.90 1.45 1.00
Sst 35.36 8.72 5.47 3.97 3.12 2.55 2.12 1.71 1.41 1.00
Sncg 40.32 9.86 5.88 4.15 3.14 2.51 2.03 1.67 1.36 1.00
Pvalb−ChC 38.94 9.58 5.85 4.23 3.29 2.63 2.16 1.70 1.37 1.00
Pvalb 34.67 8.49 5.18 3.65 2.86 2.39 1.97 1.63 1.29 1.00
PN 48.92 12.72 7.34 5.06 3.67 2.85 2.18 1.85 1.42 1.00
MSN−D2 26.13 7.33 4.95 3.74 3.03 2.51 2.09 1.76 1.47 1.00
MSN−D1 25.21 7.00 4.70 3.56 2.90 2.41 2.05 1.69 1.40 1.00
Lamp5−Lhx6 49.09 11.98 6.79 4.64 3.34 2.61 2.10 1.67 1.36 1.00
Lamp5 47.42 11.58 6.76 4.56 3.30 2.60 2.06 1.74 1.36 1.00
Foxp2 33.10 8.72 5.53 4.05 3.19 2.60 2.11 1.75 1.39 1.00
Chd7 48.03 11.66 6.61 4.50 3.38 2.60 2.08 1.64 1.38 1.00
CB 45.89 12.60 7.50 5.12 3.86 3.04 2.52 2.05 1.55 1.00
VLMC 38.36 9.94 6.19 4.56 3.65 2.95 2.34 1.85 1.39 1.00
PC 36.17 9.55 6.28 4.74 3.67 2.97 2.49 1.95 1.51 1.00
OPC 102.74 21.31 10.38 6.22 4.32 3.05 2.33 1.76 1.30 1.00
ODC 84.27 19.59 10.22 6.52 4.52 3.29 2.33 1.84 1.36 1.00
MGC 19.84 5.35 3.86 3.19 2.77 2.32 2.04 1.74 1.38 1.00
EC 24.02 6.19 4.42 3.55 2.99 2.45 2.09 1.78 1.40 1.00
ASC 68.39 15.37 8.04 5.28 3.82 2.96 2.32 1.81 1.39 1.00
0.0−0.1 0.1−0.2 0.2−0.3 0.3−0.4 0.4−0.5 0.5−0.6 0.6−0.7 0.7−0.8 0.8−0.9 0.9−1.0
Percent 0.02 0.04 0.06 0.08
PER−Pericytes−like, too few nuclei
L6b
L6−IT−Car3
L6−IT
L6−CT
L56−NP
L5−IT
L5−ET
L4−IT
L23−IT
HIP−Misc2
HIP−Misc1
DG
CA3
CA1
Amy−Exc
Vip
THM−MB
THM−Inh
THM−Exc
SubCtx−Cplx
Sst
Sncg
Pvalb−ChC
Pvalb
PN
MSN−D2
MSN−D1
Lamp5−Lhx6
Lamp5
Foxp2
Chd7
CB
VLMC
PC
OPC
ODC
MGC
EC
ASC
0.0−0.1 0.1−0.2 0.2−0.3 0.3−0.4 0.4−0.5 0.5−0.6 0.6−0.7 0.7−0.8 0.8−0.9 0.9−1.0
Percent 0.00000 0.00005 0.00010 0.00015
PRERC−Glutamatergic neurons from piriform cortex and e
L6b 34.61 9.05 5.21 3.62 2.67 2.02 1.64 1.35 1.12 1.00
L6−IT−Car3 35.01 8.41 4.79 3.19 2.36 1.87 1.47 1.23 1.05 1.00
L6−IT 50.49 12.90 6.96 4.49 3.10 2.25 1.72 1.40 1.16 1.00
L6−CT 45.56 11.02 6.00 3.96 2.87 2.14 1.69 1.38 1.11 1.00
L56−NP 23.14 5.74 3.72 2.76 2.19 1.78 1.54 1.31 1.13 1.00
L5−IT 36.70 9.19 5.15 3.39 2.52 1.89 1.58 1.28 1.11 1.00
L5−ET 46.49 10.95 6.00 3.85 2.64 2.10 1.60 1.32 1.08 1.00
L4−IT 30.31 7.94 4.72 3.28 2.53 1.92 1.59 1.39 1.24 1.00
L23−IT 46.10 11.84 6.36 4.14 2.89 2.13 1.69 1.30 1.08 1.00
HIP−Misc2 31.42 7.74 4.42 3.07 2.36 1.82 1.52 1.34 1.13 1.00
HIP−Misc1 36.36 8.85 5.06 3.40 2.59 1.90 1.65 1.38 1.10 1.00
DG 32.06 9.28 5.71 4.03 2.93 2.26 1.77 1.48 1.22 1.00
CA3 31.75 8.77 5.44 3.61 2.65 2.01 1.61 1.27 1.09 1.00
CA1 48.61 13.00 7.31 4.83 3.36 2.50 1.86 1.47 1.14 1.00
Amy−Exc 50.57 14.09 8.05 5.00 3.52 2.51 1.93 1.47 1.18 1.00
Vip 13.56 3.45 2.31 1.79 1.49 1.29 1.15 1.08 1.00 1.00
THM−MB 13.46 3.28 2.40 1.95 1.64 1.50 1.32 1.24 1.14 1.00
THM−Inh 11.95 3.11 2.24 1.85 1.57 1.39 1.24 1.14 1.09 1.00
THM−Exc 17.92 4.96 3.48 2.63 2.02 1.71 1.48 1.29 1.13 1.00
SubCtx−Cplx 18.70 4.96 3.27 2.43 1.93 1.61 1.40 1.22 1.09 1.00
Sst 17.40 4.24 2.73 2.12 1.73 1.48 1.30 1.19 1.08 1.00
Sncg 14.74 3.61 2.47 2.01 1.69 1.48 1.31 1.21 1.08 1.00
Pvalb−ChC 16.71 4.07 2.82 2.22 1.81 1.56 1.37 1.29 1.14 1.00
Pvalb 16.38 4.02 2.59 2.01 1.67 1.48 1.30 1.20 1.08 1.00
PN 18.94 4.91 3.28 2.41 1.96 1.57 1.40 1.23 1.09 1.00
MSN−D2 20.79 5.32 3.31 2.45 1.95 1.60 1.36 1.19 1.05 1.00
MSN−D1 19.17 4.98 3.18 2.31 1.82 1.55 1.35 1.20 1.04 1.00
Lamp5−Lhx6 16.82 4.06 2.67 2.04 1.61 1.37 1.22 1.13 1.04 1.00
Lamp5 16.71 4.07 2.69 2.10 1.71 1.46 1.22 1.17 1.09 1.00
Foxp2 21.72 5.30 3.23 2.30 1.80 1.51 1.32 1.19 1.07 1.00
Chd7 15.11 3.81 2.57 2.00 1.63 1.42 1.27 1.16 1.06 1.00
CB 15.23 4.36 3.00 2.27 1.90 1.69 1.48 1.29 1.17 1.00
VLMC 11.48 3.02 2.24 1.86 1.66 1.45 1.37 1.20 1.12 1.00
PC 12.06 3.21 2.43 2.02 1.73 1.55 1.42 1.28 1.16 1.00
OPC 13.77 3.61 2.47 1.95 1.64 1.44 1.30 1.23 1.06 1.00
ODC 12.89 3.46 2.45 1.95 1.63 1.44 1.33 1.16 1.05 1.00
MGC 7.75 1.94 1.61 1.42 1.33 1.26 1.15 1.13 1.07 1.00
EC 8.69 2.20 1.76 1.54 1.45 1.34 1.27 1.19 1.11 1.00
ASC 12.43 3.18 2.21 1.78 1.55 1.40 1.28 1.21 1.07 1.00
0.0−0.1 0.1−0.2 0.2−0.3 0.3−0.4 0.4−0.5 0.5−0.6 0.6−0.7 0.7−0.8 0.8−0.9 0.9−1.0
Percent 0.01 0.02 0.03 0.04

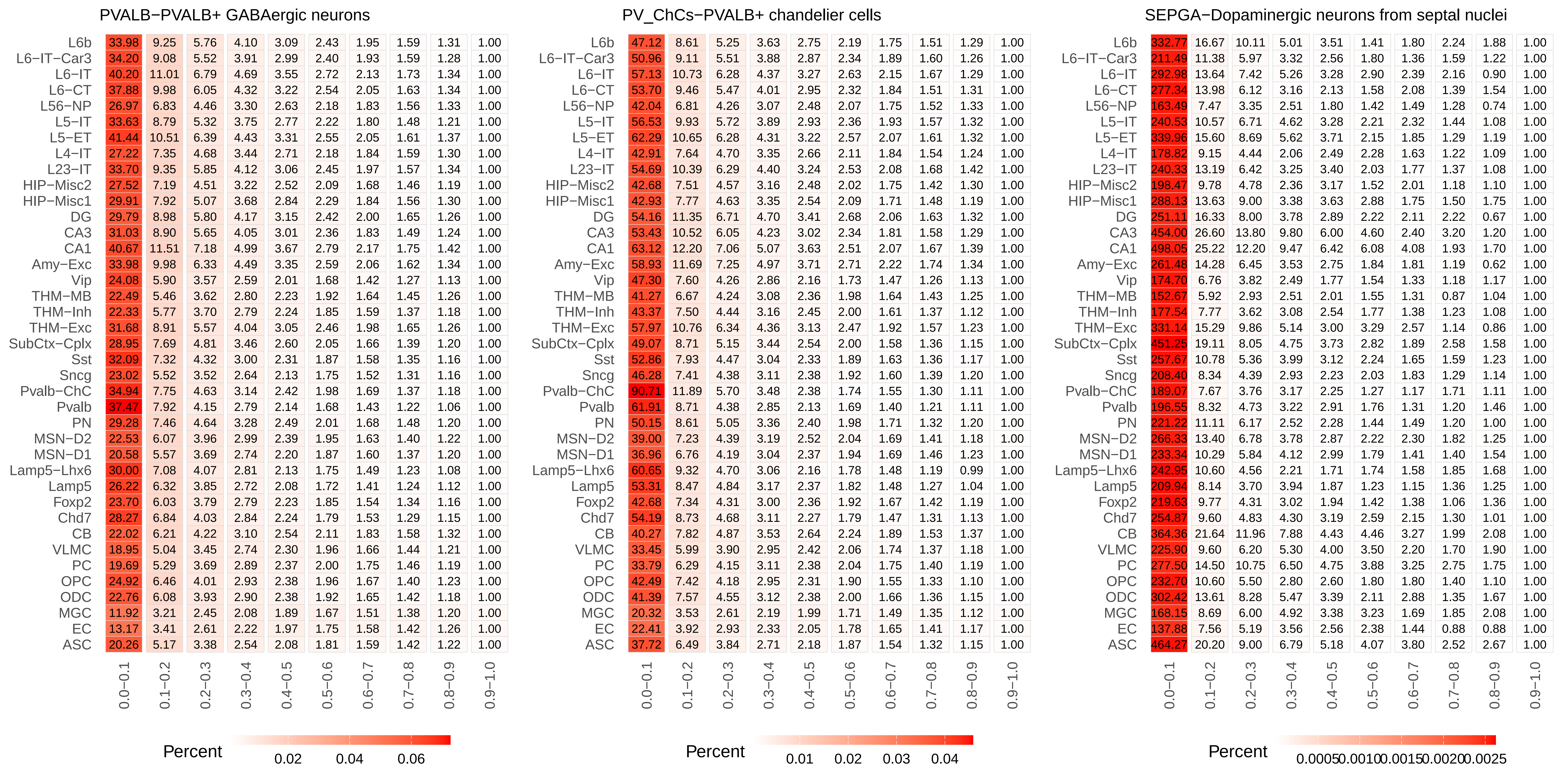
PVALB−PVALB+ GABAergic neurons
L6b 33.98 9.25 5.76 4.10 3.09 2.43 1.95 1.59 1.31 1.00
L6−IT−Car3 34.20 9.08 5.52 3.91 2.99 2.40 1.93 1.59 1.28 1.00
L6−IT 40.20 11.01 6.79 4.69 3.55 2.72 2.13 1.73 1.34 1.00
L6−CT 37.88 9.98 6.05 4.32 3.22 2.54 2.05 1.63 1.34 1.00
L56−NP 26.97 6.83 4.46 3.30 2.63 2.18 1.83 1.56 1.33 1.00
L5−IT 33.63 8.79 5.32 3.75 2.77 2.22 1.80 1.48 1.21 1.00
L5−ET 41.44 10.51 6.39 4.43 3.31 2.55 2.05 1.61 1.37 1.00
L4−IT 27.22 7.35 4.68 3.44 2.71 2.18 1.84 1.59 1.30 1.00
L23−IT 33.70 9.35 5.85 4.12 3.06 2.45 1.97 1.57 1.34 1.00
HIP−Misc2 27.52 7.19 4.51 3.22 2.52 2.09 1.68 1.46 1.19 1.00
HIP−Misc1 29.91 7.92 5.07 3.68 2.84 2.29 1.84 1.56 1.30 1.00
DG 29.79 8.98 5.80 4.17 3.15 2.42 2.00 1.65 1.26 1.00
CA3 31.03 8.90 5.65 4.05 3.01 2.36 1.83 1.49 1.24 1.00
CA1 40.67 11.51 7.18 4.99 3.67 2.79 2.17 1.75 1.42 1.00
Amy−Exc 33.98 9.98 6.33 4.49 3.35 2.59 2.06 1.62 1.34 1.00
Vip 24.08 5.90 3.57 2.59 2.01 1.68 1.42 1.27 1.13 1.00
THM−MB 22.49 5.46 3.62 2.80 2.23 1.92 1.64 1.45 1.26 1.00
THM−Inh 22.33 5.77 3.70 2.79 2.24 1.85 1.59 1.37 1.18 1.00
THM−Exc 31.68 8.91 5.57 4.04 3.05 2.46 1.98 1.65 1.26 1.00
SubCtx−Cplx 28.95 7.69 4.81 3.46 2.60 2.05 1.66 1.39 1.20 1.00
Sst 32.09 7.32 4.32 3.00 2.31 1.87 1.58 1.35 1.16 1.00
Sncg 23.02 5.52 3.52 2.64 2.13 1.75 1.52 1.31 1.16 1.00
Pvalb−ChC 34.94 7.75 4.63 3.14 2.42 1.98 1.69 1.37 1.18 1.00
Pvalb 37.47 7.92 4.15 2.79 2.14 1.68 1.43 1.22 1.06 1.00
PN 29.28 7.46 4.64 3.28 2.49 2.01 1.68 1.48 1.20 1.00
MSN−D2 22.53 6.07 3.96 2.99 2.39 1.95 1.63 1.40 1.22 1.00
MSN−D1 20.58 5.57 3.69 2.74 2.20 1.87 1.60 1.37 1.20 1.00
Lamp5−Lhx6 30.00 7.08 4.07 2.81 2.13 1.75 1.49 1.23 1.08 1.00
Lamp5 26.22 6.32 3.85 2.72 2.08 1.72 1.41 1.24 1.12 1.00
Foxp2 23.70 6.03 3.79 2.79 2.23 1.85 1.54 1.34 1.16 1.00
Chd7 28.27 6.84 4.03 2.84 2.24 1.79 1.53 1.29 1.15 1.00
CB 22.02 6.21 4.22 3.10 2.54 2.11 1.83 1.58 1.32 1.00
VLMC 18.95 5.04 3.45 2.74 2.30 1.96 1.66 1.44 1.21 1.00
PC 19.69 5.29 3.69 2.89 2.37 2.00 1.75 1.46 1.19 1.00
OPC 24.92 6.46 4.01 2.93 2.38 1.96 1.67 1.40 1.23 1.00
ODC 22.76 6.08 3.93 2.90 2.38 1.92 1.65 1.42 1.18 1.00
MGC 11.92 3.21 2.45 2.08 1.89 1.67 1.51 1.38 1.20 1.00
EC 13.17 3.41 2.61 2.22 1.97 1.75 1.58 1.42 1.26 1.00
ASC 20.26 5.17 3.38 2.54 2.08 1.81 1.59 1.42 1.22 1.00
0.0−0.1 0.1−0.2 0.2−0.3 0.3−0.4 0.4−0.5 0.5−0.6 0.6−0.7 0.7−0.8 0.8−0.9 0.9−1.0
Percent 0.02 0.04 0.06
PV_ChCs−PVALB+ chandelier cells
L6b 47.12 8.61 5.25 3.63 2.75 2.19 1.75 1.51 1.29 1.00
L6−IT−Car3 50.96 9.11 5.51 3.88 2.87 2.34 1.89 1.60 1.26 1.00
L6−IT 57.13 10.73 6.28 4.37 3.27 2.63 2.15 1.67 1.29 1.00
L6−CT 53.70 9.46 5.47 4.01 2.95 2.32 1.84 1.51 1.31 1.00
L56−NP 42.04 6.81 4.26 3.07 2.48 2.07 1.75 1.52 1.33 1.00
L5−IT 56.53 9.93 5.72 3.89 2.93 2.36 1.93 1.57 1.32 1.00
L5−ET 62.29 10.65 6.28 4.31 3.22 2.57 2.07 1.61 1.32 1.00
L4−IT 42.91 7.64 4.70 3.35 2.66 2.11 1.84 1.54 1.24 1.00
L23−IT 54.69 10.39 6.29 4.40 3.24 2.53 2.08 1.68 1.42 1.00
HIP−Misc2 42.68 7.51 4.57 3.16 2.48 2.02 1.75 1.42 1.30 1.00
HIP−Misc1 42.93 7.77 4.63 3.35 2.54 2.09 1.71 1.48 1.19 1.00
DG 54.16 11.35 6.71 4.70 3.41 2.68 2.06 1.63 1.32 1.00
CA3 53.43 10.52 6.05 4.23 3.02 2.34 1.81 1.58 1.29 1.00
CA1 63.12 12.20 7.06 5.07 3.63 2.51 2.07 1.67 1.39 1.00
Amy−Exc 58.93 11.69 7.25 4.97 3.71 2.71 2.22 1.74 1.34 1.00
Vip 47.30 7.60 4.26 2.86 2.16 1.73 1.47 1.26 1.13 1.00
THM−MB 41.27 6.67 4.24 3.08 2.36 1.98 1.64 1.43 1.25 1.00
THM−Inh 43.37 7.50 4.44 3.16 2.45 2.00 1.61 1.37 1.12 1.00
THM−Exc 57.97 10.76 6.34 4.36 3.13 2.47 1.92 1.57 1.23 1.00
SubCtx−Cplx 49.07 8.71 5.15 3.44 2.54 2.00 1.58 1.36 1.15 1.00
Sst 52.86 7.93 4.47 3.04 2.33 1.89 1.63 1.36 1.17 1.00
Sncg 46.28 7.41 4.38 3.11 2.38 1.92 1.60 1.39 1.20 1.00
Pvalb−ChC 90.71 11.89 5.70 3.48 2.38 1.74 1.55 1.30 1.11 1.00
Pvalb 61.91 8.71 4.38 2.85 2.13 1.69 1.40 1.21 1.11 1.00
PN 50.15 8.61 5.05 3.36 2.40 1.98 1.71 1.32 1.20 1.00
MSN−D2 39.00 7.23 4.39 3.19 2.52 2.04 1.69 1.41 1.18 1.00
MSN−D1 36.96 6.76 4.19 3.04 2.37 1.94 1.69 1.46 1.23 1.00
Lamp5−Lhx6 60.65 9.32 4.70 3.06 2.16 1.78 1.48 1.19 0.99 1.00
Lamp5 53.31 8.47 4.84 3.17 2.37 1.82 1.48 1.27 1.04 1.00
Foxp2 42.68 7.34 4.31 3.00 2.36 1.92 1.67 1.42 1.19 1.00
Chd7 54.19 8.73 4.68 3.11 2.27 1.79 1.47 1.31 1.13 1.00
CB 40.27 7.82 4.87 3.53 2.64 2.24 1.89 1.53 1.37 1.00
VLMC 33.45 5.99 3.90 2.95 2.42 2.06 1.74 1.37 1.18 1.00
PC 33.79 6.29 4.15 3.11 2.38 2.04 1.75 1.40 1.19 1.00
OPC 42.49 7.42 4.18 2.95 2.31 1.90 1.55 1.33 1.10 1.00
ODC 41.39 7.57 4.55 3.12 2.38 2.00 1.66 1.36 1.15 1.00
MGC 20.32 3.53 2.61 2.19 1.99 1.71 1.49 1.35 1.12 1.00
EC 22.41 3.92 2.93 2.33 2.05 1.78 1.65 1.41 1.17 1.00
ASC 37.72 6.49 3.84 2.71 2.18 1.87 1.54 1.32 1.15 1.00
0.0−0.1 0.1−0.2 0.2−0.3 0.3−0.4 0.4−0.5 0.5−0.6 0.6−0.7 0.7−0.8 0.8−0.9 0.9−1.0
Percent 0.01 0.02 0.03 0.04
SEPGA−Dopaminergic neurons from septal nuclei
L6b 332.77 16.67 10.11 5.01 3.51 1.41 1.80 2.24 1.88 1.00
L6−IT−Car3 211.49 11.38 5.97 3.32 2.56 1.80 1.36 1.59 1.22 1.00
L6−IT 292.98 13.64 7.42 5.26 3.28 2.90 2.39 2.16 0.90 1.00
L6−CT 277.34 13.98 6.12 3.16 2.13 1.58 2.08 1.39 1.54 1.00
L56−NP 163.49 7.47 3.35 2.51 1.80 1.42 1.49 1.28 0.74 1.00
L5−IT 240.53 10.57 6.71 4.62 3.28 2.21 2.32 1.44 1.08 1.00
L5−ET 339.96 15.60 8.69 5.62 3.71 2.15 1.85 1.29 1.19 1.00
L4−IT 178.82 9.15 4.44 2.06 2.49 2.28 1.63 1.22 1.09 1.00
L23−IT 240.33 13.19 6.42 3.25 3.40 2.03 1.77 1.37 1.08 1.00
HIP−Misc2 198.47 9.78 4.78 2.36 3.17 1.52 2.01 1.18 1.10 1.00
HIP−Misc1 288.13 13.63 9.00 3.38 3.63 2.88 1.75 1.50 1.75 1.00
DG 251.11 16.33 8.00 3.78 2.89 2.22 2.11 2.22 0.67 1.00
CA3 454.00 26.60 13.80 9.80 6.00 4.60 2.40 3.20 1.20 1.00
CA1 498.05 25.22 12.20 9.47 6.42 6.08 4.08 1.93 1.70 1.00
Amy−Exc 261.48 14.28 6.45 3.53 2.75 1.84 1.81 1.19 0.62 1.00
Vip 174.70 6.76 3.82 2.49 1.77 1.54 1.33 1.18 1.17 1.00
THM−MB 152.67 5.92 2.93 2.51 2.01 1.55 1.31 0.87 1.04 1.00
THM−Inh 177.54 7.77 3.62 3.08 2.54 1.77 1.38 1.23 1.08 1.00
THM−Exc 331.14 15.29 9.86 5.14 3.00 3.29 2.57 1.14 0.86 1.00
SubCtx−Cplx 451.25 19.11 8.05 4.75 3.73 2.82 1.89 2.58 1.58 1.00
Sst 257.67 10.78 5.36 3.99 3.12 2.24 1.65 1.59 1.23 1.00
Sncg 208.40 8.34 4.39 2.93 2.23 2.03 1.83 1.29 1.14 1.00
Pvalb−ChC 189.07 7.67 3.76 3.17 2.25 1.27 1.17 1.71 1.11 1.00
Pvalb 196.55 8.32 4.73 3.22 2.91 1.76 1.31 1.20 1.46 1.00
PN 221.22 11.11 6.17 2.52 2.28 1.44 1.49 1.20 1.00 1.00
MSN−D2 266.33 13.40 6.78 3.78 2.87 2.22 2.30 1.82 1.25 1.00
MSN−D1 233.34 10.29 5.84 4.12 2.99 1.79 1.41 1.40 1.54 1.00
Lamp5−Lhx6 242.95 10.60 4.56 2.21 1.71 1.74 1.58 1.85 1.68 1.00
Lamp5 209.94 8.14 3.70 3.94 1.87 1.23 1.15 1.36 1.25 1.00
Foxp2 219.63 9.77 4.31 3.02 1.94 1.42 1.38 1.06 1.36 1.00
Chd7 254.87 9.60 4.83 4.30 3.19 2.59 2.15 1.30 1.01 1.00
CB 364.36 21.64 11.96 7.88 4.43 4.46 3.27 1.99 2.08 1.00
VLMC 225.90 9.60 6.20 5.30 4.00 3.50 2.20 1.70 1.90 1.00
PC 277.50 14.50 10.75 6.50 4.75 3.88 3.25 2.75 1.75 1.00
OPC 232.70 10.60 5.50 2.80 2.60 1.80 1.80 1.40 1.10 1.00
ODC 302.42 13.61 8.28 5.47 3.39 2.11 2.88 1.35 1.67 1.00
MGC 168.15 8.69 6.00 4.92 3.38 3.23 1.69 1.85 2.08 1.00
EC 137.88 7.56 5.19 3.56 2.56 2.38 1.44 0.88 0.88 1.00
ASC 464.27 20.20 9.00 6.79 5.18 4.07 3.80 2.52 2.67 1.00
0.0−0.1 0.1−0.2 0.2−0.3 0.3−0.4 0.4−0.5 0.5−0.6 0.6−0.7 0.7−0.8 0.8−0.9 0.9−1.0
Percent 0.0005 0.0010 0.0015 0.0020 0.0025

#### SIGA−Dopaminergic neurons from Inferior colliculus and nearby nuclei −

| | 0.0−0.1 | 0.1−0.2 | 0.2−0.3 | 0.3−0.4 | 0.4−0.5 | 0.5−0.6 | 0.6−0.7 | 0.7−0.8 | 0.8−0.9 | 0.9−1.0 |
|---|---|---|---|---|---|---|---|---|---|---|
| L6b | 178.95 | 11.92 | 6.30 | 4.13 | 3.19 | 2.12 | 1.75 | 1.46 | 1.08 | 1.00 |
| L6−IT−Car3 | 151.06 | 10.23 | 5.43 | 3.45 | 2.37 | 2.25 | 1.82 | 1.50 | 1.29 | 1.00 |
| L6−IT | 198.94 | 13.85 | 7.61 | 4.52 | 3.31 | 2.71 | 2.23 | 1.56 | 1.12 | 1.00 |
| L6−CT | 264.46 | 16.55 | 7.70 | 5.60 | 4.58 | 3.00 | 1.96 | 1.87 | 1.85 | 1.00 |
| L56−NP | 121.26 | 6.97 | 3.98 | 2.25 | 1.94 | 1.65 | 1.35 | 1.13 | 0.99 | 1.00 |
| L5−IT | 183.05 | 11.19 | 6.57 | 4.48 | 3.15 | 2.12 | 1.99 | 1.46 | 1.32 | 1.00 |
| L5−ET | 280.71 | 16.55 | 9.13 | 5.77 | 3.82 | 2.61 | 2.39 | 1.81 | 1.54 | 1.00 |
| L4−IT | 151.52 | 9.33 | 4.98 | 3.93 | 2.60 | 2.50 | 1.89 | 1.64 | 1.47 | 1.00 |
| L23−IT | 225.31 | 16.19 | 7.69 | 5.06 | 3.20 | 2.94 | 2.15 | 1.71 | 1.40 | 1.00 |
| HIP−Misc2 | 124.50 | 7.93 | 3.68 | 3.10 | 2.11 | 2.18 | 1.40 | 0.88 | 0.93 | 1.00 |
| HIP−Misc1 | 238.64 | 15.12 | 7.88 | 5.20 | 3.24 | 2.92 | 1.96 | 1.88 | 1.80 | 1.00 |
| DG | 231.60 | 17.68 | 10.00 | 6.08 | 4.12 | 3.08 | 2.60 | 1.68 | 1.80 | 1.00 |
| CA3 | 188.52 | 13.84 | 7.48 | 4.52 | 3.32 | 2.68 | 1.84 | 1.32 | 1.00 | 1.00 |
| CA1 | 277.50 | 20.10 | 9.98 | 6.40 | 4.49 | 2.94 | 2.06 | 1.52 | 1.39 | 1.00 |
| Amy−Exc | 223.19 | 17.23 | 9.08 | 5.40 | 3.58 | 2.40 | 2.04 | 1.34 | 0.99 | 1.00 |
| Vip | 123.80 | 7.18 | 3.98 | 2.66 | 2.11 | 1.71 | 1.39 | 1.32 | 1.01 | 1.00 |
| THM−MB | 133.04 | 7.52 | 3.78 | 2.75 | 1.95 | 1.81 | 1.91 | 1.59 | 1.31 | 1.00 |
| THM−Inh | 126.26 | 7.80 | 5.00 | 3.17 | 2.35 | 2.15 | 1.61 | 1.41 | 1.22 | 1.00 |
| THM−Exc | 126.11 | 9.17 | 4.48 | 3.39 | 2.57 | 1.93 | 1.48 | 1.07 | 0.78 | 1.00 |
| SubCtx−Cplx | 212.03 | 13.87 | 6.47 | 3.90 | 2.52 | 1.98 | 1.58 | 1.43 | 1.47 | 1.00 |
| Sst | 141.45 | 7.62 | 3.95 | 2.74 | 1.91 | 1.54 | 1.27 | 1.03 | 1.04 | 1.00 |
| Sncg | 113.79 | 6.23 | 3.68 | 2.63 | 2.01 | 1.62 | 1.21 | 1.02 | 1.08 | 1.00 |
| Pvalb−ChC | 142.44 | 8.52 | 4.84 | 2.73 | 2.31 | 1.67 | 1.77 | 1.41 | 1.26 | 1.00 |
| Pvalb | 106.10 | 5.71 | 3.18 | 2.18 | 1.66 | 1.47 | 1.01 | 0.87 | 1.00 | 1.00 |
| PN | 138.35 | 8.53 | 4.52 | 3.19 | 2.61 | 1.80 | 2.08 | 1.38 | 1.23 | 1.00 |
| MSN−D2 | 258.45 | 15.38 | 8.40 | 4.56 | 3.03 | 2.34 | 1.54 | 0.92 | 1.24 | 1.00 |
| MSN−D1 | 214.44 | 13.12 | 7.00 | 4.67 | 2.70 | 2.05 | 1.61 | 1.06 | 0.91 | 1.00 |
| Lamp5−Lhx6 | 145.39 | 8.08 | 4.13 | 2.75 | 2.32 | 1.64 | 1.36 | 1.19 | 0.91 | 1.00 |
| Lamp5 | 133.20 | 8.28 | 4.22 | 2.67 | 1.92 | 1.45 | 1.48 | 1.18 | 0.91 | 1.00 |
| Foxp2 | 349.17 | 21.11 | 9.18 | 5.12 | 3.28 | 2.62 | 1.58 | 1.39 | 1.64 | 1.00 |
| Chd7 | 126.70 | 7.52 | 4.15 | 2.76 | 1.93 | 1.56 | 1.34 | 1.19 | 1.01 | 1.00 |
| CB | 122.56 | 8.52 | 5.56 | 3.72 | 2.61 | 2.32 | 1.75 | 1.60 | 1.65 | 1.00 |
| VLMC | 93.70 | 5.92 | 4.13 | 3.08 | 2.48 | 2.05 | 1.57 | 1.53 | 1.05 | 1.00 |
| PC | 109.86 | 7.06 | 4.37 | 3.24 | 3.08 | 2.65 | 2.25 | 1.90 | 1.67 | 1.00 |
| OPC | 135.91 | 9.09 | 4.74 | 3.07 | 3.00 | 1.88 | 1.40 | 1.30 | 1.19 | 1.00 |
| ODC | 155.22 | 10.50 | 5.95 | 4.06 | 2.82 | 2.45 | 2.20 | 1.66 | 1.43 | 1.00 |
| MGC | 58.69 | 3.11 | 2.41 | 2.03 | 1.90 | 1.57 | 1.34 | 1.19 | 1.12 | 1.00 |
| EC | 73.84 | 4.45 | 2.73 | 2.69 | 2.20 | 1.81 | 1.64 | 1.41 | 1.43 | 1.00 |
| ASC | 128.22 | 8.29 | 4.62 | 3.19 | 2.26 | 1.94 | 1.37 | 1.29 | 1.30 | 1.00 |

Percent 0.002 0.004 0.006

#### SMC−Vascular smooth muscle cells

| | 0.0−0.1 | 0.1−0.2 | 0.2−0.3 | 0.3−0.4 | 0.4−0.5 | 0.5−0.6 | 0.6−0.7 | 0.7−0.8 | 0.8−0.9 | 0.9−1.0 |
|---|---|---|---|---|---|---|---|---|---|---|
| L6b | 109.15 | 9.81 | 6.81 | 5.28 | 4.17 | 3.46 | 2.66 | 2.07 | 1.83 | 1.00 |
| L6−IT−Car3 | 147.08 | 13.24 | 8.87 | 6.69 | 4.90 | 4.06 | 3.44 | 2.61 | 1.88 | 1.00 |
| L6−IT | 101.29 | 9.25 | 6.42 | 4.87 | 3.77 | 2.92 | 2.36 | 2.07 | 1.61 | 1.00 |
| L6−CT | 114.06 | 10.47 | 7.07 | 5.10 | 4.19 | 3.20 | 2.79 | 2.29 | 1.64 | 1.00 |
| L56−NP | 89.59 | 7.66 | 5.24 | 4.14 | 3.32 | 2.76 | 2.48 | 2.04 | 1.47 | 1.00 |
| L5−IT | 91.32 | 8.33 | 5.47 | 4.19 | 3.37 | 2.82 | 2.30 | 1.95 | 1.71 | 1.00 |
| L5−ET | 91.02 | 7.97 | 5.21 | 4.24 | 3.30 | 2.90 | 2.35 | 1.97 | 1.58 | 1.00 |
| L4−IT | 99.45 | 8.90 | 5.97 | 4.66 | 3.69 | 3.21 | 2.55 | 2.06 | 1.71 | 1.00 |
| L23−IT | 110.59 | 10.03 | 6.64 | 5.29 | 4.21 | 3.41 | 2.83 | 2.21 | 1.77 | 1.00 |
| HIP−Misc2 | 106.08 | 9.30 | 6.35 | 4.87 | 4.04 | 3.22 | 2.93 | 2.20 | 1.84 | 1.00 |
| HIP−Misc1 | 105.36 | 9.29 | 6.16 | 4.74 | 3.79 | 3.21 | 2.65 | 2.22 | 1.74 | 1.00 |
| DG | 115.89 | 11.57 | 7.21 | 5.64 | 4.40 | 3.38 | 2.92 | 2.43 | 1.76 | 1.00 |
| CA3 | 107.68 | 10.14 | 7.18 | 5.55 | 4.00 | 3.30 | 2.45 | 2.25 | 1.58 | 1.00 |
| CA1 | 107.82 | 10.06 | 6.79 | 5.21 | 4.29 | 3.63 | 2.97 | 2.29 | 1.73 | 1.00 |
| Amy−Exc | 110.94 | 10.16 | 6.70 | 5.18 | 4.22 | 3.32 | 2.76 | 2.26 | 1.88 | 1.00 |
| Vip | 73.99 | 6.18 | 4.50 | 3.37 | 2.91 | 2.46 | 2.06 | 1.75 | 1.42 | 1.00 |
| THM−MB | 67.33 | 5.32 | 4.09 | 3.23 | 3.12 | 2.70 | 2.37 | 1.85 | 1.56 | 1.00 |
| THM−Inh | 82.22 | 6.83 | 4.58 | 3.98 | 3.35 | 2.75 | 2.31 | 1.83 | 1.76 | 1.00 |
| THM−Exc | 129.95 | 12.80 | 8.18 | 6.51 | 4.59 | 3.79 | 2.79 | 2.10 | 1.72 | 1.00 |
| SubCtx−Cplx | 101.01 | 8.79 | 6.14 | 4.57 | 3.72 | 2.98 | 2.48 | 2.17 | 1.60 | 1.00 |
| Sst | 88.85 | 7.25 | 4.98 | 4.07 | 3.51 | 2.90 | 2.55 | 1.98 | 1.71 | 1.00 |
| Sncg | 62.89 | 5.30 | 3.88 | 3.19 | 2.55 | 2.16 | 1.95 | 1.64 | 1.37 | 1.00 |
| Pvalb−ChC | 85.03 | 6.69 | 4.83 | 3.88 | 3.15 | 2.83 | 2.56 | 2.13 | 1.60 | 1.00 |
| Pvalb | 78.54 | 6.28 | 4.36 | 3.52 | 2.80 | 2.53 | 2.20 | 1.86 | 1.57 | 1.00 |
| PN | 106.16 | 9.64 | 6.28 | 4.59 | 3.97 | 3.18 | 2.75 | 2.33 | 1.72 | 1.00 |
| MSN−D2 | 85.82 | 7.56 | 5.00 | 3.68 | 3.21 | 2.75 | 2.31 | 1.89 | 1.70 | 1.00 |
| MSN−D1 | 85.26 | 7.34 | 5.17 | 4.07 | 3.26 | 2.51 | 2.27 | 1.92 | 1.50 | 1.00 |
| Lamp5−Lhx6 | 75.12 | 6.30 | 4.39 | 3.18 | 2.81 | 2.46 | 2.19 | 1.91 | 1.59 | 1.00 |
| Lamp5 | 63.58 | 5.08 | 3.73 | 3.01 | 2.36 | 2.20 | 2.03 | 1.69 | 1.36 | 1.00 |
| Foxp2 | 90.24 | 7.67 | 5.18 | 3.94 | 3.38 | 2.77 | 2.24 | 1.91 | 1.58 | 1.00 |
| Chd7 | 78.43 | 6.75 | 4.75 | 3.64 | 2.93 | 2.42 | 2.16 | 1.92 | 1.49 | 1.00 |
| CB | 95.27 | 9.80 | 5.89 | 4.91 | 3.81 | 2.93 | 2.58 | 2.08 | 1.56 | 1.00 |
| VLMC | 106.69 | 9.35 | 6.42 | 4.97 | 4.32 | 3.14 | 2.56 | 1.95 | 1.70 | 1.00 |
| PC | 99.84 | 9.10 | 5.93 | 4.66 | 3.75 | 2.88 | 2.51 | 1.86 | 1.35 | 1.00 |
| OPC | 110.38 | 9.95 | 6.42 | 5.06 | 4.44 | 3.40 | 2.91 | 2.06 | 1.61 | 1.00 |
| ODC | 92.98 | 8.41 | 6.00 | 4.26 | 3.65 | 3.04 | 2.50 | 2.01 | 1.46 | 1.00 |
| MGC | 116.59 | 10.36 | 6.42 | 4.95 | 3.63 | 3.08 | 2.58 | 1.59 | 1.40 | 1.00 |
| EC | 124.33 | 10.38 | 7.62 | 5.58 | 4.37 | 3.42 | 2.79 | 2.25 | 1.98 | 1.00 |
| ASC | 105.61 | 9.04 | 5.93 | 4.88 | 3.85 | 3.37 | 3.00 | 2.41 | 1.79 | 1.00 |

Percent 0.005 0.010 0.015

#### SNCG−SNCG+ GABAergic neurons

| | 0.0−0.1 | 0.1−0.2 | 0.2−0.3 | 0.3−0.4 | 0.4−0.5 | 0.5−0.6 | 0.6−0.7 | 0.7−0.8 | 0.8−0.9 | 0.9−1.0 |
|---|---|---|---|---|---|---|---|---|---|---|
| L6b | 27.84 | 6.08 | 3.81 | 2.72 | 2.18 | 1.81 | 1.50 | 1.26 | 1.12 | 1.00 |
| L6−IT−Car3 | 23.37 | 4.81 | 3.21 | 2.37 | 2.00 | 1.68 | 1.45 | 1.34 | 1.19 | 1.00 |
| L6−IT | 27.88 | 6.08 | 3.97 | 2.84 | 2.22 | 1.93 | 1.61 | 1.36 | 1.16 | 1.00 |
| L6−CT | 27.10 | 5.62 | 3.61 | 2.66 | 2.14 | 1.80 | 1.51 | 1.31 | 1.11 | 1.00 |
| L56−NP | 18.90 | 3.73 | 2.57 | 2.03 | 1.73 | 1.57 | 1.42 | 1.22 | 1.13 | 1.00 |
| L5−IT | 25.18 | 5.22 | 3.37 | 2.49 | 2.01 | 1.67 | 1.48 | 1.29 | 1.12 | 1.00 |
| L5−ET | 34.24 | 6.95 | 4.34 | 3.15 | 2.49 | 1.96 | 1.64 | 1.41 | 1.22 | 1.00 |
| L4−IT | 20.19 | 4.21 | 2.79 | 2.15 | 1.79 | 1.55 | 1.40 | 1.29 | 1.13 | 1.00 |
| L23−IT | 26.15 | 5.79 | 3.72 | 2.76 | 2.19 | 1.82 | 1.55 | 1.38 | 1.17 | 1.00 |
| HIP−Misc2 | 21.65 | 4.49 | 2.89 | 2.21 | 1.82 | 1.58 | 1.46 | 1.23 | 1.09 | 1.00 |
| HIP−Misc1 | 25.81 | 5.43 | 3.46 | 2.58 | 2.11 | 1.75 | 1.50 | 1.27 | 1.13 | 1.00 |
| DG | 32.41 | 7.70 | 4.67 | 3.44 | 2.69 | 2.16 | 1.79 | 1.41 | 1.21 | 1.00 |
| CA3 | 29.02 | 6.55 | 4.06 | 2.93 | 2.40 | 1.86 | 1.54 | 1.44 | 1.18 | 1.00 |
| CA1 | 35.70 | 8.00 | 5.03 | 3.55 | 2.75 | 2.16 | 1.81 | 1.51 | 1.31 | 1.00 |
| Amy−Exc | 28.74 | 6.58 | 4.30 | 3.17 | 2.51 | 2.01 | 1.74 | 1.44 | 1.18 | 1.00 |
| Vip | 32.79 | 6.02 | 3.35 | 2.34 | 1.81 | 1.49 | 1.27 | 1.15 | 1.03 | 1.00 |
| THM−MB | 19.78 | 3.74 | 2.56 | 2.08 | 1.77 | 1.59 | 1.37 | 1.28 | 1.17 | 1.00 |
| THM−Inh | 21.73 | 4.43 | 2.84 | 2.22 | 1.87 | 1.58 | 1.42 | 1.30 | 1.16 | 1.00 |
| THM−Exc | 22.91 | 4.94 | 3.36 | 2.43 | 2.07 | 1.72 | 1.49 | 1.28 | 1.17 | 1.00 |
| SubCtx−Cplx | 27.61 | 5.77 | 3.66 | 2.65 | 2.04 | 1.68 | 1.43 | 1.23 | 1.06 | 1.00 |
| Sst | 25.65 | 4.81 | 3.00 | 2.22 | 1.85 | 1.59 | 1.42 | 1.27 | 1.13 | 1.00 |
| Sncg | 45.84 | 8.31 | 4.38 | 2.78 | 2.09 | 1.67 | 1.35 | 1.24 | 1.05 | 1.00 |
| Pvalb−ChC | 28.99 | 5.46 | 3.32 | 2.49 | 1.95 | 1.64 | 1.44 | 1.28 | 1.19 | 1.00 |
| Pvalb | 25.49 | 4.63 | 2.92 | 2.13 | 1.76 | 1.50 | 1.38 | 1.22 | 1.11 | 1.00 |
| PN | 25.14 | 5.08 | 3.16 | 2.27 | 1.82 | 1.48 | 1.36 | 1.17 | 1.09 | 1.00 |
| MSN−D2 | 19.76 | 4.14 | 2.87 | 2.29 | 1.90 | 1.64 | 1.42 | 1.25 | 1.16 | 1.00 |
| MSN−D1 | 18.64 | 3.87 | 2.69 | 2.11 | 1.74 | 1.55 | 1.41 | 1.27 | 1.21 | 1.00 |
| Lamp5−Lhx6 | 41.33 | 7.43 | 3.92 | 2.53 | 1.89 | 1.52 | 1.24 | 1.10 | 0.94 | 1.00 |
| Lamp5 | 45.99 | 8.33 | 4.29 | 2.77 | 2.00 | 1.58 | 1.34 | 1.13 | 1.04 | 1.00 |
| Foxp2 | 23.35 | 4.77 | 3.00 | 2.27 | 1.85 | 1.54 | 1.37 | 1.22 | 1.09 | 1.00 |
| Chd7 | 32.98 | 6.21 | 3.51 | 2.49 | 1.86 | 1.59 | 1.37 | 1.20 | 1.09 | 1.00 |
| CB | 24.62 | 5.48 | 3.50 | 2.59 | 2.14 | 1.76 | 1.51 | 1.32 | 1.19 | 1.00 |
| VLMC | 17.75 | 3.71 | 2.48 | 1.95 | 1.71 | 1.53 | 1.36 | 1.21 | 1.06 | 1.00 |
| PC | 18.67 | 3.82 | 2.73 | 2.20 | 1.87 | 1.67 | 1.46 | 1.31 | 1.15 | 1.00 |
| OPC | 29.57 | 5.75 | 3.29 | 2.39 | 1.94 | 1.53 | 1.28 | 1.21 | 1.09 | 1.00 |
| ODC | 29.02 | 6.00 | 3.69 | 2.68 | 2.08 | 1.67 | 1.46 | 1.21 | 1.11 | 1.00 |
| MGC | 11.97 | 2.41 | 1.96 | 1.65 | 1.53 | 1.39 | 1.29 | 1.22 | 1.09 | 1.00 |
| EC | 14.23 | 2.92 | 2.18 | 1.89 | 1.62 | 1.51 | 1.38 | 1.27 | 1.11 | 1.00 |
| ASC | 27.35 | 5.28 | 3.17 | 2.33 | 1.84 | 1.56 | 1.36 | 1.25 | 1.10 | 1.00 |

Percent 0.01 0.02 0.03 0.04

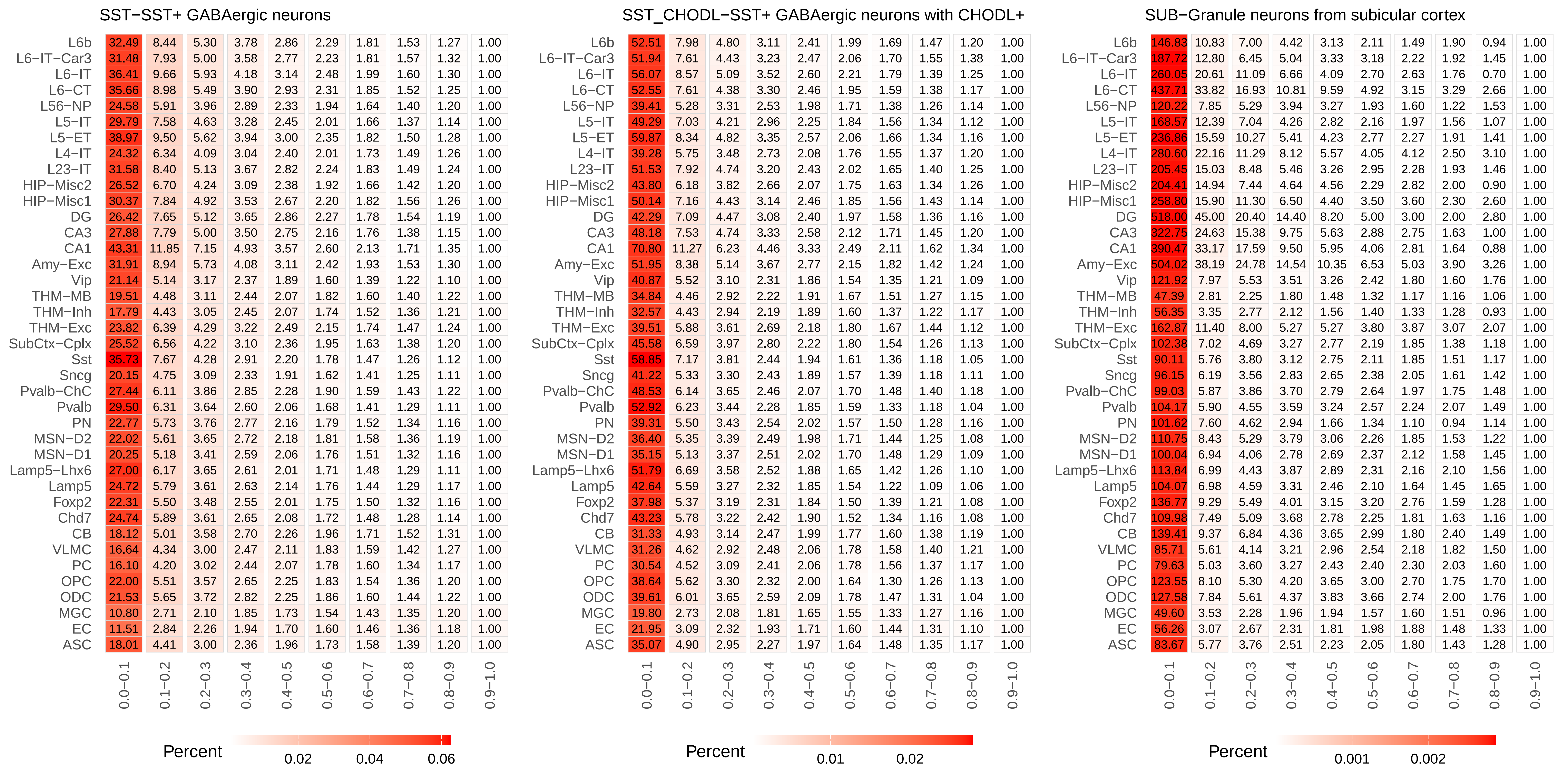
SST−SST+ GABAergic neurons
L6b 32.49 8.44 5.30 3.78 2.86 2.29 1.81 1.53 1.27 1.00
L6−IT−Car3 31.48 7.93 5.00 3.58 2.77 2.23 1.81 1.57 1.32 1.00
L6−IT 36.41 9.66 5.93 4.18 3.14 2.48 1.99 1.60 1.30 1.00
L6−CT 35.66 8.98 5.49 3.90 2.93 2.31 1.85 1.52 1.25 1.00
L56−NP 24.58 5.91 3.96 2.89 2.33 1.94 1.64 1.40 1.20 1.00
L5−IT 29.79 7.58 4.63 3.28 2.45 2.01 1.66 1.37 1.14 1.00
L5−ET 38.97 9.50 5.62 3.94 3.00 2.35 1.82 1.50 1.28 1.00
L4−IT 24.32 6.34 4.09 3.04 2.40 2.01 1.73 1.49 1.26 1.00
L23−IT 31.58 8.40 5.13 3.67 2.82 2.24 1.83 1.49 1.24 1.00
HIP−Misc2 26.52 6.70 4.24 3.09 2.38 1.92 1.66 1.42 1.20 1.00
HIP−Misc1 30.37 7.84 4.92 3.53 2.67 2.20 1.82 1.56 1.26 1.00
DG 26.42 7.65 5.12 3.65 2.86 2.27 1.78 1.54 1.19 1.00
CA3 27.88 7.79 5.00 3.50 2.75 2.16 1.76 1.38 1.15 1.00
CA1 43.31 11.85 7.15 4.93 3.57 2.60 2.13 1.71 1.35 1.00
Amy−Exc 31.91 8.94 5.73 4.08 3.11 2.42 1.93 1.53 1.30 1.00
Vip 21.14 5.14 3.17 2.37 1.89 1.60 1.39 1.22 1.10 1.00
THM−MB 19.51 4.48 3.11 2.44 2.07 1.82 1.60 1.40 1.22 1.00
THM−Inh 17.79 4.43 3.05 2.45 2.07 1.74 1.52 1.36 1.21 1.00
THM−Exc 23.82 6.39 4.29 3.22 2.49 2.15 1.74 1.47 1.24 1.00
SubCtx−Cplx 25.52 6.56 4.22 3.10 2.36 1.95 1.63 1.38 1.20 1.00
Sst 35.73 7.67 4.28 2.91 2.20 1.78 1.47 1.26 1.12 1.00
Sncg 20.15 4.75 3.09 2.33 1.91 1.62 1.41 1.25 1.11 1.00
Pvalb−ChC 27.44 6.11 3.86 2.85 2.28 1.90 1.59 1.43 1.22 1.00
Pvalb 29.50 6.31 3.64 2.60 2.06 1.68 1.41 1.29 1.11 1.00
PN 22.77 5.73 3.76 2.77 2.16 1.79 1.52 1.34 1.16 1.00
MSN−D2 22.02 5.61 3.65 2.72 2.18 1.81 1.58 1.36 1.19 1.00
MSN−D1 20.25 5.18 3.41 2.59 2.06 1.76 1.51 1.32 1.16 1.00
Lamp5−Lhx6 27.00 6.17 3.65 2.61 2.01 1.71 1.48 1.29 1.11 1.00
Lamp5 24.72 5.79 3.61 2.63 2.14 1.76 1.44 1.29 1.17 1.00
Foxp2 22.31 5.50 3.48 2.55 2.01 1.75 1.50 1.32 1.16 1.00
Chd7 24.74 5.89 3.61 2.65 2.08 1.72 1.48 1.28 1.14 1.00
CB 18.12 5.01 3.58 2.70 2.26 1.96 1.71 1.52 1.31 1.00
VLMC 16.64 4.34 3.00 2.47 2.11 1.83 1.59 1.42 1.27 1.00
PC 16.10 4.20 3.02 2.44 2.07 1.78 1.60 1.34 1.17 1.00
OPC 22.00 5.51 3.57 2.65 2.25 1.83 1.54 1.36 1.20 1.00
ODC 21.53 5.65 3.72 2.82 2.25 1.86 1.60 1.44 1.22 1.00
MGC 10.80 2.71 2.10 1.85 1.73 1.54 1.43 1.35 1.20 1.00
EC 11.51 2.84 2.26 1.94 1.70 1.60 1.46 1.36 1.18 1.00
ASC 18.01 4.41 3.00 2.36 1.96 1.73 1.58 1.39 1.20 1.00
0.0−0.1 0.1−0.2 0.2−0.3 0.3−0.4 0.4−0.5 0.5−0.6 0.6−0.7 0.7−0.8 0.8−0.9 0.9−1.0
Percent 0.02 0.04 0.06
SST_CHODL−SST+ GABAergic neurons with CHODL+
L6b 52.51 7.98 4.80 3.11 2.41 1.99 1.69 1.47 1.20 1.00
L6−IT−Car3 51.94 7.61 4.43 3.23 2.47 2.06 1.70 1.55 1.38 1.00
L6−IT 56.07 8.57 5.09 3.52 2.60 2.21 1.79 1.39 1.25 1.00
L6−CT 52.55 7.61 4.38 3.30 2.46 1.95 1.59 1.38 1.17 1.00
L56−NP 39.41 5.28 3.31 2.53 1.98 1.71 1.38 1.26 1.14 1.00
L5−IT 49.29 7.03 4.21 2.96 2.25 1.84 1.56 1.34 1.12 1.00
L5−ET 59.87 8.34 4.82 3.35 2.57 2.06 1.66 1.34 1.16 1.00
L4−IT 39.28 5.75 3.48 2.73 2.08 1.76 1.55 1.37 1.20 1.00
L23−IT 51.53 7.92 4.74 3.20 2.43 2.02 1.65 1.40 1.25 1.00
HIP−Misc2 43.80 6.18 3.82 2.66 2.07 1.75 1.63 1.34 1.26 1.00
HIP−Misc1 50.14 7.16 4.43 3.14 2.46 1.85 1.56 1.43 1.14 1.00
DG 42.29 7.09 4.47 3.08 2.40 1.97 1.58 1.36 1.16 1.00
CA3 48.18 7.53 4.74 3.33 2.58 2.12 1.71 1.45 1.20 1.00
CA1 70.80 11.27 6.23 4.46 3.33 2.49 2.11 1.62 1.34 1.00
Amy−Exc 51.95 8.38 5.14 3.67 2.77 2.15 1.82 1.42 1.24 1.00
Vip 40.87 5.52 3.10 2.31 1.86 1.54 1.35 1.21 1.09 1.00
THM−MB 34.84 4.46 2.92 2.22 1.91 1.67 1.51 1.27 1.15 1.00
THM−Inh 32.57 4.43 2.94 2.19 1.89 1.60 1.37 1.22 1.17 1.00
THM−Exc 39.51 5.88 3.61 2.69 2.18 1.80 1.67 1.44 1.12 1.00
SubCtx−Cplx 45.58 6.59 3.97 2.80 2.22 1.80 1.54 1.26 1.13 1.00
Sst 58.85 7.17 3.81 2.44 1.94 1.61 1.36 1.18 1.05 1.00
Sncg 41.22 5.33 3.30 2.43 1.89 1.57 1.39 1.18 1.11 1.00
Pvalb−ChC 48.53 6.14 3.65 2.46 2.07 1.70 1.48 1.40 1.18 1.00
Pvalb 52.92 6.23 3.44 2.28 1.85 1.59 1.33 1.18 1.04 1.00
PN 39.31 5.50 3.43 2.54 2.02 1.57 1.50 1.28 1.16 1.00
MSN−D2 36.40 5.35 3.39 2.49 1.98 1.71 1.44 1.25 1.08 1.00
MSN−D1 35.15 5.13 3.37 2.51 2.02 1.70 1.48 1.29 1.09 1.00
Lamp5−Lhx6 51.79 6.69 3.58 2.52 1.88 1.65 1.42 1.26 1.10 1.00
Lamp5 42.64 5.59 3.27 2.32 1.85 1.54 1.22 1.09 1.06 1.00
Foxp2 37.98 5.37 3.19 2.31 1.84 1.50 1.39 1.21 1.08 1.00
Chd7 43.23 5.78 3.22 2.42 1.90 1.52 1.34 1.16 1.08 1.00
CB 31.33 4.93 3.14 2.47 1.99 1.77 1.60 1.38 1.19 1.00
VLMC 31.26 4.62 2.92 2.48 2.06 1.78 1.58 1.40 1.21 1.00
PC 30.54 4.52 3.09 2.41 2.06 1.78 1.56 1.37 1.17 1.00
OPC 38.64 5.62 3.30 2.32 2.00 1.64 1.30 1.26 1.13 1.00
ODC 39.61 6.01 3.65 2.59 2.09 1.78 1.47 1.31 1.04 1.00
MGC 19.80 2.73 2.08 1.81 1.65 1.55 1.33 1.27 1.16 1.00
EC 21.95 3.09 2.32 1.93 1.71 1.60 1.44 1.31 1.10 1.00
ASC 35.07 4.90 2.95 2.27 1.97 1.64 1.48 1.35 1.17 1.00
0.0−0.1 0.1−0.2 0.2−0.3 0.3−0.4 0.4−0.5 0.5−0.6 0.6−0.7 0.7−0.8 0.8−0.9 0.9−1.0
Percent 0.01 0.02
SUB−Granule neurons from subicular cortex
L6b 146.83 10.83 7.00 4.42 3.13 2.11 1.49 1.90 0.94 1.00
L6−IT−Car3 187.72 12.80 6.45 5.04 3.33 3.18 2.22 1.92 1.45 1.00
L6−IT 260.05 20.61 11.09 6.66 4.09 2.70 2.63 1.76 0.70 1.00
L6−CT 437.71 33.82 16.93 10.81 9.59 4.92 3.15 3.29 2.66 1.00
L56−NP 120.22 7.85 5.29 3.94 3.27 1.93 1.60 1.22 1.53 1.00
L5−IT 168.57 12.39 7.04 4.26 2.82 2.16 1.97 1.56 1.07 1.00
L5−ET 236.86 15.59 10.27 5.41 4.23 2.77 2.27 1.91 1.41 1.00
L4−IT 280.60 22.16 11.29 8.12 5.57 4.05 4.12 2.50 3.10 1.00
L23−IT 205.45 15.03 8.48 5.46 3.26 2.95 2.28 1.93 1.46 1.00
HIP−Misc2 204.41 14.94 7.44 4.64 4.56 2.29 2.82 2.00 0.90 1.00
HIP−Misc1 258.80 15.90 11.30 6.50 4.40 3.50 3.60 2.30 2.60 1.00
DG 518.00 45.00 20.40 14.40 8.20 5.00 3.00 2.00 2.80 1.00
CA3 322.75 24.63 15.38 9.75 5.63 2.88 2.75 1.63 1.00 1.00
CA1 390.47 33.17 17.59 9.50 5.95 4.06 2.81 1.64 0.88 1.00
Amy−Exc 504.02 38.19 24.78 14.54 10.35 6.53 5.03 3.90 3.26 1.00
Vip 121.92 7.97 5.53 3.51 3.26 2.42 1.80 1.60 1.76 1.00
THM−MB 47.39 2.81 2.25 1.80 1.48 1.32 1.17 1.16 1.06 1.00
THM−Inh 56.35 3.35 2.77 2.12 1.56 1.40 1.33 1.28 0.93 1.00
THM−Exc 162.87 11.40 8.00 5.27 5.27 3.80 3.87 3.07 2.07 1.00
SubCtx−Cplx 102.38 7.02 4.69 3.27 2.77 2.19 1.85 1.38 1.18 1.00
Sst 90.11 5.76 3.80 3.12 2.75 2.11 1.85 1.51 1.17 1.00
Sncg 96.15 6.19 3.56 2.83 2.65 2.38 2.05 1.61 1.42 1.00
Pvalb−ChC 99.03 5.87 3.86 3.70 2.79 2.64 1.97 1.75 1.48 1.00
Pvalb 104.17 5.90 4.55 3.59 3.24 2.57 2.24 2.07 1.49 1.00
PN 101.62 7.60 4.62 2.94 1.66 1.34 1.10 0.94 1.14 1.00
MSN−D2 110.75 8.43 5.29 3.79 3.06 2.26 1.85 1.53 1.22 1.00
MSN−D1 100.04 6.94 4.06 2.78 2.69 2.37 2.12 1.58 1.45 1.00
Lamp5−Lhx6 113.84 6.99 4.43 3.87 2.89 2.31 2.16 2.10 1.56 1.00
Lamp5 104.07 6.98 4.59 3.31 2.46 2.10 1.64 1.45 1.65 1.00
Foxp2 136.77 9.29 5.49 4.01 3.15 3.20 2.76 1.59 1.28 1.00
Chd7 109.98 7.49 5.09 3.68 2.78 2.25 1.81 1.63 1.16 1.00
CB 139.41 9.37 6.84 4.36 3.65 2.99 1.80 2.40 1.49 1.00
VLMC 85.71 5.61 4.14 3.21 2.96 2.54 2.18 1.82 1.50 1.00
PC 79.63 5.03 3.60 3.27 2.43 2.40 2.30 2.03 1.60 1.00
OPC 123.55 8.10 5.30 4.20 3.65 3.00 2.70 1.75 1.70 1.00
ODC 127.58 7.84 5.61 4.37 3.83 3.66 2.74 2.00 1.76 1.00
MGC 49.60 3.53 2.28 1.96 1.94 1.57 1.60 1.51 0.96 1.00
EC 56.26 3.07 2.67 2.31 1.81 1.98 1.88 1.48 1.33 1.00
ASC 83.67 5.77 3.76 2.51 2.23 2.05 1.80 1.43 1.28 1.00
0.0−0.1 0.1−0.2 0.2−0.3 0.3−0.4 0.4−0.5 0.5−0.6 0.6−0.7 0.7−0.8 0.8−0.9 0.9−1.0
Percent 0.001 0.002

### THMGA−GABAergic neurons from thalamus

| | 0.0−0.1 | 0.1−0.2 | 0.2−0.3 | 0.3−0.4 | 0.4−0.5 | 0.5−0.6 | 0.6−0.7 | 0.7−0.8 | 0.8−0.9 | 0.9−1.0 |
|---|---|---|---|---|---|---|---|---|---|---|
| L6b | 39.69 | 5.37 | 3.47 | 2.94 | 2.56 | 2.19 | 1.84 | 1.56 | 1.42 | 1.00 |
| L6−IT−Car3 | 39.10 | 5.19 | 3.59 | 2.76 | 2.44 | 2.14 | 1.85 | 1.60 | 1.41 | 1.00 |
| L6−IT | 37.99 | 5.24 | 3.66 | 2.97 | 2.47 | 2.15 | 1.78 | 1.53 | 1.29 | 1.00 |
| L6−CT | 42.01 | 5.44 | 3.79 | 2.98 | 2.52 | 2.31 | 1.93 | 1.65 | 1.46 | 1.00 |
| L56−NP | 32.21 | 3.70 | 2.67 | 2.31 | 2.01 | 1.71 | 1.65 | 1.48 | 1.29 | 1.00 |
| L5−IT | 37.74 | 4.85 | 3.47 | 2.63 | 2.30 | 1.93 | 1.69 | 1.43 | 1.27 | 1.00 |
| L5−ET | 43.63 | 5.43 | 4.02 | 3.05 | 2.64 | 2.11 | 1.90 | 1.76 | 1.43 | 1.00 |
| L4−IT | 32.02 | 4.23 | 2.88 | 2.37 | 2.00 | 1.87 | 1.64 | 1.35 | 1.29 | 1.00 |
| L23−IT | 37.83 | 5.18 | 3.51 | 2.79 | 2.36 | 2.12 | 1.83 | 1.55 | 1.42 | 1.00 |
| HIP−Misc2 | 34.76 | 4.35 | 3.02 | 2.60 | 2.23 | 1.93 | 1.72 | 1.57 | 1.40 | 1.00 |
| HIP−Misc1 | 31.16 | 4.12 | 2.69 | 2.33 | 2.05 | 1.86 | 1.45 | 1.33 | 1.22 | 1.00 |
| DG | 34.68 | 4.98 | 3.40 | 2.96 | 2.52 | 2.07 | 1.95 | 1.47 | 1.35 | 1.00 |
| CA3 | 39.27 | 5.48 | 4.02 | 3.06 | 2.66 | 2.14 | 1.95 | 1.62 | 1.51 | 1.00 |
| CA1 | 41.74 | 5.89 | 3.94 | 3.24 | 2.72 | 2.26 | 1.99 | 1.73 | 1.37 | 1.00 |
| Amy−Exc | 37.11 | 4.98 | 3.63 | 2.98 | 2.52 | 2.08 | 1.87 | 1.61 | 1.36 | 1.00 |
| Vip | 29.56 | 3.70 | 2.56 | 1.98 | 1.70 | 1.57 | 1.40 | 1.30 | 1.25 | 1.00 |
| THM−MB | 122.31 | 12.17 | 5.23 | 2.98 | 2.12 | 1.58 | 1.30 | 1.08 | 0.95 | 1.00 |
| THM−Inh | 32.88 | 4.31 | 2.78 | 2.25 | 2.03 | 1.64 | 1.48 | 1.36 | 1.15 | 1.00 |
| THM−Exc | 55.19 | 7.60 | 5.24 | 3.81 | 3.24 | 2.70 | 2.01 | 1.77 | 1.47 | 1.00 |
| SubCtx−Cplx | 45.37 | 6.02 | 4.02 | 3.07 | 2.47 | 2.05 | 1.74 | 1.58 | 1.31 | 1.00 |
| Sst | 31.19 | 3.69 | 2.62 | 2.13 | 1.77 | 1.60 | 1.44 | 1.37 | 1.15 | 1.00 |
| Sncg | 27.96 | 3.36 | 2.38 | 1.85 | 1.66 | 1.54 | 1.42 | 1.28 | 1.18 | 1.00 |
| Pvalb−ChC | 35.28 | 4.24 | 2.82 | 2.37 | 2.03 | 1.90 | 1.48 | 1.36 | 1.15 | 1.00 |
| Pvalb | 31.96 | 3.72 | 2.64 | 2.18 | 1.81 | 1.61 | 1.42 | 1.34 | 1.16 | 1.00 |
| PN | 37.17 | 4.82 | 3.35 | 2.69 | 2.12 | 1.92 | 1.67 | 1.38 | 1.34 | 1.00 |
| MSN−D2 | 37.28 | 4.96 | 3.51 | 2.84 | 2.28 | 1.96 | 1.70 | 1.47 | 1.33 | 1.00 |
| MSN−D1 | 34.16 | 4.56 | 3.29 | 2.52 | 2.14 | 1.83 | 1.66 | 1.43 | 1.29 | 1.00 |
| Lamp5−Lhx6 | 32.18 | 4.03 | 2.79 | 2.10 | 1.73 | 1.62 | 1.44 | 1.24 | 1.25 | 1.00 |
| Lamp5 | 31.46 | 3.98 | 2.65 | 2.15 | 1.79 | 1.60 | 1.42 | 1.30 | 1.16 | 1.00 |
| Foxp2 | 37.87 | 4.94 | 3.35 | 2.64 | 2.20 | 1.86 | 1.52 | 1.39 | 1.32 | 1.00 |
| Chd7 | 33.46 | 4.06 | 2.87 | 2.36 | 1.90 | 1.70 | 1.50 | 1.36 | 1.22 | 1.00 |
| CB | 36.82 | 4.90 | 3.66 | 3.04 | 2.44 | 2.11 | 1.89 | 1.75 | 1.44 | 1.00 |
| VLMC | 27.95 | 3.51 | 2.65 | 2.15 | 1.89 | 1.82 | 1.63 | 1.54 | 1.20 | 1.00 |
| PC | 26.26 | 3.31 | 2.50 | 2.08 | 1.90 | 1.64 | 1.54 | 1.40 | 1.17 | 1.00 |
| OPC | 38.33 | 4.79 | 3.54 | 2.61 | 2.36 | 2.14 | 1.72 | 1.56 | 1.32 | 1.00 |
| ODC | 30.54 | 3.85 | 2.85 | 2.29 | 1.93 | 1.71 | 1.43 | 1.35 | 1.16 | 1.00 |
| MGC | 20.37 | 2.26 | 1.88 | 1.82 | 1.76 | 1.55 | 1.47 | 1.27 | 1.20 | 1.00 |
| EC | 22.51 | 2.61 | 2.19 | 1.87 | 1.80 | 1.66 | 1.55 | 1.43 | 1.28 | 1.00 |
| ASC | 35.04 | 4.56 | 3.04 | 2.47 | 2.10 | 1.82 | 1.61 | 1.41 | 1.23 | 1.00 |

Percent 0.004 0.008 0.012

### VIP−VIP+ GABAergic neurons

| | 0.0−0.1 | 0.1−0.2 | 0.2−0.3 | 0.3−0.4 | 0.4−0.5 | 0.5−0.6 | 0.6−0.7 | 0.7−0.8 | 0.8−0.9 | 0.9−1.0 |
|---|---|---|---|---|---|---|---|---|---|---|
| L6b | 23.23 | 6.65 | 4.37 | 3.28 | 2.59 | 2.11 | 1.76 | 1.46 | 1.26 | 1.00 |
| L6−IT−Car3 | 22.62 | 6.16 | 4.23 | 3.18 | 2.57 | 2.12 | 1.81 | 1.57 | 1.31 | 1.00 |
| L6−IT | 25.75 | 7.32 | 4.91 | 3.64 | 2.85 | 2.35 | 1.91 | 1.56 | 1.28 | 1.00 |
| L6−CT | 25.62 | 7.17 | 4.65 | 3.47 | 2.71 | 2.25 | 1.85 | 1.55 | 1.26 | 1.00 |
| L56−NP | 17.70 | 4.68 | 3.42 | 2.68 | 2.19 | 1.91 | 1.67 | 1.49 | 1.26 | 1.00 |
| L5−IT | 23.64 | 6.55 | 4.32 | 3.12 | 2.49 | 2.05 | 1.74 | 1.46 | 1.26 | 1.00 |
| L5−ET | 27.91 | 7.54 | 4.90 | 3.65 | 2.87 | 2.34 | 1.90 | 1.60 | 1.30 | 1.00 |
| L4−IT | 20.31 | 5.67 | 3.88 | 2.96 | 2.40 | 2.04 | 1.74 | 1.51 | 1.27 | 1.00 |
| L23−IT | 23.29 | 6.71 | 4.49 | 3.30 | 2.63 | 2.19 | 1.83 | 1.52 | 1.27 | 1.00 |
| HIP−Misc2 | 21.74 | 6.05 | 3.89 | 2.92 | 2.34 | 1.90 | 1.59 | 1.38 | 1.22 | 1.00 |
| HIP−Misc1 | 23.33 | 6.62 | 4.42 | 3.29 | 2.59 | 2.13 | 1.80 | 1.51 | 1.28 | 1.00 |
| DG | 34.58 | 10.58 | 6.73 | 4.60 | 3.53 | 2.66 | 2.16 | 1.69 | 1.31 | 1.00 |
| CA3 | 31.43 | 9.26 | 5.86 | 4.28 | 3.18 | 2.47 | 1.92 | 1.63 | 1.29 | 1.00 |
| CA1 | 33.17 | 9.78 | 6.33 | 4.54 | 3.47 | 2.64 | 2.10 | 1.75 | 1.43 | 1.00 |
| Amy−Exc | 24.89 | 7.57 | 5.05 | 3.72 | 2.91 | 2.30 | 1.89 | 1.56 | 1.29 | 1.00 |
| Vip | 35.43 | 8.29 | 4.62 | 3.09 | 2.27 | 1.77 | 1.47 | 1.28 | 1.09 | 1.00 |
| THM−MB | 15.47 | 3.93 | 2.84 | 2.30 | 2.01 | 1.79 | 1.59 | 1.41 | 1.21 | 1.00 |
| THM−Inh | 18.91 | 5.01 | 3.39 | 2.62 | 2.08 | 1.88 | 1.57 | 1.35 | 1.16 | 1.00 |
| THM−Exc | 21.82 | 6.35 | 4.22 | 3.23 | 2.51 | 2.08 | 1.72 | 1.43 | 1.21 | 1.00 |
| SubCtx−Cplx | 23.64 | 6.62 | 4.29 | 3.14 | 2.41 | 1.94 | 1.60 | 1.32 | 1.15 | 1.00 |
| Sst | 22.52 | 5.79 | 3.78 | 2.80 | 2.27 | 1.92 | 1.63 | 1.41 | 1.19 | 1.00 |
| Sncg | 30.75 | 7.60 | 4.54 | 3.16 | 2.46 | 2.00 | 1.67 | 1.41 | 1.21 | 1.00 |
| Pvalb−ChC | 25.55 | 6.38 | 4.06 | 2.97 | 2.30 | 1.93 | 1.65 | 1.37 | 1.17 | 1.00 |
| Pvalb | 24.03 | 5.94 | 3.73 | 2.66 | 2.17 | 1.81 | 1.56 | 1.38 | 1.19 | 1.00 |
| PN | 29.29 | 7.80 | 4.76 | 3.39 | 2.52 | 2.05 | 1.72 | 1.52 | 1.24 | 1.00 |
| MSN−D2 | 15.43 | 4.30 | 3.10 | 2.47 | 2.05 | 1.75 | 1.52 | 1.35 | 1.22 | 1.00 |
| MSN−D1 | 14.59 | 4.08 | 2.88 | 2.33 | 1.97 | 1.72 | 1.51 | 1.36 | 1.21 | 1.00 |
| Lamp5−Lhx6 | 28.86 | 7.23 | 4.23 | 2.98 | 2.31 | 1.85 | 1.55 | 1.31 | 1.08 | 1.00 |
| Lamp5 | 30.59 | 7.57 | 4.49 | 3.17 | 2.42 | 1.89 | 1.60 | 1.37 | 1.19 | 1.00 |
| Foxp2 | 18.00 | 4.89 | 3.29 | 2.50 | 2.05 | 1.73 | 1.54 | 1.33 | 1.20 | 1.00 |
| Chd7 | 30.05 | 7.56 | 4.42 | 3.10 | 2.34 | 1.84 | 1.54 | 1.30 | 1.17 | 1.00 |
| CB | 26.12 | 7.30 | 4.71 | 3.40 | 2.65 | 2.20 | 1.90 | 1.57 | 1.34 | 1.00 |
| VLMC | 18.23 | 4.99 | 3.44 | 2.63 | 2.30 | 1.97 | 1.65 | 1.47 | 1.23 | 1.00 |
| PC | 17.09 | 4.70 | 3.44 | 2.75 | 2.22 | 2.03 | 1.74 | 1.50 | 1.28 | 1.00 |
| OPC | 28.69 | 7.14 | 4.24 | 3.07 | 2.37 | 1.93 | 1.63 | 1.40 | 1.21 | 1.00 |
| ODC | 27.75 | 7.34 | 4.54 | 3.24 | 2.47 | 2.00 | 1.78 | 1.44 | 1.21 | 1.00 |
| MGC | 11.01 | 3.07 | 2.36 | 2.07 | 1.89 | 1.70 | 1.58 | 1.35 | 1.21 | 1.00 |
| EC | 12.62 | 3.44 | 2.63 | 2.27 | 1.98 | 1.75 | 1.59 | 1.40 | 1.23 | 1.00 |
| ASC | 23.62 | 6.01 | 3.75 | 2.77 | 2.23 | 1.91 | 1.66 | 1.43 | 1.18 | 1.00 |

Percent 0.02 0.04 0.06
