## Supplementary Figure 4 for "Distinct cellular DNA methylation mechanisms underlie common and rare genetic risk for brain disorders"

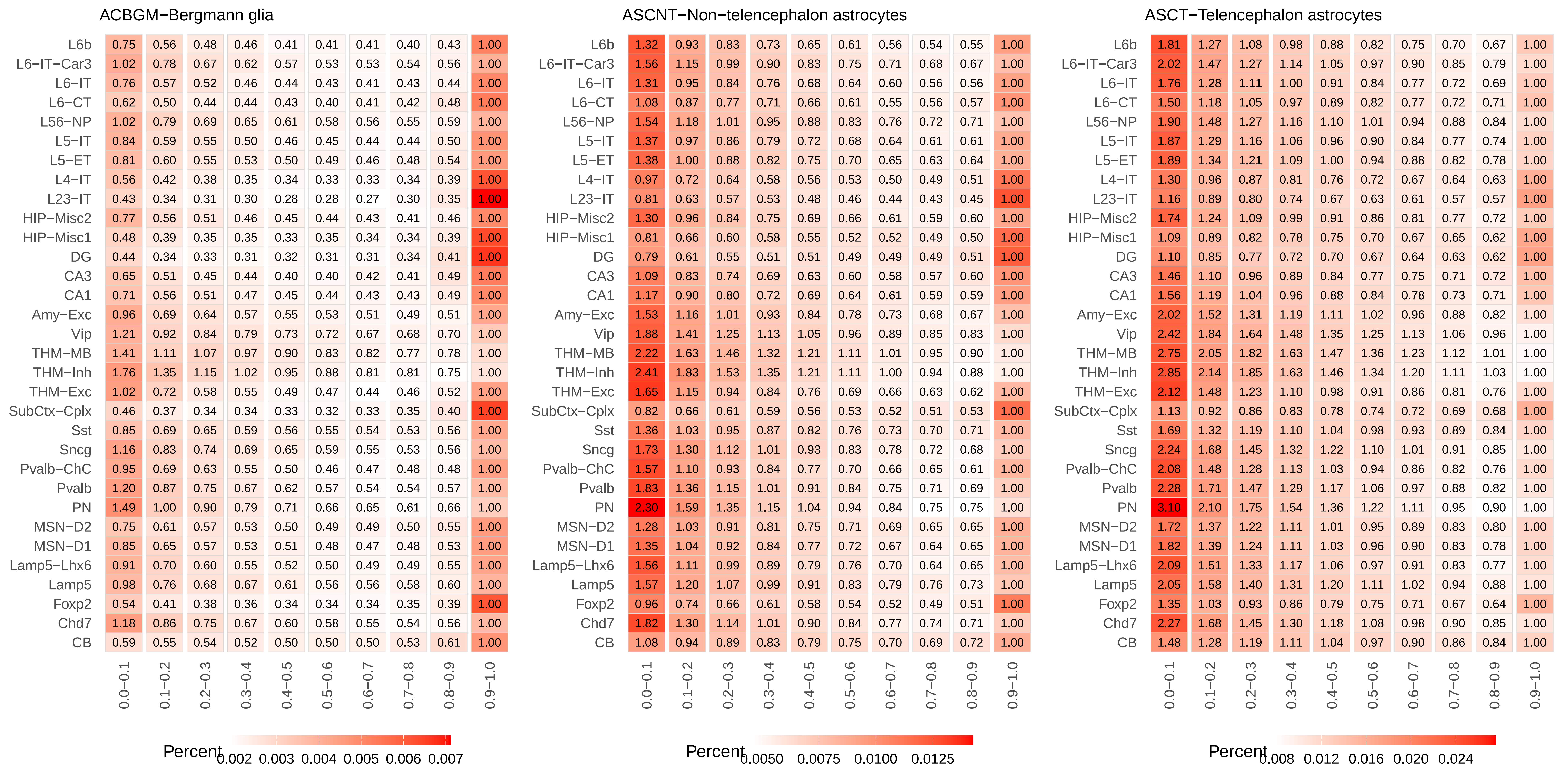
ACBGM−Bergmann glia
L6b 0.75 0.56 0.48 0.46 0.41 0.41 0.41 0.40 0.43 1.00
L6−IT−Car3 1.02 0.78 0.67 0.62 0.57 0.53 0.53 0.54 0.56 1.00
L6−IT 0.76 0.57 0.52 0.46 0.44 0.43 0.41 0.43 0.44 1.00
L6−CT 0.62 0.50 0.44 0.44 0.43 0.40 0.41 0.42 0.48 1.00
L56−NP 1.02 0.79 0.69 0.65 0.61 0.58 0.56 0.55 0.59 1.00
L5−IT 0.84 0.59 0.55 0.50 0.46 0.45 0.44 0.44 0.50 1.00
L5−ET 0.81 0.60 0.55 0.53 0.50 0.49 0.46 0.48 0.54 1.00
L4−IT 0.56 0.42 0.38 0.35 0.34 0.33 0.33 0.34 0.39 1.00
L23−IT 0.43 0.34 0.31 0.30 0.28 0.28 0.27 0.30 0.35 1.00
HIP−Misc2 0.77 0.56 0.51 0.46 0.45 0.44 0.43 0.41 0.46 1.00
HIP−Misc1 0.48 0.39 0.35 0.35 0.33 0.35 0.34 0.34 0.39 1.00
DG 0.44 0.34 0.33 0.31 0.32 0.31 0.31 0.34 0.41 1.00
CA3 0.65 0.51 0.45 0.44 0.40 0.40 0.42 0.41 0.49 1.00
CA1 0.71 0.56 0.51 0.47 0.45 0.44 0.43 0.43 0.49 1.00
Amy−Exc 0.96 0.69 0.64 0.57 0.55 0.53 0.51 0.49 0.51 1.00
Vip 1.21 0.92 0.84 0.79 0.73 0.72 0.67 0.68 0.70 1.00
THM−MB 1.41 1.11 1.07 0.97 0.90 0.83 0.82 0.77 0.78 1.00
THM−Inh 1.76 1.35 1.15 1.02 0.95 0.88 0.81 0.81 0.75 1.00
THM−Exc 1.02 0.72 0.58 0.55 0.49 0.47 0.44 0.46 0.52 1.00
SubCtx−Cplx 0.46 0.37 0.34 0.34 0.33 0.32 0.33 0.35 0.40 1.00
Sst 0.85 0.69 0.65 0.59 0.56 0.55 0.54 0.53 0.56 1.00
Sncg 1.16 0.83 0.74 0.69 0.65 0.59 0.55 0.53 0.56 1.00
Pvalb−ChC 0.95 0.69 0.63 0.55 0.50 0.46 0.47 0.48 0.48 1.00
Pvalb 1.20 0.87 0.75 0.67 0.62 0.57 0.54 0.54 0.57 1.00
PN 1.49 1.00 0.90 0.79 0.71 0.66 0.65 0.61 0.66 1.00
MSN−D2 0.75 0.61 0.57 0.53 0.50 0.49 0.49 0.50 0.55 1.00
MSN−D1 0.85 0.65 0.57 0.53 0.51 0.48 0.47 0.48 0.53 1.00
Lamp5−Lhx6 0.91 0.70 0.60 0.55 0.52 0.50 0.49 0.49 0.55 1.00
Lamp5 0.98 0.76 0.68 0.67 0.61 0.56 0.56 0.58 0.60 1.00
Foxp2 0.54 0.41 0.38 0.36 0.34 0.34 0.34 0.35 0.39 1.00
Chd7 1.18 0.86 0.75 0.67 0.60 0.58 0.55 0.54 0.56 1.00
CB 0.59 0.55 0.54 0.52 0.50 0.50 0.50 0.53 0.61 1.00
0.0−0.1 0.1−0.2 0.2−0.3 0.3−0.4 0.4−0.5 0.5−0.6 0.6−0.7 0.7−0.8 0.8−0.9 0.9−1.0
Percent
0.002 0.003 0.004 0.005 0.006 0.007
ASCNT−Non−telencephalon astrocytes
L6b 1.32 0.93 0.83 0.73 0.65 0.61 0.56 0.54 0.55 1.00
L6−IT−Car3 1.56 1.15 0.99 0.90 0.83 0.75 0.71 0.68 0.67 1.00
L6−IT 1.31 0.95 0.84 0.76 0.68 0.64 0.60 0.56 0.56 1.00
L6−CT 1.08 0.87 0.77 0.71 0.66 0.61 0.55 0.56 0.57 1.00
L56−NP 1.54 1.18 1.01 0.95 0.88 0.83 0.76 0.72 0.71 1.00
L5−IT 1.37 0.97 0.86 0.79 0.72 0.68 0.64 0.61 0.61 1.00
L5−ET 1.38 1.00 0.88 0.82 0.75 0.70 0.65 0.63 0.64 1.00
L4−IT 0.97 0.72 0.64 0.58 0.56 0.53 0.50 0.49 0.51 1.00
L23−IT 0.81 0.63 0.57 0.53 0.48 0.46 0.44 0.43 0.45 1.00
HIP−Misc2 1.30 0.96 0.84 0.75 0.69 0.66 0.61 0.59 0.60 1.00
HIP−Misc1 0.81 0.66 0.60 0.58 0.55 0.52 0.52 0.49 0.50 1.00
DG 0.79 0.61 0.55 0.51 0.51 0.49 0.49 0.49 0.51 1.00
CA3 1.09 0.83 0.74 0.69 0.63 0.60 0.58 0.57 0.60 1.00
CA1 1.17 0.90 0.80 0.72 0.69 0.64 0.61 0.59 0.59 1.00
Amy−Exc 1.53 1.16 1.01 0.93 0.84 0.78 0.73 0.68 0.67 1.00
Vip 1.88 1.41 1.25 1.13 1.05 0.96 0.89 0.85 0.83 1.00
THM−MB 2.22 1.63 1.46 1.32 1.21 1.11 1.01 0.95 0.90 1.00
THM−Inh 2.41 1.83 1.53 1.35 1.21 1.11 1.00 0.94 0.88 1.00
THM−Exc 1.65 1.15 0.94 0.84 0.76 0.69 0.66 0.63 0.62 1.00
SubCtx−Cplx 0.82 0.66 0.61 0.59 0.56 0.53 0.52 0.51 0.53 1.00
Sst 1.36 1.03 0.95 0.87 0.82 0.76 0.73 0.70 0.71 1.00
Sncg 1.73 1.30 1.12 1.01 0.93 0.83 0.78 0.72 0.68 1.00
Pvalb−ChC 1.57 1.10 0.93 0.84 0.77 0.70 0.66 0.65 0.61 1.00
Pvalb 1.83 1.36 1.15 1.01 0.91 0.84 0.75 0.71 0.69 1.00
PN 2.30 1.59 1.35 1.15 1.04 0.94 0.84 0.75 0.75 1.00
MSN−D2 1.28 1.03 0.91 0.81 0.75 0.71 0.69 0.65 0.65 1.00
MSN−D1 1.35 1.04 0.92 0.84 0.77 0.72 0.67 0.64 0.65 1.00
Lamp5−Lhx6 1.56 1.11 0.99 0.89 0.79 0.76 0.70 0.64 0.65 1.00
Lamp5 1.57 1.20 1.07 0.99 0.91 0.83 0.79 0.76 0.73 1.00
Foxp2 0.96 0.74 0.66 0.61 0.58 0.54 0.52 0.49 0.51 1.00
Chd7 1.82 1.30 1.14 1.01 0.90 0.84 0.77 0.74 0.71 1.00
CB 1.08 0.94 0.89 0.83 0.79 0.75 0.70 0.69 0.72 1.00
0.0−0.1 0.1−0.2 0.2−0.3 0.3−0.4 0.4−0.5 0.5−0.6 0.6−0.7 0.7−0.8 0.8−0.9 0.9−1.0
Percent
0.0050 0.0075 0.0100 0.0125
ASCT−Telencephalon astrocytes
L6b 1.81 1.27 1.08 0.98 0.88 0.82 0.75 0.70 0.67 1.00
L6−IT−Car3 2.02 1.47 1.27 1.14 1.05 0.97 0.90 0.85 0.79 1.00
L6−IT 1.76 1.28 1.11 1.00 0.91 0.84 0.77 0.72 0.69 1.00
L6−CT 1.50 1.18 1.05 0.97 0.89 0.82 0.77 0.72 0.71 1.00
L56−NP 1.90 1.48 1.27 1.16 1.10 1.01 0.94 0.88 0.84 1.00
L5−IT 1.87 1.29 1.16 1.06 0.96 0.90 0.84 0.77 0.74 1.00
L5−ET 1.89 1.34 1.21 1.09 1.00 0.94 0.88 0.82 0.78 1.00
L4−IT 1.30 0.96 0.87 0.81 0.76 0.72 0.67 0.64 0.63 1.00
L23−IT 1.16 0.89 0.80 0.74 0.67 0.63 0.61 0.57 0.57 1.00
HIP−Misc2 1.74 1.24 1.09 0.99 0.91 0.86 0.81 0.77 0.72 1.00
HIP−Misc1 1.09 0.89 0.82 0.78 0.75 0.70 0.67 0.65 0.62 1.00
DG 1.10 0.85 0.77 0.72 0.70 0.67 0.64 0.63 0.62 1.00
CA3 1.46 1.10 0.96 0.89 0.84 0.77 0.75 0.71 0.72 1.00
CA1 1.56 1.19 1.04 0.96 0.88 0.84 0.78 0.73 0.71 1.00
Amy−Exc 2.02 1.52 1.31 1.19 1.11 1.02 0.96 0.88 0.82 1.00
Vip 2.42 1.84 1.64 1.48 1.35 1.25 1.13 1.06 0.96 1.00
THM−MB 2.75 2.05 1.82 1.63 1.47 1.36 1.23 1.12 1.01 1.00
THM−Inh 2.85 2.14 1.85 1.63 1.46 1.34 1.20 1.11 1.03 1.00
THM−Exc 2.12 1.48 1.23 1.10 0.98 0.91 0.86 0.81 0.76 1.00
SubCtx−Cplx 1.13 0.92 0.86 0.83 0.78 0.74 0.72 0.69 0.68 1.00
Sst 1.69 1.32 1.19 1.10 1.04 0.98 0.93 0.89 0.84 1.00
Sncg 2.24 1.68 1.45 1.32 1.22 1.10 1.01 0.91 0.85 1.00
Pvalb−ChC 2.08 1.48 1.28 1.13 1.03 0.94 0.86 0.82 0.76 1.00
Pvalb 2.28 1.71 1.47 1.29 1.17 1.06 0.97 0.88 0.82 1.00
PN 3.10 2.10 1.75 1.54 1.36 1.22 1.11 0.95 0.90 1.00
MSN−D2 1.72 1.37 1.22 1.11 1.01 0.95 0.89 0.83 0.80 1.00
MSN−D1 1.82 1.39 1.24 1.11 1.03 0.96 0.90 0.83 0.78 1.00
Lamp5−Lhx6 2.09 1.51 1.33 1.17 1.06 0.97 0.91 0.83 0.77 1.00
Lamp5 2.05 1.58 1.40 1.31 1.20 1.11 1.02 0.94 0.88 1.00
Foxp2 1.35 1.03 0.93 0.86 0.79 0.75 0.71 0.67 0.64 1.00
Chd7 2.27 1.68 1.45 1.30 1.18 1.08 0.98 0.90 0.85 1.00
CB 1.48 1.28 1.19 1.11 1.04 0.97 0.90 0.86 0.84 1.00
0.0−0.1 0.1−0.2 0.2−0.3 0.3−0.4 0.4−0.5 0.5−0.6 0.6−0.7 0.7−0.8 0.8−0.9 0.9−1.0
Percent
0.008 0.012 0.016 0.020 0.024

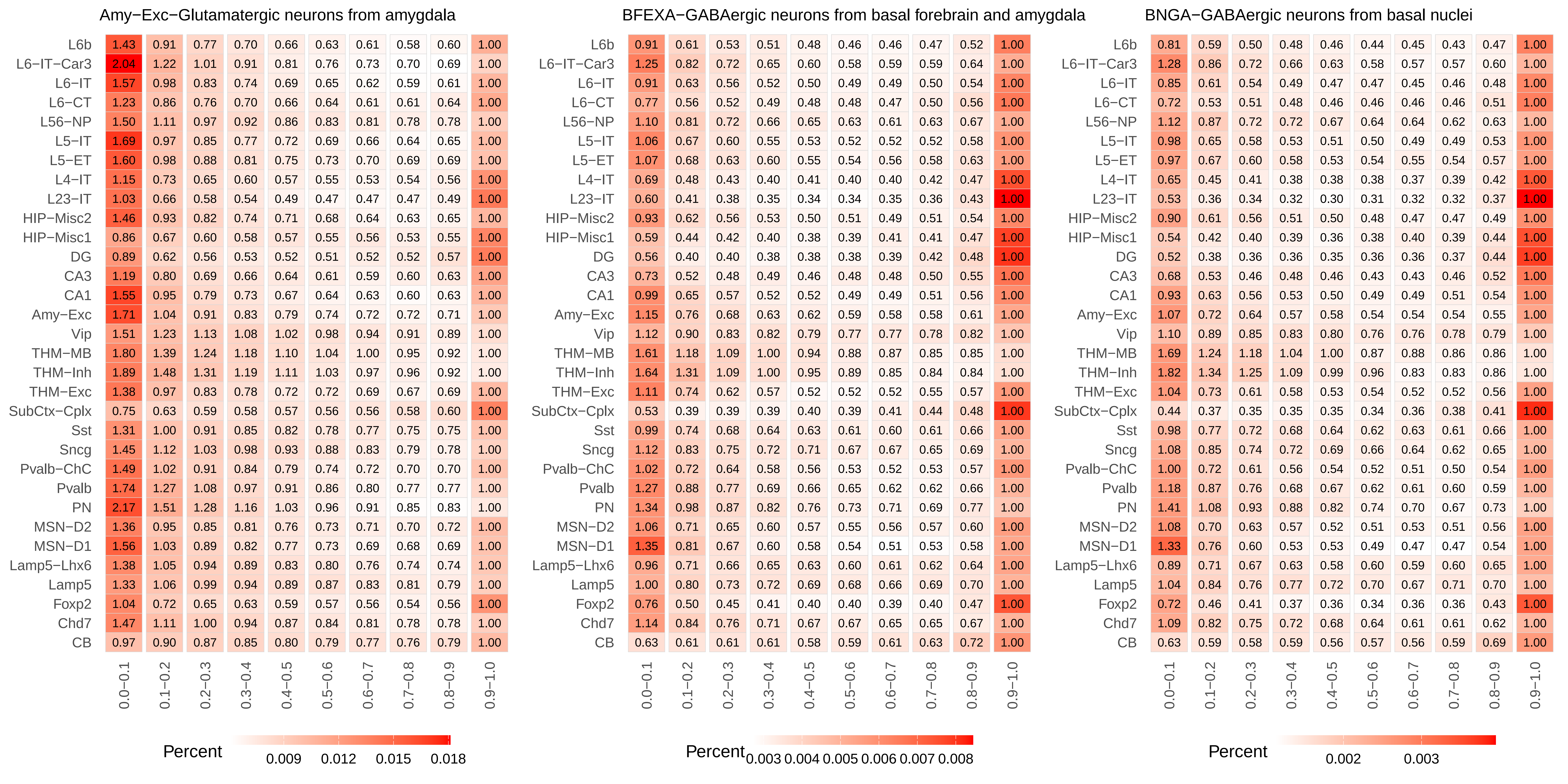
Amy−Exc−Glutamatergic neurons from amygdala
L6b 1.43 0.91 0.77 0.70 0.66 0.63 0.61 0.58 0.60 1.00
L6−IT−Car3 2.04 1.22 1.01 0.91 0.81 0.76 0.73 0.70 0.69 1.00
L6−IT 1.57 0.98 0.83 0.74 0.69 0.65 0.62 0.59 0.61 1.00
L6−CT 1.23 0.86 0.76 0.70 0.66 0.64 0.61 0.61 0.64 1.00
L56−NP 1.50 1.11 0.97 0.92 0.86 0.83 0.81 0.78 0.78 1.00
L5−IT 1.69 0.97 0.85 0.77 0.72 0.69 0.66 0.64 0.65 1.00
L5−ET 1.60 0.98 0.88 0.81 0.75 0.73 0.70 0.69 0.69 1.00
L4−IT 1.15 0.73 0.65 0.60 0.57 0.55 0.53 0.54 0.56 1.00
L23−IT 1.03 0.66 0.58 0.54 0.49 0.47 0.47 0.47 0.49 1.00
HIP−Misc2 1.46 0.93 0.82 0.74 0.71 0.68 0.64 0.63 0.65 1.00
HIP−Misc1 0.86 0.67 0.60 0.58 0.57 0.55 0.56 0.53 0.55 1.00
DG 0.89 0.62 0.56 0.53 0.52 0.51 0.52 0.52 0.57 1.00
CA3 1.19 0.80 0.69 0.66 0.64 0.61 0.59 0.60 0.63 1.00
CA1 1.55 0.95 0.79 0.73 0.67 0.64 0.63 0.60 0.63 1.00
Amy−Exc 1.71 1.04 0.91 0.83 0.79 0.74 0.72 0.72 0.71 1.00
Vip 1.51 1.23 1.13 1.08 1.02 0.98 0.94 0.91 0.89 1.00
THM−MB 1.80 1.39 1.24 1.18 1.10 1.04 1.00 0.95 0.92 1.00
THM−Inh 1.89 1.48 1.31 1.19 1.11 1.03 0.97 0.96 0.92 1.00
THM−Exc 1.38 0.97 0.83 0.78 0.72 0.72 0.69 0.67 0.69 1.00
SubCtx−Cplx 0.75 0.63 0.59 0.58 0.57 0.56 0.56 0.58 0.60 1.00
Sst 1.31 1.00 0.91 0.85 0.82 0.78 0.77 0.75 0.75 1.00
Sncg 1.45 1.12 1.03 0.98 0.93 0.88 0.83 0.79 0.78 1.00
Pvalb−ChC 1.49 1.02 0.91 0.84 0.79 0.74 0.72 0.70 0.70 1.00
Pvalb 1.74 1.27 1.08 0.97 0.91 0.86 0.80 0.77 0.77 1.00
PN 2.17 1.51 1.28 1.16 1.03 0.96 0.91 0.85 0.83 1.00
MSN−D2 1.36 0.95 0.85 0.81 0.76 0.73 0.71 0.70 0.72 1.00
MSN−D1 1.56 1.03 0.89 0.82 0.77 0.73 0.69 0.68 0.69 1.00
Lamp5−Lhx6 1.38 1.05 0.94 0.89 0.83 0.80 0.76 0.74 0.74 1.00
Lamp5 1.33 1.06 0.99 0.94 0.89 0.87 0.83 0.81 0.79 1.00
Foxp2 1.04 0.72 0.65 0.63 0.59 0.57 0.56 0.54 0.56 1.00
Chd7 1.47 1.11 1.00 0.94 0.87 0.84 0.81 0.78 0.78 1.00
CB 0.97 0.90 0.87 0.85 0.80 0.79 0.77 0.76 0.79 1.00
0.0−0.1 0.1−0.2 0.2−0.3 0.3−0.4 0.4−0.5 0.5−0.6 0.6−0.7 0.7−0.8 0.8−0.9 0.9−1.0
Percent 0.009 0.012 0.015 0.018
BFEXA−GABAergic neurons from basal forebrain and amygdala
L6b 0.91 0.61 0.53 0.51 0.48 0.46 0.46 0.47 0.52 1.00
L6−IT−Car3 1.25 0.82 0.72 0.65 0.60 0.58 0.59 0.59 0.64 1.00
L6−IT 0.91 0.63 0.56 0.52 0.50 0.49 0.49 0.50 0.54 1.00
L6−CT 0.77 0.56 0.52 0.49 0.48 0.48 0.47 0.50 0.56 1.00
L56−NP 1.10 0.81 0.72 0.66 0.65 0.63 0.61 0.63 0.67 1.00
L5−IT 1.06 0.67 0.60 0.55 0.53 0.52 0.52 0.52 0.58 1.00
L5−ET 1.07 0.68 0.63 0.60 0.55 0.54 0.56 0.58 0.63 1.00
L4−IT 0.69 0.48 0.43 0.40 0.41 0.40 0.40 0.42 0.47 1.00
L23−IT 0.60 0.41 0.38 0.35 0.34 0.34 0.35 0.36 0.43 1.00
HIP−Misc2 0.93 0.62 0.56 0.53 0.50 0.51 0.49 0.51 0.54 1.00
HIP−Misc1 0.59 0.44 0.42 0.40 0.38 0.39 0.41 0.41 0.47 1.00
DG 0.56 0.40 0.40 0.38 0.38 0.38 0.39 0.42 0.48 1.00
CA3 0.73 0.52 0.48 0.49 0.46 0.48 0.48 0.50 0.55 1.00
CA1 0.99 0.65 0.57 0.52 0.52 0.49 0.49 0.51 0.56 1.00
Amy−Exc 1.15 0.76 0.68 0.63 0.62 0.59 0.58 0.58 0.61 1.00
Vip 1.12 0.90 0.83 0.82 0.79 0.77 0.77 0.78 0.82 1.00
THM−MB 1.61 1.18 1.09 1.00 0.94 0.88 0.87 0.85 0.85 1.00
THM−Inh 1.64 1.31 1.09 1.00 0.95 0.89 0.85 0.84 0.84 1.00
THM−Exc 1.11 0.74 0.62 0.57 0.52 0.52 0.52 0.55 0.57 1.00
SubCtx−Cplx 0.53 0.39 0.39 0.39 0.40 0.39 0.41 0.44 0.48 1.00
Sst 0.99 0.74 0.68 0.64 0.63 0.61 0.60 0.61 0.66 1.00
Sncg 1.12 0.83 0.75 0.72 0.71 0.67 0.67 0.65 0.69 1.00
Pvalb−ChC 1.02 0.72 0.64 0.58 0.56 0.53 0.52 0.53 0.57 1.00
Pvalb 1.27 0.88 0.77 0.69 0.66 0.65 0.62 0.62 0.66 1.00
PN 1.34 0.98 0.87 0.82 0.76 0.73 0.71 0.69 0.77 1.00
MSN−D2 1.06 0.71 0.65 0.60 0.57 0.55 0.56 0.57 0.60 1.00
MSN−D1 1.35 0.81 0.67 0.60 0.58 0.54 0.51 0.53 0.58 1.00
Lamp5−Lhx6 0.96 0.71 0.66 0.65 0.63 0.60 0.61 0.62 0.64 1.00
Lamp5 1.00 0.80 0.73 0.72 0.69 0.68 0.66 0.69 0.70 1.00
Foxp2 0.76 0.50 0.45 0.41 0.40 0.40 0.39 0.40 0.47 1.00
Chd7 1.14 0.84 0.76 0.71 0.67 0.67 0.65 0.65 0.67 1.00
CB 0.63 0.61 0.61 0.61 0.58 0.59 0.61 0.63 0.72 1.00
0.0−0.1 0.1−0.2 0.2−0.3 0.3−0.4 0.4−0.5 0.5−0.6 0.6−0.7 0.7−0.8 0.8−0.9 0.9−1.0
Percent 0.003 0.004 0.005 0.006 0.007 0.008
BNGA−GABAergic neurons from basal nuclei
L6b 0.81 0.59 0.50 0.48 0.46 0.44 0.45 0.43 0.47 1.00
L6−IT−Car3 1.28 0.86 0.72 0.66 0.63 0.58 0.57 0.57 0.60 1.00
L6−IT 0.85 0.61 0.54 0.49 0.47 0.47 0.45 0.46 0.48 1.00
L6−CT 0.72 0.53 0.51 0.48 0.46 0.46 0.46 0.46 0.51 1.00
L56−NP 1.12 0.87 0.72 0.72 0.67 0.64 0.64 0.62 0.63 1.00
L5−IT 0.98 0.65 0.58 0.53 0.51 0.50 0.49 0.49 0.53 1.00
L5−ET 0.97 0.67 0.60 0.58 0.53 0.54 0.55 0.54 0.57 1.00
L4−IT 0.65 0.45 0.41 0.38 0.38 0.38 0.37 0.39 0.42 1.00
L23−IT 0.53 0.36 0.34 0.32 0.30 0.31 0.32 0.32 0.37 1.00
HIP−Misc2 0.90 0.61 0.56 0.51 0.50 0.48 0.47 0.47 0.49 1.00
HIP−Misc1 0.54 0.42 0.40 0.39 0.36 0.38 0.40 0.39 0.44 1.00
DG 0.52 0.38 0.36 0.36 0.35 0.36 0.36 0.37 0.44 1.00
CA3 0.68 0.53 0.46 0.48 0.46 0.43 0.43 0.46 0.52 1.00
CA1 0.93 0.63 0.56 0.53 0.50 0.49 0.49 0.51 0.54 1.00
Amy−Exc 1.07 0.72 0.64 0.57 0.58 0.54 0.54 0.54 0.55 1.00
Vip 1.10 0.89 0.85 0.83 0.80 0.76 0.76 0.78 0.79 1.00
THM−MB 1.69 1.24 1.18 1.04 1.00 0.87 0.88 0.86 0.86 1.00
THM−Inh 1.82 1.34 1.25 1.09 0.99 0.96 0.83 0.83 0.86 1.00
THM−Exc 1.04 0.73 0.61 0.58 0.53 0.54 0.52 0.52 0.56 1.00
SubCtx−Cplx 0.44 0.37 0.35 0.35 0.35 0.34 0.36 0.38 0.41 1.00
Sst 0.98 0.77 0.72 0.68 0.64 0.62 0.63 0.61 0.66 1.00
Sncg 1.08 0.85 0.74 0.72 0.69 0.66 0.64 0.62 0.65 1.00
Pvalb−ChC 1.00 0.72 0.61 0.56 0.54 0.52 0.51 0.50 0.54 1.00
Pvalb 1.18 0.87 0.76 0.68 0.67 0.62 0.61 0.60 0.59 1.00
PN 1.41 1.08 0.93 0.88 0.82 0.74 0.70 0.67 0.73 1.00
MSN−D2 1.08 0.70 0.63 0.57 0.52 0.51 0.53 0.51 0.56 1.00
MSN−D1 1.33 0.76 0.60 0.53 0.53 0.49 0.47 0.47 0.54 1.00
Lamp5−Lhx6 0.89 0.71 0.67 0.63 0.58 0.60 0.59 0.60 0.65 1.00
Lamp5 1.04 0.84 0.76 0.77 0.72 0.70 0.67 0.71 0.70 1.00
Foxp2 0.72 0.46 0.41 0.37 0.36 0.34 0.36 0.36 0.43 1.00
Chd7 1.09 0.82 0.75 0.72 0.68 0.64 0.61 0.61 0.62 1.00
CB 0.63 0.59 0.58 0.59 0.56 0.57 0.56 0.59 0.69 1.00
0.0−0.1 0.1−0.2 0.2−0.3 0.3−0.4 0.4−0.5 0.5−0.6 0.6−0.7 0.7−0.8 0.8−0.9 0.9−1.0
Percent 0.002 0.003

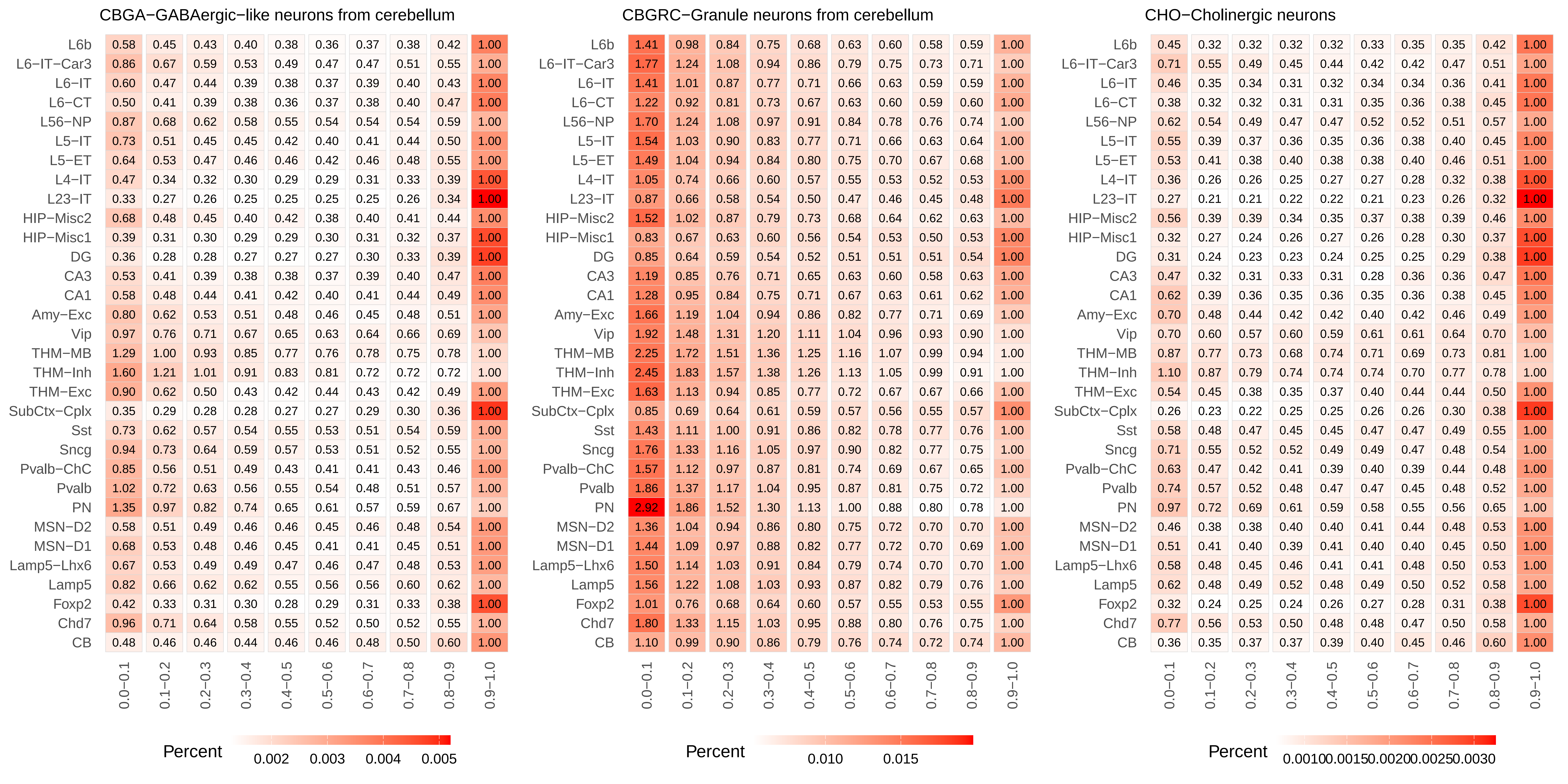
CBGA−GABAergic−like neurons from cerebellum
L6b 0.58 0.45 0.43 0.40 0.38 0.36 0.37 0.38 0.42 1.00
L6−IT−Car3 0.86 0.67 0.59 0.53 0.49 0.47 0.47 0.51 0.55 1.00
L6−IT 0.60 0.47 0.44 0.39 0.38 0.37 0.39 0.40 0.43 1.00
L6−CT 0.50 0.41 0.39 0.38 0.36 0.37 0.38 0.40 0.47 1.00
L56−NP 0.87 0.68 0.62 0.58 0.55 0.54 0.54 0.54 0.59 1.00
L5−IT 0.73 0.51 0.45 0.45 0.42 0.40 0.41 0.44 0.50 1.00
L5−ET 0.64 0.53 0.47 0.46 0.46 0.42 0.46 0.48 0.55 1.00
L4−IT 0.47 0.34 0.32 0.30 0.29 0.29 0.31 0.33 0.39 1.00
L23−IT 0.33 0.27 0.26 0.25 0.25 0.25 0.25 0.26 0.34 1.00
HIP−Misc2 0.68 0.48 0.45 0.40 0.42 0.38 0.40 0.41 0.44 1.00
HIP−Misc1 0.39 0.31 0.30 0.29 0.29 0.30 0.31 0.32 0.37 1.00
DG 0.36 0.28 0.28 0.27 0.27 0.27 0.30 0.33 0.39 1.00
CA3 0.53 0.41 0.39 0.38 0.38 0.37 0.39 0.40 0.47 1.00
CA1 0.58 0.48 0.44 0.41 0.42 0.40 0.41 0.44 0.49 1.00
Amy−Exc 0.80 0.62 0.53 0.51 0.48 0.46 0.45 0.48 0.51 1.00
Vip 0.97 0.76 0.71 0.67 0.65 0.63 0.64 0.66 0.69 1.00
THM−MB 1.29 1.00 0.93 0.85 0.77 0.76 0.78 0.75 0.78 1.00
THM−Inh 1.60 1.21 1.01 0.91 0.83 0.81 0.72 0.72 0.72 1.00
THM−Exc 0.90 0.62 0.50 0.43 0.42 0.44 0.43 0.42 0.49 1.00
SubCtx−Cplx 0.35 0.29 0.28 0.28 0.27 0.27 0.29 0.30 0.36 1.00
Sst 0.73 0.62 0.57 0.54 0.55 0.53 0.51 0.54 0.59 1.00
Sncg 0.94 0.73 0.64 0.59 0.57 0.53 0.51 0.52 0.55 1.00
Pvalb−ChC 0.85 0.56 0.51 0.49 0.43 0.41 0.41 0.43 0.46 1.00
Pvalb 1.02 0.72 0.63 0.56 0.55 0.54 0.48 0.51 0.57 1.00
PN 1.35 0.97 0.82 0.74 0.65 0.61 0.57 0.59 0.67 1.00
MSN−D2 0.58 0.51 0.49 0.46 0.46 0.45 0.46 0.48 0.54 1.00
MSN−D1 0.68 0.53 0.48 0.46 0.45 0.41 0.41 0.45 0.51 1.00
Lamp5−Lhx6 0.67 0.53 0.49 0.49 0.47 0.46 0.47 0.48 0.53 1.00
Lamp5 0.82 0.66 0.62 0.62 0.55 0.56 0.56 0.60 0.62 1.00
Foxp2 0.42 0.33 0.31 0.30 0.28 0.29 0.31 0.33 0.38 1.00
Chd7 0.96 0.71 0.64 0.58 0.55 0.52 0.50 0.52 0.55 1.00
CB 0.48 0.46 0.46 0.44 0.46 0.46 0.48 0.50 0.60 1.00
0.0−0.1 0.1−0.2 0.2−0.3 0.3−0.4 0.4−0.5 0.5−0.6 0.6−0.7 0.7−0.8 0.8−0.9 0.9−1.0
Percent 0.002 0.003 0.004 0.005
CBGRC−Granule neurons from cerebellum
L6b 1.41 0.98 0.84 0.75 0.68 0.63 0.60 0.58 0.59 1.00
L6−IT−Car3 1.77 1.24 1.08 0.94 0.86 0.79 0.75 0.73 0.71 1.00
L6−IT 1.41 1.01 0.87 0.77 0.71 0.66 0.63 0.59 0.59 1.00
L6−CT 1.22 0.92 0.81 0.73 0.67 0.63 0.60 0.59 0.60 1.00
L56−NP 1.70 1.24 1.08 0.97 0.91 0.84 0.78 0.76 0.74 1.00
L5−IT 1.54 1.03 0.90 0.83 0.77 0.71 0.66 0.63 0.64 1.00
L5−ET 1.49 1.04 0.94 0.84 0.80 0.75 0.70 0.67 0.68 1.00
L4−IT 1.05 0.74 0.66 0.60 0.57 0.55 0.53 0.52 0.53 1.00
L23−IT 0.87 0.66 0.58 0.54 0.50 0.47 0.46 0.45 0.48 1.00
HIP−Misc2 1.52 1.02 0.87 0.79 0.73 0.68 0.64 0.62 0.63 1.00
HIP−Misc1 0.83 0.67 0.63 0.60 0.56 0.54 0.53 0.50 0.53 1.00
DG 0.85 0.64 0.59 0.54 0.52 0.51 0.51 0.51 0.54 1.00
CA3 1.19 0.85 0.76 0.71 0.65 0.63 0.60 0.58 0.63 1.00
CA1 1.28 0.95 0.84 0.75 0.71 0.67 0.63 0.61 0.62 1.00
Amy−Exc 1.66 1.19 1.04 0.94 0.86 0.82 0.77 0.71 0.69 1.00
Vip 1.92 1.48 1.31 1.20 1.11 1.04 0.96 0.93 0.90 1.00
THM−MB 2.25 1.72 1.51 1.36 1.25 1.16 1.07 0.99 0.94 1.00
THM−Inh 2.45 1.83 1.57 1.38 1.26 1.13 1.05 0.99 0.91 1.00
THM−Exc 1.63 1.13 0.94 0.85 0.77 0.72 0.67 0.67 0.66 1.00
SubCtx−Cplx 0.85 0.69 0.64 0.61 0.59 0.57 0.56 0.55 0.57 1.00
Sst 1.43 1.11 1.00 0.91 0.86 0.82 0.78 0.77 0.76 1.00
Sncg 1.76 1.33 1.16 1.05 0.97 0.90 0.82 0.77 0.75 1.00
Pvalb−ChC 1.57 1.12 0.97 0.87 0.81 0.74 0.69 0.67 0.65 1.00
Pvalb 1.86 1.37 1.17 1.04 0.95 0.87 0.81 0.75 0.72 1.00
PN 2.92 1.86 1.52 1.30 1.13 1.00 0.88 0.80 0.78 1.00
MSN−D2 1.36 1.04 0.94 0.86 0.80 0.75 0.72 0.70 0.70 1.00
MSN−D1 1.44 1.09 0.97 0.88 0.82 0.77 0.72 0.70 0.69 1.00
Lamp5−Lhx6 1.50 1.14 1.03 0.91 0.84 0.79 0.74 0.70 0.70 1.00
Lamp5 1.56 1.22 1.08 1.03 0.93 0.87 0.82 0.79 0.76 1.00
Foxp2 1.01 0.76 0.68 0.64 0.60 0.57 0.55 0.53 0.55 1.00
Chd7 1.80 1.33 1.15 1.03 0.95 0.88 0.80 0.76 0.75 1.00
CB 1.10 0.99 0.90 0.86 0.79 0.76 0.74 0.72 0.74 1.00
0.0−0.1 0.1−0.2 0.2−0.3 0.3−0.4 0.4−0.5 0.5−0.6 0.6−0.7 0.7−0.8 0.8−0.9 0.9−1.0
Percent 0.010 0.015
CHO−Cholinergic neurons
L6b 0.45 0.32 0.32 0.32 0.32 0.33 0.35 0.35 0.42 1.00
L6−IT−Car3 0.71 0.55 0.49 0.45 0.44 0.42 0.42 0.47 0.51 1.00
L6−IT 0.46 0.35 0.34 0.31 0.32 0.34 0.34 0.36 0.41 1.00
L6−CT 0.38 0.32 0.32 0.31 0.31 0.35 0.36 0.38 0.45 1.00
L56−NP 0.62 0.54 0.49 0.47 0.47 0.52 0.52 0.51 0.57 1.00
L5−IT 0.55 0.39 0.37 0.36 0.35 0.36 0.38 0.40 0.45 1.00
L5−ET 0.53 0.41 0.38 0.40 0.38 0.38 0.40 0.46 0.51 1.00
L4−IT 0.36 0.26 0.26 0.25 0.27 0.27 0.28 0.32 0.38 1.00
L23−IT 0.27 0.21 0.21 0.22 0.22 0.21 0.23 0.26 0.32 1.00
HIP−Misc2 0.56 0.39 0.39 0.34 0.35 0.37 0.38 0.39 0.46 1.00
HIP−Misc1 0.32 0.27 0.24 0.26 0.27 0.26 0.28 0.30 0.37 1.00
DG 0.31 0.24 0.23 0.23 0.24 0.25 0.25 0.29 0.38 1.00
CA3 0.47 0.32 0.31 0.33 0.31 0.28 0.36 0.36 0.47 1.00
CA1 0.62 0.39 0.36 0.35 0.36 0.35 0.36 0.38 0.45 1.00
Amy−Exc 0.70 0.48 0.44 0.42 0.42 0.40 0.42 0.46 0.49 1.00
Vip 0.70 0.60 0.57 0.60 0.59 0.61 0.61 0.64 0.70 1.00
THM−MB 0.87 0.77 0.73 0.68 0.74 0.71 0.69 0.73 0.81 1.00
THM−Inh 1.10 0.87 0.79 0.74 0.74 0.74 0.70 0.77 0.78 1.00
THM−Exc 0.54 0.45 0.38 0.35 0.37 0.40 0.44 0.44 0.50 1.00
SubCtx−Cplx 0.26 0.23 0.22 0.25 0.25 0.26 0.26 0.30 0.38 1.00
Sst 0.58 0.48 0.47 0.45 0.45 0.47 0.47 0.49 0.55 1.00
Sncg 0.71 0.55 0.52 0.52 0.49 0.49 0.47 0.48 0.54 1.00
Pvalb−ChC 0.63 0.47 0.42 0.41 0.39 0.40 0.39 0.44 0.48 1.00
Pvalb 0.74 0.57 0.52 0.48 0.47 0.47 0.45 0.48 0.52 1.00
PN 0.97 0.72 0.69 0.61 0.59 0.58 0.55 0.56 0.65 1.00
MSN−D2 0.46 0.38 0.38 0.40 0.40 0.41 0.44 0.48 0.53 1.00
MSN−D1 0.51 0.41 0.40 0.39 0.41 0.40 0.40 0.45 0.50 1.00
Lamp5−Lhx6 0.58 0.48 0.45 0.46 0.41 0.41 0.48 0.50 0.53 1.00
Lamp5 0.62 0.48 0.49 0.52 0.48 0.49 0.50 0.52 0.58 1.00
Foxp2 0.32 0.24 0.25 0.24 0.26 0.27 0.28 0.31 0.38 1.00
Chd7 0.77 0.56 0.53 0.50 0.48 0.48 0.47 0.50 0.58 1.00
CB 0.36 0.35 0.37 0.37 0.39 0.40 0.45 0.46 0.60 1.00
0.0−0.1 0.1−0.2 0.2−0.3 0.3−0.4 0.4−0.5 0.5−0.6 0.6−0.7 0.7−0.8 0.8−0.9 0.9−1.0
Percent 0.00100.00150.00200.00250.0030

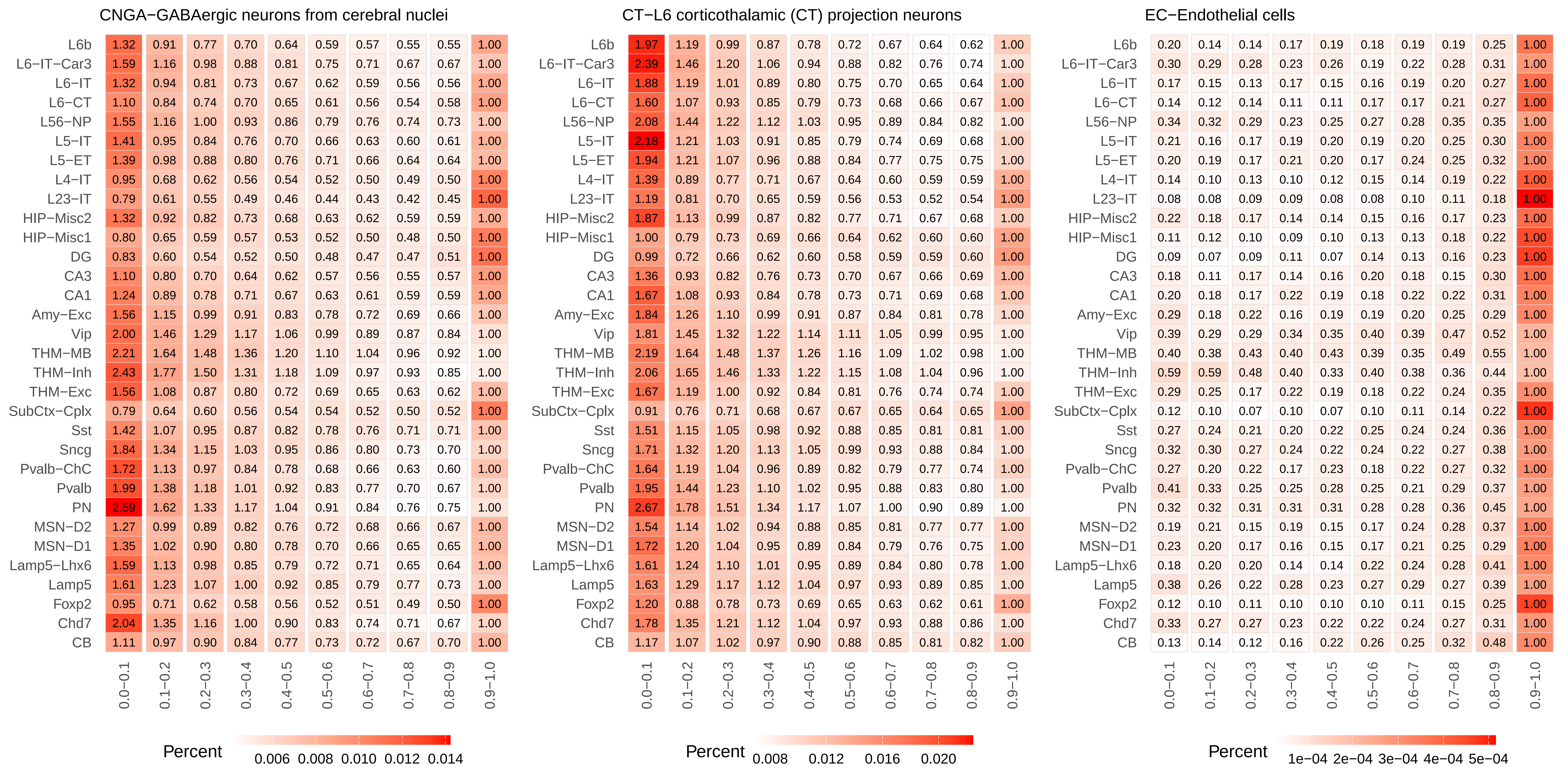
CNGA−GABAergic neurons from cerebral nuclei
CT−L6 corticothalamic (CT) projection neurons
EC−Endothelial cells
L6b
L6−IT−Car3
L6−IT
L6−CT
L56−NP
L5−IT
L5−ET
L4−IT
L23−IT
HIP−Misc2
HIP−Misc1
DG
CA3
CA1
Amy−Exc
Vip
THM−MB
THM−Inh
THM−Exc
SubCtx−Cplx
Sst
Sncg
Pvalb−ChC
Pvalb
PN
MSN−D2
MSN−D1
Lamp5−Lhx6
Lamp5
Foxp2
Chd7
CB
0.0−0.1
0.1−0.2
0.2−0.3
0.3−0.4
0.4−0.5
0.5−0.6
0.6−0.7
0.7−0.8
0.8−0.9
0.9−1.0
Percent
0.006 0.008 0.010 0.012 0.014
Percent
0.008 0.012 0.016 0.020
Percent
1e−04 2e−04 3e−04 4e−04 5e−04

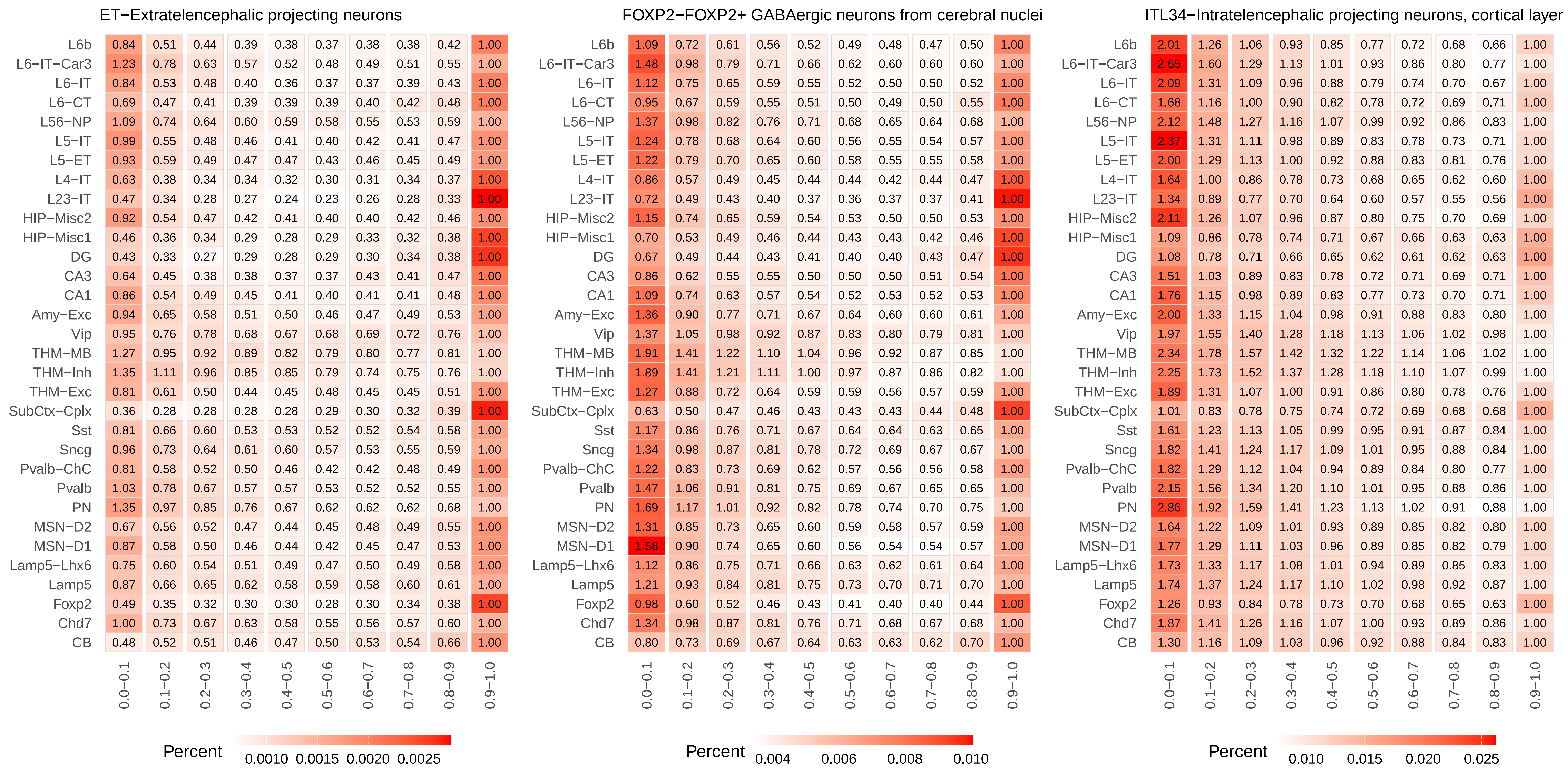
ET−Extratelencephalic projecting neurons
L6b 0.84 0.51 0.44 0.39 0.38 0.37 0.38 0.38 0.42 1.00
L6−IT−Car3 1.23 0.78 0.63 0.57 0.52 0.48 0.49 0.51 0.55 1.00
L6−IT 0.84 0.53 0.48 0.40 0.36 0.37 0.37 0.39 0.43 1.00
L6−CT 0.69 0.47 0.41 0.39 0.39 0.39 0.40 0.42 0.48 1.00
L56−NP 1.09 0.74 0.64 0.60 0.59 0.58 0.55 0.53 0.59 1.00
L5−IT 0.99 0.55 0.48 0.46 0.41 0.40 0.42 0.41 0.47 1.00
L5−ET 0.93 0.59 0.49 0.47 0.47 0.43 0.46 0.45 0.49 1.00
L4−IT 0.63 0.38 0.34 0.34 0.32 0.30 0.31 0.34 0.37 1.00
L23−IT 0.47 0.34 0.28 0.27 0.24 0.23 0.26 0.28 0.33 1.00
HIP−Misc2 0.92 0.54 0.47 0.42 0.41 0.40 0.40 0.42 0.46 1.00
HIP−Misc1 0.46 0.36 0.34 0.29 0.28 0.29 0.33 0.32 0.38 1.00
DG 0.43 0.33 0.27 0.29 0.28 0.29 0.30 0.34 0.38 1.00
CA3 0.64 0.45 0.38 0.38 0.37 0.37 0.43 0.41 0.47 1.00
CA1 0.86 0.54 0.49 0.45 0.41 0.40 0.41 0.41 0.48 1.00
Amy−Exc 0.94 0.65 0.58 0.51 0.50 0.46 0.47 0.49 0.53 1.00
Vip 0.95 0.76 0.78 0.68 0.67 0.68 0.69 0.72 0.76 1.00
THM−MB 1.27 0.95 0.92 0.89 0.82 0.79 0.80 0.77 0.81 1.00
THM−Inh 1.35 1.11 0.96 0.85 0.85 0.79 0.74 0.75 0.76 1.00
THM−Exc 0.81 0.61 0.50 0.44 0.45 0.48 0.45 0.45 0.51 1.00
SubCtx−Cplx 0.36 0.28 0.28 0.28 0.28 0.29 0.30 0.32 0.39 1.00
Sst 0.81 0.66 0.60 0.53 0.53 0.52 0.52 0.54 0.58 1.00
Sncg 0.96 0.73 0.64 0.61 0.60 0.57 0.53 0.55 0.59 1.00
Pvalb−ChC 0.81 0.58 0.52 0.50 0.46 0.42 0.42 0.48 0.49 1.00
Pvalb 1.03 0.78 0.67 0.57 0.57 0.53 0.52 0.52 0.55 1.00
PN 1.35 0.97 0.85 0.76 0.67 0.62 0.62 0.62 0.68 1.00
MSN−D2 0.67 0.56 0.52 0.47 0.44 0.45 0.48 0.49 0.55 1.00
MSN−D1 0.87 0.58 0.50 0.46 0.44 0.42 0.45 0.47 0.53 1.00
Lamp5−Lhx6 0.75 0.60 0.54 0.51 0.49 0.47 0.50 0.49 0.58 1.00
Lamp5 0.87 0.66 0.65 0.62 0.58 0.59 0.58 0.60 0.61 1.00
Foxp2 0.49 0.35 0.32 0.30 0.30 0.28 0.30 0.34 0.38 1.00
Chd7 1.00 0.73 0.67 0.63 0.58 0.55 0.56 0.57 0.60 1.00
CB 0.48 0.52 0.51 0.46 0.47 0.50 0.53 0.54 0.66 1.00
0.0−0.1 0.1−0.2 0.2−0.3 0.3−0.4 0.4−0.5 0.5−0.6 0.6−0.7 0.7−0.8 0.8−0.9 0.9−1.0
Percent
0.0010 0.0015 0.0020 0.0025
FOXP2−FOXP2+ GABAergic neurons from cerebral nuclei
L6b 1.09 0.72 0.61 0.56 0.52 0.49 0.48 0.47 0.50 1.00
L6−IT−Car3 1.48 0.98 0.79 0.71 0.66 0.62 0.60 0.60 0.60 1.00
L6−IT 1.12 0.75 0.65 0.59 0.55 0.52 0.50 0.50 0.52 1.00
L6−CT 0.95 0.67 0.59 0.55 0.51 0.50 0.49 0.50 0.55 1.00
L56−NP 1.37 0.98 0.82 0.76 0.71 0.68 0.65 0.64 0.68 1.00
L5−IT 1.24 0.78 0.68 0.64 0.60 0.56 0.55 0.54 0.57 1.00
L5−ET 1.22 0.79 0.70 0.65 0.60 0.58 0.55 0.55 0.58 1.00
L4−IT 0.86 0.57 0.49 0.45 0.44 0.44 0.42 0.44 0.47 1.00
L23−IT 0.72 0.49 0.43 0.40 0.37 0.36 0.37 0.37 0.41 1.00
HIP−Misc2 1.15 0.74 0.65 0.59 0.54 0.53 0.50 0.50 0.53 1.00
HIP−Misc1 0.70 0.53 0.49 0.46 0.44 0.43 0.43 0.42 0.46 1.00
DG 0.67 0.49 0.44 0.43 0.41 0.40 0.40 0.43 0.47 1.00
CA3 0.86 0.62 0.55 0.55 0.50 0.50 0.50 0.51 0.54 1.00
CA1 1.09 0.74 0.63 0.57 0.54 0.52 0.53 0.52 0.53 1.00
Amy−Exc 1.36 0.90 0.77 0.71 0.67 0.64 0.60 0.60 0.61 1.00
Vip 1.37 1.05 0.98 0.92 0.87 0.83 0.80 0.79 0.81 1.00
THM−MB 1.91 1.41 1.22 1.10 1.04 0.96 0.92 0.87 0.85 1.00
THM−Inh 1.89 1.41 1.21 1.11 1.00 0.97 0.87 0.86 0.82 1.00
THM−Exc 1.27 0.88 0.72 0.64 0.59 0.59 0.56 0.57 0.59 1.00
SubCtx−Cplx 0.63 0.50 0.47 0.46 0.43 0.43 0.43 0.44 0.48 1.00
Sst 1.17 0.86 0.76 0.71 0.67 0.64 0.64 0.63 0.65 1.00
Sncg 1.34 0.98 0.87 0.81 0.78 0.72 0.69 0.67 0.67 1.00
Pvalb−ChC 1.22 0.83 0.73 0.69 0.62 0.57 0.56 0.56 0.58 1.00
Pvalb 1.47 1.06 0.91 0.81 0.75 0.69 0.67 0.65 0.65 1.00
PN 1.69 1.17 1.01 0.92 0.82 0.78 0.74 0.70 0.75 1.00
MSN−D2 1.31 0.85 0.73 0.65 0.60 0.59 0.58 0.57 0.59 1.00
MSN−D1 1.58 0.90 0.74 0.65 0.60 0.56 0.54 0.54 0.57 1.00
Lamp5−Lhx6 1.12 0.86 0.75 0.71 0.66 0.63 0.62 0.61 0.64 1.00
Lamp5 1.21 0.93 0.84 0.81 0.75 0.73 0.70 0.71 0.70 1.00
Foxp2 0.98 0.60 0.52 0.46 0.43 0.41 0.40 0.40 0.44 1.00
Chd7 1.34 0.98 0.87 0.81 0.76 0.71 0.68 0.67 0.68 1.00
CB 0.80 0.73 0.69 0.67 0.64 0.63 0.63 0.62 0.70 1.00
0.0−0.1 0.1−0.2 0.2−0.3 0.3−0.4 0.4−0.5 0.5−0.6 0.6−0.7 0.7−0.8 0.8−0.9 0.9−1.0
Percent
0.004 0.006 0.008 0.010
ITL34−Intratelencephalic projecting neurons, cortical layer
L6b 2.01 1.26 1.06 0.93 0.85 0.77 0.72 0.68 0.66 1.00
L6−IT−Car3 2.65 1.60 1.29 1.13 1.01 0.93 0.86 0.80 0.77 1.00
L6−IT 2.09 1.31 1.09 0.96 0.88 0.79 0.74 0.70 0.67 1.00
L6−CT 1.68 1.16 1.00 0.90 0.82 0.78 0.72 0.69 0.71 1.00
L56−NP 2.12 1.48 1.27 1.16 1.07 0.99 0.92 0.86 0.83 1.00
L5−IT 2.37 1.31 1.11 0.98 0.89 0.83 0.78 0.73 0.71 1.00
L5−ET 2.00 1.29 1.13 1.00 0.92 0.88 0.83 0.81 0.76 1.00
L4−IT 1.64 1.00 0.86 0.78 0.73 0.68 0.65 0.62 0.60 1.00
L23−IT 1.34 0.89 0.77 0.70 0.64 0.60 0.57 0.55 0.56 1.00
HIP−Misc2 2.11 1.26 1.07 0.96 0.87 0.80 0.75 0.70 0.69 1.00
HIP−Misc1 1.09 0.86 0.78 0.74 0.71 0.67 0.66 0.63 0.63 1.00
DG 1.08 0.78 0.71 0.66 0.65 0.62 0.61 0.62 0.63 1.00
CA3 1.51 1.03 0.89 0.83 0.78 0.72 0.71 0.69 0.71 1.00
CA1 1.76 1.15 0.98 0.89 0.83 0.77 0.73 0.70 0.71 1.00
Amy−Exc 2.00 1.33 1.15 1.04 0.98 0.91 0.88 0.83 0.80 1.00
Vip 1.97 1.55 1.40 1.28 1.18 1.13 1.06 1.02 0.98 1.00
THM−MB 2.34 1.78 1.57 1.42 1.32 1.22 1.14 1.06 1.02 1.00
THM−Inh 2.25 1.73 1.52 1.37 1.28 1.18 1.10 1.07 0.99 1.00
THM−Exc 1.89 1.31 1.07 1.00 0.91 0.86 0.80 0.78 0.76 1.00
SubCtx−Cplx 1.01 0.83 0.78 0.75 0.74 0.72 0.69 0.68 0.68 1.00
Sst 1.61 1.23 1.13 1.05 0.99 0.95 0.91 0.87 0.84 1.00
Sncg 1.82 1.41 1.24 1.17 1.09 1.01 0.95 0.88 0.84 1.00
Pvalb−ChC 1.82 1.29 1.12 1.04 0.94 0.89 0.84 0.80 0.77 1.00
Pvalb 2.15 1.56 1.34 1.20 1.10 1.01 0.95 0.88 0.86 1.00
PN 2.86 1.92 1.59 1.41 1.23 1.13 1.02 0.91 0.88 1.00
MSN−D2 1.64 1.22 1.09 1.01 0.93 0.89 0.85 0.82 0.80 1.00
MSN−D1 1.77 1.29 1.11 1.03 0.96 0.89 0.85 0.82 0.79 1.00
Lamp5−Lhx6 1.73 1.33 1.17 1.08 1.01 0.94 0.89 0.85 0.83 1.00
Lamp5 1.74 1.37 1.24 1.17 1.10 1.02 0.98 0.92 0.87 1.00
Foxp2 1.26 0.93 0.84 0.78 0.73 0.70 0.68 0.65 0.63 1.00
Chd7 1.87 1.41 1.26 1.16 1.07 1.00 0.93 0.89 0.86 1.00
CB 1.30 1.16 1.09 1.03 0.96 0.92 0.88 0.84 0.83 1.00
0.0−0.1 0.1−0.2 0.2−0.3 0.3−0.4 0.4−0.5 0.5−0.6 0.6−0.7 0.7−0.8 0.8−0.9 0.9−1.0
Percent
0.010 0.015 0.020 0.025

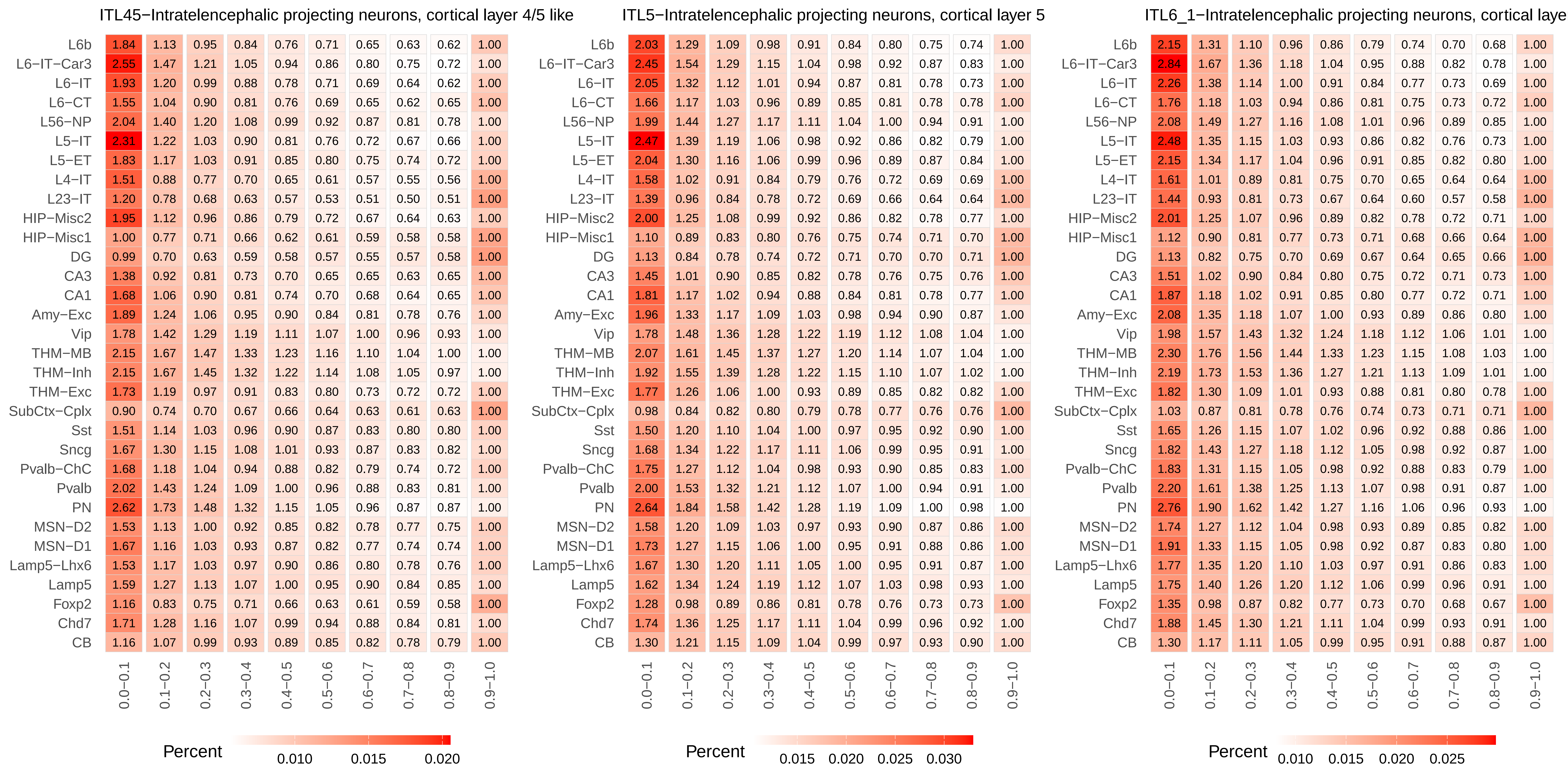
ITL45−Intratelencephalic projecting neurons, cortical layer 4/5 like
ITL5−Intratelencephalic projecting neurons, cortical layer 5
ITL6_1−Intratelencephalic projecting neurons, cortical laye
L6b
L6−IT−Car3
L6−IT
L6−CT
L56−NP
L5−IT
L5−ET
L4−IT
L23−IT
HIP−Misc2
HIP−Misc1
DG
CA3
CA1
Amy−Exc
Vip
THM−MB
THM−Inh
THM−Exc
SubCtx−Cplx
Sst
Sncg
Pvalb−ChC
Pvalb
PN
MSN−D2
MSN−D1
Lamp5−Lhx6
Lamp5
Foxp2
Chd7
CB
0.0−0.1
0.1−0.2
0.2−0.3
0.3−0.4
0.4−0.5
0.5−0.6
0.6−0.7
0.7−0.8
0.8−0.9
0.9−1.0
Percent
0.010
0.015
0.020
Percent
0.015
0.020
0.025
0.030
Percent
0.010
0.015
0.020
0.025

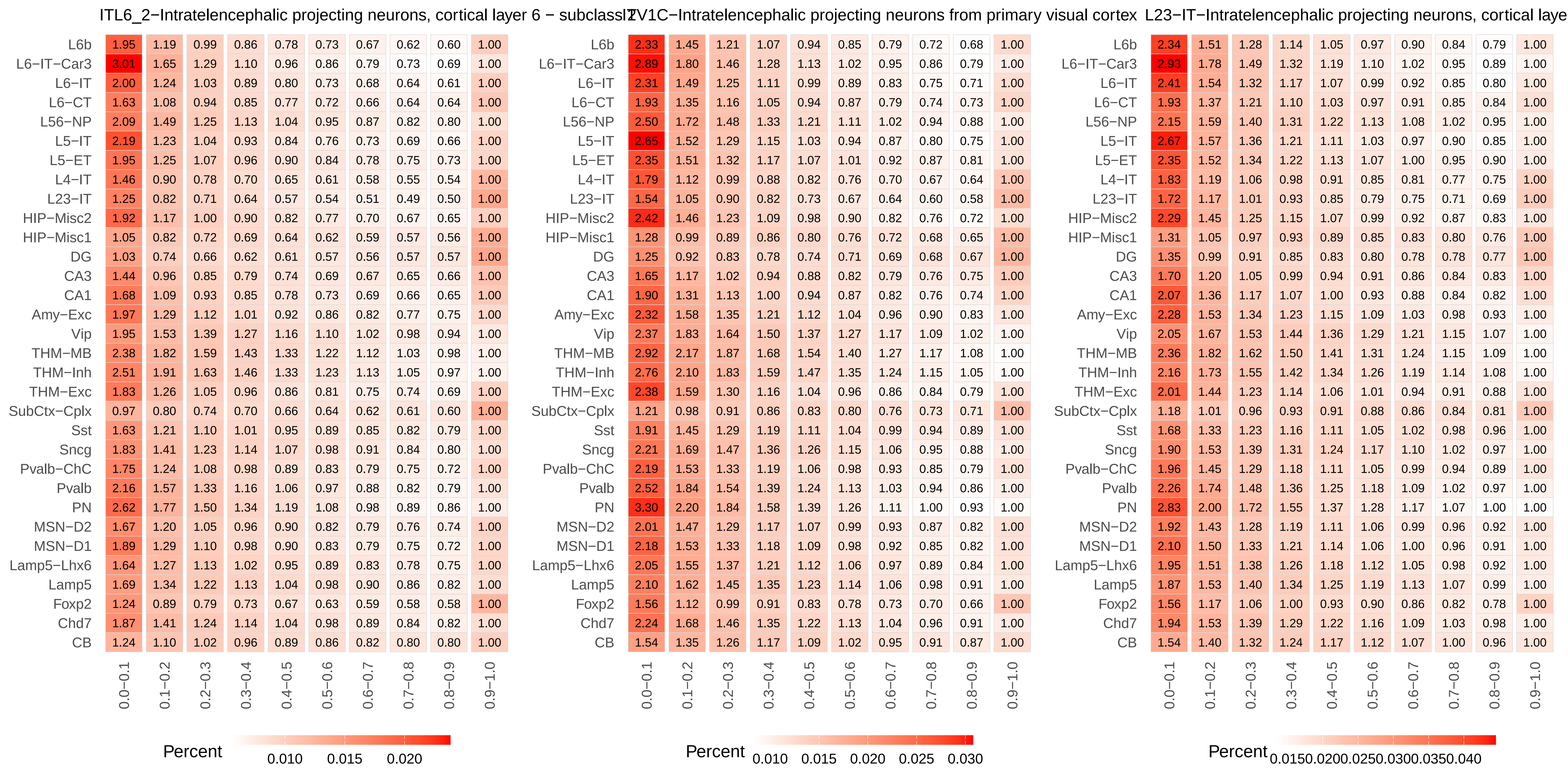
ITL6_2−Intratelencephalic projecting neurons, cortical layer 6 − subclass 2
IT V1C−Intratelencephalic projecting neurons from primary visual cortex
L23−IT−Intratelencephalic projecting neurons, cortical laye
L6b
L6−IT−Car3
L6−IT
L6−CT
L56−NP
L5−IT
L5−ET
L4−IT
L23−IT
HIP−Misc2
HIP−Misc1
DG
CA3
CA1
Amy−Exc
Vip
THM−MB
THM−Inh
THM−Exc
SubCtx−Cplx
Sst
Sncg
Pvalb−ChC
Pvalb
PN
MSN−D2
MSN−D1
Lamp5−Lhx6
Lamp5
Foxp2
Chd7
CB
0.0−0.1
0.1−0.2
0.2−0.3
0.3−0.4
0.4−0.5
0.5−0.6
0.6−0.7
0.7−0.8
0.8−0.9
0.9−1.0
Percent
0.010 0.015 0.020
Percent
0.010 0.015 0.020 0.025 0.030
Percent
0.0150.0200.0250.0300.0350.040

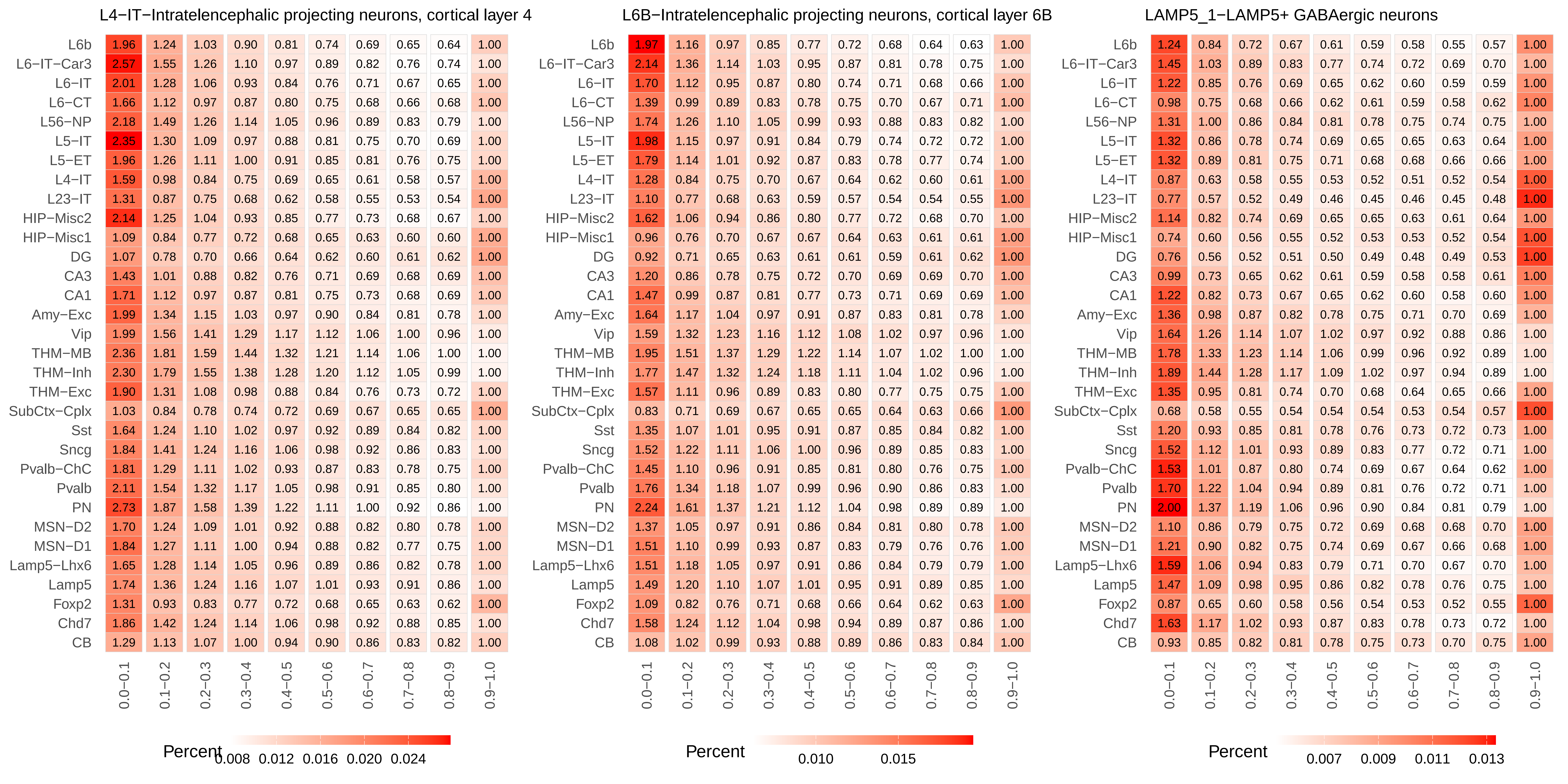
L4−IT−Intratelencephalic projecting neurons, cortical layer 4
L6B−Intratelencephalic projecting neurons, cortical layer 6B
LAMP5_1−LAMP5+ GABAergic neurons
L6b
L6−IT−Car3
L6−IT
L6−CT
L56−NP
L5−IT
L5−ET
L4−IT
L23−IT
HIP−Misc2
HIP−Misc1
DG
CA3
CA1
Amy−Exc
Vip
THM−MB
THM−Inh
THM−Exc
SubCtx−Cplx
Sst
Sncg
Pvalb−ChC
Pvalb
PN
MSN−D2
MSN−D1
Lamp5−Lhx6
Lamp5
Foxp2
Chd7
CB
0.0−0.1
0.1−0.2
0.2−0.3
0.3−0.4
0.4−0.5
0.5−0.6
0.6−0.7
0.7−0.8
0.8−0.9
0.9−1.0
Percent
0.008 0.012 0.016 0.020 0.024
Percent
0.010 0.015
Percent
0.007 0.009 0.011 0.013

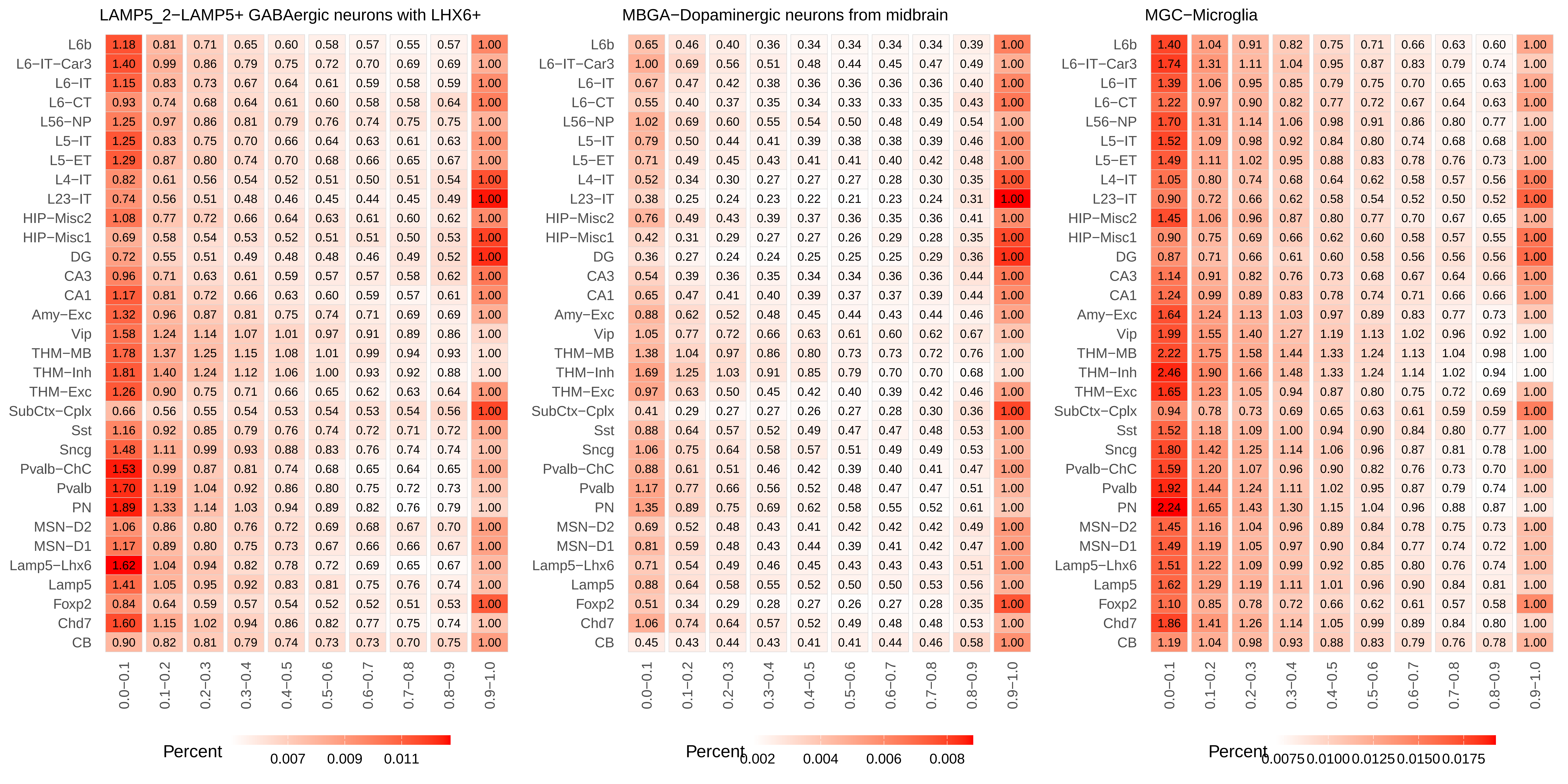
LAMP5_2−LAMP5+ GABAergic neurons with LHX6+
MBGA−Dopaminergic neurons from midbrain
MGC−Microglia
L6b
L6−IT−Car3
L6−IT
L6−CT
L56−NP
L5−IT
L5−ET
L4−IT
L23−IT
HIP−Misc2
HIP−Misc1
DG
CA3
CA1
Amy−Exc
Vip
THM−MB
THM−Inh
THM−Exc
SubCtx−Cplx
Sst
Sncg
Pvalb−ChC
Pvalb
PN
MSN−D2
MSN−D1
Lamp5−Lhx6
Lamp5
Foxp2
Chd7
CB
0.0−0.1
0.1−0.2
0.2−0.3
0.3−0.4
0.4−0.5
0.5−0.6
0.6−0.7
0.7−0.8
0.8−0.9
0.9−1.0
Percent
0.007
0.009
0.011
Percent
0.002
0.004
0.006
0.008
Percent
0.0075
0.0100
0.0125
0.0150
0.0175

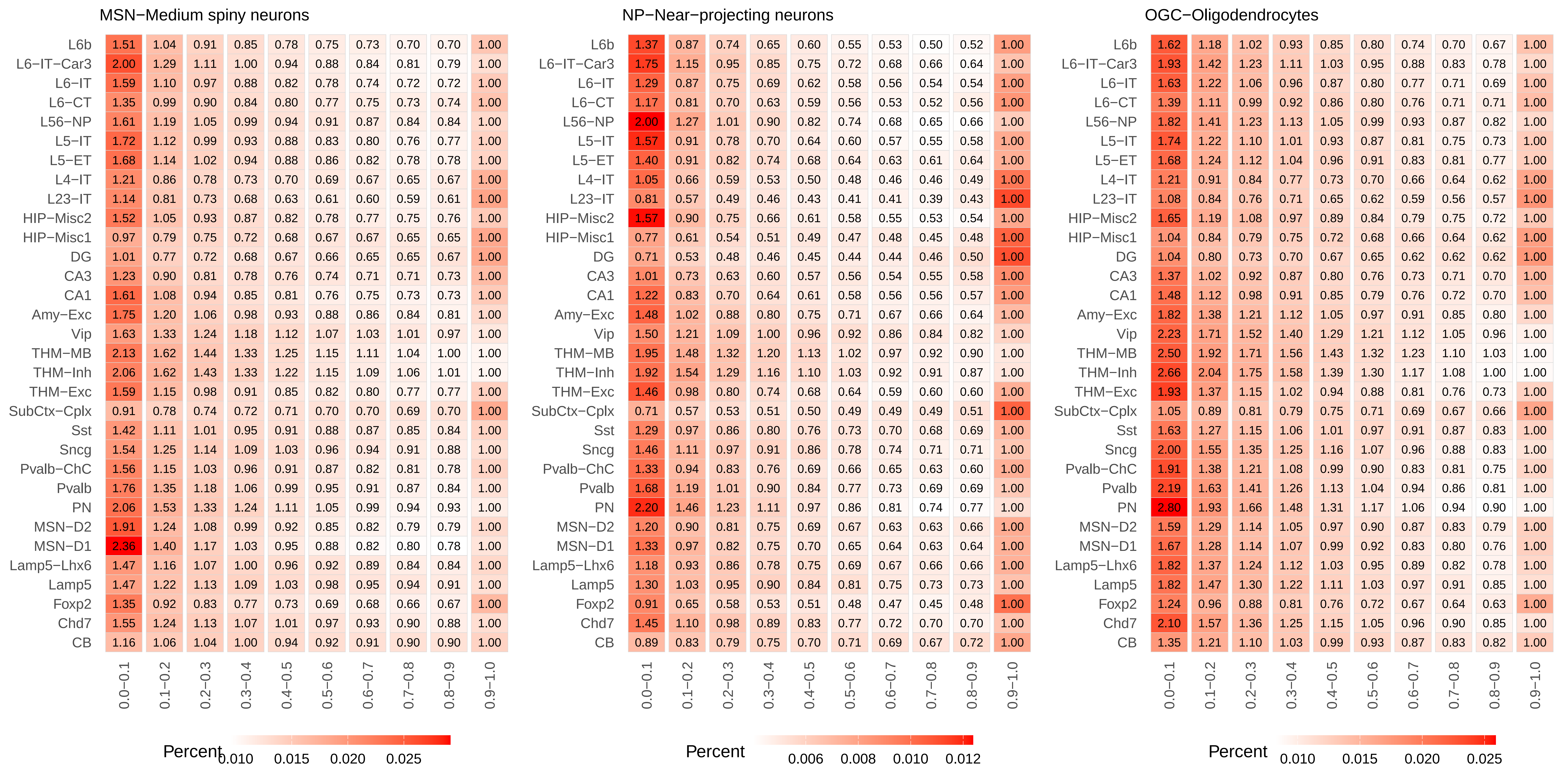
MSN−Medium spiny neurons
L6b 1.51 1.04 0.91 0.85 0.78 0.75 0.73 0.70 0.70 1.00
L6−IT−Car3 2.00 1.29 1.11 1.00 0.94 0.88 0.84 0.81 0.79 1.00
L6−IT 1.59 1.10 0.97 0.88 0.82 0.78 0.74 0.72 0.72 1.00
L6−CT 1.35 0.99 0.90 0.84 0.80 0.77 0.75 0.73 0.74 1.00
L56−NP 1.61 1.19 1.05 0.99 0.94 0.91 0.87 0.84 0.84 1.00
L5−IT 1.72 1.12 0.99 0.93 0.88 0.83 0.80 0.76 0.77 1.00
L5−ET 1.68 1.14 1.02 0.94 0.88 0.86 0.82 0.78 0.78 1.00
L4−IT 1.21 0.86 0.78 0.73 0.70 0.69 0.67 0.65 0.67 1.00
L23−IT 1.14 0.81 0.73 0.68 0.63 0.61 0.60 0.59 0.61 1.00
HIP−Misc2 1.52 1.05 0.93 0.87 0.82 0.78 0.77 0.75 0.76 1.00
HIP−Misc1 0.97 0.79 0.75 0.72 0.68 0.67 0.67 0.65 0.65 1.00
DG 1.01 0.77 0.72 0.68 0.67 0.66 0.65 0.65 0.67 1.00
CA3 1.23 0.90 0.81 0.78 0.76 0.74 0.71 0.71 0.73 1.00
CA1 1.61 1.08 0.94 0.85 0.81 0.76 0.75 0.73 0.73 1.00
Amy−Exc 1.75 1.20 1.06 0.98 0.93 0.88 0.86 0.84 0.81 1.00
Vip 1.63 1.33 1.24 1.18 1.12 1.07 1.03 1.01 0.97 1.00
THM−MB 2.13 1.62 1.44 1.33 1.25 1.15 1.11 1.04 1.00 1.00
THM−Inh 2.06 1.62 1.43 1.33 1.22 1.15 1.09 1.06 1.01 1.00
THM−Exc 1.59 1.15 0.98 0.91 0.85 0.82 0.80 0.77 0.77 1.00
SubCtx−Cplx 0.91 0.78 0.74 0.72 0.71 0.70 0.70 0.69 0.70 1.00
Sst 1.42 1.11 1.01 0.95 0.91 0.88 0.87 0.85 0.84 1.00
Sncg 1.54 1.25 1.14 1.09 1.03 0.96 0.94 0.91 0.88 1.00
Pvalb−ChC 1.56 1.15 1.03 0.96 0.91 0.87 0.82 0.81 0.78 1.00
Pvalb 1.76 1.35 1.18 1.06 0.99 0.95 0.91 0.87 0.84 1.00
PN 2.06 1.53 1.33 1.24 1.11 1.05 0.99 0.94 0.93 1.00
MSN−D2 1.91 1.24 1.08 0.99 0.92 0.85 0.82 0.79 0.79 1.00
MSN−D1 2.36 1.40 1.17 1.03 0.95 0.88 0.82 0.80 0.78 1.00
Lamp5−Lhx6 1.47 1.16 1.07 1.00 0.96 0.92 0.89 0.84 0.84 1.00
Lamp5 1.47 1.22 1.13 1.09 1.03 0.98 0.95 0.94 0.91 1.00
Foxp2 1.35 0.92 0.83 0.77 0.73 0.69 0.68 0.66 0.67 1.00
Chd7 1.55 1.24 1.13 1.07 1.01 0.97 0.93 0.90 0.88 1.00
CB 1.16 1.06 1.04 1.00 0.94 0.92 0.91 0.90 0.90 1.00
0.0−0.1 0.1−0.2 0.2−0.3 0.3−0.4 0.4−0.5 0.5−0.6 0.6−0.7 0.7−0.8 0.8−0.9 0.9−1.0
Percent 0.010 0.015 0.020 0.025
NP−Near−projecting neurons
L6b 1.37 0.87 0.74 0.65 0.60 0.55 0.53 0.50 0.52 1.00
L6−IT−Car3 1.75 1.15 0.95 0.85 0.75 0.72 0.68 0.66 0.64 1.00
L6−IT 1.29 0.87 0.75 0.69 0.62 0.58 0.56 0.54 0.54 1.00
L6−CT 1.17 0.81 0.70 0.63 0.59 0.56 0.53 0.52 0.56 1.00
L56−NP 2.00 1.27 1.01 0.90 0.82 0.74 0.68 0.65 0.66 1.00
L5−IT 1.57 0.91 0.78 0.70 0.64 0.60 0.57 0.55 0.58 1.00
L5−ET 1.40 0.91 0.82 0.74 0.68 0.64 0.63 0.61 0.64 1.00
L4−IT 1.05 0.66 0.59 0.53 0.50 0.48 0.46 0.46 0.49 1.00
L23−IT 0.81 0.57 0.49 0.46 0.43 0.41 0.41 0.39 0.43 1.00
HIP−Misc2 1.57 0.90 0.75 0.66 0.61 0.58 0.55 0.53 0.54 1.00
HIP−Misc1 0.77 0.61 0.54 0.51 0.49 0.47 0.48 0.45 0.48 1.00
DG 0.71 0.53 0.48 0.46 0.45 0.44 0.44 0.46 0.50 1.00
CA3 1.01 0.73 0.63 0.60 0.57 0.56 0.54 0.55 0.58 1.00
CA1 1.22 0.83 0.70 0.64 0.61 0.58 0.56 0.56 0.57 1.00
Amy−Exc 1.48 1.02 0.88 0.80 0.75 0.71 0.67 0.66 0.64 1.00
Vip 1.50 1.21 1.09 1.00 0.96 0.92 0.86 0.84 0.82 1.00
THM−MB 1.95 1.48 1.32 1.20 1.13 1.02 0.97 0.92 0.90 1.00
THM−Inh 1.92 1.54 1.29 1.16 1.10 1.03 0.92 0.91 0.87 1.00
THM−Exc 1.46 0.98 0.80 0.74 0.68 0.64 0.59 0.60 0.60 1.00
SubCtx−Cplx 0.71 0.57 0.53 0.51 0.50 0.49 0.49 0.49 0.51 1.00
Sst 1.29 0.97 0.86 0.80 0.76 0.73 0.70 0.68 0.69 1.00
Sncg 1.46 1.11 0.97 0.91 0.86 0.78 0.74 0.71 0.71 1.00
Pvalb−ChC 1.33 0.94 0.83 0.76 0.69 0.66 0.65 0.63 0.60 1.00
Pvalb 1.68 1.19 1.01 0.90 0.84 0.77 0.73 0.69 0.69 1.00
PN 2.20 1.46 1.23 1.11 0.97 0.86 0.81 0.74 0.77 1.00
MSN−D2 1.20 0.90 0.81 0.75 0.69 0.67 0.63 0.63 0.66 1.00
MSN−D1 1.33 0.97 0.82 0.75 0.70 0.65 0.64 0.63 0.64 1.00
Lamp5−Lhx6 1.18 0.93 0.86 0.78 0.75 0.69 0.67 0.66 0.66 1.00
Lamp5 1.30 1.03 0.95 0.90 0.84 0.81 0.75 0.73 0.73 1.00
Foxp2 0.91 0.65 0.58 0.53 0.51 0.48 0.47 0.45 0.48 1.00
Chd7 1.45 1.10 0.98 0.89 0.83 0.77 0.72 0.70 0.70 1.00
CB 0.89 0.83 0.79 0.75 0.70 0.71 0.69 0.67 0.72 1.00
0.0−0.1 0.1−0.2 0.2−0.3 0.3−0.4 0.4−0.5 0.5−0.6 0.6−0.7 0.7−0.8 0.8−0.9 0.9−1.0
Percent 0.006 0.008 0.010 0.012
OGC−Oligodendrocytes
L6b 1.62 1.18 1.02 0.93 0.85 0.80 0.74 0.70 0.67 1.00
L6−IT−Car3 1.93 1.42 1.23 1.11 1.03 0.95 0.88 0.83 0.78 1.00
L6−IT 1.63 1.22 1.06 0.96 0.87 0.80 0.77 0.71 0.69 1.00
L6−CT 1.39 1.11 0.99 0.92 0.86 0.80 0.76 0.71 0.71 1.00
L56−NP 1.82 1.41 1.23 1.13 1.05 0.99 0.93 0.87 0.82 1.00
L5−IT 1.74 1.22 1.10 1.01 0.93 0.87 0.81 0.75 0.73 1.00
L5−ET 1.68 1.24 1.12 1.04 0.96 0.91 0.83 0.81 0.77 1.00
L4−IT 1.21 0.91 0.84 0.77 0.73 0.70 0.66 0.64 0.62 1.00
L23−IT 1.08 0.84 0.76 0.71 0.65 0.62 0.59 0.56 0.57 1.00
HIP−Misc2 1.65 1.19 1.08 0.97 0.89 0.84 0.79 0.75 0.72 1.00
HIP−Misc1 1.04 0.84 0.79 0.75 0.72 0.68 0.66 0.64 0.62 1.00
DG 1.04 0.80 0.73 0.70 0.67 0.65 0.62 0.62 0.62 1.00
CA3 1.37 1.02 0.92 0.87 0.80 0.76 0.73 0.71 0.70 1.00
CA1 1.48 1.12 0.98 0.91 0.85 0.79 0.76 0.72 0.70 1.00
Amy−Exc 1.82 1.38 1.21 1.12 1.05 0.97 0.91 0.85 0.80 1.00
Vip 2.23 1.71 1.52 1.40 1.29 1.21 1.12 1.05 0.96 1.00
THM−MB 2.50 1.92 1.71 1.56 1.43 1.32 1.23 1.10 1.03 1.00
THM−Inh 2.66 2.04 1.75 1.58 1.39 1.30 1.17 1.08 1.00 1.00
THM−Exc 1.93 1.37 1.15 1.02 0.94 0.88 0.81 0.76 0.73 1.00
SubCtx−Cplx 1.05 0.89 0.81 0.79 0.75 0.71 0.69 0.67 0.66 1.00
Sst 1.63 1.27 1.15 1.06 1.01 0.97 0.91 0.87 0.83 1.00
Sncg 2.00 1.55 1.35 1.25 1.16 1.07 0.96 0.88 0.83 1.00
Pvalb−ChC 1.91 1.38 1.21 1.08 0.99 0.90 0.83 0.81 0.75 1.00
Pvalb 2.19 1.63 1.41 1.26 1.13 1.04 0.94 0.86 0.81 1.00
PN 2.80 1.93 1.66 1.48 1.31 1.17 1.06 0.94 0.90 1.00
MSN−D2 1.59 1.29 1.14 1.05 0.97 0.90 0.87 0.83 0.79 1.00
MSN−D1 1.67 1.28 1.14 1.07 0.99 0.92 0.83 0.80 0.76 1.00
Lamp5−Lhx6 1.82 1.37 1.24 1.12 1.03 0.95 0.89 0.82 0.78 1.00
Lamp5 1.82 1.47 1.30 1.22 1.11 1.03 0.97 0.91 0.85 1.00
Foxp2 1.24 0.96 0.88 0.81 0.76 0.72 0.67 0.64 0.63 1.00
Chd7 2.10 1.57 1.36 1.25 1.15 1.05 0.96 0.90 0.85 1.00
CB 1.35 1.21 1.10 1.03 0.99 0.93 0.87 0.83 0.82 1.00
0.0−0.1 0.1−0.2 0.2−0.3 0.3−0.4 0.4−0.5 0.5−0.6 0.6−0.7 0.7−0.8 0.8−0.9 0.9−1.0
Percent 0.010 0.015 0.020 0.025

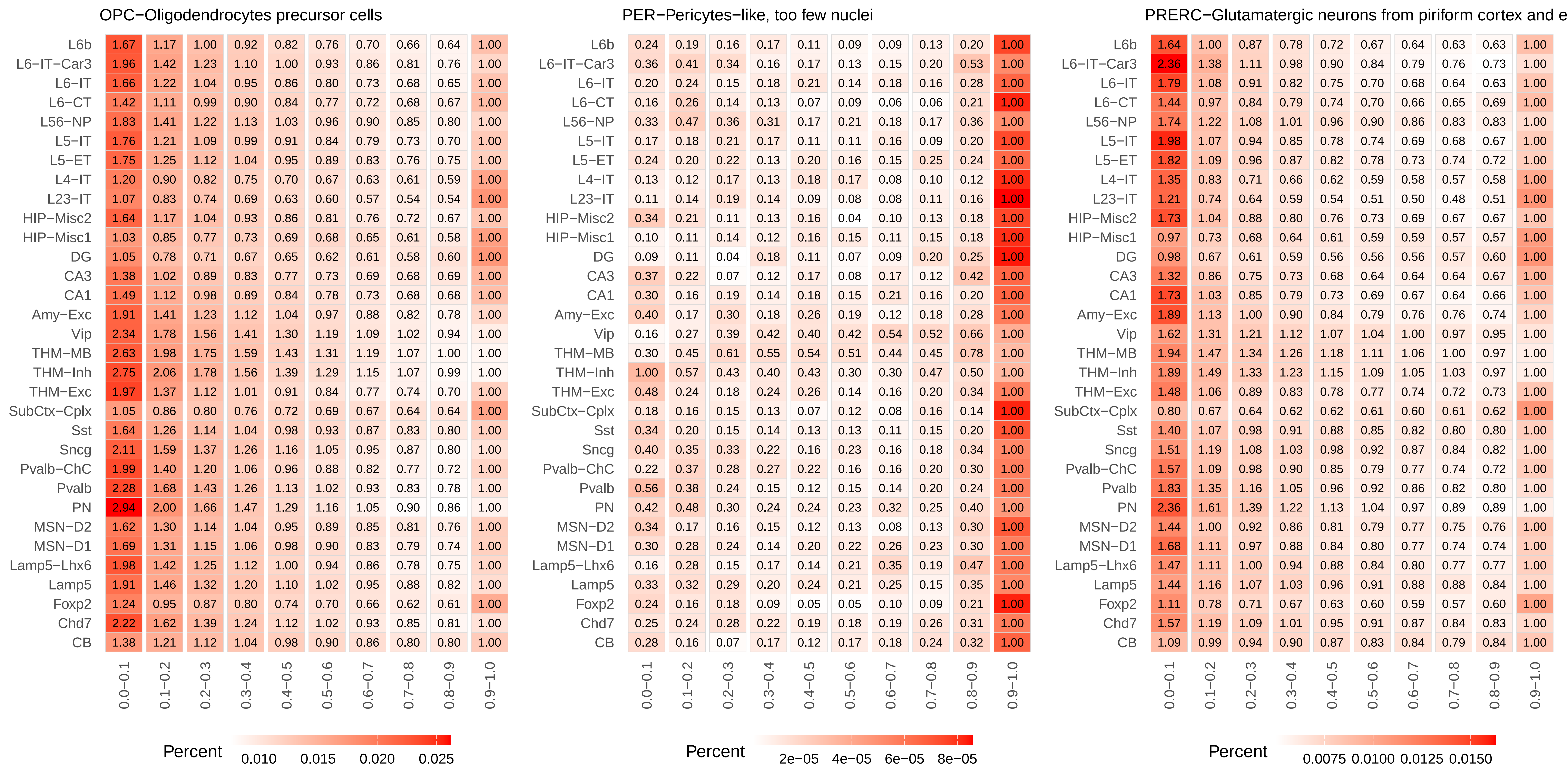
OPC−Oligodendrocytes precursor cells
L6b 1.67 1.17 1.00 0.92 0.82 0.76 0.70 0.66 0.64 1.00
L6−IT−Car3 1.96 1.42 1.23 1.10 1.00 0.93 0.86 0.81 0.76 1.00
L6−IT 1.66 1.22 1.04 0.95 0.86 0.80 0.73 0.68 0.65 1.00
L6−CT 1.42 1.11 0.99 0.90 0.84 0.77 0.72 0.68 0.67 1.00
L56−NP 1.83 1.41 1.22 1.13 1.03 0.96 0.90 0.85 0.80 1.00
L5−IT 1.76 1.21 1.09 0.99 0.91 0.84 0.79 0.73 0.70 1.00
L5−ET 1.75 1.25 1.12 1.04 0.95 0.89 0.83 0.76 0.75 1.00
L4−IT 1.20 0.90 0.82 0.75 0.70 0.67 0.63 0.61 0.59 1.00
L23−IT 1.07 0.83 0.74 0.69 0.63 0.60 0.57 0.54 0.54 1.00
HIP−Misc2 1.64 1.17 1.04 0.93 0.86 0.81 0.76 0.72 0.67 1.00
HIP−Misc1 1.03 0.85 0.77 0.73 0.69 0.68 0.65 0.61 0.58 1.00
DG 1.05 0.78 0.71 0.67 0.65 0.62 0.61 0.58 0.60 1.00
CA3 1.38 1.02 0.89 0.83 0.77 0.73 0.69 0.68 0.69 1.00
CA1 1.49 1.12 0.98 0.89 0.84 0.78 0.73 0.68 0.68 1.00
Amy−Exc 1.91 1.41 1.23 1.12 1.04 0.97 0.88 0.82 0.78 1.00
Vip 2.34 1.78 1.56 1.41 1.30 1.19 1.09 1.02 0.94 1.00
THM−MB 2.63 1.98 1.75 1.59 1.43 1.31 1.19 1.07 1.00 1.00
THM−Inh 2.75 2.06 1.78 1.56 1.39 1.29 1.15 1.07 0.99 1.00
THM−Exc 1.97 1.37 1.12 1.01 0.91 0.84 0.77 0.74 0.70 1.00
SubCtx−Cplx 1.05 0.86 0.80 0.76 0.72 0.69 0.67 0.64 0.64 1.00
Sst 1.64 1.26 1.14 1.04 0.98 0.93 0.87 0.83 0.80 1.00
Sncg 2.11 1.59 1.37 1.26 1.16 1.05 0.95 0.87 0.80 1.00
Pvalb−ChC 1.99 1.40 1.20 1.06 0.96 0.88 0.82 0.77 0.72 1.00
Pvalb 2.28 1.68 1.43 1.26 1.13 1.02 0.93 0.83 0.78 1.00
PN 2.94 2.00 1.66 1.47 1.29 1.16 1.05 0.90 0.86 1.00
MSN−D2 1.62 1.30 1.14 1.04 0.95 0.89 0.85 0.81 0.76 1.00
MSN−D1 1.69 1.31 1.15 1.06 0.98 0.90 0.83 0.79 0.74 1.00
Lamp5−Lhx6 1.98 1.42 1.25 1.12 1.00 0.94 0.86 0.78 0.75 1.00
Lamp5 1.91 1.46 1.32 1.20 1.10 1.02 0.95 0.88 0.82 1.00
Foxp2 1.24 0.95 0.87 0.80 0.74 0.70 0.66 0.62 0.61 1.00
Chd7 2.22 1.62 1.39 1.24 1.12 1.02 0.93 0.85 0.81 1.00
CB 1.38 1.21 1.12 1.04 0.98 0.90 0.86 0.80 0.80 1.00
0.0−0.1 0.1−0.2 0.2−0.3 0.3−0.4 0.4−0.5 0.5−0.6 0.6−0.7 0.7−0.8 0.8−0.9 0.9−1.0
Percent
0.010 0.015 0.020 0.025
PER−Pericytes−like, too few nuclei
L6b 0.24 0.19 0.16 0.17 0.11 0.09 0.09 0.13 0.20 1.00
L6−IT−Car3 0.36 0.41 0.34 0.16 0.17 0.13 0.15 0.20 0.53 1.00
L6−IT 0.20 0.24 0.15 0.18 0.21 0.14 0.18 0.16 0.28 1.00
L6−CT 0.16 0.26 0.14 0.13 0.07 0.09 0.06 0.06 0.21 1.00
L56−NP 0.33 0.47 0.36 0.31 0.17 0.21 0.18 0.17 0.36 1.00
L5−IT 0.17 0.18 0.21 0.17 0.11 0.11 0.16 0.09 0.20 1.00
L5−ET 0.24 0.20 0.22 0.13 0.20 0.16 0.15 0.25 0.24 1.00
L4−IT 0.13 0.12 0.17 0.13 0.18 0.17 0.08 0.10 0.12 1.00
L23−IT 0.11 0.14 0.19 0.14 0.09 0.08 0.08 0.11 0.16 1.00
HIP−Misc2 0.34 0.21 0.11 0.13 0.16 0.04 0.10 0.13 0.18 1.00
HIP−Misc1 0.10 0.11 0.14 0.12 0.16 0.15 0.11 0.15 0.18 1.00
DG 0.09 0.11 0.04 0.18 0.11 0.07 0.09 0.20 0.25 1.00
CA3 0.37 0.22 0.07 0.12 0.17 0.08 0.17 0.12 0.42 1.00
CA1 0.30 0.16 0.19 0.14 0.18 0.15 0.21 0.16 0.20 1.00
Amy−Exc 0.40 0.17 0.30 0.18 0.26 0.19 0.12 0.18 0.28 1.00
Vip 0.16 0.27 0.39 0.42 0.40 0.42 0.54 0.52 0.66 1.00
THM−MB 0.30 0.45 0.61 0.55 0.54 0.51 0.44 0.45 0.78 1.00
THM−Inh 1.00 0.57 0.43 0.40 0.43 0.30 0.30 0.47 0.50 1.00
THM−Exc 0.48 0.24 0.18 0.24 0.26 0.14 0.16 0.20 0.34 1.00
SubCtx−Cplx 0.18 0.16 0.15 0.13 0.07 0.12 0.08 0.16 0.14 1.00
Sst 0.34 0.20 0.15 0.14 0.13 0.13 0.11 0.15 0.20 1.00
Sncg 0.40 0.35 0.33 0.22 0.16 0.23 0.16 0.18 0.34 1.00
Pvalb−ChC 0.22 0.37 0.28 0.27 0.22 0.16 0.16 0.20 0.30 1.00
Pvalb 0.56 0.38 0.24 0.15 0.12 0.15 0.14 0.20 0.24 1.00
PN 0.42 0.48 0.30 0.24 0.24 0.23 0.32 0.25 0.40 1.00
MSN−D2 0.34 0.17 0.16 0.15 0.12 0.13 0.08 0.13 0.30 1.00
MSN−D1 0.30 0.28 0.24 0.14 0.20 0.22 0.26 0.23 0.30 1.00
Lamp5−Lhx6 0.16 0.28 0.15 0.17 0.14 0.21 0.35 0.19 0.47 1.00
Lamp5 0.33 0.32 0.29 0.20 0.24 0.21 0.25 0.15 0.35 1.00
Foxp2 0.24 0.16 0.18 0.09 0.05 0.05 0.10 0.09 0.21 1.00
Chd7 0.25 0.24 0.28 0.22 0.19 0.18 0.19 0.26 0.31 1.00
CB 0.28 0.16 0.07 0.17 0.12 0.17 0.18 0.24 0.32 1.00
0.0−0.1 0.1−0.2 0.2−0.3 0.3−0.4 0.4−0.5 0.5−0.6 0.6−0.7 0.7−0.8 0.8−0.9 0.9−1.0
Percent
2e−05 4e−05 6e−05 8e−05
PRERC−Glutamatergic neurons from piriform cortex and e
L6b 1.64 1.00 0.87 0.78 0.72 0.67 0.64 0.63 0.63 1.00
L6−IT−Car3 2.36 1.38 1.11 0.98 0.90 0.84 0.79 0.76 0.73 1.00
L6−IT 1.79 1.08 0.91 0.82 0.75 0.70 0.68 0.64 0.63 1.00
L6−CT 1.44 0.97 0.84 0.79 0.74 0.70 0.66 0.65 0.69 1.00
L56−NP 1.74 1.22 1.08 1.01 0.96 0.90 0.86 0.83 0.83 1.00
L5−IT 1.98 1.07 0.94 0.85 0.78 0.74 0.69 0.68 0.67 1.00
L5−ET 1.82 1.09 0.96 0.87 0.82 0.78 0.73 0.74 0.72 1.00
L4−IT 1.35 0.83 0.71 0.66 0.62 0.59 0.58 0.57 0.58 1.00
L23−IT 1.21 0.74 0.64 0.59 0.54 0.51 0.50 0.48 0.51 1.00
HIP−Misc2 1.73 1.04 0.88 0.80 0.76 0.73 0.69 0.67 0.67 1.00
HIP−Misc1 0.97 0.73 0.68 0.64 0.61 0.59 0.59 0.57 0.57 1.00
DG 0.98 0.67 0.61 0.59 0.56 0.56 0.56 0.57 0.60 1.00
CA3 1.32 0.86 0.75 0.73 0.68 0.64 0.64 0.64 0.67 1.00
CA1 1.73 1.03 0.85 0.79 0.73 0.69 0.67 0.64 0.66 1.00
Amy−Exc 1.89 1.13 1.00 0.90 0.84 0.79 0.76 0.76 0.74 1.00
Vip 1.62 1.31 1.21 1.12 1.07 1.04 1.00 0.97 0.95 1.00
THM−MB 1.94 1.47 1.34 1.26 1.18 1.11 1.06 1.00 0.97 1.00
THM−Inh 1.89 1.49 1.33 1.23 1.15 1.09 1.05 1.03 0.97 1.00
THM−Exc 1.48 1.06 0.89 0.83 0.78 0.77 0.74 0.72 0.73 1.00
SubCtx−Cplx 0.80 0.67 0.64 0.62 0.62 0.61 0.60 0.61 0.62 1.00
Sst 1.40 1.07 0.98 0.91 0.88 0.85 0.82 0.80 0.80 1.00
Sncg 1.51 1.19 1.08 1.03 0.98 0.92 0.87 0.84 0.82 1.00
Pvalb−ChC 1.57 1.09 0.98 0.90 0.85 0.79 0.77 0.74 0.72 1.00
Pvalb 1.83 1.35 1.16 1.05 0.96 0.92 0.86 0.82 0.80 1.00
PN 2.36 1.61 1.39 1.22 1.13 1.04 0.97 0.89 0.89 1.00
MSN−D2 1.44 1.00 0.92 0.86 0.81 0.79 0.77 0.75 0.76 1.00
MSN−D1 1.68 1.11 0.97 0.88 0.84 0.80 0.77 0.74 0.74 1.00
Lamp5−Lhx6 1.47 1.11 1.00 0.94 0.88 0.84 0.80 0.77 0.77 1.00
Lamp5 1.44 1.16 1.07 1.03 0.96 0.91 0.88 0.88 0.84 1.00
Foxp2 1.11 0.78 0.71 0.67 0.63 0.60 0.59 0.57 0.60 1.00
Chd7 1.57 1.19 1.09 1.01 0.95 0.91 0.87 0.84 0.83 1.00
CB 1.09 0.99 0.94 0.90 0.87 0.83 0.84 0.79 0.84 1.00
0.0−0.1 0.1−0.2 0.2−0.3 0.3−0.4 0.4−0.5 0.5−0.6 0.6−0.7 0.7−0.8 0.8−0.9 0.9−1.0
Percent
0.0075 0.0100 0.0125 0.0150

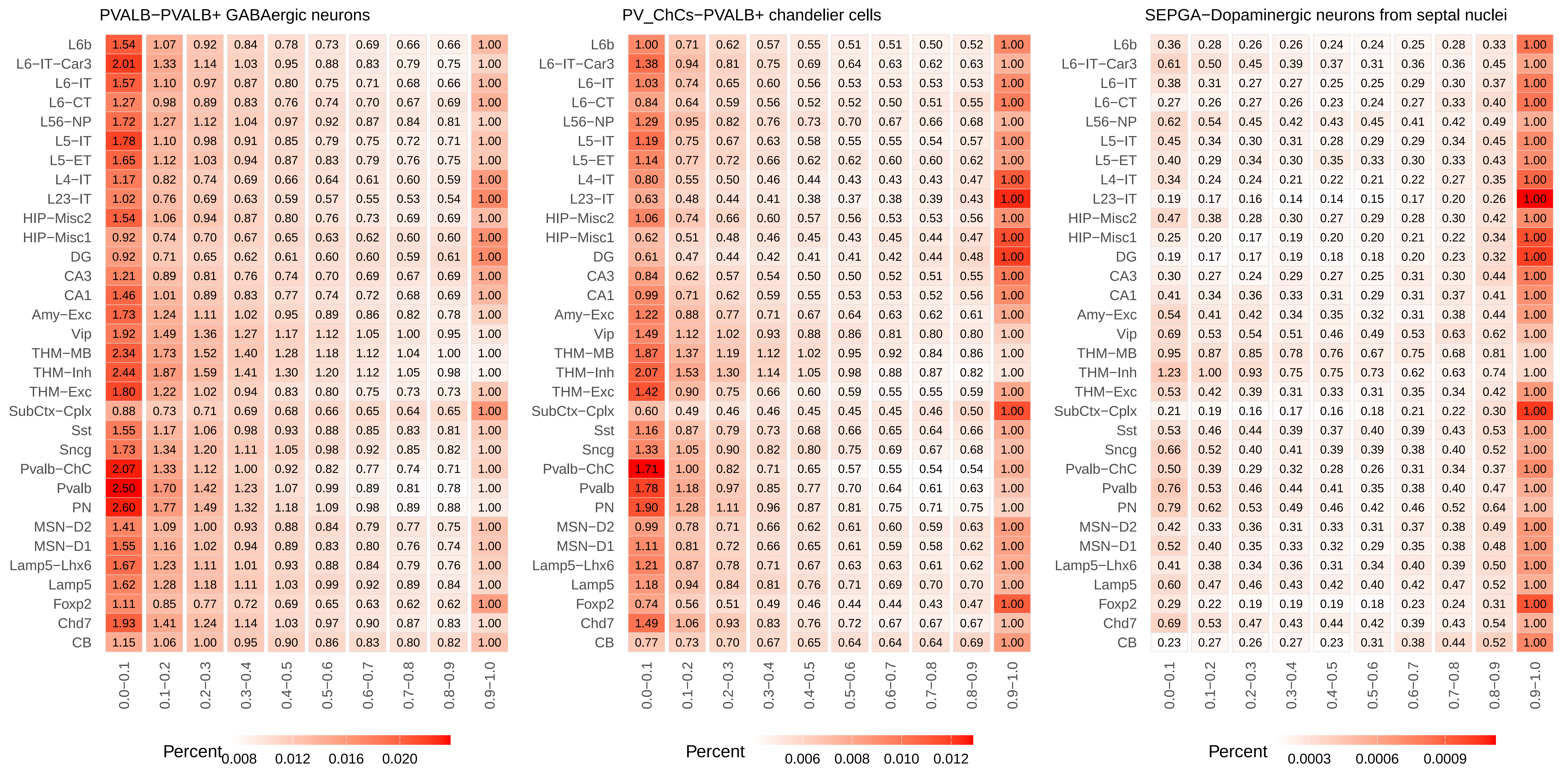
PVALB−PVALB+ GABAergic neurons
L6b 1.54 1.07 0.92 0.84 0.78 0.73 0.69 0.66 0.66 1.00
L6−IT−Car3 2.01 1.33 1.14 1.03 0.95 0.88 0.83 0.79 0.75 1.00
L6−IT 1.57 1.10 0.97 0.87 0.80 0.75 0.71 0.68 0.66 1.00
L6−CT 1.27 0.98 0.89 0.83 0.76 0.74 0.70 0.67 0.69 1.00
L56−NP 1.72 1.27 1.12 1.04 0.97 0.92 0.87 0.84 0.81 1.00
L5−IT 1.78 1.10 0.98 0.91 0.85 0.79 0.75 0.72 0.71 1.00
L5−ET 1.65 1.12 1.03 0.94 0.87 0.83 0.79 0.76 0.75 1.00
L4−IT 1.17 0.82 0.74 0.69 0.66 0.64 0.61 0.60 0.59 1.00
L23−IT 1.02 0.76 0.69 0.63 0.59 0.57 0.55 0.53 0.54 1.00
HIP−Misc2 1.54 1.06 0.94 0.87 0.80 0.76 0.73 0.69 0.69 1.00
HIP−Misc1 0.92 0.74 0.70 0.67 0.65 0.63 0.62 0.60 0.60 1.00
DG 0.92 0.71 0.65 0.62 0.61 0.60 0.60 0.59 0.61 1.00
CA3 1.21 0.89 0.81 0.76 0.74 0.70 0.69 0.67 0.69 1.00
CA1 1.46 1.01 0.89 0.83 0.77 0.74 0.72 0.68 0.69 1.00
Amy−Exc 1.73 1.24 1.11 1.02 0.95 0.89 0.86 0.82 0.78 1.00
Vip 1.92 1.49 1.36 1.27 1.17 1.12 1.05 1.00 0.95 1.00
THM−MB 2.34 1.73 1.52 1.40 1.28 1.18 1.12 1.04 1.00 1.00
THM−Inh 2.44 1.87 1.59 1.41 1.30 1.20 1.12 1.05 0.98 1.00
THM−Exc 1.80 1.22 1.02 0.94 0.83 0.80 0.75 0.73 0.73 1.00
SubCtx−Cplx 0.88 0.73 0.71 0.69 0.68 0.66 0.65 0.64 0.65 1.00
Sst 1.55 1.17 1.06 0.98 0.93 0.88 0.85 0.83 0.81 1.00
Sncg 1.73 1.34 1.20 1.11 1.05 0.98 0.92 0.85 0.82 1.00
Pvalb−ChC 2.07 1.33 1.12 1.00 0.92 0.82 0.77 0.74 0.71 1.00
Pvalb 2.50 1.70 1.42 1.23 1.07 0.99 0.89 0.81 0.78 1.00
PN 2.60 1.77 1.49 1.32 1.18 1.09 0.98 0.89 0.88 1.00
MSN−D2 1.41 1.09 1.00 0.93 0.88 0.84 0.79 0.77 0.75 1.00
MSN−D1 1.55 1.16 1.02 0.94 0.89 0.83 0.80 0.76 0.74 1.00
Lamp5−Lhx6 1.67 1.23 1.11 1.01 0.93 0.88 0.84 0.79 0.76 1.00
Lamp5 1.62 1.28 1.18 1.11 1.03 0.99 0.92 0.89 0.84 1.00
Foxp2 1.11 0.85 0.77 0.72 0.69 0.65 0.63 0.62 0.62 1.00
Chd7 1.93 1.41 1.24 1.14 1.03 0.97 0.90 0.87 0.83 1.00
CB 1.15 1.06 1.00 0.95 0.90 0.86 0.83 0.80 0.82 1.00
0.0−0.1 0.1−0.2 0.2−0.3 0.3−0.4 0.4−0.5 0.5−0.6 0.6−0.7 0.7−0.8 0.8−0.9 0.9−1.0
Percent 0.008 0.012 0.016 0.020
PV_ChCs−PVALB+ chandelier cells
L6b 1.00 0.71 0.62 0.57 0.55 0.51 0.51 0.50 0.52 1.00
L6−IT−Car3 1.38 0.94 0.81 0.75 0.69 0.64 0.63 0.62 0.63 1.00
L6−IT 1.03 0.74 0.65 0.60 0.56 0.53 0.53 0.53 0.53 1.00
L6−CT 0.84 0.64 0.59 0.56 0.52 0.52 0.50 0.51 0.55 1.00
L56−NP 1.29 0.95 0.82 0.76 0.73 0.70 0.67 0.66 0.68 1.00
L5−IT 1.19 0.75 0.67 0.63 0.58 0.55 0.55 0.54 0.57 1.00
L5−ET 1.14 0.77 0.72 0.66 0.62 0.62 0.60 0.60 0.62 1.00
L4−IT 0.80 0.55 0.50 0.46 0.44 0.43 0.43 0.43 0.47 1.00
L23−IT 0.63 0.48 0.44 0.41 0.38 0.37 0.38 0.39 0.43 1.00
HIP−Misc2 1.06 0.74 0.66 0.60 0.57 0.56 0.53 0.53 0.56 1.00
HIP−Misc1 0.62 0.51 0.48 0.46 0.45 0.43 0.45 0.44 0.47 1.00
DG 0.61 0.47 0.44 0.42 0.41 0.41 0.42 0.44 0.48 1.00
CA3 0.84 0.62 0.57 0.54 0.50 0.50 0.52 0.51 0.55 1.00
CA1 0.99 0.71 0.62 0.59 0.55 0.53 0.53 0.52 0.56 1.00
Amy−Exc 1.22 0.88 0.77 0.71 0.67 0.64 0.63 0.62 0.61 1.00
Vip 1.49 1.12 1.02 0.93 0.88 0.86 0.81 0.80 0.80 1.00
THM−MB 1.87 1.37 1.19 1.12 1.02 0.95 0.92 0.84 0.86 1.00
THM−Inh 2.07 1.53 1.30 1.14 1.05 0.98 0.88 0.87 0.82 1.00
THM−Exc 1.42 0.90 0.75 0.66 0.60 0.59 0.55 0.55 0.59 1.00
SubCtx−Cplx 0.60 0.49 0.46 0.46 0.45 0.45 0.45 0.46 0.50 1.00
Sst 1.16 0.87 0.79 0.73 0.68 0.66 0.65 0.64 0.66 1.00
Sncg 1.33 1.05 0.90 0.82 0.80 0.75 0.69 0.67 0.68 1.00
Pvalb−ChC 1.71 1.00 0.82 0.71 0.65 0.57 0.55 0.54 0.54 1.00
Pvalb 1.78 1.18 0.97 0.85 0.77 0.70 0.64 0.61 0.63 1.00
PN 1.90 1.28 1.11 0.96 0.87 0.81 0.75 0.71 0.75 1.00
MSN−D2 0.99 0.78 0.71 0.66 0.62 0.61 0.60 0.59 0.63 1.00
MSN−D1 1.11 0.81 0.72 0.66 0.65 0.61 0.59 0.58 0.62 1.00
Lamp5−Lhx6 1.21 0.87 0.78 0.71 0.67 0.63 0.63 0.61 0.62 1.00
Lamp5 1.18 0.94 0.84 0.81 0.76 0.71 0.69 0.70 0.70 1.00
Foxp2 0.74 0.56 0.51 0.49 0.46 0.44 0.44 0.43 0.47 1.00
Chd7 1.49 1.06 0.93 0.83 0.76 0.72 0.67 0.67 0.67 1.00
CB 0.77 0.73 0.70 0.67 0.65 0.64 0.64 0.64 0.69 1.00
0.0−0.1 0.1−0.2 0.2−0.3 0.3−0.4 0.4−0.5 0.5−0.6 0.6−0.7 0.7−0.8 0.8−0.9 0.9−1.0
Percent 0.006 0.008 0.010 0.012
SEPGA−Dopaminergic neurons from septal nuclei
L6b 0.36 0.28 0.26 0.26 0.24 0.24 0.25 0.28 0.33 1.00
L6−IT−Car3 0.61 0.50 0.45 0.39 0.37 0.31 0.36 0.36 0.45 1.00
L6−IT 0.38 0.31 0.27 0.27 0.25 0.25 0.29 0.30 0.37 1.00
L6−CT 0.27 0.26 0.27 0.26 0.23 0.24 0.27 0.33 0.40 1.00
L56−NP 0.62 0.54 0.45 0.42 0.43 0.45 0.41 0.42 0.49 1.00
L5−IT 0.45 0.34 0.30 0.31 0.28 0.29 0.29 0.34 0.45 1.00
L5−ET 0.40 0.29 0.34 0.30 0.35 0.33 0.30 0.33 0.43 1.00
L4−IT 0.34 0.24 0.24 0.21 0.22 0.21 0.22 0.27 0.35 1.00
L23−IT 0.19 0.17 0.16 0.14 0.14 0.15 0.17 0.20 0.26 1.00
HIP−Misc2 0.47 0.38 0.28 0.30 0.27 0.29 0.28 0.30 0.42 1.00
HIP−Misc1 0.25 0.20 0.17 0.19 0.20 0.20 0.21 0.22 0.34 1.00
DG 0.19 0.17 0.17 0.19 0.18 0.18 0.20 0.23 0.32 1.00
CA3 0.30 0.27 0.24 0.29 0.27 0.25 0.31 0.30 0.44 1.00
CA1 0.41 0.34 0.36 0.33 0.31 0.29 0.31 0.37 0.41 1.00
Amy−Exc 0.54 0.41 0.42 0.34 0.35 0.32 0.31 0.38 0.44 1.00
Vip 0.69 0.53 0.54 0.51 0.46 0.49 0.53 0.63 0.62 1.00
THM−MB 0.95 0.87 0.85 0.78 0.76 0.67 0.75 0.68 0.81 1.00
THM−Inh 1.23 1.00 0.93 0.75 0.75 0.73 0.62 0.63 0.74 1.00
THM−Exc 0.53 0.42 0.39 0.31 0.33 0.31 0.35 0.34 0.42 1.00
SubCtx−Cplx 0.21 0.19 0.16 0.17 0.16 0.18 0.21 0.22 0.30 1.00
Sst 0.53 0.46 0.44 0.39 0.37 0.40 0.39 0.43 0.53 1.00
Sncg 0.66 0.52 0.40 0.41 0.39 0.39 0.38 0.40 0.52 1.00
Pvalb−ChC 0.50 0.39 0.29 0.32 0.28 0.26 0.31 0.34 0.37 1.00
Pvalb 0.76 0.53 0.46 0.44 0.41 0.35 0.38 0.40 0.47 1.00
PN 0.79 0.62 0.53 0.49 0.46 0.42 0.46 0.52 0.64 1.00
MSN−D2 0.42 0.33 0.36 0.31 0.33 0.31 0.37 0.38 0.49 1.00
MSN−D1 0.52 0.40 0.35 0.33 0.32 0.29 0.35 0.38 0.48 1.00
Lamp5−Lhx6 0.41 0.38 0.34 0.36 0.31 0.34 0.40 0.39 0.50 1.00
Lamp5 0.60 0.47 0.46 0.43 0.42 0.40 0.42 0.47 0.52 1.00
Foxp2 0.29 0.22 0.19 0.19 0.19 0.18 0.23 0.24 0.31 1.00
Chd7 0.69 0.53 0.47 0.43 0.44 0.42 0.39 0.43 0.54 1.00
CB 0.23 0.27 0.26 0.27 0.23 0.31 0.38 0.44 0.52 1.00
0.0−0.1 0.1−0.2 0.2−0.3 0.3−0.4 0.4−0.5 0.5−0.6 0.6−0.7 0.7−0.8 0.8−0.9 0.9−1.0
Percent 0.0003 0.0006 0.0009

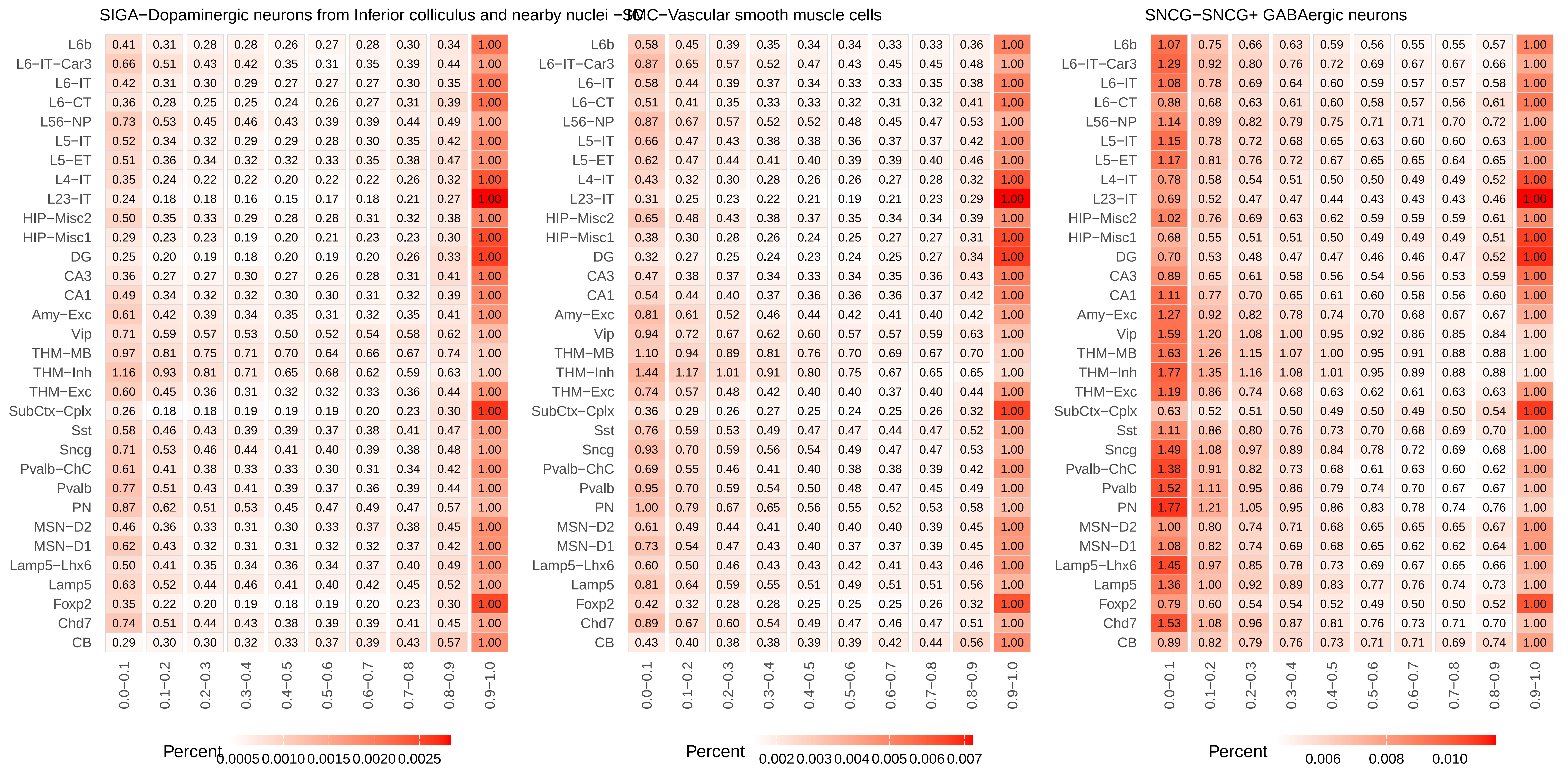
SIGA−Dopaminergic neurons from Inferior colliculus and nearby nuclei
SMC−Vascular smooth muscle cells
SNCG−SNCG+ GABAergic neurons
0.0−0.1 0.1−0.2 0.2−0.3 0.3−0.4 0.4−0.5 0.5−0.6 0.6−0.7 0.7−0.8 0.8−0.9 0.9−1.0
L6b 0.41 0.31 0.28 0.28 0.26 0.27 0.28 0.30 0.34 1.00
L6−IT−Car3 0.66 0.51 0.43 0.42 0.35 0.31 0.35 0.39 0.44 1.00
L6−IT 0.42 0.31 0.30 0.29 0.27 0.27 0.27 0.30 0.35 1.00
L6−CT 0.36 0.28 0.25 0.25 0.24 0.26 0.27 0.31 0.39 1.00
L56−NP 0.73 0.53 0.45 0.46 0.43 0.39 0.39 0.44 0.49 1.00
L5−IT 0.52 0.34 0.32 0.29 0.29 0.28 0.30 0.35 0.42 1.00
L5−ET 0.51 0.36 0.34 0.32 0.32 0.33 0.35 0.38 0.47 1.00
L4−IT 0.35 0.24 0.22 0.22 0.20 0.22 0.22 0.26 0.32 1.00
L23−IT 0.24 0.18 0.18 0.16 0.15 0.17 0.18 0.21 0.27 1.00
HIP−Misc2 0.50 0.35 0.33 0.29 0.28 0.28 0.31 0.32 0.38 1.00
HIP−Misc1 0.29 0.23 0.23 0.19 0.20 0.21 0.23 0.23 0.30 1.00
DG 0.25 0.20 0.19 0.18 0.20 0.19 0.20 0.26 0.33 1.00
CA3 0.36 0.27 0.27 0.30 0.27 0.26 0.28 0.31 0.41 1.00
CA1 0.49 0.34 0.32 0.32 0.30 0.30 0.31 0.32 0.39 1.00
Amy−Exc 0.61 0.42 0.39 0.34 0.35 0.31 0.32 0.35 0.41 1.00
Vip 0.71 0.59 0.57 0.53 0.50 0.52 0.54 0.58 0.62 1.00
THM−MB 0.97 0.81 0.75 0.71 0.70 0.64 0.66 0.67 0.74 1.00
THM−Inh 1.16 0.93 0.81 0.71 0.65 0.68 0.62 0.59 0.63 1.00
THM−Exc 0.60 0.45 0.36 0.31 0.32 0.32 0.33 0.36 0.44 1.00
SubCtx−Cplx 0.26 0.18 0.18 0.19 0.19 0.19 0.20 0.23 0.30 1.00
Sst 0.58 0.46 0.43 0.39 0.39 0.37 0.38 0.41 0.47 1.00
Sncg 0.71 0.53 0.46 0.44 0.41 0.40 0.39 0.38 0.48 1.00
Pvalb−ChC 0.61 0.41 0.38 0.33 0.33 0.30 0.31 0.34 0.42 1.00
Pvalb 0.77 0.51 0.43 0.41 0.39 0.37 0.36 0.39 0.44 1.00
PN 0.87 0.62 0.51 0.53 0.45 0.47 0.49 0.47 0.57 1.00
MSN−D2 0.46 0.36 0.33 0.31 0.30 0.33 0.37 0.38 0.45 1.00
MSN−D1 0.62 0.43 0.32 0.31 0.31 0.32 0.32 0.37 0.42 1.00
Lamp5−Lhx6 0.50 0.41 0.35 0.34 0.36 0.34 0.37 0.40 0.49 1.00
Lamp5 0.63 0.52 0.44 0.46 0.41 0.40 0.42 0.45 0.52 1.00
Foxp2 0.35 0.22 0.20 0.19 0.18 0.19 0.20 0.23 0.30 1.00
Chd7 0.74 0.51 0.44 0.43 0.38 0.39 0.39 0.41 0.45 1.00
CB 0.29 0.30 0.30 0.32 0.33 0.37 0.39 0.43 0.57 1.00
Percent 0.0005 0.0010 0.0015 0.0020 0.0025
L6b 0.58 0.45 0.39 0.35 0.34 0.34 0.33 0.33 0.36 1.00
L6−IT−Car3 0.87 0.65 0.57 0.52 0.47 0.43 0.45 0.45 0.48 1.00
L6−IT 0.58 0.44 0.39 0.37 0.34 0.33 0.33 0.35 0.38 1.00
L6−CT 0.51 0.41 0.35 0.33 0.33 0.32 0.31 0.32 0.41 1.00
L56−NP 0.87 0.67 0.57 0.52 0.52 0.48 0.45 0.47 0.53 1.00
L5−IT 0.66 0.47 0.43 0.38 0.38 0.36 0.37 0.37 0.42 1.00
L5−ET 0.62 0.47 0.44 0.41 0.40 0.39 0.39 0.40 0.46 1.00
L4−IT 0.43 0.32 0.30 0.28 0.26 0.26 0.27 0.28 0.32 1.00
L23−IT 0.31 0.25 0.23 0.22 0.21 0.19 0.21 0.23 0.29 1.00
HIP−Misc2 0.65 0.48 0.43 0.38 0.37 0.35 0.34 0.34 0.39 1.00
HIP−Misc1 0.38 0.30 0.28 0.26 0.24 0.25 0.27 0.27 0.31 1.00
DG 0.32 0.27 0.25 0.24 0.23 0.24 0.25 0.27 0.34 1.00
CA3 0.47 0.38 0.37 0.34 0.33 0.34 0.35 0.36 0.43 1.00
CA1 0.54 0.44 0.40 0.37 0.36 0.36 0.36 0.37 0.42 1.00
Amy−Exc 0.81 0.61 0.52 0.46 0.44 0.42 0.41 0.40 0.42 1.00
Vip 0.94 0.72 0.67 0.62 0.60 0.57 0.57 0.59 0.63 1.00
THM−MB 1.10 0.94 0.89 0.81 0.76 0.70 0.69 0.67 0.70 1.00
THM−Inh 1.44 1.17 1.01 0.91 0.80 0.75 0.67 0.65 0.65 1.00
THM−Exc 0.74 0.57 0.48 0.42 0.40 0.40 0.37 0.40 0.44 1.00
SubCtx−Cplx 0.36 0.29 0.26 0.27 0.25 0.24 0.25 0.26 0.32 1.00
Sst 0.76 0.59 0.53 0.49 0.47 0.47 0.44 0.47 0.52 1.00
Sncg 0.93 0.70 0.59 0.56 0.54 0.49 0.47 0.47 0.53 1.00
Pvalb−ChC 0.69 0.55 0.46 0.41 0.40 0.38 0.38 0.39 0.42 1.00
Pvalb 0.95 0.70 0.59 0.54 0.50 0.48 0.47 0.45 0.49 1.00
PN 1.00 0.79 0.67 0.65 0.56 0.55 0.52 0.53 0.58 1.00
MSN−D2 0.61 0.49 0.44 0.41 0.40 0.40 0.40 0.39 0.45 1.00
MSN−D1 0.73 0.54 0.47 0.43 0.40 0.37 0.37 0.39 0.45 1.00
Lamp5−Lhx6 0.60 0.50 0.46 0.43 0.43 0.42 0.41 0.43 0.46 1.00
Lamp5 0.81 0.64 0.59 0.55 0.51 0.49 0.51 0.51 0.56 1.00
Foxp2 0.42 0.32 0.28 0.28 0.25 0.25 0.25 0.26 0.32 1.00
Chd7 0.89 0.67 0.60 0.54 0.49 0.47 0.46 0.47 0.51 1.00
CB 0.43 0.40 0.38 0.38 0.39 0.39 0.42 0.44 0.56 1.00
Percent 0.002 0.003 0.004 0.005 0.006 0.007
L6b 1.07 0.75 0.66 0.63 0.59 0.56 0.55 0.55 0.57 1.00
L6−IT−Car3 1.29 0.92 0.80 0.76 0.72 0.69 0.67 0.67 0.66 1.00
L6−IT 1.08 0.78 0.69 0.64 0.60 0.59 0.57 0.57 0.58 1.00
L6−CT 0.88 0.68 0.63 0.61 0.60 0.58 0.57 0.56 0.61 1.00
L56−NP 1.14 0.89 0.82 0.79 0.75 0.71 0.71 0.70 0.72 1.00
L5−IT 1.15 0.78 0.72 0.68 0.65 0.63 0.60 0.60 0.63 1.00
L5−ET 1.17 0.81 0.76 0.72 0.67 0.65 0.65 0.64 0.65 1.00
L4−IT 0.78 0.58 0.54 0.51 0.50 0.50 0.49 0.49 0.52 1.00
L23−IT 0.69 0.52 0.47 0.47 0.44 0.43 0.43 0.43 0.46 1.00
HIP−Misc2 1.02 0.76 0.69 0.63 0.62 0.59 0.59 0.59 0.61 1.00
HIP−Misc1 0.68 0.55 0.51 0.51 0.50 0.49 0.49 0.49 0.51 1.00
DG 0.70 0.53 0.48 0.47 0.47 0.46 0.46 0.47 0.52 1.00
CA3 0.89 0.65 0.61 0.58 0.56 0.54 0.56 0.53 0.59 1.00
CA1 1.11 0.77 0.70 0.65 0.61 0.60 0.58 0.56 0.60 1.00
Amy−Exc 1.27 0.92 0.82 0.78 0.74 0.70 0.68 0.67 0.67 1.00
Vip 1.59 1.20 1.08 1.00 0.95 0.92 0.86 0.85 0.84 1.00
THM−MB 1.63 1.26 1.15 1.07 1.00 0.95 0.91 0.88 0.88 1.00
THM−Inh 1.77 1.35 1.16 1.08 1.01 0.95 0.89 0.88 0.88 1.00
THM−Exc 1.19 0.86 0.74 0.68 0.63 0.62 0.61 0.63 0.63 1.00
SubCtx−Cplx 0.63 0.52 0.51 0.50 0.49 0.50 0.49 0.50 0.54 1.00
Sst 1.11 0.86 0.80 0.76 0.73 0.70 0.68 0.69 0.70 1.00
Sncg 1.49 1.08 0.97 0.89 0.84 0.78 0.72 0.69 0.68 1.00
Pvalb−ChC 1.38 0.91 0.82 0.73 0.68 0.61 0.63 0.60 0.62 1.00
Pvalb 1.52 1.11 0.95 0.86 0.79 0.74 0.70 0.67 0.67 1.00
PN 1.77 1.21 1.05 0.95 0.86 0.83 0.78 0.74 0.76 1.00
MSN−D2 1.00 0.80 0.74 0.71 0.68 0.65 0.65 0.65 0.67 1.00
MSN−D1 1.08 0.82 0.74 0.69 0.68 0.65 0.62 0.62 0.64 1.00
Lamp5−Lhx6 1.45 0.97 0.85 0.78 0.73 0.69 0.67 0.65 0.66 1.00
Lamp5 1.36 1.00 0.92 0.89 0.83 0.77 0.76 0.74 0.73 1.00
Foxp2 0.79 0.60 0.54 0.54 0.52 0.49 0.50 0.50 0.52 1.00
Chd7 1.53 1.08 0.96 0.87 0.81 0.76 0.73 0.71 0.70 1.00
CB 0.89 0.82 0.79 0.76 0.73 0.71 0.71 0.69 0.74 1.00
Percent 0.006 0.008 0.010

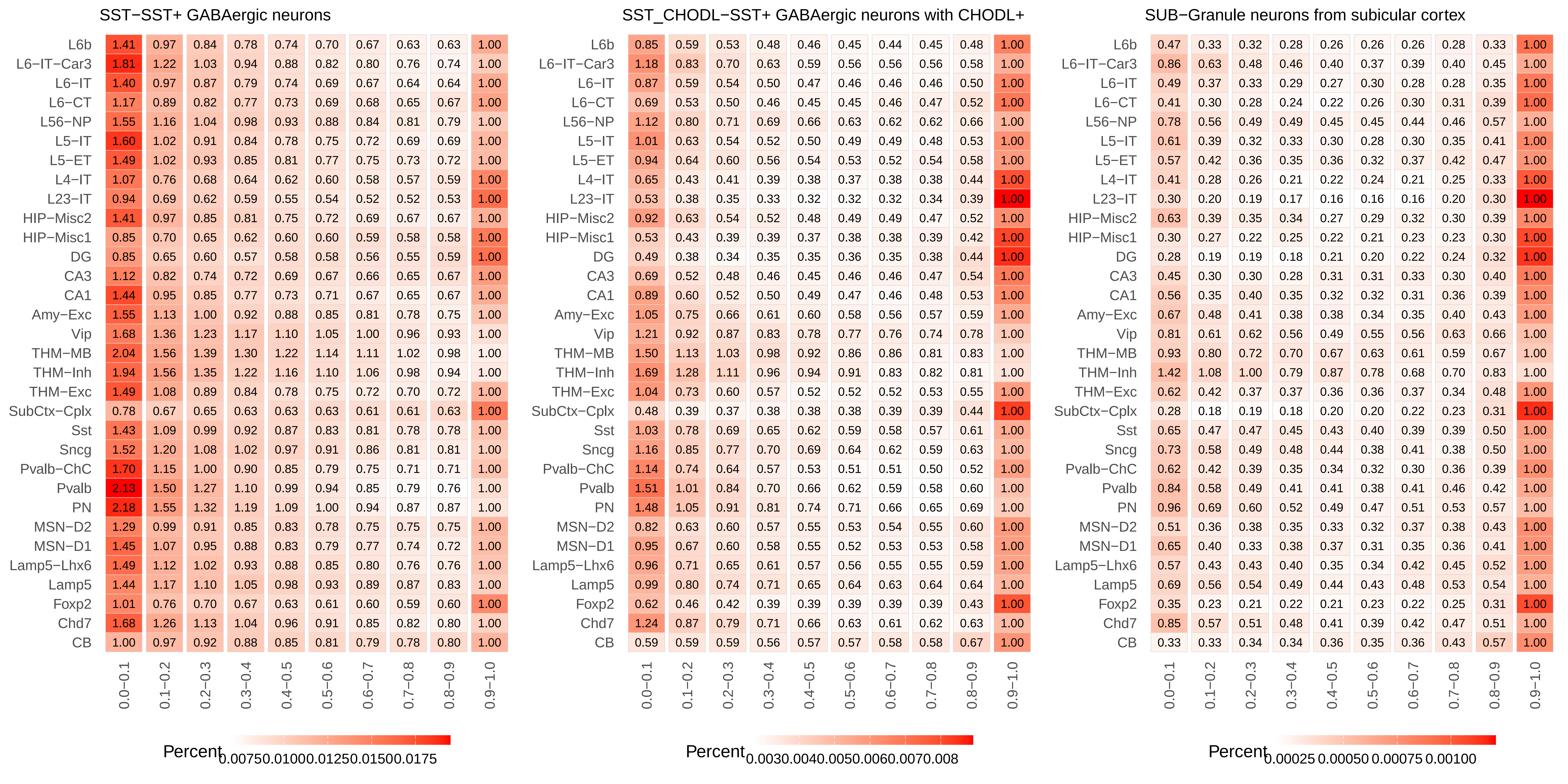
SST−SST+ GABAergic neurons
SST_CHODL−SST+ GABAergic neurons with CHODL+
SUB−Granule neurons from subicular cortex
L6b
L6−IT−Car3
L6−IT
L6−CT
L56−NP
L5−IT
L5−ET
L4−IT
L23−IT
HIP−Misc2
HIP−Misc1
DG
CA3
CA1
Amy−Exc
Vip
THM−MB
THM−Inh
THM−Exc
SubCtx−Cplx
Sst
Sncg
Pvalb−ChC
Pvalb
PN
MSN−D2
MSN−D1
Lamp5−Lhx6
Lamp5
Foxp2
Chd7
CB
0.0−0.1
0.1−0.2
0.2−0.3
0.3−0.4
0.4−0.5
0.5−0.6
0.6−0.7
0.7−0.8
0.8−0.9
0.9−1.0
Percent
0.0075 0.0100 0.0125 0.0150 0.0175
Percent
0.003 0.004 0.005 0.006 0.007 0.008
Percent
0.00025 0.00050 0.00075 0.00100

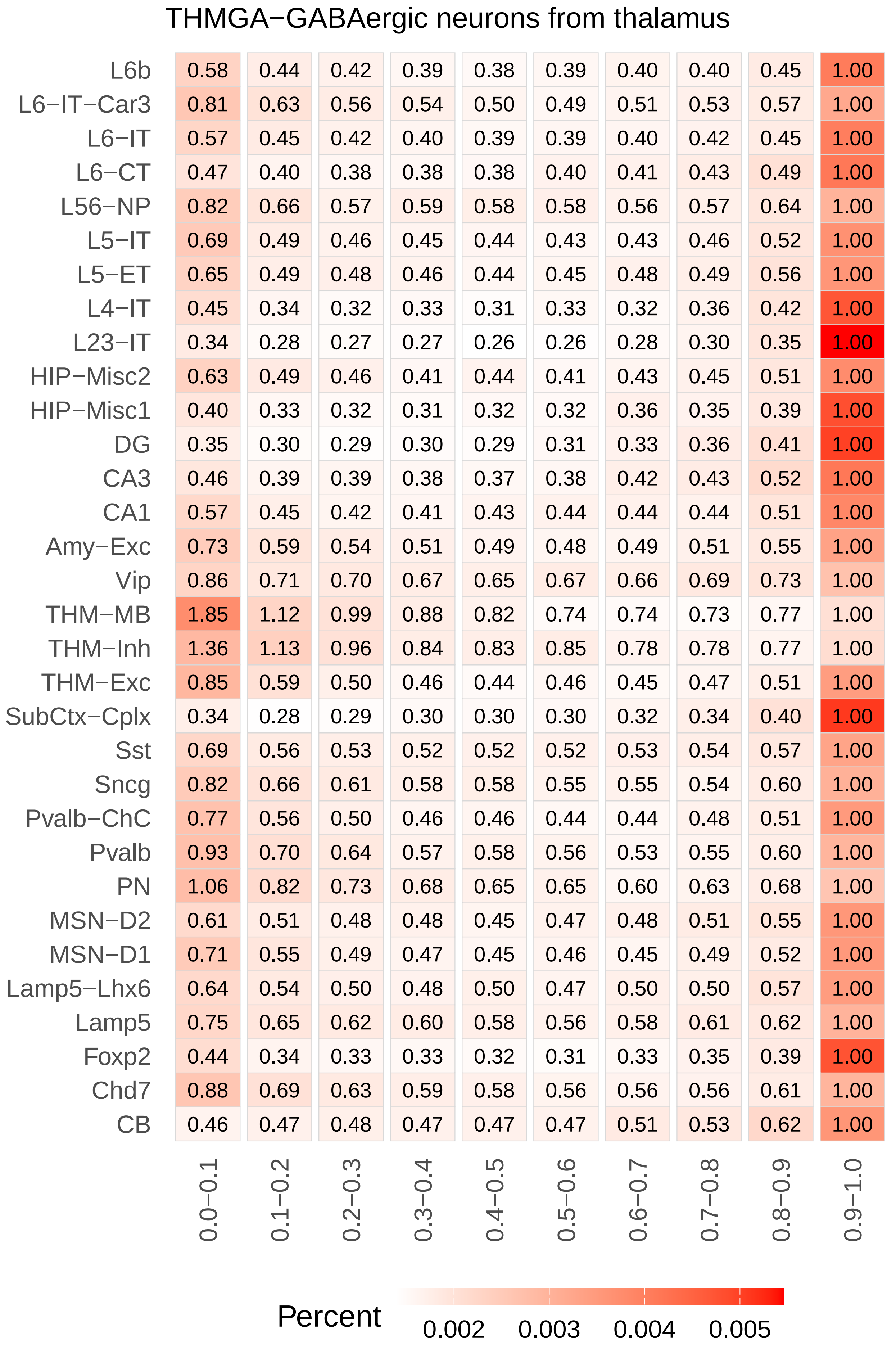
THMGA−GABAergic neurons from thalamus
L6b 0.58 0.44 0.42 0.39 0.38 0.39 0.40 0.40 0.45 1.00
L6−IT−Car3 0.81 0.63 0.56 0.54 0.50 0.49 0.51 0.53 0.57 1.00
L6−IT 0.57 0.45 0.42 0.40 0.39 0.39 0.40 0.42 0.45 1.00
L6−CT 0.47 0.40 0.38 0.38 0.38 0.40 0.41 0.43 0.49 1.00
L56−NP 0.82 0.66 0.57 0.59 0.58 0.58 0.56 0.57 0.64 1.00
L5−IT 0.69 0.49 0.46 0.45 0.44 0.43 0.43 0.46 0.52 1.00
L5−ET 0.65 0.49 0.48 0.46 0.44 0.45 0.48 0.49 0.56 1.00
L4−IT 0.45 0.34 0.32 0.33 0.31 0.33 0.32 0.36 0.42 1.00
L23−IT 0.34 0.28 0.27 0.27 0.26 0.26 0.28 0.30 0.35 1.00
HIP−Misc2 0.63 0.49 0.46 0.41 0.44 0.41 0.43 0.45 0.51 1.00
HIP−Misc1 0.40 0.33 0.32 0.31 0.32 0.32 0.36 0.35 0.39 1.00
DG 0.35 0.30 0.29 0.30 0.29 0.31 0.33 0.36 0.41 1.00
CA3 0.46 0.39 0.39 0.38 0.37 0.38 0.42 0.43 0.52 1.00
CA1 0.57 0.45 0.42 0.41 0.43 0.44 0.44 0.44 0.51 1.00
Amy−Exc 0.73 0.59 0.54 0.51 0.49 0.48 0.49 0.51 0.55 1.00
Vip 0.86 0.71 0.70 0.67 0.65 0.67 0.66 0.69 0.73 1.00
THM−MB 1.85 1.12 0.99 0.88 0.82 0.74 0.74 0.73 0.77 1.00
THM−Inh 1.36 1.13 0.96 0.84 0.83 0.85 0.78 0.78 0.77 1.00
THM−Exc 0.85 0.59 0.50 0.46 0.44 0.46 0.45 0.47 0.51 1.00
SubCtx−Cplx 0.34 0.28 0.29 0.30 0.30 0.30 0.32 0.34 0.40 1.00
Sst 0.69 0.56 0.53 0.52 0.52 0.52 0.53 0.54 0.57 1.00
Sncg 0.82 0.66 0.61 0.58 0.58 0.55 0.55 0.54 0.60 1.00
Pvalb−ChC 0.77 0.56 0.50 0.46 0.46 0.44 0.44 0.48 0.51 1.00
Pvalb 0.93 0.70 0.64 0.57 0.58 0.56 0.53 0.55 0.60 1.00
PN 1.06 0.82 0.73 0.68 0.65 0.65 0.60 0.63 0.68 1.00
MSN−D2 0.61 0.51 0.48 0.48 0.45 0.47 0.48 0.51 0.55 1.00
MSN−D1 0.71 0.55 0.49 0.47 0.45 0.46 0.45 0.49 0.52 1.00
Lamp5−Lhx6 0.64 0.54 0.50 0.48 0.50 0.47 0.50 0.50 0.57 1.00
Lamp5 0.75 0.65 0.62 0.60 0.58 0.56 0.58 0.61 0.62 1.00
Foxp2 0.44 0.34 0.33 0.33 0.32 0.31 0.33 0.35 0.39 1.00
Chd7 0.88 0.69 0.63 0.59 0.58 0.56 0.56 0.56 0.61 1.00
CB 0.46 0.47 0.48 0.47 0.47 0.47 0.51 0.53 0.62 1.00
0.0−0.1 0.1−0.2 0.2−0.3 0.3−0.4 0.4−0.5 0.5−0.6 0.6−0.7 0.7−0.8 0.8−0.9 0.9−1.0
Percent 0.002 0.003 0.004 0.005

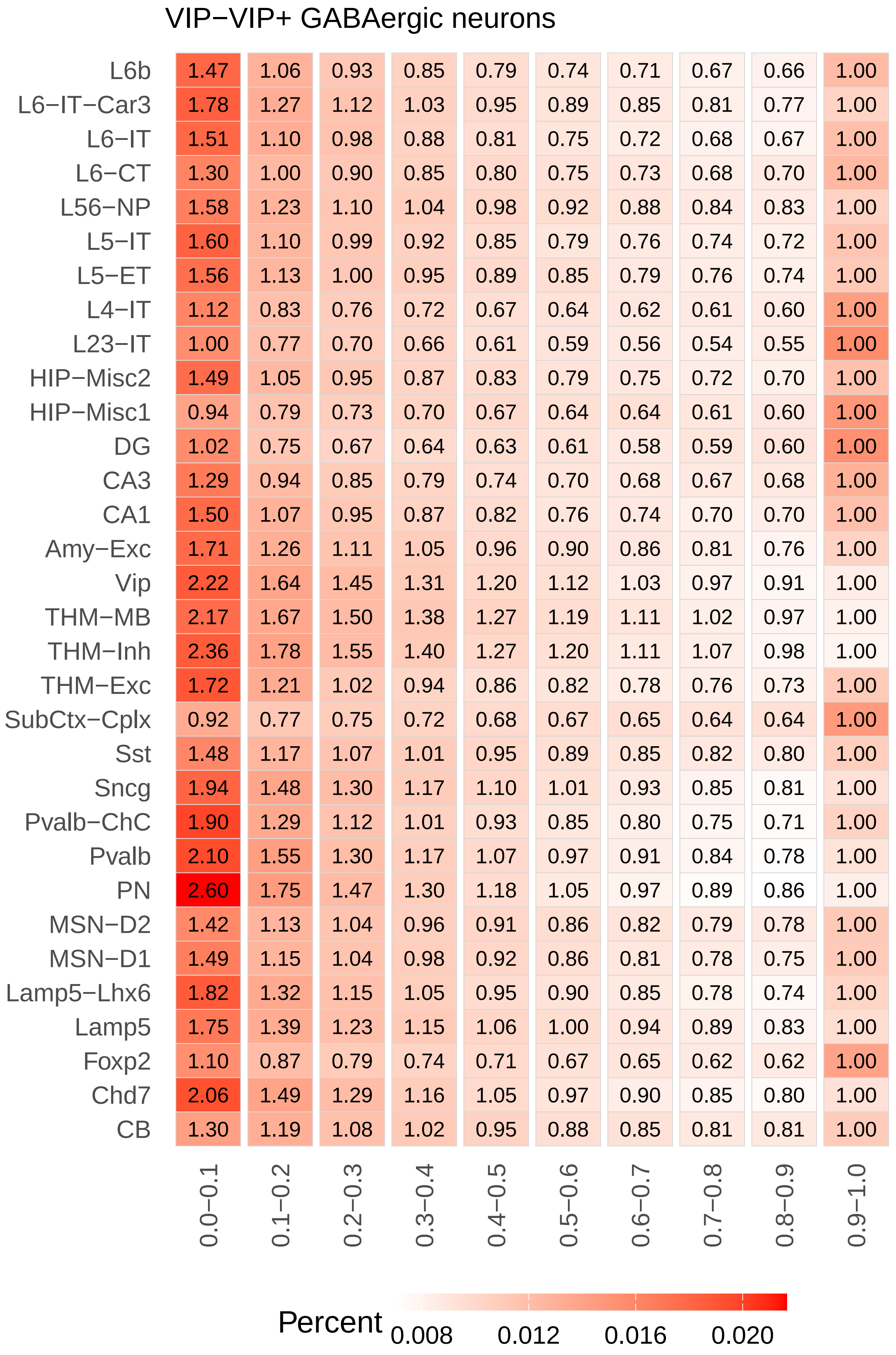
VIP−VIP+ GABAergic neurons
L6b 1.47 1.06 0.93 0.85 0.79 0.74 0.71 0.67 0.66 1.00
L6−IT−Car3 1.78 1.27 1.12 1.03 0.95 0.89 0.85 0.81 0.77 1.00
L6−IT 1.51 1.10 0.98 0.88 0.81 0.75 0.72 0.68 0.67 1.00
L6−CT 1.30 1.00 0.90 0.85 0.80 0.75 0.73 0.68 0.70 1.00
L56−NP 1.58 1.23 1.10 1.04 0.98 0.92 0.88 0.84 0.83 1.00
L5−IT 1.60 1.10 0.99 0.92 0.85 0.79 0.76 0.74 0.72 1.00
L5−ET 1.56 1.13 1.00 0.95 0.89 0.85 0.79 0.76 0.74 1.00
L4−IT 1.12 0.83 0.76 0.72 0.67 0.64 0.62 0.61 0.60 1.00
L23−IT 1.00 0.77 0.70 0.66 0.61 0.59 0.56 0.54 0.55 1.00
HIP−Misc2 1.49 1.05 0.95 0.87 0.83 0.79 0.75 0.72 0.70 1.00
HIP−Misc1 0.94 0.79 0.73 0.70 0.67 0.64 0.64 0.61 0.60 1.00
DG 1.02 0.75 0.67 0.64 0.63 0.61 0.58 0.59 0.60 1.00
CA3 1.29 0.94 0.85 0.79 0.74 0.70 0.68 0.67 0.68 1.00
CA1 1.50 1.07 0.95 0.87 0.82 0.76 0.74 0.70 0.70 1.00
Amy−Exc 1.71 1.26 1.11 1.05 0.96 0.90 0.86 0.81 0.76 1.00
Vip 2.22 1.64 1.45 1.31 1.20 1.12 1.03 0.97 0.91 1.00
THM−MB 2.17 1.67 1.50 1.38 1.27 1.19 1.11 1.02 0.97 1.00
THM−Inh 2.36 1.78 1.55 1.40 1.27 1.20 1.11 1.07 0.98 1.00
THM−Exc 1.72 1.21 1.02 0.94 0.86 0.82 0.78 0.76 0.73 1.00
SubCtx−Cplx 0.92 0.77 0.75 0.72 0.68 0.67 0.65 0.64 0.64 1.00
Sst 1.48 1.17 1.07 1.01 0.95 0.89 0.85 0.82 0.80 1.00
Sncg 1.94 1.48 1.30 1.17 1.10 1.01 0.93 0.85 0.81 1.00
Pvalb−ChC 1.90 1.29 1.12 1.01 0.93 0.85 0.80 0.75 0.71 1.00
Pvalb 2.10 1.55 1.30 1.17 1.07 0.97 0.91 0.84 0.78 1.00
PN 2.60 1.75 1.47 1.30 1.18 1.05 0.97 0.89 0.86 1.00
MSN−D2 1.42 1.13 1.04 0.96 0.91 0.86 0.82 0.79 0.78 1.00
MSN−D1 1.49 1.15 1.04 0.98 0.92 0.86 0.81 0.78 0.75 1.00
Lamp5−Lhx6 1.82 1.32 1.15 1.05 0.95 0.90 0.85 0.78 0.74 1.00
Lamp5 1.75 1.39 1.23 1.15 1.06 1.00 0.94 0.89 0.83 1.00
Foxp2 1.10 0.87 0.79 0.74 0.71 0.67 0.65 0.62 0.62 1.00
Chd7 2.06 1.49 1.29 1.16 1.05 0.97 0.90 0.85 0.80 1.00
CB 1.30 1.19 1.08 1.02 0.95 0.88 0.85 0.81 0.81 1.00
0.0−0.1 0.1−0.2 0.2−0.3 0.3−0.4 0.4−0.5 0.5−0.6 0.6−0.7 0.7−0.8 0.8−0.9 0.9−1.0
Percent 0.008 0.012 0.016 0.020
