## Supplementary Figure 5 for "Distinct cellular DNA methylation mechanisms underlie common and rare genetic risk for brain disorders"

### ACBGM−Bergmann glia

| | 0.0−0.1 | 0.1−0.2 | 0.2−0.3 | 0.3−0.4 | 0.4−0.5 | 0.5−0.6 | 0.6−0.7 | 0.7−0.8 | 0.8−0.9 | 0.9−1.0 |
|---|---|---|---|---|---|---|---|---|---|---|
| L6b | 5.82 | 3.39 | 2.78 | 2.41 | 2.09 | 1.91 | 1.70 | 1.56 | 1.33 | 1.00 |
| L6−IT−Car3 | 6.94 | 4.22 | 3.45 | 2.89 | 2.67 | 2.40 | 2.06 | 1.77 | 1.55 | 1.00 |
| L6−IT | 0.88 | 0.62 | 0.58 | 0.52 | 0.51 | 0.48 | 0.46 | 0.46 | 0.49 | 1.00 |
| L6−CT | 0.50 | 0.34 | 0.31 | 0.30 | 0.29 | 0.27 | 0.27 | 0.29 | 0.33 | 1.00 |
| L56−NP | 1.20 | 0.84 | 0.75 | 0.68 | 0.63 | 0.62 | 0.58 | 0.57 | 0.57 | 1.00 |
| L5−IT | 0.95 | 0.60 | 0.50 | 0.43 | 0.40 | 0.38 | 0.37 | 0.36 | 0.40 | 1.00 |
| L5−ET | 2.99 | 1.99 | 1.77 | 1.56 | 1.49 | 1.36 | 1.33 | 1.28 | 1.17 | 1.00 |
| L4−IT | 2.94 | 1.84 | 1.67 | 1.53 | 1.45 | 1.36 | 1.35 | 1.25 | 1.20 | 1.00 |
| HIP−Misc2 | 0.17 | 0.15 | 0.16 | 0.17 | 0.17 | 0.17 | 0.19 | 0.20 | 0.24 | 1.00 |
| HIP−Misc1 | 1.98 | 1.29 | 1.18 | 1.12 | 1.04 | 1.05 | 1.03 | 1.01 | 1.02 | 1.00 |
| DG | 1.21 | 1.02 | 0.94 | 0.88 | 0.89 | 0.85 | 0.84 | 0.88 | 0.93 | 1.00 |
| CA3 | 3.91 | 2.71 | 2.32 | 2.08 | 1.93 | 1.86 | 1.71 | 1.52 | 1.34 | 1.00 |
| CA1 | 3.28 | 2.16 | 1.93 | 1.78 | 1.69 | 1.59 | 1.48 | 1.38 | 1.26 | 1.00 |
| Amy−Exc | 0.94 | 0.65 | 0.56 | 0.51 | 0.48 | 0.46 | 0.44 | 0.43 | 0.48 | 1.00 |
| Vip | 0.86 | 0.61 | 0.52 | 0.51 | 0.49 | 0.45 | 0.45 | 0.46 | 0.50 | 1.00 |
| THM−MB | 0.66 | 0.47 | 0.38 | 0.35 | 0.32 | 0.30 | 0.30 | 0.30 | 0.34 | 1.00 |
| THM−Inh | 0.68 | 0.48 | 0.41 | 0.38 | 0.34 | 0.32 | 0.31 | 0.32 | 0.37 | 1.00 |
| THM−Exc | 9.08 | 4.57 | 3.42 | 2.80 | 2.38 | 2.14 | 1.88 | 1.62 | 1.41 | 1.00 |
| SubCtx−Cplx | 0.66 | 0.49 | 0.42 | 0.41 | 0.39 | 0.37 | 0.35 | 0.37 | 0.41 | 1.00 |
| Sst | 0.58 | 0.40 | 0.34 | 0.32 | 0.28 | 0.28 | 0.27 | 0.28 | 0.32 | 1.00 |
| Sncg | 0.87 | 0.58 | 0.51 | 0.45 | 0.43 | 0.41 | 0.40 | 0.39 | 0.44 | 1.00 |
| Pvalb−ChC | 0.73 | 0.56 | 0.49 | 0.45 | 0.43 | 0.41 | 0.41 | 0.40 | 0.42 | 1.00 |
| Pvalb | 0.56 | 0.39 | 0.34 | 0.31 | 0.29 | 0.27 | 0.27 | 0.29 | 0.31 | 1.00 |
| PN | 1.02 | 0.61 | 0.49 | 0.43 | 0.40 | 0.36 | 0.34 | 0.35 | 0.39 | 1.00 |
| MSN−D2 | 0.63 | 0.50 | 0.46 | 0.44 | 0.40 | 0.37 | 0.39 | 0.41 | 0.44 | 1.00 |
| MSN−D1 | 0.42 | 0.32 | 0.28 | 0.26 | 0.25 | 0.24 | 0.24 | 0.27 | 0.31 | 1.00 |
| Lamp5−Lhx6 | 0.54 | 0.41 | 0.38 | 0.34 | 0.33 | 0.31 | 0.32 | 0.34 | 0.37 | 1.00 |
| Lamp5 | 1.00 | 0.72 | 0.65 | 0.59 | 0.54 | 0.53 | 0.51 | 0.52 | 0.55 | 1.00 |
| Foxp2 | 0.51 | 0.42 | 0.39 | 0.37 | 0.35 | 0.36 | 0.34 | 0.36 | 0.43 | 1.00 |
| Chd7 | 0.61 | 0.44 | 0.40 | 0.37 | 0.34 | 0.34 | 0.33 | 0.34 | 0.38 | 1.00 |
| CB | 0.55 | 0.44 | 0.42 | 0.39 | 0.38 | 0.36 | 0.36 | 0.36 | 0.43 | 1.00 |

Percent 0.0025 0.0050 0.0075 0.0100

### ASCNT−Non−telencephalon astrocytes

| | 0.0−0.1 | 0.1−0.2 | 0.2−0.3 | 0.3−0.4 | 0.4−0.5 | 0.5−0.6 | 0.6−0.7 | 0.7−0.8 | 0.8−0.9 | 0.9−1.0 |
|---|---|---|---|---|---|---|---|---|---|---|
| L6b | 5.44 | 3.50 | 2.88 | 2.52 | 2.27 | 2.03 | 1.82 | 1.64 | 1.38 | 1.00 |
| L6−IT−Car3 | 5.97 | 3.98 | 3.34 | 2.86 | 2.58 | 2.38 | 1.99 | 1.78 | 1.50 | 1.00 |
| L6−IT | 1.45 | 1.06 | 0.92 | 0.84 | 0.77 | 0.71 | 0.67 | 0.62 | 0.61 | 1.00 |
| L6−CT | 0.92 | 0.66 | 0.58 | 0.53 | 0.49 | 0.46 | 0.44 | 0.43 | 0.42 | 1.00 |
| L56−NP | 1.88 | 1.32 | 1.14 | 1.04 | 0.95 | 0.88 | 0.83 | 0.76 | 0.70 | 1.00 |
| L5−IT | 1.67 | 1.07 | 0.87 | 0.76 | 0.65 | 0.57 | 0.53 | 0.50 | 0.51 | 1.00 |
| L5−ET | 3.21 | 2.26 | 2.02 | 1.87 | 1.68 | 1.59 | 1.47 | 1.36 | 1.21 | 1.00 |
| L4−IT | 3.10 | 2.22 | 1.98 | 1.83 | 1.69 | 1.61 | 1.49 | 1.40 | 1.26 | 1.00 |
| HIP−Misc2 | 0.29 | 0.26 | 0.27 | 0.27 | 0.28 | 0.29 | 0.30 | 0.32 | 0.36 | 1.00 |
| HIP−Misc1 | 2.28 | 1.72 | 1.53 | 1.44 | 1.35 | 1.32 | 1.22 | 1.17 | 1.08 | 1.00 |
| DG | 1.74 | 1.49 | 1.33 | 1.26 | 1.22 | 1.16 | 1.10 | 1.06 | 1.06 | 1.00 |
| CA3 | 3.80 | 2.84 | 2.50 | 2.25 | 2.07 | 1.91 | 1.72 | 1.58 | 1.35 | 1.00 |
| CA1 | 3.41 | 2.59 | 2.22 | 2.02 | 1.91 | 1.78 | 1.65 | 1.51 | 1.28 | 1.00 |
| Amy−Exc | 1.56 | 1.12 | 0.95 | 0.82 | 0.75 | 0.69 | 0.64 | 0.59 | 0.57 | 1.00 |
| Vip | 1.52 | 1.08 | 0.94 | 0.86 | 0.79 | 0.72 | 0.67 | 0.62 | 0.61 | 1.00 |
| THM−MB | 1.29 | 0.88 | 0.72 | 0.62 | 0.54 | 0.49 | 0.45 | 0.42 | 0.43 | 1.00 |
| THM−Inh | 1.30 | 0.91 | 0.74 | 0.67 | 0.60 | 0.54 | 0.50 | 0.48 | 0.48 | 1.00 |
| THM−Exc | 7.35 | 4.24 | 3.35 | 2.85 | 2.47 | 2.19 | 1.95 | 1.77 | 1.49 | 1.00 |
| SubCtx−Cplx | 1.19 | 0.90 | 0.78 | 0.72 | 0.65 | 0.60 | 0.55 | 0.53 | 0.52 | 1.00 |
| Sst | 1.05 | 0.75 | 0.64 | 0.57 | 0.51 | 0.47 | 0.43 | 0.40 | 0.40 | 1.00 |
| Sncg | 1.57 | 1.09 | 0.92 | 0.81 | 0.73 | 0.66 | 0.59 | 0.55 | 0.55 | 1.00 |
| Pvalb−ChC | 1.30 | 0.98 | 0.86 | 0.77 | 0.69 | 0.62 | 0.59 | 0.55 | 0.54 | 1.00 |
| Pvalb | 1.04 | 0.76 | 0.65 | 0.57 | 0.53 | 0.47 | 0.43 | 0.41 | 0.41 | 1.00 |
| PN | 1.82 | 1.09 | 0.87 | 0.75 | 0.67 | 0.58 | 0.53 | 0.50 | 0.50 | 1.00 |
| MSN−D2 | 1.20 | 0.93 | 0.84 | 0.76 | 0.68 | 0.61 | 0.58 | 0.55 | 0.56 | 1.00 |
| MSN−D1 | 0.82 | 0.64 | 0.56 | 0.52 | 0.46 | 0.41 | 0.39 | 0.38 | 0.40 | 1.00 |
| Lamp5−Lhx6 | 1.04 | 0.79 | 0.70 | 0.64 | 0.59 | 0.54 | 0.49 | 0.48 | 0.47 | 1.00 |
| Lamp5 | 1.76 | 1.28 | 1.10 | 1.00 | 0.88 | 0.82 | 0.74 | 0.70 | 0.67 | 1.00 |
| Foxp2 | 0.99 | 0.81 | 0.72 | 0.66 | 0.63 | 0.57 | 0.54 | 0.51 | 0.53 | 1.00 |
| Chd7 | 1.19 | 0.87 | 0.74 | 0.67 | 0.61 | 0.55 | 0.51 | 0.48 | 0.49 | 1.00 |
| CB | 1.07 | 0.84 | 0.77 | 0.71 | 0.66 | 0.60 | 0.57 | 0.52 | 0.55 | 1.00 |

Percent 0.005 0.010 0.015 0.020

### ASCT−Telencephalon astrocytes

| | 0.0−0.1 | 0.1−0.2 | 0.2−0.3 | 0.3−0.4 | 0.4−0.5 | 0.5−0.6 | 0.6−0.7 | 0.7−0.8 | 0.8−0.9 | 0.9−1.0 |
|---|---|---|---|---|---|---|---|---|---|---|
| L6b | 4.73 | 3.29 | 2.86 | 2.53 | 2.28 | 2.05 | 1.83 | 1.61 | 1.38 | 1.00 |
| L6−IT−Car3 | 5.01 | 3.61 | 3.10 | 2.73 | 2.50 | 2.23 | 2.01 | 1.74 | 1.48 | 1.00 |
| L6−IT | 1.93 | 1.40 | 1.23 | 1.12 | 1.04 | 0.95 | 0.86 | 0.79 | 0.73 | 1.00 |
| L6−CT | 1.29 | 0.94 | 0.82 | 0.76 | 0.71 | 0.65 | 0.61 | 0.56 | 0.53 | 1.00 |
| L56−NP | 2.33 | 1.70 | 1.50 | 1.36 | 1.25 | 1.17 | 1.06 | 0.97 | 0.86 | 1.00 |
| L5−IT | 2.31 | 1.51 | 1.23 | 1.06 | 0.93 | 0.82 | 0.73 | 0.66 | 0.62 | 1.00 |
| L5−ET | 3.14 | 2.36 | 2.16 | 1.99 | 1.81 | 1.71 | 1.57 | 1.45 | 1.29 | 1.00 |
| L4−IT | 3.08 | 2.38 | 2.18 | 2.03 | 1.86 | 1.75 | 1.62 | 1.48 | 1.30 | 1.00 |
| HIP−Misc2 | 0.41 | 0.37 | 0.37 | 0.38 | 0.38 | 0.39 | 0.40 | 0.41 | 0.45 | 1.00 |
| HIP−Misc1 | 2.46 | 1.97 | 1.79 | 1.69 | 1.57 | 1.51 | 1.39 | 1.34 | 1.18 | 1.00 |
| DG | 2.06 | 1.77 | 1.64 | 1.53 | 1.45 | 1.37 | 1.31 | 1.22 | 1.15 | 1.00 |
| CA3 | 3.50 | 2.75 | 2.49 | 2.28 | 2.08 | 1.93 | 1.77 | 1.57 | 1.36 | 1.00 |
| CA1 | 3.32 | 2.59 | 2.30 | 2.12 | 2.02 | 1.87 | 1.71 | 1.55 | 1.35 | 1.00 |
| Amy−Exc | 2.08 | 1.50 | 1.28 | 1.14 | 1.04 | 0.95 | 0.85 | 0.78 | 0.71 | 1.00 |
| Vip | 2.10 | 1.55 | 1.35 | 1.23 | 1.11 | 1.03 | 0.93 | 0.82 | 0.75 | 1.00 |
| THM−MB | 1.85 | 1.29 | 1.06 | 0.92 | 0.81 | 0.72 | 0.64 | 0.57 | 0.54 | 1.00 |
| THM−Inh | 1.78 | 1.29 | 1.07 | 0.97 | 0.86 | 0.78 | 0.70 | 0.64 | 0.60 | 1.00 |
| THM−Exc | 5.86 | 3.76 | 3.00 | 2.68 | 2.35 | 2.16 | 1.91 | 1.73 | 1.44 | 1.00 |
| SubCtx−Cplx | 1.67 | 1.29 | 1.15 | 1.06 | 0.96 | 0.87 | 0.79 | 0.72 | 0.67 | 1.00 |
| Sst | 1.46 | 1.11 | 0.95 | 0.84 | 0.77 | 0.69 | 0.62 | 0.55 | 0.51 | 1.00 |
| Sncg | 2.18 | 1.58 | 1.37 | 1.21 | 1.07 | 0.95 | 0.84 | 0.76 | 0.69 | 1.00 |
| Pvalb−ChC | 1.80 | 1.39 | 1.21 | 1.08 | 0.98 | 0.89 | 0.81 | 0.73 | 0.67 | 1.00 |
| Pvalb | 1.46 | 1.08 | 0.95 | 0.84 | 0.76 | 0.69 | 0.62 | 0.57 | 0.52 | 1.00 |
| PN | 2.50 | 1.54 | 1.25 | 1.06 | 0.96 | 0.84 | 0.75 | 0.68 | 0.62 | 1.00 |
| MSN−D2 | 1.68 | 1.35 | 1.18 | 1.08 | 0.98 | 0.88 | 0.79 | 0.72 | 0.68 | 1.00 |
| MSN−D1 | 1.20 | 0.95 | 0.84 | 0.76 | 0.68 | 0.61 | 0.56 | 0.53 | 0.50 | 1.00 |
| Lamp5−Lhx6 | 1.50 | 1.16 | 1.03 | 0.94 | 0.86 | 0.77 | 0.70 | 0.64 | 0.60 | 1.00 |
| Lamp5 | 2.46 | 1.82 | 1.58 | 1.43 | 1.27 | 1.15 | 1.01 | 0.92 | 0.82 | 1.00 |
| Foxp2 | 1.45 | 1.19 | 1.08 | 0.98 | 0.92 | 0.84 | 0.76 | 0.71 | 0.67 | 1.00 |
| Chd7 | 1.68 | 1.27 | 1.10 | 0.98 | 0.87 | 0.79 | 0.72 | 0.64 | 0.60 | 1.00 |
| CB | 1.49 | 1.22 | 1.10 | 1.02 | 0.93 | 0.86 | 0.78 | 0.70 | 0.68 | 1.00 |

Percent 0.010 0.015 0.020 0.025 0.030

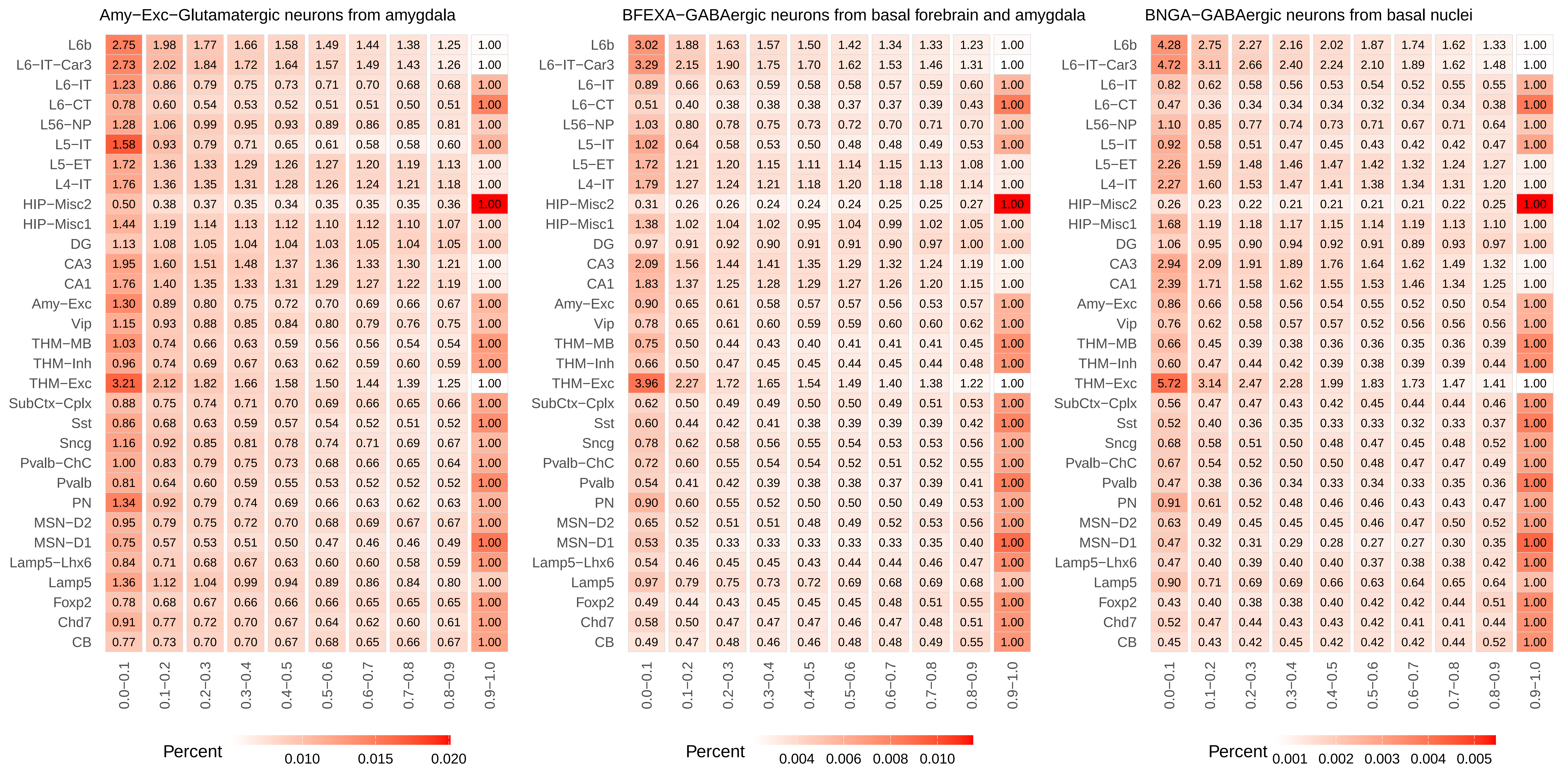
Amy−Exc−Glutamatergic neurons from amygdala
L6b 2.75 1.98 1.77 1.66 1.58 1.49 1.44 1.38 1.25 1.00
L6−IT−Car3 2.73 2.02 1.84 1.72 1.64 1.57 1.49 1.43 1.26 1.00
L6−IT 1.23 0.86 0.79 0.75 0.73 0.71 0.70 0.68 0.68 1.00
L6−CT 0.78 0.60 0.54 0.53 0.52 0.51 0.51 0.50 0.51 1.00
L56−NP 1.28 1.06 0.99 0.95 0.93 0.89 0.86 0.85 0.81 1.00
L5−IT 1.58 0.93 0.79 0.71 0.65 0.61 0.58 0.58 0.60 1.00
L5−ET 1.72 1.36 1.33 1.29 1.26 1.27 1.20 1.19 1.13 1.00
L4−IT 1.76 1.36 1.35 1.31 1.28 1.26 1.24 1.21 1.18 1.00
HIP−Misc2 0.50 0.38 0.37 0.35 0.34 0.35 0.35 0.35 0.36 1.00
HIP−Misc1 1.44 1.19 1.14 1.13 1.12 1.10 1.12 1.10 1.07 1.00
DG 1.13 1.08 1.05 1.04 1.04 1.03 1.05 1.04 1.05 1.00
CA3 1.95 1.60 1.51 1.48 1.37 1.36 1.33 1.30 1.21 1.00
CA1 1.76 1.40 1.35 1.33 1.31 1.29 1.27 1.22 1.19 1.00
Amy−Exc 1.30 0.89 0.80 0.75 0.72 0.70 0.69 0.66 0.67 1.00
Vip 1.15 0.93 0.88 0.85 0.84 0.80 0.79 0.76 0.75 1.00
THM−MB 1.03 0.74 0.66 0.63 0.59 0.56 0.56 0.54 0.54 1.00
THM−Inh 0.96 0.74 0.69 0.67 0.63 0.62 0.59 0.60 0.59 1.00
THM−Exc 3.21 2.12 1.82 1.66 1.58 1.50 1.44 1.39 1.25 1.00
SubCtx−Cplx 0.88 0.75 0.74 0.71 0.70 0.69 0.66 0.65 0.66 1.00
Sst 0.86 0.68 0.63 0.59 0.57 0.54 0.52 0.51 0.52 1.00
Sncg 1.16 0.92 0.85 0.81 0.78 0.74 0.71 0.69 0.67 1.00
Pvalb−ChC 1.00 0.83 0.79 0.75 0.73 0.68 0.66 0.65 0.64 1.00
Pvalb 0.81 0.64 0.60 0.59 0.55 0.53 0.52 0.52 0.52 1.00
PN 1.34 0.92 0.79 0.74 0.69 0.66 0.63 0.62 0.63 1.00
MSN−D2 0.95 0.79 0.75 0.72 0.70 0.68 0.69 0.67 0.67 1.00
MSN−D1 0.75 0.57 0.53 0.51 0.50 0.47 0.46 0.46 0.49 1.00
Lamp5−Lhx6 0.84 0.71 0.68 0.67 0.63 0.60 0.60 0.58 0.59 1.00
Lamp5 1.36 1.12 1.04 0.99 0.94 0.89 0.86 0.84 0.80 1.00
Foxp2 0.78 0.68 0.67 0.66 0.66 0.66 0.65 0.65 0.65 1.00
Chd7 0.91 0.77 0.72 0.70 0.67 0.64 0.62 0.60 0.61 1.00
CB 0.77 0.73 0.70 0.70 0.67 0.68 0.65 0.66 0.67 1.00
0.0−0.1 0.1−0.2 0.2−0.3 0.3−0.4 0.4−0.5 0.5−0.6 0.6−0.7 0.7−0.8 0.8−0.9 0.9−1.0
Percent 0.010 0.015 0.020
BFEXA−GABAergic neurons from basal forebrain and amygdala
L6b 3.02 1.88 1.63 1.57 1.50 1.42 1.34 1.33 1.23 1.00
L6−IT−Car3 3.29 2.15 1.90 1.75 1.70 1.62 1.53 1.46 1.31 1.00
L6−IT 0.89 0.66 0.63 0.59 0.58 0.58 0.57 0.59 0.60 1.00
L6−CT 0.51 0.40 0.38 0.38 0.38 0.37 0.37 0.39 0.43 1.00
L56−NP 1.03 0.80 0.78 0.75 0.73 0.72 0.70 0.71 0.70 1.00
L5−IT 1.02 0.64 0.58 0.53 0.50 0.48 0.48 0.49 0.53 1.00
L5−ET 1.72 1.21 1.20 1.15 1.11 1.14 1.15 1.13 1.08 1.00
L4−IT 1.79 1.27 1.24 1.21 1.18 1.20 1.18 1.18 1.14 1.00
HIP−Misc2 0.31 0.26 0.26 0.24 0.24 0.24 0.25 0.25 0.27 1.00
HIP−Misc1 1.38 1.02 1.04 1.02 0.95 1.04 0.99 1.02 1.05 1.00
DG 0.97 0.91 0.92 0.90 0.91 0.91 0.90 0.97 1.00 1.00
CA3 2.09 1.56 1.44 1.41 1.35 1.29 1.32 1.24 1.19 1.00
CA1 1.83 1.37 1.25 1.28 1.29 1.27 1.26 1.20 1.15 1.00
Amy−Exc 0.90 0.65 0.61 0.58 0.57 0.57 0.56 0.53 0.57 1.00
Vip 0.78 0.65 0.61 0.60 0.59 0.59 0.60 0.60 0.62 1.00
THM−MB 0.75 0.50 0.44 0.43 0.40 0.41 0.41 0.41 0.45 1.00
THM−Inh 0.66 0.50 0.47 0.45 0.45 0.44 0.45 0.44 0.48 1.00
THM−Exc 3.96 2.27 1.72 1.65 1.54 1.49 1.40 1.38 1.22 1.00
SubCtx−Cplx 0.62 0.50 0.49 0.49 0.50 0.50 0.49 0.51 0.53 1.00
Sst 0.60 0.44 0.42 0.41 0.38 0.39 0.39 0.39 0.42 1.00
Sncg 0.78 0.62 0.58 0.56 0.55 0.54 0.53 0.53 0.56 1.00
Pvalb−ChC 0.72 0.60 0.55 0.54 0.54 0.52 0.51 0.52 0.55 1.00
Pvalb 0.54 0.41 0.42 0.39 0.38 0.38 0.37 0.39 0.41 1.00
PN 0.90 0.60 0.55 0.52 0.50 0.50 0.50 0.49 0.53 1.00
MSN−D2 0.65 0.52 0.51 0.51 0.48 0.49 0.52 0.53 0.56 1.00
MSN−D1 0.53 0.35 0.33 0.33 0.33 0.33 0.33 0.35 0.40 1.00
Lamp5−Lhx6 0.54 0.46 0.45 0.45 0.43 0.44 0.44 0.46 0.47 1.00
Lamp5 0.97 0.79 0.75 0.73 0.72 0.69 0.68 0.69 0.68 1.00
Foxp2 0.49 0.44 0.43 0.45 0.45 0.45 0.48 0.51 0.55 1.00
Chd7 0.58 0.50 0.47 0.47 0.47 0.46 0.47 0.48 0.51 1.00
CB 0.49 0.47 0.48 0.46 0.46 0.48 0.48 0.49 0.55 1.00
0.0−0.1 0.1−0.2 0.2−0.3 0.3−0.4 0.4−0.5 0.5−0.6 0.6−0.7 0.7−0.8 0.8−0.9 0.9−1.0
Percent 0.004 0.006 0.008 0.010
BNGA−GABAergic neurons from basal nuclei
L6b 4.28 2.75 2.27 2.16 2.02 1.87 1.74 1.62 1.33 1.00
L6−IT−Car3 4.72 3.11 2.66 2.40 2.24 2.10 1.89 1.62 1.48 1.00
L6−IT 0.82 0.62 0.58 0.56 0.53 0.54 0.52 0.55 0.55 1.00
L6−CT 0.47 0.36 0.34 0.34 0.34 0.32 0.34 0.34 0.38 1.00
L56−NP 1.10 0.85 0.77 0.74 0.73 0.71 0.67 0.71 0.64 1.00
L5−IT 0.92 0.58 0.51 0.47 0.45 0.43 0.42 0.42 0.47 1.00
L5−ET 2.26 1.59 1.48 1.46 1.47 1.42 1.32 1.24 1.27 1.00
L4−IT 2.27 1.60 1.53 1.47 1.41 1.38 1.34 1.31 1.20 1.00
HIP−Misc2 0.26 0.23 0.22 0.21 0.21 0.21 0.21 0.22 0.25 1.00
HIP−Misc1 1.68 1.19 1.18 1.17 1.15 1.14 1.19 1.13 1.10 1.00
DG 1.06 0.95 0.90 0.94 0.92 0.91 0.89 0.93 0.97 1.00
CA3 2.94 2.09 1.91 1.89 1.76 1.64 1.62 1.49 1.32 1.00
CA1 2.39 1.71 1.58 1.62 1.55 1.53 1.46 1.34 1.25 1.00
Amy−Exc 0.86 0.66 0.58 0.56 0.54 0.55 0.52 0.50 0.54 1.00
Vip 0.76 0.62 0.58 0.57 0.57 0.52 0.56 0.56 0.56 1.00
THM−MB 0.66 0.45 0.39 0.38 0.36 0.36 0.35 0.36 0.39 1.00
THM−Inh 0.60 0.47 0.44 0.42 0.39 0.38 0.39 0.39 0.44 1.00
THM−Exc 5.72 3.14 2.47 2.28 1.99 1.83 1.73 1.47 1.41 1.00
SubCtx−Cplx 0.56 0.47 0.47 0.43 0.42 0.45 0.44 0.44 0.46 1.00
Sst 0.52 0.40 0.36 0.35 0.33 0.33 0.32 0.33 0.37 1.00
Sncg 0.68 0.58 0.51 0.50 0.48 0.47 0.45 0.48 0.52 1.00
Pvalb−ChC 0.67 0.54 0.52 0.50 0.50 0.48 0.47 0.47 0.49 1.00
Pvalb 0.47 0.38 0.36 0.34 0.33 0.34 0.33 0.35 0.36 1.00
PN 0.91 0.61 0.52 0.48 0.46 0.46 0.43 0.43 0.47 1.00
MSN−D2 0.63 0.49 0.45 0.45 0.45 0.46 0.47 0.50 0.52 1.00
MSN−D1 0.47 0.32 0.31 0.29 0.28 0.27 0.27 0.30 0.35 1.00
Lamp5−Lhx6 0.47 0.40 0.39 0.40 0.40 0.37 0.38 0.38 0.42 1.00
Lamp5 0.90 0.71 0.69 0.69 0.66 0.63 0.64 0.65 0.64 1.00
Foxp2 0.43 0.40 0.38 0.38 0.40 0.42 0.42 0.44 0.51 1.00
Chd7 0.52 0.47 0.44 0.43 0.43 0.42 0.41 0.41 0.44 1.00
CB 0.45 0.43 0.42 0.45 0.42 0.42 0.42 0.44 0.52 1.00
0.0−0.1 0.1−0.2 0.2−0.3 0.3−0.4 0.4−0.5 0.5−0.6 0.6−0.7 0.7−0.8 0.8−0.9 0.9−1.0
Percent 0.001 0.002 0.003 0.004 0.005

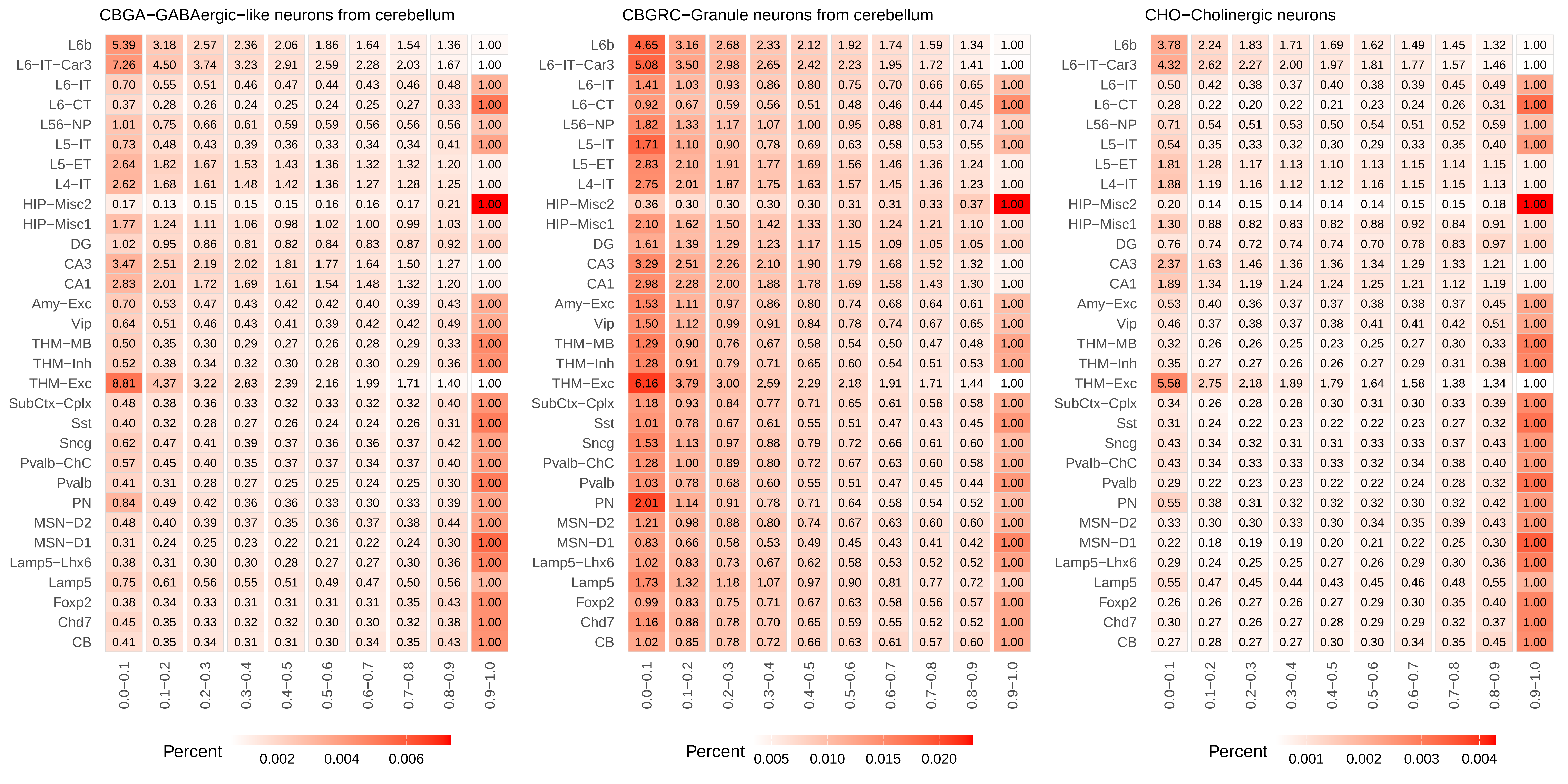
CBGA−GABAergic−like neurons from cerebellum
L6b 5.39 3.18 2.57 2.36 2.06 1.86 1.64 1.54 1.36 1.00
L6−IT−Car3 7.26 4.50 3.74 3.23 2.91 2.59 2.28 2.03 1.67 1.00
L6−IT 0.70 0.55 0.51 0.46 0.47 0.44 0.43 0.46 0.48 1.00
L6−CT 0.37 0.28 0.26 0.24 0.25 0.24 0.25 0.27 0.33 1.00
L56−NP 1.01 0.75 0.66 0.61 0.59 0.59 0.56 0.56 0.56 1.00
L5−IT 0.73 0.48 0.43 0.39 0.36 0.33 0.34 0.34 0.41 1.00
L5−ET 2.64 1.82 1.67 1.53 1.43 1.36 1.32 1.32 1.20 1.00
L4−IT 2.62 1.68 1.61 1.48 1.42 1.36 1.27 1.28 1.25 1.00
HIP−Misc2 0.17 0.13 0.15 0.15 0.15 0.16 0.16 0.17 0.21 1.00
HIP−Misc1 1.77 1.24 1.11 1.06 0.98 1.02 1.00 0.99 1.03 1.00
DG 1.02 0.95 0.86 0.81 0.82 0.84 0.83 0.87 0.92 1.00
CA3 3.47 2.51 2.19 2.02 1.81 1.77 1.64 1.50 1.27 1.00
CA1 2.83 2.01 1.72 1.69 1.61 1.54 1.48 1.32 1.20 1.00
Amy−Exc 0.70 0.53 0.47 0.43 0.42 0.42 0.40 0.39 0.43 1.00
Vip 0.64 0.51 0.46 0.43 0.41 0.39 0.42 0.42 0.49 1.00
THM−MB 0.50 0.35 0.30 0.29 0.27 0.26 0.28 0.29 0.33 1.00
THM−Inh 0.52 0.38 0.34 0.32 0.30 0.28 0.30 0.29 0.36 1.00
THM−Exc 8.81 4.37 3.22 2.83 2.39 2.16 1.99 1.71 1.40 1.00
SubCtx−Cplx 0.48 0.38 0.36 0.33 0.32 0.33 0.32 0.32 0.40 1.00
Sst 0.40 0.32 0.28 0.27 0.26 0.24 0.24 0.26 0.31 1.00
Sncg 0.62 0.47 0.41 0.39 0.37 0.36 0.36 0.37 0.42 1.00
Pvalb−ChC 0.57 0.45 0.40 0.35 0.37 0.37 0.34 0.37 0.40 1.00
Pvalb 0.41 0.31 0.28 0.27 0.25 0.25 0.24 0.25 0.30 1.00
PN 0.84 0.49 0.42 0.36 0.36 0.33 0.30 0.33 0.39 1.00
MSN−D2 0.48 0.40 0.39 0.37 0.35 0.36 0.37 0.38 0.44 1.00
MSN−D1 0.31 0.24 0.25 0.23 0.22 0.21 0.22 0.24 0.30 1.00
Lamp5−Lhx6 0.38 0.31 0.30 0.30 0.28 0.27 0.27 0.30 0.36 1.00
Lamp5 0.75 0.61 0.56 0.55 0.51 0.49 0.47 0.50 0.56 1.00
Foxp2 0.38 0.34 0.33 0.31 0.31 0.31 0.31 0.35 0.43 1.00
Chd7 0.45 0.35 0.33 0.32 0.32 0.30 0.30 0.32 0.38 1.00
CB 0.41 0.35 0.34 0.31 0.31 0.30 0.34 0.35 0.43 1.00
0.0−0.1 0.1−0.2 0.2−0.3 0.3−0.4 0.4−0.5 0.5−0.6 0.6−0.7 0.7−0.8 0.8−0.9 0.9−1.0
Percent 0.002 0.004 0.006
CBGRC−Granule neurons from cerebellum
L6b 4.65 3.16 2.68 2.33 2.12 1.92 1.74 1.59 1.34 1.00
L6−IT−Car3 5.08 3.50 2.98 2.65 2.42 2.23 1.95 1.72 1.41 1.00
L6−IT 1.41 1.03 0.93 0.86 0.80 0.75 0.70 0.66 0.65 1.00
L6−CT 0.92 0.67 0.59 0.56 0.51 0.48 0.46 0.44 0.45 1.00
L56−NP 1.82 1.33 1.17 1.07 1.00 0.95 0.88 0.81 0.74 1.00
L5−IT 1.71 1.10 0.90 0.78 0.69 0.63 0.58 0.53 0.55 1.00
L5−ET 2.83 2.10 1.91 1.77 1.69 1.56 1.46 1.36 1.24 1.00
L4−IT 2.75 2.01 1.87 1.75 1.63 1.57 1.45 1.36 1.23 1.00
HIP−Misc2 0.36 0.30 0.30 0.30 0.30 0.31 0.31 0.33 0.37 1.00
HIP−Misc1 2.10 1.62 1.50 1.42 1.33 1.30 1.24 1.21 1.10 1.00
DG 1.61 1.39 1.29 1.23 1.17 1.15 1.09 1.05 1.05 1.00
CA3 3.29 2.51 2.26 2.10 1.90 1.79 1.68 1.52 1.32 1.00
CA1 2.98 2.28 2.00 1.88 1.78 1.69 1.58 1.43 1.30 1.00
Amy−Exc 1.53 1.11 0.97 0.86 0.80 0.74 0.68 0.64 0.61 1.00
Vip 1.50 1.12 0.99 0.91 0.84 0.78 0.74 0.67 0.65 1.00
THM−MB 1.29 0.90 0.76 0.67 0.58 0.54 0.50 0.47 0.48 1.00
THM−Inh 1.28 0.91 0.79 0.71 0.65 0.60 0.54 0.51 0.53 1.00
THM−Exc 6.16 3.79 3.00 2.59 2.29 2.18 1.91 1.71 1.44 1.00
SubCtx−Cplx 1.18 0.93 0.84 0.77 0.71 0.65 0.61 0.58 0.58 1.00
Sst 1.01 0.78 0.67 0.61 0.55 0.51 0.47 0.43 0.45 1.00
Sncg 1.53 1.13 0.97 0.88 0.79 0.72 0.66 0.61 0.60 1.00
Pvalb−ChC 1.28 1.00 0.89 0.80 0.72 0.67 0.63 0.60 0.58 1.00
Pvalb 1.03 0.78 0.68 0.60 0.55 0.51 0.47 0.45 0.44 1.00
PN 2.01 1.14 0.91 0.78 0.71 0.64 0.58 0.54 0.52 1.00
MSN−D2 1.21 0.98 0.88 0.80 0.74 0.67 0.63 0.60 0.60 1.00
MSN−D1 0.83 0.66 0.58 0.53 0.49 0.45 0.43 0.41 0.42 1.00
Lamp5−Lhx6 1.02 0.83 0.73 0.67 0.62 0.58 0.53 0.52 0.52 1.00
Lamp5 1.73 1.32 1.18 1.07 0.97 0.90 0.81 0.77 0.72 1.00
Foxp2 0.99 0.83 0.75 0.71 0.67 0.63 0.58 0.56 0.57 1.00
Chd7 1.16 0.88 0.78 0.70 0.65 0.59 0.55 0.52 0.52 1.00
CB 1.02 0.85 0.78 0.72 0.66 0.63 0.61 0.57 0.60 1.00
0.0−0.1 0.1−0.2 0.2−0.3 0.3−0.4 0.4−0.5 0.5−0.6 0.6−0.7 0.7−0.8 0.8−0.9 0.9−1.0
Percent 0.005 0.010 0.015 0.020
CHO−Cholinergic neurons
L6b 3.78 2.24 1.83 1.71 1.69 1.62 1.49 1.45 1.32 1.00
L6−IT−Car3 4.32 2.62 2.27 2.00 1.97 1.81 1.77 1.57 1.46 1.00
L6−IT 0.50 0.42 0.38 0.37 0.40 0.38 0.39 0.45 0.49 1.00
L6−CT 0.28 0.22 0.20 0.22 0.21 0.23 0.24 0.26 0.31 1.00
L56−NP 0.71 0.54 0.51 0.53 0.50 0.54 0.51 0.52 0.59 1.00
L5−IT 0.54 0.35 0.33 0.32 0.30 0.29 0.33 0.35 0.40 1.00
L5−ET 1.81 1.28 1.17 1.13 1.10 1.13 1.15 1.14 1.15 1.00
L4−IT 1.88 1.19 1.16 1.12 1.12 1.16 1.15 1.15 1.13 1.00
HIP−Misc2 0.20 0.14 0.15 0.14 0.14 0.14 0.15 0.15 0.18 1.00
HIP−Misc1 1.30 0.88 0.82 0.83 0.82 0.88 0.92 0.84 0.91 1.00
DG 0.76 0.74 0.72 0.74 0.74 0.70 0.78 0.83 0.97 1.00
CA3 2.37 1.63 1.46 1.36 1.36 1.34 1.29 1.33 1.21 1.00
CA1 1.89 1.34 1.19 1.24 1.24 1.25 1.21 1.12 1.19 1.00
Amy−Exc 0.53 0.40 0.36 0.37 0.37 0.38 0.38 0.37 0.45 1.00
Vip 0.46 0.37 0.38 0.37 0.38 0.41 0.41 0.42 0.51 1.00
THM−MB 0.32 0.26 0.26 0.25 0.23 0.25 0.27 0.30 0.33 1.00
THM−Inh 0.35 0.27 0.27 0.26 0.26 0.27 0.29 0.31 0.38 1.00
THM−Exc 5.58 2.75 2.18 1.89 1.79 1.64 1.58 1.38 1.34 1.00
SubCtx−Cplx 0.34 0.26 0.28 0.28 0.30 0.31 0.30 0.33 0.39 1.00
Sst 0.31 0.24 0.22 0.23 0.22 0.22 0.23 0.27 0.32 1.00
Sncg 0.43 0.34 0.32 0.31 0.31 0.33 0.33 0.37 0.43 1.00
Pvalb−ChC 0.43 0.34 0.33 0.33 0.33 0.32 0.34 0.38 0.40 1.00
Pvalb 0.29 0.22 0.23 0.23 0.22 0.22 0.24 0.28 0.32 1.00
PN 0.55 0.38 0.31 0.32 0.32 0.32 0.30 0.32 0.42 1.00
MSN−D2 0.33 0.30 0.30 0.33 0.30 0.34 0.35 0.39 0.43 1.00
MSN−D1 0.22 0.18 0.19 0.19 0.20 0.21 0.22 0.25 0.30 1.00
Lamp5−Lhx6 0.29 0.24 0.25 0.25 0.27 0.26 0.29 0.30 0.36 1.00
Lamp5 0.55 0.47 0.45 0.44 0.43 0.45 0.46 0.48 0.55 1.00
Foxp2 0.26 0.26 0.27 0.26 0.27 0.29 0.30 0.35 0.40 1.00
Chd7 0.30 0.27 0.26 0.27 0.28 0.29 0.29 0.32 0.37 1.00
CB 0.27 0.28 0.27 0.27 0.30 0.30 0.34 0.35 0.45 1.00
0.0−0.1 0.1−0.2 0.2−0.3 0.3−0.4 0.4−0.5 0.5−0.6 0.6−0.7 0.7−0.8 0.8−0.9 0.9−1.0
Percent 0.001 0.002 0.003 0.004

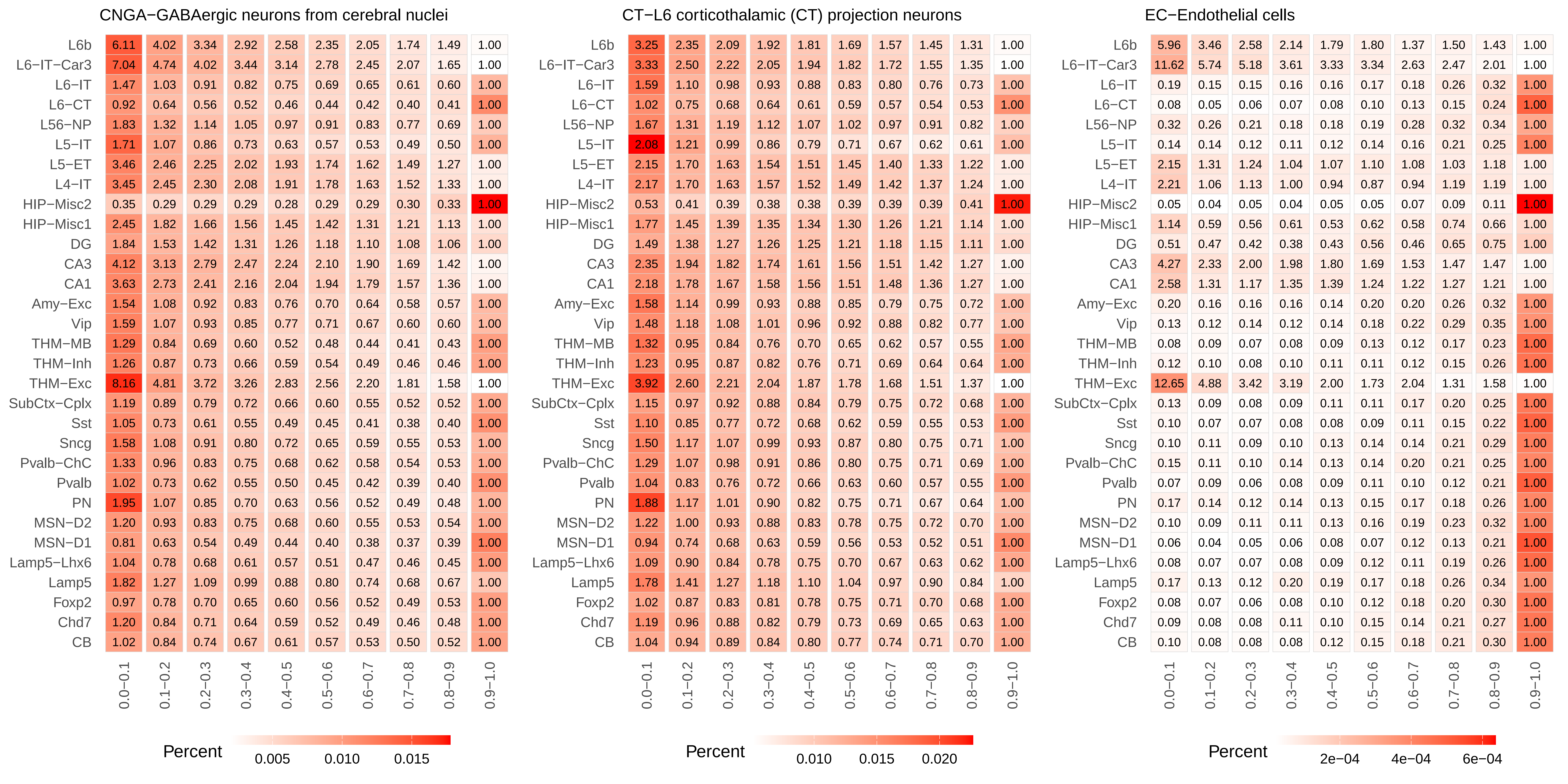
CNGA−GABAergic neurons from cerebral nuclei
L6b 6.11 4.02 3.34 2.92 2.58 2.35 2.05 1.74 1.49 1.00
L6−IT−Car3 7.04 4.74 4.02 3.44 3.14 2.78 2.45 2.07 1.65 1.00
L6−IT 1.47 1.03 0.91 0.82 0.75 0.69 0.65 0.61 0.60 1.00
L6−CT 0.92 0.64 0.56 0.52 0.46 0.44 0.42 0.40 0.41 1.00
L56−NP 1.83 1.32 1.14 1.05 0.97 0.91 0.83 0.77 0.69 1.00
L5−IT 1.71 1.07 0.86 0.73 0.63 0.57 0.53 0.49 0.50 1.00
L5−ET 3.46 2.46 2.25 2.02 1.93 1.74 1.62 1.49 1.27 1.00
L4−IT 3.45 2.45 2.30 2.08 1.91 1.78 1.63 1.52 1.33 1.00
HIP−Misc2 0.35 0.29 0.29 0.29 0.28 0.29 0.29 0.30 0.33 1.00
HIP−Misc1 2.45 1.82 1.66 1.56 1.45 1.42 1.31 1.21 1.13 1.00
DG 1.84 1.53 1.42 1.31 1.26 1.18 1.10 1.08 1.06 1.00
CA3 4.12 3.13 2.79 2.47 2.24 2.10 1.90 1.69 1.42 1.00
CA1 3.63 2.73 2.41 2.16 2.04 1.94 1.79 1.57 1.36 1.00
Amy−Exc 1.54 1.08 0.92 0.83 0.76 0.70 0.64 0.58 0.57 1.00
Vip 1.59 1.07 0.93 0.85 0.77 0.71 0.67 0.60 0.60 1.00
THM−MB 1.29 0.84 0.69 0.60 0.52 0.48 0.44 0.41 0.43 1.00
THM−Inh 1.26 0.87 0.73 0.66 0.59 0.54 0.49 0.46 0.46 1.00
THM−Exc 8.16 4.81 3.72 3.26 2.83 2.56 2.20 1.81 1.58 1.00
SubCtx−Cplx 1.19 0.89 0.79 0.72 0.66 0.60 0.55 0.52 0.52 1.00
Sst 1.05 0.73 0.61 0.55 0.49 0.45 0.41 0.38 0.40 1.00
Sncg 1.58 1.08 0.91 0.80 0.72 0.65 0.59 0.55 0.53 1.00
Pvalb−ChC 1.33 0.96 0.83 0.75 0.68 0.62 0.58 0.54 0.53 1.00
Pvalb 1.02 0.73 0.62 0.55 0.50 0.45 0.42 0.39 0.40 1.00
PN 1.95 1.07 0.85 0.70 0.63 0.56 0.52 0.49 0.48 1.00
MSN−D2 1.20 0.93 0.83 0.75 0.68 0.60 0.55 0.53 0.54 1.00
MSN−D1 0.81 0.63 0.54 0.49 0.44 0.40 0.38 0.37 0.39 1.00
Lamp5−Lhx6 1.04 0.78 0.68 0.61 0.57 0.51 0.47 0.46 0.45 1.00
Lamp5 1.82 1.27 1.09 0.99 0.88 0.80 0.74 0.68 0.67 1.00
Foxp2 0.97 0.78 0.70 0.65 0.60 0.56 0.52 0.49 0.53 1.00
Chd7 1.20 0.84 0.71 0.64 0.59 0.52 0.49 0.46 0.48 1.00
CB 1.02 0.84 0.74 0.67 0.61 0.57 0.53 0.50 0.52 1.00
0.0−0.1 0.1−0.2 0.2−0.3 0.3−0.4 0.4−0.5 0.5−0.6 0.6−0.7 0.7−0.8 0.8−0.9 0.9−1.0
Percent
0.005 0.010 0.015
CT−L6 corticothalamic (CT) projection neurons
L6b 3.25 2.35 2.09 1.92 1.81 1.69 1.57 1.45 1.31 1.00
L6−IT−Car3 3.33 2.50 2.22 2.05 1.94 1.82 1.72 1.55 1.35 1.00
L6−IT 1.59 1.10 0.98 0.93 0.88 0.83 0.80 0.76 0.73 1.00
L6−CT 1.02 0.75 0.68 0.64 0.61 0.59 0.57 0.54 0.53 1.00
L56−NP 1.67 1.31 1.19 1.12 1.07 1.02 0.97 0.91 0.82 1.00
L5−IT 2.08 1.21 0.99 0.86 0.79 0.71 0.67 0.62 0.61 1.00
L5−ET 2.15 1.70 1.63 1.54 1.51 1.45 1.40 1.33 1.22 1.00
L4−IT 2.17 1.70 1.63 1.57 1.52 1.49 1.42 1.37 1.24 1.00
HIP−Misc2 0.53 0.41 0.39 0.38 0.38 0.39 0.39 0.39 0.41 1.00
HIP−Misc1 1.77 1.45 1.39 1.35 1.34 1.30 1.26 1.21 1.14 1.00
DG 1.49 1.38 1.27 1.26 1.25 1.21 1.18 1.15 1.11 1.00
CA3 2.35 1.94 1.82 1.74 1.61 1.56 1.51 1.42 1.27 1.00
CA1 2.18 1.78 1.67 1.58 1.56 1.51 1.48 1.36 1.27 1.00
Amy−Exc 1.58 1.14 0.99 0.93 0.88 0.85 0.79 0.75 0.72 1.00
Vip 1.48 1.18 1.08 1.01 0.96 0.92 0.88 0.82 0.77 1.00
THM−MB 1.32 0.95 0.84 0.76 0.70 0.65 0.62 0.57 0.55 1.00
THM−Inh 1.23 0.95 0.87 0.82 0.76 0.71 0.69 0.64 0.64 1.00
THM−Exc 3.92 2.60 2.21 2.04 1.87 1.78 1.68 1.51 1.37 1.00
SubCtx−Cplx 1.15 0.97 0.92 0.88 0.84 0.79 0.75 0.72 0.68 1.00
Sst 1.10 0.85 0.77 0.72 0.68 0.62 0.59 0.55 0.53 1.00
Sncg 1.50 1.17 1.07 0.99 0.93 0.87 0.80 0.75 0.71 1.00
Pvalb−ChC 1.29 1.07 0.98 0.91 0.86 0.80 0.75 0.71 0.69 1.00
Pvalb 1.04 0.83 0.76 0.72 0.66 0.63 0.60 0.57 0.55 1.00
PN 1.88 1.17 1.01 0.90 0.82 0.75 0.71 0.67 0.64 1.00
MSN−D2 1.22 1.00 0.93 0.88 0.83 0.78 0.75 0.72 0.70 1.00
MSN−D1 0.94 0.74 0.68 0.63 0.59 0.56 0.53 0.52 0.51 1.00
Lamp5−Lhx6 1.09 0.90 0.84 0.78 0.75 0.70 0.67 0.63 0.62 1.00
Lamp5 1.78 1.41 1.27 1.18 1.10 1.04 0.97 0.90 0.84 1.00
Foxp2 1.02 0.87 0.83 0.81 0.78 0.75 0.71 0.70 0.68 1.00
Chd7 1.19 0.96 0.88 0.82 0.79 0.73 0.69 0.65 0.63 1.00
CB 1.04 0.94 0.89 0.84 0.80 0.77 0.74 0.71 0.70 1.00
0.0−0.1 0.1−0.2 0.2−0.3 0.3−0.4 0.4−0.5 0.5−0.6 0.6−0.7 0.7−0.8 0.8−0.9 0.9−1.0
Percent
0.010 0.015 0.020
EC−Endothelial cells
L6b 5.96 3.46 2.58 2.14 1.79 1.80 1.37 1.50 1.43 1.00
L6−IT−Car3 11.62 5.74 5.18 3.61 3.33 3.34 2.63 2.47 2.01 1.00
L6−IT 0.19 0.15 0.15 0.16 0.16 0.17 0.18 0.26 0.32 1.00
L6−CT 0.08 0.05 0.06 0.07 0.08 0.10 0.13 0.15 0.24 1.00
L56−NP 0.32 0.26 0.21 0.18 0.18 0.19 0.28 0.32 0.34 1.00
L5−IT 0.14 0.14 0.12 0.11 0.12 0.14 0.16 0.21 0.25 1.00
L5−ET 2.15 1.31 1.24 1.04 1.07 1.10 1.08 1.03 1.18 1.00
L4−IT 2.21 1.06 1.13 1.00 0.94 0.87 0.94 1.19 1.19 1.00
HIP−Misc2 0.05 0.04 0.05 0.04 0.05 0.05 0.07 0.09 0.11 1.00
HIP−Misc1 1.14 0.59 0.56 0.61 0.53 0.62 0.58 0.74 0.66 1.00
DG 0.51 0.47 0.42 0.38 0.43 0.56 0.46 0.65 0.75 1.00
CA3 4.27 2.33 2.00 1.98 1.80 1.69 1.53 1.47 1.47 1.00
CA1 2.58 1.31 1.17 1.35 1.39 1.24 1.22 1.27 1.21 1.00
Amy−Exc 0.20 0.16 0.16 0.16 0.14 0.20 0.20 0.26 0.32 1.00
Vip 0.13 0.12 0.14 0.12 0.14 0.18 0.22 0.29 0.35 1.00
THM−MB 0.08 0.09 0.07 0.08 0.09 0.13 0.12 0.17 0.23 1.00
THM−Inh 0.12 0.10 0.08 0.10 0.11 0.11 0.12 0.15 0.26 1.00
THM−Exc 12.65 4.88 3.42 3.19 2.00 1.73 2.04 1.31 1.58 1.00
SubCtx−Cplx 0.13 0.09 0.08 0.09 0.11 0.11 0.17 0.20 0.25 1.00
Sst 0.10 0.07 0.07 0.08 0.08 0.09 0.11 0.15 0.22 1.00
Sncg 0.10 0.11 0.09 0.10 0.13 0.14 0.14 0.21 0.29 1.00
Pvalb−ChC 0.15 0.11 0.10 0.14 0.13 0.14 0.20 0.21 0.25 1.00
Pvalb 0.07 0.09 0.06 0.08 0.09 0.11 0.10 0.12 0.21 1.00
PN 0.17 0.14 0.12 0.14 0.13 0.15 0.17 0.18 0.26 1.00
MSN−D2 0.10 0.09 0.11 0.11 0.13 0.16 0.19 0.23 0.32 1.00
MSN−D1 0.06 0.04 0.05 0.06 0.08 0.07 0.12 0.13 0.21 1.00
Lamp5−Lhx6 0.08 0.07 0.07 0.08 0.09 0.12 0.11 0.19 0.26 1.00
Lamp5 0.17 0.13 0.12 0.20 0.19 0.17 0.18 0.26 0.34 1.00
Foxp2 0.08 0.07 0.06 0.08 0.10 0.12 0.18 0.20 0.30 1.00
Chd7 0.09 0.08 0.08 0.11 0.10 0.15 0.14 0.21 0.27 1.00
CB 0.10 0.08 0.08 0.08 0.12 0.15 0.18 0.21 0.30 1.00
0.0−0.1 0.1−0.2 0.2−0.3 0.3−0.4 0.4−0.5 0.5−0.6 0.6−0.7 0.7−0.8 0.8−0.9 0.9−1.0
Percent
2e−04 4e−04 6e−04

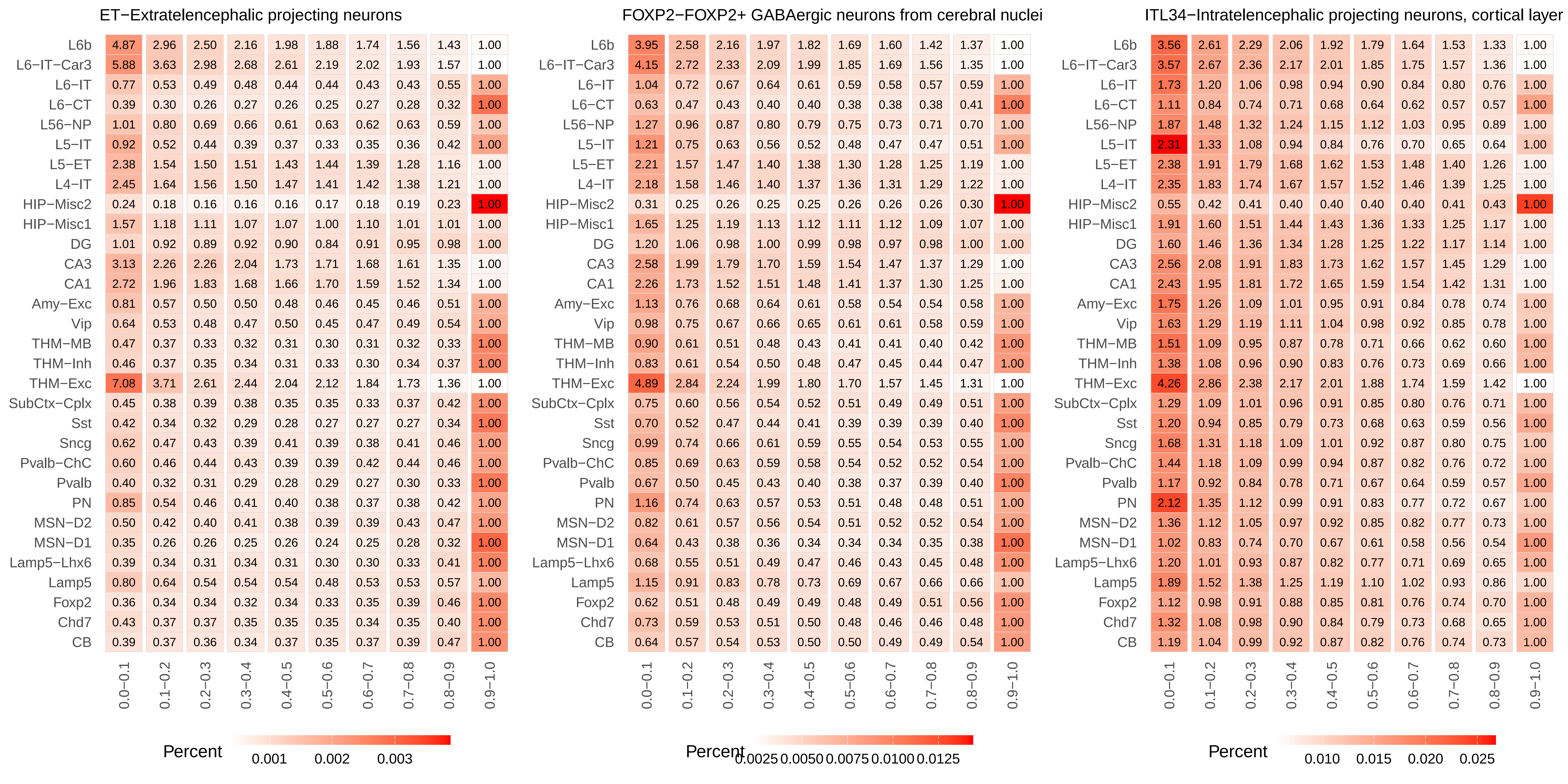
ET−Extratelencephalic projecting neurons
L6b 4.87 2.96 2.50 2.16 1.98 1.88 1.74 1.56 1.43 1.00
L6−IT−Car3 5.88 3.63 2.98 2.68 2.61 2.19 2.02 1.93 1.57 1.00
L6−IT 0.77 0.53 0.49 0.48 0.44 0.44 0.43 0.43 0.55 1.00
L6−CT 0.39 0.30 0.26 0.27 0.26 0.25 0.27 0.28 0.32 1.00
L56−NP 1.01 0.80 0.69 0.66 0.61 0.63 0.62 0.63 0.59 1.00
L5−IT 0.92 0.52 0.44 0.39 0.37 0.33 0.35 0.36 0.42 1.00
L5−ET 2.38 1.54 1.50 1.51 1.43 1.44 1.39 1.28 1.16 1.00
L4−IT 2.45 1.64 1.56 1.50 1.47 1.41 1.42 1.38 1.21 1.00
HIP−Misc2 0.24 0.18 0.16 0.16 0.16 0.17 0.18 0.19 0.23 1.00
HIP−Misc1 1.57 1.18 1.11 1.07 1.07 1.00 1.10 1.01 1.01 1.00
DG 1.01 0.92 0.89 0.92 0.90 0.84 0.91 0.95 0.98 1.00
CA3 3.13 2.26 2.26 2.04 1.73 1.71 1.68 1.61 1.35 1.00
CA1 2.72 1.96 1.83 1.68 1.66 1.70 1.59 1.52 1.34 1.00
Amy−Exc 0.81 0.57 0.50 0.50 0.48 0.46 0.45 0.46 0.51 1.00
Vip 0.64 0.53 0.48 0.47 0.50 0.45 0.47 0.49 0.54 1.00
THM−MB 0.47 0.37 0.33 0.32 0.31 0.30 0.31 0.32 0.33 1.00
THM−Inh 0.46 0.37 0.35 0.34 0.31 0.33 0.30 0.34 0.37 1.00
THM−Exc 7.08 3.71 2.61 2.44 2.04 2.12 1.84 1.73 1.36 1.00
SubCtx−Cplx 0.45 0.38 0.39 0.38 0.35 0.35 0.33 0.37 0.42 1.00
Sst 0.42 0.34 0.32 0.29 0.28 0.27 0.27 0.27 0.34 1.00
Sncg 0.62 0.47 0.43 0.39 0.41 0.39 0.38 0.41 0.46 1.00
Pvalb−ChC 0.60 0.46 0.44 0.43 0.39 0.39 0.42 0.44 0.46 1.00
Pvalb 0.40 0.32 0.31 0.29 0.28 0.29 0.27 0.30 0.33 1.00
PN 0.85 0.54 0.46 0.41 0.40 0.38 0.37 0.38 0.42 1.00
MSN−D2 0.50 0.42 0.40 0.41 0.38 0.39 0.39 0.43 0.47 1.00
MSN−D1 0.35 0.26 0.26 0.25 0.26 0.24 0.25 0.28 0.32 1.00
Lamp5−Lhx6 0.39 0.34 0.31 0.34 0.31 0.30 0.30 0.33 0.41 1.00
Lamp5 0.80 0.64 0.54 0.54 0.54 0.48 0.53 0.53 0.57 1.00
Foxp2 0.36 0.34 0.34 0.32 0.34 0.33 0.35 0.39 0.46 1.00
Chd7 0.43 0.37 0.37 0.35 0.35 0.35 0.34 0.35 0.40 1.00
CB 0.39 0.37 0.36 0.34 0.37 0.35 0.37 0.39 0.47 1.00
0.0−0.1 0.1−0.2 0.2−0.3 0.3−0.4 0.4−0.5 0.5−0.6 0.6−0.7 0.7−0.8 0.8−0.9 0.9−1.0
Percent 0.001 0.002 0.003
FOXP2−FOXP2+ GABAergic neurons from cerebral nuclei
L6b 3.95 2.58 2.16 1.97 1.82 1.69 1.60 1.42 1.37 1.00
L6−IT−Car3 4.15 2.72 2.33 2.09 1.99 1.85 1.69 1.56 1.35 1.00
L6−IT 1.04 0.72 0.67 0.64 0.61 0.59 0.58 0.57 0.59 1.00
L6−CT 0.63 0.47 0.43 0.40 0.40 0.38 0.38 0.38 0.41 1.00
L56−NP 1.27 0.96 0.87 0.80 0.79 0.75 0.73 0.71 0.70 1.00
L5−IT 1.21 0.75 0.63 0.56 0.52 0.48 0.47 0.47 0.51 1.00
L5−ET 2.21 1.57 1.47 1.40 1.38 1.30 1.28 1.25 1.19 1.00
L4−IT 2.18 1.58 1.46 1.40 1.37 1.36 1.31 1.29 1.22 1.00
HIP−Misc2 0.31 0.25 0.26 0.25 0.25 0.26 0.26 0.26 0.30 1.00
HIP−Misc1 1.65 1.25 1.19 1.13 1.12 1.11 1.12 1.09 1.07 1.00
DG 1.20 1.06 0.98 1.00 0.99 0.98 0.97 0.98 1.00 1.00
CA3 2.58 1.99 1.79 1.70 1.59 1.54 1.47 1.37 1.29 1.00
CA1 2.26 1.73 1.52 1.51 1.48 1.41 1.37 1.30 1.25 1.00
Amy−Exc 1.13 0.76 0.68 0.64 0.61 0.58 0.54 0.54 0.58 1.00
Vip 0.98 0.75 0.67 0.66 0.65 0.61 0.61 0.58 0.59 1.00
THM−MB 0.90 0.61 0.51 0.48 0.43 0.41 0.41 0.40 0.42 1.00
THM−Inh 0.83 0.61 0.54 0.50 0.48 0.47 0.45 0.44 0.47 1.00
THM−Exc 4.89 2.84 2.24 1.99 1.80 1.70 1.57 1.45 1.31 1.00
SubCtx−Cplx 0.75 0.60 0.56 0.54 0.52 0.51 0.49 0.49 0.51 1.00
Sst 0.70 0.52 0.47 0.44 0.41 0.39 0.39 0.39 0.40 1.00
Sncg 0.99 0.74 0.66 0.61 0.59 0.55 0.54 0.53 0.55 1.00
Pvalb−ChC 0.85 0.69 0.63 0.59 0.58 0.54 0.52 0.52 0.54 1.00
Pvalb 0.67 0.50 0.45 0.43 0.40 0.38 0.37 0.39 0.40 1.00
PN 1.16 0.74 0.63 0.57 0.53 0.51 0.48 0.48 0.51 1.00
MSN−D2 0.82 0.61 0.57 0.56 0.54 0.51 0.52 0.52 0.54 1.00
MSN−D1 0.64 0.43 0.38 0.36 0.34 0.34 0.34 0.35 0.38 1.00
Lamp5−Lhx6 0.68 0.55 0.51 0.49 0.47 0.46 0.43 0.45 0.48 1.00
Lamp5 1.15 0.91 0.83 0.78 0.73 0.69 0.67 0.66 0.66 1.00
Foxp2 0.62 0.51 0.48 0.49 0.49 0.48 0.49 0.51 0.56 1.00
Chd7 0.73 0.59 0.53 0.51 0.50 0.48 0.46 0.46 0.48 1.00
CB 0.64 0.57 0.54 0.53 0.50 0.50 0.49 0.49 0.54 1.00
0.0−0.1 0.1−0.2 0.2−0.3 0.3−0.4 0.4−0.5 0.5−0.6 0.6−0.7 0.7−0.8 0.8−0.9 0.9−1.0
Percent 0.0025 0.0050 0.0075 0.0100 0.0125
ITL34−Intratelencephalic projecting neurons, cortical layer
L6b 3.56 2.61 2.29 2.06 1.92 1.79 1.64 1.53 1.33 1.00
L6−IT−Car3 3.57 2.67 2.36 2.17 2.01 1.85 1.75 1.57 1.36 1.00
L6−IT 1.73 1.20 1.06 0.98 0.94 0.90 0.84 0.80 0.76 1.00
L6−CT 1.11 0.84 0.74 0.71 0.68 0.64 0.62 0.57 0.57 1.00
L56−NP 1.87 1.48 1.32 1.24 1.15 1.12 1.03 0.95 0.89 1.00
L5−IT 2.31 1.33 1.08 0.94 0.84 0.76 0.70 0.65 0.64 1.00
L5−ET 2.38 1.91 1.79 1.68 1.62 1.53 1.48 1.40 1.26 1.00
L4−IT 2.35 1.83 1.74 1.67 1.57 1.52 1.46 1.39 1.25 1.00
HIP−Misc2 0.55 0.42 0.41 0.40 0.40 0.40 0.40 0.41 0.43 1.00
HIP−Misc1 1.91 1.60 1.51 1.44 1.43 1.36 1.33 1.25 1.17 1.00
DG 1.60 1.46 1.36 1.34 1.28 1.25 1.22 1.17 1.14 1.00
CA3 2.56 2.08 1.91 1.83 1.73 1.62 1.57 1.45 1.29 1.00
CA1 2.43 1.95 1.81 1.72 1.65 1.59 1.54 1.42 1.31 1.00
Amy−Exc 1.75 1.26 1.09 1.01 0.95 0.91 0.84 0.78 0.74 1.00
Vip 1.63 1.29 1.19 1.11 1.04 0.98 0.92 0.85 0.78 1.00
THM−MB 1.51 1.09 0.95 0.87 0.78 0.71 0.66 0.62 0.60 1.00
THM−Inh 1.38 1.08 0.96 0.90 0.83 0.76 0.73 0.69 0.66 1.00
THM−Exc 4.26 2.86 2.38 2.17 2.01 1.88 1.74 1.59 1.42 1.00
SubCtx−Cplx 1.29 1.09 1.01 0.96 0.91 0.85 0.80 0.76 0.71 1.00
Sst 1.20 0.94 0.85 0.79 0.73 0.68 0.63 0.59 0.56 1.00
Sncg 1.68 1.31 1.18 1.09 1.01 0.92 0.87 0.80 0.75 1.00
Pvalb−ChC 1.44 1.18 1.09 0.99 0.94 0.87 0.82 0.76 0.72 1.00
Pvalb 1.17 0.92 0.84 0.78 0.71 0.67 0.64 0.59 0.57 1.00
PN 2.12 1.35 1.12 0.99 0.91 0.83 0.77 0.72 0.67 1.00
MSN−D2 1.36 1.12 1.05 0.97 0.92 0.85 0.82 0.77 0.73 1.00
MSN−D1 1.02 0.83 0.74 0.70 0.67 0.61 0.58 0.56 0.54 1.00
Lamp5−Lhx6 1.20 1.01 0.93 0.87 0.82 0.77 0.71 0.69 0.65 1.00
Lamp5 1.89 1.52 1.38 1.25 1.19 1.10 1.02 0.93 0.86 1.00
Foxp2 1.12 0.98 0.91 0.88 0.85 0.81 0.76 0.74 0.70 1.00
Chd7 1.32 1.08 0.98 0.90 0.84 0.79 0.73 0.68 0.65 1.00
CB 1.19 1.04 0.99 0.92 0.87 0.82 0.76 0.74 0.73 1.00
0.0−0.1 0.1−0.2 0.2−0.3 0.3−0.4 0.4−0.5 0.5−0.6 0.6−0.7 0.7−0.8 0.8−0.9 0.9−1.0
Percent 0.010 0.015 0.020 0.025

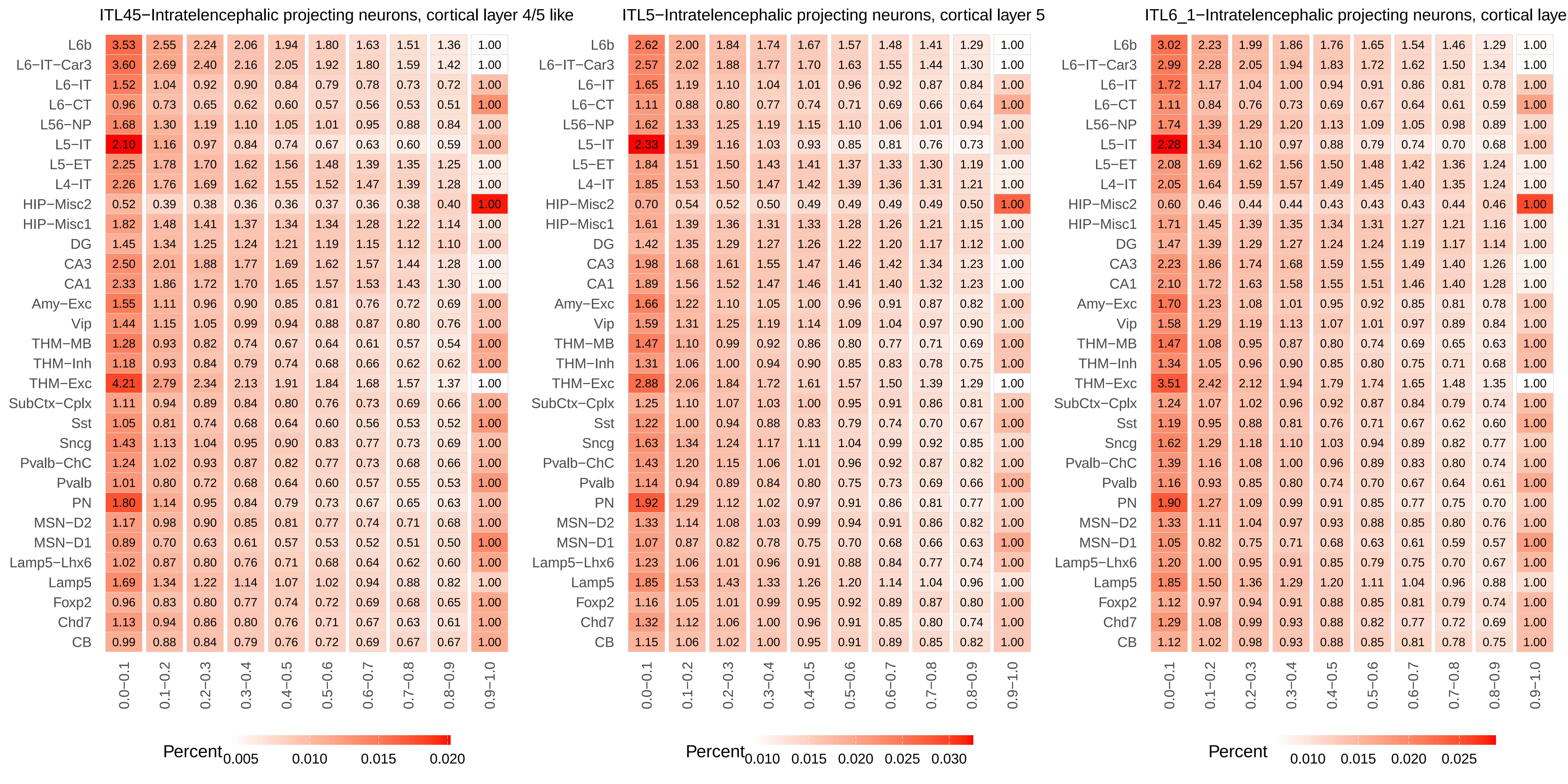
ITL45−Intratelencephalic projecting neurons, cortical layer 4/5 like
L6b 3.53 2.55 2.24 2.06 1.94 1.80 1.63 1.51 1.36 1.00
L6−IT−Car3 3.60 2.69 2.40 2.16 2.05 1.92 1.80 1.59 1.42 1.00
L6−IT 1.52 1.04 0.92 0.90 0.84 0.79 0.78 0.73 0.72 1.00
L6−CT 0.96 0.73 0.65 0.62 0.60 0.57 0.56 0.53 0.51 1.00
L56−NP 1.68 1.30 1.19 1.10 1.05 1.01 0.95 0.88 0.84 1.00
L5−IT 2.10 1.16 0.97 0.84 0.74 0.67 0.63 0.60 0.59 1.00
L5−ET 2.25 1.78 1.70 1.62 1.56 1.48 1.39 1.35 1.25 1.00
L4−IT 2.26 1.76 1.69 1.62 1.55 1.52 1.47 1.39 1.28 1.00
HIP−Misc2 0.52 0.39 0.38 0.36 0.36 0.37 0.36 0.38 0.40 1.00
HIP−Misc1 1.82 1.48 1.41 1.37 1.34 1.34 1.28 1.22 1.14 1.00
DG 1.45 1.34 1.25 1.24 1.21 1.19 1.15 1.12 1.10 1.00
CA3 2.50 2.01 1.88 1.77 1.69 1.62 1.57 1.44 1.28 1.00
CA1 2.33 1.86 1.72 1.70 1.65 1.57 1.53 1.43 1.30 1.00
Amy−Exc 1.55 1.11 0.96 0.90 0.85 0.81 0.76 0.72 0.69 1.00
Vip 1.44 1.15 1.05 0.99 0.94 0.88 0.87 0.80 0.76 1.00
THM−MB 1.28 0.93 0.82 0.74 0.67 0.64 0.61 0.57 0.54 1.00
THM−Inh 1.18 0.93 0.84 0.79 0.74 0.68 0.66 0.62 0.62 1.00
THM−Exc 4.21 2.79 2.34 2.13 1.91 1.84 1.68 1.57 1.37 1.00
SubCtx−Cplx 1.11 0.94 0.89 0.84 0.80 0.76 0.73 0.69 0.66 1.00
Sst 1.05 0.81 0.74 0.68 0.64 0.60 0.56 0.53 0.52 1.00
Sncg 1.43 1.13 1.04 0.95 0.90 0.83 0.77 0.73 0.69 1.00
Pvalb−ChC 1.24 1.02 0.93 0.87 0.82 0.77 0.73 0.68 0.66 1.00
Pvalb 1.01 0.80 0.72 0.68 0.64 0.60 0.57 0.55 0.53 1.00
PN 1.80 1.14 0.95 0.84 0.79 0.73 0.67 0.65 0.63 1.00
MSN−D2 1.17 0.98 0.90 0.85 0.81 0.77 0.74 0.71 0.68 1.00
MSN−D1 0.89 0.70 0.63 0.61 0.57 0.53 0.52 0.51 0.50 1.00
Lamp5−Lhx6 1.02 0.87 0.80 0.76 0.71 0.68 0.64 0.62 0.60 1.00
Lamp5 1.69 1.34 1.22 1.14 1.07 1.02 0.94 0.88 0.82 1.00
Foxp2 0.96 0.83 0.80 0.77 0.74 0.72 0.69 0.68 0.65 1.00
Chd7 1.13 0.94 0.86 0.80 0.76 0.71 0.67 0.63 0.61 1.00
CB 0.99 0.88 0.84 0.79 0.76 0.72 0.69 0.67 0.67 1.00
0.0−0.1 0.1−0.2 0.2−0.3 0.3−0.4 0.4−0.5 0.5−0.6 0.6−0.7 0.7−0.8 0.8−0.9 0.9−1.0
Percent 0.005 0.010 0.015 0.020
ITL5−Intratelencephalic projecting neurons, cortical layer 5
L6b 2.62 2.00 1.84 1.74 1.67 1.57 1.48 1.41 1.29 1.00
L6−IT−Car3 2.57 2.02 1.88 1.77 1.70 1.63 1.55 1.44 1.30 1.00
L6−IT 1.65 1.19 1.10 1.04 1.01 0.96 0.92 0.87 0.84 1.00
L6−CT 1.11 0.88 0.80 0.77 0.74 0.71 0.69 0.66 0.64 1.00
L56−NP 1.62 1.33 1.25 1.19 1.15 1.10 1.06 1.01 0.94 1.00
L5−IT 2.33 1.39 1.16 1.03 0.93 0.85 0.81 0.76 0.73 1.00
L5−ET 1.84 1.51 1.50 1.43 1.41 1.37 1.33 1.30 1.19 1.00
L4−IT 1.85 1.53 1.50 1.47 1.42 1.39 1.36 1.31 1.21 1.00
HIP−Misc2 0.70 0.54 0.52 0.50 0.49 0.49 0.49 0.49 0.50 1.00
HIP−Misc1 1.61 1.39 1.36 1.31 1.33 1.28 1.26 1.21 1.15 1.00
DG 1.42 1.35 1.29 1.27 1.26 1.22 1.20 1.17 1.12 1.00
CA3 1.98 1.68 1.61 1.55 1.47 1.46 1.42 1.34 1.23 1.00
CA1 1.89 1.56 1.52 1.47 1.46 1.41 1.40 1.32 1.23 1.00
Amy−Exc 1.66 1.22 1.10 1.05 1.00 0.96 0.91 0.87 0.82 1.00
Vip 1.59 1.31 1.25 1.19 1.14 1.09 1.04 0.97 0.90 1.00
THM−MB 1.47 1.10 0.99 0.92 0.86 0.80 0.77 0.71 0.69 1.00
THM−Inh 1.31 1.06 1.00 0.94 0.90 0.85 0.83 0.78 0.75 1.00
THM−Exc 2.88 2.06 1.84 1.72 1.61 1.57 1.50 1.39 1.29 1.00
SubCtx−Cplx 1.25 1.10 1.07 1.03 1.00 0.95 0.91 0.86 0.81 1.00
Sst 1.22 1.00 0.94 0.88 0.83 0.79 0.74 0.70 0.67 1.00
Sncg 1.63 1.34 1.24 1.17 1.11 1.04 0.99 0.92 0.85 1.00
Pvalb−ChC 1.43 1.20 1.15 1.06 1.01 0.96 0.92 0.87 0.82 1.00
Pvalb 1.14 0.94 0.89 0.84 0.80 0.75 0.73 0.69 0.66 1.00
PN 1.92 1.29 1.12 1.02 0.97 0.91 0.86 0.81 0.77 1.00
MSN−D2 1.33 1.14 1.08 1.03 0.99 0.94 0.91 0.86 0.82 1.00
MSN−D1 1.07 0.87 0.82 0.78 0.75 0.70 0.68 0.66 0.63 1.00
Lamp5−Lhx6 1.23 1.06 1.01 0.96 0.91 0.88 0.84 0.77 0.74 1.00
Lamp5 1.85 1.53 1.43 1.33 1.26 1.20 1.14 1.04 0.96 1.00
Foxp2 1.16 1.05 1.01 0.99 0.95 0.92 0.89 0.87 0.80 1.00
Chd7 1.32 1.12 1.06 1.00 0.96 0.91 0.85 0.80 0.74 1.00
CB 1.15 1.06 1.02 1.00 0.95 0.91 0.89 0.85 0.82 1.00
0.0−0.1 0.1−0.2 0.2−0.3 0.3−0.4 0.4−0.5 0.5−0.6 0.6−0.7 0.7−0.8 0.8−0.9 0.9−1.0
Percent 0.010 0.015 0.020 0.025 0.030
ITL6_1−Intratelencephalic projecting neurons, cortical laye
L6b 3.02 2.23 1.99 1.86 1.76 1.65 1.54 1.46 1.29 1.00
L6−IT−Car3 2.99 2.28 2.05 1.94 1.83 1.72 1.62 1.50 1.34 1.00
L6−IT 1.72 1.17 1.04 1.00 0.94 0.91 0.86 0.81 0.78 1.00
L6−CT 1.11 0.84 0.76 0.73 0.69 0.67 0.64 0.61 0.59 1.00
L56−NP 1.74 1.39 1.29 1.20 1.13 1.09 1.05 0.98 0.89 1.00
L5−IT 2.28 1.34 1.10 0.97 0.88 0.79 0.74 0.70 0.68 1.00
L5−ET 2.08 1.69 1.62 1.56 1.50 1.48 1.42 1.36 1.24 1.00
L4−IT 2.05 1.64 1.59 1.57 1.49 1.45 1.40 1.35 1.24 1.00
HIP−Misc2 0.60 0.46 0.44 0.44 0.43 0.43 0.43 0.44 0.46 1.00
HIP−Misc1 1.71 1.45 1.39 1.35 1.34 1.31 1.27 1.21 1.16 1.00
DG 1.47 1.39 1.29 1.27 1.24 1.24 1.19 1.17 1.14 1.00
CA3 2.23 1.86 1.74 1.68 1.59 1.55 1.49 1.40 1.26 1.00
CA1 2.10 1.72 1.63 1.58 1.55 1.51 1.46 1.40 1.28 1.00
Amy−Exc 1.70 1.23 1.08 1.01 0.95 0.92 0.85 0.81 0.78 1.00
Vip 1.58 1.29 1.19 1.13 1.07 1.01 0.97 0.89 0.84 1.00
THM−MB 1.47 1.08 0.95 0.87 0.80 0.74 0.69 0.65 0.63 1.00
THM−Inh 1.34 1.05 0.96 0.90 0.85 0.80 0.75 0.71 0.68 1.00
THM−Exc 3.51 2.42 2.12 1.94 1.79 1.74 1.65 1.48 1.35 1.00
SubCtx−Cplx 1.24 1.07 1.02 0.96 0.92 0.87 0.84 0.79 0.74 1.00
Sst 1.19 0.95 0.88 0.81 0.76 0.71 0.67 0.62 0.60 1.00
Sncg 1.62 1.29 1.18 1.10 1.03 0.94 0.89 0.82 0.77 1.00
Pvalb−ChC 1.39 1.16 1.08 1.00 0.96 0.89 0.83 0.80 0.74 1.00
Pvalb 1.16 0.93 0.85 0.80 0.74 0.70 0.67 0.64 0.61 1.00
PN 1.90 1.27 1.09 0.99 0.91 0.85 0.77 0.75 0.70 1.00
MSN−D2 1.33 1.11 1.04 0.97 0.93 0.88 0.85 0.80 0.76 1.00
MSN−D1 1.05 0.82 0.75 0.71 0.68 0.63 0.61 0.59 0.57 1.00
Lamp5−Lhx6 1.20 1.00 0.95 0.91 0.85 0.79 0.75 0.70 0.67 1.00
Lamp5 1.85 1.50 1.36 1.29 1.20 1.11 1.04 0.96 0.88 1.00
Foxp2 1.12 0.97 0.94 0.91 0.88 0.85 0.81 0.79 0.74 1.00
Chd7 1.29 1.08 0.99 0.93 0.88 0.82 0.77 0.72 0.69 1.00
CB 1.12 1.02 0.98 0.93 0.88 0.85 0.81 0.78 0.75 1.00
0.0−0.1 0.1−0.2 0.2−0.3 0.3−0.4 0.4−0.5 0.5−0.6 0.6−0.7 0.7−0.8 0.8−0.9 0.9−1.0
Percent 0.010 0.015 0.020 0.025

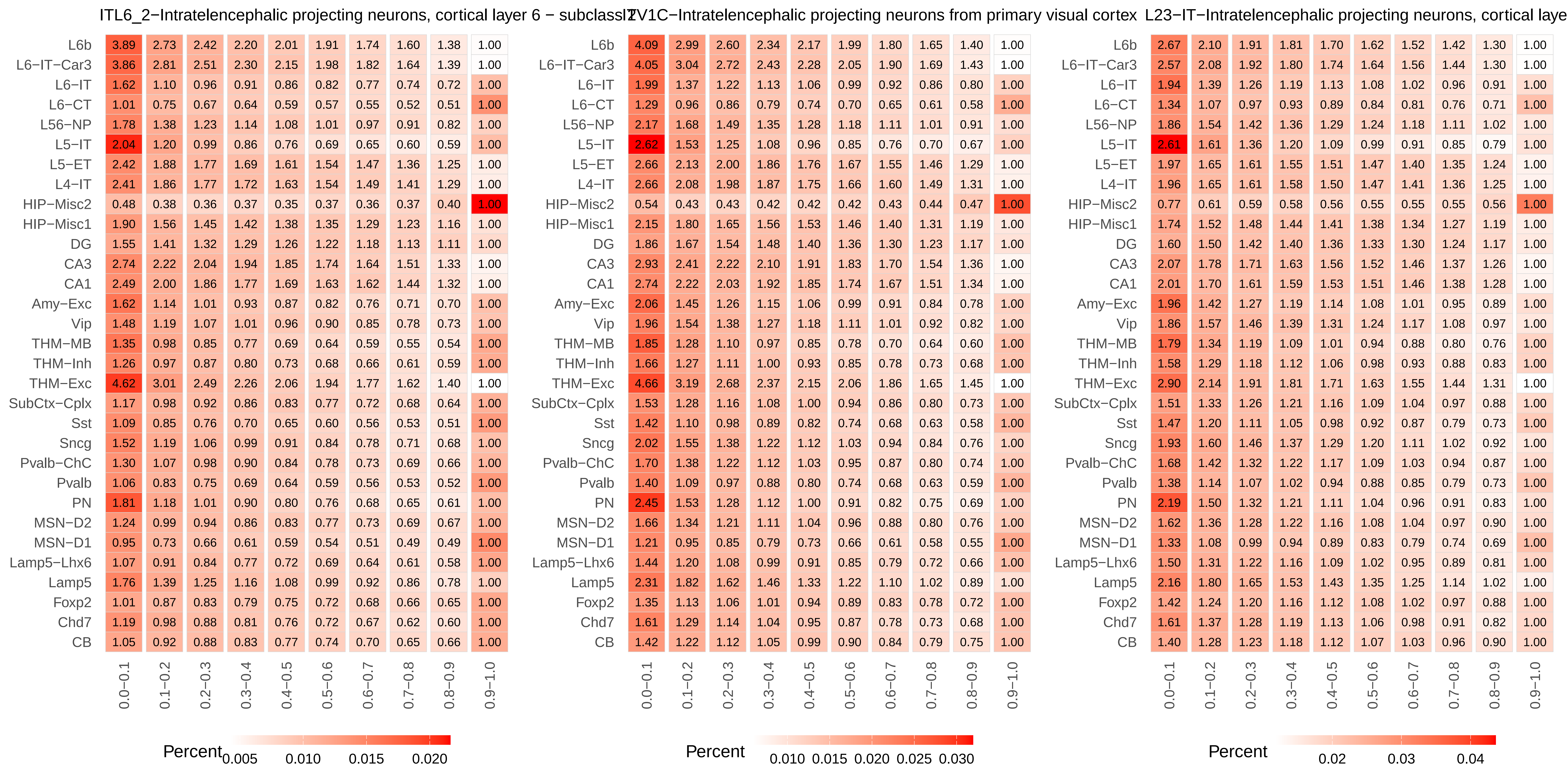
ITL6_2−Intratelencephalic projecting neurons, cortical layer 6 − subclass
L6b 3.89 2.73 2.42 2.20 2.01 1.91 1.74 1.60 1.38 1.00
L6−IT−Car3 3.86 2.81 2.51 2.30 2.15 1.98 1.82 1.64 1.39 1.00
L6−IT 1.62 1.10 0.96 0.91 0.86 0.82 0.77 0.74 0.72 1.00
L6−CT 1.01 0.75 0.67 0.64 0.59 0.57 0.55 0.52 0.51 1.00
L56−NP 1.78 1.38 1.23 1.14 1.08 1.01 0.97 0.91 0.82 1.00
L5−IT 2.04 1.20 0.99 0.86 0.76 0.69 0.65 0.60 0.59 1.00
L5−ET 2.42 1.88 1.77 1.69 1.61 1.54 1.47 1.36 1.25 1.00
L4−IT 2.41 1.86 1.77 1.72 1.63 1.54 1.49 1.41 1.29 1.00
HIP−Misc2 0.48 0.38 0.36 0.37 0.35 0.37 0.36 0.37 0.40 1.00
HIP−Misc1 1.90 1.56 1.45 1.42 1.38 1.35 1.29 1.23 1.16 1.00
DG 1.55 1.41 1.32 1.29 1.26 1.22 1.18 1.13 1.11 1.00
CA3 2.74 2.22 2.04 1.94 1.85 1.74 1.64 1.51 1.33 1.00
CA1 2.49 2.00 1.86 1.77 1.69 1.63 1.62 1.44 1.32 1.00
Amy−Exc 1.62 1.14 1.01 0.93 0.87 0.82 0.76 0.71 0.70 1.00
Vip 1.48 1.19 1.07 1.01 0.96 0.90 0.85 0.78 0.73 1.00
THM−MB 1.35 0.98 0.85 0.77 0.69 0.64 0.59 0.55 0.54 1.00
THM−Inh 1.26 0.97 0.87 0.80 0.73 0.68 0.66 0.61 0.59 1.00
THM−Exc 4.62 3.01 2.49 2.26 2.06 1.94 1.77 1.62 1.40 1.00
SubCtx−Cplx 1.17 0.98 0.92 0.86 0.83 0.77 0.72 0.68 0.64 1.00
Sst 1.09 0.85 0.76 0.70 0.65 0.60 0.56 0.53 0.51 1.00
Sncg 1.52 1.19 1.06 0.99 0.91 0.84 0.78 0.71 0.68 1.00
Pvalb−ChC 1.30 1.07 0.98 0.90 0.84 0.78 0.73 0.69 0.66 1.00
Pvalb 1.06 0.83 0.75 0.69 0.64 0.59 0.56 0.53 0.52 1.00
PN 1.81 1.18 1.01 0.90 0.80 0.76 0.68 0.65 0.61 1.00
MSN−D2 1.24 0.99 0.94 0.86 0.83 0.77 0.73 0.69 0.67 1.00
MSN−D1 0.95 0.73 0.66 0.61 0.59 0.54 0.51 0.49 0.49 1.00
Lamp5−Lhx6 1.07 0.91 0.84 0.77 0.72 0.69 0.64 0.61 0.58 1.00
Lamp5 1.76 1.39 1.25 1.16 1.08 0.99 0.92 0.86 0.78 1.00
Foxp2 1.01 0.87 0.83 0.79 0.75 0.72 0.68 0.66 0.65 1.00
Chd7 1.19 0.98 0.88 0.81 0.76 0.72 0.67 0.62 0.60 1.00
CB 1.05 0.92 0.88 0.83 0.77 0.74 0.70 0.65 0.66 1.00
0.0−0.1 0.1−0.2 0.2−0.3 0.3−0.4 0.4−0.5 0.5−0.6 0.6−0.7 0.7−0.8 0.8−0.9 0.9−1.0
Percent 0.005 0.010 0.015 0.020
ITV1C−Intratelencephalic projecting neurons from primary visual cortex
L6b 4.09 2.99 2.60 2.34 2.17 1.99 1.80 1.65 1.40 1.00
L6−IT−Car3 4.05 3.04 2.72 2.43 2.28 2.05 1.90 1.69 1.43 1.00
L6−IT 1.99 1.37 1.22 1.13 1.06 0.99 0.92 0.86 0.80 1.00
L6−CT 1.29 0.96 0.86 0.79 0.74 0.70 0.65 0.61 0.58 1.00
L56−NP 2.17 1.68 1.49 1.35 1.28 1.18 1.11 1.01 0.91 1.00
L5−IT 2.62 1.53 1.25 1.08 0.96 0.85 0.76 0.70 0.67 1.00
L5−ET 2.66 2.13 2.00 1.86 1.76 1.67 1.55 1.46 1.29 1.00
L4−IT 2.66 2.08 1.98 1.87 1.75 1.66 1.60 1.49 1.31 1.00
HIP−Misc2 0.54 0.43 0.43 0.42 0.42 0.42 0.43 0.44 0.47 1.00
HIP−Misc1 2.15 1.80 1.65 1.56 1.53 1.46 1.40 1.31 1.19 1.00
DG 1.86 1.67 1.54 1.48 1.40 1.36 1.30 1.23 1.17 1.00
CA3 2.93 2.41 2.22 2.10 1.91 1.83 1.70 1.54 1.36 1.00
CA1 2.74 2.22 2.03 1.92 1.85 1.74 1.67 1.51 1.34 1.00
Amy−Exc 2.06 1.45 1.26 1.15 1.06 0.99 0.91 0.84 0.78 1.00
Vip 1.96 1.54 1.38 1.27 1.18 1.11 1.01 0.92 0.82 1.00
THM−MB 1.85 1.28 1.10 0.97 0.85 0.78 0.70 0.64 0.60 1.00
THM−Inh 1.66 1.27 1.11 1.00 0.93 0.85 0.78 0.73 0.68 1.00
THM−Exc 4.66 3.19 2.68 2.37 2.15 2.06 1.86 1.65 1.45 1.00
SubCtx−Cplx 1.53 1.28 1.16 1.08 1.00 0.94 0.86 0.80 0.73 1.00
Sst 1.42 1.10 0.98 0.89 0.82 0.74 0.68 0.63 0.58 1.00
Sncg 2.02 1.55 1.38 1.22 1.12 1.03 0.94 0.84 0.76 1.00
Pvalb−ChC 1.70 1.38 1.22 1.12 1.03 0.95 0.87 0.80 0.74 1.00
Pvalb 1.40 1.09 0.97 0.88 0.80 0.74 0.68 0.63 0.59 1.00
PN 2.45 1.53 1.28 1.12 1.00 0.91 0.82 0.75 0.69 1.00
MSN−D2 1.66 1.34 1.21 1.11 1.04 0.96 0.88 0.80 0.76 1.00
MSN−D1 1.21 0.95 0.85 0.79 0.73 0.66 0.61 0.58 0.55 1.00
Lamp5−Lhx6 1.44 1.20 1.08 0.99 0.91 0.85 0.79 0.72 0.66 1.00
Lamp5 2.31 1.82 1.62 1.46 1.33 1.22 1.10 1.02 0.89 1.00
Foxp2 1.35 1.13 1.06 1.01 0.94 0.89 0.83 0.78 0.72 1.00
Chd7 1.61 1.29 1.14 1.04 0.95 0.87 0.78 0.73 0.68 1.00
CB 1.42 1.22 1.12 1.05 0.99 0.90 0.84 0.79 0.75 1.00
0.0−0.1 0.1−0.2 0.2−0.3 0.3−0.4 0.4−0.5 0.5−0.6 0.6−0.7 0.7−0.8 0.8−0.9 0.9−1.0
Percent 0.010 0.015 0.020 0.025 0.030
L23−IT−Intratelencephalic projecting neurons, cortical laye
L6b 2.67 2.10 1.91 1.81 1.70 1.62 1.52 1.42 1.30 1.00
L6−IT−Car3 2.57 2.08 1.92 1.80 1.74 1.64 1.56 1.44 1.30 1.00
L6−IT 1.94 1.39 1.26 1.19 1.13 1.08 1.02 0.96 0.91 1.00
L6−CT 1.34 1.07 0.97 0.93 0.89 0.84 0.81 0.76 0.71 1.00
L56−NP 1.86 1.54 1.42 1.36 1.29 1.24 1.18 1.11 1.02 1.00
L5−IT 2.61 1.61 1.36 1.20 1.09 0.99 0.91 0.85 0.79 1.00
L5−ET 1.97 1.65 1.61 1.55 1.51 1.47 1.40 1.35 1.24 1.00
L4−IT 1.96 1.65 1.61 1.58 1.50 1.47 1.41 1.36 1.25 1.00
HIP−Misc2 0.77 0.61 0.59 0.58 0.56 0.55 0.55 0.55 0.56 1.00
HIP−Misc1 1.74 1.52 1.48 1.44 1.41 1.38 1.34 1.27 1.19 1.00
DG 1.60 1.50 1.42 1.40 1.36 1.33 1.30 1.24 1.17 1.00
CA3 2.07 1.78 1.71 1.63 1.56 1.52 1.46 1.37 1.26 1.00
CA1 2.01 1.70 1.61 1.59 1.53 1.51 1.46 1.38 1.28 1.00
Amy−Exc 1.96 1.42 1.27 1.19 1.14 1.08 1.01 0.95 0.89 1.00
Vip 1.86 1.57 1.46 1.39 1.31 1.24 1.17 1.08 0.97 1.00
THM−MB 1.79 1.34 1.19 1.09 1.01 0.94 0.88 0.80 0.76 1.00
THM−Inh 1.58 1.29 1.18 1.12 1.06 0.98 0.93 0.88 0.83 1.00
THM−Exc 2.90 2.14 1.91 1.81 1.71 1.63 1.55 1.44 1.31 1.00
SubCtx−Cplx 1.51 1.33 1.26 1.21 1.16 1.09 1.04 0.97 0.88 1.00
Sst 1.47 1.20 1.11 1.05 0.98 0.92 0.87 0.79 0.73 1.00
Sncg 1.93 1.60 1.46 1.37 1.29 1.20 1.11 1.02 0.92 1.00
Pvalb−ChC 1.68 1.42 1.32 1.22 1.17 1.09 1.03 0.94 0.87 1.00
Pvalb 1.38 1.14 1.07 1.02 0.94 0.88 0.85 0.79 0.73 1.00
PN 2.19 1.50 1.32 1.21 1.11 1.04 0.96 0.91 0.83 1.00
MSN−D2 1.62 1.36 1.28 1.22 1.16 1.08 1.04 0.97 0.90 1.00
MSN−D1 1.33 1.08 0.99 0.94 0.89 0.83 0.79 0.74 0.69 1.00
Lamp5−Lhx6 1.50 1.31 1.22 1.16 1.09 1.02 0.95 0.89 0.81 1.00
Lamp5 2.16 1.80 1.65 1.53 1.43 1.35 1.25 1.14 1.02 1.00
Foxp2 1.42 1.24 1.20 1.16 1.12 1.08 1.02 0.97 0.88 1.00
Chd7 1.61 1.37 1.28 1.19 1.13 1.06 0.98 0.91 0.82 1.00
CB 1.40 1.28 1.23 1.18 1.12 1.07 1.03 0.96 0.90 1.00
0.0−0.1 0.1−0.2 0.2−0.3 0.3−0.4 0.4−0.5 0.5−0.6 0.6−0.7 0.7−0.8 0.8−0.9 0.9−1.0
Percent 0.02 0.03 0.04

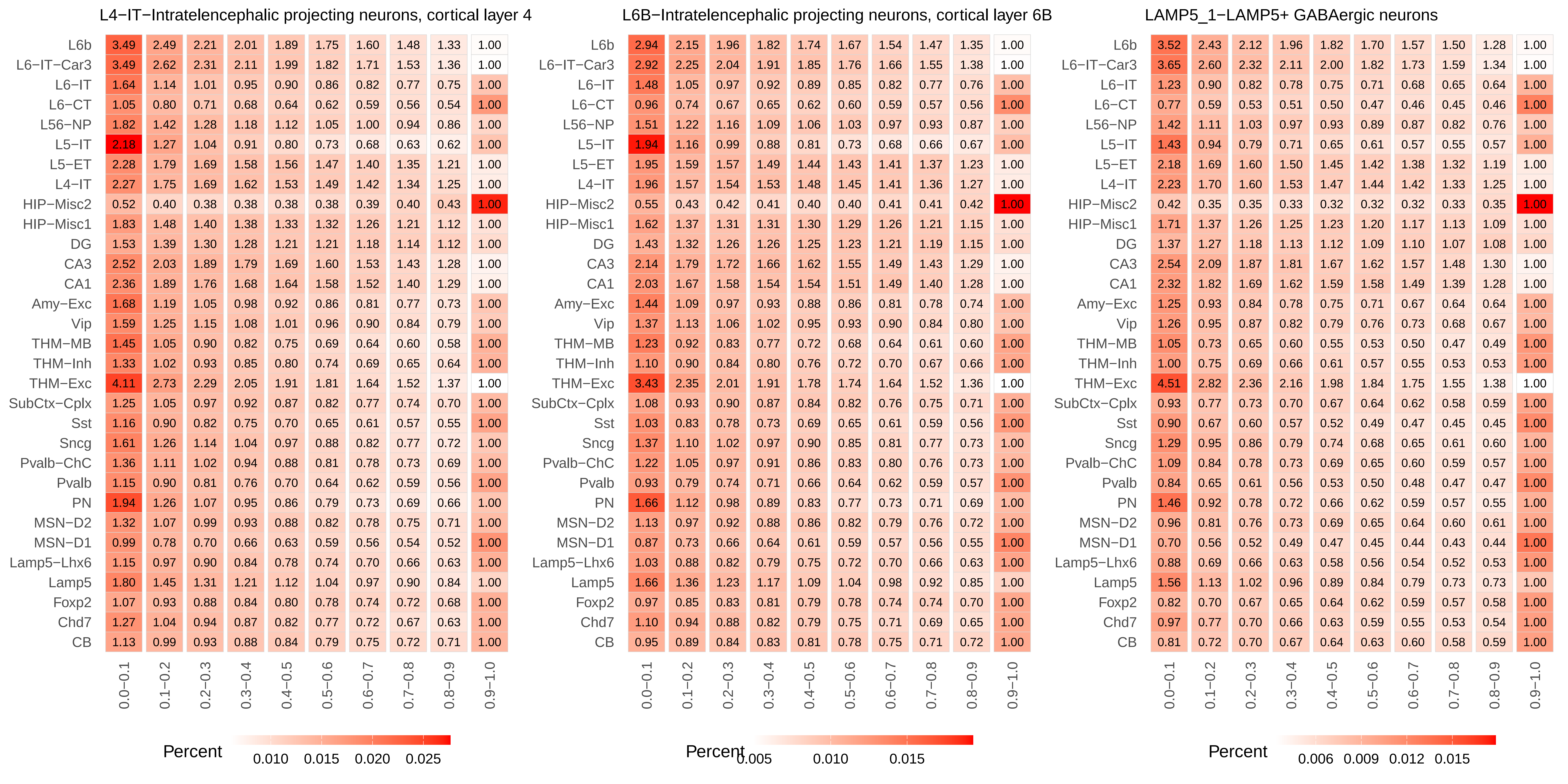
L4−IT−Intratelencephalic projecting neurons, cortical layer 4
L6B−Intratelencephalic projecting neurons, cortical layer 6B
LAMP5_1−LAMP5+ GABAergic neurons
L6b
L6−IT−Car3
L6−IT
L6−CT
L56−NP
L5−IT
L5−ET
L4−IT
HIP−Misc2
HIP−Misc1
DG
CA3
CA1
Amy−Exc
Vip
THM−MB
THM−Inh
THM−Exc
SubCtx−Cplx
Sst
Sncg
Pvalb−ChC
Pvalb
PN
MSN−D2
MSN−D1
Lamp5−Lhx6
Lamp5
Foxp2
Chd7
CB
0.0−0.1
0.1−0.2
0.2−0.3
0.3−0.4
0.4−0.5
0.5−0.6
0.6−0.7
0.7−0.8
0.8−0.9
0.9−1.0
Percent
0.010 0.015 0.020 0.025
Percent
0.005 0.010 0.015
Percent
0.006 0.009 0.012 0.015

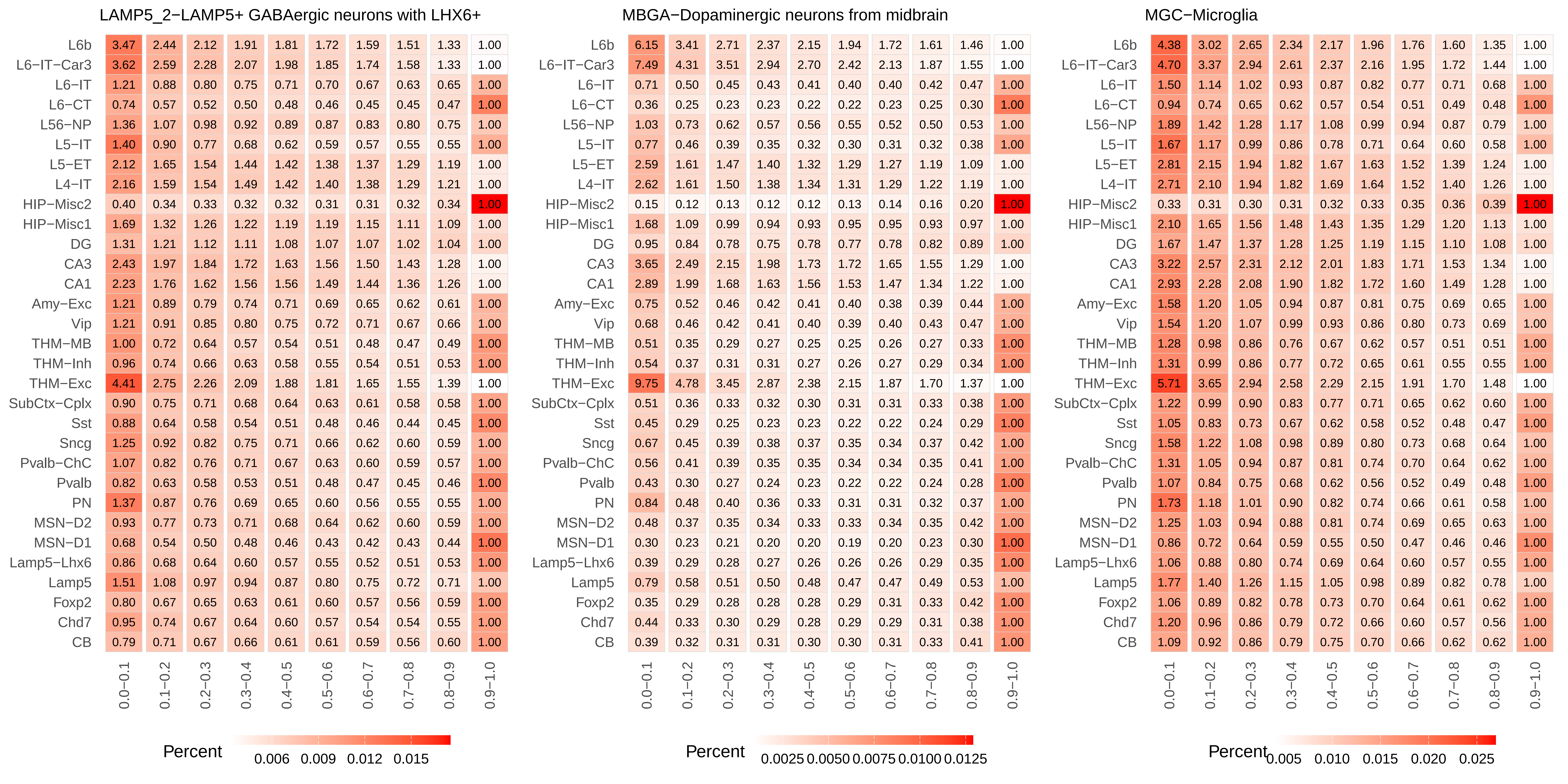
LAMP5_2−LAMP5+ GABAergic neurons with LHX6+
MBGA−Dopaminergic neurons from midbrain
MGC−Microglia
L6b
L6−IT−Car3
L6−IT
L6−CT
L56−NP
L5−IT
L5−ET
L4−IT
HIP−Misc2
HIP−Misc1
DG
CA3
CA1
Amy−Exc
Vip
THM−MB
THM−Inh
THM−Exc
SubCtx−Cplx
Sst
Sncg
Pvalb−ChC
Pvalb
PN
MSN−D2
MSN−D1
Lamp5−Lhx6
Lamp5
Foxp2
Chd7
CB
0.0−0.1
0.1−0.2
0.2−0.3
0.3−0.4
0.4−0.5
0.5−0.6
0.6−0.7
0.7−0.8
0.8−0.9
0.9−1.0
Percent
0.006 0.009 0.012 0.015
Percent
0.0025 0.0050 0.0075 0.0100 0.0125
Percent
0.005 0.010 0.015 0.020 0.025

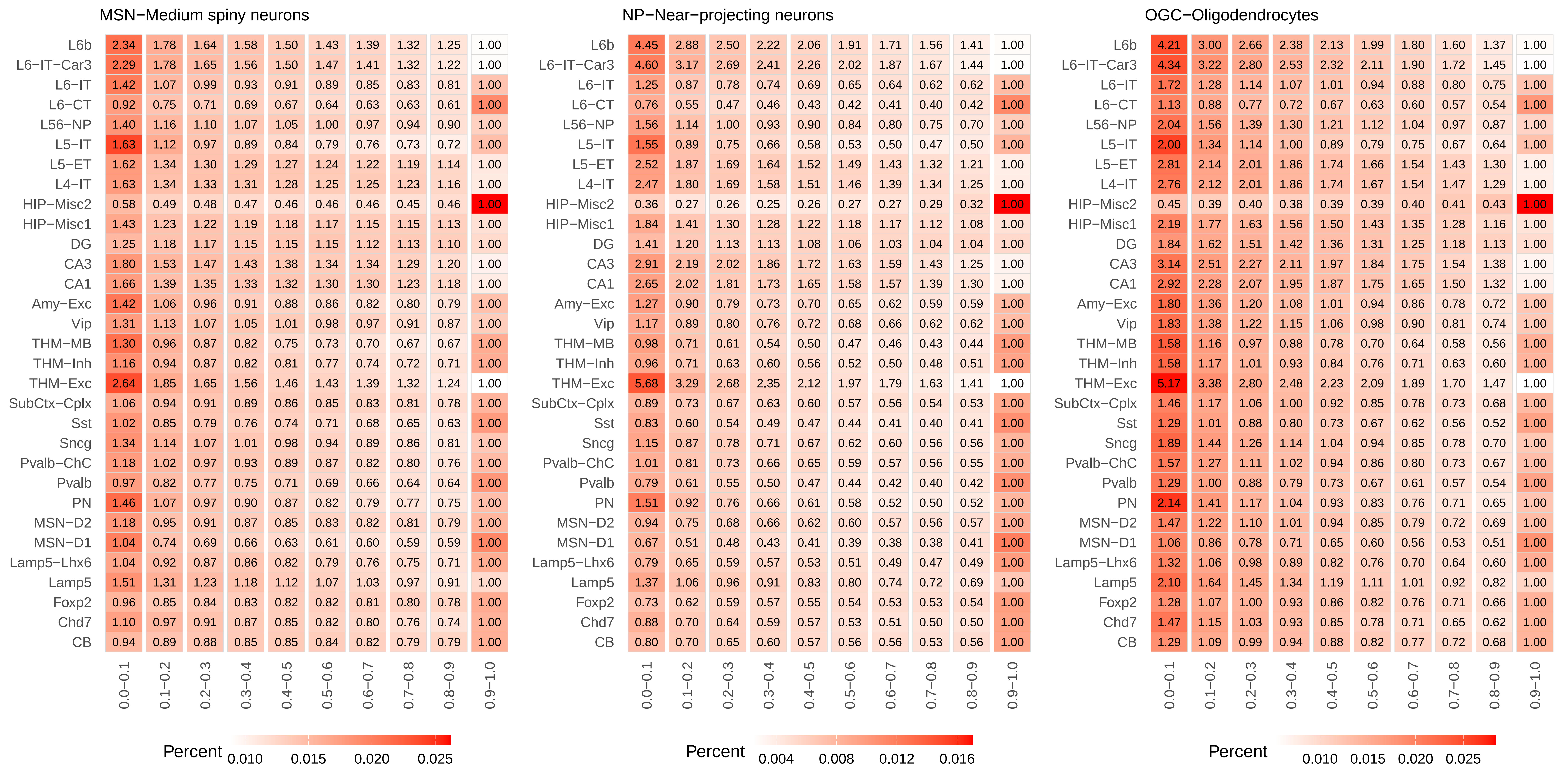
MSN−Medium spiny neurons
L6b 2.34 1.78 1.64 1.58 1.50 1.43 1.39 1.32 1.25 1.00
L6−IT−Car3 2.29 1.78 1.65 1.56 1.50 1.47 1.41 1.32 1.22 1.00
L6−IT 1.42 1.07 0.99 0.93 0.91 0.89 0.85 0.83 0.81 1.00
L6−CT 0.92 0.75 0.71 0.69 0.67 0.64 0.63 0.63 0.61 1.00
L56−NP 1.40 1.16 1.10 1.07 1.05 1.00 0.97 0.94 0.90 1.00
L5−IT 1.63 1.12 0.97 0.89 0.84 0.79 0.76 0.73 0.72 1.00
L5−ET 1.62 1.34 1.30 1.29 1.27 1.24 1.22 1.19 1.14 1.00
L4−IT 1.63 1.34 1.33 1.31 1.28 1.25 1.25 1.23 1.16 1.00
HIP−Misc2 0.58 0.49 0.48 0.47 0.46 0.46 0.46 0.45 0.46 1.00
HIP−Misc1 1.43 1.23 1.22 1.19 1.18 1.17 1.15 1.15 1.13 1.00
DG 1.25 1.18 1.17 1.15 1.15 1.15 1.12 1.13 1.10 1.00
CA3 1.80 1.53 1.47 1.43 1.38 1.34 1.34 1.29 1.20 1.00
CA1 1.66 1.39 1.35 1.33 1.32 1.30 1.30 1.23 1.18 1.00
Amy−Exc 1.42 1.06 0.96 0.91 0.88 0.86 0.82 0.80 0.79 1.00
Vip 1.31 1.13 1.07 1.05 1.01 0.98 0.97 0.91 0.87 1.00
THM−MB 1.30 0.96 0.87 0.82 0.75 0.73 0.70 0.67 0.67 1.00
THM−Inh 1.16 0.94 0.87 0.82 0.81 0.77 0.74 0.72 0.71 1.00
THM−Exc 2.64 1.85 1.65 1.56 1.46 1.43 1.39 1.32 1.24 1.00
SubCtx−Cplx 1.06 0.94 0.91 0.89 0.86 0.85 0.83 0.81 0.78 1.00
Sst 1.02 0.85 0.79 0.76 0.74 0.71 0.68 0.65 0.63 1.00
Sncg 1.34 1.14 1.07 1.01 0.98 0.94 0.89 0.86 0.81 1.00
Pvalb−ChC 1.18 1.02 0.97 0.93 0.89 0.87 0.82 0.80 0.76 1.00
Pvalb 0.97 0.82 0.77 0.75 0.71 0.69 0.66 0.64 0.64 1.00
PN 1.46 1.07 0.97 0.90 0.87 0.82 0.79 0.77 0.75 1.00
MSN−D2 1.18 0.95 0.91 0.87 0.85 0.83 0.82 0.81 0.79 1.00
MSN−D1 1.04 0.74 0.69 0.66 0.63 0.61 0.60 0.59 0.59 1.00
Lamp5−Lhx6 1.04 0.92 0.87 0.86 0.82 0.79 0.76 0.75 0.71 1.00
Lamp5 1.51 1.31 1.23 1.18 1.12 1.07 1.03 0.97 0.91 1.00
Foxp2 0.96 0.85 0.84 0.83 0.82 0.82 0.81 0.80 0.78 1.00
Chd7 1.10 0.97 0.91 0.87 0.85 0.82 0.80 0.76 0.74 1.00
CB 0.94 0.89 0.88 0.85 0.85 0.84 0.82 0.79 0.79 1.00
0.0−0.1 0.1−0.2 0.2−0.3 0.3−0.4 0.4−0.5 0.5−0.6 0.6−0.7 0.7−0.8 0.8−0.9 0.9−1.0
Percent 0.010 0.015 0.020 0.025
NP−Near−projecting neurons
L6b 4.45 2.88 2.50 2.22 2.06 1.91 1.71 1.56 1.41 1.00
L6−IT−Car3 4.60 3.17 2.69 2.41 2.26 2.02 1.87 1.67 1.44 1.00
L6−IT 1.25 0.87 0.78 0.74 0.69 0.65 0.64 0.62 0.62 1.00
L6−CT 0.76 0.55 0.47 0.46 0.43 0.42 0.41 0.40 0.42 1.00
L56−NP 1.56 1.14 1.00 0.93 0.90 0.84 0.80 0.75 0.70 1.00
L5−IT 1.55 0.89 0.75 0.66 0.58 0.53 0.50 0.47 0.50 1.00
L5−ET 2.52 1.87 1.69 1.64 1.52 1.49 1.43 1.32 1.21 1.00
L4−IT 2.47 1.80 1.69 1.58 1.51 1.46 1.39 1.34 1.25 1.00
HIP−Misc2 0.36 0.27 0.26 0.25 0.26 0.27 0.27 0.29 0.32 1.00
HIP−Misc1 1.84 1.41 1.30 1.28 1.22 1.18 1.17 1.12 1.08 1.00
DG 1.41 1.20 1.13 1.13 1.08 1.06 1.03 1.04 1.04 1.00
CA3 2.91 2.19 2.02 1.86 1.72 1.63 1.59 1.43 1.25 1.00
CA1 2.65 2.02 1.81 1.73 1.65 1.58 1.57 1.39 1.30 1.00
Amy−Exc 1.27 0.90 0.79 0.73 0.70 0.65 0.62 0.59 0.59 1.00
Vip 1.17 0.89 0.80 0.76 0.72 0.68 0.66 0.62 0.62 1.00
THM−MB 0.98 0.71 0.61 0.54 0.50 0.47 0.46 0.43 0.44 1.00
THM−Inh 0.96 0.71 0.63 0.60 0.56 0.52 0.50 0.48 0.51 1.00
THM−Exc 5.68 3.29 2.68 2.35 2.12 1.97 1.79 1.63 1.41 1.00
SubCtx−Cplx 0.89 0.73 0.67 0.63 0.60 0.57 0.56 0.54 0.53 1.00
Sst 0.83 0.60 0.54 0.49 0.47 0.44 0.41 0.40 0.41 1.00
Sncg 1.15 0.87 0.78 0.71 0.67 0.62 0.60 0.56 0.56 1.00
Pvalb−ChC 1.01 0.81 0.73 0.66 0.65 0.59 0.57 0.56 0.55 1.00
Pvalb 0.79 0.61 0.55 0.50 0.47 0.44 0.42 0.40 0.42 1.00
PN 1.51 0.92 0.76 0.66 0.61 0.58 0.52 0.50 0.52 1.00
MSN−D2 0.94 0.75 0.68 0.66 0.62 0.60 0.57 0.56 0.57 1.00
MSN−D1 0.67 0.51 0.48 0.43 0.41 0.39 0.38 0.38 0.41 1.00
Lamp5−Lhx6 0.79 0.65 0.59 0.57 0.53 0.51 0.49 0.47 0.49 1.00
Lamp5 1.37 1.06 0.96 0.91 0.83 0.80 0.74 0.72 0.69 1.00
Foxp2 0.73 0.62 0.59 0.57 0.55 0.54 0.53 0.53 0.54 1.00
Chd7 0.88 0.70 0.64 0.59 0.57 0.53 0.51 0.50 0.50 1.00
CB 0.80 0.70 0.65 0.60 0.57 0.56 0.56 0.53 0.56 1.00
0.0−0.1 0.1−0.2 0.2−0.3 0.3−0.4 0.4−0.5 0.5−0.6 0.6−0.7 0.7−0.8 0.8−0.9 0.9−1.0
Percent 0.004 0.008 0.012 0.016
OGC−Oligodendrocytes
L6b 4.21 3.00 2.66 2.38 2.13 1.99 1.80 1.60 1.37 1.00
L6−IT−Car3 4.34 3.22 2.80 2.53 2.32 2.11 1.90 1.72 1.45 1.00
L6−IT 1.72 1.28 1.14 1.07 1.01 0.94 0.88 0.80 0.75 1.00
L6−CT 1.13 0.88 0.77 0.72 0.67 0.63 0.60 0.57 0.54 1.00
L56−NP 2.04 1.56 1.39 1.30 1.21 1.12 1.04 0.97 0.87 1.00
L5−IT 2.00 1.34 1.14 1.00 0.89 0.79 0.75 0.67 0.64 1.00
L5−ET 2.81 2.14 2.01 1.86 1.74 1.66 1.54 1.43 1.30 1.00
L4−IT 2.76 2.12 2.01 1.86 1.74 1.67 1.54 1.47 1.29 1.00
HIP−Misc2 0.45 0.39 0.40 0.38 0.39 0.39 0.40 0.41 0.43 1.00
HIP−Misc1 2.19 1.77 1.63 1.56 1.50 1.43 1.35 1.28 1.16 1.00
DG 1.84 1.62 1.51 1.42 1.36 1.31 1.25 1.18 1.13 1.00
CA3 3.14 2.51 2.27 2.11 1.97 1.84 1.75 1.54 1.38 1.00
CA1 2.92 2.28 2.07 1.95 1.87 1.75 1.65 1.50 1.32 1.00
Amy−Exc 1.80 1.36 1.20 1.08 1.01 0.94 0.86 0.78 0.72 1.00
Vip 1.83 1.38 1.22 1.15 1.06 0.98 0.90 0.81 0.74 1.00
THM−MB 1.58 1.16 0.97 0.88 0.78 0.70 0.64 0.58 0.56 1.00
THM−Inh 1.58 1.17 1.01 0.93 0.84 0.76 0.71 0.63 0.60 1.00
THM−Exc 5.17 3.38 2.80 2.48 2.23 2.09 1.89 1.70 1.47 1.00
SubCtx−Cplx 1.46 1.17 1.06 1.00 0.92 0.85 0.78 0.73 0.68 1.00
Sst 1.29 1.01 0.88 0.80 0.73 0.67 0.62 0.56 0.52 1.00
Sncg 1.89 1.44 1.26 1.14 1.04 0.94 0.85 0.78 0.70 1.00
Pvalb−ChC 1.57 1.27 1.11 1.02 0.94 0.86 0.80 0.73 0.67 1.00
Pvalb 1.29 1.00 0.88 0.79 0.73 0.67 0.61 0.57 0.54 1.00
PN 2.14 1.41 1.17 1.04 0.93 0.83 0.76 0.71 0.65 1.00
MSN−D2 1.47 1.22 1.10 1.01 0.94 0.85 0.79 0.72 0.69 1.00
MSN−D1 1.06 0.86 0.78 0.71 0.65 0.60 0.56 0.53 0.51 1.00
Lamp5−Lhx6 1.32 1.06 0.98 0.89 0.82 0.76 0.70 0.64 0.60 1.00
Lamp5 2.10 1.64 1.45 1.34 1.19 1.11 1.01 0.92 0.82 1.00
Foxp2 1.28 1.07 1.00 0.93 0.86 0.82 0.76 0.71 0.66 1.00
Chd7 1.47 1.15 1.03 0.93 0.85 0.78 0.71 0.65 0.62 1.00
CB 1.29 1.09 0.99 0.94 0.88 0.82 0.77 0.72 0.68 1.00
0.0−0.1 0.1−0.2 0.2−0.3 0.3−0.4 0.4−0.5 0.5−0.6 0.6−0.7 0.7−0.8 0.8−0.9 0.9−1.0
Percent 0.010 0.015 0.020 0.025

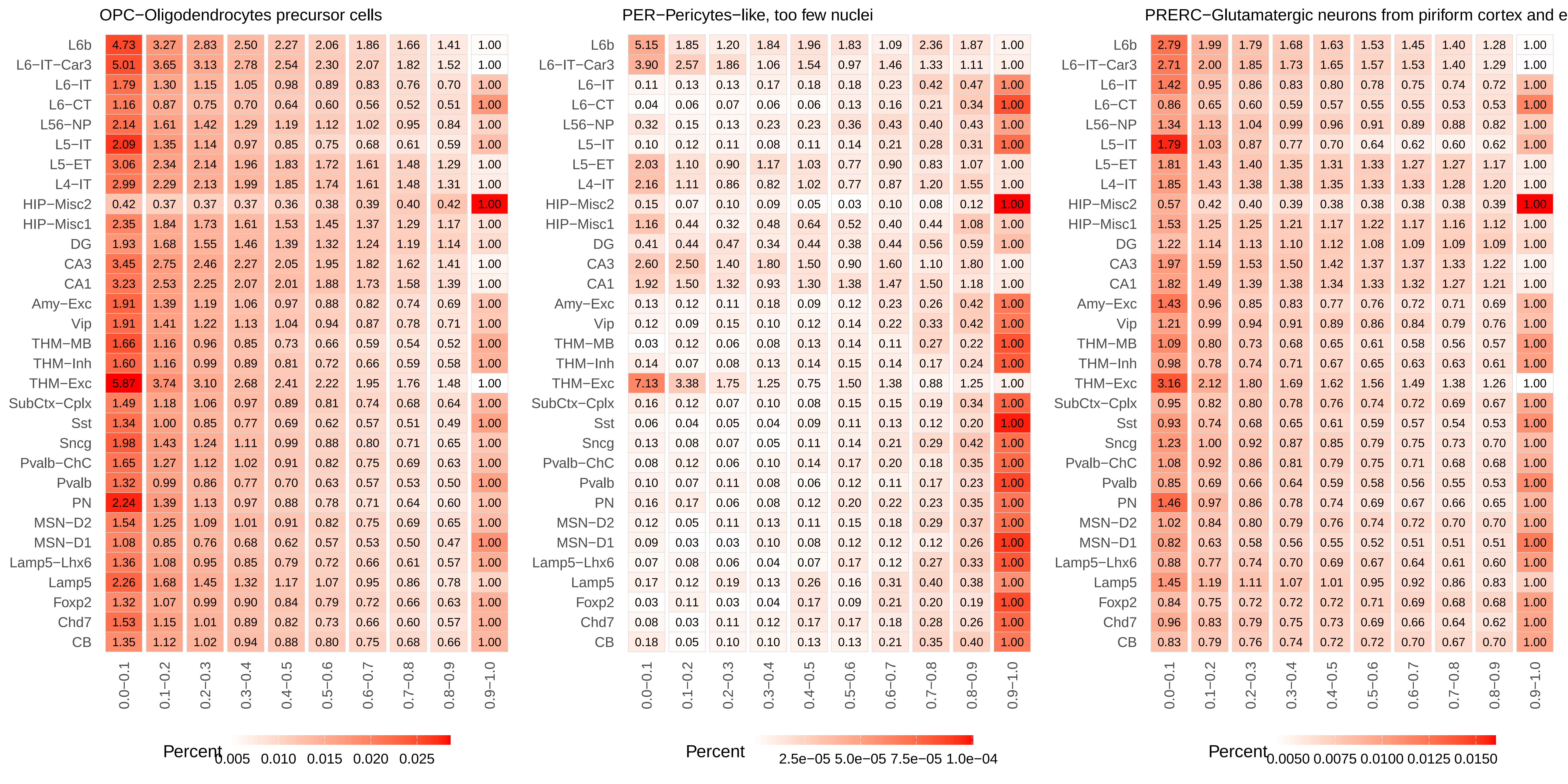

OPC−Oligodendrocytes precursor cells
PER−Pericytes−like, too few nuclei
PRERC−Glutamatergic neurons from piriform cortex and e
L6b
L6−IT−Car3
L6−IT
L6−CT
L56−NP
L5−IT
L5−ET
L4−IT
HIP−Misc2
HIP−Misc1
DG
CA3
CA1
Amy−Exc
Vip
THM−MB
THM−Inh
THM−Exc
SubCtx−Cplx
Sst
Sncg
Pvalb−ChC
Pvalb
PN
MSN−D2
MSN−D1
Lamp5−Lhx6
Lamp5
Foxp2
Chd7
CB
0.0−0.1
0.1−0.2
0.2−0.3
0.3−0.4
0.4−0.5
0.5−0.6
0.6−0.7
0.7−0.8
0.8−0.9
0.9−1.0
Percent
0.005 0.010 0.015 0.020 0.025
Percent
2.5e−05 5.0e−05 7.5e−05 1.0e−04
Percent
0.0050 0.0075 0.0100 0.0125 0.0150

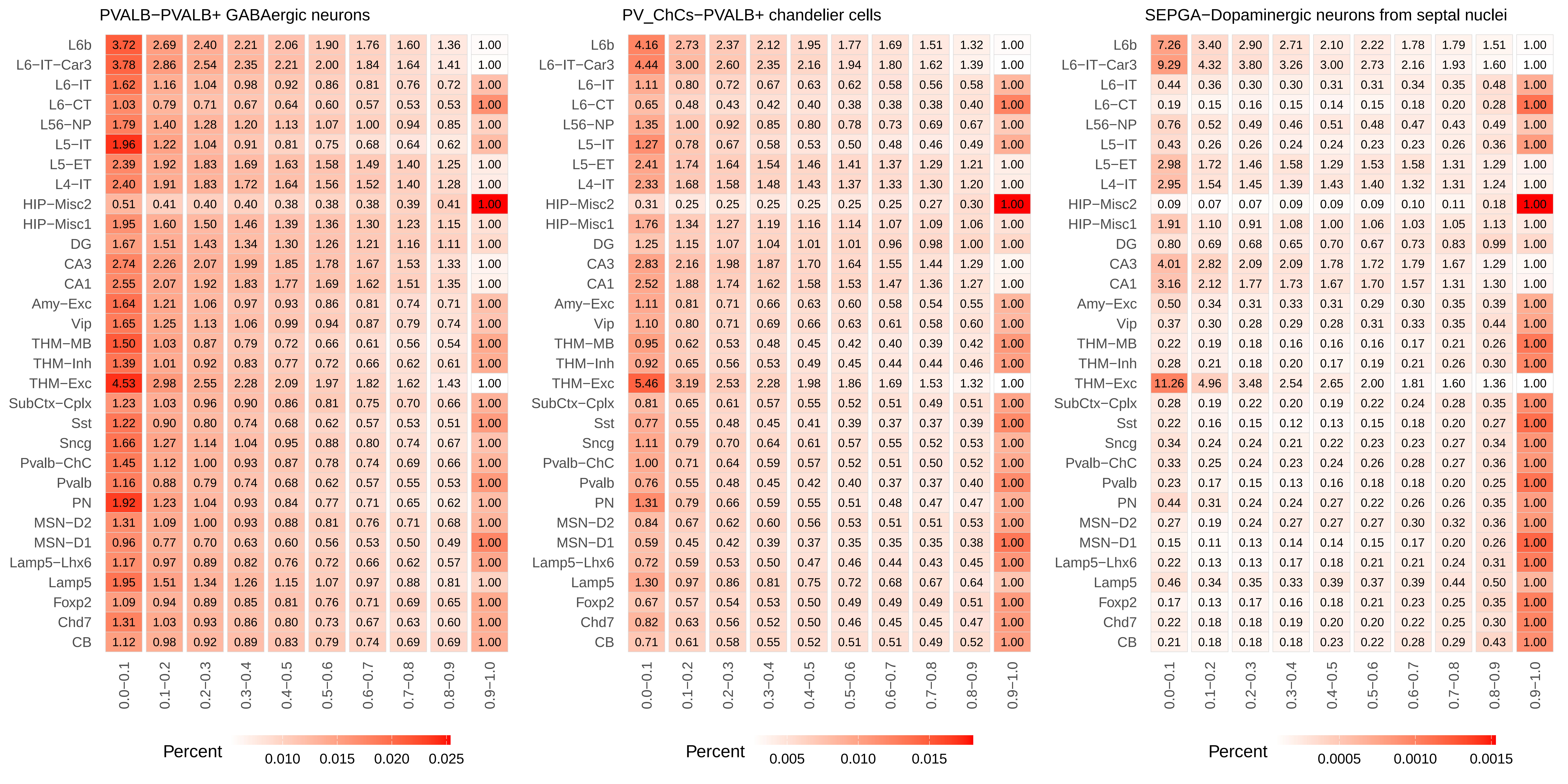
PVALB−PVALB+ GABAergic neurons
L6b 3.72 2.69 2.40 2.21 2.06 1.90 1.76 1.60 1.36 1.00
L6−IT−Car3 3.78 2.86 2.54 2.35 2.21 2.00 1.84 1.64 1.41 1.00
L6−IT 1.62 1.16 1.04 0.98 0.92 0.86 0.81 0.76 0.72 1.00
L6−CT 1.03 0.79 0.71 0.67 0.64 0.60 0.57 0.53 0.53 1.00
L56−NP 1.79 1.40 1.28 1.20 1.13 1.07 1.00 0.94 0.85 1.00
L5−IT 1.96 1.22 1.04 0.91 0.81 0.75 0.68 0.64 0.62 1.00
L5−ET 2.39 1.92 1.83 1.69 1.63 1.58 1.49 1.40 1.25 1.00
L4−IT 2.40 1.91 1.83 1.72 1.64 1.56 1.52 1.40 1.28 1.00
HIP−Misc2 0.51 0.41 0.40 0.40 0.38 0.38 0.38 0.39 0.41 1.00
HIP−Misc1 1.95 1.60 1.50 1.46 1.39 1.36 1.30 1.23 1.15 1.00
DG 1.67 1.51 1.43 1.34 1.30 1.26 1.21 1.16 1.11 1.00
CA3 2.74 2.26 2.07 1.99 1.85 1.78 1.67 1.53 1.33 1.00
CA1 2.55 2.07 1.92 1.83 1.77 1.69 1.62 1.51 1.35 1.00
Amy−Exc 1.64 1.21 1.06 0.97 0.93 0.86 0.81 0.74 0.71 1.00
Vip 1.65 1.25 1.13 1.06 0.99 0.94 0.87 0.79 0.74 1.00
THM−MB 1.50 1.03 0.87 0.79 0.72 0.66 0.61 0.56 0.54 1.00
THM−Inh 1.39 1.01 0.92 0.83 0.77 0.72 0.66 0.62 0.61 1.00
THM−Exc 4.53 2.98 2.55 2.28 2.09 1.97 1.82 1.62 1.43 1.00
SubCtx−Cplx 1.23 1.03 0.96 0.90 0.86 0.81 0.75 0.70 0.66 1.00
Sst 1.22 0.90 0.80 0.74 0.68 0.62 0.57 0.53 0.51 1.00
Sncg 1.66 1.27 1.14 1.04 0.95 0.88 0.80 0.74 0.67 1.00
Pvalb−ChC 1.45 1.12 1.00 0.93 0.87 0.78 0.74 0.69 0.66 1.00
Pvalb 1.16 0.88 0.79 0.74 0.68 0.62 0.57 0.55 0.53 1.00
PN 1.92 1.23 1.04 0.93 0.84 0.77 0.71 0.65 0.62 1.00
MSN−D2 1.31 1.09 1.00 0.93 0.88 0.81 0.76 0.71 0.68 1.00
MSN−D1 0.96 0.77 0.70 0.63 0.60 0.56 0.53 0.50 0.49 1.00
Lamp5−Lhx6 1.17 0.97 0.89 0.82 0.76 0.72 0.66 0.62 0.57 1.00
Lamp5 1.95 1.51 1.34 1.26 1.15 1.07 0.97 0.88 0.81 1.00
Foxp2 1.09 0.94 0.89 0.85 0.81 0.76 0.71 0.69 0.65 1.00
Chd7 1.31 1.03 0.93 0.86 0.80 0.73 0.67 0.63 0.60 1.00
CB 1.12 0.98 0.92 0.89 0.83 0.79 0.74 0.69 0.69 1.00
0.0−0.1 0.1−0.2 0.2−0.3 0.3−0.4 0.4−0.5 0.5−0.6 0.6−0.7 0.7−0.8 0.8−0.9 0.9−1.0
Percent 0.010 0.015 0.020 0.025
PV_ChCs−PVALB+ chandelier cells
L6b 4.16 2.73 2.37 2.12 1.95 1.77 1.69 1.51 1.32 1.00
L6−IT−Car3 4.44 3.00 2.60 2.35 2.16 1.94 1.80 1.62 1.39 1.00
L6−IT 1.11 0.80 0.72 0.67 0.63 0.62 0.58 0.56 0.58 1.00
L6−CT 0.65 0.48 0.43 0.42 0.40 0.38 0.38 0.38 0.40 1.00
L56−NP 1.35 1.00 0.92 0.85 0.80 0.78 0.73 0.69 0.67 1.00
L5−IT 1.27 0.78 0.67 0.58 0.53 0.50 0.48 0.46 0.49 1.00
L5−ET 2.41 1.74 1.64 1.54 1.46 1.41 1.37 1.29 1.21 1.00
L4−IT 2.33 1.68 1.58 1.48 1.43 1.37 1.33 1.30 1.20 1.00
HIP−Misc2 0.31 0.25 0.25 0.25 0.25 0.25 0.25 0.27 0.30 1.00
HIP−Misc1 1.76 1.34 1.27 1.19 1.16 1.14 1.07 1.09 1.06 1.00
DG 1.25 1.15 1.07 1.04 1.01 1.01 0.96 0.98 1.00 1.00
CA3 2.83 2.16 1.98 1.87 1.70 1.64 1.55 1.44 1.29 1.00
CA1 2.52 1.88 1.74 1.62 1.58 1.53 1.47 1.36 1.27 1.00
Amy−Exc 1.11 0.81 0.71 0.66 0.63 0.60 0.58 0.54 0.55 1.00
Vip 1.10 0.80 0.71 0.69 0.66 0.63 0.61 0.58 0.60 1.00
THM−MB 0.95 0.62 0.53 0.48 0.45 0.42 0.40 0.39 0.42 1.00
THM−Inh 0.92 0.65 0.56 0.53 0.49 0.45 0.44 0.44 0.46 1.00
THM−Exc 5.46 3.19 2.53 2.28 1.98 1.86 1.69 1.53 1.32 1.00
SubCtx−Cplx 0.81 0.65 0.61 0.57 0.55 0.52 0.51 0.49 0.51 1.00
Sst 0.77 0.55 0.48 0.45 0.41 0.39 0.37 0.37 0.39 1.00
Sncg 1.11 0.79 0.70 0.64 0.61 0.57 0.55 0.52 0.53 1.00
Pvalb−ChC 1.00 0.71 0.64 0.59 0.57 0.52 0.51 0.50 0.52 1.00
Pvalb 0.76 0.55 0.48 0.45 0.42 0.40 0.37 0.37 0.40 1.00
PN 1.31 0.79 0.66 0.59 0.55 0.51 0.48 0.47 0.47 1.00
MSN−D2 0.84 0.67 0.62 0.60 0.56 0.53 0.51 0.51 0.53 1.00
MSN−D1 0.59 0.45 0.42 0.39 0.37 0.35 0.35 0.35 0.38 1.00
Lamp5−Lhx6 0.72 0.59 0.53 0.50 0.47 0.46 0.44 0.43 0.45 1.00
Lamp5 1.30 0.97 0.86 0.81 0.75 0.72 0.68 0.67 0.64 1.00
Foxp2 0.67 0.57 0.54 0.53 0.50 0.49 0.49 0.49 0.51 1.00
Chd7 0.82 0.63 0.56 0.52 0.50 0.46 0.45 0.45 0.47 1.00
CB 0.71 0.61 0.58 0.55 0.52 0.51 0.51 0.49 0.52 1.00
0.0−0.1 0.1−0.2 0.2−0.3 0.3−0.4 0.4−0.5 0.5−0.6 0.6−0.7 0.7−0.8 0.8−0.9 0.9−1.0
Percent 0.005 0.010 0.015
SEPGA−Dopaminergic neurons from septal nuclei
L6b 7.26 3.40 2.90 2.71 2.10 2.22 1.78 1.79 1.51 1.00
L6−IT−Car3 9.29 4.32 3.80 3.26 3.00 2.73 2.16 1.93 1.60 1.00
L6−IT 0.44 0.36 0.30 0.30 0.31 0.31 0.34 0.35 0.48 1.00
L6−CT 0.19 0.15 0.16 0.15 0.14 0.15 0.18 0.20 0.28 1.00
L56−NP 0.76 0.52 0.49 0.46 0.51 0.48 0.47 0.43 0.49 1.00
L5−IT 0.43 0.26 0.26 0.24 0.24 0.23 0.23 0.26 0.36 1.00
L5−ET 2.98 1.72 1.46 1.58 1.29 1.53 1.58 1.31 1.29 1.00
L4−IT 2.95 1.54 1.45 1.39 1.43 1.40 1.32 1.31 1.24 1.00
HIP−Misc2 0.09 0.07 0.07 0.09 0.09 0.09 0.10 0.11 0.18 1.00
HIP−Misc1 1.91 1.10 0.91 1.08 1.00 1.06 1.03 1.05 1.13 1.00
DG 0.80 0.69 0.68 0.65 0.70 0.67 0.73 0.83 0.99 1.00
CA3 4.01 2.82 2.09 2.09 1.78 1.72 1.79 1.67 1.29 1.00
CA1 3.16 2.12 1.77 1.73 1.67 1.70 1.57 1.31 1.30 1.00
Amy−Exc 0.50 0.34 0.31 0.33 0.31 0.29 0.30 0.35 0.39 1.00
Vip 0.37 0.30 0.28 0.29 0.28 0.31 0.33 0.35 0.44 1.00
THM−MB 0.22 0.19 0.18 0.16 0.16 0.16 0.17 0.21 0.26 1.00
THM−Inh 0.28 0.21 0.18 0.20 0.17 0.19 0.21 0.26 0.30 1.00
THM−Exc 11.26 4.96 3.48 2.54 2.65 2.00 1.81 1.60 1.36 1.00
SubCtx−Cplx 0.28 0.19 0.22 0.20 0.19 0.22 0.24 0.28 0.35 1.00
Sst 0.22 0.16 0.15 0.12 0.13 0.15 0.18 0.20 0.27 1.00
Sncg 0.34 0.24 0.24 0.21 0.22 0.23 0.23 0.27 0.34 1.00
Pvalb−ChC 0.33 0.25 0.24 0.23 0.24 0.26 0.28 0.27 0.36 1.00
Pvalb 0.23 0.17 0.15 0.13 0.16 0.18 0.18 0.20 0.25 1.00
PN 0.44 0.31 0.24 0.24 0.27 0.22 0.26 0.26 0.35 1.00
MSN−D2 0.27 0.19 0.24 0.27 0.27 0.27 0.30 0.32 0.36 1.00
MSN−D1 0.15 0.11 0.13 0.14 0.14 0.15 0.17 0.20 0.26 1.00
Lamp5−Lhx6 0.22 0.13 0.13 0.17 0.18 0.21 0.21 0.24 0.31 1.00
Lamp5 0.46 0.34 0.35 0.33 0.39 0.37 0.39 0.44 0.50 1.00
Foxp2 0.17 0.13 0.17 0.16 0.18 0.21 0.23 0.25 0.35 1.00
Chd7 0.22 0.18 0.18 0.19 0.20 0.20 0.22 0.25 0.30 1.00
CB 0.21 0.18 0.18 0.18 0.23 0.22 0.28 0.29 0.43 1.00
0.0−0.1 0.1−0.2 0.2−0.3 0.3−0.4 0.4−0.5 0.5−0.6 0.6−0.7 0.7−0.8 0.8−0.9 0.9−1.0
Percent 0.0005 0.0010 0.0015

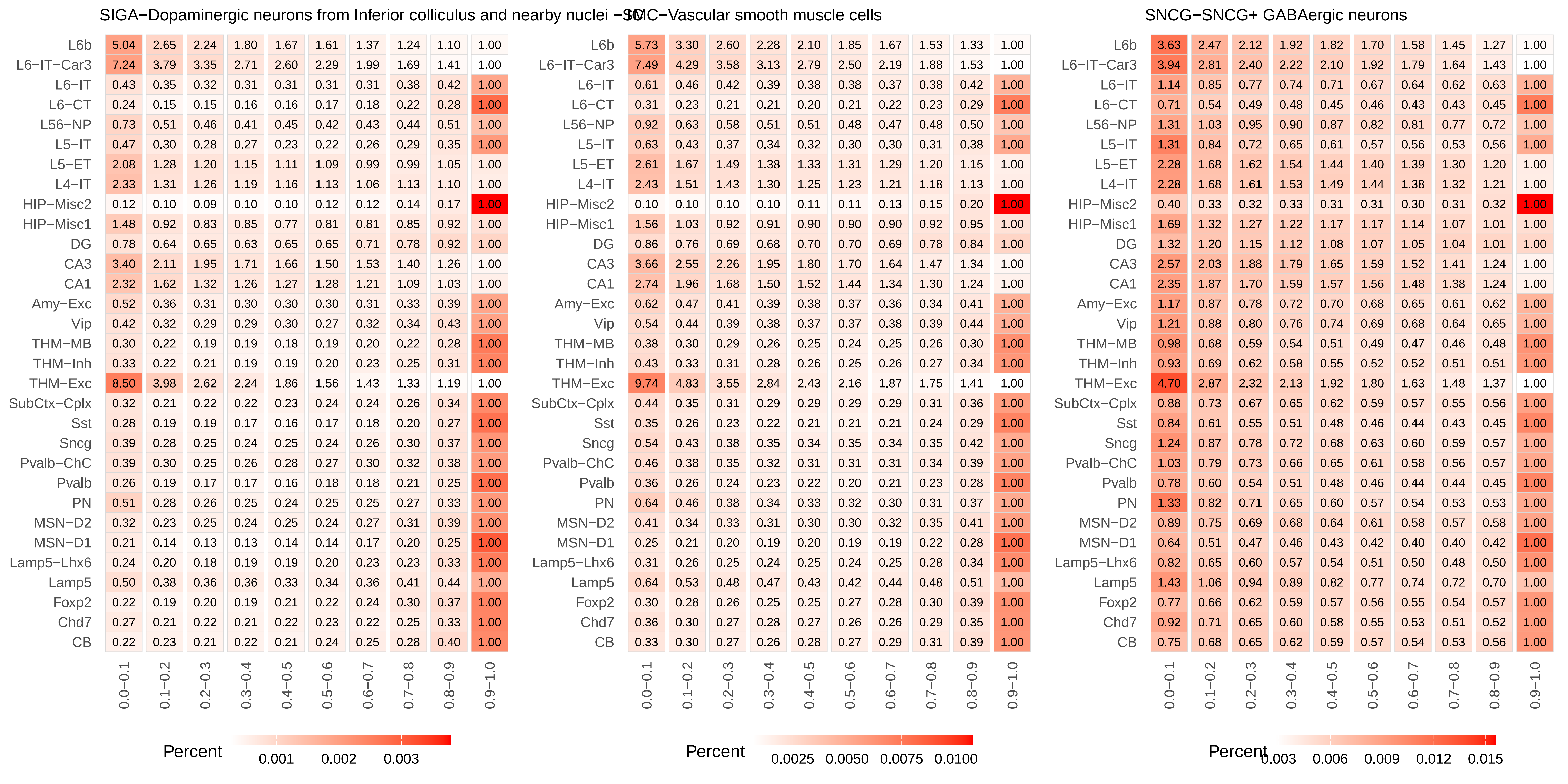
SIGA−Dopaminergic neurons from Inferior colliculus and nearby nuclei
L6b 5.04 2.65 2.24 1.80 1.67 1.61 1.37 1.24 1.10 1.00
L6−IT−Car3 7.24 3.79 3.35 2.71 2.60 2.29 1.99 1.69 1.41 1.00
L6−IT 0.43 0.35 0.32 0.31 0.31 0.31 0.31 0.38 0.42 1.00
L6−CT 0.24 0.15 0.15 0.16 0.16 0.17 0.18 0.22 0.28 1.00
L56−NP 0.73 0.51 0.46 0.41 0.45 0.42 0.43 0.44 0.51 1.00
L5−IT 0.47 0.30 0.28 0.27 0.23 0.22 0.26 0.29 0.35 1.00
L5−ET 2.08 1.28 1.20 1.15 1.11 1.09 0.99 0.99 1.05 1.00
L4−IT 2.33 1.31 1.26 1.19 1.16 1.13 1.06 1.13 1.10 1.00
HIP−Misc2 0.12 0.10 0.09 0.10 0.10 0.12 0.12 0.14 0.17 1.00
HIP−Misc1 1.48 0.92 0.83 0.85 0.77 0.81 0.81 0.85 0.92 1.00
DG 0.78 0.64 0.65 0.63 0.65 0.65 0.71 0.78 0.92 1.00
CA3 3.40 2.11 1.95 1.71 1.66 1.50 1.53 1.40 1.26 1.00
CA1 2.32 1.62 1.32 1.26 1.27 1.28 1.21 1.09 1.03 1.00
Amy−Exc 0.52 0.36 0.31 0.30 0.30 0.30 0.31 0.33 0.39 1.00
Vip 0.42 0.32 0.29 0.29 0.30 0.27 0.32 0.34 0.43 1.00
THM−MB 0.30 0.22 0.19 0.19 0.18 0.19 0.20 0.22 0.28 1.00
THM−Inh 0.33 0.22 0.21 0.19 0.19 0.20 0.23 0.25 0.31 1.00
THM−Exc 8.50 3.98 2.62 2.24 1.86 1.56 1.43 1.33 1.19 1.00
SubCtx−Cplx 0.32 0.21 0.22 0.22 0.23 0.24 0.24 0.26 0.34 1.00
Sst 0.28 0.19 0.19 0.17 0.16 0.17 0.18 0.20 0.27 1.00
Sncg 0.39 0.28 0.25 0.24 0.25 0.24 0.26 0.30 0.37 1.00
Pvalb−ChC 0.39 0.30 0.25 0.26 0.28 0.27 0.30 0.32 0.38 1.00
Pvalb 0.26 0.19 0.17 0.17 0.16 0.18 0.18 0.21 0.25 1.00
PN 0.51 0.28 0.26 0.25 0.24 0.25 0.25 0.27 0.33 1.00
MSN−D2 0.32 0.23 0.25 0.24 0.25 0.24 0.27 0.31 0.39 1.00
MSN−D1 0.21 0.14 0.13 0.13 0.14 0.14 0.17 0.20 0.25 1.00
Lamp5−Lhx6 0.24 0.20 0.18 0.19 0.19 0.20 0.23 0.23 0.33 1.00
Lamp5 0.50 0.38 0.36 0.36 0.33 0.34 0.36 0.41 0.44 1.00
Foxp2 0.22 0.19 0.20 0.19 0.21 0.22 0.24 0.30 0.37 1.00
Chd7 0.27 0.21 0.22 0.21 0.22 0.23 0.22 0.25 0.33 1.00
CB 0.22 0.23 0.21 0.22 0.21 0.24 0.25 0.28 0.40 1.00
0.0−0.1 0.1−0.2 0.2−0.3 0.3−0.4 0.4−0.5 0.5−0.6 0.6−0.7 0.7−0.8 0.8−0.9 0.9−1.0
Percent 0.001 0.002 0.003
SMC−Vascular smooth muscle cells
L6b 5.73 3.30 2.60 2.28 2.10 1.85 1.67 1.53 1.33 1.00
L6−IT−Car3 7.49 4.29 3.58 3.13 2.79 2.50 2.19 1.88 1.53 1.00
L6−IT 0.61 0.46 0.42 0.39 0.38 0.38 0.37 0.38 0.42 1.00
L6−CT 0.31 0.23 0.21 0.21 0.20 0.21 0.22 0.23 0.29 1.00
L56−NP 0.92 0.63 0.58 0.51 0.51 0.48 0.47 0.48 0.50 1.00
L5−IT 0.63 0.43 0.37 0.34 0.32 0.30 0.30 0.31 0.38 1.00
L5−ET 2.61 1.67 1.49 1.38 1.33 1.31 1.29 1.20 1.15 1.00
L4−IT 2.43 1.51 1.43 1.30 1.25 1.23 1.21 1.18 1.13 1.00
HIP−Misc2 0.10 0.10 0.10 0.10 0.11 0.11 0.13 0.15 0.20 1.00
HIP−Misc1 1.56 1.03 0.92 0.91 0.90 0.90 0.90 0.92 0.95 1.00
DG 0.86 0.76 0.69 0.68 0.70 0.70 0.69 0.78 0.84 1.00
CA3 3.66 2.55 2.26 1.95 1.80 1.70 1.64 1.47 1.34 1.00
CA1 2.74 1.96 1.68 1.50 1.52 1.44 1.34 1.30 1.24 1.00
Amy−Exc 0.62 0.47 0.41 0.39 0.38 0.37 0.36 0.34 0.41 1.00
Vip 0.54 0.44 0.39 0.38 0.37 0.37 0.38 0.39 0.44 1.00
THM−MB 0.38 0.30 0.29 0.26 0.25 0.24 0.25 0.26 0.30 1.00
THM−Inh 0.43 0.33 0.31 0.28 0.26 0.25 0.26 0.27 0.34 1.00
THM−Exc 9.74 4.83 3.55 2.84 2.43 2.16 1.87 1.75 1.41 1.00
SubCtx−Cplx 0.44 0.35 0.31 0.29 0.29 0.29 0.29 0.31 0.36 1.00
Sst 0.35 0.26 0.23 0.22 0.21 0.21 0.21 0.24 0.29 1.00
Sncg 0.54 0.43 0.38 0.35 0.34 0.35 0.34 0.35 0.42 1.00
Pvalb−ChC 0.46 0.38 0.35 0.32 0.31 0.31 0.31 0.34 0.39 1.00
Pvalb 0.36 0.26 0.24 0.23 0.22 0.20 0.21 0.23 0.28 1.00
PN 0.64 0.46 0.38 0.34 0.33 0.32 0.30 0.31 0.37 1.00
MSN−D2 0.41 0.34 0.33 0.31 0.30 0.30 0.32 0.35 0.41 1.00
MSN−D1 0.25 0.21 0.20 0.19 0.20 0.19 0.19 0.22 0.28 1.00
Lamp5−Lhx6 0.31 0.26 0.25 0.24 0.25 0.24 0.25 0.28 0.34 1.00
Lamp5 0.64 0.53 0.48 0.47 0.43 0.42 0.44 0.48 0.51 1.00
Foxp2 0.30 0.28 0.26 0.25 0.25 0.27 0.28 0.30 0.39 1.00
Chd7 0.36 0.30 0.27 0.28 0.27 0.26 0.26 0.29 0.35 1.00
CB 0.33 0.30 0.27 0.26 0.28 0.27 0.29 0.31 0.39 1.00
0.0−0.1 0.1−0.2 0.2−0.3 0.3−0.4 0.4−0.5 0.5−0.6 0.6−0.7 0.7−0.8 0.8−0.9 0.9−1.0
Percent 0.0025 0.0050 0.0075 0.0100
SNCG−SNCG+ GABAergic neurons
L6b 3.63 2.47 2.12 1.92 1.82 1.70 1.58 1.45 1.27 1.00
L6−IT−Car3 3.94 2.81 2.40 2.22 2.10 1.92 1.79 1.64 1.43 1.00
L6−IT 1.14 0.85 0.77 0.74 0.71 0.67 0.64 0.62 0.63 1.00
L6−CT 0.71 0.54 0.49 0.48 0.45 0.46 0.43 0.43 0.45 1.00
L56−NP 1.31 1.03 0.95 0.90 0.87 0.82 0.81 0.77 0.72 1.00
L5−IT 1.31 0.84 0.72 0.65 0.61 0.57 0.56 0.53 0.56 1.00
L5−ET 2.28 1.68 1.62 1.54 1.44 1.40 1.39 1.30 1.20 1.00
L4−IT 2.28 1.68 1.61 1.53 1.49 1.44 1.38 1.32 1.21 1.00
HIP−Misc2 0.40 0.33 0.32 0.33 0.31 0.31 0.30 0.31 0.32 1.00
HIP−Misc1 1.69 1.32 1.27 1.22 1.17 1.17 1.14 1.07 1.01 1.00
DG 1.32 1.20 1.15 1.12 1.08 1.07 1.05 1.04 1.01 1.00
CA3 2.57 2.03 1.88 1.79 1.65 1.59 1.52 1.41 1.24 1.00
CA1 2.35 1.87 1.70 1.59 1.57 1.56 1.48 1.38 1.24 1.00
Amy−Exc 1.17 0.87 0.78 0.72 0.70 0.68 0.65 0.61 0.62 1.00
Vip 1.21 0.88 0.80 0.76 0.74 0.69 0.68 0.64 0.65 1.00
THM−MB 0.98 0.68 0.59 0.54 0.51 0.49 0.47 0.46 0.48 1.00
THM−Inh 0.93 0.69 0.62 0.58 0.55 0.52 0.52 0.51 0.51 1.00
THM−Exc 4.70 2.87 2.32 2.13 1.92 1.80 1.63 1.48 1.37 1.00
SubCtx−Cplx 0.88 0.73 0.67 0.65 0.62 0.59 0.57 0.55 0.56 1.00
Sst 0.84 0.61 0.55 0.51 0.48 0.46 0.44 0.43 0.45 1.00
Sncg 1.24 0.87 0.78 0.72 0.68 0.63 0.60 0.59 0.57 1.00
Pvalb−ChC 1.03 0.79 0.73 0.66 0.65 0.61 0.58 0.56 0.57 1.00
Pvalb 0.78 0.60 0.54 0.51 0.48 0.46 0.44 0.44 0.45 1.00
PN 1.33 0.82 0.71 0.65 0.60 0.57 0.54 0.53 0.53 1.00
MSN−D2 0.89 0.75 0.69 0.68 0.64 0.61 0.58 0.57 0.58 1.00
MSN−D1 0.64 0.51 0.47 0.46 0.43 0.42 0.40 0.40 0.42 1.00
Lamp5−Lhx6 0.82 0.65 0.60 0.57 0.54 0.51 0.50 0.48 0.50 1.00
Lamp5 1.43 1.06 0.94 0.89 0.82 0.77 0.74 0.72 0.70 1.00
Foxp2 0.77 0.66 0.62 0.59 0.57 0.56 0.55 0.54 0.57 1.00
Chd7 0.92 0.71 0.65 0.60 0.58 0.55 0.53 0.51 0.52 1.00
CB 0.75 0.68 0.65 0.62 0.59 0.57 0.54 0.53 0.56 1.00
0.0−0.1 0.1−0.2 0.2−0.3 0.3−0.4 0.4−0.5 0.5−0.6 0.6−0.7 0.7−0.8 0.8−0.9 0.9−1.0
Percent 0.003 0.006 0.009 0.012 0.015

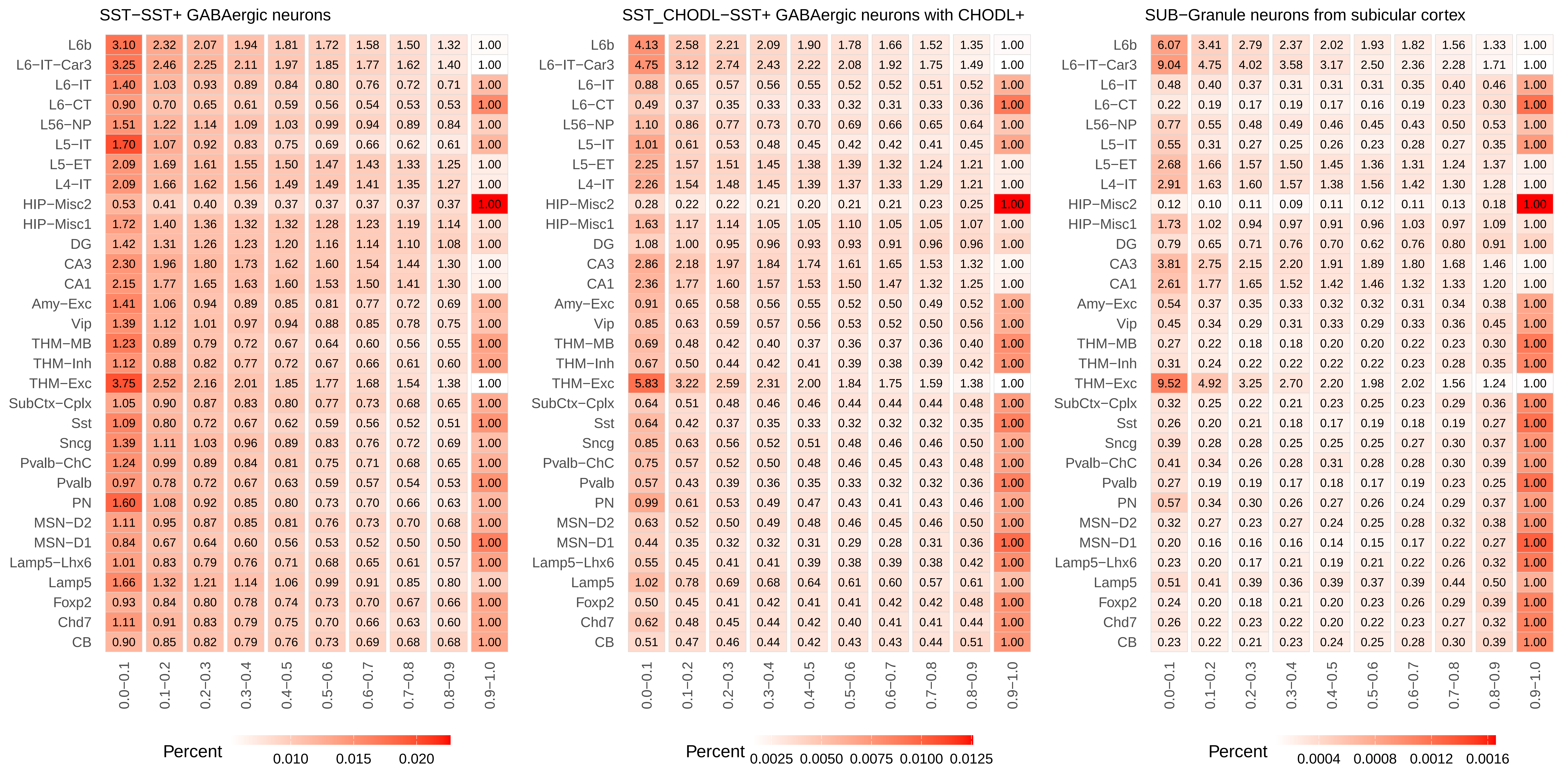
SST−SST+ GABAergic neurons
SST_CHODL−SST+ GABAergic neurons with CHODL+
SUB−Granule neurons from subicular cortex
L6b
L6−IT−Car3
L6−IT
L6−CT
L56−NP
L5−IT
L5−ET
L4−IT
HIP−Misc2
HIP−Misc1
DG
CA3
CA1
Amy−Exc
Vip
THM−MB
THM−Inh
THM−Exc
SubCtx−Cplx
Sst
Sncg
Pvalb−ChC
Pvalb
PN
MSN−D2
MSN−D1
Lamp5−Lhx6
Lamp5
Foxp2
Chd7
CB
0.0−0.1
0.1−0.2
0.2−0.3
0.3−0.4
0.4−0.5
0.5−0.6
0.6−0.7
0.7−0.8
0.8−0.9
0.9−1.0
Percent
0.010 0.015 0.020
Percent
0.0025 0.0050 0.0075 0.0100 0.0125
Percent
0.0004 0.0008 0.0012 0.0016

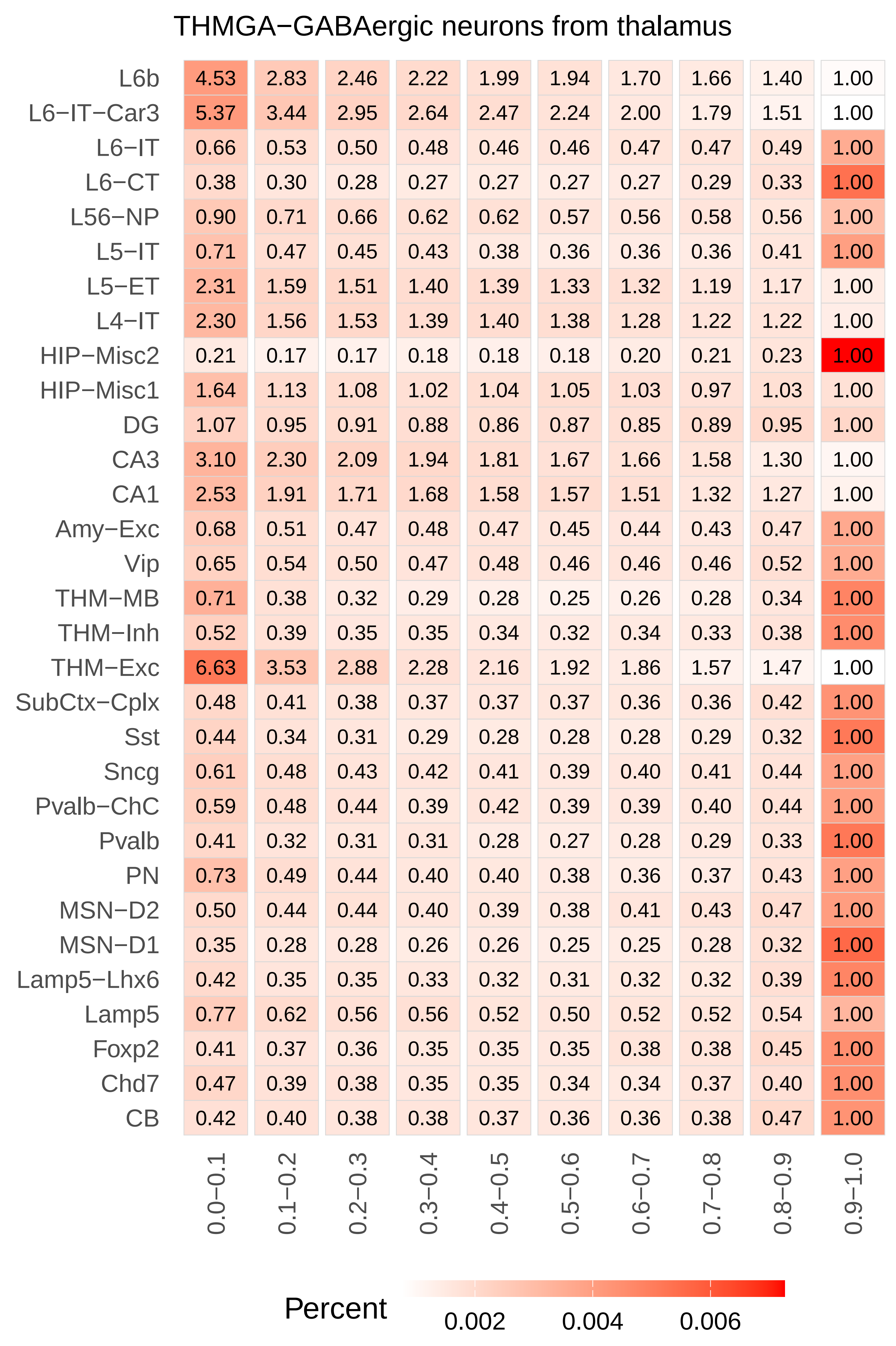
THMGA−GABAergic neurons from thalamus
L6b 4.53 2.83 2.46 2.22 1.99 1.94 1.70 1.66 1.40 1.00
L6−IT−Car3 5.37 3.44 2.95 2.64 2.47 2.24 2.00 1.79 1.51 1.00
L6−IT 0.66 0.53 0.50 0.48 0.46 0.46 0.47 0.47 0.49 1.00
L6−CT 0.38 0.30 0.28 0.27 0.27 0.27 0.27 0.29 0.33 1.00
L56−NP 0.90 0.71 0.66 0.62 0.62 0.57 0.56 0.58 0.56 1.00
L5−IT 0.71 0.47 0.45 0.43 0.38 0.36 0.36 0.36 0.41 1.00
L5−ET 2.31 1.59 1.51 1.40 1.39 1.33 1.32 1.19 1.17 1.00
L4−IT 2.30 1.56 1.53 1.39 1.40 1.38 1.28 1.22 1.22 1.00
HIP−Misc2 0.21 0.17 0.17 0.18 0.18 0.18 0.20 0.21 0.23 1.00
HIP−Misc1 1.64 1.13 1.08 1.02 1.04 1.05 1.03 0.97 1.03 1.00
DG 1.07 0.95 0.91 0.88 0.86 0.87 0.85 0.89 0.95 1.00
CA3 3.10 2.30 2.09 1.94 1.81 1.67 1.66 1.58 1.30 1.00
CA1 2.53 1.91 1.71 1.68 1.58 1.57 1.51 1.32 1.27 1.00
Amy−Exc 0.68 0.51 0.47 0.48 0.47 0.45 0.44 0.43 0.47 1.00
Vip 0.65 0.54 0.50 0.47 0.48 0.46 0.46 0.46 0.52 1.00
THM−MB 0.71 0.38 0.32 0.29 0.28 0.25 0.26 0.28 0.34 1.00
THM−Inh 0.52 0.39 0.35 0.35 0.34 0.32 0.34 0.33 0.38 1.00
THM−Exc 6.63 3.53 2.88 2.28 2.16 1.92 1.86 1.57 1.47 1.00
SubCtx−Cplx 0.48 0.41 0.38 0.37 0.37 0.37 0.36 0.36 0.42 1.00
Sst 0.44 0.34 0.31 0.29 0.28 0.28 0.28 0.29 0.32 1.00
Sncg 0.61 0.48 0.43 0.42 0.41 0.39 0.40 0.41 0.44 1.00
Pvalb−ChC 0.59 0.48 0.44 0.39 0.42 0.39 0.39 0.40 0.44 1.00
Pvalb 0.41 0.32 0.31 0.31 0.28 0.27 0.28 0.29 0.33 1.00
PN 0.73 0.49 0.44 0.40 0.40 0.38 0.36 0.37 0.43 1.00
MSN−D2 0.50 0.44 0.44 0.40 0.39 0.38 0.41 0.43 0.47 1.00
MSN−D1 0.35 0.28 0.28 0.26 0.26 0.25 0.25 0.28 0.32 1.00
Lamp5−Lhx6 0.42 0.35 0.35 0.33 0.32 0.31 0.32 0.32 0.39 1.00
Lamp5 0.77 0.62 0.56 0.56 0.52 0.50 0.52 0.52 0.54 1.00
Foxp2 0.41 0.37 0.36 0.35 0.35 0.35 0.38 0.38 0.45 1.00
Chd7 0.47 0.39 0.38 0.35 0.35 0.34 0.34 0.37 0.40 1.00
CB 0.42 0.40 0.38 0.38 0.37 0.36 0.36 0.38 0.47 1.00
0.0−0.1 0.1−0.2 0.2−0.3 0.3−0.4 0.4−0.5 0.5−0.6 0.6−0.7 0.7−0.8 0.8−0.9 0.9−1.0
Percent
0.002 0.004 0.006

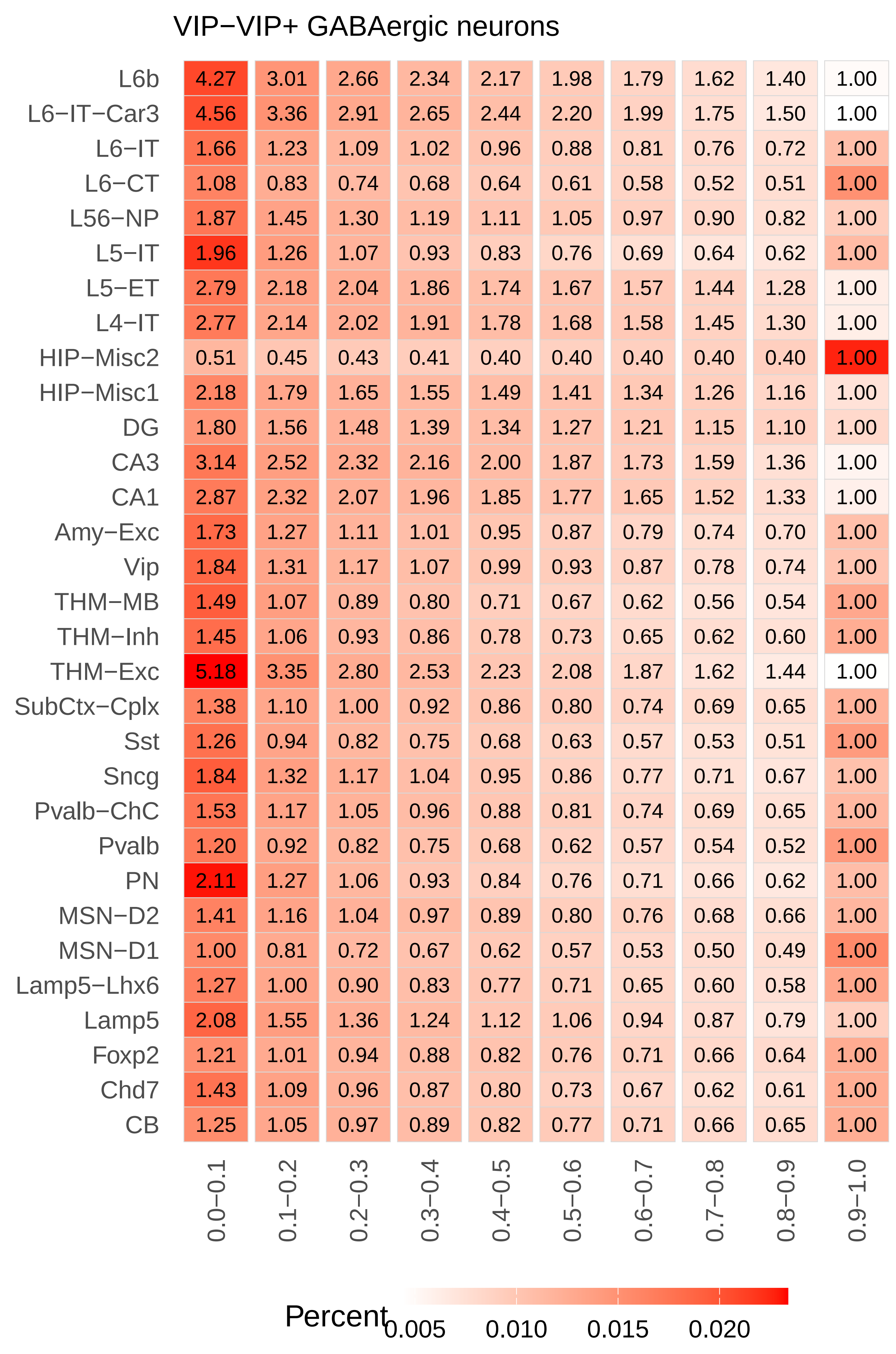
VIP−VIP+ GABAergic neurons
L6b 4.27 3.01 2.66 2.34 2.17 1.98 1.79 1.62 1.40 1.00
L6−IT−Car3 4.56 3.36 2.91 2.65 2.44 2.20 1.99 1.75 1.50 1.00
L6−IT 1.66 1.23 1.09 1.02 0.96 0.88 0.81 0.76 0.72 1.00
L6−CT 1.08 0.83 0.74 0.68 0.64 0.61 0.58 0.52 0.51 1.00
L56−NP 1.87 1.45 1.30 1.19 1.11 1.05 0.97 0.90 0.82 1.00
L5−IT 1.96 1.26 1.07 0.93 0.83 0.76 0.69 0.64 0.62 1.00
L5−ET 2.79 2.18 2.04 1.86 1.74 1.67 1.57 1.44 1.28 1.00
L4−IT 2.77 2.14 2.02 1.91 1.78 1.68 1.58 1.45 1.30 1.00
HIP−Misc2 0.51 0.45 0.43 0.41 0.40 0.40 0.40 0.40 0.40 1.00
HIP−Misc1 2.18 1.79 1.65 1.55 1.49 1.41 1.34 1.26 1.16 1.00
DG 1.80 1.56 1.48 1.39 1.34 1.27 1.21 1.15 1.10 1.00
CA3 3.14 2.52 2.32 2.16 2.00 1.87 1.73 1.59 1.36 1.00
CA1 2.87 2.32 2.07 1.96 1.85 1.77 1.65 1.52 1.33 1.00
Amy−Exc 1.73 1.27 1.11 1.01 0.95 0.87 0.79 0.74 0.70 1.00
Vip 1.84 1.31 1.17 1.07 0.99 0.93 0.87 0.78 0.74 1.00
THM−MB 1.49 1.07 0.89 0.80 0.71 0.67 0.62 0.56 0.54 1.00
THM−Inh 1.45 1.06 0.93 0.86 0.78 0.73 0.65 0.62 0.60 1.00
THM−Exc 5.18 3.35 2.80 2.53 2.23 2.08 1.87 1.62 1.44 1.00
SubCtx−Cplx 1.38 1.10 1.00 0.92 0.86 0.80 0.74 0.69 0.65 1.00
Sst 1.26 0.94 0.82 0.75 0.68 0.63 0.57 0.53 0.51 1.00
Sncg 1.84 1.32 1.17 1.04 0.95 0.86 0.77 0.71 0.67 1.00
Pvalb−ChC 1.53 1.17 1.05 0.96 0.88 0.81 0.74 0.69 0.65 1.00
Pvalb 1.20 0.92 0.82 0.75 0.68 0.62 0.57 0.54 0.52 1.00
PN 2.11 1.27 1.06 0.93 0.84 0.76 0.71 0.66 0.62 1.00
MSN−D2 1.41 1.16 1.04 0.97 0.89 0.80 0.76 0.68 0.66 1.00
MSN−D1 1.00 0.81 0.72 0.67 0.62 0.57 0.53 0.50 0.49 1.00
Lamp5−Lhx6 1.27 1.00 0.90 0.83 0.77 0.71 0.65 0.60 0.58 1.00
Lamp5 2.08 1.55 1.36 1.24 1.12 1.06 0.94 0.87 0.79 1.00
Foxp2 1.21 1.01 0.94 0.88 0.82 0.76 0.71 0.66 0.64 1.00
Chd7 1.43 1.09 0.96 0.87 0.80 0.73 0.67 0.62 0.61 1.00
CB 1.25 1.05 0.97 0.89 0.82 0.77 0.71 0.66 0.65 1.00
0.0−0.1 0.1−0.2 0.2−0.3 0.3−0.4 0.4−0.5 0.5−0.6 0.6−0.7 0.7−0.8 0.8−0.9 0.9−1.0
Percent
0.005 0.010 0.015 0.020
