## Supplementary Figure 6 for "Distinct cellular DNA methylation mechanisms underlie common and rare genetic risk for brain disorders"

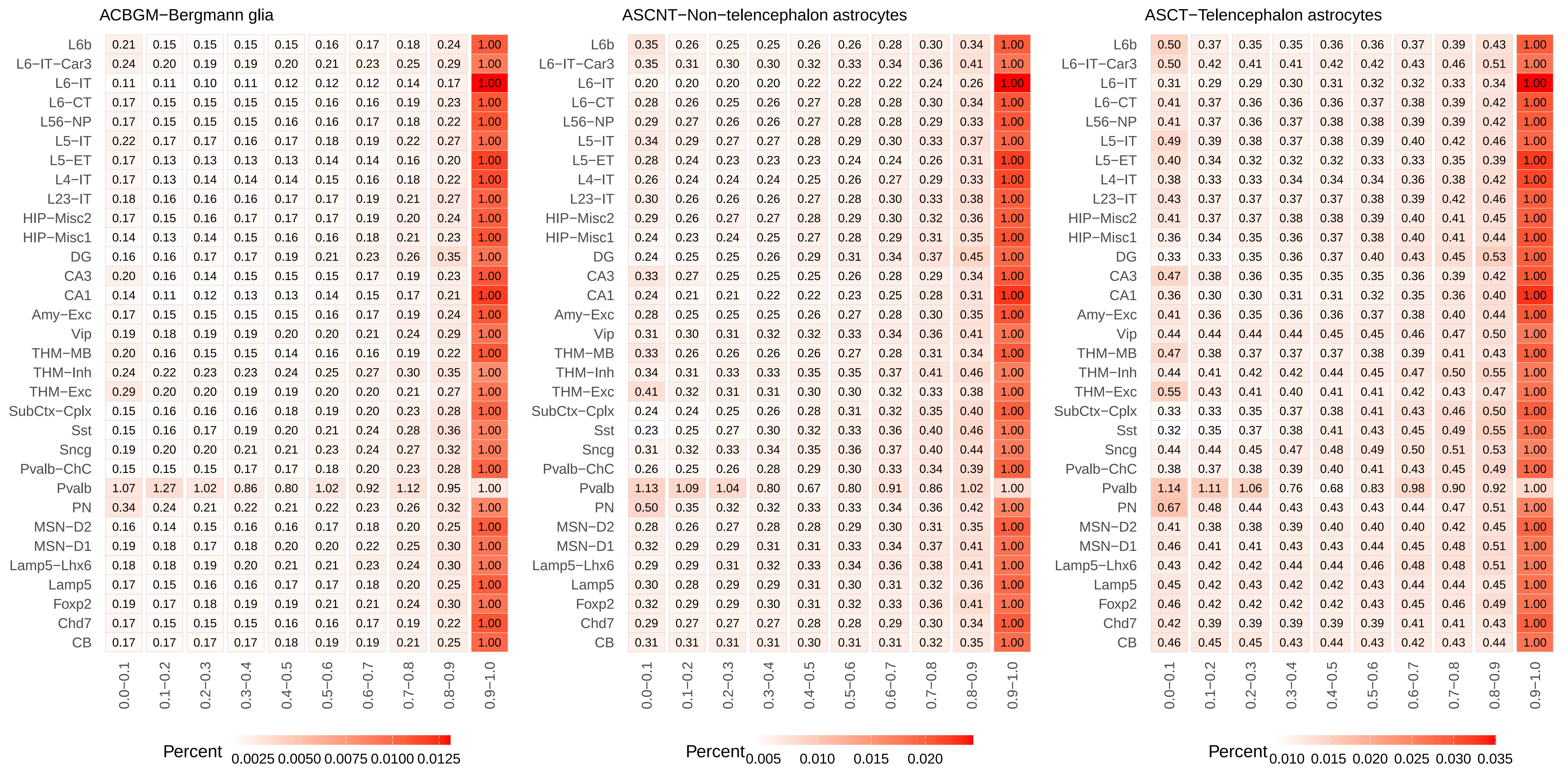
ACBGM−Bergmann glia
L6b 0.21 0.15 0.15 0.15 0.15 0.16 0.17 0.18 0.24 1.00
L6−IT−Car3 0.24 0.20 0.19 0.19 0.20 0.21 0.23 0.25 0.29 1.00
L6−IT 0.11 0.11 0.10 0.11 0.12 0.12 0.12 0.14 0.17 1.00
L6−CT 0.17 0.15 0.15 0.15 0.15 0.16 0.16 0.19 0.23 1.00
L56−NP 0.17 0.15 0.15 0.15 0.16 0.16 0.17 0.18 0.22 1.00
L5−IT 0.22 0.17 0.17 0.16 0.17 0.18 0.19 0.22 0.27 1.00
L5−ET 0.17 0.13 0.13 0.13 0.13 0.14 0.14 0.16 0.20 1.00
L4−IT 0.17 0.13 0.14 0.14 0.14 0.15 0.16 0.18 0.22 1.00
L23−IT 0.18 0.16 0.16 0.16 0.17 0.17 0.19 0.21 0.27 1.00
HIP−Misc2 0.17 0.15 0.16 0.17 0.17 0.17 0.19 0.20 0.24 1.00
HIP−Misc1 0.14 0.13 0.14 0.15 0.16 0.16 0.18 0.21 0.23 1.00
DG 0.16 0.16 0.17 0.17 0.19 0.21 0.23 0.26 0.35 1.00
CA3 0.20 0.16 0.14 0.15 0.15 0.15 0.17 0.19 0.23 1.00
CA1 0.14 0.11 0.12 0.13 0.13 0.14 0.15 0.17 0.21 1.00
Amy−Exc 0.17 0.15 0.15 0.15 0.15 0.16 0.17 0.19 0.24 1.00
Vip 0.19 0.18 0.19 0.19 0.20 0.20 0.21 0.24 0.29 1.00
THM−MB 0.20 0.16 0.15 0.15 0.14 0.16 0.16 0.19 0.22 1.00
THM−Inh 0.24 0.22 0.23 0.23 0.24 0.25 0.27 0.30 0.35 1.00
THM−Exc 0.29 0.20 0.20 0.19 0.19 0.20 0.20 0.21 0.27 1.00
SubCtx−Cplx 0.15 0.16 0.16 0.16 0.18 0.19 0.20 0.23 0.28 1.00
Sst 0.15 0.16 0.17 0.19 0.20 0.21 0.24 0.28 0.36 1.00
Sncg 0.19 0.20 0.20 0.21 0.21 0.23 0.24 0.27 0.32 1.00
Pvalb−ChC 0.15 0.15 0.15 0.17 0.17 0.18 0.20 0.23 0.28 1.00
Pvalb 1.07 1.27 1.02 0.86 0.80 1.02 0.92 1.12 0.95 1.00
PN 0.34 0.24 0.21 0.22 0.21 0.22 0.23 0.26 0.32 1.00
MSN−D2 0.16 0.14 0.15 0.16 0.16 0.17 0.18 0.20 0.25 1.00
MSN−D1 0.19 0.18 0.17 0.18 0.20 0.20 0.22 0.25 0.30 1.00
Lamp5−Lhx6 0.18 0.18 0.19 0.20 0.21 0.21 0.23 0.24 0.30 1.00
Lamp5 0.17 0.15 0.16 0.16 0.17 0.17 0.18 0.20 0.25 1.00
Foxp2 0.19 0.17 0.18 0.19 0.19 0.21 0.21 0.24 0.30 1.00
Chd7 0.17 0.15 0.15 0.15 0.16 0.16 0.17 0.19 0.22 1.00
CB 0.17 0.17 0.17 0.17 0.18 0.19 0.19 0.21 0.25 1.00
0.0−0.1 0.1−0.2 0.2−0.3 0.3−0.4 0.4−0.5 0.5−0.6 0.6−0.7 0.7−0.8 0.8−0.9 0.9−1.0
Percent
0.0025 0.0050 0.0075 0.0100 0.0125
ASCNT−Non−telencephalon astrocytes
L6b 0.35 0.26 0.25 0.25 0.26 0.26 0.28 0.30 0.34 1.00
L6−IT−Car3 0.35 0.31 0.30 0.30 0.32 0.33 0.34 0.36 0.41 1.00
L6−IT 0.20 0.20 0.20 0.20 0.22 0.22 0.22 0.24 0.26 1.00
L6−CT 0.28 0.26 0.25 0.26 0.27 0.28 0.28 0.30 0.34 1.00
L56−NP 0.29 0.27 0.26 0.26 0.27 0.28 0.28 0.29 0.33 1.00
L5−IT 0.34 0.29 0.27 0.27 0.28 0.29 0.30 0.33 0.37 1.00
L5−ET 0.28 0.24 0.23 0.23 0.23 0.24 0.24 0.26 0.31 1.00
L4−IT 0.26 0.24 0.24 0.24 0.25 0.26 0.27 0.29 0.33 1.00
L23−IT 0.30 0.26 0.26 0.26 0.27 0.28 0.30 0.33 0.38 1.00
HIP−Misc2 0.29 0.26 0.27 0.27 0.28 0.29 0.30 0.32 0.36 1.00
HIP−Misc1 0.24 0.23 0.24 0.25 0.27 0.28 0.29 0.31 0.35 1.00
DG 0.24 0.25 0.25 0.26 0.29 0.31 0.34 0.37 0.45 1.00
CA3 0.33 0.27 0.25 0.25 0.25 0.26 0.28 0.29 0.34 1.00
CA1 0.24 0.21 0.21 0.22 0.22 0.23 0.25 0.28 0.31 1.00
Amy−Exc 0.28 0.25 0.25 0.25 0.26 0.27 0.28 0.30 0.35 1.00
Vip 0.31 0.30 0.31 0.32 0.32 0.33 0.34 0.36 0.41 1.00
THM−MB 0.33 0.26 0.26 0.26 0.26 0.27 0.28 0.31 0.34 1.00
THM−Inh 0.34 0.31 0.33 0.33 0.35 0.35 0.37 0.41 0.46 1.00
THM−Exc 0.41 0.32 0.31 0.31 0.30 0.30 0.32 0.33 0.38 1.00
SubCtx−Cplx 0.24 0.24 0.25 0.26 0.28 0.31 0.32 0.35 0.40 1.00
Sst 0.23 0.25 0.27 0.30 0.32 0.33 0.36 0.40 0.46 1.00
Sncg 0.31 0.32 0.33 0.34 0.35 0.36 0.37 0.40 0.44 1.00
Pvalb−ChC 0.26 0.25 0.26 0.28 0.29 0.30 0.33 0.34 0.39 1.00
Pvalb 1.13 1.09 1.04 0.80 0.67 0.80 0.91 0.86 1.02 1.00
PN 0.50 0.35 0.32 0.32 0.33 0.33 0.34 0.36 0.42 1.00
MSN−D2 0.28 0.26 0.27 0.28 0.28 0.29 0.30 0.31 0.35 1.00
MSN−D1 0.32 0.29 0.29 0.31 0.31 0.33 0.34 0.37 0.41 1.00
Lamp5−Lhx6 0.29 0.29 0.31 0.32 0.33 0.34 0.36 0.38 0.41 1.00
Lamp5 0.30 0.28 0.29 0.29 0.31 0.30 0.31 0.32 0.36 1.00
Foxp2 0.32 0.29 0.29 0.30 0.31 0.32 0.33 0.36 0.41 1.00
Chd7 0.29 0.27 0.27 0.27 0.28 0.28 0.29 0.30 0.34 1.00
CB 0.31 0.31 0.31 0.31 0.30 0.31 0.31 0.32 0.35 1.00
0.0−0.1 0.1−0.2 0.2−0.3 0.3−0.4 0.4−0.5 0.5−0.6 0.6−0.7 0.7−0.8 0.8−0.9 0.9−1.0
Percent
0.005 0.010 0.015 0.020
ASCT−Telencephalon astrocytes
L6b 0.50 0.37 0.35 0.35 0.36 0.36 0.37 0.39 0.43 1.00
L6−IT−Car3 0.50 0.42 0.41 0.41 0.42 0.42 0.43 0.46 0.51 1.00
L6−IT 0.31 0.29 0.29 0.30 0.31 0.32 0.32 0.33 0.34 1.00
L6−CT 0.41 0.37 0.36 0.36 0.36 0.37 0.38 0.39 0.42 1.00
L56−NP 0.41 0.37 0.36 0.37 0.38 0.38 0.39 0.39 0.42 1.00
L5−IT 0.49 0.39 0.38 0.37 0.38 0.39 0.40 0.42 0.46 1.00
L5−ET 0.40 0.34 0.32 0.32 0.32 0.33 0.33 0.35 0.39 1.00
L4−IT 0.38 0.33 0.33 0.34 0.34 0.34 0.36 0.38 0.42 1.00
L23−IT 0.43 0.37 0.37 0.37 0.37 0.38 0.39 0.42 0.46 1.00
HIP−Misc2 0.41 0.37 0.37 0.38 0.38 0.39 0.40 0.41 0.45 1.00
HIP−Misc1 0.36 0.34 0.35 0.36 0.37 0.38 0.40 0.41 0.44 1.00
DG 0.33 0.33 0.35 0.36 0.37 0.40 0.43 0.45 0.53 1.00
CA3 0.47 0.38 0.36 0.35 0.35 0.35 0.36 0.39 0.42 1.00
CA1 0.36 0.30 0.30 0.31 0.31 0.32 0.35 0.36 0.40 1.00
Amy−Exc 0.41 0.36 0.35 0.36 0.36 0.37 0.38 0.40 0.44 1.00
Vip 0.44 0.44 0.44 0.44 0.45 0.45 0.46 0.47 0.50 1.00
THM−MB 0.47 0.38 0.37 0.37 0.37 0.38 0.39 0.41 0.43 1.00
THM−Inh 0.44 0.41 0.42 0.42 0.44 0.45 0.47 0.50 0.55 1.00
THM−Exc 0.55 0.43 0.41 0.40 0.41 0.41 0.42 0.43 0.47 1.00
SubCtx−Cplx 0.33 0.33 0.35 0.37 0.38 0.41 0.43 0.46 0.50 1.00
Sst 0.32 0.35 0.37 0.38 0.41 0.43 0.45 0.49 0.55 1.00
Sncg 0.44 0.44 0.45 0.47 0.48 0.49 0.50 0.51 0.53 1.00
Pvalb−ChC 0.38 0.37 0.38 0.39 0.40 0.41 0.43 0.45 0.49 1.00
Pvalb 1.14 1.11 1.06 0.76 0.68 0.83 0.98 0.90 0.92 1.00
PN 0.67 0.48 0.44 0.43 0.43 0.43 0.44 0.47 0.51 1.00
MSN−D2 0.41 0.38 0.38 0.39 0.40 0.40 0.40 0.42 0.45 1.00
MSN−D1 0.46 0.41 0.41 0.43 0.43 0.44 0.45 0.48 0.51 1.00
Lamp5−Lhx6 0.43 0.42 0.42 0.44 0.44 0.46 0.48 0.48 0.51 1.00
Lamp5 0.45 0.42 0.43 0.42 0.42 0.43 0.44 0.44 0.45 1.00
Foxp2 0.46 0.42 0.42 0.42 0.42 0.43 0.45 0.46 0.49 1.00
Chd7 0.42 0.39 0.39 0.39 0.39 0.39 0.41 0.41 0.43 1.00
CB 0.46 0.45 0.45 0.43 0.44 0.43 0.42 0.43 0.44 1.00
0.0−0.1 0.1−0.2 0.2−0.3 0.3−0.4 0.4−0.5 0.5−0.6 0.6−0.7 0.7−0.8 0.8−0.9 0.9−1.0
Percent
0.010 0.015 0.020 0.025 0.030 0.035

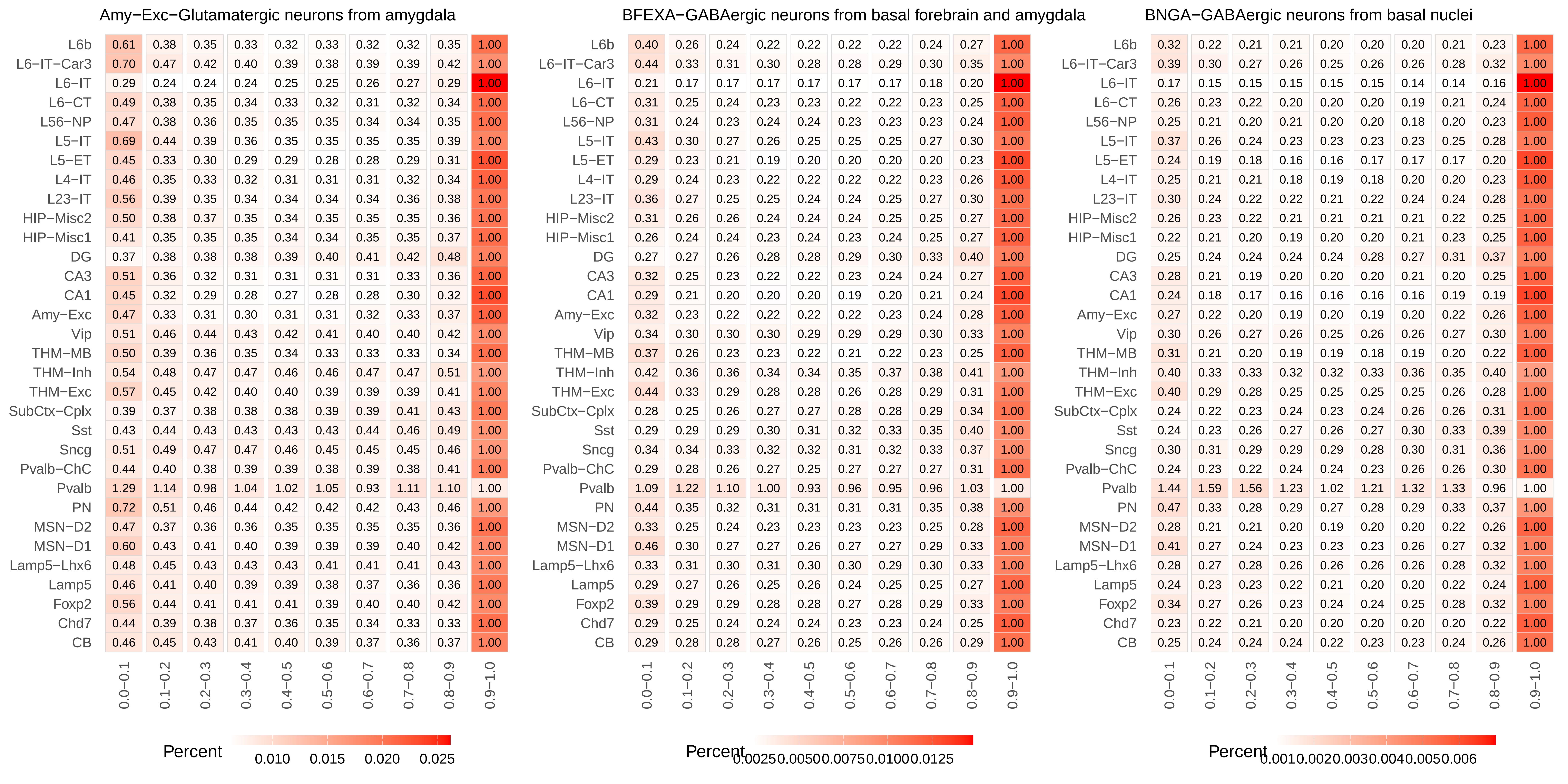
Amy−Exc−Glutamatergic neurons from amygdala
L6b 0.61 0.38 0.35 0.33 0.32 0.33 0.32 0.32 0.35 1.00
L6−IT−Car3 0.70 0.47 0.42 0.40 0.39 0.38 0.39 0.39 0.42 1.00
L6−IT 0.29 0.24 0.24 0.24 0.25 0.25 0.26 0.27 0.29 1.00
L6−CT 0.49 0.38 0.35 0.34 0.33 0.32 0.31 0.32 0.34 1.00
L56−NP 0.47 0.38 0.36 0.35 0.35 0.35 0.34 0.34 0.35 1.00
L5−IT 0.69 0.44 0.39 0.36 0.35 0.35 0.35 0.35 0.39 1.00
L5−ET 0.45 0.33 0.30 0.29 0.29 0.28 0.28 0.29 0.31 1.00
L4−IT 0.46 0.35 0.33 0.32 0.31 0.31 0.31 0.32 0.34 1.00
L23−IT 0.56 0.39 0.35 0.34 0.34 0.34 0.34 0.36 0.38 1.00
HIP−Misc2 0.50 0.38 0.37 0.35 0.34 0.35 0.35 0.35 0.36 1.00
HIP−Misc1 0.41 0.35 0.35 0.35 0.34 0.34 0.35 0.35 0.37 1.00
DG 0.37 0.38 0.38 0.38 0.39 0.40 0.41 0.42 0.48 1.00
CA3 0.51 0.36 0.32 0.31 0.31 0.31 0.31 0.33 0.36 1.00
CA1 0.45 0.32 0.29 0.28 0.27 0.28 0.28 0.30 0.32 1.00
Amy−Exc 0.47 0.33 0.31 0.30 0.31 0.31 0.32 0.33 0.37 1.00
Vip 0.51 0.46 0.44 0.43 0.42 0.41 0.40 0.40 0.42 1.00
THM−MB 0.50 0.39 0.36 0.35 0.34 0.33 0.33 0.33 0.34 1.00
THM−Inh 0.54 0.48 0.47 0.47 0.46 0.46 0.47 0.47 0.51 1.00
THM−Exc 0.57 0.45 0.42 0.40 0.40 0.39 0.39 0.39 0.41 1.00
SubCtx−Cplx 0.39 0.37 0.38 0.38 0.38 0.39 0.39 0.41 0.43 1.00
Sst 0.43 0.44 0.43 0.43 0.43 0.43 0.44 0.46 0.49 1.00
Sncg 0.51 0.49 0.47 0.47 0.46 0.45 0.45 0.45 0.46 1.00
Pvalb−ChC 0.44 0.40 0.38 0.39 0.39 0.38 0.39 0.38 0.41 1.00
Pvalb 1.29 1.14 0.98 1.04 1.02 1.05 0.93 1.11 1.10 1.00
PN 0.72 0.51 0.46 0.44 0.42 0.42 0.42 0.43 0.46 1.00
MSN−D2 0.47 0.37 0.36 0.36 0.35 0.35 0.35 0.35 0.36 1.00
MSN−D1 0.60 0.43 0.41 0.40 0.39 0.39 0.39 0.40 0.42 1.00
Lamp5−Lhx6 0.48 0.45 0.43 0.43 0.43 0.41 0.41 0.41 0.43 1.00
Lamp5 0.46 0.41 0.40 0.39 0.39 0.38 0.37 0.36 0.36 1.00
Foxp2 0.56 0.44 0.41 0.41 0.41 0.39 0.40 0.40 0.42 1.00
Chd7 0.44 0.39 0.38 0.37 0.36 0.35 0.34 0.33 0.33 1.00
CB 0.46 0.45 0.43 0.41 0.40 0.39 0.37 0.36 0.37 1.00
0.0−0.1 0.1−0.2 0.2−0.3 0.3−0.4 0.4−0.5 0.5−0.6 0.6−0.7 0.7−0.8 0.8−0.9 0.9−1.0
Percent 0.010 0.015 0.020 0.025
BFEXA−GABAergic neurons from basal forebrain and amygdala
L6b 0.40 0.26 0.24 0.22 0.22 0.22 0.22 0.24 0.27 1.00
L6−IT−Car3 0.44 0.33 0.31 0.30 0.28 0.28 0.29 0.30 0.35 1.00
L6−IT 0.21 0.17 0.17 0.17 0.17 0.17 0.17 0.18 0.20 1.00
L6−CT 0.31 0.25 0.24 0.23 0.23 0.22 0.22 0.23 0.25 1.00
L56−NP 0.31 0.24 0.23 0.24 0.24 0.23 0.23 0.23 0.24 1.00
L5−IT 0.43 0.30 0.27 0.26 0.25 0.25 0.25 0.27 0.30 1.00
L5−ET 0.29 0.23 0.21 0.19 0.20 0.20 0.20 0.20 0.23 1.00
L4−IT 0.29 0.24 0.23 0.22 0.22 0.22 0.22 0.23 0.26 1.00
L23−IT 0.36 0.27 0.25 0.25 0.24 0.24 0.25 0.27 0.30 1.00
HIP−Misc2 0.31 0.26 0.26 0.24 0.24 0.24 0.25 0.25 0.27 1.00
HIP−Misc1 0.26 0.24 0.24 0.23 0.24 0.23 0.24 0.25 0.27 1.00
DG 0.27 0.27 0.26 0.28 0.28 0.29 0.30 0.33 0.40 1.00
CA3 0.32 0.25 0.23 0.22 0.22 0.23 0.24 0.24 0.27 1.00
CA1 0.29 0.21 0.20 0.20 0.20 0.19 0.20 0.21 0.24 1.00
Amy−Exc 0.32 0.23 0.22 0.22 0.22 0.22 0.23 0.24 0.28 1.00
Vip 0.34 0.30 0.30 0.30 0.29 0.29 0.29 0.30 0.33 1.00
THM−MB 0.37 0.26 0.23 0.23 0.22 0.21 0.22 0.23 0.25 1.00
THM−Inh 0.42 0.36 0.36 0.34 0.34 0.35 0.37 0.38 0.41 1.00
THM−Exc 0.44 0.33 0.29 0.28 0.28 0.26 0.28 0.29 0.31 1.00
SubCtx−Cplx 0.28 0.25 0.26 0.27 0.27 0.28 0.28 0.29 0.34 1.00
Sst 0.29 0.29 0.29 0.30 0.31 0.32 0.33 0.35 0.40 1.00
Sncg 0.34 0.34 0.33 0.32 0.32 0.31 0.32 0.33 0.37 1.00
Pvalb−ChC 0.29 0.28 0.26 0.27 0.25 0.27 0.27 0.27 0.31 1.00
Pvalb 1.09 1.22 1.10 1.00 0.93 0.96 0.95 0.96 1.03 1.00
PN 0.44 0.35 0.32 0.31 0.31 0.31 0.31 0.35 0.38 1.00
MSN−D2 0.33 0.25 0.24 0.23 0.23 0.23 0.23 0.25 0.28 1.00
MSN−D1 0.46 0.30 0.27 0.27 0.26 0.27 0.27 0.29 0.33 1.00
Lamp5−Lhx6 0.33 0.31 0.30 0.31 0.30 0.30 0.29 0.30 0.33 1.00
Lamp5 0.29 0.27 0.26 0.25 0.26 0.24 0.25 0.25 0.27 1.00
Foxp2 0.39 0.29 0.29 0.28 0.28 0.27 0.28 0.29 0.33 1.00
Chd7 0.29 0.25 0.24 0.24 0.24 0.23 0.23 0.24 0.25 1.00
CB 0.29 0.28 0.28 0.27 0.26 0.25 0.26 0.26 0.29 1.00
0.0−0.1 0.1−0.2 0.2−0.3 0.3−0.4 0.4−0.5 0.5−0.6 0.6−0.7 0.7−0.8 0.8−0.9 0.9−1.0
Percent 0.0025 0.0050 0.0075 0.0100 0.0125
BNGA−GABAergic neurons from basal nuclei
L6b 0.32 0.22 0.21 0.21 0.20 0.20 0.20 0.21 0.23 1.00
L6−IT−Car3 0.39 0.30 0.27 0.26 0.25 0.26 0.26 0.28 0.32 1.00
L6−IT 0.17 0.15 0.15 0.15 0.15 0.15 0.14 0.14 0.16 1.00
L6−CT 0.26 0.23 0.22 0.20 0.20 0.20 0.19 0.21 0.24 1.00
L56−NP 0.25 0.21 0.20 0.21 0.20 0.20 0.18 0.20 0.23 1.00
L5−IT 0.37 0.26 0.24 0.23 0.23 0.23 0.23 0.25 0.28 1.00
L5−ET 0.24 0.19 0.18 0.16 0.16 0.17 0.17 0.17 0.20 1.00
L4−IT 0.25 0.21 0.21 0.18 0.19 0.18 0.20 0.20 0.23 1.00
L23−IT 0.30 0.24 0.22 0.22 0.21 0.22 0.24 0.24 0.28 1.00
HIP−Misc2 0.26 0.23 0.22 0.21 0.21 0.21 0.21 0.22 0.25 1.00
HIP−Misc1 0.22 0.21 0.20 0.19 0.20 0.20 0.21 0.23 0.25 1.00
DG 0.25 0.24 0.24 0.24 0.24 0.28 0.27 0.31 0.37 1.00
CA3 0.28 0.21 0.19 0.20 0.20 0.20 0.21 0.20 0.25 1.00
CA1 0.24 0.18 0.17 0.16 0.16 0.16 0.16 0.19 0.19 1.00
Amy−Exc 0.27 0.22 0.20 0.19 0.20 0.19 0.20 0.22 0.26 1.00
Vip 0.30 0.26 0.27 0.26 0.25 0.26 0.26 0.27 0.30 1.00
THM−MB 0.31 0.21 0.20 0.19 0.19 0.18 0.19 0.20 0.22 1.00
THM−Inh 0.40 0.33 0.33 0.32 0.32 0.33 0.36 0.35 0.40 1.00
THM−Exc 0.40 0.29 0.28 0.25 0.25 0.25 0.25 0.26 0.28 1.00
SubCtx−Cplx 0.24 0.22 0.23 0.24 0.23 0.24 0.26 0.26 0.31 1.00
Sst 0.24 0.23 0.26 0.27 0.26 0.27 0.30 0.33 0.39 1.00
Sncg 0.30 0.31 0.29 0.29 0.29 0.28 0.30 0.31 0.36 1.00
Pvalb−ChC 0.24 0.23 0.22 0.24 0.24 0.23 0.26 0.26 0.30 1.00
Pvalb 1.44 1.59 1.56 1.23 1.02 1.21 1.32 1.33 0.96 1.00
PN 0.47 0.33 0.28 0.29 0.27 0.28 0.29 0.33 0.37 1.00
MSN−D2 0.28 0.21 0.21 0.20 0.19 0.20 0.20 0.22 0.26 1.00
MSN−D1 0.41 0.27 0.24 0.23 0.23 0.23 0.26 0.27 0.32 1.00
Lamp5−Lhx6 0.28 0.27 0.28 0.26 0.26 0.26 0.26 0.28 0.32 1.00
Lamp5 0.24 0.23 0.23 0.22 0.21 0.20 0.20 0.22 0.24 1.00
Foxp2 0.34 0.27 0.26 0.23 0.24 0.24 0.25 0.28 0.32 1.00
Chd7 0.23 0.22 0.21 0.20 0.20 0.20 0.20 0.20 0.22 1.00
CB 0.25 0.24 0.24 0.24 0.22 0.23 0.23 0.24 0.26 1.00
0.0−0.1 0.1−0.2 0.2−0.3 0.3−0.4 0.4−0.5 0.5−0.6 0.6−0.7 0.7−0.8 0.8−0.9 0.9−1.0
Percent 0.001 0.002 0.003 0.004 0.005 0.006

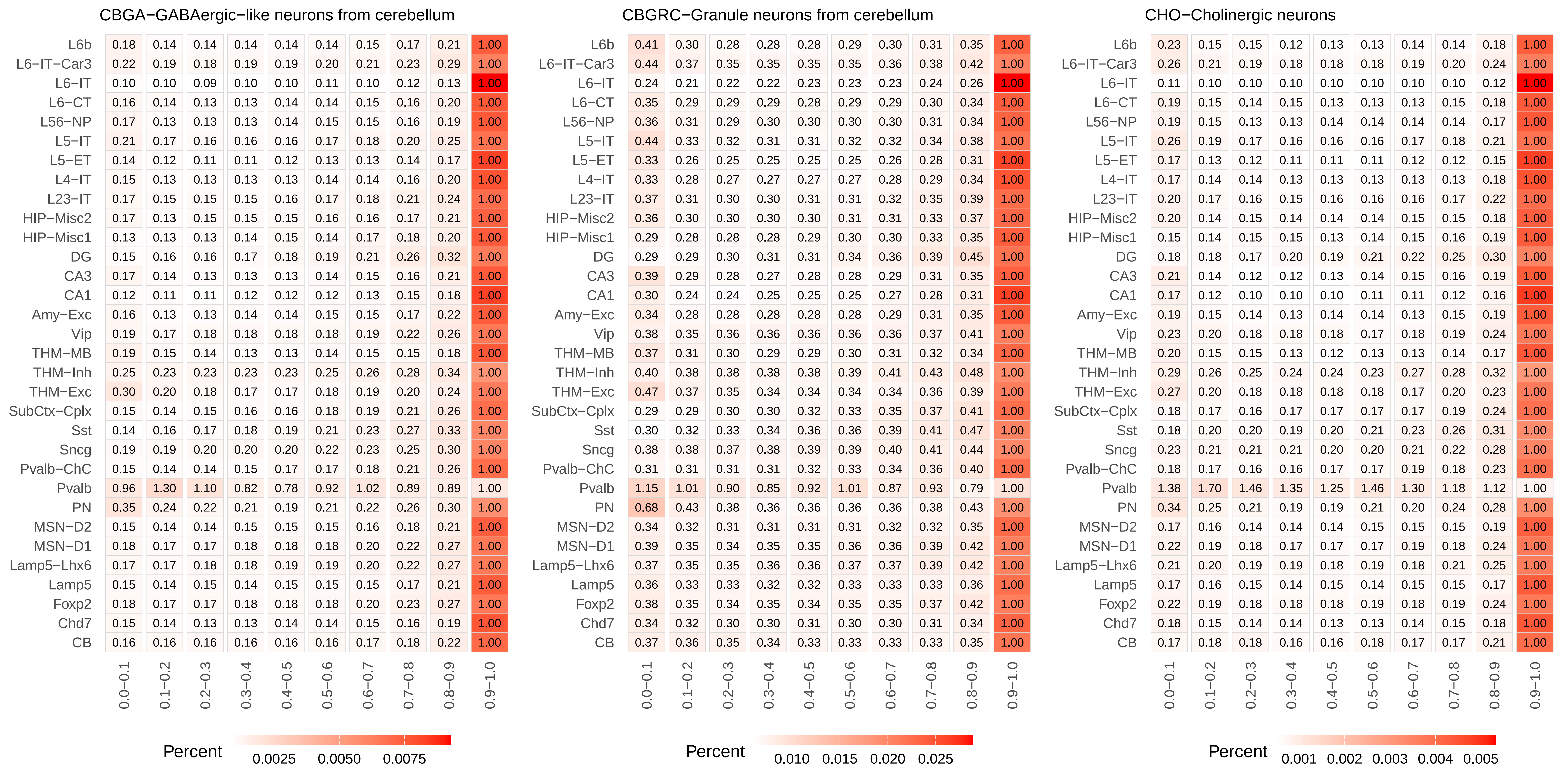
CBGA−GABAergic−like neurons from cerebellum
L6b 0.18 0.14 0.14 0.14 0.14 0.14 0.15 0.17 0.21 1.00
L6−IT−Car3 0.22 0.19 0.18 0.19 0.19 0.20 0.21 0.23 0.29 1.00
L6−IT 0.10 0.10 0.09 0.10 0.10 0.11 0.10 0.12 0.13 1.00
L6−CT 0.16 0.14 0.13 0.13 0.14 0.14 0.15 0.16 0.20 1.00
L56−NP 0.17 0.13 0.13 0.13 0.14 0.15 0.15 0.16 0.19 1.00
L5−IT 0.21 0.17 0.16 0.16 0.16 0.17 0.18 0.20 0.25 1.00
L5−ET 0.14 0.12 0.11 0.11 0.12 0.13 0.13 0.14 0.17 1.00
L4−IT 0.15 0.13 0.13 0.13 0.13 0.14 0.14 0.16 0.20 1.00
L23−IT 0.17 0.15 0.15 0.15 0.16 0.17 0.18 0.21 0.24 1.00
HIP−Misc2 0.17 0.13 0.15 0.15 0.15 0.16 0.16 0.17 0.21 1.00
HIP−Misc1 0.13 0.13 0.13 0.14 0.15 0.14 0.17 0.18 0.20 1.00
DG 0.15 0.16 0.16 0.17 0.18 0.19 0.21 0.26 0.32 1.00
CA3 0.17 0.14 0.13 0.13 0.13 0.14 0.15 0.16 0.21 1.00
CA1 0.12 0.11 0.11 0.12 0.12 0.12 0.13 0.15 0.18 1.00
Amy−Exc 0.16 0.13 0.13 0.14 0.14 0.15 0.15 0.17 0.22 1.00
Vip 0.19 0.17 0.18 0.18 0.18 0.18 0.19 0.22 0.26 1.00
THM−MB 0.19 0.15 0.14 0.13 0.13 0.14 0.15 0.15 0.18 1.00
THM−Inh 0.25 0.23 0.23 0.23 0.23 0.25 0.26 0.28 0.34 1.00
THM−Exc 0.30 0.20 0.18 0.17 0.17 0.18 0.19 0.20 0.24 1.00
SubCtx−Cplx 0.15 0.14 0.15 0.16 0.16 0.18 0.19 0.21 0.26 1.00
Sst 0.14 0.16 0.17 0.18 0.19 0.21 0.23 0.27 0.33 1.00
Sncg 0.19 0.19 0.20 0.20 0.20 0.22 0.23 0.25 0.30 1.00
Pvalb−ChC 0.15 0.14 0.14 0.15 0.17 0.17 0.18 0.21 0.26 1.00
Pvalb 0.96 1.30 1.10 0.82 0.78 0.92 1.02 0.89 0.89 1.00
PN 0.35 0.24 0.22 0.21 0.19 0.21 0.22 0.26 0.30 1.00
MSN−D2 0.15 0.14 0.14 0.15 0.15 0.15 0.16 0.18 0.21 1.00
MSN−D1 0.18 0.17 0.17 0.18 0.18 0.18 0.20 0.22 0.27 1.00
Lamp5−Lhx6 0.17 0.17 0.18 0.18 0.19 0.19 0.20 0.22 0.27 1.00
Lamp5 0.15 0.14 0.15 0.14 0.15 0.15 0.15 0.17 0.21 1.00
Foxp2 0.18 0.17 0.17 0.18 0.18 0.18 0.20 0.23 0.27 1.00
Chd7 0.15 0.14 0.13 0.13 0.14 0.14 0.15 0.16 0.19 1.00
CB 0.16 0.16 0.16 0.16 0.16 0.16 0.17 0.18 0.22 1.00
0.0−0.1 0.1−0.2 0.2−0.3 0.3−0.4 0.4−0.5 0.5−0.6 0.6−0.7 0.7−0.8 0.8−0.9 0.9−1.0
Percent
0.0025 0.0050 0.0075
CBGRC−Granule neurons from cerebellum
L6b 0.41 0.30 0.28 0.28 0.28 0.29 0.30 0.31 0.35 1.00
L6−IT−Car3 0.44 0.37 0.35 0.35 0.35 0.35 0.36 0.38 0.42 1.00
L6−IT 0.24 0.21 0.22 0.22 0.23 0.23 0.23 0.24 0.26 1.00
L6−CT 0.35 0.29 0.29 0.29 0.28 0.29 0.29 0.30 0.34 1.00
L56−NP 0.36 0.31 0.29 0.30 0.30 0.30 0.30 0.31 0.34 1.00
L5−IT 0.44 0.33 0.32 0.31 0.31 0.32 0.32 0.34 0.38 1.00
L5−ET 0.33 0.26 0.25 0.25 0.25 0.25 0.26 0.28 0.31 1.00
L4−IT 0.33 0.28 0.27 0.27 0.27 0.27 0.28 0.29 0.34 1.00
L23−IT 0.37 0.31 0.30 0.30 0.31 0.31 0.32 0.35 0.39 1.00
HIP−Misc2 0.36 0.30 0.30 0.30 0.30 0.31 0.31 0.33 0.37 1.00
HIP−Misc1 0.29 0.28 0.28 0.28 0.29 0.30 0.30 0.33 0.35 1.00
DG 0.29 0.29 0.30 0.31 0.31 0.34 0.36 0.39 0.45 1.00
CA3 0.39 0.29 0.28 0.27 0.28 0.28 0.29 0.31 0.35 1.00
CA1 0.30 0.24 0.24 0.25 0.25 0.25 0.27 0.28 0.31 1.00
Amy−Exc 0.34 0.28 0.28 0.28 0.28 0.28 0.29 0.31 0.35 1.00
Vip 0.38 0.35 0.36 0.36 0.36 0.36 0.36 0.37 0.41 1.00
THM−MB 0.37 0.31 0.30 0.29 0.29 0.30 0.31 0.32 0.34 1.00
THM−Inh 0.40 0.38 0.38 0.38 0.38 0.39 0.41 0.43 0.48 1.00
THM−Exc 0.47 0.37 0.35 0.34 0.34 0.34 0.34 0.36 0.39 1.00
SubCtx−Cplx 0.29 0.29 0.30 0.30 0.32 0.33 0.35 0.37 0.41 1.00
Sst 0.30 0.32 0.33 0.34 0.36 0.36 0.39 0.41 0.47 1.00
Sncg 0.38 0.38 0.37 0.38 0.39 0.39 0.40 0.41 0.44 1.00
Pvalb−ChC 0.31 0.31 0.31 0.31 0.32 0.33 0.34 0.36 0.40 1.00
Pvalb 1.15 1.01 0.90 0.85 0.92 1.01 0.87 0.93 0.79 1.00
PN 0.68 0.43 0.38 0.36 0.36 0.36 0.36 0.38 0.43 1.00
MSN−D2 0.34 0.32 0.31 0.31 0.31 0.31 0.32 0.32 0.35 1.00
MSN−D1 0.39 0.35 0.34 0.35 0.35 0.36 0.36 0.39 0.42 1.00
Lamp5−Lhx6 0.37 0.35 0.35 0.36 0.36 0.37 0.37 0.39 0.42 1.00
Lamp5 0.36 0.33 0.33 0.32 0.32 0.33 0.33 0.33 0.36 1.00
Foxp2 0.38 0.35 0.34 0.35 0.34 0.35 0.35 0.37 0.42 1.00
Chd7 0.34 0.32 0.30 0.30 0.31 0.30 0.30 0.31 0.34 1.00
CB 0.37 0.36 0.35 0.34 0.33 0.33 0.33 0.33 0.35 1.00
0.0−0.1 0.1−0.2 0.2−0.3 0.3−0.4 0.4−0.5 0.5−0.6 0.6−0.7 0.7−0.8 0.8−0.9 0.9−1.0
Percent
0.010 0.015 0.020 0.025
CHO−Cholinergic neurons
L6b 0.23 0.15 0.15 0.12 0.13 0.13 0.14 0.14 0.18 1.00
L6−IT−Car3 0.26 0.21 0.19 0.18 0.18 0.18 0.19 0.20 0.24 1.00
L6−IT 0.11 0.10 0.10 0.10 0.10 0.10 0.10 0.10 0.12 1.00
L6−CT 0.19 0.15 0.14 0.15 0.13 0.13 0.13 0.15 0.18 1.00
L56−NP 0.19 0.15 0.13 0.13 0.14 0.14 0.14 0.14 0.17 1.00
L5−IT 0.26 0.19 0.17 0.16 0.16 0.16 0.17 0.18 0.21 1.00
L5−ET 0.17 0.13 0.12 0.11 0.11 0.11 0.12 0.12 0.15 1.00
L4−IT 0.17 0.14 0.14 0.13 0.13 0.13 0.13 0.13 0.18 1.00
L23−IT 0.20 0.17 0.16 0.15 0.16 0.16 0.16 0.17 0.22 1.00
HIP−Misc2 0.20 0.14 0.15 0.14 0.14 0.14 0.15 0.15 0.18 1.00
HIP−Misc1 0.15 0.14 0.15 0.15 0.13 0.14 0.15 0.16 0.19 1.00
DG 0.18 0.18 0.17 0.20 0.19 0.21 0.22 0.25 0.30 1.00
CA3 0.21 0.14 0.12 0.12 0.13 0.14 0.15 0.16 0.19 1.00
CA1 0.17 0.12 0.10 0.10 0.10 0.11 0.11 0.12 0.16 1.00
Amy−Exc 0.19 0.15 0.14 0.13 0.14 0.14 0.13 0.15 0.19 1.00
Vip 0.23 0.20 0.18 0.18 0.18 0.17 0.18 0.19 0.24 1.00
THM−MB 0.20 0.15 0.15 0.13 0.12 0.13 0.13 0.14 0.17 1.00
THM−Inh 0.29 0.26 0.25 0.24 0.24 0.23 0.27 0.28 0.32 1.00
THM−Exc 0.27 0.20 0.18 0.18 0.18 0.18 0.17 0.20 0.23 1.00
SubCtx−Cplx 0.18 0.17 0.16 0.17 0.17 0.17 0.17 0.19 0.24 1.00
Sst 0.18 0.20 0.20 0.19 0.20 0.21 0.23 0.26 0.31 1.00
Sncg 0.23 0.21 0.21 0.21 0.20 0.20 0.21 0.22 0.28 1.00
Pvalb−ChC 0.18 0.17 0.16 0.16 0.17 0.17 0.19 0.18 0.23 1.00
Pvalb 1.38 1.70 1.46 1.35 1.25 1.46 1.30 1.18 1.12 1.00
PN 0.34 0.25 0.21 0.19 0.19 0.21 0.20 0.24 0.28 1.00
MSN−D2 0.17 0.16 0.14 0.14 0.14 0.15 0.15 0.15 0.19 1.00
MSN−D1 0.22 0.19 0.18 0.17 0.17 0.17 0.19 0.18 0.24 1.00
Lamp5−Lhx6 0.21 0.20 0.19 0.19 0.18 0.19 0.18 0.21 0.25 1.00
Lamp5 0.17 0.16 0.15 0.14 0.15 0.14 0.15 0.15 0.17 1.00
Foxp2 0.22 0.19 0.18 0.18 0.18 0.19 0.18 0.19 0.24 1.00
Chd7 0.18 0.15 0.14 0.14 0.13 0.13 0.14 0.15 0.18 1.00
CB 0.17 0.18 0.18 0.16 0.16 0.18 0.17 0.17 0.21 1.00
0.0−0.1 0.1−0.2 0.2−0.3 0.3−0.4 0.4−0.5 0.5−0.6 0.6−0.7 0.7−0.8 0.8−0.9 0.9−1.0
Percent
0.001 0.002 0.003 0.004 0.005

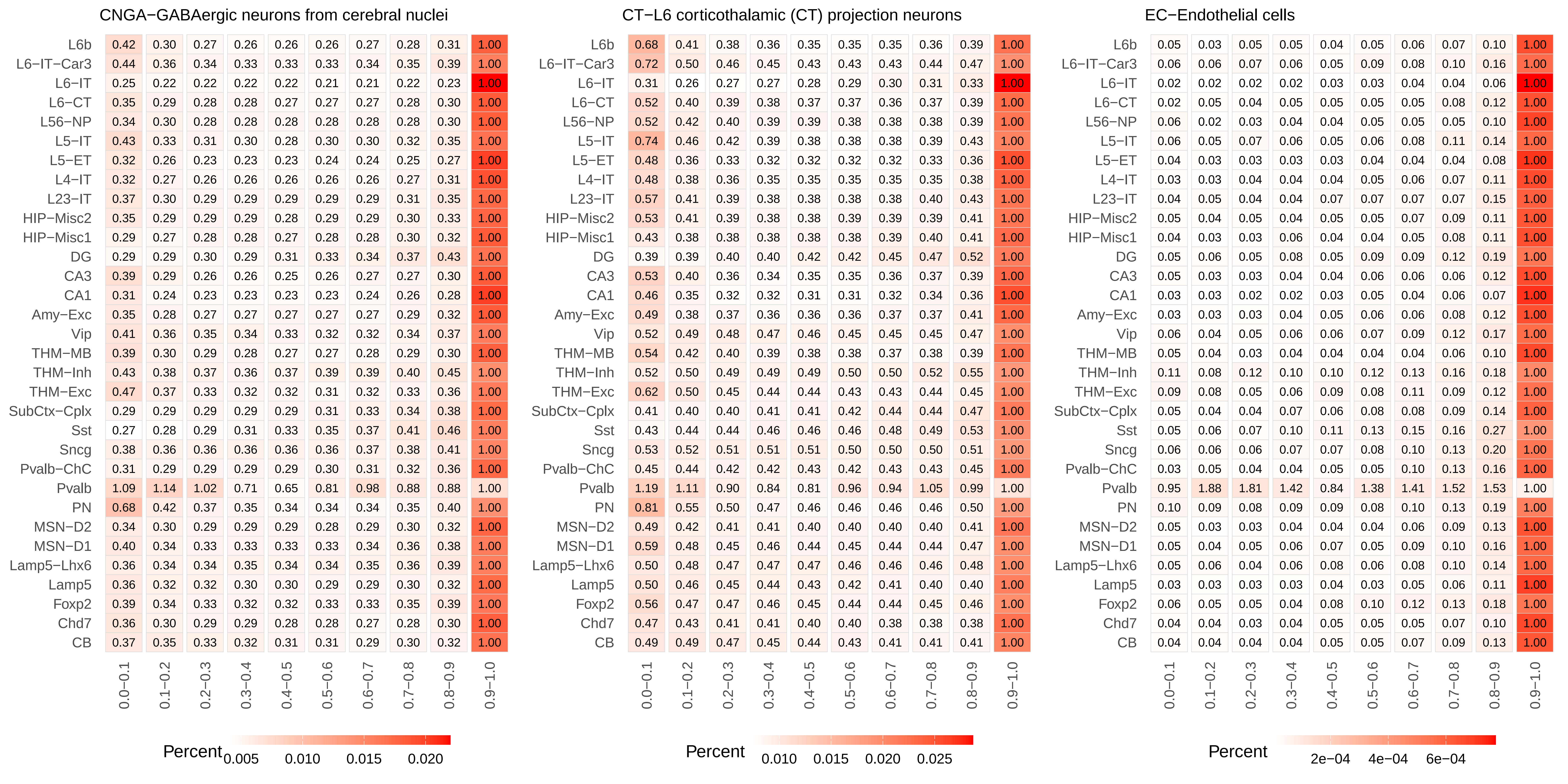

CNGA−GABAergic neurons from cerebral nuclei
L6b 0.42 0.30 0.27 0.26 0.26 0.26 0.27 0.28 0.31 1.00
L6−IT−Car3 0.44 0.36 0.34 0.33 0.33 0.33 0.34 0.35 0.39 1.00
L6−IT 0.25 0.22 0.22 0.22 0.22 0.21 0.21 0.22 0.23 1.00
L6−CT 0.35 0.29 0.28 0.28 0.27 0.27 0.27 0.28 0.30 1.00
L56−NP 0.34 0.30 0.28 0.28 0.28 0.28 0.28 0.28 0.30 1.00
L5−IT 0.43 0.33 0.31 0.30 0.28 0.30 0.30 0.32 0.35 1.00
L5−ET 0.32 0.26 0.23 0.23 0.23 0.24 0.24 0.25 0.27 1.00
L4−IT 0.32 0.27 0.26 0.26 0.26 0.26 0.26 0.27 0.31 1.00
L23−IT 0.37 0.30 0.29 0.29 0.29 0.29 0.29 0.31 0.35 1.00
HIP−Misc2 0.35 0.29 0.29 0.29 0.28 0.29 0.29 0.30 0.33 1.00
HIP−Misc1 0.29 0.27 0.28 0.28 0.27 0.28 0.28 0.30 0.32 1.00
DG 0.29 0.29 0.30 0.29 0.31 0.33 0.34 0.37 0.43 1.00
CA3 0.39 0.29 0.26 0.26 0.25 0.26 0.27 0.27 0.30 1.00
CA1 0.31 0.24 0.23 0.23 0.23 0.23 0.24 0.26 0.28 1.00
Amy−Exc 0.35 0.28 0.27 0.27 0.27 0.27 0.27 0.29 0.32 1.00
Vip 0.41 0.36 0.35 0.34 0.33 0.32 0.32 0.34 0.37 1.00
THM−MB 0.39 0.30 0.29 0.28 0.27 0.27 0.28 0.29 0.30 1.00
THM−Inh 0.43 0.38 0.37 0.36 0.37 0.39 0.39 0.40 0.45 1.00
THM−Exc 0.47 0.37 0.33 0.32 0.32 0.31 0.32 0.33 0.36 1.00
SubCtx−Cplx 0.29 0.29 0.29 0.29 0.29 0.31 0.33 0.34 0.38 1.00
Sst 0.27 0.28 0.29 0.31 0.33 0.35 0.37 0.41 0.46 1.00
Sncg 0.38 0.36 0.36 0.36 0.36 0.36 0.37 0.38 0.41 1.00
Pvalb−ChC 0.31 0.29 0.29 0.29 0.29 0.30 0.31 0.32 0.36 1.00
Pvalb 1.09 1.14 1.02 0.71 0.65 0.81 0.98 0.88 0.88 1.00
PN 0.68 0.42 0.37 0.35 0.34 0.34 0.34 0.35 0.40 1.00
MSN−D2 0.34 0.30 0.29 0.29 0.29 0.28 0.29 0.30 0.32 1.00
MSN−D1 0.40 0.34 0.33 0.33 0.33 0.33 0.34 0.36 0.38 1.00
Lamp5−Lhx6 0.36 0.34 0.34 0.35 0.34 0.34 0.35 0.36 0.39 1.00
Lamp5 0.36 0.32 0.32 0.30 0.30 0.29 0.29 0.30 0.32 1.00
Foxp2 0.39 0.34 0.33 0.32 0.32 0.33 0.33 0.35 0.39 1.00
Chd7 0.36 0.30 0.29 0.29 0.28 0.28 0.27 0.28 0.30 1.00
CB 0.37 0.35 0.33 0.32 0.31 0.31 0.29 0.30 0.32 1.00
0.0−0.1 0.1−0.2 0.2−0.3 0.3−0.4 0.4−0.5 0.5−0.6 0.6−0.7 0.7−0.8 0.8−0.9 0.9−1.0
Percent 0.005 0.010 0.015 0.020
CT−L6 corticothalamic (CT) projection neurons
L6b 0.68 0.41 0.38 0.36 0.35 0.35 0.35 0.36 0.39 1.00
L6−IT−Car3 0.72 0.50 0.46 0.45 0.43 0.43 0.43 0.44 0.47 1.00
L6−IT 0.31 0.26 0.27 0.27 0.28 0.29 0.30 0.31 0.33 1.00
L6−CT 0.52 0.40 0.39 0.38 0.37 0.37 0.36 0.37 0.39 1.00
L56−NP 0.52 0.42 0.40 0.39 0.39 0.38 0.38 0.38 0.39 1.00
L5−IT 0.74 0.46 0.42 0.39 0.38 0.38 0.38 0.39 0.43 1.00
L5−ET 0.48 0.36 0.33 0.32 0.32 0.32 0.32 0.33 0.36 1.00
L4−IT 0.48 0.38 0.36 0.35 0.35 0.35 0.35 0.35 0.38 1.00
L23−IT 0.57 0.41 0.39 0.38 0.38 0.38 0.38 0.40 0.43 1.00
HIP−Misc2 0.53 0.41 0.39 0.38 0.38 0.39 0.39 0.39 0.41 1.00
HIP−Misc1 0.43 0.38 0.38 0.38 0.38 0.38 0.39 0.40 0.41 1.00
DG 0.39 0.39 0.40 0.40 0.42 0.42 0.45 0.47 0.52 1.00
CA3 0.53 0.40 0.36 0.34 0.35 0.35 0.36 0.37 0.39 1.00
CA1 0.46 0.35 0.32 0.32 0.31 0.31 0.32 0.34 0.36 1.00
Amy−Exc 0.49 0.38 0.37 0.36 0.36 0.36 0.37 0.37 0.41 1.00
Vip 0.52 0.49 0.48 0.47 0.46 0.45 0.45 0.45 0.47 1.00
THM−MB 0.54 0.42 0.40 0.39 0.38 0.38 0.37 0.38 0.39 1.00
THM−Inh 0.52 0.50 0.49 0.49 0.49 0.50 0.50 0.52 0.55 1.00
THM−Exc 0.62 0.50 0.45 0.44 0.44 0.43 0.43 0.44 0.45 1.00
SubCtx−Cplx 0.41 0.40 0.40 0.41 0.41 0.42 0.44 0.44 0.47 1.00
Sst 0.43 0.44 0.44 0.46 0.46 0.46 0.48 0.49 0.53 1.00
Sncg 0.53 0.52 0.51 0.51 0.51 0.50 0.50 0.50 0.51 1.00
Pvalb−ChC 0.45 0.44 0.42 0.42 0.43 0.42 0.43 0.43 0.45 1.00
Pvalb 1.19 1.11 0.90 0.84 0.81 0.96 0.94 1.05 0.99 1.00
PN 0.81 0.55 0.50 0.47 0.46 0.46 0.46 0.46 0.50 1.00
MSN−D2 0.49 0.42 0.41 0.41 0.40 0.40 0.40 0.40 0.41 1.00
MSN−D1 0.59 0.48 0.45 0.46 0.44 0.45 0.44 0.44 0.47 1.00
Lamp5−Lhx6 0.50 0.48 0.47 0.47 0.47 0.46 0.46 0.46 0.48 1.00
Lamp5 0.50 0.46 0.45 0.44 0.43 0.42 0.41 0.40 0.40 1.00
Foxp2 0.56 0.47 0.47 0.46 0.45 0.44 0.44 0.45 0.46 1.00
Chd7 0.47 0.43 0.41 0.41 0.40 0.40 0.38 0.38 0.38 1.00
CB 0.49 0.49 0.47 0.45 0.44 0.43 0.41 0.41 0.41 1.00
0.0−0.1 0.1−0.2 0.2−0.3 0.3−0.4 0.4−0.5 0.5−0.6 0.6−0.7 0.7−0.8 0.8−0.9 0.9−1.0
Percent 0.010 0.015 0.020 0.025
EC−Endothelial cells
L6b 0.05 0.03 0.05 0.05 0.04 0.05 0.06 0.07 0.10 1.00
L6−IT−Car3 0.06 0.06 0.07 0.06 0.05 0.09 0.08 0.10 0.16 1.00
L6−IT 0.02 0.02 0.02 0.02 0.03 0.03 0.04 0.04 0.06 1.00
L6−CT 0.02 0.05 0.04 0.05 0.05 0.05 0.05 0.08 0.12 1.00
L56−NP 0.06 0.02 0.03 0.04 0.04 0.05 0.05 0.05 0.10 1.00
L5−IT 0.06 0.05 0.07 0.06 0.05 0.06 0.08 0.11 0.14 1.00
L5−ET 0.04 0.03 0.03 0.03 0.03 0.04 0.04 0.04 0.08 1.00
L4−IT 0.03 0.03 0.04 0.04 0.04 0.05 0.06 0.07 0.11 1.00
L23−IT 0.04 0.05 0.04 0.04 0.07 0.07 0.07 0.07 0.15 1.00
HIP−Misc2 0.05 0.04 0.05 0.04 0.05 0.05 0.07 0.09 0.11 1.00
HIP−Misc1 0.04 0.03 0.03 0.06 0.04 0.04 0.05 0.08 0.11 1.00
DG 0.05 0.06 0.05 0.08 0.05 0.09 0.09 0.12 0.19 1.00
CA3 0.05 0.03 0.03 0.04 0.04 0.06 0.06 0.06 0.12 1.00
CA1 0.03 0.03 0.02 0.02 0.03 0.05 0.04 0.06 0.07 1.00
Amy−Exc 0.03 0.03 0.03 0.04 0.05 0.06 0.06 0.08 0.12 1.00
Vip 0.06 0.04 0.05 0.06 0.06 0.07 0.09 0.12 0.17 1.00
THM−MB 0.05 0.04 0.03 0.04 0.04 0.04 0.04 0.06 0.10 1.00
THM−Inh 0.11 0.08 0.12 0.10 0.10 0.12 0.13 0.16 0.18 1.00
THM−Exc 0.09 0.08 0.05 0.06 0.09 0.08 0.11 0.09 0.12 1.00
SubCtx−Cplx 0.05 0.04 0.04 0.07 0.06 0.08 0.08 0.09 0.14 1.00
Sst 0.05 0.06 0.07 0.10 0.11 0.13 0.15 0.16 0.27 1.00
Sncg 0.06 0.06 0.06 0.07 0.07 0.08 0.10 0.13 0.20 1.00
Pvalb−ChC 0.03 0.05 0.04 0.04 0.05 0.05 0.10 0.13 0.16 1.00
Pvalb 0.95 1.88 1.81 1.42 0.84 1.38 1.41 1.52 1.53 1.00
PN 0.10 0.09 0.08 0.09 0.09 0.08 0.10 0.13 0.19 1.00
MSN−D2 0.05 0.03 0.03 0.04 0.04 0.04 0.06 0.09 0.13 1.00
MSN−D1 0.05 0.04 0.05 0.06 0.07 0.05 0.09 0.10 0.16 1.00
Lamp5−Lhx6 0.05 0.06 0.04 0.06 0.08 0.06 0.08 0.10 0.14 1.00
Lamp5 0.03 0.03 0.03 0.03 0.04 0.03 0.05 0.06 0.11 1.00
Foxp2 0.06 0.05 0.05 0.04 0.08 0.10 0.12 0.13 0.18 1.00
Chd7 0.04 0.04 0.03 0.04 0.05 0.05 0.05 0.07 0.10 1.00
CB 0.04 0.04 0.04 0.04 0.05 0.05 0.07 0.09 0.13 1.00
0.0−0.1 0.1−0.2 0.2−0.3 0.3−0.4 0.4−0.5 0.5−0.6 0.6−0.7 0.7−0.8 0.8−0.9 0.9−1.0
Percent 2e−04 4e−04 6e−04

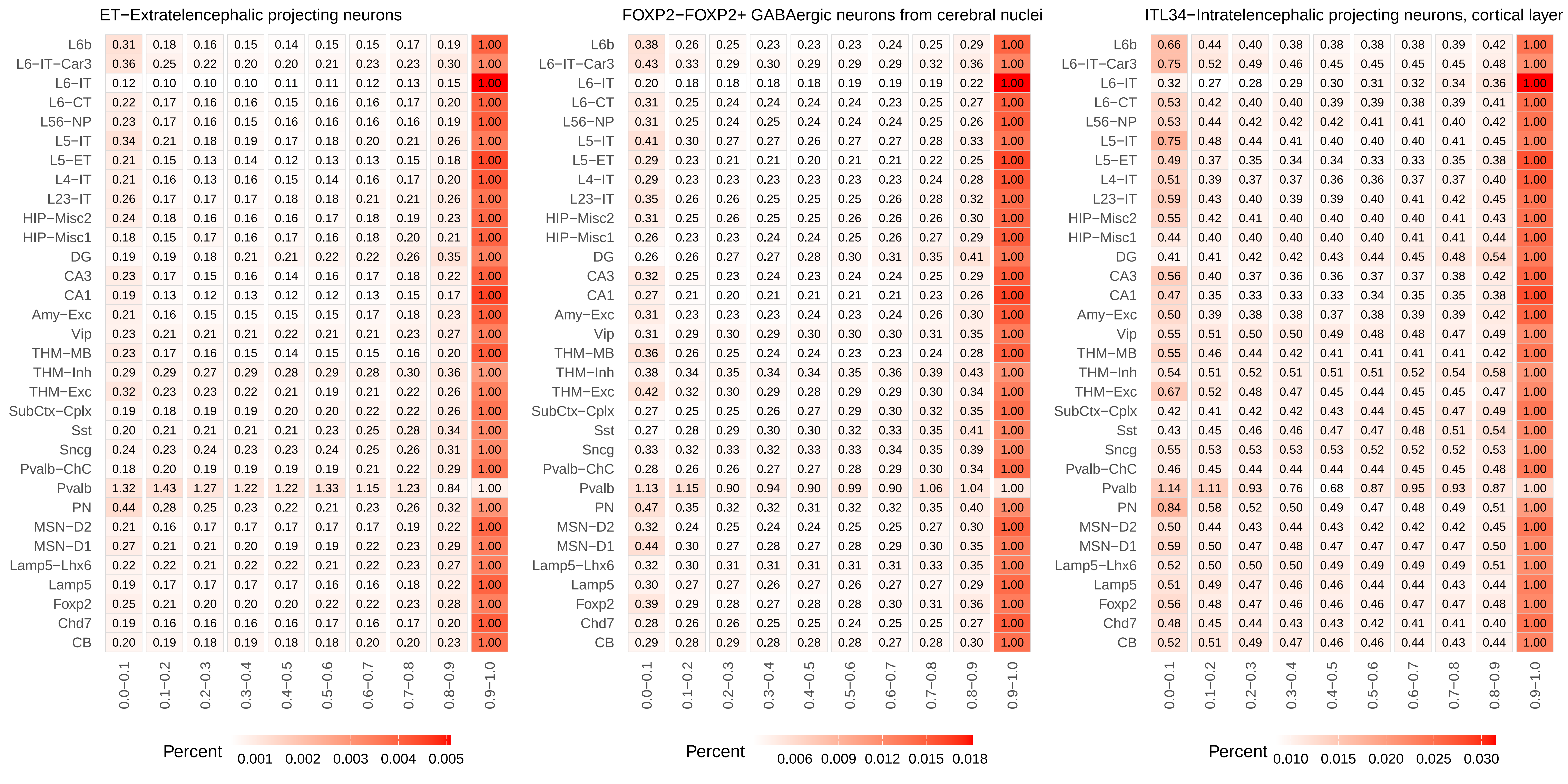
ET−Extratelencephalic projecting neurons
L6b 0.31 0.18 0.16 0.15 0.14 0.15 0.15 0.17 0.19 1.00
L6−IT−Car3 0.36 0.25 0.22 0.20 0.20 0.21 0.23 0.23 0.30 1.00
L6−IT 0.12 0.10 0.10 0.10 0.11 0.11 0.12 0.13 0.15 1.00
L6−CT 0.22 0.17 0.16 0.16 0.15 0.16 0.16 0.17 0.20 1.00
L56−NP 0.23 0.17 0.16 0.15 0.16 0.16 0.16 0.16 0.19 1.00
L5−IT 0.34 0.21 0.18 0.19 0.17 0.18 0.20 0.21 0.26 1.00
L5−ET 0.21 0.15 0.13 0.14 0.12 0.13 0.13 0.15 0.18 1.00
L4−IT 0.21 0.16 0.13 0.16 0.15 0.14 0.16 0.17 0.20 1.00
L23−IT 0.26 0.17 0.17 0.17 0.18 0.18 0.21 0.21 0.26 1.00
HIP−Misc2 0.24 0.18 0.16 0.16 0.16 0.17 0.18 0.19 0.23 1.00
HIP−Misc1 0.18 0.15 0.17 0.16 0.17 0.16 0.18 0.20 0.21 1.00
DG 0.19 0.19 0.18 0.21 0.21 0.22 0.22 0.26 0.35 1.00
CA3 0.23 0.17 0.15 0.16 0.14 0.16 0.17 0.18 0.22 1.00
CA1 0.19 0.13 0.12 0.13 0.12 0.12 0.13 0.15 0.17 1.00
Amy−Exc 0.21 0.16 0.15 0.15 0.15 0.15 0.17 0.18 0.23 1.00
Vip 0.23 0.21 0.21 0.21 0.22 0.21 0.21 0.23 0.27 1.00
THM−MB 0.23 0.17 0.16 0.15 0.14 0.15 0.15 0.16 0.20 1.00
THM−Inh 0.29 0.29 0.27 0.29 0.28 0.29 0.28 0.30 0.36 1.00
THM−Exc 0.32 0.23 0.23 0.22 0.21 0.19 0.21 0.22 0.26 1.00
SubCtx−Cplx 0.19 0.18 0.19 0.19 0.20 0.20 0.22 0.22 0.26 1.00
Sst 0.20 0.21 0.21 0.21 0.21 0.23 0.25 0.28 0.34 1.00
Sncg 0.24 0.23 0.24 0.23 0.23 0.24 0.25 0.26 0.31 1.00
Pvalb−ChC 0.18 0.20 0.19 0.19 0.19 0.19 0.21 0.22 0.29 1.00
Pvalb 1.32 1.43 1.27 1.22 1.22 1.33 1.15 1.23 0.84 1.00
PN 0.44 0.28 0.25 0.23 0.22 0.21 0.23 0.26 0.32 1.00
MSN−D2 0.21 0.16 0.17 0.17 0.17 0.17 0.17 0.19 0.22 1.00
MSN−D1 0.27 0.21 0.21 0.20 0.19 0.19 0.22 0.23 0.29 1.00
Lamp5−Lhx6 0.22 0.22 0.21 0.22 0.22 0.21 0.22 0.23 0.27 1.00
Lamp5 0.19 0.17 0.17 0.17 0.17 0.16 0.16 0.18 0.22 1.00
Foxp2 0.25 0.21 0.20 0.20 0.20 0.22 0.22 0.23 0.28 1.00
Chd7 0.19 0.16 0.16 0.16 0.16 0.17 0.16 0.17 0.20 1.00
CB 0.20 0.19 0.18 0.19 0.18 0.18 0.20 0.20 0.23 1.00
0.0−0.1 0.1−0.2 0.2−0.3 0.3−0.4 0.4−0.5 0.5−0.6 0.6−0.7 0.7−0.8 0.8−0.9 0.9−1.0
Percent 0.001 0.002 0.003 0.004 0.005
FOXP2−FOXP2+ GABAergic neurons from cerebral nuclei
L6b 0.38 0.26 0.25 0.23 0.23 0.23 0.24 0.25 0.29 1.00
L6−IT−Car3 0.43 0.33 0.29 0.30 0.29 0.29 0.29 0.32 0.36 1.00
L6−IT 0.20 0.18 0.18 0.18 0.18 0.19 0.19 0.19 0.22 1.00
L6−CT 0.31 0.25 0.24 0.24 0.24 0.24 0.23 0.25 0.27 1.00
L56−NP 0.31 0.25 0.24 0.25 0.24 0.24 0.24 0.25 0.26 1.00
L5−IT 0.41 0.30 0.27 0.27 0.26 0.27 0.27 0.28 0.33 1.00
L5−ET 0.29 0.23 0.21 0.21 0.20 0.21 0.21 0.22 0.25 1.00
L4−IT 0.29 0.23 0.23 0.23 0.23 0.23 0.23 0.24 0.28 1.00
L23−IT 0.35 0.26 0.26 0.25 0.25 0.25 0.26 0.28 0.32 1.00
HIP−Misc2 0.31 0.25 0.26 0.25 0.25 0.26 0.26 0.26 0.30 1.00
HIP−Misc1 0.26 0.23 0.23 0.24 0.24 0.25 0.26 0.27 0.29 1.00
DG 0.26 0.26 0.27 0.27 0.28 0.30 0.31 0.35 0.41 1.00
CA3 0.32 0.25 0.23 0.24 0.23 0.24 0.24 0.25 0.29 1.00
CA1 0.27 0.21 0.20 0.21 0.21 0.21 0.21 0.23 0.26 1.00
Amy−Exc 0.31 0.23 0.23 0.23 0.24 0.23 0.24 0.26 0.30 1.00
Vip 0.31 0.29 0.30 0.29 0.30 0.30 0.30 0.31 0.35 1.00
THM−MB 0.36 0.26 0.25 0.24 0.24 0.23 0.23 0.24 0.28 1.00
THM−Inh 0.38 0.34 0.35 0.34 0.34 0.35 0.36 0.39 0.43 1.00
THM−Exc 0.42 0.32 0.30 0.29 0.28 0.29 0.29 0.30 0.34 1.00
SubCtx−Cplx 0.27 0.25 0.25 0.26 0.27 0.29 0.30 0.32 0.35 1.00
Sst 0.27 0.28 0.29 0.30 0.30 0.32 0.33 0.35 0.41 1.00
Sncg 0.33 0.32 0.33 0.32 0.33 0.33 0.34 0.35 0.39 1.00
Pvalb−ChC 0.28 0.26 0.26 0.27 0.27 0.28 0.29 0.30 0.34 1.00
Pvalb 1.13 1.15 0.90 0.94 0.90 0.99 0.90 1.06 1.04 1.00
PN 0.47 0.35 0.32 0.32 0.31 0.32 0.32 0.35 0.40 1.00
MSN−D2 0.32 0.24 0.25 0.24 0.24 0.25 0.25 0.27 0.30 1.00
MSN−D1 0.44 0.30 0.27 0.28 0.27 0.28 0.29 0.30 0.35 1.00
Lamp5−Lhx6 0.32 0.30 0.31 0.31 0.31 0.31 0.31 0.33 0.35 1.00
Lamp5 0.30 0.27 0.27 0.26 0.27 0.26 0.27 0.27 0.29 1.00
Foxp2 0.39 0.29 0.28 0.27 0.28 0.28 0.30 0.31 0.36 1.00
Chd7 0.28 0.26 0.26 0.25 0.25 0.24 0.25 0.25 0.27 1.00
CB 0.29 0.28 0.29 0.28 0.28 0.28 0.27 0.28 0.30 1.00
0.0−0.1 0.1−0.2 0.2−0.3 0.3−0.4 0.4−0.5 0.5−0.6 0.6−0.7 0.7−0.8 0.8−0.9 0.9−1.0
Percent 0.006 0.009 0.012 0.015 0.018
ITL34−Intratelencephalic projecting neurons, cortical layer
L6b 0.66 0.44 0.40 0.38 0.38 0.38 0.38 0.39 0.42 1.00
L6−IT−Car3 0.75 0.52 0.49 0.46 0.45 0.45 0.45 0.45 0.48 1.00
L6−IT 0.32 0.27 0.28 0.29 0.30 0.31 0.32 0.34 0.36 1.00
L6−CT 0.53 0.42 0.40 0.40 0.39 0.39 0.38 0.39 0.41 1.00
L56−NP 0.53 0.44 0.42 0.42 0.42 0.41 0.41 0.40 0.42 1.00
L5−IT 0.75 0.48 0.44 0.41 0.40 0.40 0.40 0.41 0.45 1.00
L5−ET 0.49 0.37 0.35 0.34 0.34 0.33 0.33 0.35 0.38 1.00
L4−IT 0.51 0.39 0.37 0.37 0.36 0.36 0.37 0.37 0.40 1.00
L23−IT 0.59 0.43 0.40 0.39 0.39 0.40 0.41 0.42 0.45 1.00
HIP−Misc2 0.55 0.42 0.41 0.40 0.40 0.40 0.40 0.41 0.43 1.00
HIP−Misc1 0.44 0.40 0.40 0.40 0.40 0.40 0.41 0.41 0.44 1.00
DG 0.41 0.41 0.42 0.42 0.43 0.44 0.45 0.48 0.54 1.00
CA3 0.56 0.40 0.37 0.36 0.36 0.37 0.37 0.38 0.42 1.00
CA1 0.47 0.35 0.33 0.33 0.33 0.34 0.35 0.35 0.38 1.00
Amy−Exc 0.50 0.39 0.38 0.38 0.37 0.38 0.39 0.39 0.42 1.00
Vip 0.55 0.51 0.50 0.50 0.49 0.48 0.48 0.47 0.49 1.00
THM−MB 0.55 0.46 0.44 0.42 0.41 0.41 0.41 0.41 0.42 1.00
THM−Inh 0.54 0.51 0.52 0.51 0.51 0.51 0.52 0.54 0.58 1.00
THM−Exc 0.67 0.52 0.48 0.47 0.45 0.44 0.45 0.45 0.47 1.00
SubCtx−Cplx 0.42 0.41 0.42 0.42 0.43 0.44 0.45 0.47 0.49 1.00
Sst 0.43 0.45 0.46 0.46 0.47 0.47 0.48 0.51 0.54 1.00
Sncg 0.55 0.53 0.53 0.53 0.53 0.52 0.52 0.52 0.53 1.00
Pvalb−ChC 0.46 0.45 0.44 0.44 0.44 0.44 0.45 0.45 0.48 1.00
Pvalb 1.14 1.11 0.93 0.76 0.68 0.87 0.95 0.93 0.87 1.00
PN 0.84 0.58 0.52 0.50 0.49 0.47 0.48 0.49 0.51 1.00
MSN−D2 0.50 0.44 0.43 0.44 0.43 0.42 0.42 0.42 0.45 1.00
MSN−D1 0.59 0.50 0.47 0.48 0.47 0.47 0.47 0.47 0.50 1.00
Lamp5−Lhx6 0.52 0.50 0.50 0.50 0.49 0.49 0.49 0.49 0.51 1.00
Lamp5 0.51 0.49 0.47 0.46 0.46 0.44 0.44 0.43 0.44 1.00
Foxp2 0.56 0.48 0.47 0.46 0.46 0.46 0.47 0.47 0.48 1.00
Chd7 0.48 0.45 0.44 0.43 0.43 0.42 0.41 0.41 0.40 1.00
CB 0.52 0.51 0.49 0.47 0.46 0.46 0.44 0.43 0.44 1.00
0.0−0.1 0.1−0.2 0.2−0.3 0.3−0.4 0.4−0.5 0.5−0.6 0.6−0.7 0.7−0.8 0.8−0.9 0.9−1.0
Percent 0.010 0.015 0.020 0.025 0.030

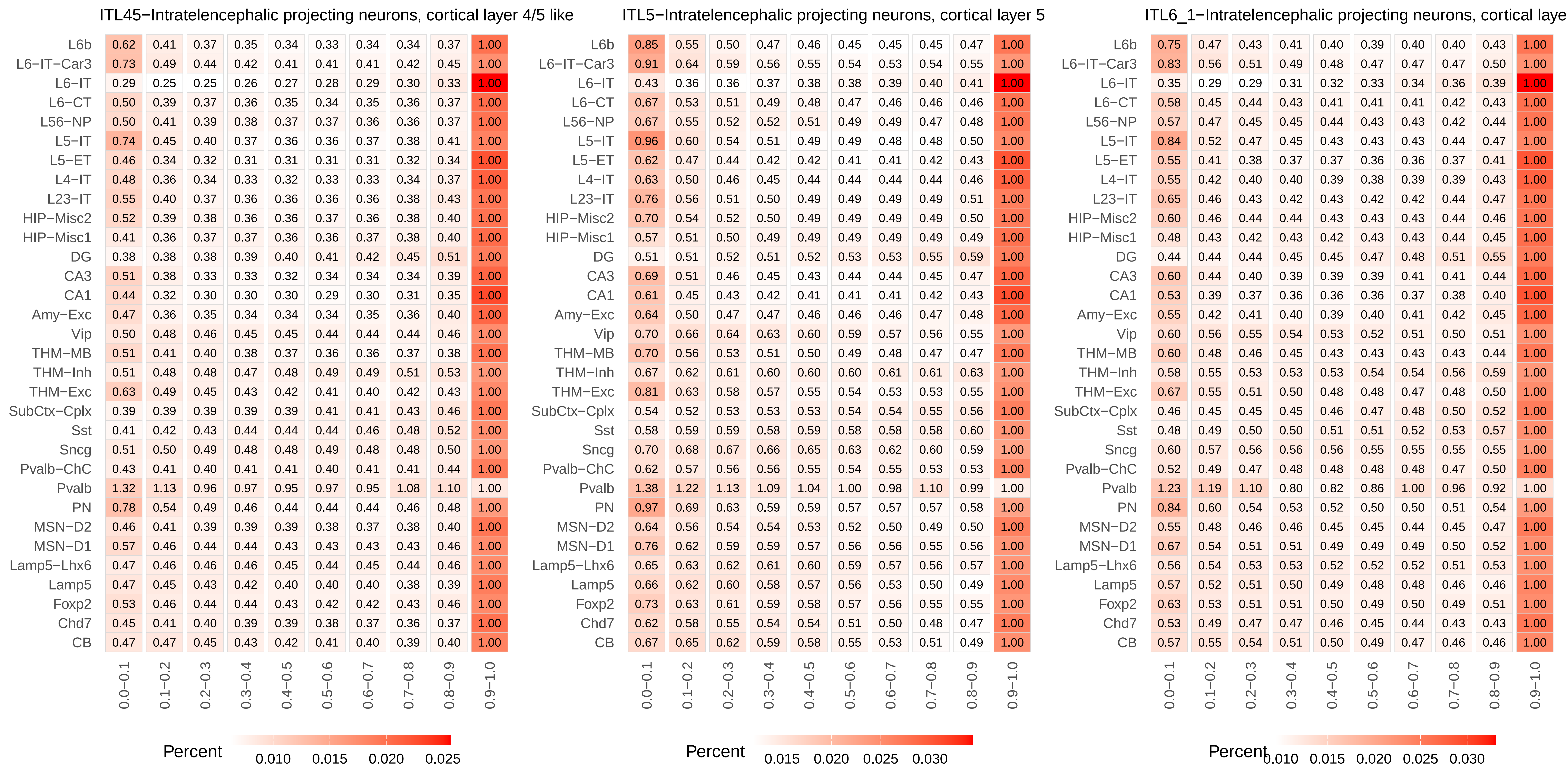
ITL45−Intratelencephalic projecting neurons, cortical layer 4/5 like
L6b 0.62 0.41 0.37 0.35 0.34 0.33 0.34 0.34 0.37 1.00
L6−IT−Car3 0.73 0.49 0.44 0.42 0.41 0.41 0.41 0.42 0.45 1.00
L6−IT 0.29 0.25 0.25 0.26 0.27 0.28 0.29 0.30 0.33 1.00
L6−CT 0.50 0.39 0.37 0.36 0.35 0.34 0.35 0.36 0.37 1.00
L56−NP 0.50 0.41 0.39 0.38 0.37 0.37 0.36 0.36 0.37 1.00
L5−IT 0.74 0.45 0.40 0.37 0.36 0.36 0.37 0.38 0.41 1.00
L5−ET 0.46 0.34 0.32 0.31 0.31 0.31 0.31 0.32 0.34 1.00
L4−IT 0.48 0.36 0.34 0.33 0.32 0.33 0.33 0.34 0.37 1.00
L23−IT 0.55 0.40 0.37 0.36 0.36 0.36 0.36 0.38 0.43 1.00
HIP−Misc2 0.52 0.39 0.38 0.36 0.36 0.37 0.36 0.38 0.40 1.00
HIP−Misc1 0.41 0.36 0.37 0.37 0.36 0.36 0.37 0.38 0.40 1.00
DG 0.38 0.38 0.38 0.39 0.40 0.41 0.42 0.45 0.51 1.00
CA3 0.51 0.38 0.33 0.33 0.32 0.34 0.34 0.34 0.39 1.00
CA1 0.44 0.32 0.30 0.30 0.30 0.29 0.30 0.31 0.35 1.00
Amy−Exc 0.47 0.36 0.35 0.34 0.34 0.34 0.35 0.36 0.40 1.00
Vip 0.50 0.48 0.46 0.45 0.45 0.44 0.44 0.44 0.46 1.00
THM−MB 0.51 0.41 0.40 0.38 0.37 0.36 0.36 0.37 0.38 1.00
THM−Inh 0.51 0.48 0.48 0.47 0.48 0.49 0.49 0.51 0.53 1.00
THM−Exc 0.63 0.49 0.45 0.43 0.42 0.41 0.40 0.42 0.43 1.00
SubCtx−Cplx 0.39 0.39 0.39 0.39 0.39 0.41 0.41 0.43 0.46 1.00
Sst 0.41 0.42 0.43 0.44 0.44 0.44 0.46 0.48 0.52 1.00
Sncg 0.51 0.50 0.49 0.48 0.48 0.49 0.48 0.48 0.50 1.00
Pvalb−ChC 0.43 0.41 0.40 0.41 0.41 0.40 0.41 0.41 0.44 1.00
Pvalb 1.32 1.13 0.96 0.97 0.95 0.97 0.95 1.08 1.10 1.00
PN 0.78 0.54 0.49 0.46 0.44 0.44 0.44 0.46 0.48 1.00
MSN−D2 0.46 0.41 0.39 0.39 0.39 0.38 0.37 0.38 0.40 1.00
MSN−D1 0.57 0.46 0.44 0.44 0.43 0.43 0.43 0.43 0.46 1.00
Lamp5−Lhx6 0.47 0.46 0.46 0.46 0.45 0.44 0.45 0.44 0.46 1.00
Lamp5 0.47 0.45 0.43 0.42 0.40 0.40 0.40 0.38 0.39 1.00
Foxp2 0.53 0.46 0.44 0.44 0.43 0.42 0.42 0.43 0.46 1.00
Chd7 0.45 0.41 0.40 0.39 0.39 0.38 0.37 0.36 0.37 1.00
CB 0.47 0.47 0.45 0.43 0.42 0.41 0.40 0.39 0.40 1.00
0.0−0.1 0.1−0.2 0.2−0.3 0.3−0.4 0.4−0.5 0.5−0.6 0.6−0.7 0.7−0.8 0.8−0.9 0.9−1.0
Percent
0.010 0.015 0.020 0.025
ITL5−Intratelencephalic projecting neurons, cortical layer 5
L6b 0.85 0.55 0.50 0.47 0.46 0.45 0.45 0.45 0.47 1.00
L6−IT−Car3 0.91 0.64 0.59 0.56 0.55 0.54 0.53 0.54 0.55 1.00
L6−IT 0.43 0.36 0.36 0.37 0.38 0.38 0.39 0.40 0.41 1.00
L6−CT 0.67 0.53 0.51 0.49 0.48 0.47 0.46 0.46 0.46 1.00
L56−NP 0.67 0.55 0.52 0.52 0.51 0.49 0.49 0.47 0.48 1.00
L5−IT 0.96 0.60 0.54 0.51 0.49 0.49 0.48 0.48 0.50 1.00
L5−ET 0.62 0.47 0.44 0.42 0.42 0.41 0.41 0.42 0.43 1.00
L4−IT 0.63 0.50 0.46 0.45 0.44 0.44 0.44 0.44 0.46 1.00
L23−IT 0.76 0.56 0.51 0.50 0.49 0.49 0.49 0.49 0.51 1.00
HIP−Misc2 0.70 0.54 0.52 0.50 0.49 0.49 0.49 0.49 0.50 1.00
HIP−Misc1 0.57 0.51 0.50 0.49 0.49 0.49 0.49 0.49 0.49 1.00
DG 0.51 0.51 0.52 0.51 0.52 0.53 0.53 0.55 0.59 1.00
CA3 0.69 0.51 0.46 0.45 0.43 0.44 0.44 0.45 0.47 1.00
CA1 0.61 0.45 0.43 0.42 0.41 0.41 0.41 0.42 0.43 1.00
Amy−Exc 0.64 0.50 0.47 0.47 0.46 0.46 0.46 0.47 0.48 1.00
Vip 0.70 0.66 0.64 0.63 0.60 0.59 0.57 0.56 0.55 1.00
THM−MB 0.70 0.56 0.53 0.51 0.50 0.49 0.48 0.47 0.47 1.00
THM−Inh 0.67 0.62 0.61 0.60 0.60 0.60 0.61 0.61 0.63 1.00
THM−Exc 0.81 0.63 0.58 0.57 0.55 0.54 0.53 0.53 0.55 1.00
SubCtx−Cplx 0.54 0.52 0.53 0.53 0.53 0.54 0.54 0.55 0.56 1.00
Sst 0.58 0.59 0.59 0.58 0.59 0.58 0.58 0.58 0.60 1.00
Sncg 0.70 0.68 0.67 0.66 0.65 0.63 0.62 0.60 0.59 1.00
Pvalb−ChC 0.62 0.57 0.56 0.56 0.55 0.54 0.55 0.53 0.53 1.00
Pvalb 1.38 1.22 1.13 1.09 1.04 1.00 0.98 1.10 0.99 1.00
PN 0.97 0.69 0.63 0.59 0.59 0.57 0.57 0.57 0.58 1.00
MSN−D2 0.64 0.56 0.54 0.54 0.53 0.52 0.50 0.49 0.50 1.00
MSN−D1 0.76 0.62 0.59 0.59 0.57 0.56 0.56 0.55 0.56 1.00
Lamp5−Lhx6 0.65 0.63 0.62 0.61 0.60 0.59 0.57 0.56 0.57 1.00
Lamp5 0.66 0.62 0.60 0.58 0.57 0.56 0.53 0.50 0.49 1.00
Foxp2 0.73 0.63 0.61 0.59 0.58 0.57 0.56 0.55 0.55 1.00
Chd7 0.62 0.58 0.55 0.54 0.54 0.51 0.50 0.48 0.47 1.00
CB 0.67 0.65 0.62 0.59 0.58 0.55 0.53 0.51 0.49 1.00
0.0−0.1 0.1−0.2 0.2−0.3 0.3−0.4 0.4−0.5 0.5−0.6 0.6−0.7 0.7−0.8 0.8−0.9 0.9−1.0
Percent
0.015 0.020 0.025 0.030
ITL6_1−Intratelencephalic projecting neurons, cortical laye
L6b 0.75 0.47 0.43 0.41 0.40 0.39 0.40 0.40 0.43 1.00
L6−IT−Car3 0.83 0.56 0.51 0.49 0.48 0.47 0.47 0.47 0.50 1.00
L6−IT 0.35 0.29 0.29 0.31 0.32 0.33 0.34 0.36 0.39 1.00
L6−CT 0.58 0.45 0.44 0.43 0.41 0.41 0.41 0.42 0.43 1.00
L56−NP 0.57 0.47 0.45 0.45 0.44 0.43 0.43 0.42 0.44 1.00
L5−IT 0.84 0.52 0.47 0.45 0.43 0.43 0.43 0.44 0.47 1.00
L5−ET 0.55 0.41 0.38 0.37 0.37 0.36 0.36 0.37 0.41 1.00
L4−IT 0.55 0.42 0.40 0.40 0.39 0.38 0.39 0.39 0.43 1.00
L23−IT 0.65 0.46 0.43 0.42 0.43 0.42 0.42 0.44 0.47 1.00
HIP−Misc2 0.60 0.46 0.44 0.44 0.43 0.43 0.43 0.44 0.46 1.00
HIP−Misc1 0.48 0.43 0.42 0.43 0.42 0.43 0.43 0.44 0.45 1.00
DG 0.44 0.44 0.44 0.45 0.45 0.47 0.48 0.51 0.55 1.00
CA3 0.60 0.44 0.40 0.39 0.39 0.39 0.41 0.41 0.44 1.00
CA1 0.53 0.39 0.37 0.36 0.36 0.36 0.37 0.38 0.40 1.00
Amy−Exc 0.55 0.42 0.41 0.40 0.39 0.40 0.41 0.42 0.45 1.00
Vip 0.60 0.56 0.55 0.54 0.53 0.52 0.51 0.50 0.51 1.00
THM−MB 0.60 0.48 0.46 0.45 0.43 0.43 0.43 0.43 0.44 1.00
THM−Inh 0.58 0.55 0.53 0.53 0.53 0.54 0.54 0.56 0.59 1.00
THM−Exc 0.67 0.55 0.51 0.50 0.48 0.48 0.47 0.48 0.50 1.00
SubCtx−Cplx 0.46 0.45 0.45 0.45 0.46 0.47 0.48 0.50 0.52 1.00
Sst 0.48 0.49 0.50 0.50 0.51 0.51 0.52 0.53 0.57 1.00
Sncg 0.60 0.57 0.56 0.56 0.56 0.55 0.55 0.55 0.55 1.00
Pvalb−ChC 0.52 0.49 0.47 0.48 0.48 0.48 0.48 0.47 0.50 1.00
Pvalb 1.23 1.19 1.10 0.80 0.82 0.86 1.00 0.96 0.92 1.00
PN 0.84 0.60 0.54 0.53 0.52 0.50 0.50 0.51 0.54 1.00
MSN−D2 0.55 0.48 0.46 0.46 0.45 0.45 0.44 0.45 0.47 1.00
MSN−D1 0.67 0.54 0.51 0.51 0.49 0.49 0.49 0.50 0.52 1.00
Lamp5−Lhx6 0.56 0.54 0.53 0.53 0.52 0.52 0.52 0.51 0.53 1.00
Lamp5 0.57 0.52 0.51 0.50 0.49 0.48 0.48 0.46 0.46 1.00
Foxp2 0.63 0.53 0.51 0.51 0.50 0.49 0.50 0.49 0.51 1.00
Chd7 0.53 0.49 0.47 0.47 0.46 0.45 0.44 0.43 0.43 1.00
CB 0.57 0.55 0.54 0.51 0.50 0.49 0.47 0.46 0.46 1.00
0.0−0.1 0.1−0.2 0.2−0.3 0.3−0.4 0.4−0.5 0.5−0.6 0.6−0.7 0.7−0.8 0.8−0.9 0.9−1.0
Percent
0.010 0.015 0.020 0.025 0.030

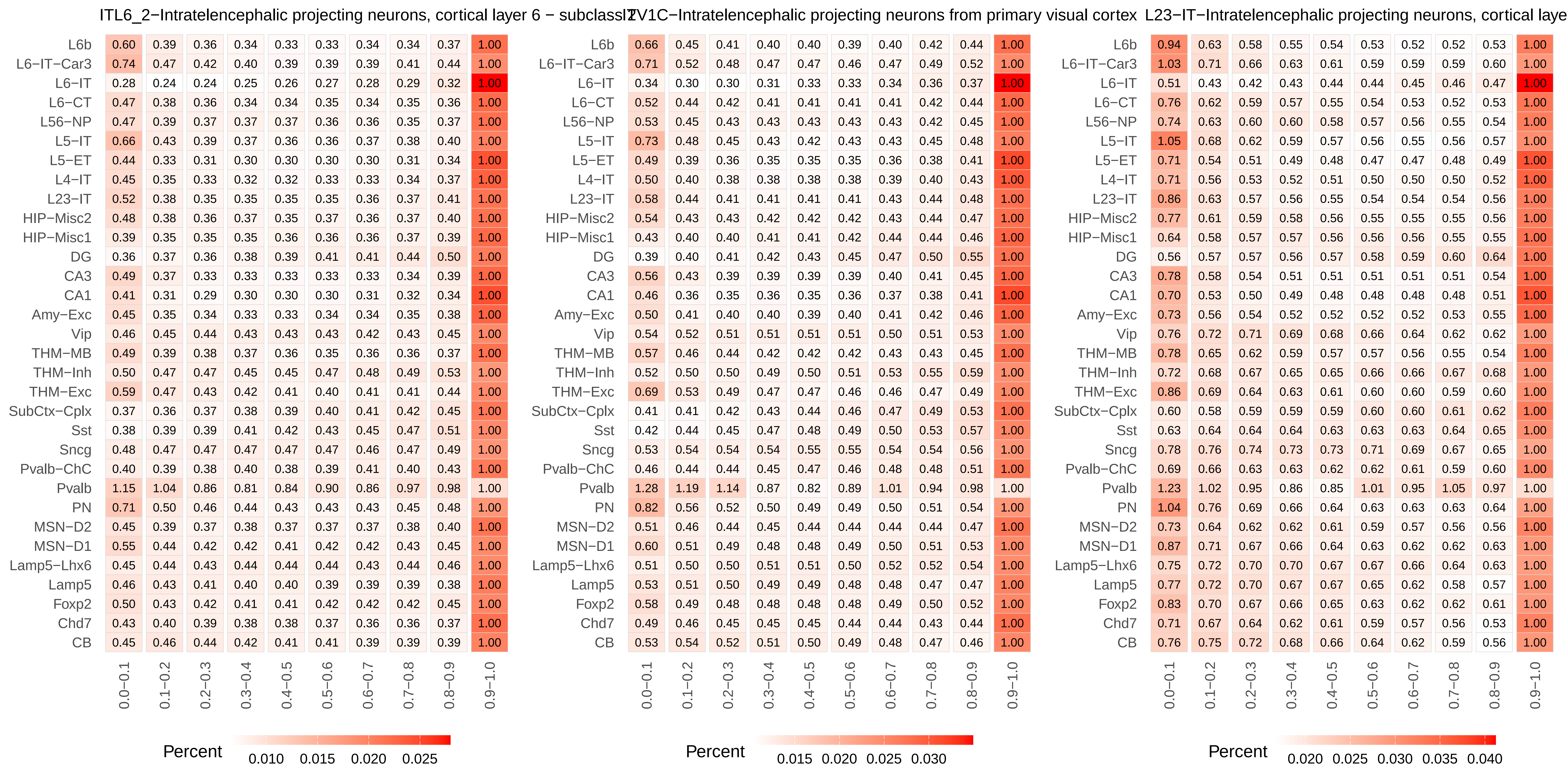
ITL6_2−Intratelencephalic projecting neurons, cortical layer 6 − subclass
ITV1C−Intratelencephalic projecting neurons from primary visual cortex
L23−IT−Intratelencephalic projecting neurons, cortical laye
0.0−0.1 0.1−0.2 0.2−0.3 0.3−0.4 0.4−0.5 0.5−0.6 0.6−0.7 0.7−0.8 0.8−0.9 0.9−1.0
L6b 0.60 0.39 0.36 0.34 0.33 0.33 0.34 0.34 0.37 1.00
L6−IT−Car3 0.74 0.47 0.42 0.40 0.39 0.39 0.39 0.41 0.44 1.00
L6−IT 0.28 0.24 0.24 0.25 0.26 0.27 0.28 0.29 0.32 1.00
L6−CT 0.47 0.38 0.36 0.34 0.34 0.35 0.34 0.35 0.36 1.00
L56−NP 0.47 0.39 0.37 0.37 0.37 0.36 0.36 0.35 0.37 1.00
L5−IT 0.66 0.43 0.39 0.37 0.36 0.36 0.37 0.38 0.40 1.00
L5−ET 0.44 0.33 0.31 0.30 0.30 0.30 0.30 0.31 0.34 1.00
L4−IT 0.45 0.35 0.33 0.32 0.32 0.33 0.33 0.34 0.37 1.00
L23−IT 0.52 0.38 0.35 0.35 0.35 0.35 0.36 0.37 0.41 1.00
HIP−Misc2 0.48 0.38 0.36 0.37 0.35 0.37 0.36 0.37 0.40 1.00
HIP−Misc1 0.39 0.35 0.35 0.35 0.36 0.36 0.36 0.37 0.39 1.00
DG 0.36 0.37 0.36 0.38 0.39 0.41 0.41 0.44 0.50 1.00
CA3 0.49 0.37 0.33 0.33 0.33 0.33 0.33 0.34 0.39 1.00
CA1 0.41 0.31 0.29 0.30 0.30 0.30 0.31 0.32 0.34 1.00
Amy−Exc 0.45 0.35 0.34 0.33 0.33 0.34 0.34 0.35 0.38 1.00
Vip 0.46 0.45 0.44 0.43 0.43 0.43 0.42 0.43 0.45 1.00
THM−MB 0.49 0.39 0.38 0.37 0.36 0.35 0.36 0.36 0.37 1.00
THM−Inh 0.50 0.47 0.47 0.45 0.45 0.47 0.48 0.49 0.53 1.00
THM−Exc 0.59 0.47 0.43 0.42 0.41 0.40 0.41 0.41 0.44 1.00
SubCtx−Cplx 0.37 0.36 0.37 0.38 0.39 0.40 0.41 0.42 0.45 1.00
Sst 0.38 0.39 0.39 0.41 0.42 0.43 0.45 0.47 0.51 1.00
Sncg 0.48 0.47 0.47 0.47 0.47 0.47 0.46 0.47 0.49 1.00
Pvalb−ChC 0.40 0.39 0.38 0.40 0.38 0.39 0.41 0.40 0.43 1.00
Pvalb 1.15 1.04 0.86 0.81 0.84 0.90 0.86 0.97 0.98 1.00
PN 0.71 0.50 0.46 0.44 0.43 0.43 0.43 0.45 0.48 1.00
MSN−D2 0.45 0.39 0.37 0.38 0.37 0.37 0.37 0.38 0.40 1.00
MSN−D1 0.55 0.44 0.42 0.42 0.41 0.42 0.42 0.43 0.45 1.00
Lamp5−Lhx6 0.45 0.44 0.43 0.44 0.44 0.44 0.43 0.44 0.46 1.00
Lamp5 0.46 0.43 0.41 0.40 0.40 0.39 0.39 0.39 0.38 1.00
Foxp2 0.50 0.43 0.42 0.41 0.41 0.42 0.42 0.42 0.45 1.00
Chd7 0.43 0.40 0.39 0.38 0.38 0.37 0.36 0.36 0.37 1.00
CB 0.45 0.46 0.44 0.42 0.41 0.41 0.39 0.39 0.39 1.00
Percent 0.010 0.015 0.020 0.025
L6b 0.66 0.45 0.41 0.40 0.40 0.39 0.40 0.42 0.44 1.00
L6−IT−Car3 0.71 0.52 0.48 0.47 0.47 0.46 0.47 0.49 0.52 1.00
L6−IT 0.34 0.30 0.30 0.31 0.33 0.33 0.34 0.36 0.37 1.00
L6−CT 0.52 0.44 0.42 0.41 0.41 0.41 0.41 0.42 0.44 1.00
L56−NP 0.53 0.45 0.43 0.43 0.43 0.43 0.43 0.42 0.45 1.00
L5−IT 0.73 0.48 0.45 0.43 0.42 0.43 0.43 0.45 0.48 1.00
L5−ET 0.49 0.39 0.36 0.35 0.35 0.35 0.36 0.38 0.41 1.00
L4−IT 0.50 0.40 0.38 0.38 0.38 0.38 0.39 0.40 0.43 1.00
L23−IT 0.58 0.44 0.41 0.41 0.41 0.41 0.43 0.44 0.48 1.00
HIP−Misc2 0.54 0.43 0.43 0.42 0.42 0.42 0.43 0.44 0.47 1.00
HIP−Misc1 0.43 0.40 0.40 0.41 0.41 0.42 0.44 0.44 0.46 1.00
DG 0.39 0.40 0.41 0.42 0.43 0.45 0.47 0.50 0.55 1.00
CA3 0.56 0.43 0.39 0.39 0.39 0.39 0.40 0.41 0.45 1.00
CA1 0.46 0.36 0.35 0.36 0.35 0.36 0.37 0.38 0.41 1.00
Amy−Exc 0.50 0.41 0.40 0.40 0.39 0.40 0.41 0.42 0.46 1.00
Vip 0.54 0.52 0.51 0.51 0.51 0.51 0.50 0.51 0.53 1.00
THM−MB 0.57 0.46 0.44 0.42 0.42 0.42 0.43 0.43 0.45 1.00
THM−Inh 0.52 0.50 0.50 0.49 0.50 0.51 0.53 0.55 0.59 1.00
THM−Exc 0.69 0.53 0.49 0.47 0.47 0.46 0.46 0.47 0.49 1.00
SubCtx−Cplx 0.41 0.41 0.42 0.43 0.44 0.46 0.47 0.49 0.53 1.00
Sst 0.42 0.44 0.45 0.47 0.48 0.49 0.50 0.53 0.57 1.00
Sncg 0.53 0.54 0.54 0.54 0.55 0.55 0.54 0.54 0.56 1.00
Pvalb−ChC 0.46 0.44 0.44 0.45 0.47 0.46 0.48 0.48 0.51 1.00
Pvalb 1.28 1.19 1.14 0.87 0.82 0.89 1.01 0.94 0.98 1.00
PN 0.82 0.56 0.52 0.50 0.49 0.49 0.50 0.51 0.54 1.00
MSN−D2 0.51 0.46 0.44 0.45 0.44 0.44 0.44 0.44 0.47 1.00
MSN−D1 0.60 0.51 0.49 0.48 0.48 0.49 0.50 0.51 0.53 1.00
Lamp5−Lhx6 0.51 0.50 0.50 0.51 0.51 0.50 0.52 0.52 0.54 1.00
Lamp5 0.53 0.51 0.50 0.49 0.49 0.48 0.48 0.47 0.47 1.00
Foxp2 0.58 0.49 0.48 0.48 0.48 0.48 0.49 0.50 0.52 1.00
Chd7 0.49 0.46 0.45 0.45 0.45 0.44 0.44 0.43 0.44 1.00
CB 0.53 0.54 0.52 0.51 0.50 0.49 0.48 0.47 0.46 1.00
Percent 0.015 0.020 0.025 0.030
L6b 0.94 0.63 0.58 0.55 0.54 0.53 0.52 0.52 0.53 1.00
L6−IT−Car3 1.03 0.71 0.66 0.63 0.61 0.59 0.59 0.59 0.60 1.00
L6−IT 0.51 0.43 0.42 0.43 0.44 0.44 0.45 0.46 0.47 1.00
L6−CT 0.76 0.62 0.59 0.57 0.55 0.54 0.53 0.52 0.53 1.00
L56−NP 0.74 0.63 0.60 0.60 0.58 0.57 0.56 0.55 0.54 1.00
L5−IT 1.05 0.68 0.62 0.59 0.57 0.56 0.55 0.56 0.57 1.00
L5−ET 0.71 0.54 0.51 0.49 0.48 0.47 0.47 0.48 0.49 1.00
L4−IT 0.71 0.56 0.53 0.52 0.51 0.50 0.50 0.50 0.52 1.00
L23−IT 0.86 0.63 0.57 0.56 0.55 0.54 0.54 0.54 0.56 1.00
HIP−Misc2 0.77 0.61 0.59 0.58 0.56 0.55 0.55 0.55 0.56 1.00
HIP−Misc1 0.64 0.58 0.57 0.57 0.56 0.56 0.56 0.55 0.55 1.00
DG 0.56 0.57 0.57 0.56 0.57 0.58 0.59 0.60 0.64 1.00
CA3 0.78 0.58 0.54 0.51 0.51 0.51 0.51 0.51 0.54 1.00
CA1 0.70 0.53 0.50 0.49 0.48 0.48 0.48 0.48 0.51 1.00
Amy−Exc 0.73 0.56 0.54 0.52 0.52 0.52 0.52 0.53 0.55 1.00
Vip 0.76 0.72 0.71 0.69 0.68 0.66 0.64 0.62 0.62 1.00
THM−MB 0.78 0.65 0.62 0.59 0.57 0.57 0.56 0.55 0.54 1.00
THM−Inh 0.72 0.68 0.67 0.65 0.65 0.66 0.66 0.67 0.68 1.00
THM−Exc 0.86 0.69 0.64 0.63 0.61 0.60 0.60 0.59 0.60 1.00
SubCtx−Cplx 0.60 0.58 0.59 0.59 0.59 0.60 0.60 0.61 0.62 1.00
Sst 0.63 0.64 0.64 0.64 0.63 0.63 0.63 0.64 0.65 1.00
Sncg 0.78 0.76 0.74 0.73 0.73 0.71 0.69 0.67 0.65 1.00
Pvalb−ChC 0.69 0.66 0.63 0.63 0.62 0.62 0.61 0.59 0.60 1.00
Pvalb 1.23 1.02 0.95 0.86 0.85 1.01 0.95 1.05 0.97 1.00
PN 1.04 0.76 0.69 0.66 0.64 0.63 0.63 0.63 0.64 1.00
MSN−D2 0.73 0.64 0.62 0.62 0.61 0.59 0.57 0.56 0.56 1.00
MSN−D1 0.87 0.71 0.67 0.66 0.64 0.63 0.62 0.62 0.63 1.00
Lamp5−Lhx6 0.75 0.72 0.70 0.70 0.67 0.67 0.66 0.64 0.63 1.00
Lamp5 0.77 0.72 0.70 0.67 0.67 0.65 0.62 0.58 0.57 1.00
Foxp2 0.83 0.70 0.67 0.66 0.65 0.63 0.62 0.62 0.61 1.00
Chd7 0.71 0.67 0.64 0.62 0.61 0.59 0.57 0.56 0.53 1.00
CB 0.76 0.75 0.72 0.68 0.66 0.64 0.62 0.59 0.56 1.00
Percent 0.020 0.025 0.030 0.035 0.040

### L4−IT−Intratelencephalic projecting neurons, cortical layer 4

| | 0.0−0.1 | 0.1−0.2 | 0.2−0.3 | 0.3−0.4 | 0.4−0.5 | 0.5−0.6 | 0.6−0.7 | 0.7−0.8 | 0.8−0.9 | 0.9−1.0 |
|---|---|---|---|---|---|---|---|---|---|---|
| L6b | 0.60 | 0.41 | 0.38 | 0.36 | 0.36 | 0.36 | 0.36 | 0.38 | 0.41 | 1.00 |
| L6−IT−Car3 | 0.69 | 0.49 | 0.45 | 0.44 | 0.43 | 0.43 | 0.43 | 0.45 | 0.49 | 1.00 |
| L6−IT | 0.29 | 0.25 | 0.26 | 0.27 | 0.28 | 0.29 | 0.31 | 0.32 | 0.35 | 1.00 |
| L6−CT | 0.49 | 0.39 | 0.37 | 0.37 | 0.37 | 0.36 | 0.37 | 0.38 | 0.41 | 1.00 |
| L56−NP | 0.50 | 0.42 | 0.40 | 0.39 | 0.39 | 0.39 | 0.39 | 0.40 | 0.41 | 1.00 |
| L5−IT | 0.70 | 0.45 | 0.41 | 0.39 | 0.38 | 0.39 | 0.39 | 0.41 | 0.45 | 1.00 |
| L5−ET | 0.45 | 0.35 | 0.33 | 0.33 | 0.33 | 0.32 | 0.33 | 0.34 | 0.38 | 1.00 |
| L4−IT | 0.48 | 0.36 | 0.35 | 0.34 | 0.34 | 0.34 | 0.35 | 0.37 | 0.40 | 1.00 |
| L23−IT | 0.54 | 0.40 | 0.38 | 0.38 | 0.37 | 0.37 | 0.39 | 0.41 | 0.45 | 1.00 |
| HIP−Misc2 | 0.52 | 0.40 | 0.38 | 0.38 | 0.38 | 0.38 | 0.39 | 0.40 | 0.43 | 1.00 |
| HIP−Misc1 | 0.41 | 0.37 | 0.37 | 0.37 | 0.37 | 0.38 | 0.39 | 0.41 | 0.42 | 1.00 |
| DG | 0.38 | 0.38 | 0.38 | 0.39 | 0.40 | 0.42 | 0.44 | 0.46 | 0.53 | 1.00 |
| CA3 | 0.51 | 0.39 | 0.35 | 0.35 | 0.35 | 0.35 | 0.37 | 0.37 | 0.41 | 1.00 |
| CA1 | 0.43 | 0.33 | 0.32 | 0.32 | 0.32 | 0.32 | 0.33 | 0.36 | 0.38 | 1.00 |
| Amy−Exc | 0.47 | 0.36 | 0.35 | 0.35 | 0.35 | 0.36 | 0.37 | 0.39 | 0.42 | 1.00 |
| Vip | 0.50 | 0.48 | 0.47 | 0.47 | 0.46 | 0.46 | 0.46 | 0.47 | 0.49 | 1.00 |
| THM−MB | 0.51 | 0.42 | 0.41 | 0.39 | 0.39 | 0.38 | 0.39 | 0.40 | 0.41 | 1.00 |
| THM−Inh | 0.50 | 0.47 | 0.47 | 0.47 | 0.48 | 0.49 | 0.50 | 0.53 | 0.56 | 1.00 |
| THM−Exc | 0.63 | 0.50 | 0.45 | 0.44 | 0.42 | 0.43 | 0.43 | 0.44 | 0.47 | 1.00 |
| SubCtx−Cplx | 0.40 | 0.39 | 0.39 | 0.40 | 0.41 | 0.42 | 0.44 | 0.45 | 0.49 | 1.00 |
| Sst | 0.41 | 0.43 | 0.43 | 0.44 | 0.45 | 0.46 | 0.46 | 0.49 | 0.53 | 1.00 |
| Sncg | 0.50 | 0.50 | 0.49 | 0.50 | 0.49 | 0.50 | 0.50 | 0.50 | 0.52 | 1.00 |
| Pvalb−ChC | 0.43 | 0.41 | 0.41 | 0.42 | 0.42 | 0.42 | 0.43 | 0.43 | 0.47 | 1.00 |
| Pvalb | 1.17 | 1.17 | 1.00 | 0.79 | 0.69 | 0.88 | 0.98 | 0.95 | 0.90 | 1.00 |
| PN | 0.74 | 0.53 | 0.48 | 0.47 | 0.46 | 0.45 | 0.46 | 0.48 | 0.51 | 1.00 |
| MSN−D2 | 0.47 | 0.41 | 0.40 | 0.40 | 0.41 | 0.40 | 0.40 | 0.41 | 0.43 | 1.00 |
| MSN−D1 | 0.56 | 0.47 | 0.45 | 0.45 | 0.44 | 0.45 | 0.46 | 0.46 | 0.49 | 1.00 |
| Lamp5−Lhx6 | 0.48 | 0.48 | 0.47 | 0.47 | 0.47 | 0.47 | 0.47 | 0.48 | 0.50 | 1.00 |
| Lamp5 | 0.48 | 0.46 | 0.45 | 0.44 | 0.43 | 0.43 | 0.43 | 0.42 | 0.44 | 1.00 |
| Foxp2 | 0.53 | 0.45 | 0.44 | 0.44 | 0.44 | 0.44 | 0.44 | 0.45 | 0.48 | 1.00 |
| Chd7 | 0.45 | 0.42 | 0.41 | 0.41 | 0.40 | 0.40 | 0.40 | 0.40 | 0.41 | 1.00 |
| CB | 0.48 | 0.48 | 0.46 | 0.45 | 0.44 | 0.44 | 0.42 | 0.42 | 0.43 | 1.00 |

Percent 0.010 0.015 0.020 0.025 0.030

### L6B−Intratelencephalic projecting neurons, cortical layer 6B

| | 0.0−0.1 | 0.1−0.2 | 0.2−0.3 | 0.3−0.4 | 0.4−0.5 | 0.5−0.6 | 0.6−0.7 | 0.7−0.8 | 0.8−0.9 | 0.9−1.0 |
|---|---|---|---|---|---|---|---|---|---|---|
| L6b | 0.75 | 0.44 | 0.39 | 0.37 | 0.35 | 0.36 | 0.35 | 0.36 | 0.39 | 1.00 |
| L6−IT−Car3 | 0.74 | 0.52 | 0.48 | 0.47 | 0.45 | 0.44 | 0.45 | 0.46 | 0.47 | 1.00 |
| L6−IT | 0.35 | 0.30 | 0.29 | 0.29 | 0.30 | 0.31 | 0.31 | 0.33 | 0.34 | 1.00 |
| L6−CT | 0.53 | 0.43 | 0.40 | 0.40 | 0.38 | 0.38 | 0.38 | 0.38 | 0.39 | 1.00 |
| L56−NP | 0.54 | 0.44 | 0.41 | 0.41 | 0.40 | 0.39 | 0.39 | 0.38 | 0.40 | 1.00 |
| L5−IT | 0.78 | 0.49 | 0.45 | 0.42 | 0.41 | 0.40 | 0.40 | 0.41 | 0.44 | 1.00 |
| L5−ET | 0.50 | 0.38 | 0.35 | 0.34 | 0.33 | 0.33 | 0.33 | 0.34 | 0.36 | 1.00 |
| L4−IT | 0.49 | 0.40 | 0.38 | 0.37 | 0.36 | 0.36 | 0.37 | 0.37 | 0.39 | 1.00 |
| L23−IT | 0.61 | 0.45 | 0.41 | 0.40 | 0.40 | 0.40 | 0.41 | 0.41 | 0.44 | 1.00 |
| HIP−Misc2 | 0.55 | 0.43 | 0.42 | 0.41 | 0.40 | 0.40 | 0.41 | 0.41 | 0.42 | 1.00 |
| HIP−Misc1 | 0.45 | 0.40 | 0.40 | 0.40 | 0.40 | 0.40 | 0.41 | 0.41 | 0.41 | 1.00 |
| DG | 0.41 | 0.42 | 0.42 | 0.43 | 0.44 | 0.44 | 0.45 | 0.48 | 0.53 | 1.00 |
| CA3 | 0.54 | 0.41 | 0.38 | 0.37 | 0.37 | 0.37 | 0.38 | 0.37 | 0.40 | 1.00 |
| CA1 | 0.48 | 0.37 | 0.35 | 0.34 | 0.34 | 0.33 | 0.34 | 0.34 | 0.37 | 1.00 |
| Amy−Exc | 0.51 | 0.41 | 0.39 | 0.38 | 0.38 | 0.38 | 0.38 | 0.39 | 0.42 | 1.00 |
| Vip | 0.55 | 0.52 | 0.51 | 0.50 | 0.49 | 0.48 | 0.47 | 0.47 | 0.48 | 1.00 |
| THM−MB | 0.56 | 0.44 | 0.43 | 0.41 | 0.40 | 0.39 | 0.39 | 0.39 | 0.40 | 1.00 |
| THM−Inh | 0.55 | 0.52 | 0.51 | 0.51 | 0.51 | 0.52 | 0.53 | 0.55 | 0.56 | 1.00 |
| THM−Exc | 0.66 | 0.51 | 0.48 | 0.47 | 0.45 | 0.44 | 0.45 | 0.45 | 0.47 | 1.00 |
| SubCtx−Cplx | 0.43 | 0.42 | 0.43 | 0.43 | 0.43 | 0.45 | 0.46 | 0.46 | 0.49 | 1.00 |
| Sst | 0.47 | 0.48 | 0.48 | 0.48 | 0.49 | 0.50 | 0.50 | 0.52 | 0.54 | 1.00 |
| Sncg | 0.56 | 0.54 | 0.54 | 0.53 | 0.53 | 0.52 | 0.51 | 0.51 | 0.52 | 1.00 |
| Pvalb−ChC | 0.49 | 0.47 | 0.44 | 0.46 | 0.45 | 0.45 | 0.45 | 0.46 | 0.47 | 1.00 |
| Pvalb | 1.30 | 1.13 | 0.99 | 1.00 | 0.99 | 1.07 | 1.00 | 1.18 | 0.91 | 1.00 |
| PN | 0.82 | 0.58 | 0.53 | 0.50 | 0.49 | 0.49 | 0.49 | 0.49 | 0.52 | 1.00 |
| MSN−D2 | 0.51 | 0.44 | 0.44 | 0.43 | 0.43 | 0.42 | 0.41 | 0.41 | 0.43 | 1.00 |
| MSN−D1 | 0.61 | 0.49 | 0.49 | 0.48 | 0.46 | 0.46 | 0.46 | 0.47 | 0.49 | 1.00 |
| Lamp5−Lhx6 | 0.53 | 0.50 | 0.49 | 0.49 | 0.48 | 0.47 | 0.47 | 0.48 | 0.48 | 1.00 |
| Lamp5 | 0.53 | 0.48 | 0.45 | 0.45 | 0.44 | 0.44 | 0.43 | 0.42 | 0.42 | 1.00 |
| Foxp2 | 0.59 | 0.51 | 0.50 | 0.48 | 0.47 | 0.47 | 0.47 | 0.47 | 0.47 | 1.00 |
| Chd7 | 0.48 | 0.45 | 0.44 | 0.42 | 0.42 | 0.41 | 0.40 | 0.39 | 0.39 | 1.00 |
| CB | 0.52 | 0.50 | 0.49 | 0.47 | 0.46 | 0.45 | 0.44 | 0.43 | 0.42 | 1.00 |

Percent 0.010 0.015 0.020

### LAMP5_1−LAMP5+ GABAergic neurons

| | 0.0−0.1 | 0.1−0.2 | 0.2−0.3 | 0.3−0.4 | 0.4−0.5 | 0.5−0.6 | 0.6−0.7 | 0.7−0.8 | 0.8−0.9 | 0.9−1.0 |
|---|---|---|---|---|---|---|---|---|---|---|
| L6b | 0.52 | 0.35 | 0.31 | 0.30 | 0.29 | 0.29 | 0.29 | 0.30 | 0.33 | 1.00 |
| L6−IT−Car3 | 0.52 | 0.41 | 0.38 | 0.38 | 0.37 | 0.37 | 0.36 | 0.38 | 0.40 | 1.00 |
| L6−IT | 0.29 | 0.25 | 0.24 | 0.24 | 0.24 | 0.24 | 0.23 | 0.24 | 0.25 | 1.00 |
| L6−CT | 0.42 | 0.34 | 0.32 | 0.31 | 0.30 | 0.30 | 0.29 | 0.30 | 0.32 | 1.00 |
| L56−NP | 0.41 | 0.35 | 0.33 | 0.32 | 0.32 | 0.31 | 0.30 | 0.30 | 0.32 | 1.00 |
| L5−IT | 0.53 | 0.39 | 0.35 | 0.34 | 0.32 | 0.32 | 0.33 | 0.34 | 0.36 | 1.00 |
| L5−ET | 0.39 | 0.31 | 0.28 | 0.27 | 0.26 | 0.26 | 0.26 | 0.27 | 0.29 | 1.00 |
| L4−IT | 0.37 | 0.32 | 0.30 | 0.30 | 0.29 | 0.29 | 0.29 | 0.30 | 0.32 | 1.00 |
| L23−IT | 0.46 | 0.36 | 0.33 | 0.33 | 0.32 | 0.32 | 0.32 | 0.33 | 0.36 | 1.00 |
| HIP−Misc2 | 0.42 | 0.35 | 0.35 | 0.33 | 0.32 | 0.32 | 0.32 | 0.33 | 0.35 | 1.00 |
| HIP−Misc1 | 0.35 | 0.31 | 0.32 | 0.32 | 0.31 | 0.31 | 0.31 | 0.32 | 0.33 | 1.00 |
| DG | 0.35 | 0.35 | 0.35 | 0.35 | 0.36 | 0.36 | 0.38 | 0.39 | 0.45 | 1.00 |
| CA3 | 0.44 | 0.32 | 0.30 | 0.29 | 0.28 | 0.29 | 0.29 | 0.30 | 0.33 | 1.00 |
| CA1 | 0.39 | 0.29 | 0.27 | 0.27 | 0.26 | 0.26 | 0.26 | 0.27 | 0.29 | 1.00 |
| Amy−Exc | 0.41 | 0.33 | 0.31 | 0.30 | 0.30 | 0.30 | 0.30 | 0.31 | 0.33 | 1.00 |
| Vip | 0.48 | 0.43 | 0.41 | 0.39 | 0.38 | 0.37 | 0.36 | 0.36 | 0.38 | 1.00 |
| THM−MB | 0.45 | 0.35 | 0.33 | 0.31 | 0.31 | 0.30 | 0.29 | 0.30 | 0.31 | 1.00 |
| THM−Inh | 0.49 | 0.44 | 0.43 | 0.42 | 0.41 | 0.41 | 0.42 | 0.43 | 0.48 | 1.00 |
| THM−Exc | 0.54 | 0.41 | 0.38 | 0.37 | 0.35 | 0.34 | 0.35 | 0.35 | 0.38 | 1.00 |
| SubCtx−Cplx | 0.35 | 0.34 | 0.33 | 0.33 | 0.34 | 0.35 | 0.36 | 0.37 | 0.39 | 1.00 |
| Sst | 0.35 | 0.37 | 0.37 | 0.38 | 0.38 | 0.39 | 0.41 | 0.43 | 0.48 | 1.00 |
| Sncg | 0.46 | 0.42 | 0.43 | 0.42 | 0.41 | 0.40 | 0.40 | 0.41 | 0.43 | 1.00 |
| Pvalb−ChC | 0.40 | 0.36 | 0.34 | 0.35 | 0.34 | 0.34 | 0.34 | 0.35 | 0.38 | 1.00 |
| Pvalb | 1.23 | 1.21 | 1.09 | 0.92 | 0.88 | 0.89 | 0.96 | 0.97 | 0.97 | 1.00 |
| PN | 0.68 | 0.46 | 0.41 | 0.39 | 0.38 | 0.37 | 0.38 | 0.39 | 0.44 | 1.00 |
| MSN−D2 | 0.41 | 0.35 | 0.34 | 0.33 | 0.33 | 0.32 | 0.32 | 0.31 | 0.33 | 1.00 |
| MSN−D1 | 0.48 | 0.40 | 0.37 | 0.37 | 0.37 | 0.37 | 0.37 | 0.37 | 0.39 | 1.00 |
| Lamp5−Lhx6 | 0.43 | 0.38 | 0.38 | 0.39 | 0.38 | 0.37 | 0.37 | 0.38 | 0.40 | 1.00 |
| Lamp5 | 0.42 | 0.37 | 0.35 | 0.34 | 0.34 | 0.33 | 0.32 | 0.32 | 0.33 | 1.00 |
| Foxp2 | 0.48 | 0.39 | 0.39 | 0.37 | 0.36 | 0.36 | 0.36 | 0.36 | 0.39 | 1.00 |
| Chd7 | 0.40 | 0.35 | 0.34 | 0.32 | 0.32 | 0.31 | 0.30 | 0.30 | 0.31 | 1.00 |
| CB | 0.42 | 0.40 | 0.40 | 0.37 | 0.35 | 0.35 | 0.33 | 0.32 | 0.34 | 1.00 |

Percent 0.010 0.015 0.020

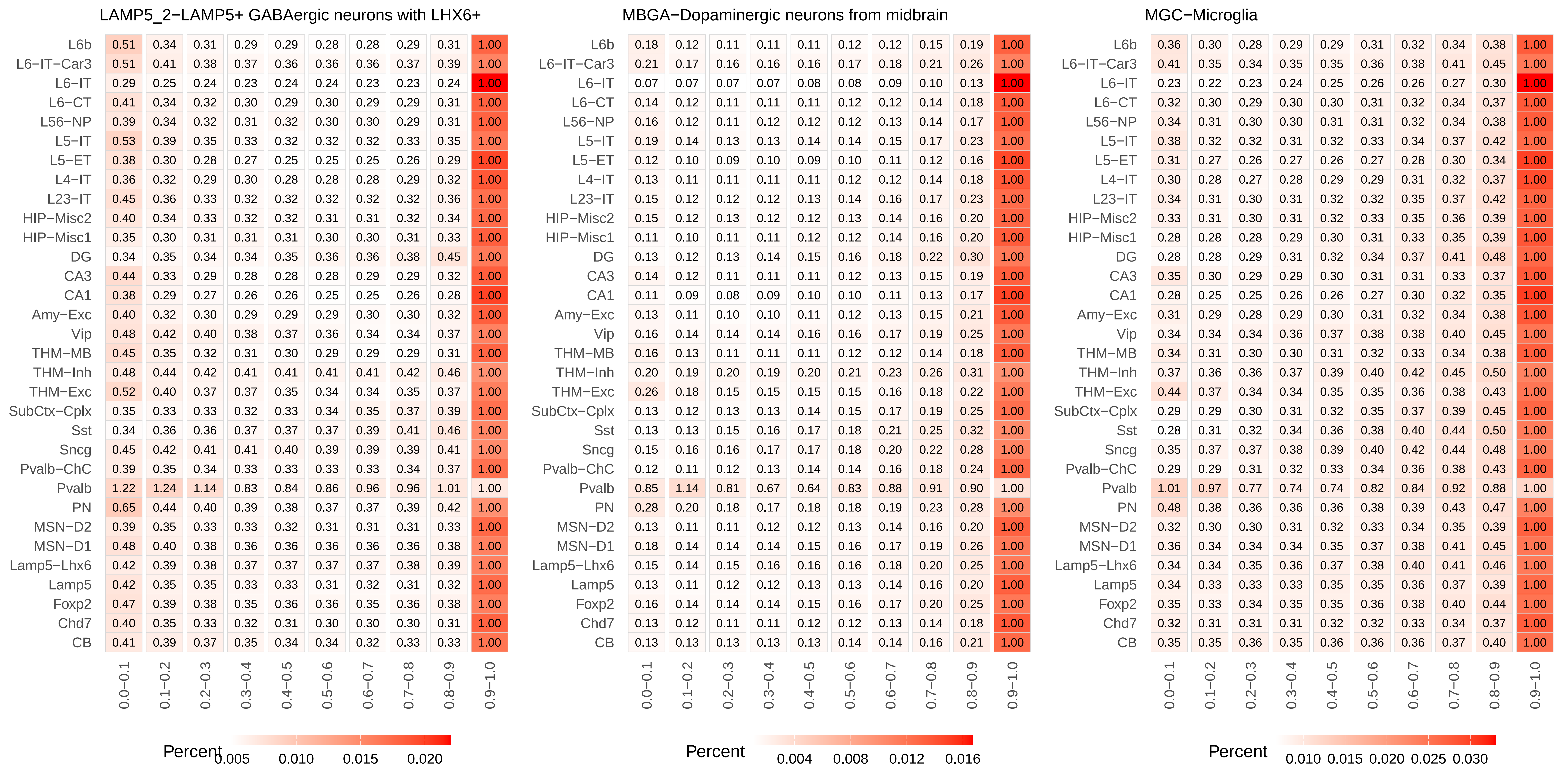
LAMP5_2−LAMP5+ GABAergic neurons with LHX6+
0.0−0.1 0.1−0.2 0.2−0.3 0.3−0.4 0.4−0.5 0.5−0.6 0.6−0.7 0.7−0.8 0.8−0.9 0.9−1.0
L6b 0.51 0.34 0.31 0.29 0.29 0.28 0.28 0.29 0.31 1.00
L6−IT−Car3 0.51 0.41 0.38 0.37 0.36 0.36 0.36 0.37 0.39 1.00
L6−IT 0.29 0.25 0.24 0.23 0.24 0.24 0.23 0.23 0.24 1.00
L6−CT 0.41 0.34 0.32 0.30 0.29 0.30 0.29 0.29 0.31 1.00
L56−NP 0.39 0.34 0.32 0.31 0.32 0.30 0.30 0.29 0.31 1.00
L5−IT 0.53 0.39 0.35 0.33 0.32 0.32 0.32 0.33 0.35 1.00
L5−ET 0.38 0.30 0.28 0.27 0.25 0.25 0.25 0.26 0.29 1.00
L4−IT 0.36 0.32 0.29 0.30 0.28 0.28 0.28 0.29 0.32 1.00
L23−IT 0.45 0.36 0.33 0.32 0.32 0.32 0.32 0.32 0.36 1.00
HIP−Misc2 0.40 0.34 0.33 0.32 0.32 0.31 0.31 0.32 0.34 1.00
HIP−Misc1 0.35 0.30 0.31 0.31 0.31 0.30 0.30 0.31 0.33 1.00
DG 0.34 0.35 0.34 0.34 0.35 0.36 0.36 0.38 0.45 1.00
CA3 0.44 0.33 0.29 0.28 0.28 0.28 0.29 0.29 0.32 1.00
CA1 0.38 0.29 0.27 0.26 0.26 0.25 0.25 0.26 0.28 1.00
Amy−Exc 0.40 0.32 0.30 0.29 0.29 0.29 0.30 0.30 0.32 1.00
Vip 0.48 0.42 0.40 0.38 0.37 0.36 0.34 0.34 0.37 1.00
THM−MB 0.45 0.35 0.32 0.31 0.30 0.29 0.29 0.29 0.31 1.00
THM−Inh 0.48 0.44 0.42 0.41 0.41 0.41 0.41 0.42 0.46 1.00
THM−Exc 0.52 0.40 0.37 0.37 0.35 0.34 0.34 0.35 0.37 1.00
SubCtx−Cplx 0.35 0.33 0.33 0.32 0.33 0.34 0.35 0.37 0.39 1.00
Sst 0.34 0.36 0.36 0.37 0.37 0.37 0.39 0.41 0.46 1.00
Sncg 0.45 0.42 0.41 0.41 0.40 0.39 0.39 0.39 0.41 1.00
Pvalb−ChC 0.39 0.35 0.34 0.33 0.33 0.33 0.33 0.34 0.37 1.00
Pvalb 1.22 1.24 1.14 0.83 0.84 0.86 0.96 0.96 1.01 1.00
PN 0.65 0.44 0.40 0.39 0.38 0.37 0.37 0.39 0.42 1.00
MSN−D2 0.39 0.35 0.33 0.33 0.32 0.31 0.31 0.31 0.33 1.00
MSN−D1 0.48 0.40 0.38 0.36 0.36 0.36 0.36 0.36 0.38 1.00
Lamp5−Lhx6 0.42 0.39 0.38 0.37 0.37 0.37 0.37 0.38 0.39 1.00
Lamp5 0.42 0.35 0.35 0.33 0.33 0.31 0.32 0.31 0.32 1.00
Foxp2 0.47 0.39 0.38 0.35 0.36 0.36 0.35 0.36 0.38 1.00
Chd7 0.40 0.35 0.33 0.32 0.31 0.30 0.30 0.30 0.31 1.00
CB 0.41 0.39 0.37 0.35 0.34 0.34 0.32 0.33 0.33 1.00
Percent
0.005 0.010 0.015 0.020
MBGA−Dopaminergic neurons from midbrain
0.0−0.1 0.1−0.2 0.2−0.3 0.3−0.4 0.4−0.5 0.5−0.6 0.6−0.7 0.7−0.8 0.8−0.9 0.9−1.0
L6b 0.18 0.12 0.11 0.11 0.11 0.12 0.12 0.15 0.19 1.00
L6−IT−Car3 0.21 0.17 0.16 0.16 0.16 0.17 0.18 0.21 0.26 1.00
L6−IT 0.07 0.07 0.07 0.07 0.08 0.08 0.09 0.10 0.13 1.00
L6−CT 0.14 0.12 0.11 0.11 0.11 0.12 0.12 0.14 0.18 1.00
L56−NP 0.16 0.12 0.11 0.12 0.12 0.12 0.13 0.14 0.17 1.00
L5−IT 0.19 0.14 0.13 0.13 0.14 0.14 0.15 0.17 0.23 1.00
L5−ET 0.12 0.10 0.09 0.10 0.09 0.10 0.11 0.12 0.16 1.00
L4−IT 0.13 0.11 0.11 0.11 0.11 0.12 0.12 0.14 0.18 1.00
L23−IT 0.15 0.12 0.12 0.12 0.13 0.14 0.16 0.17 0.23 1.00
HIP−Misc2 0.15 0.12 0.13 0.12 0.12 0.13 0.14 0.16 0.20 1.00
HIP−Misc1 0.11 0.10 0.11 0.11 0.12 0.12 0.14 0.16 0.20 1.00
DG 0.13 0.12 0.13 0.14 0.15 0.16 0.18 0.22 0.30 1.00
CA3 0.14 0.12 0.11 0.11 0.11 0.12 0.13 0.15 0.19 1.00
CA1 0.11 0.09 0.08 0.09 0.10 0.10 0.11 0.13 0.17 1.00
Amy−Exc 0.13 0.11 0.10 0.10 0.11 0.12 0.13 0.15 0.21 1.00
Vip 0.16 0.14 0.14 0.14 0.16 0.16 0.17 0.19 0.25 1.00
THM−MB 0.16 0.13 0.11 0.11 0.11 0.12 0.12 0.14 0.18 1.00
THM−Inh 0.20 0.19 0.20 0.19 0.20 0.21 0.23 0.26 0.31 1.00
THM−Exc 0.26 0.18 0.15 0.15 0.15 0.15 0.16 0.18 0.22 1.00
SubCtx−Cplx 0.13 0.12 0.13 0.13 0.14 0.15 0.17 0.19 0.25 1.00
Sst 0.13 0.13 0.15 0.16 0.17 0.18 0.21 0.25 0.32 1.00
Sncg 0.15 0.16 0.16 0.17 0.17 0.18 0.20 0.22 0.28 1.00
Pvalb−ChC 0.12 0.11 0.12 0.13 0.14 0.14 0.16 0.18 0.24 1.00
Pvalb 0.85 1.14 0.81 0.67 0.64 0.83 0.88 0.91 0.90 1.00
PN 0.28 0.20 0.18 0.17 0.18 0.18 0.19 0.23 0.28 1.00
MSN−D2 0.13 0.11 0.11 0.12 0.12 0.13 0.14 0.16 0.20 1.00
MSN−D1 0.18 0.14 0.14 0.14 0.15 0.16 0.17 0.19 0.26 1.00
Lamp5−Lhx6 0.15 0.14 0.15 0.16 0.16 0.16 0.18 0.20 0.25 1.00
Lamp5 0.13 0.11 0.12 0.12 0.13 0.13 0.14 0.16 0.20 1.00
Foxp2 0.16 0.14 0.14 0.14 0.15 0.16 0.17 0.20 0.25 1.00
Chd7 0.13 0.12 0.11 0.11 0.12 0.12 0.13 0.14 0.18 1.00
CB 0.13 0.13 0.13 0.13 0.13 0.14 0.14 0.16 0.21 1.00
Percent
0.004 0.008 0.012 0.016
MGC−Microglia
0.0−0.1 0.1−0.2 0.2−0.3 0.3−0.4 0.4−0.5 0.5−0.6 0.6−0.7 0.7−0.8 0.8−0.9 0.9−1.0
L6b 0.36 0.30 0.28 0.29 0.29 0.31 0.32 0.34 0.38 1.00
L6−IT−Car3 0.41 0.35 0.34 0.35 0.35 0.36 0.38 0.41 0.45 1.00
L6−IT 0.23 0.22 0.23 0.24 0.25 0.26 0.26 0.27 0.30 1.00
L6−CT 0.32 0.30 0.29 0.30 0.30 0.31 0.32 0.34 0.37 1.00
L56−NP 0.34 0.31 0.30 0.30 0.31 0.31 0.32 0.34 0.38 1.00
L5−IT 0.38 0.32 0.32 0.31 0.32 0.33 0.34 0.37 0.42 1.00
L5−ET 0.31 0.27 0.26 0.27 0.26 0.27 0.28 0.30 0.34 1.00
L4−IT 0.30 0.28 0.27 0.28 0.29 0.29 0.31 0.32 0.37 1.00
L23−IT 0.34 0.31 0.30 0.31 0.32 0.32 0.35 0.37 0.42 1.00
HIP−Misc2 0.33 0.31 0.30 0.31 0.32 0.33 0.35 0.36 0.39 1.00
HIP−Misc1 0.28 0.28 0.28 0.29 0.30 0.31 0.33 0.35 0.39 1.00
DG 0.28 0.28 0.29 0.31 0.32 0.34 0.37 0.41 0.48 1.00
CA3 0.35 0.30 0.29 0.29 0.30 0.31 0.31 0.33 0.37 1.00
CA1 0.28 0.25 0.25 0.26 0.26 0.27 0.30 0.32 0.35 1.00
Amy−Exc 0.31 0.29 0.28 0.29 0.30 0.31 0.32 0.34 0.38 1.00
Vip 0.34 0.34 0.34 0.36 0.37 0.38 0.38 0.40 0.45 1.00
THM−MB 0.34 0.31 0.30 0.30 0.31 0.32 0.33 0.34 0.38 1.00
THM−Inh 0.37 0.36 0.36 0.37 0.39 0.40 0.42 0.45 0.50 1.00
THM−Exc 0.44 0.37 0.34 0.34 0.35 0.35 0.36 0.38 0.43 1.00
SubCtx−Cplx 0.29 0.29 0.30 0.31 0.32 0.35 0.37 0.39 0.45 1.00
Sst 0.28 0.31 0.32 0.34 0.36 0.38 0.40 0.44 0.50 1.00
Sncg 0.35 0.37 0.37 0.38 0.39 0.40 0.42 0.44 0.48 1.00
Pvalb−ChC 0.29 0.29 0.31 0.32 0.33 0.34 0.36 0.38 0.43 1.00
Pvalb 1.01 0.97 0.77 0.74 0.74 0.82 0.84 0.92 0.88 1.00
PN 0.48 0.38 0.36 0.36 0.36 0.38 0.39 0.43 0.47 1.00
MSN−D2 0.32 0.30 0.30 0.31 0.32 0.33 0.34 0.35 0.39 1.00
MSN−D1 0.36 0.34 0.34 0.34 0.35 0.37 0.38 0.41 0.45 1.00
Lamp5−Lhx6 0.34 0.34 0.35 0.36 0.37 0.38 0.40 0.41 0.46 1.00
Lamp5 0.34 0.33 0.33 0.33 0.35 0.35 0.36 0.37 0.39 1.00
Foxp2 0.35 0.33 0.34 0.35 0.35 0.36 0.38 0.40 0.44 1.00
Chd7 0.32 0.31 0.31 0.31 0.32 0.32 0.33 0.34 0.37 1.00
CB 0.35 0.35 0.36 0.35 0.36 0.36 0.36 0.37 0.40 1.00
Percent
0.010 0.015 0.020 0.025 0.030

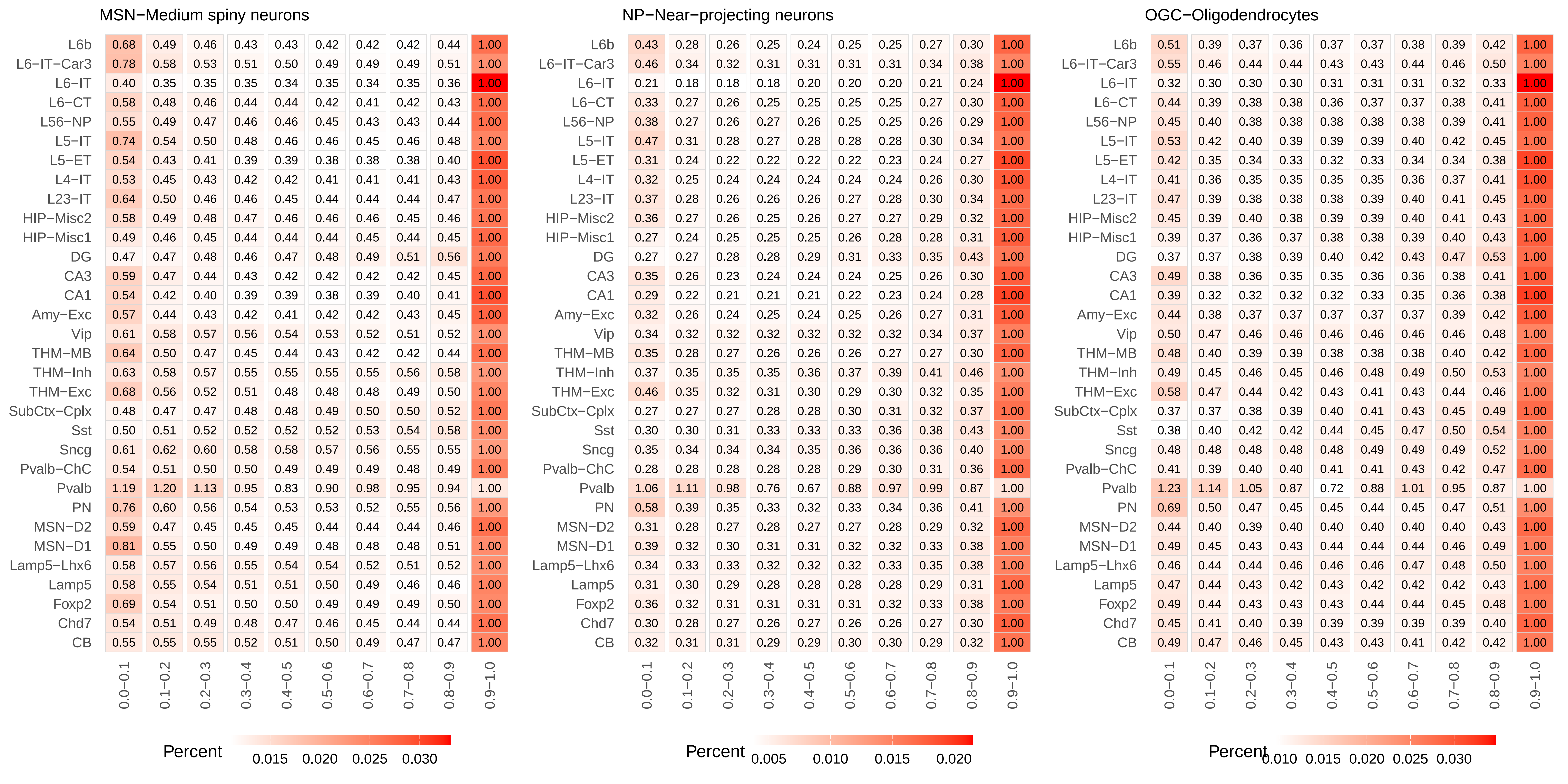
MSN−Medium spiny neurons
L6b 0.68 0.49 0.46 0.43 0.43 0.42 0.42 0.42 0.44 1.00
L6−IT−Car3 0.78 0.58 0.53 0.51 0.50 0.49 0.49 0.49 0.51 1.00
L6−IT 0.40 0.35 0.35 0.35 0.34 0.35 0.34 0.35 0.36 1.00
L6−CT 0.58 0.48 0.46 0.44 0.44 0.42 0.41 0.42 0.43 1.00
L56−NP 0.55 0.49 0.47 0.46 0.46 0.45 0.43 0.43 0.44 1.00
L5−IT 0.74 0.54 0.50 0.48 0.46 0.46 0.45 0.46 0.48 1.00
L5−ET 0.54 0.43 0.41 0.39 0.39 0.38 0.38 0.38 0.40 1.00
L4−IT 0.53 0.45 0.43 0.42 0.42 0.41 0.41 0.41 0.43 1.00
L23−IT 0.64 0.50 0.46 0.46 0.45 0.44 0.44 0.44 0.47 1.00
HIP−Misc2 0.58 0.49 0.48 0.47 0.46 0.46 0.46 0.45 0.46 1.00
HIP−Misc1 0.49 0.46 0.45 0.44 0.44 0.44 0.45 0.44 0.45 1.00
DG 0.47 0.47 0.48 0.46 0.47 0.48 0.49 0.51 0.56 1.00
CA3 0.59 0.47 0.44 0.43 0.42 0.42 0.42 0.42 0.45 1.00
CA1 0.54 0.42 0.40 0.39 0.39 0.38 0.39 0.40 0.41 1.00
Amy−Exc 0.57 0.44 0.43 0.42 0.41 0.42 0.42 0.43 0.45 1.00
Vip 0.61 0.58 0.57 0.56 0.54 0.53 0.52 0.51 0.52 1.00
THM−MB 0.64 0.50 0.47 0.45 0.44 0.43 0.42 0.42 0.44 1.00
THM−Inh 0.63 0.58 0.57 0.55 0.55 0.55 0.55 0.56 0.58 1.00
THM−Exc 0.68 0.56 0.52 0.51 0.48 0.48 0.48 0.49 0.50 1.00
SubCtx−Cplx 0.48 0.47 0.47 0.48 0.48 0.49 0.50 0.50 0.52 1.00
Sst 0.50 0.51 0.52 0.52 0.52 0.52 0.53 0.54 0.58 1.00
Sncg 0.61 0.62 0.60 0.58 0.58 0.57 0.56 0.55 0.55 1.00
Pvalb−ChC 0.54 0.51 0.50 0.50 0.49 0.49 0.49 0.48 0.49 1.00
Pvalb 1.19 1.20 1.13 0.95 0.83 0.90 0.98 0.95 0.94 1.00
PN 0.76 0.60 0.56 0.54 0.53 0.53 0.52 0.55 0.56 1.00
MSN−D2 0.59 0.47 0.45 0.45 0.45 0.44 0.44 0.44 0.46 1.00
MSN−D1 0.81 0.55 0.50 0.49 0.49 0.48 0.48 0.48 0.51 1.00
Lamp5−Lhx6 0.58 0.57 0.56 0.55 0.54 0.54 0.52 0.51 0.52 1.00
Lamp5 0.58 0.55 0.54 0.51 0.51 0.50 0.49 0.46 0.46 1.00
Foxp2 0.69 0.54 0.51 0.50 0.50 0.49 0.49 0.49 0.50 1.00
Chd7 0.54 0.51 0.49 0.48 0.47 0.46 0.45 0.44 0.44 1.00
CB 0.55 0.55 0.55 0.52 0.51 0.50 0.49 0.47 0.47 1.00
0.0−0.1 0.1−0.2 0.2−0.3 0.3−0.4 0.4−0.5 0.5−0.6 0.6−0.7 0.7−0.8 0.8−0.9 0.9−1.0
Percent 0.015 0.020 0.025 0.030
NP−Near−projecting neurons
L6b 0.43 0.28 0.26 0.25 0.24 0.25 0.25 0.27 0.30 1.00
L6−IT−Car3 0.46 0.34 0.32 0.31 0.31 0.31 0.31 0.34 0.38 1.00
L6−IT 0.21 0.18 0.18 0.18 0.20 0.20 0.20 0.21 0.24 1.00
L6−CT 0.33 0.27 0.26 0.25 0.25 0.25 0.25 0.27 0.30 1.00
L56−NP 0.38 0.27 0.26 0.27 0.26 0.25 0.25 0.26 0.29 1.00
L5−IT 0.47 0.31 0.28 0.27 0.28 0.28 0.28 0.30 0.34 1.00
L5−ET 0.31 0.24 0.22 0.22 0.22 0.22 0.23 0.24 0.27 1.00
L4−IT 0.32 0.25 0.24 0.24 0.24 0.24 0.24 0.26 0.30 1.00
L23−IT 0.37 0.28 0.26 0.26 0.26 0.27 0.28 0.30 0.34 1.00
HIP−Misc2 0.36 0.27 0.26 0.25 0.26 0.27 0.27 0.29 0.32 1.00
HIP−Misc1 0.27 0.24 0.25 0.25 0.25 0.26 0.28 0.28 0.31 1.00
DG 0.27 0.27 0.28 0.28 0.29 0.31 0.33 0.35 0.43 1.00
CA3 0.35 0.26 0.23 0.24 0.24 0.24 0.25 0.26 0.30 1.00
CA1 0.29 0.22 0.21 0.21 0.21 0.22 0.23 0.24 0.28 1.00
Amy−Exc 0.32 0.26 0.24 0.25 0.24 0.25 0.26 0.27 0.31 1.00
Vip 0.34 0.32 0.32 0.32 0.32 0.32 0.32 0.34 0.37 1.00
THM−MB 0.35 0.28 0.27 0.26 0.26 0.26 0.27 0.27 0.30 1.00
THM−Inh 0.37 0.35 0.35 0.35 0.36 0.37 0.39 0.41 0.46 1.00
THM−Exc 0.46 0.35 0.32 0.31 0.30 0.29 0.30 0.32 0.35 1.00
SubCtx−Cplx 0.27 0.27 0.27 0.28 0.28 0.30 0.31 0.32 0.37 1.00
Sst 0.30 0.30 0.31 0.33 0.33 0.33 0.36 0.38 0.43 1.00
Sncg 0.35 0.34 0.34 0.34 0.35 0.36 0.36 0.36 0.40 1.00
Pvalb−ChC 0.28 0.28 0.28 0.28 0.28 0.29 0.30 0.31 0.36 1.00
Pvalb 1.06 1.11 0.98 0.76 0.67 0.88 0.97 0.99 0.87 1.00
PN 0.58 0.39 0.35 0.33 0.32 0.33 0.34 0.36 0.41 1.00
MSN−D2 0.31 0.28 0.27 0.28 0.27 0.27 0.28 0.29 0.32 1.00
MSN−D1 0.39 0.32 0.30 0.31 0.31 0.32 0.32 0.33 0.38 1.00
Lamp5−Lhx6 0.34 0.33 0.33 0.32 0.32 0.32 0.33 0.35 0.38 1.00
Lamp5 0.31 0.30 0.29 0.28 0.28 0.28 0.28 0.29 0.31 1.00
Foxp2 0.36 0.32 0.31 0.31 0.31 0.31 0.32 0.33 0.38 1.00
Chd7 0.30 0.28 0.27 0.26 0.27 0.26 0.26 0.27 0.30 1.00
CB 0.32 0.31 0.31 0.29 0.29 0.30 0.30 0.29 0.32 1.00
0.0−0.1 0.1−0.2 0.2−0.3 0.3−0.4 0.4−0.5 0.5−0.6 0.6−0.7 0.7−0.8 0.8−0.9 0.9−1.0
Percent 0.005 0.010 0.015 0.020
OGC−Oligodendrocytes
L6b 0.51 0.39 0.37 0.36 0.37 0.37 0.38 0.39 0.42 1.00
L6−IT−Car3 0.55 0.46 0.44 0.44 0.43 0.43 0.44 0.46 0.50 1.00
L6−IT 0.32 0.30 0.30 0.30 0.31 0.31 0.31 0.32 0.33 1.00
L6−CT 0.44 0.39 0.38 0.38 0.36 0.37 0.37 0.38 0.41 1.00
L56−NP 0.45 0.40 0.38 0.38 0.38 0.38 0.38 0.39 0.41 1.00
L5−IT 0.53 0.42 0.40 0.39 0.39 0.39 0.40 0.42 0.45 1.00
L5−ET 0.42 0.35 0.34 0.33 0.32 0.33 0.34 0.34 0.38 1.00
L4−IT 0.41 0.36 0.35 0.35 0.35 0.35 0.36 0.37 0.41 1.00
L23−IT 0.47 0.39 0.38 0.38 0.38 0.39 0.40 0.41 0.45 1.00
HIP−Misc2 0.45 0.39 0.40 0.38 0.39 0.39 0.40 0.41 0.43 1.00
HIP−Misc1 0.39 0.37 0.36 0.37 0.38 0.38 0.39 0.40 0.43 1.00
DG 0.37 0.37 0.38 0.39 0.40 0.42 0.43 0.47 0.53 1.00
CA3 0.49 0.38 0.36 0.35 0.35 0.36 0.36 0.38 0.41 1.00
CA1 0.39 0.32 0.32 0.32 0.32 0.33 0.35 0.36 0.38 1.00
Amy−Exc 0.44 0.38 0.37 0.37 0.37 0.37 0.37 0.39 0.42 1.00
Vip 0.50 0.47 0.46 0.46 0.46 0.46 0.46 0.46 0.48 1.00
THM−MB 0.48 0.40 0.39 0.39 0.38 0.38 0.38 0.40 0.42 1.00
THM−Inh 0.49 0.45 0.46 0.45 0.46 0.48 0.49 0.50 0.53 1.00
THM−Exc 0.58 0.47 0.44 0.42 0.43 0.41 0.43 0.44 0.46 1.00
SubCtx−Cplx 0.37 0.37 0.38 0.39 0.40 0.41 0.43 0.45 0.49 1.00
Sst 0.38 0.40 0.42 0.42 0.44 0.45 0.47 0.50 0.54 1.00
Sncg 0.48 0.48 0.48 0.48 0.48 0.49 0.49 0.49 0.52 1.00
Pvalb−ChC 0.41 0.39 0.40 0.40 0.41 0.41 0.43 0.42 0.47 1.00
Pvalb 1.23 1.14 1.05 0.87 0.72 0.88 1.01 0.95 0.87 1.00
PN 0.69 0.50 0.47 0.45 0.45 0.44 0.45 0.47 0.51 1.00
MSN−D2 0.44 0.40 0.39 0.40 0.40 0.40 0.40 0.40 0.43 1.00
MSN−D1 0.49 0.45 0.43 0.43 0.44 0.44 0.44 0.46 0.49 1.00
Lamp5−Lhx6 0.46 0.44 0.44 0.46 0.46 0.46 0.47 0.48 0.50 1.00
Lamp5 0.47 0.44 0.43 0.42 0.43 0.42 0.42 0.42 0.43 1.00
Foxp2 0.49 0.44 0.43 0.43 0.43 0.44 0.44 0.45 0.48 1.00
Chd7 0.45 0.41 0.40 0.39 0.39 0.39 0.39 0.39 0.40 1.00
CB 0.49 0.47 0.46 0.45 0.43 0.43 0.41 0.42 0.42 1.00
0.0−0.1 0.1−0.2 0.2−0.3 0.3−0.4 0.4−0.5 0.5−0.6 0.6−0.7 0.7−0.8 0.8−0.9 0.9−1.0
Percent 0.010 0.015 0.020 0.025 0.030

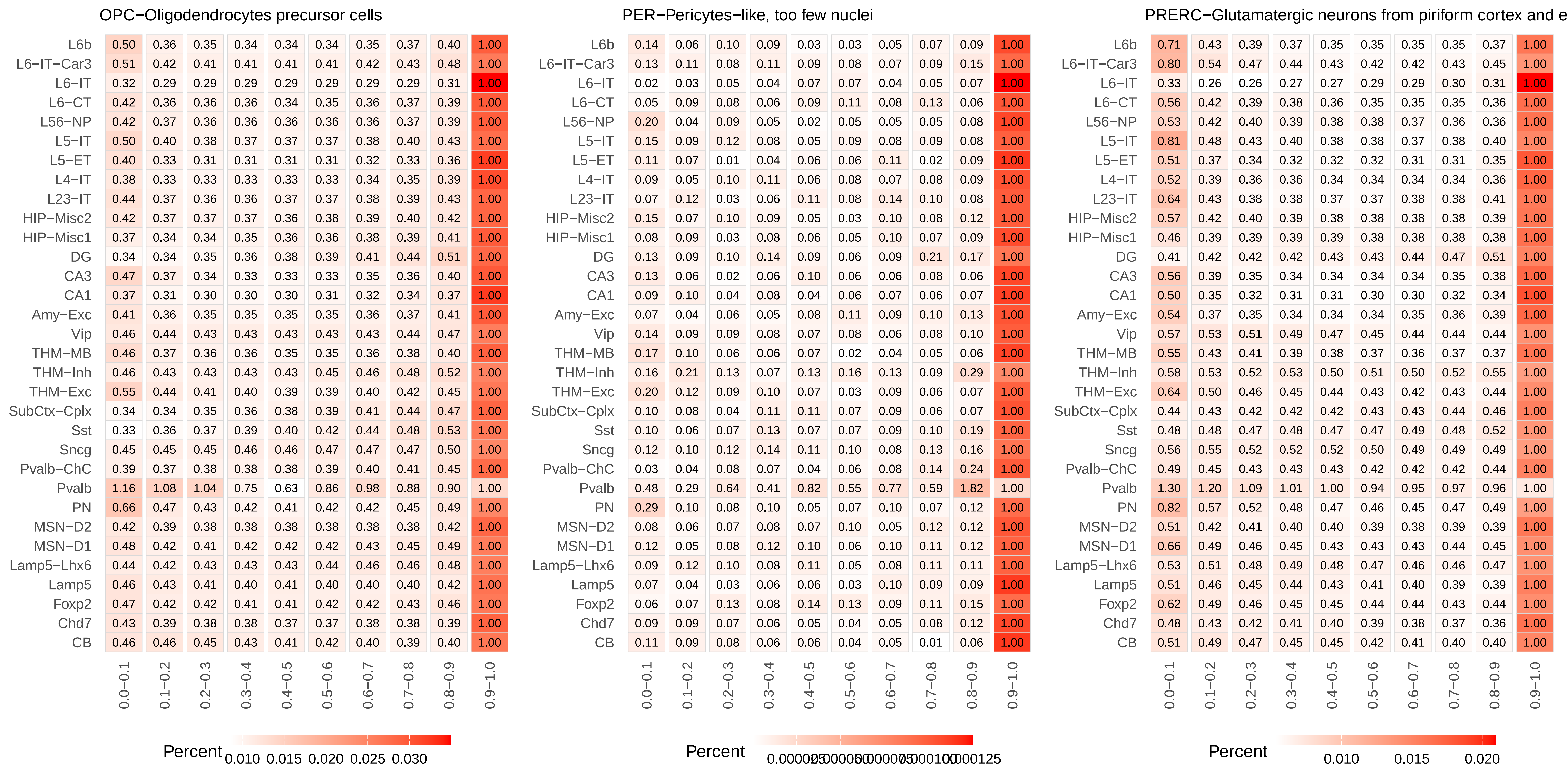
OPC−Oligodendrocytes precursor cells
L6b 0.50 0.36 0.35 0.34 0.34 0.34 0.35 0.37 0.40 1.00
L6−IT−Car3 0.51 0.42 0.41 0.41 0.41 0.41 0.42 0.43 0.48 1.00
L6−IT 0.32 0.29 0.29 0.29 0.29 0.29 0.29 0.29 0.31 1.00
L6−CT 0.42 0.36 0.36 0.36 0.34 0.35 0.36 0.37 0.39 1.00
L56−NP 0.42 0.37 0.36 0.36 0.36 0.36 0.36 0.37 0.39 1.00
L5−IT 0.50 0.40 0.38 0.37 0.37 0.37 0.38 0.40 0.43 1.00
L5−ET 0.40 0.33 0.31 0.31 0.31 0.31 0.32 0.33 0.36 1.00
L4−IT 0.38 0.33 0.33 0.33 0.33 0.33 0.34 0.35 0.39 1.00
L23−IT 0.44 0.37 0.36 0.36 0.37 0.37 0.38 0.39 0.43 1.00
HIP−Misc2 0.42 0.37 0.37 0.37 0.36 0.38 0.39 0.40 0.42 1.00
HIP−Misc1 0.37 0.34 0.34 0.35 0.36 0.36 0.38 0.39 0.41 1.00
DG 0.34 0.34 0.35 0.36 0.38 0.39 0.41 0.44 0.51 1.00
CA3 0.47 0.37 0.34 0.33 0.33 0.33 0.35 0.36 0.40 1.00
CA1 0.37 0.31 0.30 0.30 0.30 0.31 0.32 0.34 0.37 1.00
Amy−Exc 0.41 0.36 0.35 0.35 0.35 0.35 0.36 0.37 0.41 1.00
Vip 0.46 0.44 0.43 0.43 0.43 0.43 0.43 0.44 0.47 1.00
THM−MB 0.46 0.37 0.36 0.36 0.35 0.35 0.36 0.38 0.40 1.00
THM−Inh 0.46 0.43 0.43 0.43 0.43 0.45 0.46 0.48 0.52 1.00
THM−Exc 0.55 0.44 0.41 0.40 0.39 0.39 0.40 0.42 0.45 1.00
SubCtx−Cplx 0.34 0.34 0.35 0.36 0.38 0.39 0.41 0.44 0.47 1.00
Sst 0.33 0.36 0.37 0.39 0.40 0.42 0.44 0.48 0.53 1.00
Sncg 0.45 0.45 0.45 0.46 0.46 0.47 0.47 0.47 0.50 1.00
Pvalb−ChC 0.39 0.37 0.38 0.38 0.38 0.39 0.40 0.41 0.45 1.00
Pvalb 1.16 1.08 1.04 0.75 0.63 0.86 0.98 0.88 0.90 1.00
PN 0.66 0.47 0.43 0.42 0.41 0.42 0.42 0.45 0.49 1.00
MSN−D2 0.42 0.39 0.38 0.38 0.38 0.38 0.38 0.38 0.42 1.00
MSN−D1 0.48 0.42 0.41 0.42 0.42 0.42 0.43 0.45 0.49 1.00
Lamp5−Lhx6 0.44 0.42 0.43 0.43 0.43 0.44 0.46 0.46 0.48 1.00
Lamp5 0.46 0.43 0.41 0.40 0.41 0.40 0.40 0.40 0.42 1.00
Foxp2 0.47 0.42 0.42 0.41 0.41 0.42 0.42 0.43 0.46 1.00
Chd7 0.43 0.39 0.38 0.38 0.37 0.37 0.38 0.38 0.39 1.00
CB 0.46 0.46 0.45 0.43 0.41 0.42 0.40 0.39 0.40 1.00
0.0−0.1 0.1−0.2 0.2−0.3 0.3−0.4 0.4−0.5 0.5−0.6 0.6−0.7 0.7−0.8 0.8−0.9 0.9−1.0
Percent
0.010 0.015 0.020 0.025 0.030
PER−Pericytes−like, too few nuclei
L6b 0.14 0.06 0.10 0.09 0.03 0.03 0.05 0.07 0.09 1.00
L6−IT−Car3 0.13 0.11 0.08 0.11 0.09 0.08 0.07 0.09 0.15 1.00
L6−IT 0.02 0.03 0.05 0.04 0.07 0.07 0.04 0.05 0.07 1.00
L6−CT 0.05 0.09 0.08 0.06 0.09 0.11 0.08 0.13 0.06 1.00
L56−NP 0.20 0.04 0.09 0.05 0.02 0.05 0.05 0.05 0.08 1.00
L5−IT 0.15 0.09 0.12 0.08 0.05 0.09 0.08 0.09 0.08 1.00
L5−ET 0.11 0.07 0.01 0.04 0.06 0.06 0.11 0.02 0.09 1.00
L4−IT 0.09 0.05 0.10 0.11 0.06 0.08 0.07 0.08 0.09 1.00
L23−IT 0.07 0.12 0.03 0.06 0.11 0.08 0.14 0.10 0.08 1.00
HIP−Misc2 0.15 0.07 0.10 0.09 0.05 0.03 0.10 0.08 0.12 1.00
HIP−Misc1 0.08 0.09 0.03 0.08 0.06 0.05 0.10 0.07 0.09 1.00
DG 0.13 0.09 0.10 0.14 0.09 0.06 0.09 0.21 0.17 1.00
CA3 0.13 0.06 0.02 0.06 0.10 0.06 0.06 0.08 0.06 1.00
CA1 0.09 0.10 0.04 0.08 0.04 0.06 0.07 0.06 0.07 1.00
Amy−Exc 0.07 0.04 0.06 0.05 0.08 0.11 0.09 0.10 0.13 1.00
Vip 0.14 0.09 0.09 0.08 0.07 0.08 0.06 0.08 0.10 1.00
THM−MB 0.17 0.10 0.06 0.06 0.07 0.02 0.04 0.05 0.06 1.00
THM−Inh 0.16 0.21 0.13 0.07 0.13 0.16 0.13 0.09 0.29 1.00
THM−Exc 0.20 0.12 0.09 0.10 0.07 0.03 0.09 0.06 0.07 1.00
SubCtx−Cplx 0.10 0.08 0.04 0.11 0.11 0.07 0.09 0.06 0.07 1.00
Sst 0.10 0.06 0.07 0.13 0.07 0.07 0.09 0.10 0.19 1.00
Sncg 0.12 0.10 0.12 0.14 0.11 0.10 0.08 0.13 0.16 1.00
Pvalb−ChC 0.03 0.04 0.08 0.07 0.04 0.06 0.08 0.14 0.24 1.00
Pvalb 0.48 0.29 0.64 0.41 0.82 0.55 0.77 0.59 1.82 1.00
PN 0.29 0.10 0.08 0.10 0.05 0.07 0.10 0.07 0.12 1.00
MSN−D2 0.08 0.06 0.07 0.08 0.07 0.10 0.05 0.12 0.12 1.00
MSN−D1 0.12 0.05 0.08 0.12 0.10 0.06 0.10 0.11 0.12 1.00
Lamp5−Lhx6 0.09 0.12 0.10 0.08 0.11 0.05 0.08 0.11 0.11 1.00
Lamp5 0.07 0.04 0.03 0.06 0.06 0.03 0.10 0.09 0.09 1.00
Foxp2 0.06 0.07 0.13 0.08 0.14 0.13 0.09 0.11 0.15 1.00
Chd7 0.09 0.09 0.07 0.06 0.05 0.04 0.05 0.08 0.12 1.00
CB 0.11 0.09 0.08 0.06 0.06 0.04 0.05 0.01 0.06 1.00
0.0−0.1 0.1−0.2 0.2−0.3 0.3−0.4 0.4−0.5 0.5−0.6 0.6−0.7 0.7−0.8 0.8−0.9 0.9−1.0
Percent
0.000025 0.000050 0.000075 0.000100 0.000125
PRERC−Glutamatergic neurons from piriform cortex and e
L6b 0.71 0.43 0.39 0.37 0.35 0.35 0.35 0.35 0.37 1.00
L6−IT−Car3 0.80 0.54 0.47 0.44 0.43 0.42 0.42 0.43 0.45 1.00
L6−IT 0.33 0.26 0.26 0.27 0.27 0.29 0.29 0.30 0.31 1.00
L6−CT 0.56 0.42 0.39 0.38 0.36 0.35 0.35 0.35 0.36 1.00
L56−NP 0.53 0.42 0.40 0.39 0.38 0.38 0.37 0.36 0.36 1.00
L5−IT 0.81 0.48 0.43 0.40 0.38 0.38 0.37 0.38 0.40 1.00
L5−ET 0.51 0.37 0.34 0.32 0.32 0.32 0.31 0.31 0.35 1.00
L4−IT 0.52 0.39 0.36 0.36 0.34 0.34 0.34 0.34 0.36 1.00
L23−IT 0.64 0.43 0.38 0.38 0.37 0.37 0.38 0.38 0.41 1.00
HIP−Misc2 0.57 0.42 0.40 0.39 0.38 0.38 0.38 0.38 0.39 1.00
HIP−Misc1 0.46 0.39 0.39 0.39 0.39 0.38 0.38 0.38 0.38 1.00
DG 0.41 0.42 0.42 0.42 0.43 0.43 0.44 0.47 0.51 1.00
CA3 0.56 0.39 0.35 0.34 0.34 0.34 0.34 0.35 0.38 1.00
CA1 0.50 0.35 0.32 0.31 0.31 0.30 0.30 0.32 0.34 1.00
Amy−Exc 0.54 0.37 0.35 0.34 0.34 0.34 0.35 0.36 0.39 1.00
Vip 0.57 0.53 0.51 0.49 0.47 0.45 0.44 0.44 0.44 1.00
THM−MB 0.55 0.43 0.41 0.39 0.38 0.37 0.36 0.37 0.37 1.00
THM−Inh 0.58 0.53 0.52 0.53 0.50 0.51 0.50 0.52 0.55 1.00
THM−Exc 0.64 0.50 0.46 0.45 0.44 0.43 0.42 0.43 0.44 1.00
SubCtx−Cplx 0.44 0.43 0.42 0.42 0.42 0.43 0.43 0.44 0.46 1.00
Sst 0.48 0.48 0.47 0.48 0.47 0.47 0.49 0.48 0.52 1.00
Sncg 0.56 0.55 0.52 0.52 0.52 0.50 0.49 0.49 0.49 1.00
Pvalb−ChC 0.49 0.45 0.43 0.43 0.43 0.42 0.42 0.42 0.44 1.00
Pvalb 1.30 1.20 1.09 1.01 1.00 0.94 0.95 0.97 0.96 1.00
PN 0.82 0.57 0.52 0.48 0.47 0.46 0.45 0.47 0.49 1.00
MSN−D2 0.51 0.42 0.41 0.40 0.40 0.39 0.38 0.39 0.39 1.00
MSN−D1 0.66 0.49 0.46 0.45 0.43 0.43 0.43 0.44 0.45 1.00
Lamp5−Lhx6 0.53 0.51 0.48 0.49 0.48 0.47 0.46 0.46 0.47 1.00
Lamp5 0.51 0.46 0.45 0.44 0.43 0.41 0.40 0.39 0.39 1.00
Foxp2 0.62 0.49 0.46 0.45 0.45 0.44 0.44 0.43 0.44 1.00
Chd7 0.48 0.43 0.42 0.41 0.40 0.39 0.38 0.37 0.36 1.00
CB 0.51 0.49 0.47 0.45 0.45 0.42 0.41 0.40 0.40 1.00
0.0−0.1 0.1−0.2 0.2−0.3 0.3−0.4 0.4−0.5 0.5−0.6 0.6−0.7 0.7−0.8 0.8−0.9 0.9−1.0
Percent
0.010 0.015 0.020

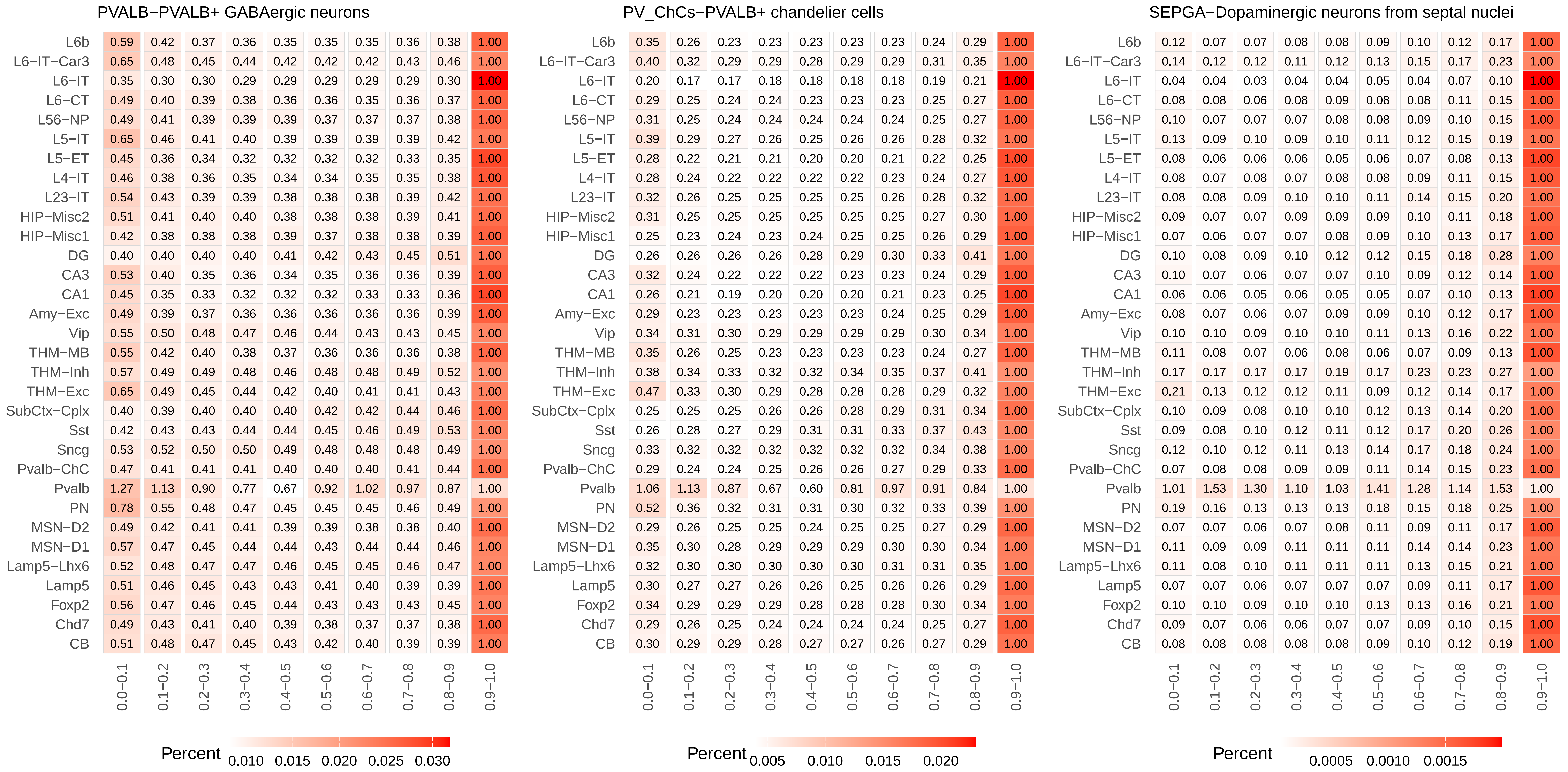
PVALB−PVALB+ GABAergic neurons
L6b 0.59 0.42 0.37 0.36 0.35 0.35 0.35 0.36 0.38 1.00
L6−IT−Car3 0.65 0.48 0.45 0.44 0.42 0.42 0.42 0.43 0.46 1.00
L6−IT 0.35 0.30 0.30 0.29 0.29 0.29 0.29 0.29 0.30 1.00
L6−CT 0.49 0.40 0.39 0.38 0.36 0.36 0.35 0.36 0.37 1.00
L56−NP 0.49 0.41 0.39 0.39 0.39 0.37 0.37 0.37 0.38 1.00
L5−IT 0.65 0.46 0.41 0.40 0.39 0.39 0.39 0.39 0.42 1.00
L5−ET 0.45 0.36 0.34 0.32 0.32 0.32 0.32 0.33 0.35 1.00
L4−IT 0.46 0.38 0.36 0.35 0.34 0.34 0.35 0.35 0.38 1.00
L23−IT 0.54 0.43 0.39 0.39 0.38 0.38 0.38 0.39 0.42 1.00
HIP−Misc2 0.51 0.41 0.40 0.40 0.38 0.38 0.38 0.39 0.41 1.00
HIP−Misc1 0.42 0.38 0.38 0.38 0.39 0.37 0.38 0.38 0.39 1.00
DG 0.40 0.40 0.40 0.40 0.41 0.42 0.43 0.45 0.51 1.00
CA3 0.53 0.40 0.35 0.36 0.34 0.35 0.36 0.36 0.39 1.00
CA1 0.45 0.35 0.33 0.32 0.32 0.32 0.33 0.33 0.36 1.00
Amy−Exc 0.49 0.39 0.37 0.36 0.36 0.36 0.36 0.36 0.39 1.00
Vip 0.55 0.50 0.48 0.47 0.46 0.44 0.43 0.43 0.45 1.00
THM−MB 0.55 0.42 0.40 0.38 0.37 0.36 0.36 0.36 0.38 1.00
THM−Inh 0.57 0.49 0.49 0.48 0.46 0.48 0.48 0.49 0.52 1.00
THM−Exc 0.65 0.49 0.45 0.44 0.42 0.40 0.41 0.41 0.43 1.00
SubCtx−Cplx 0.40 0.39 0.40 0.40 0.40 0.42 0.42 0.44 0.46 1.00
Sst 0.42 0.43 0.43 0.44 0.44 0.45 0.46 0.49 0.53 1.00
Sncg 0.53 0.52 0.50 0.50 0.49 0.48 0.48 0.48 0.49 1.00
Pvalb−ChC 0.47 0.41 0.41 0.41 0.40 0.40 0.40 0.41 0.44 1.00
Pvalb 1.27 1.13 0.90 0.77 0.67 0.92 1.02 0.97 0.87 1.00
PN 0.78 0.55 0.48 0.47 0.45 0.45 0.45 0.46 0.49 1.00
MSN−D2 0.49 0.42 0.41 0.41 0.39 0.39 0.38 0.38 0.40 1.00
MSN−D1 0.57 0.47 0.45 0.44 0.44 0.43 0.44 0.44 0.46 1.00
Lamp5−Lhx6 0.52 0.48 0.47 0.47 0.46 0.45 0.45 0.46 0.47 1.00
Lamp5 0.51 0.46 0.45 0.43 0.43 0.41 0.40 0.39 0.39 1.00
Foxp2 0.56 0.47 0.46 0.45 0.44 0.43 0.43 0.43 0.45 1.00
Chd7 0.49 0.43 0.41 0.40 0.39 0.38 0.37 0.37 0.38 1.00
CB 0.51 0.48 0.47 0.45 0.43 0.42 0.40 0.39 0.39 1.00
0.0−0.1 0.1−0.2 0.2−0.3 0.3−0.4 0.4−0.5 0.5−0.6 0.6−0.7 0.7−0.8 0.8−0.9 0.9−1.0
Percent 0.010 0.015 0.020 0.025 0.030
PV_ChCs−PVALB+ chandelier cells
L6b 0.35 0.26 0.23 0.23 0.23 0.23 0.23 0.24 0.29 1.00
L6−IT−Car3 0.40 0.32 0.29 0.29 0.28 0.29 0.29 0.31 0.35 1.00
L6−IT 0.20 0.17 0.17 0.18 0.18 0.18 0.18 0.19 0.21 1.00
L6−CT 0.29 0.25 0.24 0.24 0.23 0.23 0.23 0.25 0.27 1.00
L56−NP 0.31 0.25 0.24 0.24 0.24 0.24 0.24 0.25 0.27 1.00
L5−IT 0.39 0.29 0.27 0.26 0.25 0.26 0.26 0.28 0.32 1.00
L5−ET 0.28 0.22 0.21 0.21 0.20 0.20 0.21 0.22 0.25 1.00
L4−IT 0.28 0.24 0.22 0.22 0.22 0.22 0.23 0.24 0.27 1.00
L23−IT 0.32 0.26 0.25 0.25 0.25 0.25 0.26 0.28 0.32 1.00
HIP−Misc2 0.31 0.25 0.25 0.25 0.25 0.25 0.25 0.27 0.30 1.00
HIP−Misc1 0.25 0.23 0.24 0.23 0.24 0.25 0.25 0.26 0.29 1.00
DG 0.26 0.26 0.26 0.26 0.28 0.29 0.30 0.33 0.41 1.00
CA3 0.32 0.24 0.22 0.22 0.22 0.23 0.23 0.24 0.29 1.00
CA1 0.26 0.21 0.19 0.20 0.20 0.20 0.21 0.23 0.25 1.00
Amy−Exc 0.29 0.23 0.23 0.23 0.23 0.23 0.24 0.25 0.29 1.00
Vip 0.34 0.31 0.30 0.29 0.29 0.29 0.29 0.30 0.34 1.00
THM−MB 0.35 0.26 0.25 0.23 0.23 0.23 0.23 0.24 0.27 1.00
THM−Inh 0.38 0.34 0.33 0.32 0.32 0.34 0.35 0.37 0.41 1.00
THM−Exc 0.47 0.33 0.30 0.29 0.28 0.28 0.28 0.29 0.32 1.00
SubCtx−Cplx 0.25 0.25 0.25 0.26 0.26 0.28 0.29 0.31 0.34 1.00
Sst 0.26 0.28 0.27 0.29 0.31 0.31 0.33 0.37 0.43 1.00
Sncg 0.33 0.32 0.32 0.32 0.32 0.32 0.32 0.34 0.38 1.00
Pvalb−ChC 0.29 0.24 0.24 0.25 0.26 0.26 0.27 0.29 0.33 1.00
Pvalb 1.06 1.13 0.87 0.67 0.60 0.81 0.97 0.91 0.84 1.00
PN 0.52 0.36 0.32 0.31 0.31 0.30 0.32 0.33 0.39 1.00
MSN−D2 0.29 0.26 0.25 0.25 0.24 0.25 0.25 0.27 0.29 1.00
MSN−D1 0.35 0.30 0.28 0.29 0.29 0.29 0.30 0.30 0.34 1.00
Lamp5−Lhx6 0.32 0.30 0.30 0.30 0.30 0.30 0.31 0.31 0.35 1.00
Lamp5 0.30 0.27 0.27 0.26 0.26 0.25 0.26 0.26 0.29 1.00
Foxp2 0.34 0.29 0.29 0.29 0.28 0.28 0.28 0.30 0.34 1.00
Chd7 0.29 0.26 0.25 0.24 0.24 0.24 0.24 0.25 0.27 1.00
CB 0.30 0.29 0.29 0.28 0.27 0.27 0.26 0.27 0.29 1.00
0.0−0.1 0.1−0.2 0.2−0.3 0.3−0.4 0.4−0.5 0.5−0.6 0.6−0.7 0.7−0.8 0.8−0.9 0.9−1.0
Percent 0.005 0.010 0.015 0.020
SEPGA−Dopaminergic neurons from septal nuclei
L6b 0.12 0.07 0.07 0.08 0.08 0.09 0.10 0.12 0.17 1.00
L6−IT−Car3 0.14 0.12 0.12 0.11 0.12 0.13 0.15 0.17 0.23 1.00
L6−IT 0.04 0.04 0.03 0.04 0.04 0.05 0.04 0.07 0.10 1.00
L6−CT 0.08 0.08 0.06 0.08 0.09 0.08 0.08 0.11 0.15 1.00
L56−NP 0.10 0.07 0.07 0.07 0.08 0.08 0.09 0.10 0.15 1.00
L5−IT 0.13 0.09 0.10 0.09 0.10 0.11 0.12 0.15 0.19 1.00
L5−ET 0.08 0.06 0.06 0.06 0.05 0.06 0.07 0.08 0.13 1.00
L4−IT 0.08 0.07 0.08 0.07 0.08 0.08 0.09 0.11 0.15 1.00
L23−IT 0.08 0.08 0.09 0.10 0.10 0.11 0.14 0.15 0.20 1.00
HIP−Misc2 0.09 0.07 0.07 0.09 0.09 0.09 0.10 0.11 0.18 1.00
HIP−Misc1 0.07 0.06 0.07 0.07 0.08 0.09 0.10 0.13 0.17 1.00
DG 0.10 0.08 0.09 0.10 0.12 0.12 0.15 0.18 0.28 1.00
CA3 0.10 0.07 0.06 0.07 0.07 0.10 0.09 0.12 0.14 1.00
CA1 0.06 0.06 0.05 0.06 0.05 0.05 0.07 0.10 0.13 1.00
Amy−Exc 0.08 0.07 0.06 0.07 0.09 0.09 0.10 0.12 0.17 1.00
Vip 0.10 0.10 0.09 0.10 0.10 0.11 0.13 0.16 0.22 1.00
THM−MB 0.11 0.08 0.07 0.06 0.08 0.06 0.07 0.09 0.13 1.00
THM−Inh 0.17 0.17 0.17 0.17 0.19 0.17 0.23 0.23 0.27 1.00
THM−Exc 0.21 0.13 0.12 0.12 0.11 0.09 0.12 0.14 0.17 1.00
SubCtx−Cplx 0.10 0.09 0.08 0.10 0.10 0.12 0.13 0.14 0.20 1.00
Sst 0.09 0.08 0.10 0.12 0.11 0.12 0.17 0.20 0.26 1.00
Sncg 0.12 0.10 0.12 0.11 0.13 0.14 0.17 0.18 0.24 1.00
Pvalb−ChC 0.07 0.08 0.08 0.09 0.09 0.11 0.14 0.15 0.23 1.00
Pvalb 1.01 1.53 1.30 1.10 1.03 1.41 1.28 1.14 1.53 1.00
PN 0.19 0.16 0.13 0.13 0.13 0.18 0.15 0.18 0.25 1.00
MSN−D2 0.07 0.07 0.06 0.07 0.08 0.11 0.09 0.11 0.17 1.00
MSN−D1 0.11 0.09 0.09 0.11 0.11 0.11 0.14 0.14 0.23 1.00
Lamp5−Lhx6 0.11 0.08 0.10 0.11 0.11 0.11 0.13 0.15 0.21 1.00
Lamp5 0.07 0.07 0.06 0.07 0.07 0.07 0.09 0.11 0.17 1.00
Foxp2 0.10 0.10 0.09 0.10 0.10 0.13 0.13 0.16 0.21 1.00
Chd7 0.09 0.07 0.06 0.06 0.07 0.07 0.09 0.10 0.15 1.00
CB 0.08 0.08 0.08 0.08 0.08 0.09 0.10 0.12 0.19 1.00
0.0−0.1 0.1−0.2 0.2−0.3 0.3−0.4 0.4−0.5 0.5−0.6 0.6−0.7 0.7−0.8 0.8−0.9 0.9−1.0
Percent 0.0005 0.0010 0.0015

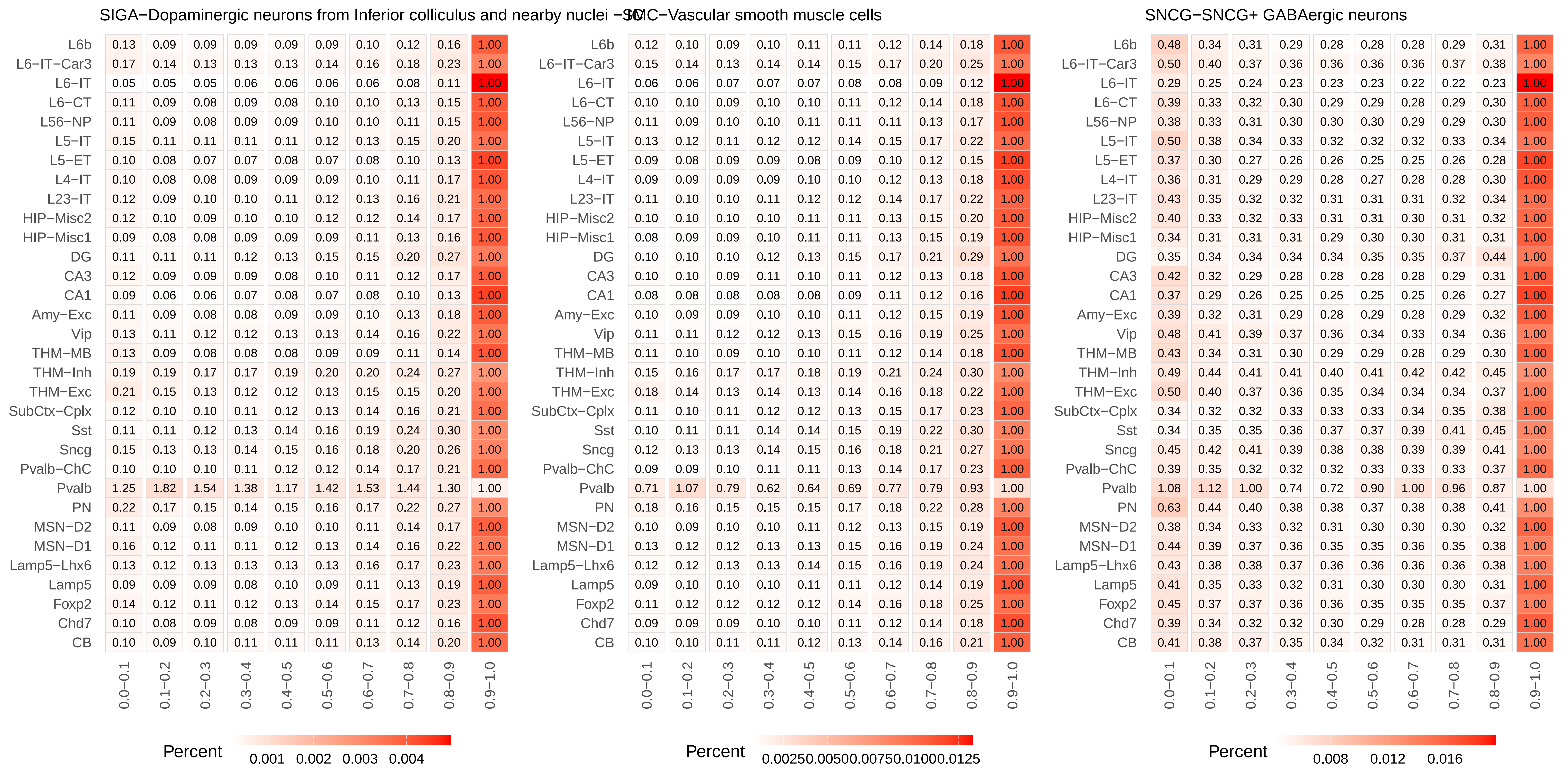
SIGA−Dopaminergic neurons from Inferior colliculus and nearby nuclei −SC
L6b 0.13 0.09 0.09 0.09 0.09 0.09 0.10 0.12 0.16 1.00
L6−IT−Car3 0.17 0.14 0.13 0.13 0.13 0.14 0.16 0.18 0.23 1.00
L6−IT 0.05 0.05 0.05 0.06 0.06 0.06 0.06 0.08 0.11 1.00
L6−CT 0.11 0.09 0.08 0.09 0.08 0.10 0.10 0.13 0.15 1.00
L56−NP 0.11 0.09 0.08 0.09 0.09 0.10 0.10 0.11 0.15 1.00
L5−IT 0.15 0.11 0.11 0.11 0.11 0.12 0.13 0.15 0.20 1.00
L5−ET 0.10 0.08 0.07 0.07 0.08 0.07 0.08 0.10 0.13 1.00
L4−IT 0.10 0.08 0.08 0.09 0.09 0.09 0.10 0.11 0.17 1.00
L23−IT 0.12 0.09 0.10 0.10 0.11 0.12 0.13 0.16 0.21 1.00
HIP−Misc2 0.12 0.10 0.09 0.10 0.10 0.12 0.12 0.14 0.17 1.00
HIP−Misc1 0.09 0.08 0.08 0.09 0.09 0.09 0.11 0.13 0.16 1.00
DG 0.11 0.11 0.11 0.12 0.13 0.15 0.15 0.20 0.27 1.00
CA3 0.12 0.09 0.09 0.09 0.08 0.10 0.11 0.12 0.17 1.00
CA1 0.09 0.06 0.06 0.07 0.08 0.07 0.08 0.10 0.13 1.00
Amy−Exc 0.11 0.09 0.08 0.08 0.09 0.09 0.10 0.13 0.18 1.00
Vip 0.13 0.11 0.12 0.12 0.13 0.13 0.14 0.16 0.22 1.00
THM−MB 0.13 0.09 0.08 0.08 0.08 0.09 0.09 0.11 0.14 1.00
THM−Inh 0.19 0.19 0.17 0.17 0.19 0.20 0.20 0.24 0.27 1.00
THM−Exc 0.21 0.15 0.13 0.12 0.12 0.13 0.15 0.15 0.20 1.00
SubCtx−Cplx 0.12 0.10 0.10 0.11 0.12 0.13 0.14 0.16 0.21 1.00
Sst 0.11 0.11 0.12 0.13 0.14 0.16 0.19 0.24 0.30 1.00
Sncg 0.15 0.13 0.13 0.14 0.15 0.16 0.18 0.20 0.26 1.00
Pvalb−ChC 0.10 0.10 0.10 0.11 0.12 0.12 0.14 0.17 0.21 1.00
Pvalb 1.25 1.82 1.54 1.38 1.17 1.42 1.53 1.44 1.30 1.00
PN 0.22 0.17 0.15 0.14 0.15 0.16 0.17 0.22 0.27 1.00
MSN−D2 0.11 0.09 0.08 0.09 0.10 0.10 0.11 0.14 0.17 1.00
MSN−D1 0.16 0.12 0.11 0.11 0.12 0.13 0.14 0.16 0.22 1.00
Lamp5−Lhx6 0.13 0.12 0.13 0.13 0.13 0.13 0.16 0.17 0.23 1.00
Lamp5 0.09 0.09 0.09 0.08 0.10 0.09 0.11 0.13 0.19 1.00
Foxp2 0.14 0.12 0.11 0.12 0.13 0.14 0.15 0.17 0.23 1.00
Chd7 0.10 0.08 0.09 0.08 0.09 0.09 0.11 0.12 0.16 1.00
CB 0.10 0.09 0.10 0.11 0.11 0.11 0.13 0.14 0.20 1.00
0.0−0.1 0.1−0.2 0.2−0.3 0.3−0.4 0.4−0.5 0.5−0.6 0.6−0.7 0.7−0.8 0.8−0.9 0.9−1.0
Percent
0.001 0.002 0.003 0.004
VSMC−Vascular smooth muscle cells
L6b 0.12 0.10 0.09 0.10 0.11 0.11 0.12 0.14 0.18 1.00
L6−IT−Car3 0.15 0.14 0.13 0.14 0.14 0.15 0.17 0.20 0.25 1.00
L6−IT 0.06 0.06 0.07 0.07 0.07 0.08 0.08 0.09 0.12 1.00
L6−CT 0.10 0.10 0.09 0.10 0.10 0.11 0.12 0.14 0.18 1.00
L56−NP 0.11 0.09 0.10 0.10 0.11 0.11 0.11 0.13 0.17 1.00
L5−IT 0.13 0.12 0.11 0.12 0.12 0.14 0.15 0.17 0.22 1.00
L5−ET 0.09 0.08 0.09 0.09 0.08 0.09 0.10 0.12 0.15 1.00
L4−IT 0.09 0.09 0.09 0.09 0.10 0.10 0.12 0.13 0.18 1.00
L23−IT 0.11 0.10 0.10 0.11 0.12 0.12 0.14 0.17 0.22 1.00
HIP−Misc2 0.10 0.10 0.10 0.10 0.11 0.11 0.13 0.15 0.20 1.00
HIP−Misc1 0.08 0.09 0.09 0.10 0.11 0.11 0.13 0.15 0.19 1.00
DG 0.10 0.10 0.10 0.12 0.13 0.15 0.17 0.21 0.29 1.00
CA3 0.10 0.10 0.09 0.11 0.10 0.11 0.12 0.13 0.18 1.00
CA1 0.08 0.08 0.08 0.08 0.08 0.09 0.11 0.12 0.16 1.00
Amy−Exc 0.10 0.09 0.09 0.10 0.10 0.11 0.12 0.15 0.19 1.00
Vip 0.11 0.11 0.12 0.12 0.13 0.15 0.16 0.19 0.25 1.00
THM−MB 0.11 0.10 0.09 0.10 0.10 0.11 0.12 0.14 0.18 1.00
THM−Inh 0.15 0.16 0.17 0.17 0.18 0.19 0.21 0.24 0.30 1.00
THM−Exc 0.18 0.14 0.13 0.14 0.13 0.14 0.16 0.18 0.22 1.00
SubCtx−Cplx 0.11 0.10 0.11 0.12 0.12 0.13 0.15 0.17 0.23 1.00
Sst 0.10 0.11 0.11 0.14 0.14 0.15 0.19 0.22 0.30 1.00
Sncg 0.12 0.13 0.13 0.14 0.15 0.16 0.18 0.21 0.27 1.00
Pvalb−ChC 0.09 0.09 0.10 0.11 0.11 0.13 0.14 0.17 0.23 1.00
Pvalb 0.71 1.07 0.79 0.62 0.64 0.69 0.77 0.79 0.93 1.00
PN 0.18 0.16 0.15 0.15 0.15 0.17 0.18 0.22 0.28 1.00
MSN−D2 0.10 0.09 0.10 0.10 0.11 0.12 0.13 0.15 0.19 1.00
MSN−D1 0.13 0.12 0.12 0.13 0.13 0.15 0.16 0.19 0.24 1.00
Lamp5−Lhx6 0.12 0.12 0.13 0.13 0.14 0.15 0.16 0.19 0.24 1.00
Lamp5 0.09 0.10 0.10 0.10 0.11 0.11 0.12 0.14 0.19 1.00
Foxp2 0.11 0.12 0.12 0.12 0.12 0.14 0.16 0.18 0.25 1.00
Chd7 0.09 0.09 0.09 0.10 0.10 0.11 0.12 0.14 0.18 1.00
CB 0.10 0.10 0.11 0.11 0.12 0.13 0.14 0.16 0.21 1.00
0.0−0.1 0.1−0.2 0.2−0.3 0.3−0.4 0.4−0.5 0.5−0.6 0.6−0.7 0.7−0.8 0.8−0.9 0.9−1.0
Percent
0.00250.00500.00750.01000.0125
SNCG−SNCG+ GABAergic neurons
L6b 0.48 0.34 0.31 0.29 0.28 0.28 0.28 0.29 0.31 1.00
L6−IT−Car3 0.50 0.40 0.37 0.36 0.36 0.36 0.36 0.37 0.38 1.00
L6−IT 0.29 0.25 0.24 0.23 0.23 0.23 0.22 0.22 0.23 1.00
L6−CT 0.39 0.33 0.32 0.30 0.29 0.29 0.28 0.29 0.30 1.00
L56−NP 0.38 0.33 0.31 0.30 0.30 0.30 0.29 0.29 0.30 1.00
L5−IT 0.50 0.38 0.34 0.33 0.32 0.32 0.32 0.33 0.34 1.00
L5−ET 0.37 0.30 0.27 0.26 0.26 0.25 0.25 0.26 0.28 1.00
L4−IT 0.36 0.31 0.29 0.29 0.28 0.27 0.28 0.28 0.30 1.00
L23−IT 0.43 0.35 0.32 0.32 0.31 0.31 0.31 0.32 0.34 1.00
HIP−Misc2 0.40 0.33 0.32 0.33 0.31 0.31 0.30 0.31 0.32 1.00
HIP−Misc1 0.34 0.31 0.31 0.31 0.29 0.30 0.30 0.31 0.31 1.00
DG 0.35 0.34 0.34 0.34 0.34 0.35 0.35 0.37 0.44 1.00
CA3 0.42 0.32 0.29 0.28 0.28 0.28 0.28 0.29 0.31 1.00
CA1 0.37 0.29 0.26 0.25 0.25 0.25 0.25 0.26 0.27 1.00
Amy−Exc 0.39 0.32 0.31 0.29 0.28 0.29 0.28 0.29 0.32 1.00
Vip 0.48 0.41 0.39 0.37 0.36 0.34 0.33 0.34 0.36 1.00
THM−MB 0.43 0.34 0.31 0.30 0.29 0.29 0.28 0.29 0.30 1.00
THM−Inh 0.49 0.44 0.41 0.41 0.40 0.41 0.42 0.42 0.45 1.00
THM−Exc 0.50 0.40 0.37 0.36 0.35 0.34 0.34 0.34 0.37 1.00
SubCtx−Cplx 0.34 0.32 0.32 0.33 0.33 0.33 0.34 0.35 0.38 1.00
Sst 0.34 0.35 0.35 0.36 0.37 0.37 0.39 0.41 0.45 1.00
Sncg 0.45 0.42 0.41 0.39 0.38 0.38 0.39 0.39 0.41 1.00
Pvalb−ChC 0.39 0.35 0.32 0.32 0.32 0.33 0.33 0.33 0.37 1.00
Pvalb 1.08 1.12 1.00 0.74 0.72 0.90 1.00 0.96 0.87 1.00
PN 0.63 0.44 0.40 0.38 0.38 0.37 0.38 0.38 0.41 1.00
MSN−D2 0.38 0.34 0.33 0.32 0.31 0.30 0.30 0.30 0.32 1.00
MSN−D1 0.44 0.39 0.37 0.36 0.35 0.35 0.36 0.35 0.38 1.00
Lamp5−Lhx6 0.43 0.38 0.38 0.37 0.36 0.36 0.36 0.36 0.38 1.00
Lamp5 0.41 0.35 0.33 0.32 0.31 0.30 0.30 0.30 0.31 1.00
Foxp2 0.45 0.37 0.37 0.36 0.36 0.35 0.35 0.35 0.37 1.00
Chd7 0.39 0.34 0.32 0.32 0.30 0.29 0.28 0.28 0.29 1.00
CB 0.41 0.38 0.37 0.35 0.34 0.32 0.31 0.31 0.31 1.00
0.0−0.1 0.1−0.2 0.2−0.3 0.3−0.4 0.4−0.5 0.5−0.6 0.6−0.7 0.7−0.8 0.8−0.9 0.9−1.0
Percent
0.008 0.012 0.016

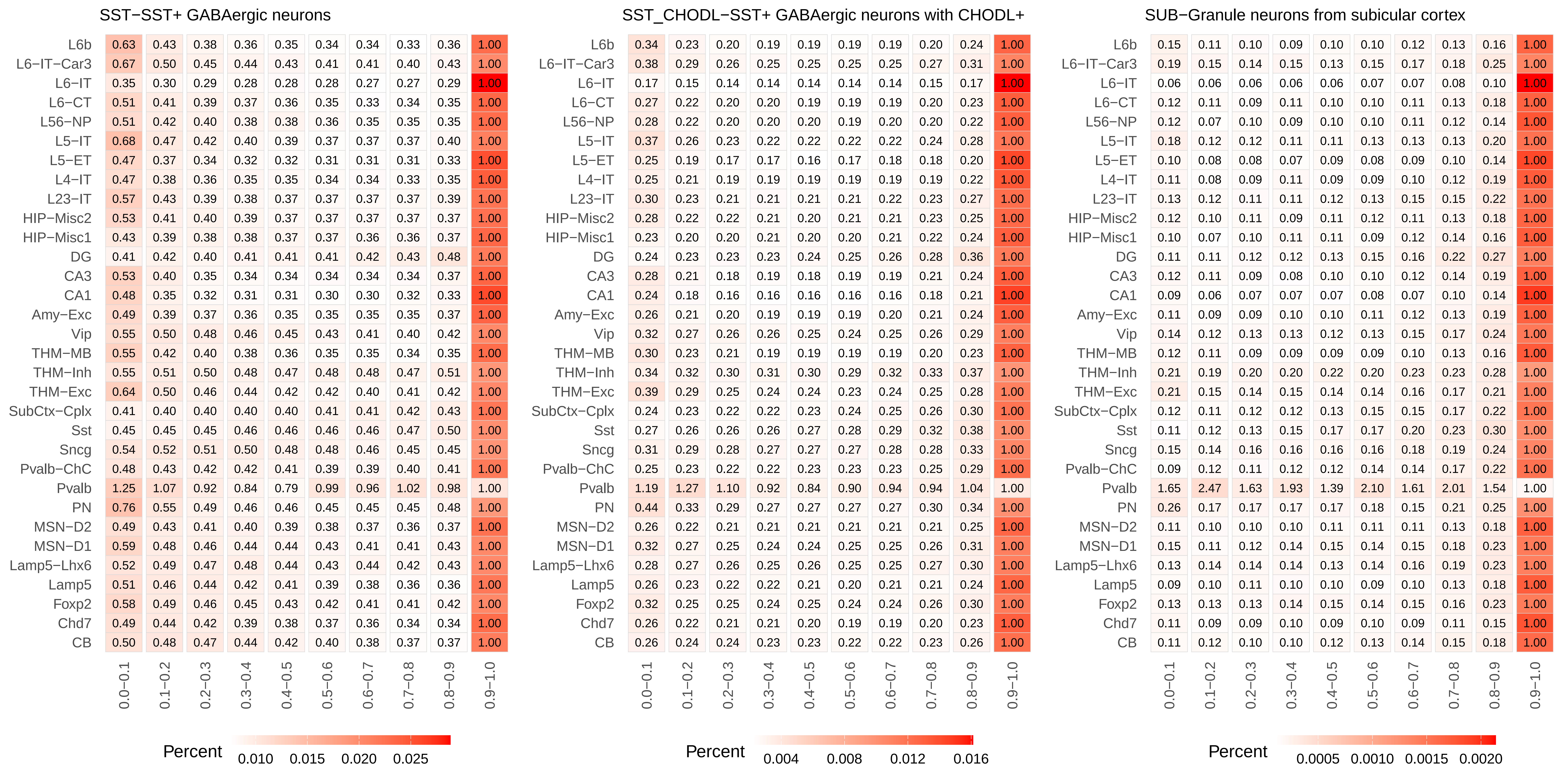
SST−SST+ GABAergic neurons
SST_CHODL−SST+ GABAergic neurons with CHODL+
SUB−Granule neurons from subicular cortex
L6b
L6−IT−Car3
L6−IT
L6−CT
L56−NP
L5−IT
L5−ET
L4−IT
L23−IT
HIP−Misc2
HIP−Misc1
DG
CA3
CA1
Amy−Exc
Vip
THM−MB
THM−Inh
THM−Exc
SubCtx−Cplx
Sst
Sncg
Pvalb−ChC
Pvalb
PN
MSN−D2
MSN−D1
Lamp5−Lhx6
Lamp5
Foxp2
Chd7
CB
0.0−0.1
0.1−0.2
0.2−0.3
0.3−0.4
0.4−0.5
0.5−0.6
0.6−0.7
0.7−0.8
0.8−0.9
0.9−1.0
Percent
0.010 0.015 0.020 0.025
Percent
0.004 0.008 0.012 0.016
Percent
0.0005 0.0010 0.0015 0.0020

### THMGA−GABAergic neurons from thalamus

| | 0.0−0.1 | 0.1−0.2 | 0.2−0.3 | 0.3−0.4 | 0.4−0.5 | 0.5−0.6 | 0.6−0.7 | 0.7−0.8 | 0.8−0.9 | 0.9−1.0 |
|---|---|---|---|---|---|---|---|---|---|---|
| L6b | 0.23 | 0.18 | 0.17 | 0.16 | 0.17 | 0.16 | 0.17 | 0.18 | 0.22 | 1.00 |
| L6−IT−Car3 | 0.27 | 0.24 | 0.22 | 0.23 | 0.22 | 0.22 | 0.23 | 0.26 | 0.31 | 1.00 |
| L6−IT | 0.15 | 0.14 | 0.13 | 0.12 | 0.13 | 0.13 | 0.12 | 0.12 | 0.14 | 1.00 |
| L6−CT | 0.20 | 0.18 | 0.17 | 0.17 | 0.16 | 0.17 | 0.17 | 0.18 | 0.21 | 1.00 |
| L56−NP | 0.20 | 0.17 | 0.16 | 0.17 | 0.17 | 0.17 | 0.17 | 0.19 | 0.21 | 1.00 |
| L5−IT | 0.25 | 0.21 | 0.20 | 0.20 | 0.19 | 0.19 | 0.20 | 0.22 | 0.26 | 1.00 |
| L5−ET | 0.17 | 0.15 | 0.15 | 0.15 | 0.14 | 0.14 | 0.15 | 0.16 | 0.18 | 1.00 |
| L4−IT | 0.18 | 0.16 | 0.16 | 0.16 | 0.16 | 0.16 | 0.18 | 0.18 | 0.21 | 1.00 |
| L23−IT | 0.22 | 0.19 | 0.18 | 0.18 | 0.20 | 0.19 | 0.21 | 0.22 | 0.27 | 1.00 |
| HIP−Misc2 | 0.21 | 0.17 | 0.17 | 0.18 | 0.18 | 0.18 | 0.20 | 0.21 | 0.23 | 1.00 |
| HIP−Misc1 | 0.18 | 0.16 | 0.17 | 0.16 | 0.17 | 0.17 | 0.19 | 0.20 | 0.23 | 1.00 |
| DG | 0.20 | 0.20 | 0.20 | 0.20 | 0.21 | 0.23 | 0.23 | 0.28 | 0.35 | 1.00 |
| CA3 | 0.20 | 0.17 | 0.16 | 0.16 | 0.16 | 0.17 | 0.18 | 0.19 | 0.22 | 1.00 |
| CA1 | 0.17 | 0.14 | 0.14 | 0.15 | 0.14 | 0.15 | 0.16 | 0.17 | 0.19 | 1.00 |
| Amy−Exc | 0.20 | 0.17 | 0.17 | 0.17 | 0.17 | 0.18 | 0.17 | 0.19 | 0.22 | 1.00 |
| Vip | 0.23 | 0.21 | 0.21 | 0.21 | 0.21 | 0.22 | 0.22 | 0.23 | 0.28 | 1.00 |
| THM−MB | 0.34 | 0.17 | 0.15 | 0.13 | 0.14 | 0.13 | 0.14 | 0.16 | 0.19 | 1.00 |
| THM−Inh | 0.30 | 0.28 | 0.28 | 0.27 | 0.26 | 0.28 | 0.30 | 0.32 | 0.36 | 1.00 |
| THM−Exc | 0.35 | 0.23 | 0.22 | 0.21 | 0.20 | 0.19 | 0.22 | 0.22 | 0.26 | 1.00 |
| SubCtx−Cplx | 0.19 | 0.18 | 0.17 | 0.19 | 0.20 | 0.21 | 0.22 | 0.24 | 0.28 | 1.00 |
| Sst | 0.21 | 0.22 | 0.23 | 0.23 | 0.24 | 0.25 | 0.26 | 0.29 | 0.36 | 1.00 |
| Sncg | 0.24 | 0.23 | 0.25 | 0.24 | 0.24 | 0.25 | 0.26 | 0.27 | 0.31 | 1.00 |
| Pvalb−ChC | 0.19 | 0.18 | 0.19 | 0.19 | 0.20 | 0.20 | 0.21 | 0.23 | 0.28 | 1.00 |
| Pvalb | 1.25 | 1.48 | 1.29 | 1.07 | 1.20 | 1.13 | 1.16 | 1.07 | 0.96 | 1.00 |
| PN | 0.34 | 0.26 | 0.24 | 0.24 | 0.24 | 0.24 | 0.25 | 0.28 | 0.35 | 1.00 |
| MSN−D2 | 0.20 | 0.18 | 0.17 | 0.17 | 0.18 | 0.18 | 0.18 | 0.19 | 0.22 | 1.00 |
| MSN−D1 | 0.26 | 0.21 | 0.20 | 0.22 | 0.21 | 0.21 | 0.22 | 0.24 | 0.29 | 1.00 |
| Lamp5−Lhx6 | 0.22 | 0.21 | 0.22 | 0.22 | 0.22 | 0.23 | 0.22 | 0.25 | 0.28 | 1.00 |
| Lamp5 | 0.19 | 0.18 | 0.18 | 0.18 | 0.18 | 0.18 | 0.18 | 0.19 | 0.21 | 1.00 |
| Foxp2 | 0.24 | 0.22 | 0.21 | 0.21 | 0.21 | 0.22 | 0.22 | 0.24 | 0.30 | 1.00 |
| Chd7 | 0.19 | 0.18 | 0.17 | 0.17 | 0.17 | 0.16 | 0.17 | 0.18 | 0.21 | 1.00 |
| CB | 0.20 | 0.20 | 0.20 | 0.20 | 0.19 | 0.20 | 0.20 | 0.20 | 0.24 | 1.00 |

Percent 0.0025 0.0050 0.0075

### VIP−VIP+ GABAergic neurons

| | 0.0−0.1 | 0.1−0.2 | 0.2−0.3 | 0.3−0.4 | 0.4−0.5 | 0.5−0.6 | 0.6−0.7 | 0.7−0.8 | 0.8−0.9 | 0.9−1.0 |
|---|---|---|---|---|---|---|---|---|---|---|
| L6b | 0.59 | 0.44 | 0.40 | 0.39 | 0.38 | 0.38 | 0.38 | 0.38 | 0.39 | 1.00 |
| L6−IT−Car3 | 0.63 | 0.51 | 0.47 | 0.47 | 0.46 | 0.45 | 0.44 | 0.46 | 0.48 | 1.00 |
| L6−IT | 0.38 | 0.34 | 0.33 | 0.32 | 0.32 | 0.31 | 0.31 | 0.31 | 0.31 | 1.00 |
| L6−CT | 0.51 | 0.43 | 0.41 | 0.41 | 0.39 | 0.38 | 0.37 | 0.38 | 0.38 | 1.00 |
| L56−NP | 0.49 | 0.44 | 0.42 | 0.41 | 0.40 | 0.40 | 0.39 | 0.37 | 0.39 | 1.00 |
| L5−IT | 0.62 | 0.47 | 0.44 | 0.43 | 0.41 | 0.41 | 0.40 | 0.41 | 0.44 | 1.00 |
| L5−ET | 0.47 | 0.38 | 0.36 | 0.34 | 0.33 | 0.34 | 0.33 | 0.34 | 0.36 | 1.00 |
| L4−IT | 0.47 | 0.40 | 0.39 | 0.37 | 0.36 | 0.36 | 0.36 | 0.36 | 0.39 | 1.00 |
| L23−IT | 0.54 | 0.44 | 0.42 | 0.41 | 0.40 | 0.40 | 0.40 | 0.40 | 0.43 | 1.00 |
| HIP−Misc2 | 0.51 | 0.45 | 0.43 | 0.41 | 0.40 | 0.40 | 0.40 | 0.40 | 0.40 | 1.00 |
| HIP−Misc1 | 0.43 | 0.41 | 0.41 | 0.39 | 0.39 | 0.39 | 0.39 | 0.39 | 0.41 | 1.00 |
| DG | 0.42 | 0.43 | 0.42 | 0.41 | 0.43 | 0.44 | 0.45 | 0.46 | 0.51 | 1.00 |
| CA3 | 0.56 | 0.41 | 0.38 | 0.37 | 0.37 | 0.36 | 0.36 | 0.37 | 0.39 | 1.00 |
| CA1 | 0.47 | 0.37 | 0.34 | 0.34 | 0.33 | 0.33 | 0.33 | 0.34 | 0.36 | 1.00 |
| Amy−Exc | 0.50 | 0.41 | 0.40 | 0.39 | 0.38 | 0.38 | 0.38 | 0.38 | 0.40 | 1.00 |
| Vip | 0.61 | 0.53 | 0.51 | 0.48 | 0.47 | 0.45 | 0.44 | 0.44 | 0.45 | 1.00 |
| THM−MB | 0.54 | 0.45 | 0.42 | 0.40 | 0.39 | 0.39 | 0.38 | 0.38 | 0.38 | 1.00 |
| THM−Inh | 0.58 | 0.53 | 0.52 | 0.50 | 0.50 | 0.50 | 0.50 | 0.52 | 0.54 | 1.00 |
| THM−Exc | 0.64 | 0.51 | 0.47 | 0.46 | 0.44 | 0.43 | 0.42 | 0.43 | 0.45 | 1.00 |
| SubCtx−Cplx | 0.42 | 0.41 | 0.41 | 0.41 | 0.42 | 0.43 | 0.44 | 0.45 | 0.47 | 1.00 |
| Sst | 0.38 | 0.41 | 0.42 | 0.43 | 0.44 | 0.45 | 0.47 | 0.49 | 0.54 | 1.00 |
| Sncg | 0.56 | 0.54 | 0.52 | 0.51 | 0.51 | 0.50 | 0.49 | 0.48 | 0.50 | 1.00 |
| Pvalb−ChC | 0.48 | 0.43 | 0.42 | 0.42 | 0.43 | 0.43 | 0.42 | 0.42 | 0.45 | 1.00 |
| Pvalb | 1.17 | 1.04 | 0.85 | 0.81 | 0.77 | 0.96 | 0.94 | 0.98 | 0.98 | 1.00 |
| PN | 0.83 | 0.57 | 0.50 | 0.48 | 0.47 | 0.46 | 0.45 | 0.47 | 0.49 | 1.00 |
| MSN−D2 | 0.50 | 0.46 | 0.43 | 0.43 | 0.42 | 0.41 | 0.40 | 0.39 | 0.41 | 1.00 |
| MSN−D1 | 0.57 | 0.50 | 0.48 | 0.47 | 0.46 | 0.46 | 0.46 | 0.46 | 0.48 | 1.00 |
| Lamp5−Lhx6 | 0.54 | 0.50 | 0.49 | 0.48 | 0.48 | 0.47 | 0.47 | 0.47 | 0.48 | 1.00 |
| Lamp5 | 0.54 | 0.48 | 0.46 | 0.45 | 0.44 | 0.42 | 0.41 | 0.41 | 0.40 | 1.00 |
| Foxp2 | 0.56 | 0.48 | 0.48 | 0.46 | 0.45 | 0.45 | 0.44 | 0.45 | 0.46 | 1.00 |
| Chd7 | 0.51 | 0.45 | 0.43 | 0.42 | 0.40 | 0.39 | 0.39 | 0.38 | 0.39 | 1.00 |
| CB | 0.55 | 0.52 | 0.50 | 0.46 | 0.45 | 0.43 | 0.41 | 0.40 | 0.40 | 1.00 |

Percent 0.010 0.015 0.020 0.025
