## Supplementary Figure 7 for "Distinct cellular DNA methylation mechanisms underlie common and rare genetic risk for brain disorders"

### ACBGM−Bergmann glia

| | 0.0−0.1 | 0.1−0.2 | 0.2−0.3 | 0.3−0.4 | 0.4−0.5 | 0.5−0.6 | 0.6−0.7 | 0.7−0.8 | 0.8−0.9 | 0.9−1.0 |
|---|---|---|---|---|---|---|---|---|---|---|
| L6b | 1.15 | 0.82 | 0.71 | 0.63 | 0.60 | 0.57 | 0.57 | 0.58 | 0.63 | 1.00 |
| L6−IT−Car3 | 0.69 | 0.50 | 0.48 | 0.45 | 0.45 | 0.42 | 0.41 | 0.42 | 0.47 | 1.00 |
| L6−IT | 0.80 | 0.59 | 0.52 | 0.48 | 0.46 | 0.44 | 0.42 | 0.42 | 0.45 | 1.00 |
| L6−CT | 1.01 | 0.83 | 0.78 | 0.73 | 0.71 | 0.71 | 0.74 | 0.72 | 0.80 | 1.00 |
| L56−NP | 1.44 | 1.08 | 0.92 | 0.87 | 0.81 | 0.74 | 0.69 | 0.67 | 0.68 | 1.00 |
| L5−IT | 0.88 | 0.66 | 0.58 | 0.58 | 0.55 | 0.54 | 0.51 | 0.53 | 0.57 | 1.00 |
| L5−ET | 0.81 | 0.55 | 0.51 | 0.49 | 0.44 | 0.42 | 0.43 | 0.44 | 0.49 | 1.00 |
| L4−IT | 0.49 | 0.37 | 0.35 | 0.33 | 0.32 | 0.32 | 0.32 | 0.34 | 0.39 | 1.00 |
| L23−IT | 0.67 | 0.48 | 0.45 | 0.42 | 0.42 | 0.42 | 0.41 | 0.41 | 0.47 | 1.00 |
| HIP−Misc2 | 0.85 | 0.64 | 0.58 | 0.54 | 0.51 | 0.47 | 0.47 | 0.47 | 0.54 | 1.00 |
| HIP−Misc1 | 6.36 | 4.64 | 4.07 | 3.66 | 3.31 | 2.99 | 2.54 | 2.13 | 1.62 | 1.00 |
| DG | 0.60 | 0.46 | 0.43 | 0.42 | 0.41 | 0.42 | 0.42 | 0.44 | 0.49 | 1.00 |
| CA3 | 0.81 | 0.71 | 0.63 | 0.61 | 0.59 | 0.56 | 0.56 | 0.56 | 0.60 | 1.00 |
| CA1 | 0.56 | 0.40 | 0.37 | 0.35 | 0.31 | 0.32 | 0.33 | 0.36 | 0.41 | 1.00 |
| Amy−Exc | 0.67 | 0.53 | 0.49 | 0.44 | 0.45 | 0.44 | 0.43 | 0.44 | 0.49 | 1.00 |
| Vip | 1.30 | 1.00 | 0.92 | 0.84 | 0.79 | 0.77 | 0.72 | 0.71 | 0.71 | 1.00 |
| THM−MB | 1.42 | 1.04 | 0.91 | 0.85 | 0.78 | 0.73 | 0.70 | 0.70 | 0.71 | 1.00 |
| THM−Inh | 5.45 | 3.60 | 3.20 | 2.78 | 2.54 | 2.15 | 2.00 | 1.82 | 1.55 | 1.00 |
| THM−Exc | 1.44 | 1.26 | 1.19 | 1.11 | 1.05 | 1.00 | 0.94 | 0.90 | 0.85 | 1.00 |
| SubCtx−Cplx | 0.70 | 0.61 | 0.55 | 0.56 | 0.55 | 0.55 | 0.54 | 0.53 | 0.57 | 1.00 |
| Sst | 0.92 | 0.75 | 0.69 | 0.65 | 0.61 | 0.57 | 0.58 | 0.56 | 0.57 | 1.00 |
| Sncg | 0.86 | 0.68 | 0.61 | 0.59 | 0.56 | 0.51 | 0.50 | 0.49 | 0.54 | 1.00 |
| Pvalb−ChC | 0.83 | 0.67 | 0.61 | 0.61 | 0.55 | 0.55 | 0.53 | 0.50 | 0.58 | 1.00 |
| Pvalb | 0.63 | 0.54 | 0.52 | 0.54 | 0.53 | 0.45 | 0.43 | 0.42 | 0.43 | 1.00 |
| PN | 1.23 | 0.71 | 0.60 | 0.55 | 0.50 | 0.48 | 0.47 | 0.47 | 0.52 | 1.00 |
| MSN−D2 | 0.68 | 0.58 | 0.51 | 0.48 | 0.50 | 0.47 | 0.46 | 0.45 | 0.51 | 1.00 |
| MSN−D1 | 0.74 | 0.60 | 0.56 | 0.52 | 0.50 | 0.49 | 0.49 | 0.48 | 0.50 | 1.00 |
| Lamp5−Lhx6 | 1.19 | 0.95 | 0.87 | 0.84 | 0.80 | 0.77 | 0.75 | 0.73 | 0.75 | 1.00 |
| Lamp5 | 1.01 | 0.75 | 0.69 | 0.68 | 0.67 | 0.62 | 0.61 | 0.60 | 0.65 | 1.00 |
| Foxp2 | 0.80 | 0.63 | 0.58 | 0.57 | 0.56 | 0.53 | 0.53 | 0.52 | 0.55 | 1.00 |
| Chd7 | 0.68 | 0.48 | 0.44 | 0.40 | 0.38 | 0.37 | 0.36 | 0.37 | 0.42 | 1.00 |
| CB | 5.63 | 4.24 | 3.91 | 3.68 | 3.34 | 2.99 | 2.70 | 2.13 | 1.85 | 1.00 |

Percent 0.01 0.02 0.03

### ASCNT−Non−telencephalon astrocytes

| | 0.0−0.1 | 0.1−0.2 | 0.2−0.3 | 0.3−0.4 | 0.4−0.5 | 0.5−0.6 | 0.6−0.7 | 0.7−0.8 | 0.8−0.9 | 0.9−1.0 |
|---|---|---|---|---|---|---|---|---|---|---|
| L6b | 1.85 | 1.33 | 1.12 | 0.99 | 0.91 | 0.83 | 0.78 | 0.75 | 0.73 | 1.00 |
| L6−IT−Car3 | 1.24 | 0.92 | 0.84 | 0.78 | 0.73 | 0.67 | 0.63 | 0.59 | 0.60 | 1.00 |
| L6−IT | 1.32 | 0.98 | 0.89 | 0.79 | 0.73 | 0.67 | 0.60 | 0.58 | 0.56 | 1.00 |
| L6−CT | 1.75 | 1.39 | 1.26 | 1.16 | 1.08 | 1.02 | 1.00 | 0.89 | 0.90 | 1.00 |
| L56−NP | 2.17 | 1.63 | 1.39 | 1.25 | 1.17 | 1.05 | 0.94 | 0.88 | 0.82 | 1.00 |
| L5−IT | 1.50 | 1.11 | 0.99 | 0.92 | 0.83 | 0.78 | 0.73 | 0.68 | 0.68 | 1.00 |
| L5−ET | 1.39 | 0.98 | 0.86 | 0.79 | 0.71 | 0.66 | 0.63 | 0.59 | 0.60 | 1.00 |
| L4−IT | 0.88 | 0.69 | 0.64 | 0.59 | 0.55 | 0.51 | 0.49 | 0.48 | 0.49 | 1.00 |
| L23−IT | 1.15 | 0.87 | 0.78 | 0.71 | 0.67 | 0.62 | 0.58 | 0.56 | 0.56 | 1.00 |
| HIP−Misc2 | 1.45 | 1.06 | 0.96 | 0.87 | 0.82 | 0.75 | 0.70 | 0.65 | 0.65 | 1.00 |
| HIP−Misc1 | 4.30 | 3.52 | 3.19 | 2.94 | 2.65 | 2.44 | 2.15 | 1.88 | 1.55 | 1.00 |
| DG | 1.04 | 0.82 | 0.73 | 0.69 | 0.68 | 0.66 | 0.63 | 0.61 | 0.62 | 1.00 |
| CA3 | 1.21 | 1.06 | 0.95 | 0.92 | 0.87 | 0.82 | 0.78 | 0.75 | 0.73 | 1.00 |
| CA1 | 1.03 | 0.75 | 0.66 | 0.60 | 0.55 | 0.51 | 0.49 | 0.48 | 0.52 | 1.00 |
| Amy−Exc | 1.16 | 0.93 | 0.83 | 0.77 | 0.71 | 0.67 | 0.64 | 0.61 | 0.60 | 1.00 |
| Vip | 2.04 | 1.58 | 1.40 | 1.28 | 1.16 | 1.08 | 0.98 | 0.91 | 0.86 | 1.00 |
| THM−MB | 2.30 | 1.70 | 1.42 | 1.27 | 1.14 | 1.04 | 0.92 | 0.87 | 0.83 | 1.00 |
| THM−Inh | 4.09 | 3.14 | 2.85 | 2.53 | 2.45 | 2.16 | 1.96 | 1.82 | 1.48 | 1.00 |
| THM−Exc | 1.82 | 1.63 | 1.51 | 1.42 | 1.34 | 1.25 | 1.18 | 1.09 | 0.99 | 1.00 |
| SubCtx−Cplx | 1.14 | 1.00 | 0.91 | 0.89 | 0.85 | 0.80 | 0.76 | 0.72 | 0.70 | 1.00 |
| Sst | 1.49 | 1.21 | 1.06 | 0.98 | 0.92 | 0.86 | 0.81 | 0.74 | 0.72 | 1.00 |
| Sncg | 1.47 | 1.17 | 1.04 | 0.96 | 0.88 | 0.79 | 0.72 | 0.70 | 0.66 | 1.00 |
| Pvalb−ChC | 1.36 | 1.12 | 0.96 | 0.90 | 0.83 | 0.78 | 0.72 | 0.67 | 0.66 | 1.00 |
| Pvalb | 1.04 | 0.80 | 0.70 | 0.75 | 0.93 | 0.79 | 0.70 | 0.65 | 0.59 | 1.00 |
| PN | 2.02 | 1.24 | 1.01 | 0.89 | 0.82 | 0.72 | 0.68 | 0.63 | 0.64 | 1.00 |
| MSN−D2 | 1.26 | 1.03 | 0.91 | 0.82 | 0.78 | 0.70 | 0.67 | 0.64 | 0.62 | 1.00 |
| MSN−D1 | 1.30 | 1.03 | 0.95 | 0.86 | 0.80 | 0.75 | 0.71 | 0.67 | 0.65 | 1.00 |
| Lamp5−Lhx6 | 1.77 | 1.43 | 1.33 | 1.23 | 1.16 | 1.09 | 1.02 | 0.95 | 0.89 | 1.00 |
| Lamp5 | 1.51 | 1.22 | 1.07 | 1.01 | 0.96 | 0.88 | 0.81 | 0.78 | 0.75 | 1.00 |
| Foxp2 | 1.32 | 1.05 | 0.97 | 0.89 | 0.84 | 0.79 | 0.75 | 0.69 | 0.68 | 1.00 |
| Chd7 | 1.23 | 0.90 | 0.78 | 0.70 | 0.64 | 0.60 | 0.55 | 0.52 | 0.54 | 1.00 |
| CB | 4.74 | 3.79 | 3.43 | 3.23 | 2.86 | 2.67 | 2.32 | 1.96 | 1.62 | 1.00 |

Percent 0.02 0.04 0.06 0.08

### ASCT−Telencephalon astrocytes

| | 0.0−0.1 | 0.1−0.2 | 0.2−0.3 | 0.3−0.4 | 0.4−0.5 | 0.5−0.6 | 0.6−0.7 | 0.7−0.8 | 0.8−0.9 | 0.9−1.0 |
|---|---|---|---|---|---|---|---|---|---|---|
| L6b | 2.39 | 1.72 | 1.46 | 1.29 | 1.17 | 1.08 | 1.00 | 0.91 | 0.85 | 1.00 |
| L6−IT−Car3 | 1.67 | 1.25 | 1.14 | 1.04 | 0.96 | 0.88 | 0.82 | 0.76 | 0.72 | 1.00 |
| L6−IT | 1.76 | 1.29 | 1.17 | 1.05 | 0.99 | 0.90 | 0.81 | 0.75 | 0.70 | 1.00 |
| L6−CT | 2.33 | 1.84 | 1.66 | 1.54 | 1.44 | 1.33 | 1.26 | 1.13 | 1.05 | 1.00 |
| L56−NP | 2.61 | 2.02 | 1.77 | 1.64 | 1.49 | 1.35 | 1.21 | 1.09 | 0.98 | 1.00 |
| L5−IT | 1.98 | 1.48 | 1.31 | 1.21 | 1.11 | 1.03 | 0.94 | 0.86 | 0.81 | 1.00 |
| L5−ET | 1.92 | 1.36 | 1.18 | 1.06 | 0.96 | 0.87 | 0.80 | 0.76 | 0.71 | 1.00 |
| L4−IT | 1.21 | 0.95 | 0.87 | 0.81 | 0.74 | 0.70 | 0.66 | 0.62 | 0.60 | 1.00 |
| L23−IT | 1.57 | 1.18 | 1.07 | 0.97 | 0.90 | 0.85 | 0.77 | 0.72 | 0.69 | 1.00 |
| HIP−Misc2 | 1.88 | 1.42 | 1.27 | 1.17 | 1.07 | 0.97 | 0.91 | 0.83 | 0.78 | 1.00 |
| HIP−Misc1 | 3.30 | 2.87 | 2.65 | 2.45 | 2.30 | 2.12 | 1.96 | 1.73 | 1.44 | 1.00 |
| DG | 1.45 | 1.11 | 1.01 | 0.95 | 0.89 | 0.85 | 0.82 | 0.78 | 0.74 | 1.00 |
| CA3 | 1.49 | 1.32 | 1.22 | 1.15 | 1.09 | 1.01 | 0.98 | 0.91 | 0.84 | 1.00 |
| CA1 | 1.44 | 1.04 | 0.94 | 0.84 | 0.78 | 0.71 | 0.66 | 0.64 | 0.63 | 1.00 |
| Amy−Exc | 1.57 | 1.25 | 1.12 | 1.05 | 0.97 | 0.90 | 0.84 | 0.78 | 0.73 | 1.00 |
| Vip | 2.56 | 2.04 | 1.84 | 1.68 | 1.55 | 1.41 | 1.27 | 1.14 | 1.01 | 1.00 |
| THM−MB | 2.92 | 2.18 | 1.87 | 1.66 | 1.48 | 1.36 | 1.19 | 1.08 | 0.97 | 1.00 |
| THM−Inh | 3.28 | 2.67 | 2.53 | 2.33 | 2.19 | 2.05 | 1.88 | 1.71 | 1.45 | 1.00 |
| THM−Exc | 1.86 | 1.68 | 1.58 | 1.50 | 1.42 | 1.34 | 1.26 | 1.18 | 1.06 | 1.00 |
| SubCtx−Cplx | 1.47 | 1.32 | 1.22 | 1.18 | 1.12 | 1.07 | 1.00 | 0.91 | 0.84 | 1.00 |
| Sst | 1.96 | 1.60 | 1.44 | 1.34 | 1.25 | 1.17 | 1.07 | 0.96 | 0.87 | 1.00 |
| Sncg | 1.96 | 1.61 | 1.46 | 1.33 | 1.20 | 1.10 | 1.00 | 0.91 | 0.82 | 1.00 |
| Pvalb−ChC | 1.78 | 1.47 | 1.31 | 1.22 | 1.11 | 1.03 | 0.94 | 0.86 | 0.81 | 1.00 |
| Pvalb | 1.31 | 1.01 | 0.89 | 0.89 | 1.19 | 1.03 | 0.93 | 0.82 | 0.74 | 1.00 |
| PN | 2.66 | 1.70 | 1.41 | 1.22 | 1.10 | 1.00 | 0.91 | 0.84 | 0.76 | 1.00 |
| MSN−D2 | 1.74 | 1.43 | 1.27 | 1.15 | 1.06 | 0.96 | 0.88 | 0.80 | 0.76 | 1.00 |
| MSN−D1 | 1.70 | 1.40 | 1.27 | 1.17 | 1.09 | 1.01 | 0.93 | 0.85 | 0.78 | 1.00 |
| Lamp5−Lhx6 | 2.16 | 1.81 | 1.68 | 1.56 | 1.46 | 1.36 | 1.26 | 1.15 | 1.03 | 1.00 |
| Lamp5 | 1.94 | 1.58 | 1.42 | 1.32 | 1.25 | 1.14 | 1.07 | 0.98 | 0.88 | 1.00 |
| Foxp2 | 1.72 | 1.42 | 1.29 | 1.22 | 1.14 | 1.06 | 0.98 | 0.89 | 0.81 | 1.00 |
| Chd7 | 1.69 | 1.26 | 1.10 | 0.99 | 0.91 | 0.83 | 0.76 | 0.69 | 0.67 | 1.00 |
| CB | 3.99 | 3.36 | 3.10 | 2.92 | 2.65 | 2.40 | 2.19 | 1.85 | 1.55 | 1.00 |

Percent 0.05 0.10

#### Amy−Exc−Glutamatergic neurons from amygdala

| | 0.0−0.1 | 0.1−0.2 | 0.2−0.3 | 0.3−0.4 | 0.4−0.5 | 0.5−0.6 | 0.6−0.7 | 0.7−0.8 | 0.8−0.9 | 0.9−1.0 |
|---|---|---|---|---|---|---|---|---|---|---|
| L6b | 1.72 | 1.13 | 0.99 | 0.90 | 0.85 | 0.82 | 0.80 | 0.79 | 0.79 | 1.00 |
| L6−IT−Car3 | 1.26 | 0.81 | 0.74 | 0.70 | 0.67 | 0.66 | 0.66 | 0.64 | 0.67 | 1.00 |
| L6−IT | 1.36 | 0.87 | 0.78 | 0.73 | 0.70 | 0.65 | 0.65 | 0.64 | 0.65 | 1.00 |
| L6−CT | 1.53 | 1.16 | 1.10 | 1.05 | 1.01 | 1.01 | 0.98 | 0.93 | 0.96 | 1.00 |
| L56−NP | 1.70 | 1.27 | 1.16 | 1.07 | 1.03 | 0.99 | 0.96 | 0.91 | 0.88 | 1.00 |
| L5−IT | 1.48 | 0.95 | 0.85 | 0.80 | 0.78 | 0.75 | 0.75 | 0.73 | 0.74 | 1.00 |
| L5−ET | 1.47 | 0.91 | 0.81 | 0.74 | 0.69 | 0.65 | 0.67 | 0.64 | 0.65 | 1.00 |
| L4−IT | 0.89 | 0.64 | 0.58 | 0.56 | 0.54 | 0.54 | 0.53 | 0.52 | 0.56 | 1.00 |
| L23−IT | 1.23 | 0.81 | 0.73 | 0.68 | 0.64 | 0.64 | 0.61 | 0.62 | 0.62 | 1.00 |
| HIP−Misc2 | 1.38 | 0.93 | 0.83 | 0.78 | 0.76 | 0.76 | 0.72 | 0.71 | 0.74 | 1.00 |
| HIP−Misc1 | 2.32 | 1.97 | 1.85 | 1.73 | 1.69 | 1.62 | 1.50 | 1.40 | 1.28 | 1.00 |
| DG | 0.99 | 0.73 | 0.68 | 0.65 | 0.65 | 0.66 | 0.66 | 0.66 | 0.68 | 1.00 |
| CA3 | 0.99 | 0.89 | 0.86 | 0.85 | 0.84 | 0.82 | 0.83 | 0.80 | 0.82 | 1.00 |
| CA1 | 1.07 | 0.67 | 0.61 | 0.57 | 0.55 | 0.53 | 0.53 | 0.55 | 0.58 | 1.00 |
| Amy−Exc | 1.21 | 0.82 | 0.75 | 0.71 | 0.69 | 0.67 | 0.67 | 0.66 | 0.67 | 1.00 |
| Vip | 1.45 | 1.26 | 1.19 | 1.14 | 1.12 | 1.05 | 1.03 | 1.01 | 0.95 | 1.00 |
| THM−MB | 1.65 | 1.27 | 1.14 | 1.09 | 1.04 | 1.00 | 0.93 | 0.91 | 0.90 | 1.00 |
| THM−Inh | 2.21 | 1.72 | 1.65 | 1.58 | 1.53 | 1.48 | 1.43 | 1.35 | 1.23 | 1.00 |
| THM−Exc | 1.34 | 1.25 | 1.19 | 1.16 | 1.12 | 1.10 | 1.06 | 1.02 | 0.98 | 1.00 |
| SubCtx−Cplx | 0.93 | 0.86 | 0.83 | 0.83 | 0.83 | 0.83 | 0.81 | 0.80 | 0.78 | 1.00 |
| Sst | 1.27 | 1.03 | 0.97 | 0.94 | 0.91 | 0.88 | 0.84 | 0.82 | 0.80 | 1.00 |
| Sncg | 1.14 | 0.98 | 0.92 | 0.90 | 0.87 | 0.84 | 0.82 | 0.80 | 0.78 | 1.00 |
| Pvalb−ChC | 1.10 | 0.95 | 0.89 | 0.86 | 0.84 | 0.83 | 0.79 | 0.78 | 0.76 | 1.00 |
| Pvalb | 0.85 | 0.80 | 0.78 | 0.76 | 0.78 | 0.71 | 0.70 | 0.67 | 0.67 | 1.00 |
| PN | 1.63 | 1.07 | 0.91 | 0.84 | 0.80 | 0.75 | 0.74 | 0.74 | 0.73 | 1.00 |
| MSN−D2 | 1.17 | 0.90 | 0.82 | 0.80 | 0.76 | 0.75 | 0.73 | 0.72 | 0.71 | 1.00 |
| MSN−D1 | 1.20 | 0.92 | 0.84 | 0.80 | 0.78 | 0.75 | 0.75 | 0.74 | 0.74 | 1.00 |
| Lamp5−Lhx6 | 1.33 | 1.19 | 1.11 | 1.11 | 1.11 | 1.03 | 1.04 | 0.99 | 0.94 | 1.00 |
| Lamp5 | 1.23 | 1.04 | 0.99 | 0.97 | 0.94 | 0.91 | 0.89 | 0.84 | 0.84 | 1.00 |
| Foxp2 | 1.14 | 0.91 | 0.86 | 0.85 | 0.85 | 0.80 | 0.79 | 0.77 | 0.76 | 1.00 |
| Chd7 | 0.98 | 0.78 | 0.72 | 0.69 | 0.66 | 0.65 | 0.64 | 0.63 | 0.64 | 1.00 |
| CB | 2.57 | 2.23 | 2.12 | 2.04 | 1.94 | 1.81 | 1.72 | 1.53 | 1.37 | 1.00 |

Percent 0.025 0.050 0.075

#### BFEXA−GABAergic neurons from basal forebrain and amygdala

| | 0.0−0.1 | 0.1−0.2 | 0.2−0.3 | 0.3−0.4 | 0.4−0.5 | 0.5−0.6 | 0.6−0.7 | 0.7−0.8 | 0.8−0.9 | 0.9−1.0 |
|---|---|---|---|---|---|---|---|---|---|---|
| L6b | 1.23 | 0.85 | 0.74 | 0.72 | 0.69 | 0.65 | 0.68 | 0.68 | 0.71 | 1.00 |
| L6−IT−Car3 | 0.79 | 0.54 | 0.52 | 0.51 | 0.50 | 0.50 | 0.50 | 0.53 | 0.58 | 1.00 |
| L6−IT | 0.87 | 0.59 | 0.56 | 0.53 | 0.53 | 0.52 | 0.50 | 0.53 | 0.54 | 1.00 |
| L6−CT | 1.05 | 0.85 | 0.84 | 0.82 | 0.81 | 0.81 | 0.82 | 0.82 | 0.86 | 1.00 |
| L56−NP | 1.40 | 1.00 | 0.90 | 0.85 | 0.83 | 0.79 | 0.77 | 0.76 | 0.77 | 1.00 |
| L5−IT | 0.95 | 0.68 | 0.62 | 0.62 | 0.61 | 0.61 | 0.60 | 0.62 | 0.65 | 1.00 |
| L5−ET | 0.99 | 0.62 | 0.59 | 0.56 | 0.54 | 0.52 | 0.52 | 0.51 | 0.58 | 1.00 |
| L4−IT | 0.56 | 0.43 | 0.41 | 0.40 | 0.39 | 0.39 | 0.40 | 0.41 | 0.46 | 1.00 |
| L23−IT | 0.80 | 0.55 | 0.50 | 0.50 | 0.48 | 0.49 | 0.49 | 0.51 | 0.55 | 1.00 |
| HIP−Misc2 | 0.95 | 0.66 | 0.61 | 0.62 | 0.60 | 0.57 | 0.58 | 0.60 | 0.64 | 1.00 |
| HIP−Misc1 | 2.97 | 2.39 | 2.15 | 2.01 | 1.89 | 1.78 | 1.63 | 1.53 | 1.31 | 1.00 |
| DG | 0.65 | 0.52 | 0.51 | 0.49 | 0.48 | 0.51 | 0.52 | 0.55 | 0.58 | 1.00 |
| CA3 | 0.74 | 0.68 | 0.67 | 0.65 | 0.67 | 0.67 | 0.66 | 0.68 | 0.71 | 1.00 |
| CA1 | 0.66 | 0.47 | 0.42 | 0.39 | 0.40 | 0.40 | 0.42 | 0.44 | 0.50 | 1.00 |
| Amy−Exc | 0.78 | 0.58 | 0.54 | 0.52 | 0.52 | 0.52 | 0.53 | 0.54 | 0.59 | 1.00 |
| Vip | 1.14 | 0.96 | 0.93 | 0.86 | 0.86 | 0.86 | 0.85 | 0.84 | 0.82 | 1.00 |
| THM−MB | 1.38 | 1.00 | 0.91 | 0.89 | 0.81 | 0.83 | 0.77 | 0.78 | 0.78 | 1.00 |
| THM−Inh | 2.83 | 2.00 | 1.88 | 1.74 | 1.63 | 1.56 | 1.48 | 1.42 | 1.32 | 1.00 |
| THM−Exc | 1.27 | 1.14 | 1.10 | 1.07 | 1.03 | 1.00 | 0.97 | 0.94 | 0.92 | 1.00 |
| SubCtx−Cplx | 0.70 | 0.63 | 0.60 | 0.62 | 0.62 | 0.61 | 0.62 | 0.62 | 0.66 | 1.00 |
| Sst | 0.97 | 0.81 | 0.74 | 0.73 | 0.70 | 0.69 | 0.70 | 0.67 | 0.69 | 1.00 |
| Sncg | 0.79 | 0.68 | 0.64 | 0.64 | 0.65 | 0.63 | 0.63 | 0.62 | 0.66 | 1.00 |
| Pvalb−ChC | 0.79 | 0.67 | 0.65 | 0.65 | 0.63 | 0.63 | 0.62 | 0.61 | 0.67 | 1.00 |
| Pvalb | 0.65 | 0.62 | 0.61 | 0.63 | 0.56 | 0.51 | 0.52 | 0.51 | 0.54 | 1.00 |
| PN | 1.05 | 0.72 | 0.64 | 0.61 | 0.59 | 0.58 | 0.58 | 0.60 | 0.62 | 1.00 |
| MSN−D2 | 0.90 | 0.64 | 0.61 | 0.58 | 0.57 | 0.54 | 0.55 | 0.57 | 0.61 | 1.00 |
| MSN−D1 | 0.97 | 0.68 | 0.62 | 0.58 | 0.58 | 0.55 | 0.55 | 0.56 | 0.61 | 1.00 |
| Lamp5−Lhx6 | 1.09 | 0.95 | 0.87 | 0.89 | 0.86 | 0.88 | 0.84 | 0.84 | 0.84 | 1.00 |
| Lamp5 | 0.95 | 0.79 | 0.75 | 0.74 | 0.74 | 0.73 | 0.73 | 0.71 | 0.71 | 1.00 |
| Foxp2 | 0.91 | 0.69 | 0.65 | 0.64 | 0.65 | 0.62 | 0.63 | 0.64 | 0.66 | 1.00 |
| Chd7 | 0.67 | 0.51 | 0.48 | 0.48 | 0.47 | 0.48 | 0.47 | 0.49 | 0.54 | 1.00 |
| CB | 2.97 | 2.38 | 2.30 | 2.18 | 2.04 | 1.93 | 1.83 | 1.60 | 1.46 | 1.00 |

Percent 0.01 0.02 0.03 0.04

#### BNGA−GABAergic neurons from basal nuclei

| | 0.0−0.1 | 0.1−0.2 | 0.2−0.3 | 0.3−0.4 | 0.4−0.5 | 0.5−0.6 | 0.6−0.7 | 0.7−0.8 | 0.8−0.9 | 0.9−1.0 |
|---|---|---|---|---|---|---|---|---|---|---|
| L6b | 1.16 | 0.84 | 0.73 | 0.68 | 0.65 | 0.62 | 0.63 | 0.64 | 0.66 | 1.00 |
| L6−IT−Car3 | 0.76 | 0.52 | 0.49 | 0.48 | 0.44 | 0.44 | 0.46 | 0.48 | 0.52 | 1.00 |
| L6−IT | 0.84 | 0.58 | 0.53 | 0.50 | 0.48 | 0.48 | 0.43 | 0.45 | 0.49 | 1.00 |
| L6−CT | 1.00 | 0.78 | 0.80 | 0.74 | 0.75 | 0.76 | 0.71 | 0.72 | 0.76 | 1.00 |
| L56−NP | 1.43 | 1.04 | 0.94 | 0.86 | 0.81 | 0.78 | 0.75 | 0.72 | 0.72 | 1.00 |
| L5−IT | 0.95 | 0.64 | 0.63 | 0.59 | 0.58 | 0.59 | 0.58 | 0.56 | 0.64 | 1.00 |
| L5−ET | 0.92 | 0.60 | 0.55 | 0.53 | 0.48 | 0.46 | 0.45 | 0.48 | 0.55 | 1.00 |
| L4−IT | 0.53 | 0.38 | 0.39 | 0.36 | 0.34 | 0.35 | 0.36 | 0.37 | 0.41 | 1.00 |
| L23−IT | 0.76 | 0.53 | 0.48 | 0.47 | 0.45 | 0.46 | 0.42 | 0.47 | 0.50 | 1.00 |
| HIP−Misc2 | 0.95 | 0.66 | 0.61 | 0.58 | 0.56 | 0.53 | 0.54 | 0.56 | 0.59 | 1.00 |
| HIP−Misc1 | 4.37 | 3.49 | 3.01 | 2.81 | 2.63 | 2.44 | 1.98 | 1.78 | 1.51 | 1.00 |
| DG | 0.65 | 0.50 | 0.46 | 0.45 | 0.46 | 0.48 | 0.46 | 0.48 | 0.54 | 1.00 |
| CA3 | 0.79 | 0.66 | 0.70 | 0.65 | 0.63 | 0.65 | 0.63 | 0.61 | 0.68 | 1.00 |
| CA1 | 0.62 | 0.44 | 0.38 | 0.36 | 0.35 | 0.36 | 0.37 | 0.41 | 0.45 | 1.00 |
| Amy−Exc | 0.75 | 0.54 | 0.52 | 0.49 | 0.48 | 0.48 | 0.48 | 0.46 | 0.53 | 1.00 |
| Vip | 1.13 | 0.99 | 0.92 | 0.87 | 0.81 | 0.82 | 0.81 | 0.77 | 0.78 | 1.00 |
| THM−MB | 1.37 | 0.99 | 0.92 | 0.84 | 0.78 | 0.78 | 0.73 | 0.73 | 0.75 | 1.00 |
| THM−Inh | 3.97 | 2.84 | 2.50 | 2.31 | 2.10 | 1.92 | 1.84 | 1.71 | 1.40 | 1.00 |
| THM−Exc | 1.20 | 1.11 | 1.07 | 1.03 | 0.99 | 0.97 | 0.94 | 0.92 | 0.88 | 1.00 |
| SubCtx−Cplx | 0.69 | 0.62 | 0.60 | 0.60 | 0.59 | 0.58 | 0.57 | 0.56 | 0.61 | 1.00 |
| Sst | 0.93 | 0.76 | 0.72 | 0.68 | 0.67 | 0.63 | 0.63 | 0.60 | 0.62 | 1.00 |
| Sncg | 0.72 | 0.66 | 0.60 | 0.59 | 0.56 | 0.55 | 0.55 | 0.55 | 0.62 | 1.00 |
| Pvalb−ChC | 0.80 | 0.69 | 0.62 | 0.63 | 0.59 | 0.58 | 0.59 | 0.59 | 0.63 | 1.00 |
| Pvalb | 0.66 | 0.63 | 0.67 | 0.66 | 0.54 | 0.48 | 0.46 | 0.46 | 0.50 | 1.00 |
| PN | 1.15 | 0.71 | 0.61 | 0.56 | 0.54 | 0.52 | 0.51 | 0.54 | 0.56 | 1.00 |
| MSN−D2 | 0.87 | 0.60 | 0.55 | 0.53 | 0.51 | 0.48 | 0.51 | 0.50 | 0.53 | 1.00 |
| MSN−D1 | 0.92 | 0.63 | 0.57 | 0.55 | 0.52 | 0.51 | 0.50 | 0.50 | 0.55 | 1.00 |
| Lamp5−Lhx6 | 1.09 | 0.93 | 0.88 | 0.86 | 0.88 | 0.85 | 0.83 | 0.80 | 0.79 | 1.00 |
| Lamp5 | 0.93 | 0.76 | 0.75 | 0.71 | 0.71 | 0.70 | 0.68 | 0.65 | 0.70 | 1.00 |
| Foxp2 | 0.86 | 0.67 | 0.63 | 0.60 | 0.59 | 0.57 | 0.57 | 0.57 | 0.64 | 1.00 |
| Chd7 | 0.60 | 0.45 | 0.44 | 0.43 | 0.41 | 0.40 | 0.40 | 0.40 | 0.48 | 1.00 |
| CB | 4.02 | 3.25 | 3.07 | 2.91 | 2.68 | 2.48 | 2.29 | 1.89 | 1.62 | 1.00 |

Percent 0.004 0.008 0.012 0.016

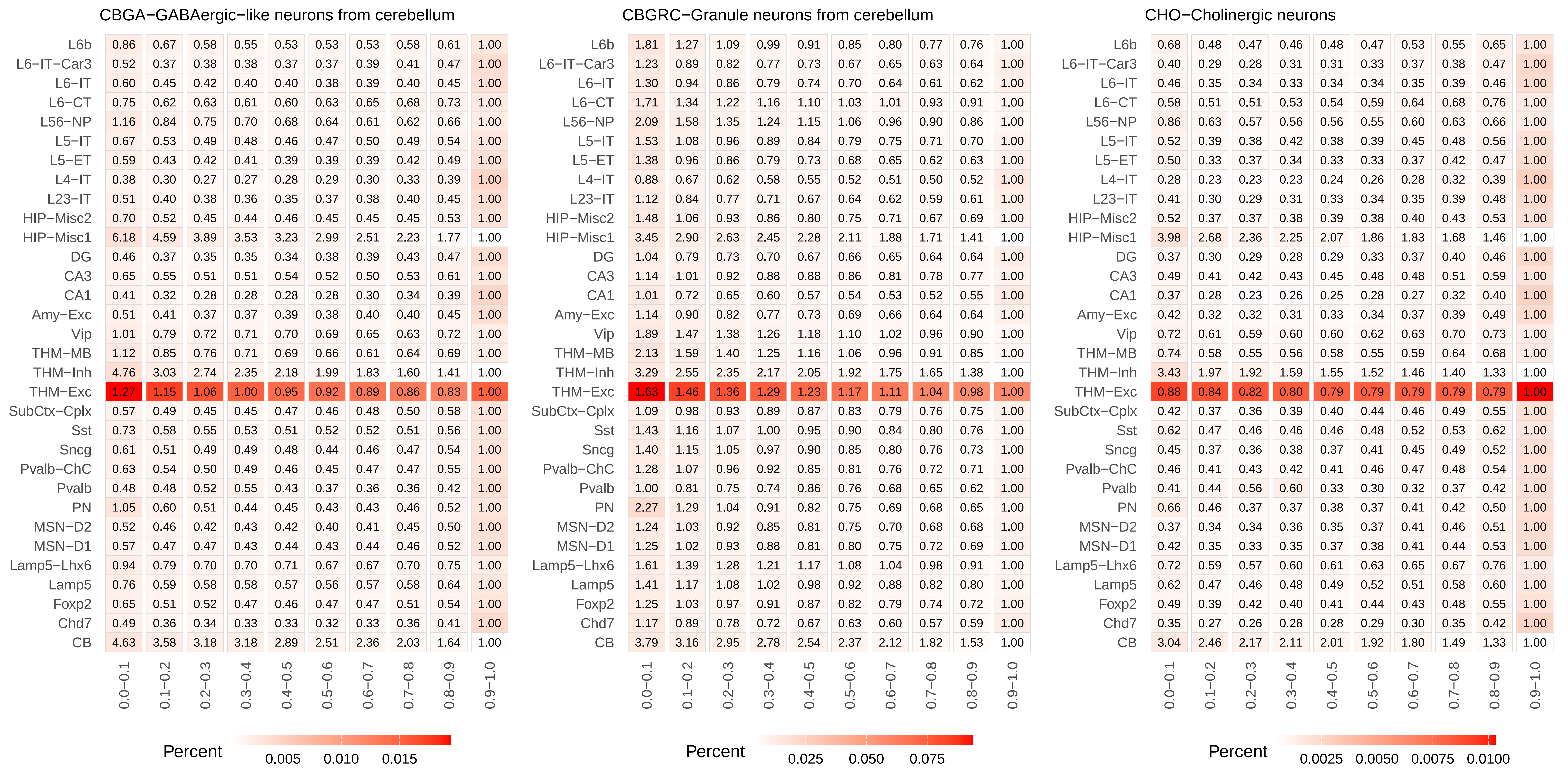

CBGA−GABAergic−like neurons from cerebellum
CBGRC−Granule neurons from cerebellum
CHO−Cholinergic neurons
L6b
L6−IT−Car3
L6−IT
L6−CT
L56−NP
L5−IT
L5−ET
L4−IT
L23−IT
HIP−Misc2
HIP−Misc1
DG
CA3
CA1
Amy−Exc
Vip
THM−MB
THM−Inh
THM−Exc
SubCtx−Cplx
Sst
Sncg
Pvalb−ChC
Pvalb
PN
MSN−D2
MSN−D1
Lamp5−Lhx6
Lamp5
Foxp2
Chd7
CB
0.0−0.1
0.1−0.2
0.2−0.3
0.3−0.4
0.4−0.5
0.5−0.6
0.6−0.7
0.7−0.8
0.8−0.9
0.9−1.0
Percent
0.005
0.010
0.015
Percent
0.025
0.050
0.075
Percent
0.0025
0.0050
0.0075
0.0100

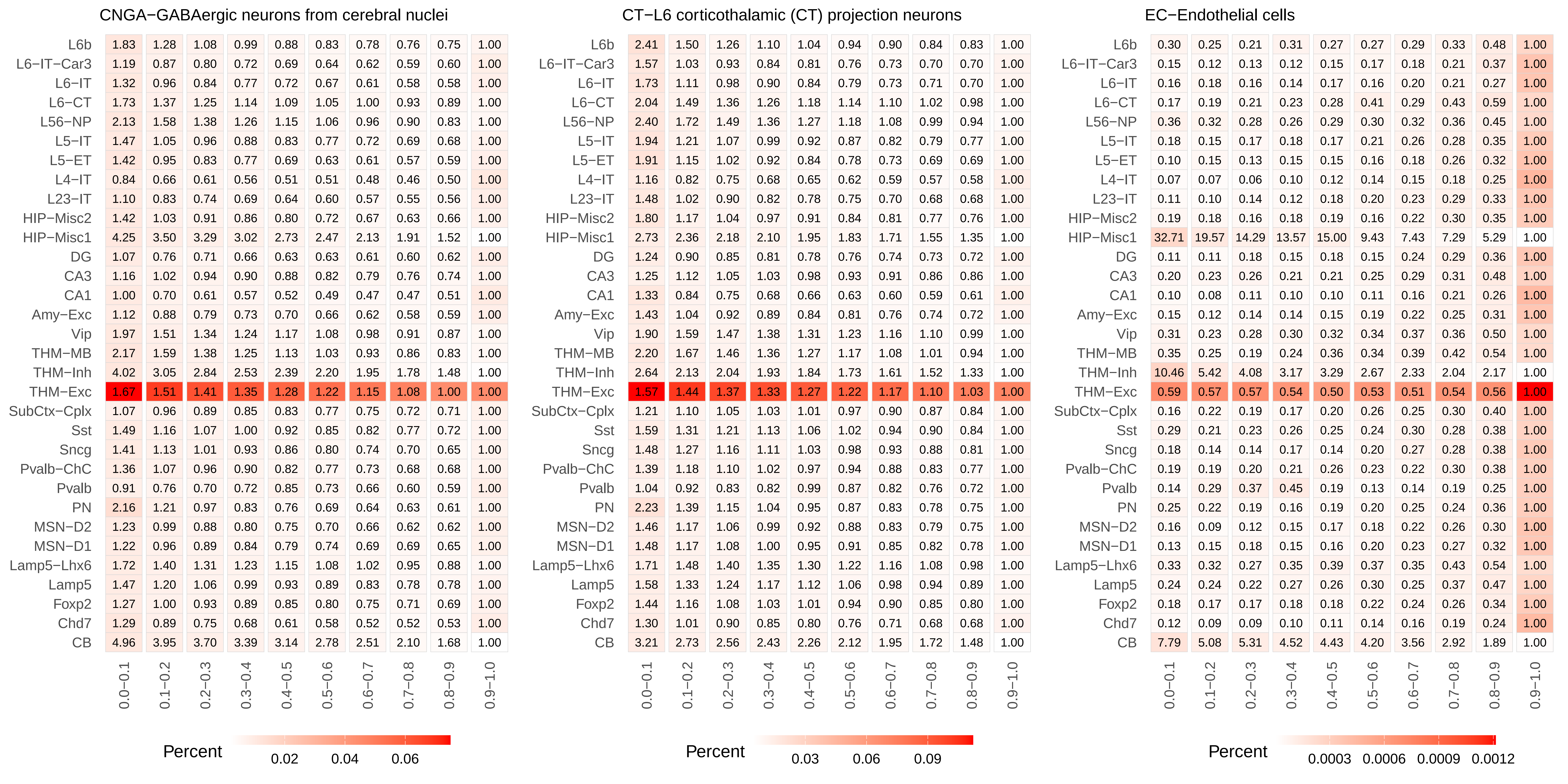
CNGA−GABAergic neurons from cerebral nuclei
L6b 1.83 1.28 1.08 0.99 0.88 0.83 0.78 0.76 0.75 1.00
L6−IT−Car3 1.19 0.87 0.80 0.72 0.69 0.64 0.62 0.59 0.60 1.00
L6−IT 1.32 0.96 0.84 0.77 0.72 0.67 0.61 0.58 0.58 1.00
L6−CT 1.73 1.37 1.25 1.14 1.09 1.05 1.00 0.93 0.89 1.00
L56−NP 2.13 1.58 1.38 1.26 1.15 1.06 0.96 0.90 0.83 1.00
L5−IT 1.47 1.05 0.96 0.88 0.83 0.77 0.72 0.69 0.68 1.00
L5−ET 1.42 0.95 0.83 0.77 0.69 0.63 0.61 0.57 0.59 1.00
L4−IT 0.84 0.66 0.61 0.56 0.51 0.51 0.48 0.46 0.50 1.00
L23−IT 1.10 0.83 0.74 0.69 0.64 0.60 0.57 0.55 0.56 1.00
HIP−Misc2 1.42 1.03 0.91 0.86 0.80 0.72 0.67 0.63 0.66 1.00
HIP−Misc1 4.25 3.50 3.29 3.02 2.73 2.47 2.13 1.91 1.52 1.00
DG 1.07 0.76 0.71 0.66 0.63 0.63 0.61 0.60 0.62 1.00
CA3 1.16 1.02 0.94 0.90 0.88 0.82 0.79 0.76 0.74 1.00
CA1 1.00 0.70 0.61 0.57 0.52 0.49 0.47 0.47 0.51 1.00
Amy−Exc 1.12 0.88 0.79 0.73 0.70 0.66 0.62 0.58 0.59 1.00
Vip 1.97 1.51 1.34 1.24 1.17 1.08 0.98 0.91 0.87 1.00
THM−MB 2.17 1.59 1.38 1.25 1.13 1.03 0.93 0.86 0.83 1.00
THM−Inh 4.02 3.05 2.84 2.53 2.39 2.20 1.95 1.78 1.48 1.00
THM−Exc 1.67 1.51 1.41 1.35 1.28 1.22 1.15 1.08 1.00 1.00
SubCtx−Cplx 1.07 0.96 0.89 0.85 0.83 0.77 0.75 0.72 0.71 1.00
Sst 1.49 1.16 1.07 1.00 0.92 0.85 0.82 0.77 0.72 1.00
Sncg 1.41 1.13 1.01 0.93 0.86 0.80 0.74 0.70 0.65 1.00
Pvalb−ChC 1.36 1.07 0.96 0.90 0.82 0.77 0.73 0.68 0.68 1.00
Pvalb 0.91 0.76 0.70 0.72 0.85 0.73 0.66 0.60 0.59 1.00
PN 2.16 1.21 0.97 0.83 0.76 0.69 0.64 0.63 0.61 1.00
MSN−D2 1.23 0.99 0.88 0.80 0.75 0.70 0.66 0.62 0.62 1.00
MSN−D1 1.22 0.96 0.89 0.84 0.79 0.74 0.69 0.69 0.65 1.00
Lamp5−Lhx6 1.72 1.40 1.31 1.23 1.15 1.08 1.02 0.95 0.88 1.00
Lamp5 1.47 1.20 1.06 0.99 0.93 0.89 0.83 0.78 0.78 1.00
Foxp2 1.27 1.00 0.93 0.89 0.85 0.80 0.75 0.71 0.69 1.00
Chd7 1.29 0.89 0.75 0.68 0.61 0.58 0.52 0.52 0.53 1.00
CB 4.96 3.95 3.70 3.39 3.14 2.78 2.51 2.10 1.68 1.00
0.0−0.1 0.1−0.2 0.2−0.3 0.3−0.4 0.4−0.5 0.5−0.6 0.6−0.7 0.7−0.8 0.8−0.9 0.9−1.0
Percent
0.02 0.04 0.06
CT−L6 corticothalamic (CT) projection neurons
L6b 2.41 1.50 1.26 1.10 1.04 0.94 0.90 0.84 0.83 1.00
L6−IT−Car3 1.57 1.03 0.93 0.84 0.81 0.76 0.73 0.70 0.70 1.00
L6−IT 1.73 1.11 0.98 0.90 0.84 0.79 0.73 0.71 0.70 1.00
L6−CT 2.04 1.49 1.36 1.26 1.18 1.14 1.10 1.02 0.98 1.00
L56−NP 2.40 1.72 1.49 1.36 1.27 1.18 1.08 0.99 0.94 1.00
L5−IT 1.94 1.21 1.07 0.99 0.92 0.87 0.82 0.79 0.77 1.00
L5−ET 1.91 1.15 1.02 0.92 0.84 0.78 0.73 0.69 0.69 1.00
L4−IT 1.16 0.82 0.75 0.68 0.65 0.62 0.59 0.57 0.58 1.00
L23−IT 1.48 1.02 0.90 0.82 0.78 0.75 0.70 0.68 0.68 1.00
HIP−Misc2 1.80 1.17 1.04 0.97 0.91 0.84 0.81 0.77 0.76 1.00
HIP−Misc1 2.73 2.36 2.18 2.10 1.95 1.83 1.71 1.55 1.35 1.00
DG 1.24 0.90 0.85 0.81 0.78 0.76 0.74 0.73 0.72 1.00
CA3 1.25 1.12 1.05 1.03 0.98 0.93 0.91 0.86 0.86 1.00
CA1 1.33 0.84 0.75 0.68 0.66 0.63 0.60 0.59 0.61 1.00
Amy−Exc 1.43 1.04 0.92 0.89 0.84 0.81 0.76 0.74 0.72 1.00
Vip 1.90 1.59 1.47 1.38 1.31 1.23 1.16 1.10 0.99 1.00
THM−MB 2.20 1.67 1.46 1.36 1.27 1.17 1.08 1.01 0.94 1.00
THM−Inh 2.64 2.13 2.04 1.93 1.84 1.73 1.61 1.52 1.33 1.00
THM−Exc 1.57 1.44 1.37 1.33 1.27 1.22 1.17 1.10 1.03 1.00
SubCtx−Cplx 1.21 1.10 1.05 1.03 1.01 0.97 0.90 0.87 0.84 1.00
Sst 1.59 1.31 1.21 1.13 1.06 1.02 0.94 0.90 0.84 1.00
Sncg 1.48 1.27 1.16 1.11 1.03 0.98 0.93 0.88 0.81 1.00
Pvalb−ChC 1.39 1.18 1.10 1.02 0.97 0.94 0.88 0.83 0.77 1.00
Pvalb 1.04 0.92 0.83 0.82 0.99 0.87 0.82 0.76 0.72 1.00
PN 2.23 1.39 1.15 1.04 0.95 0.87 0.83 0.78 0.75 1.00
MSN−D2 1.46 1.17 1.06 0.99 0.92 0.88 0.83 0.79 0.75 1.00
MSN−D1 1.48 1.17 1.08 1.00 0.95 0.91 0.85 0.82 0.78 1.00
Lamp5−Lhx6 1.71 1.48 1.40 1.35 1.30 1.22 1.16 1.08 0.98 1.00
Lamp5 1.58 1.33 1.24 1.17 1.12 1.06 0.98 0.94 0.89 1.00
Foxp2 1.44 1.16 1.08 1.03 1.01 0.94 0.90 0.85 0.80 1.00
Chd7 1.30 1.01 0.90 0.85 0.80 0.76 0.71 0.68 0.68 1.00
CB 3.21 2.73 2.56 2.43 2.26 2.12 1.95 1.72 1.48 1.00
0.0−0.1 0.1−0.2 0.2−0.3 0.3−0.4 0.4−0.5 0.5−0.6 0.6−0.7 0.7−0.8 0.8−0.9 0.9−1.0
Percent
0.03 0.06 0.09
EC−Endothelial cells
L6b 0.30 0.25 0.21 0.31 0.27 0.27 0.29 0.33 0.48 1.00
L6−IT−Car3 0.15 0.12 0.13 0.12 0.15 0.17 0.18 0.21 0.37 1.00
L6−IT 0.16 0.18 0.16 0.14 0.17 0.16 0.20 0.21 0.27 1.00
L6−CT 0.17 0.19 0.21 0.23 0.28 0.41 0.29 0.43 0.59 1.00
L56−NP 0.36 0.32 0.28 0.26 0.29 0.30 0.32 0.36 0.45 1.00
L5−IT 0.18 0.15 0.17 0.18 0.17 0.21 0.26 0.28 0.35 1.00
L5−ET 0.10 0.15 0.13 0.15 0.15 0.16 0.18 0.26 0.32 1.00
L4−IT 0.07 0.07 0.06 0.10 0.12 0.14 0.15 0.18 0.25 1.00
L23−IT 0.11 0.10 0.14 0.12 0.18 0.20 0.23 0.29 0.33 1.00
HIP−Misc2 0.19 0.18 0.16 0.18 0.19 0.16 0.22 0.30 0.35 1.00
HIP−Misc1 32.71 19.57 14.29 13.57 15.00 9.43 7.43 7.29 5.29 1.00
DG 0.11 0.11 0.18 0.15 0.18 0.15 0.24 0.29 0.36 1.00
CA3 0.20 0.23 0.26 0.21 0.21 0.25 0.29 0.31 0.48 1.00
CA1 0.10 0.08 0.11 0.10 0.10 0.11 0.16 0.21 0.26 1.00
Amy−Exc 0.15 0.12 0.14 0.14 0.15 0.19 0.22 0.25 0.31 1.00
Vip 0.31 0.23 0.28 0.30 0.32 0.34 0.37 0.36 0.50 1.00
THM−MB 0.35 0.25 0.19 0.24 0.36 0.34 0.39 0.42 0.54 1.00
THM−Inh 10.46 5.42 4.08 3.17 3.29 2.67 2.33 2.04 2.17 1.00
THM−Exc 0.59 0.57 0.57 0.54 0.50 0.53 0.51 0.54 0.56 1.00
SubCtx−Cplx 0.16 0.22 0.19 0.17 0.20 0.26 0.25 0.30 0.40 1.00
Sst 0.29 0.21 0.23 0.26 0.25 0.24 0.30 0.28 0.38 1.00
Sncg 0.18 0.14 0.14 0.17 0.14 0.20 0.27 0.28 0.38 1.00
Pvalb−ChC 0.19 0.19 0.20 0.21 0.26 0.23 0.22 0.30 0.38 1.00
Pvalb 0.14 0.29 0.37 0.45 0.19 0.13 0.14 0.19 0.25 1.00
PN 0.25 0.22 0.19 0.16 0.19 0.20 0.25 0.24 0.36 1.00
MSN−D2 0.16 0.09 0.12 0.15 0.17 0.18 0.22 0.26 0.30 1.00
MSN−D1 0.13 0.15 0.18 0.15 0.16 0.20 0.23 0.27 0.32 1.00
Lamp5−Lhx6 0.33 0.32 0.27 0.35 0.39 0.37 0.35 0.43 0.54 1.00
Lamp5 0.24 0.24 0.22 0.27 0.26 0.30 0.25 0.37 0.47 1.00
Foxp2 0.18 0.17 0.17 0.18 0.18 0.22 0.24 0.26 0.34 1.00
Chd7 0.12 0.09 0.09 0.10 0.11 0.14 0.16 0.19 0.24 1.00
CB 7.79 5.08 5.31 4.52 4.43 4.20 3.56 2.92 1.89 1.00
0.0−0.1 0.1−0.2 0.2−0.3 0.3−0.4 0.4−0.5 0.5−0.6 0.6−0.7 0.7−0.8 0.8−0.9 0.9−1.0
Percent
0.0003 0.0006 0.0009 0.0012

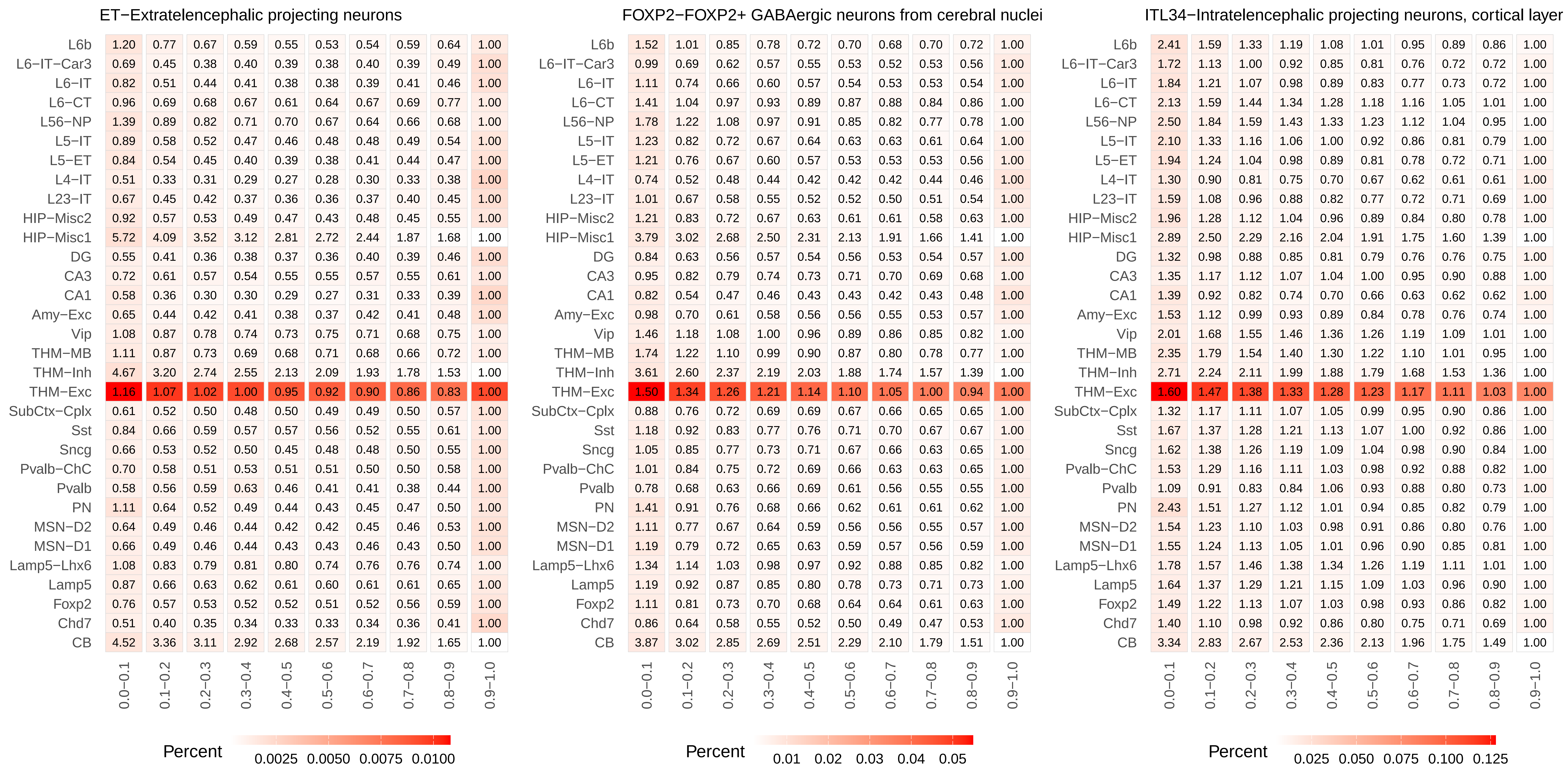
ET−Extratelencephalic projecting neurons
L6b 1.20 0.77 0.67 0.59 0.55 0.53 0.54 0.59 0.64 1.00
L6−IT−Car3 0.69 0.45 0.38 0.40 0.39 0.38 0.40 0.39 0.49 1.00
L6−IT 0.82 0.51 0.44 0.41 0.38 0.38 0.39 0.41 0.46 1.00
L6−CT 0.96 0.69 0.68 0.67 0.61 0.64 0.67 0.69 0.77 1.00
L56−NP 1.39 0.89 0.82 0.71 0.70 0.67 0.64 0.66 0.68 1.00
L5−IT 0.89 0.58 0.52 0.47 0.46 0.48 0.48 0.49 0.54 1.00
L5−ET 0.84 0.54 0.45 0.40 0.39 0.38 0.41 0.44 0.47 1.00
L4−IT 0.51 0.33 0.31 0.29 0.27 0.28 0.30 0.33 0.38 1.00
L23−IT 0.67 0.45 0.42 0.37 0.36 0.36 0.37 0.40 0.45 1.00
HIP−Misc2 0.92 0.57 0.53 0.49 0.47 0.43 0.48 0.45 0.55 1.00
HIP−Misc1 5.72 4.09 3.52 3.12 2.81 2.72 2.44 1.87 1.68 1.00
DG 0.55 0.41 0.36 0.38 0.37 0.36 0.40 0.39 0.46 1.00
CA3 0.72 0.61 0.57 0.54 0.55 0.55 0.57 0.55 0.61 1.00
CA1 0.58 0.36 0.30 0.30 0.29 0.27 0.31 0.33 0.39 1.00
Amy−Exc 0.65 0.44 0.42 0.41 0.38 0.37 0.42 0.41 0.48 1.00
Vip 1.08 0.87 0.78 0.74 0.73 0.75 0.71 0.68 0.75 1.00
THM−MB 1.11 0.87 0.73 0.69 0.68 0.71 0.68 0.66 0.72 1.00
THM−Inh 4.67 3.20 2.74 2.55 2.13 2.09 1.93 1.78 1.53 1.00
THM−Exc 1.16 1.07 1.02 1.00 0.95 0.92 0.90 0.86 0.83 1.00
SubCtx−Cplx 0.61 0.52 0.50 0.48 0.50 0.49 0.49 0.50 0.57 1.00
Sst 0.84 0.66 0.59 0.57 0.57 0.56 0.52 0.55 0.61 1.00
Sncg 0.66 0.53 0.52 0.50 0.45 0.48 0.48 0.50 0.55 1.00
Pvalb−ChC 0.70 0.58 0.51 0.53 0.51 0.51 0.50 0.50 0.58 1.00
Pvalb 0.58 0.56 0.59 0.63 0.46 0.41 0.41 0.38 0.44 1.00
PN 1.11 0.64 0.52 0.49 0.44 0.43 0.45 0.47 0.50 1.00
MSN−D2 0.64 0.49 0.46 0.44 0.42 0.42 0.45 0.46 0.53 1.00
MSN−D1 0.66 0.49 0.46 0.44 0.43 0.43 0.46 0.43 0.50 1.00
Lamp5−Lhx6 1.08 0.83 0.79 0.81 0.80 0.74 0.76 0.76 0.74 1.00
Lamp5 0.87 0.66 0.63 0.62 0.61 0.60 0.61 0.61 0.65 1.00
Foxp2 0.76 0.57 0.53 0.52 0.52 0.51 0.52 0.56 0.59 1.00
Chd7 0.51 0.40 0.35 0.34 0.33 0.33 0.34 0.36 0.41 1.00
CB 4.52 3.36 3.11 2.92 2.68 2.57 2.19 1.92 1.65 1.00
0.0−0.1 0.1−0.2 0.2−0.3 0.3−0.4 0.4−0.5 0.5−0.6 0.6−0.7 0.7−0.8 0.8−0.9 0.9−1.0
Percent 0.0025 0.0050 0.0075 0.0100
FOXP2−FOXP2+ GABAergic neurons from cerebral nuclei
L6b 1.52 1.01 0.85 0.78 0.72 0.70 0.68 0.70 0.72 1.00
L6−IT−Car3 0.99 0.69 0.62 0.57 0.55 0.53 0.52 0.53 0.56 1.00
L6−IT 1.11 0.74 0.66 0.60 0.57 0.54 0.53 0.53 0.54 1.00
L6−CT 1.41 1.04 0.97 0.93 0.89 0.87 0.88 0.84 0.86 1.00
L56−NP 1.78 1.22 1.08 0.97 0.91 0.85 0.82 0.77 0.78 1.00
L5−IT 1.23 0.82 0.72 0.67 0.64 0.63 0.63 0.61 0.64 1.00
L5−ET 1.21 0.76 0.67 0.60 0.57 0.53 0.53 0.53 0.56 1.00
L4−IT 0.74 0.52 0.48 0.44 0.42 0.42 0.42 0.44 0.46 1.00
L23−IT 1.01 0.67 0.58 0.55 0.52 0.52 0.50 0.51 0.54 1.00
HIP−Misc2 1.21 0.83 0.72 0.67 0.63 0.61 0.61 0.58 0.63 1.00
HIP−Misc1 3.79 3.02 2.68 2.50 2.31 2.13 1.91 1.66 1.41 1.00
DG 0.84 0.63 0.56 0.57 0.54 0.56 0.53 0.54 0.57 1.00
CA3 0.95 0.82 0.79 0.74 0.73 0.71 0.70 0.69 0.68 1.00
CA1 0.82 0.54 0.47 0.46 0.43 0.43 0.42 0.43 0.48 1.00
Amy−Exc 0.98 0.70 0.61 0.58 0.56 0.56 0.55 0.53 0.57 1.00
Vip 1.46 1.18 1.08 1.00 0.96 0.89 0.86 0.85 0.82 1.00
THM−MB 1.74 1.22 1.10 0.99 0.90 0.87 0.80 0.78 0.77 1.00
THM−Inh 3.61 2.60 2.37 2.19 2.03 1.88 1.74 1.57 1.39 1.00
THM−Exc 1.50 1.34 1.26 1.21 1.14 1.10 1.05 1.00 0.94 1.00
SubCtx−Cplx 0.88 0.76 0.72 0.69 0.69 0.67 0.66 0.65 0.65 1.00
Sst 1.18 0.92 0.83 0.77 0.76 0.71 0.70 0.67 0.67 1.00
Sncg 1.05 0.85 0.77 0.73 0.71 0.67 0.66 0.63 0.65 1.00
Pvalb−ChC 1.01 0.84 0.75 0.72 0.69 0.66 0.63 0.63 0.65 1.00
Pvalb 0.78 0.68 0.63 0.66 0.69 0.61 0.56 0.55 0.55 1.00
PN 1.41 0.91 0.76 0.68 0.66 0.62 0.61 0.61 0.62 1.00
MSN−D2 1.11 0.77 0.67 0.64 0.59 0.56 0.56 0.55 0.57 1.00
MSN−D1 1.19 0.79 0.72 0.65 0.63 0.59 0.57 0.56 0.59 1.00
Lamp5−Lhx6 1.34 1.14 1.03 0.98 0.97 0.92 0.88 0.85 0.82 1.00
Lamp5 1.19 0.92 0.87 0.85 0.80 0.78 0.73 0.71 0.73 1.00
Foxp2 1.11 0.81 0.73 0.70 0.68 0.64 0.64 0.61 0.63 1.00
Chd7 0.86 0.64 0.58 0.55 0.52 0.50 0.49 0.47 0.53 1.00
CB 3.87 3.02 2.85 2.69 2.51 2.29 2.10 1.79 1.51 1.00
0.0−0.1 0.1−0.2 0.2−0.3 0.3−0.4 0.4−0.5 0.5−0.6 0.6−0.7 0.7−0.8 0.8−0.9 0.9−1.0
Percent 0.01 0.02 0.03 0.04 0.05
ITL34−Intratelencephalic projecting neurons, cortical layer
L6b 2.41 1.59 1.33 1.19 1.08 1.01 0.95 0.89 0.86 1.00
L6−IT−Car3 1.72 1.13 1.00 0.92 0.85 0.81 0.76 0.72 0.72 1.00
L6−IT 1.84 1.21 1.07 0.98 0.89 0.83 0.77 0.73 0.72 1.00
L6−CT 2.13 1.59 1.44 1.34 1.28 1.18 1.16 1.05 1.01 1.00
L56−NP 2.50 1.84 1.59 1.43 1.33 1.23 1.12 1.04 0.95 1.00
L5−IT 2.10 1.33 1.16 1.06 1.00 0.92 0.86 0.81 0.79 1.00
L5−ET 1.94 1.24 1.04 0.98 0.89 0.81 0.78 0.72 0.71 1.00
L4−IT 1.30 0.90 0.81 0.75 0.70 0.67 0.62 0.61 0.61 1.00
L23−IT 1.59 1.08 0.96 0.88 0.82 0.77 0.72 0.71 0.69 1.00
HIP−Misc2 1.96 1.28 1.12 1.04 0.96 0.89 0.84 0.80 0.78 1.00
HIP−Misc1 2.89 2.50 2.29 2.16 2.04 1.91 1.75 1.60 1.39 1.00
DG 1.32 0.98 0.88 0.85 0.81 0.79 0.76 0.76 0.75 1.00
CA3 1.35 1.17 1.12 1.07 1.04 1.00 0.95 0.90 0.88 1.00
CA1 1.39 0.92 0.82 0.74 0.70 0.66 0.63 0.62 0.62 1.00
Amy−Exc 1.53 1.12 0.99 0.93 0.89 0.84 0.78 0.76 0.74 1.00
Vip 2.01 1.68 1.55 1.46 1.36 1.26 1.19 1.09 1.01 1.00
THM−MB 2.35 1.79 1.54 1.40 1.30 1.22 1.10 1.01 0.95 1.00
THM−Inh 2.71 2.24 2.11 1.99 1.88 1.79 1.68 1.53 1.36 1.00
THM−Exc 1.60 1.47 1.38 1.33 1.28 1.23 1.17 1.11 1.03 1.00
SubCtx−Cplx 1.32 1.17 1.11 1.07 1.05 0.99 0.95 0.90 0.86 1.00
Sst 1.67 1.37 1.28 1.21 1.13 1.07 1.00 0.92 0.86 1.00
Sncg 1.62 1.38 1.26 1.19 1.09 1.04 0.98 0.90 0.84 1.00
Pvalb−ChC 1.53 1.29 1.16 1.11 1.03 0.98 0.92 0.88 0.82 1.00
Pvalb 1.09 0.91 0.83 0.84 1.06 0.93 0.88 0.80 0.73 1.00
PN 2.43 1.51 1.27 1.12 1.01 0.94 0.85 0.82 0.79 1.00
MSN−D2 1.54 1.23 1.10 1.03 0.98 0.91 0.86 0.80 0.76 1.00
MSN−D1 1.55 1.24 1.13 1.05 1.01 0.96 0.90 0.85 0.81 1.00
Lamp5−Lhx6 1.78 1.57 1.46 1.38 1.34 1.26 1.19 1.11 1.01 1.00
Lamp5 1.64 1.37 1.29 1.21 1.15 1.09 1.03 0.96 0.90 1.00
Foxp2 1.49 1.22 1.13 1.07 1.03 0.98 0.93 0.86 0.82 1.00
Chd7 1.40 1.10 0.98 0.92 0.86 0.80 0.75 0.71 0.69 1.00
CB 3.34 2.83 2.67 2.53 2.36 2.13 1.96 1.75 1.49 1.00
0.0−0.1 0.1−0.2 0.2−0.3 0.3−0.4 0.4−0.5 0.5−0.6 0.6−0.7 0.7−0.8 0.8−0.9 0.9−1.0
Percent 0.025 0.050 0.075 0.100 0.125

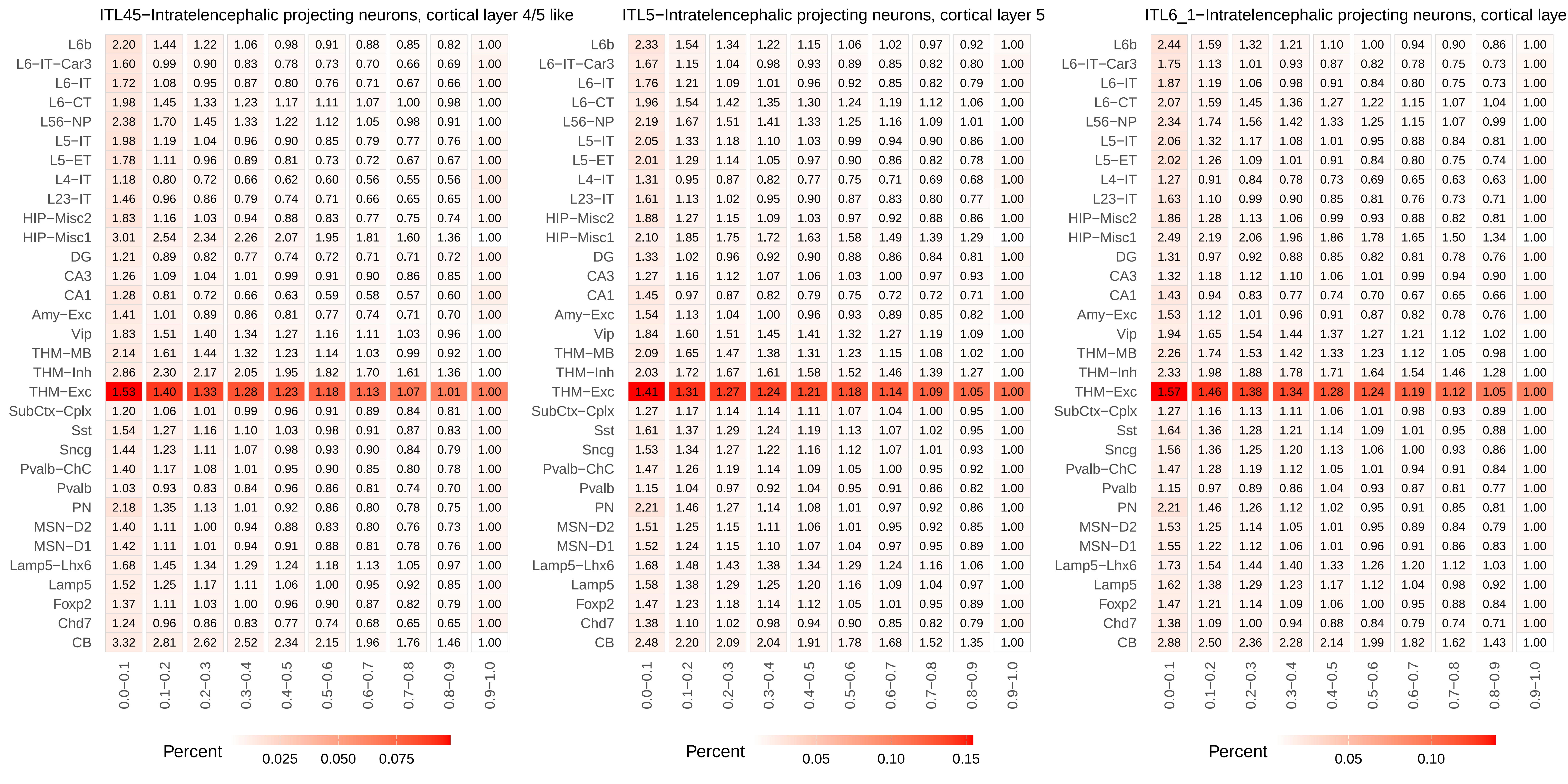

ITL45−Intratelencephalic projecting neurons, cortical layer 4/5 like
ITL5−Intratelencephalic projecting neurons, cortical layer 5
ITL6_1−Intratelencephalic projecting neurons, cortical laye
L6b
L6−IT−Car3
L6−IT
L6−CT
L56−NP
L5−IT
L5−ET
L4−IT
L23−IT
HIP−Misc2
HIP−Misc1
DG
CA3
CA1
Amy−Exc
Vip
THM−MB
THM−Inh
THM−Exc
SubCtx−Cplx
Sst
Sncg
Pvalb−ChC
Pvalb
PN
MSN−D2
MSN−D1
Lamp5−Lhx6
Lamp5
Foxp2
Chd7
CB
0.0−0.1
0.1−0.2
0.2−0.3
0.3−0.4
0.4−0.5
0.5−0.6
0.6−0.7
0.7−0.8
0.8−0.9
0.9−1.0
Percent
0.025
0.050
0.075
Percent
0.05
0.10
0.15
Percent
0.05
0.10

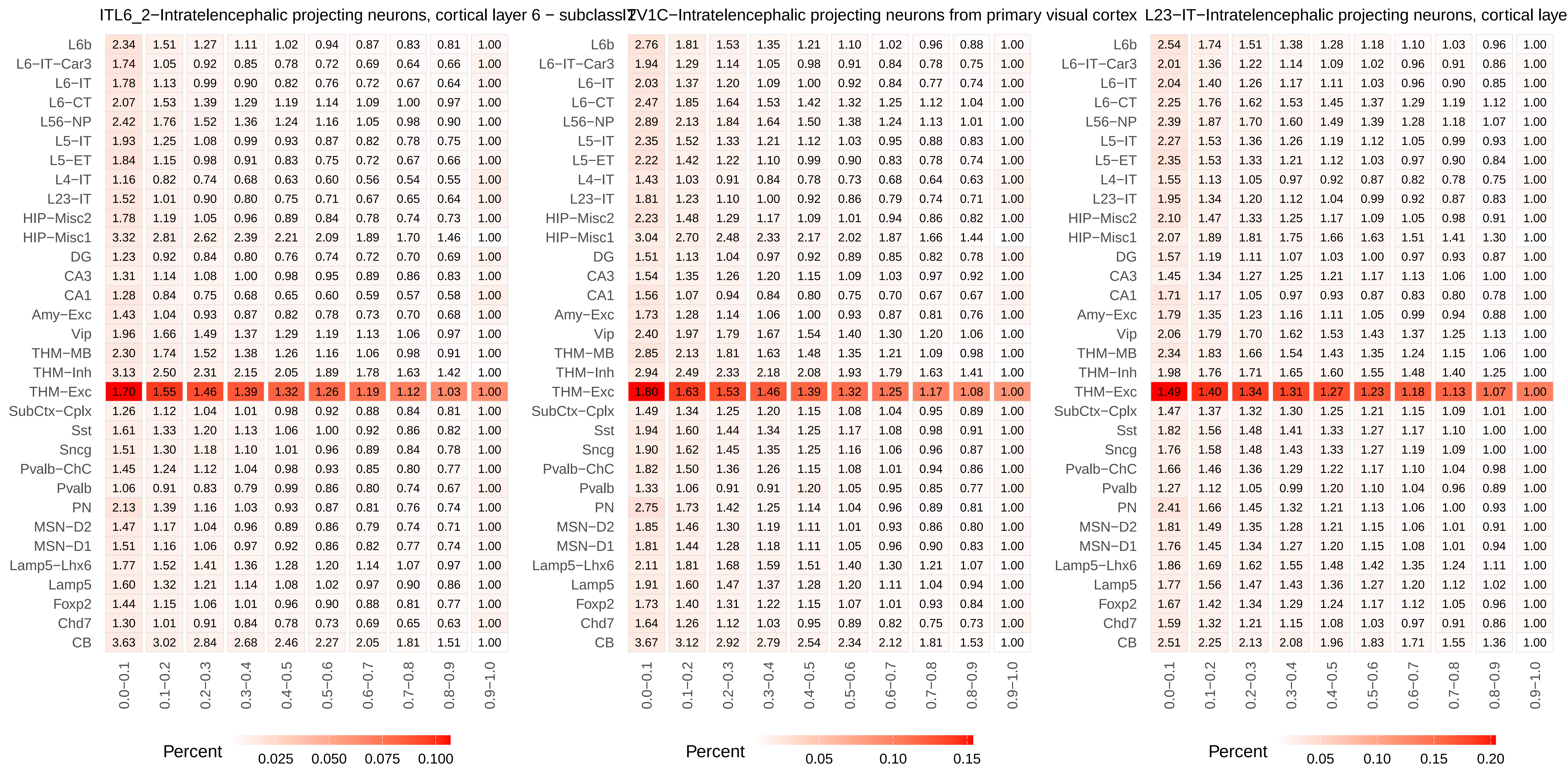
ITL6_2−Intratelencephalic projecting neurons, cortical layer 6 − subclassI2V1C−Intratelencephalic projecting neurons from primary visual cortex
L23−IT−Intratelencephalic projecting neurons, cortical laye
L6b 2.34 1.51 1.27 1.11 1.02 0.94 0.87 0.83 0.81 1.00
L6−IT−Car3 1.74 1.05 0.92 0.85 0.78 0.72 0.69 0.64 0.66 1.00
L6−IT 1.78 1.13 0.99 0.90 0.82 0.76 0.72 0.67 0.64 1.00
L6−CT 2.07 1.53 1.39 1.29 1.19 1.14 1.09 1.00 0.97 1.00
L56−NP 2.42 1.76 1.52 1.36 1.24 1.16 1.05 0.98 0.90 1.00
L5−IT 1.93 1.25 1.08 0.99 0.93 0.87 0.82 0.78 0.75 1.00
L5−ET 1.84 1.15 0.98 0.91 0.83 0.75 0.72 0.67 0.66 1.00
L4−IT 1.16 0.82 0.74 0.68 0.63 0.60 0.56 0.54 0.55 1.00
L23−IT 1.52 1.01 0.90 0.80 0.75 0.71 0.67 0.65 0.64 1.00
HIP−Misc2 1.78 1.19 1.05 0.96 0.89 0.84 0.78 0.74 0.73 1.00
HIP−Misc1 3.32 2.81 2.62 2.39 2.21 2.09 1.89 1.70 1.46 1.00
DG 1.23 0.92 0.84 0.80 0.76 0.74 0.72 0.70 0.69 1.00
CA3 1.31 1.14 1.08 1.00 0.98 0.95 0.89 0.86 0.83 1.00
CA1 1.28 0.84 0.75 0.68 0.65 0.60 0.59 0.57 0.58 1.00
Amy−Exc 1.43 1.04 0.93 0.87 0.82 0.78 0.73 0.70 0.68 1.00
Vip 1.96 1.66 1.49 1.37 1.29 1.19 1.13 1.06 0.97 1.00
THM−MB 2.30 1.74 1.52 1.38 1.26 1.16 1.06 0.98 0.91 1.00
THM−Inh 3.13 2.50 2.31 2.15 2.05 1.89 1.78 1.63 1.42 1.00
THM−Exc 1.70 1.55 1.46 1.39 1.32 1.26 1.19 1.12 1.03 1.00
SubCtx−Cplx 1.26 1.12 1.04 1.01 0.98 0.92 0.88 0.84 0.81 1.00
Sst 1.61 1.33 1.20 1.13 1.06 1.00 0.92 0.86 0.82 1.00
Sncg 1.51 1.30 1.18 1.10 1.01 0.96 0.89 0.84 0.78 1.00
Pvalb−ChC 1.45 1.24 1.12 1.04 0.98 0.93 0.85 0.80 0.77 1.00
Pvalb 1.06 0.91 0.83 0.79 0.99 0.86 0.80 0.74 0.67 1.00
PN 2.13 1.39 1.16 1.03 0.93 0.87 0.81 0.76 0.74 1.00
MSN−D2 1.47 1.17 1.04 0.96 0.89 0.86 0.79 0.74 0.71 1.00
MSN−D1 1.51 1.16 1.06 0.97 0.92 0.86 0.82 0.77 0.74 1.00
Lamp5−Lhx6 1.77 1.52 1.41 1.36 1.28 1.20 1.14 1.07 0.97 1.00
Lamp5 1.60 1.32 1.21 1.14 1.08 1.02 0.97 0.90 0.86 1.00
Foxp2 1.44 1.15 1.06 1.01 0.96 0.90 0.88 0.81 0.77 1.00
Chd7 1.30 1.01 0.91 0.84 0.78 0.73 0.69 0.65 0.63 1.00
CB 3.63 3.02 2.84 2.68 2.46 2.27 2.05 1.81 1.51 1.00
0.0−0.1 0.1−0.2 0.2−0.3 0.3−0.4 0.4−0.5 0.5−0.6 0.6−0.7 0.7−0.8 0.8−0.9 0.9−1.0
Percent
0.025 0.050 0.075 0.100
L6b 2.76 1.81 1.53 1.35 1.21 1.10 1.02 0.96 0.88 1.00
L6−IT−Car3 1.94 1.29 1.14 1.05 0.98 0.91 0.84 0.78 0.75 1.00
L6−IT 2.03 1.37 1.20 1.09 1.00 0.92 0.84 0.77 0.74 1.00
L6−CT 2.47 1.85 1.64 1.53 1.42 1.32 1.25 1.12 1.04 1.00
L56−NP 2.89 2.13 1.84 1.64 1.50 1.38 1.24 1.13 1.01 1.00
L5−IT 2.35 1.52 1.33 1.21 1.12 1.03 0.95 0.88 0.83 1.00
L5−ET 2.22 1.42 1.22 1.10 0.99 0.90 0.83 0.78 0.74 1.00
L4−IT 1.43 1.03 0.91 0.84 0.78 0.73 0.68 0.64 0.63 1.00
L23−IT 1.81 1.23 1.10 1.00 0.92 0.86 0.79 0.74 0.71 1.00
HIP−Misc2 2.23 1.48 1.29 1.17 1.09 1.01 0.94 0.86 0.82 1.00
HIP−Misc1 3.04 2.70 2.48 2.33 2.17 2.02 1.87 1.66 1.44 1.00
DG 1.51 1.13 1.04 0.97 0.92 0.89 0.85 0.82 0.78 1.00
CA3 1.54 1.35 1.26 1.20 1.15 1.09 1.03 0.97 0.92 1.00
CA1 1.56 1.07 0.94 0.84 0.80 0.75 0.70 0.67 0.67 1.00
Amy−Exc 1.73 1.28 1.14 1.06 1.00 0.93 0.87 0.81 0.76 1.00
Vip 2.40 1.97 1.79 1.67 1.54 1.40 1.30 1.20 1.06 1.00
THM−MB 2.85 2.13 1.81 1.63 1.48 1.35 1.21 1.09 0.98 1.00
THM−Inh 2.94 2.49 2.33 2.18 2.08 1.93 1.79 1.63 1.41 1.00
THM−Exc 1.80 1.63 1.53 1.46 1.39 1.32 1.25 1.17 1.08 1.00
SubCtx−Cplx 1.49 1.34 1.25 1.20 1.15 1.08 1.04 0.95 0.89 1.00
Sst 1.94 1.60 1.44 1.34 1.25 1.17 1.08 0.98 0.91 1.00
Sncg 1.90 1.62 1.45 1.35 1.25 1.16 1.06 0.96 0.87 1.00
Pvalb−ChC 1.82 1.50 1.36 1.26 1.15 1.08 1.01 0.94 0.86 1.00
Pvalb 1.33 1.06 0.91 0.91 1.20 1.05 0.95 0.85 0.77 1.00
PN 2.75 1.73 1.42 1.25 1.14 1.04 0.96 0.89 0.81 1.00
MSN−D2 1.85 1.46 1.30 1.19 1.11 1.01 0.93 0.86 0.80 1.00
MSN−D1 1.81 1.44 1.28 1.18 1.11 1.05 0.96 0.90 0.83 1.00
Lamp5−Lhx6 2.11 1.81 1.68 1.59 1.51 1.40 1.30 1.21 1.07 1.00
Lamp5 1.91 1.60 1.47 1.37 1.28 1.20 1.11 1.04 0.94 1.00
Foxp2 1.73 1.40 1.31 1.22 1.15 1.07 1.01 0.93 0.84 1.00
Chd7 1.64 1.26 1.12 1.03 0.95 0.89 0.82 0.75 0.73 1.00
CB 3.67 3.12 2.92 2.79 2.54 2.34 2.12 1.81 1.53 1.00
0.0−0.1 0.1−0.2 0.2−0.3 0.3−0.4 0.4−0.5 0.5−0.6 0.6−0.7 0.7−0.8 0.8−0.9 0.9−1.0
Percent
0.05 0.10 0.15
L6b 2.54 1.74 1.51 1.38 1.28 1.18 1.10 1.03 0.96 1.00
L6−IT−Car3 2.01 1.36 1.22 1.14 1.09 1.02 0.96 0.91 0.86 1.00
L6−IT 2.04 1.40 1.26 1.17 1.11 1.03 0.96 0.90 0.85 1.00
L6−CT 2.25 1.76 1.62 1.53 1.45 1.37 1.29 1.19 1.12 1.00
L56−NP 2.39 1.87 1.70 1.60 1.49 1.39 1.28 1.18 1.07 1.00
L5−IT 2.27 1.53 1.36 1.26 1.19 1.12 1.05 0.99 0.93 1.00
L5−ET 2.35 1.53 1.33 1.21 1.12 1.03 0.97 0.90 0.84 1.00
L4−IT 1.55 1.13 1.05 0.97 0.92 0.87 0.82 0.78 0.75 1.00
L23−IT 1.95 1.34 1.20 1.12 1.04 0.99 0.92 0.87 0.83 1.00
HIP−Misc2 2.10 1.47 1.33 1.25 1.17 1.09 1.05 0.98 0.91 1.00
HIP−Misc1 2.07 1.89 1.81 1.75 1.66 1.63 1.51 1.41 1.30 1.00
DG 1.57 1.19 1.11 1.07 1.03 1.00 0.97 0.93 0.87 1.00
CA3 1.45 1.34 1.27 1.25 1.21 1.17 1.13 1.06 1.00 1.00
CA1 1.71 1.17 1.05 0.97 0.93 0.87 0.83 0.80 0.78 1.00
Amy−Exc 1.79 1.35 1.23 1.16 1.11 1.05 0.99 0.94 0.88 1.00
Vip 2.06 1.79 1.70 1.62 1.53 1.43 1.37 1.25 1.13 1.00
THM−MB 2.34 1.83 1.66 1.54 1.43 1.35 1.24 1.15 1.06 1.00
THM−Inh 1.98 1.76 1.71 1.65 1.60 1.55 1.48 1.40 1.25 1.00
THM−Exc 1.49 1.40 1.34 1.31 1.27 1.23 1.18 1.13 1.07 1.00
SubCtx−Cplx 1.47 1.37 1.32 1.30 1.25 1.21 1.15 1.09 1.01 1.00
Sst 1.82 1.56 1.48 1.41 1.33 1.27 1.17 1.10 1.00 1.00
Sncg 1.76 1.58 1.48 1.43 1.33 1.27 1.19 1.09 1.00 1.00
Pvalb−ChC 1.66 1.46 1.36 1.29 1.22 1.17 1.10 1.04 0.98 1.00
Pvalb 1.27 1.12 1.05 0.99 1.20 1.10 1.04 0.96 0.89 1.00
PN 2.41 1.66 1.45 1.32 1.21 1.13 1.06 1.00 0.93 1.00
MSN−D2 1.81 1.49 1.35 1.28 1.21 1.15 1.06 1.01 0.91 1.00
MSN−D1 1.76 1.45 1.34 1.27 1.20 1.15 1.08 1.01 0.94 1.00
Lamp5−Lhx6 1.86 1.69 1.62 1.55 1.48 1.42 1.35 1.24 1.11 1.00
Lamp5 1.77 1.56 1.47 1.43 1.36 1.27 1.20 1.12 1.02 1.00
Foxp2 1.67 1.42 1.34 1.29 1.24 1.17 1.12 1.05 0.96 1.00
Chd7 1.59 1.32 1.21 1.15 1.08 1.03 0.97 0.91 0.86 1.00
CB 2.51 2.25 2.13 2.08 1.96 1.83 1.71 1.55 1.36 1.00
0.0−0.1 0.1−0.2 0.2−0.3 0.3−0.4 0.4−0.5 0.5−0.6 0.6−0.7 0.7−0.8 0.8−0.9 0.9−1.0
Percent
0.05 0.10 0.15 0.20

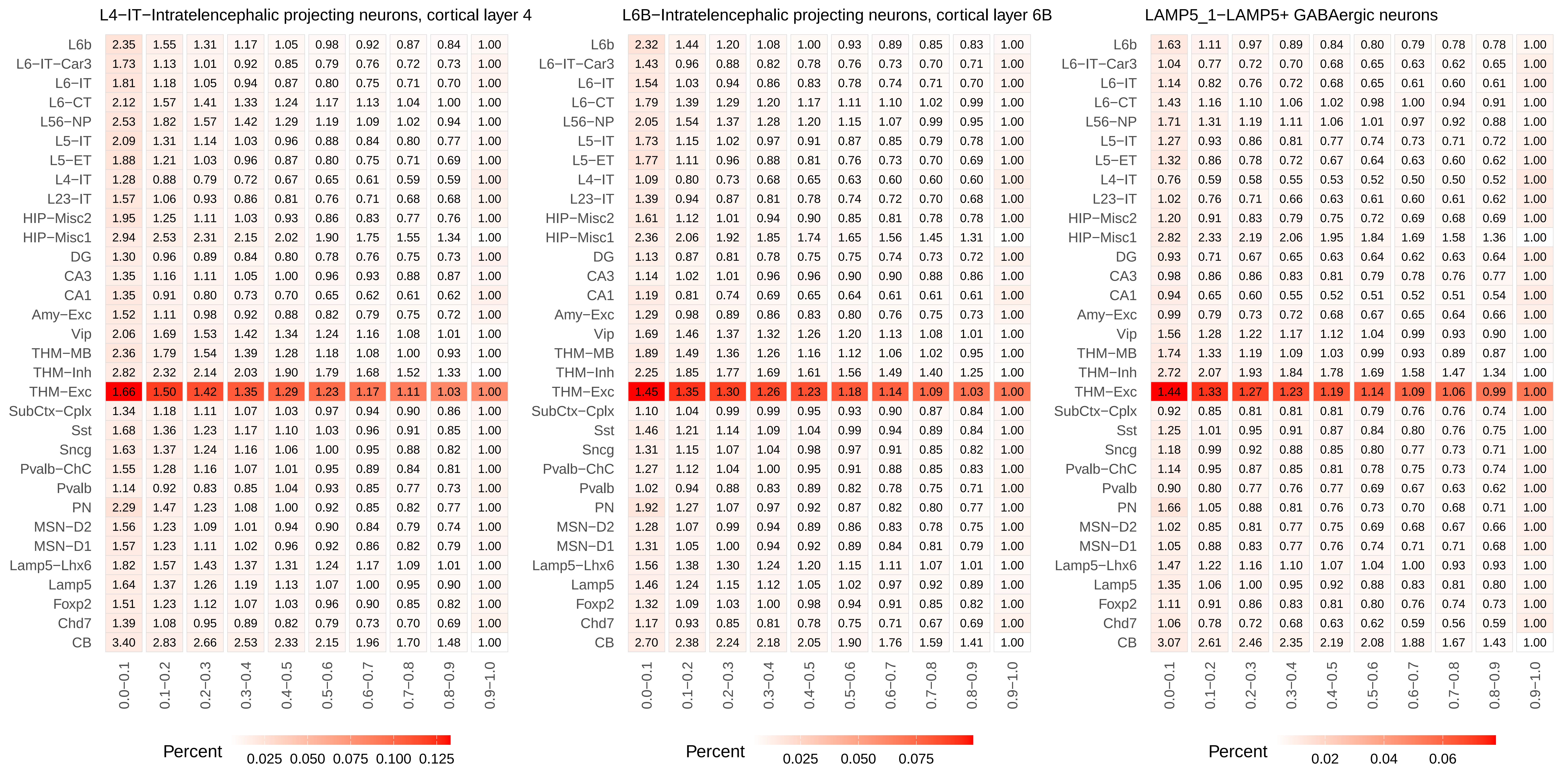

L4−IT−Intratelencephalic projecting neurons, cortical layer 4
L6B−Intratelencephalic projecting neurons, cortical layer 6B
LAMP5_1−LAMP5+ GABAergic neurons
L6b
L6−IT−Car3
L6−IT
L6−CT
L56−NP
L5−IT
L5−ET
L4−IT
L23−IT
HIP−Misc2
HIP−Misc1
DG
CA3
CA1
Amy−Exc
Vip
THM−MB
THM−Inh
THM−Exc
SubCtx−Cplx
Sst
Sncg
Pvalb−ChC
Pvalb
PN
MSN−D2
MSN−D1
Lamp5−Lhx6
Lamp5
Foxp2
Chd7
CB
0.0−0.1
0.1−0.2
0.2−0.3
0.3−0.4
0.4−0.5
0.5−0.6
0.6−0.7
0.7−0.8
0.8−0.9
0.9−1.0
Percent
0.025 0.050 0.075 0.100 0.125
Percent
0.025 0.050 0.075
Percent
0.02 0.04 0.06

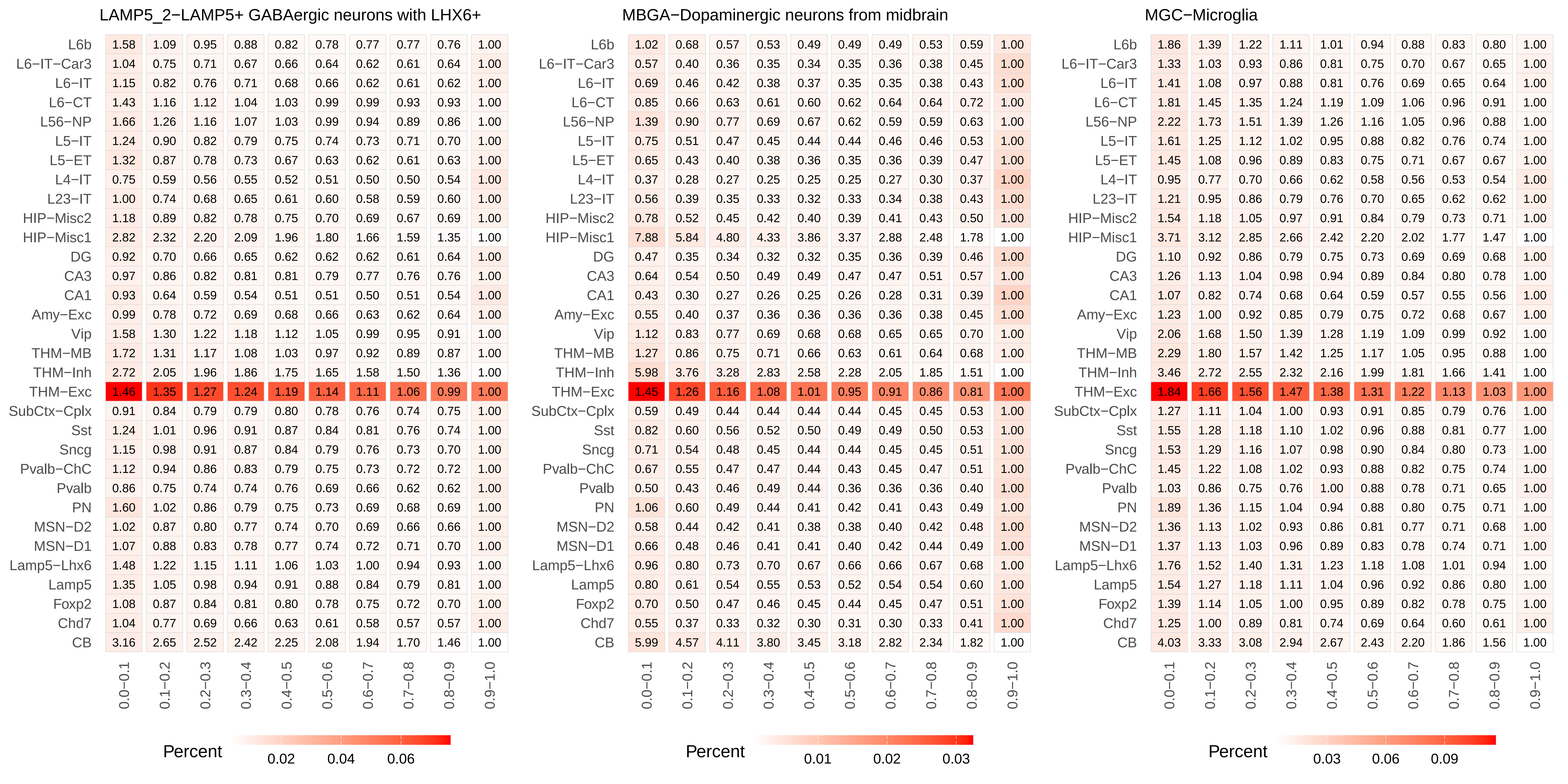
LAMP5_2−LAMP5+ GABAergic neurons with LHX6+
L6b 1.58 1.09 0.95 0.88 0.82 0.78 0.77 0.77 0.76 1.00
L6−IT−Car3 1.04 0.75 0.71 0.67 0.66 0.64 0.62 0.61 0.64 1.00
L6−IT 1.15 0.82 0.76 0.71 0.68 0.66 0.62 0.61 0.62 1.00
L6−CT 1.43 1.16 1.12 1.04 1.03 0.99 0.99 0.93 0.93 1.00
L56−NP 1.66 1.26 1.16 1.07 1.03 0.99 0.94 0.89 0.86 1.00
L5−IT 1.24 0.90 0.82 0.79 0.75 0.74 0.73 0.71 0.70 1.00
L5−ET 1.32 0.87 0.78 0.73 0.67 0.63 0.62 0.61 0.63 1.00
L4−IT 0.75 0.59 0.56 0.55 0.52 0.51 0.50 0.50 0.54 1.00
L23−IT 1.00 0.74 0.68 0.65 0.61 0.60 0.58 0.59 0.60 1.00
HIP−Misc2 1.18 0.89 0.82 0.78 0.75 0.70 0.69 0.67 0.69 1.00
HIP−Misc1 2.82 2.32 2.20 2.09 1.96 1.80 1.66 1.59 1.35 1.00
DG 0.92 0.70 0.66 0.65 0.62 0.62 0.62 0.61 0.64 1.00
CA3 0.97 0.86 0.82 0.81 0.81 0.79 0.77 0.76 0.76 1.00
CA1 0.93 0.64 0.59 0.54 0.51 0.51 0.50 0.51 0.54 1.00
Amy−Exc 0.99 0.78 0.72 0.69 0.68 0.66 0.63 0.62 0.64 1.00
Vip 1.58 1.30 1.22 1.18 1.12 1.05 0.99 0.95 0.91 1.00
THM−MB 1.72 1.31 1.17 1.08 1.03 0.97 0.92 0.89 0.87 1.00
THM−Inh 2.72 2.05 1.96 1.86 1.75 1.65 1.58 1.50 1.36 1.00
THM−Exc 1.46 1.35 1.27 1.24 1.19 1.14 1.11 1.06 0.99 1.00
SubCtx−Cplx 0.91 0.84 0.79 0.79 0.80 0.78 0.76 0.74 0.75 1.00
Sst 1.24 1.01 0.96 0.91 0.87 0.84 0.81 0.76 0.74 1.00
Sncg 1.15 0.98 0.91 0.87 0.84 0.79 0.76 0.73 0.70 1.00
Pvalb−ChC 1.12 0.94 0.86 0.83 0.79 0.75 0.73 0.72 0.72 1.00
Pvalb 0.86 0.75 0.74 0.74 0.76 0.69 0.66 0.62 0.62 1.00
PN 1.60 1.02 0.86 0.79 0.75 0.73 0.69 0.68 0.69 1.00
MSN−D2 1.02 0.87 0.80 0.77 0.74 0.70 0.69 0.66 0.66 1.00
MSN−D1 1.07 0.88 0.83 0.78 0.77 0.74 0.72 0.71 0.70 1.00
Lamp5−Lhx6 1.48 1.22 1.15 1.11 1.06 1.03 1.00 0.94 0.93 1.00
Lamp5 1.35 1.05 0.98 0.94 0.91 0.88 0.84 0.79 0.81 1.00
Foxp2 1.08 0.87 0.84 0.81 0.80 0.78 0.75 0.72 0.70 1.00
Chd7 1.04 0.77 0.69 0.66 0.63 0.61 0.58 0.57 0.57 1.00
CB 3.16 2.65 2.52 2.42 2.25 2.08 1.94 1.70 1.46 1.00
0.0−0.1 0.1−0.2 0.2−0.3 0.3−0.4 0.4−0.5 0.5−0.6 0.6−0.7 0.7−0.8 0.8−0.9 0.9−1.0
Percent 0.02 0.04 0.06
MBGA−Dopaminergic neurons from midbrain
L6b 1.02 0.68 0.57 0.53 0.49 0.49 0.49 0.53 0.59 1.00
L6−IT−Car3 0.57 0.40 0.36 0.35 0.34 0.35 0.36 0.38 0.45 1.00
L6−IT 0.69 0.46 0.42 0.38 0.37 0.35 0.35 0.38 0.43 1.00
L6−CT 0.85 0.66 0.63 0.61 0.60 0.62 0.64 0.64 0.72 1.00
L56−NP 1.39 0.90 0.77 0.69 0.67 0.62 0.59 0.59 0.63 1.00
L5−IT 0.75 0.51 0.47 0.45 0.44 0.44 0.46 0.46 0.53 1.00
L5−ET 0.65 0.43 0.40 0.38 0.36 0.35 0.36 0.39 0.47 1.00
L4−IT 0.37 0.28 0.27 0.25 0.25 0.25 0.27 0.30 0.37 1.00
L23−IT 0.56 0.39 0.35 0.33 0.32 0.33 0.34 0.38 0.43 1.00
HIP−Misc2 0.78 0.52 0.45 0.42 0.40 0.39 0.41 0.43 0.50 1.00
HIP−Misc1 7.88 5.84 4.80 4.33 3.86 3.37 2.88 2.48 1.78 1.00
DG 0.47 0.35 0.34 0.32 0.32 0.35 0.36 0.39 0.46 1.00
CA3 0.64 0.54 0.50 0.49 0.49 0.47 0.47 0.51 0.57 1.00
CA1 0.43 0.30 0.27 0.26 0.25 0.26 0.28 0.31 0.39 1.00
Amy−Exc 0.55 0.40 0.37 0.36 0.36 0.36 0.36 0.38 0.45 1.00
Vip 1.12 0.83 0.77 0.69 0.68 0.68 0.65 0.65 0.70 1.00
THM−MB 1.27 0.86 0.75 0.71 0.66 0.63 0.61 0.64 0.68 1.00
THM−Inh 5.98 3.76 3.28 2.83 2.58 2.28 2.05 1.85 1.51 1.00
THM−Exc 1.45 1.26 1.16 1.08 1.01 0.95 0.91 0.86 0.81 1.00
SubCtx−Cplx 0.59 0.49 0.44 0.44 0.44 0.44 0.45 0.45 0.53 1.00
Sst 0.82 0.60 0.56 0.52 0.50 0.49 0.49 0.50 0.53 1.00
Sncg 0.71 0.54 0.48 0.45 0.44 0.44 0.45 0.45 0.51 1.00
Pvalb−ChC 0.67 0.55 0.47 0.47 0.44 0.43 0.45 0.47 0.51 1.00
Pvalb 0.50 0.43 0.46 0.49 0.44 0.36 0.36 0.36 0.40 1.00
PN 1.06 0.60 0.49 0.44 0.41 0.42 0.41 0.43 0.49 1.00
MSN−D2 0.58 0.44 0.42 0.41 0.38 0.38 0.40 0.42 0.48 1.00
MSN−D1 0.66 0.48 0.46 0.41 0.41 0.40 0.42 0.44 0.49 1.00
Lamp5−Lhx6 0.96 0.80 0.73 0.70 0.67 0.66 0.66 0.67 0.68 1.00
Lamp5 0.80 0.61 0.54 0.55 0.53 0.52 0.54 0.54 0.60 1.00
Foxp2 0.70 0.50 0.47 0.46 0.45 0.44 0.45 0.47 0.51 1.00
Chd7 0.55 0.37 0.33 0.32 0.30 0.31 0.30 0.33 0.41 1.00
CB 5.99 4.57 4.11 3.80 3.45 3.18 2.82 2.34 1.82 1.00
0.0−0.1 0.1−0.2 0.2−0.3 0.3−0.4 0.4−0.5 0.5−0.6 0.6−0.7 0.7−0.8 0.8−0.9 0.9−1.0
Percent 0.01 0.02 0.03
MGC−Microglia
L6b 1.86 1.39 1.22 1.11 1.01 0.94 0.88 0.83 0.80 1.00
L6−IT−Car3 1.33 1.03 0.93 0.86 0.81 0.75 0.70 0.67 0.65 1.00
L6−IT 1.41 1.08 0.97 0.88 0.81 0.76 0.69 0.65 0.64 1.00
L6−CT 1.81 1.45 1.35 1.24 1.19 1.09 1.06 0.96 0.91 1.00
L56−NP 2.22 1.73 1.51 1.39 1.26 1.16 1.05 0.96 0.88 1.00
L5−IT 1.61 1.25 1.12 1.02 0.95 0.88 0.82 0.76 0.74 1.00
L5−ET 1.45 1.08 0.96 0.89 0.83 0.75 0.71 0.67 0.67 1.00
L4−IT 0.95 0.77 0.70 0.66 0.62 0.58 0.56 0.53 0.54 1.00
L23−IT 1.21 0.95 0.86 0.79 0.76 0.70 0.65 0.62 0.62 1.00
HIP−Misc2 1.54 1.18 1.05 0.97 0.91 0.84 0.79 0.73 0.71 1.00
HIP−Misc1 3.71 3.12 2.85 2.66 2.42 2.20 2.02 1.77 1.47 1.00
DG 1.10 0.92 0.86 0.79 0.75 0.73 0.69 0.69 0.68 1.00
CA3 1.26 1.13 1.04 0.98 0.94 0.89 0.84 0.80 0.78 1.00
CA1 1.07 0.82 0.74 0.68 0.64 0.59 0.57 0.55 0.56 1.00
Amy−Exc 1.23 1.00 0.92 0.85 0.79 0.75 0.72 0.68 0.67 1.00
Vip 2.06 1.68 1.50 1.39 1.28 1.19 1.09 0.99 0.92 1.00
THM−MB 2.29 1.80 1.57 1.42 1.25 1.17 1.05 0.95 0.88 1.00
THM−Inh 3.46 2.72 2.55 2.32 2.16 1.99 1.81 1.66 1.41 1.00
THM−Exc 1.84 1.66 1.56 1.47 1.38 1.31 1.22 1.13 1.03 1.00
SubCtx−Cplx 1.27 1.11 1.04 1.00 0.93 0.91 0.85 0.79 0.76 1.00
Sst 1.55 1.28 1.18 1.10 1.02 0.96 0.88 0.81 0.77 1.00
Sncg 1.53 1.29 1.16 1.07 0.98 0.90 0.84 0.80 0.73 1.00
Pvalb−ChC 1.45 1.22 1.08 1.02 0.93 0.88 0.82 0.75 0.74 1.00
Pvalb 1.03 0.86 0.75 0.76 1.00 0.88 0.78 0.71 0.65 1.00
PN 1.89 1.36 1.15 1.04 0.94 0.88 0.80 0.75 0.71 1.00
MSN−D2 1.36 1.13 1.02 0.93 0.86 0.81 0.77 0.71 0.68 1.00
MSN−D1 1.37 1.13 1.03 0.96 0.89 0.83 0.78 0.74 0.71 1.00
Lamp5−Lhx6 1.76 1.52 1.40 1.31 1.23 1.18 1.08 1.01 0.94 1.00
Lamp5 1.54 1.27 1.18 1.11 1.04 0.96 0.92 0.86 0.80 1.00
Foxp2 1.39 1.14 1.05 1.00 0.95 0.89 0.82 0.78 0.75 1.00
Chd7 1.25 1.00 0.89 0.81 0.74 0.69 0.64 0.60 0.61 1.00
CB 4.03 3.33 3.08 2.94 2.67 2.43 2.20 1.86 1.56 1.00
0.0−0.1 0.1−0.2 0.2−0.3 0.3−0.4 0.4−0.5 0.5−0.6 0.6−0.7 0.7−0.8 0.8−0.9 0.9−1.0
Percent 0.03 0.06 0.09

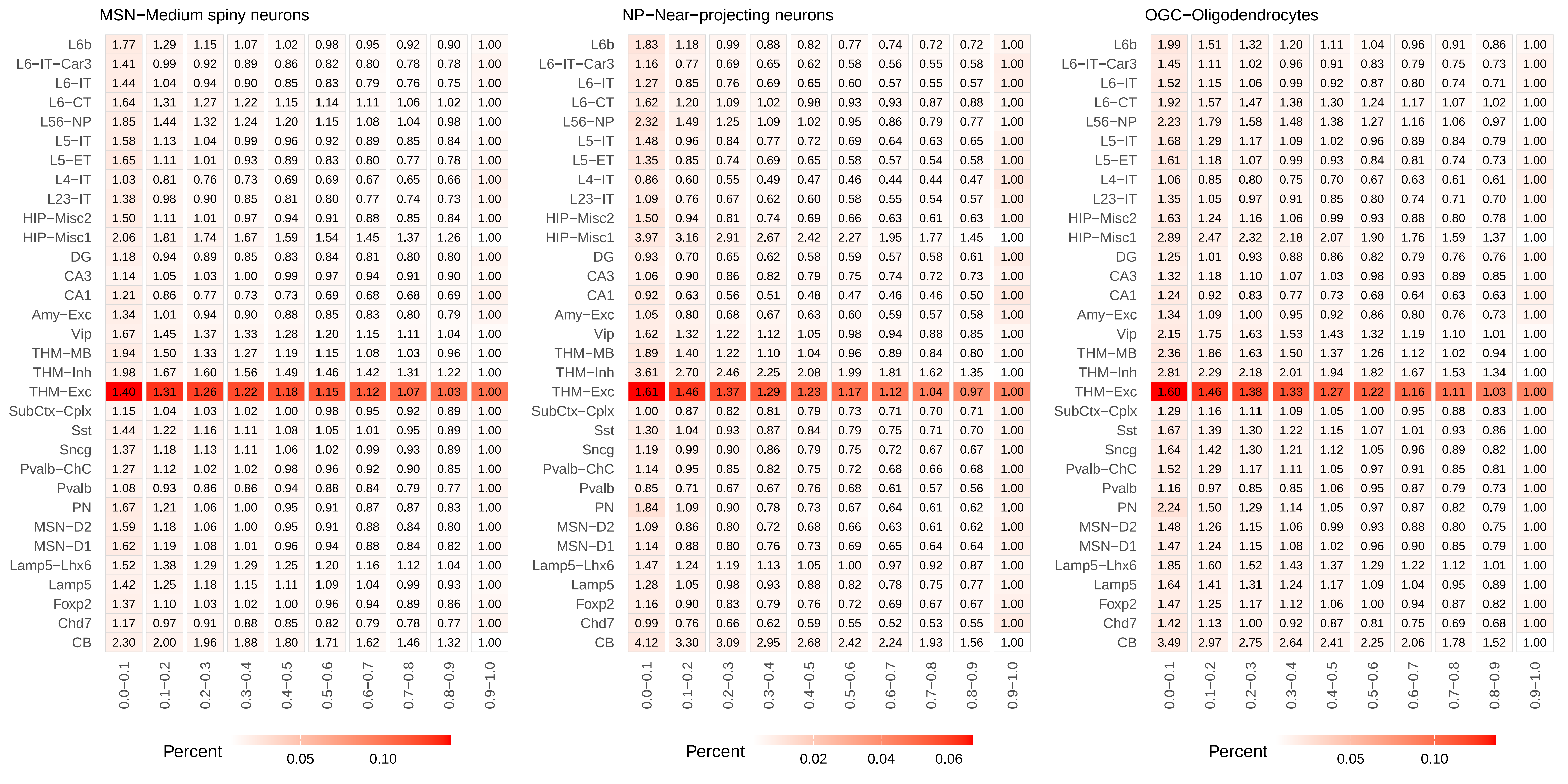
MSN−Medium spiny neurons
L6b 1.77 1.29 1.15 1.07 1.02 0.98 0.95 0.92 0.90 1.00
L6−IT−Car3 1.41 0.99 0.92 0.89 0.86 0.82 0.80 0.78 0.78 1.00
L6−IT 1.44 1.04 0.94 0.90 0.85 0.83 0.79 0.76 0.75 1.00
L6−CT 1.64 1.31 1.27 1.22 1.15 1.14 1.11 1.06 1.02 1.00
L56−NP 1.85 1.44 1.32 1.24 1.20 1.15 1.08 1.04 0.98 1.00
L5−IT 1.58 1.13 1.04 0.99 0.96 0.92 0.89 0.85 0.84 1.00
L5−ET 1.65 1.11 1.01 0.93 0.89 0.83 0.80 0.77 0.78 1.00
L4−IT 1.03 0.81 0.76 0.73 0.69 0.69 0.67 0.65 0.66 1.00
L23−IT 1.38 0.98 0.90 0.85 0.81 0.80 0.77 0.74 0.73 1.00
HIP−Misc2 1.50 1.11 1.01 0.97 0.94 0.91 0.88 0.85 0.84 1.00
HIP−Misc1 2.06 1.81 1.74 1.67 1.59 1.54 1.45 1.37 1.26 1.00
DG 1.18 0.94 0.89 0.85 0.83 0.84 0.81 0.80 0.80 1.00
CA3 1.14 1.05 1.03 1.00 0.99 0.97 0.94 0.91 0.90 1.00
CA1 1.21 0.86 0.77 0.73 0.73 0.69 0.68 0.68 0.69 1.00
Amy−Exc 1.34 1.01 0.94 0.90 0.88 0.85 0.83 0.80 0.79 1.00
Vip 1.67 1.45 1.37 1.33 1.28 1.20 1.15 1.11 1.04 1.00
THM−MB 1.94 1.50 1.33 1.27 1.19 1.15 1.08 1.03 0.96 1.00
THM−Inh 1.98 1.67 1.60 1.56 1.49 1.46 1.42 1.31 1.22 1.00
THM−Exc 1.40 1.31 1.26 1.22 1.18 1.15 1.12 1.07 1.03 1.00
SubCtx−Cplx 1.15 1.04 1.03 1.02 1.00 0.98 0.95 0.92 0.89 1.00
Sst 1.44 1.22 1.16 1.11 1.08 1.05 1.01 0.95 0.89 1.00
Sncg 1.37 1.18 1.13 1.11 1.06 1.02 0.99 0.93 0.89 1.00
Pvalb−ChC 1.27 1.12 1.02 1.02 0.98 0.96 0.92 0.90 0.85 1.00
Pvalb 1.08 0.93 0.86 0.86 0.94 0.88 0.84 0.79 0.77 1.00
PN 1.67 1.21 1.06 1.00 0.95 0.91 0.87 0.87 0.83 1.00
MSN−D2 1.59 1.18 1.06 1.00 0.95 0.91 0.88 0.84 0.80 1.00
MSN−D1 1.62 1.19 1.08 1.01 0.96 0.94 0.88 0.84 0.82 1.00
Lamp5−Lhx6 1.52 1.38 1.29 1.29 1.25 1.20 1.16 1.12 1.04 1.00
Lamp5 1.42 1.25 1.18 1.15 1.11 1.09 1.04 0.99 0.93 1.00
Foxp2 1.37 1.10 1.03 1.02 1.00 0.96 0.94 0.89 0.86 1.00
Chd7 1.17 0.97 0.91 0.88 0.85 0.82 0.79 0.78 0.77 1.00
CB 2.30 2.00 1.96 1.88 1.80 1.71 1.62 1.46 1.32 1.00
0.0−0.1 0.1−0.2 0.2−0.3 0.3−0.4 0.4−0.5 0.5−0.6 0.6−0.7 0.7−0.8 0.8−0.9 0.9−1.0
Percent 0.05 0.10
NP−Near−projecting neurons
L6b 1.83 1.18 0.99 0.88 0.82 0.77 0.74 0.72 0.72 1.00
L6−IT−Car3 1.16 0.77 0.69 0.65 0.62 0.58 0.56 0.55 0.58 1.00
L6−IT 1.27 0.85 0.76 0.69 0.65 0.60 0.57 0.55 0.57 1.00
L6−CT 1.62 1.20 1.09 1.02 0.98 0.93 0.93 0.87 0.88 1.00
L56−NP 2.32 1.49 1.25 1.09 1.02 0.95 0.86 0.79 0.77 1.00
L5−IT 1.48 0.96 0.84 0.77 0.72 0.69 0.64 0.63 0.65 1.00
L5−ET 1.35 0.85 0.74 0.69 0.65 0.58 0.57 0.54 0.58 1.00
L4−IT 0.86 0.60 0.55 0.49 0.47 0.46 0.44 0.44 0.47 1.00
L23−IT 1.09 0.76 0.67 0.62 0.60 0.58 0.55 0.54 0.57 1.00
HIP−Misc2 1.50 0.94 0.81 0.74 0.69 0.66 0.63 0.61 0.63 1.00
HIP−Misc1 3.97 3.16 2.91 2.67 2.42 2.27 1.95 1.77 1.45 1.00
DG 0.93 0.70 0.65 0.62 0.58 0.59 0.57 0.58 0.61 1.00
CA3 1.06 0.90 0.86 0.82 0.79 0.75 0.74 0.72 0.73 1.00
CA1 0.92 0.63 0.56 0.51 0.48 0.47 0.46 0.46 0.50 1.00
Amy−Exc 1.05 0.80 0.68 0.67 0.63 0.60 0.59 0.57 0.58 1.00
Vip 1.62 1.32 1.22 1.12 1.05 0.98 0.94 0.88 0.85 1.00
THM−MB 1.89 1.40 1.22 1.10 1.04 0.96 0.89 0.84 0.80 1.00
THM−Inh 3.61 2.70 2.46 2.25 2.08 1.99 1.81 1.62 1.35 1.00
THM−Exc 1.61 1.46 1.37 1.29 1.23 1.17 1.12 1.04 0.97 1.00
SubCtx−Cplx 1.00 0.87 0.82 0.81 0.79 0.73 0.71 0.70 0.71 1.00
Sst 1.30 1.04 0.93 0.87 0.84 0.79 0.75 0.71 0.70 1.00
Sncg 1.19 0.99 0.90 0.86 0.79 0.75 0.72 0.67 0.67 1.00
Pvalb−ChC 1.14 0.95 0.85 0.82 0.75 0.72 0.68 0.66 0.68 1.00
Pvalb 0.85 0.71 0.67 0.67 0.76 0.68 0.61 0.57 0.56 1.00
PN 1.84 1.09 0.90 0.78 0.73 0.67 0.64 0.61 0.62 1.00
MSN−D2 1.09 0.86 0.80 0.72 0.68 0.66 0.63 0.61 0.62 1.00
MSN−D1 1.14 0.88 0.80 0.76 0.73 0.69 0.65 0.64 0.64 1.00
Lamp5−Lhx6 1.47 1.24 1.19 1.13 1.05 1.00 0.97 0.92 0.87 1.00
Lamp5 1.28 1.05 0.98 0.93 0.88 0.82 0.78 0.75 0.77 1.00
Foxp2 1.16 0.90 0.83 0.79 0.76 0.72 0.69 0.67 0.67 1.00
Chd7 0.99 0.76 0.66 0.62 0.59 0.55 0.52 0.53 0.55 1.00
CB 4.12 3.30 3.09 2.95 2.68 2.42 2.24 1.93 1.56 1.00
0.0−0.1 0.1−0.2 0.2−0.3 0.3−0.4 0.4−0.5 0.5−0.6 0.6−0.7 0.7−0.8 0.8−0.9 0.9−1.0
Percent 0.02 0.04 0.06
OGC−Oligodendrocytes
L6b 1.99 1.51 1.32 1.20 1.11 1.04 0.96 0.91 0.86 1.00
L6−IT−Car3 1.45 1.11 1.02 0.96 0.91 0.83 0.79 0.75 0.73 1.00
L6−IT 1.52 1.15 1.06 0.99 0.92 0.87 0.80 0.74 0.71 1.00
L6−CT 1.92 1.57 1.47 1.38 1.30 1.24 1.17 1.07 1.02 1.00
L56−NP 2.23 1.79 1.58 1.48 1.38 1.27 1.16 1.06 0.97 1.00
L5−IT 1.68 1.29 1.17 1.09 1.02 0.96 0.89 0.84 0.79 1.00
L5−ET 1.61 1.18 1.07 0.99 0.93 0.84 0.81 0.74 0.73 1.00
L4−IT 1.06 0.85 0.80 0.75 0.70 0.67 0.63 0.61 0.61 1.00
L23−IT 1.35 1.05 0.97 0.91 0.85 0.80 0.74 0.71 0.70 1.00
HIP−Misc2 1.63 1.24 1.16 1.06 0.99 0.93 0.88 0.80 0.78 1.00
HIP−Misc1 2.89 2.47 2.32 2.18 2.07 1.90 1.76 1.59 1.37 1.00
DG 1.25 1.01 0.93 0.88 0.86 0.82 0.79 0.76 0.76 1.00
CA3 1.32 1.18 1.10 1.07 1.03 0.98 0.93 0.89 0.85 1.00
CA1 1.24 0.92 0.83 0.77 0.73 0.68 0.64 0.63 0.63 1.00
Amy−Exc 1.34 1.09 1.00 0.95 0.92 0.86 0.80 0.76 0.73 1.00
Vip 2.15 1.75 1.63 1.53 1.43 1.32 1.19 1.10 1.01 1.00
THM−MB 2.36 1.86 1.63 1.50 1.37 1.26 1.12 1.02 0.94 1.00
THM−Inh 2.81 2.29 2.18 2.01 1.94 1.82 1.67 1.53 1.34 1.00
THM−Exc 1.60 1.46 1.38 1.33 1.27 1.22 1.16 1.11 1.03 1.00
SubCtx−Cplx 1.29 1.16 1.11 1.09 1.05 1.00 0.95 0.88 0.83 1.00
Sst 1.67 1.39 1.30 1.22 1.15 1.07 1.01 0.93 0.86 1.00
Sncg 1.64 1.42 1.30 1.21 1.12 1.05 0.96 0.89 0.82 1.00
Pvalb−ChC 1.52 1.29 1.17 1.11 1.05 0.97 0.91 0.85 0.81 1.00
Pvalb 1.16 0.97 0.85 0.85 1.06 0.95 0.87 0.79 0.73 1.00
PN 2.24 1.50 1.29 1.14 1.05 0.97 0.87 0.82 0.79 1.00
MSN−D2 1.48 1.26 1.15 1.06 0.99 0.93 0.88 0.80 0.75 1.00
MSN−D1 1.47 1.24 1.15 1.08 1.02 0.96 0.90 0.85 0.79 1.00
Lamp5−Lhx6 1.85 1.60 1.52 1.43 1.37 1.29 1.22 1.12 1.01 1.00
Lamp5 1.64 1.41 1.31 1.24 1.17 1.09 1.04 0.95 0.89 1.00
Foxp2 1.47 1.25 1.17 1.12 1.06 1.00 0.94 0.87 0.82 1.00
Chd7 1.42 1.13 1.00 0.92 0.87 0.81 0.75 0.69 0.68 1.00
CB 3.49 2.97 2.75 2.64 2.41 2.25 2.06 1.78 1.52 1.00
0.0−0.1 0.1−0.2 0.2−0.3 0.3−0.4 0.4−0.5 0.5−0.6 0.6−0.7 0.7−0.8 0.8−0.9 0.9−1.0
Percent 0.05 0.10

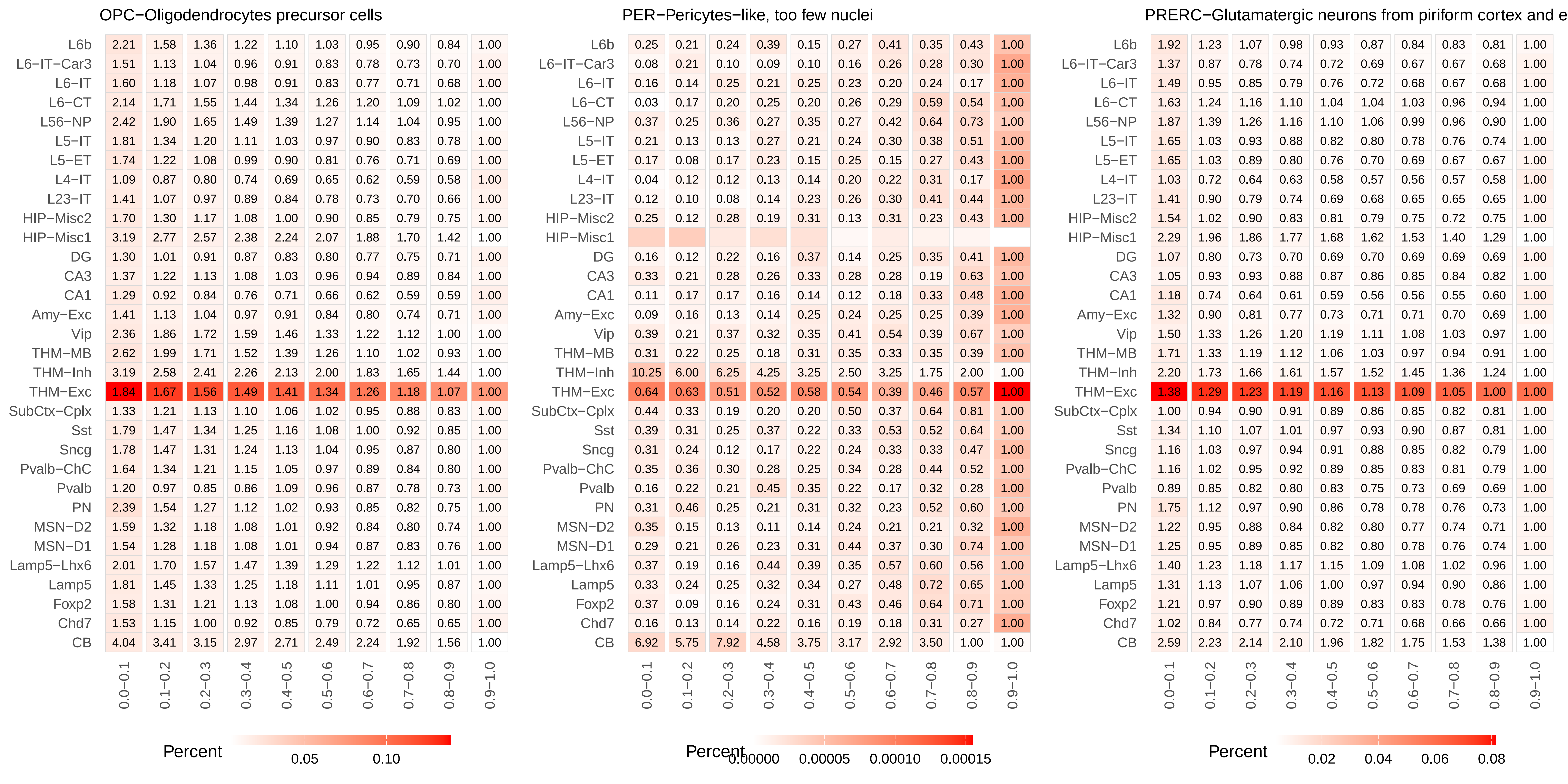
OPC−Oligodendrocytes precursor cells
L6b 2.21 1.58 1.36 1.22 1.10 1.03 0.95 0.90 0.84 1.00
L6−IT−Car3 1.51 1.13 1.04 0.96 0.91 0.83 0.78 0.73 0.70 1.00
L6−IT 1.60 1.18 1.07 0.98 0.91 0.83 0.77 0.71 0.68 1.00
L6−CT 2.14 1.71 1.55 1.44 1.34 1.26 1.20 1.09 1.02 1.00
L56−NP 2.42 1.90 1.65 1.49 1.39 1.27 1.14 1.04 0.95 1.00
L5−IT 1.81 1.34 1.20 1.11 1.03 0.97 0.90 0.83 0.78 1.00
L5−ET 1.74 1.22 1.08 0.99 0.90 0.81 0.76 0.71 0.69 1.00
L4−IT 1.09 0.87 0.80 0.74 0.69 0.65 0.62 0.59 0.58 1.00
L23−IT 1.41 1.07 0.97 0.89 0.84 0.78 0.73 0.70 0.66 1.00
HIP−Misc2 1.70 1.30 1.17 1.08 1.00 0.90 0.85 0.79 0.75 1.00
HIP−Misc1 3.19 2.77 2.57 2.38 2.24 2.07 1.88 1.70 1.42 1.00
DG 1.30 1.01 0.91 0.87 0.83 0.80 0.77 0.75 0.71 1.00
CA3 1.37 1.22 1.13 1.08 1.03 0.96 0.94 0.89 0.84 1.00
CA1 1.29 0.92 0.84 0.76 0.71 0.66 0.62 0.59 0.59 1.00
Amy−Exc 1.41 1.13 1.04 0.97 0.91 0.84 0.80 0.74 0.71 1.00
Vip 2.36 1.86 1.72 1.59 1.46 1.33 1.22 1.12 1.00 1.00
THM−MB 2.62 1.99 1.71 1.52 1.39 1.26 1.10 1.02 0.93 1.00
THM−Inh 3.19 2.58 2.41 2.26 2.13 2.00 1.83 1.65 1.44 1.00
THM−Exc 1.84 1.67 1.56 1.49 1.41 1.34 1.26 1.18 1.07 1.00
SubCtx−Cplx 1.33 1.21 1.13 1.10 1.06 1.02 0.95 0.88 0.83 1.00
Sst 1.79 1.47 1.34 1.25 1.16 1.08 1.00 0.92 0.85 1.00
Sncg 1.78 1.47 1.31 1.24 1.13 1.04 0.95 0.87 0.80 1.00
Pvalb−ChC 1.64 1.34 1.21 1.15 1.05 0.97 0.89 0.84 0.80 1.00
Pvalb 1.20 0.97 0.85 0.86 1.09 0.96 0.87 0.78 0.73 1.00
PN 2.39 1.54 1.27 1.12 1.02 0.93 0.85 0.82 0.75 1.00
MSN−D2 1.59 1.32 1.18 1.08 1.01 0.92 0.84 0.80 0.74 1.00
MSN−D1 1.54 1.28 1.18 1.08 1.01 0.94 0.87 0.83 0.76 1.00
Lamp5−Lhx6 2.01 1.70 1.57 1.47 1.39 1.29 1.22 1.12 1.01 1.00
Lamp5 1.81 1.45 1.33 1.25 1.18 1.11 1.01 0.95 0.87 1.00
Foxp2 1.58 1.31 1.21 1.13 1.08 1.00 0.94 0.86 0.80 1.00
Chd7 1.53 1.15 1.00 0.92 0.85 0.79 0.72 0.65 0.65 1.00
CB 4.04 3.41 3.15 2.97 2.71 2.49 2.24 1.92 1.56 1.00
0.0−0.1 0.1−0.2 0.2−0.3 0.3−0.4 0.4−0.5 0.5−0.6 0.6−0.7 0.7−0.8 0.8−0.9 0.9−1.0
Percent 0.05 0.10
PER−Pericytes−like, too few nuclei
L6b 0.25 0.21 0.24 0.39 0.15 0.27 0.41 0.35 0.43 1.00
L6−IT−Car3 0.08 0.21 0.10 0.09 0.10 0.16 0.26 0.28 0.30 1.00
L6−IT 0.16 0.14 0.25 0.21 0.25 0.23 0.20 0.24 0.17 1.00
L6−CT 0.03 0.17 0.20 0.25 0.20 0.26 0.29 0.59 0.54 1.00
L56−NP 0.37 0.25 0.36 0.27 0.35 0.27 0.42 0.64 0.73 1.00
L5−IT 0.21 0.13 0.13 0.27 0.21 0.24 0.30 0.38 0.51 1.00
L5−ET 0.17 0.08 0.17 0.23 0.15 0.25 0.15 0.27 0.43 1.00
L4−IT 0.04 0.12 0.12 0.13 0.14 0.20 0.22 0.31 0.17 1.00
L23−IT 0.12 0.10 0.08 0.14 0.23 0.26 0.30 0.41 0.44 1.00
HIP−Misc2 0.25 0.12 0.28 0.19 0.31 0.13 0.31 0.23 0.43 1.00
HIP−Misc1
DG 0.16 0.12 0.22 0.16 0.37 0.14 0.25 0.35 0.41 1.00
CA3 0.33 0.21 0.28 0.26 0.33 0.28 0.28 0.19 0.63 1.00
CA1 0.11 0.17 0.17 0.16 0.14 0.12 0.18 0.33 0.48 1.00
Amy−Exc 0.09 0.16 0.13 0.14 0.25 0.24 0.25 0.25 0.39 1.00
Vip 0.39 0.21 0.37 0.32 0.35 0.41 0.54 0.39 0.67 1.00
THM−MB 0.31 0.22 0.25 0.18 0.31 0.35 0.33 0.35 0.39 1.00
THM−Inh 10.25 6.00 6.25 4.25 3.25 2.50 3.25 1.75 2.00 1.00
THM−Exc 0.64 0.63 0.51 0.52 0.58 0.54 0.39 0.46 0.57 1.00
SubCtx−Cplx 0.44 0.33 0.19 0.20 0.20 0.50 0.37 0.64 0.81 1.00
Sst 0.39 0.31 0.25 0.37 0.22 0.33 0.53 0.52 0.64 1.00
Sncg 0.31 0.24 0.12 0.17 0.22 0.24 0.33 0.33 0.47 1.00
Pvalb−ChC 0.35 0.36 0.30 0.28 0.25 0.34 0.28 0.44 0.52 1.00
Pvalb 0.16 0.22 0.21 0.45 0.35 0.22 0.17 0.32 0.28 1.00
PN 0.31 0.46 0.25 0.21 0.31 0.32 0.23 0.52 0.60 1.00
MSN−D2 0.35 0.15 0.13 0.11 0.14 0.24 0.21 0.21 0.32 1.00
MSN−D1 0.29 0.21 0.26 0.23 0.31 0.44 0.37 0.30 0.74 1.00
Lamp5−Lhx6 0.37 0.19 0.16 0.44 0.39 0.35 0.57 0.60 0.56 1.00
Lamp5 0.33 0.24 0.25 0.32 0.34 0.27 0.48 0.72 0.65 1.00
Foxp2 0.37 0.09 0.16 0.24 0.31 0.43 0.46 0.64 0.71 1.00
Chd7 0.16 0.13 0.14 0.22 0.16 0.19 0.18 0.31 0.27 1.00
CB 6.92 5.75 7.92 4.58 3.75 3.17 2.92 3.50 1.00 1.00
0.0−0.1 0.1−0.2 0.2−0.3 0.3−0.4 0.4−0.5 0.5−0.6 0.6−0.7 0.7−0.8 0.8−0.9 0.9−1.0
Percent 0.00000 0.00005 0.00010 0.00015
PRERC−Glutamatergic neurons from piriform cortex and e
L6b 1.92 1.23 1.07 0.98 0.93 0.87 0.84 0.83 0.81 1.00
L6−IT−Car3 1.37 0.87 0.78 0.74 0.72 0.69 0.67 0.67 0.68 1.00
L6−IT 1.49 0.95 0.85 0.79 0.76 0.72 0.68 0.67 0.68 1.00
L6−CT 1.63 1.24 1.16 1.10 1.04 1.04 1.03 0.96 0.94 1.00
L56−NP 1.87 1.39 1.26 1.16 1.10 1.06 0.99 0.96 0.90 1.00
L5−IT 1.65 1.03 0.93 0.88 0.82 0.80 0.78 0.76 0.74 1.00
L5−ET 1.65 1.03 0.89 0.80 0.76 0.70 0.69 0.67 0.67 1.00
L4−IT 1.03 0.72 0.64 0.63 0.58 0.57 0.56 0.57 0.58 1.00
L23−IT 1.41 0.90 0.79 0.74 0.69 0.68 0.65 0.65 0.65 1.00
HIP−Misc2 1.54 1.02 0.90 0.83 0.81 0.79 0.75 0.72 0.75 1.00
HIP−Misc1 2.29 1.96 1.86 1.77 1.68 1.62 1.53 1.40 1.29 1.00
DG 1.07 0.80 0.73 0.70 0.69 0.70 0.69 0.69 0.69 1.00
CA3 1.05 0.93 0.93 0.88 0.87 0.86 0.85 0.84 0.82 1.00
CA1 1.18 0.74 0.64 0.61 0.59 0.56 0.56 0.55 0.60 1.00
Amy−Exc 1.32 0.90 0.81 0.77 0.73 0.71 0.71 0.70 0.69 1.00
Vip 1.50 1.33 1.26 1.20 1.19 1.11 1.08 1.03 0.97 1.00
THM−MB 1.71 1.33 1.19 1.12 1.06 1.03 0.97 0.94 0.91 1.00
THM−Inh 2.20 1.73 1.66 1.61 1.57 1.52 1.45 1.36 1.24 1.00
THM−Exc 1.38 1.29 1.23 1.19 1.16 1.13 1.09 1.05 1.00 1.00
SubCtx−Cplx 1.00 0.94 0.90 0.91 0.89 0.86 0.85 0.82 0.81 1.00
Sst 1.34 1.10 1.07 1.01 0.97 0.93 0.90 0.87 0.81 1.00
Sncg 1.16 1.03 0.97 0.94 0.91 0.88 0.85 0.82 0.79 1.00
Pvalb−ChC 1.16 1.02 0.95 0.92 0.89 0.85 0.83 0.81 0.79 1.00
Pvalb 0.89 0.85 0.82 0.80 0.83 0.75 0.73 0.69 0.69 1.00
PN 1.75 1.12 0.97 0.90 0.86 0.78 0.78 0.76 0.73 1.00
MSN−D2 1.22 0.95 0.88 0.84 0.82 0.80 0.77 0.74 0.71 1.00
MSN−D1 1.25 0.95 0.89 0.85 0.82 0.80 0.78 0.76 0.74 1.00
Lamp5−Lhx6 1.40 1.23 1.18 1.17 1.15 1.09 1.08 1.02 0.96 1.00
Lamp5 1.31 1.13 1.07 1.06 1.00 0.97 0.94 0.90 0.86 1.00
Foxp2 1.21 0.97 0.90 0.89 0.89 0.83 0.83 0.78 0.76 1.00
Chd7 1.02 0.84 0.77 0.74 0.72 0.71 0.68 0.66 0.66 1.00
CB 2.59 2.23 2.14 2.10 1.96 1.82 1.75 1.53 1.38 1.00
0.0−0.1 0.1−0.2 0.2−0.3 0.3−0.4 0.4−0.5 0.5−0.6 0.6−0.7 0.7−0.8 0.8−0.9 0.9−1.0
Percent 0.02 0.04 0.06 0.08

### PVALB−PVALB+ GABAergic neurons

| | 0.0−0.1 | 0.1−0.2 | 0.2−0.3 | 0.3−0.4 | 0.4−0.5 | 0.5−0.6 | 0.6−0.7 | 0.7−0.8 | 0.8−0.9 | 0.9−1.0 |
|---|---|---|---|---|---|---|---|---|---|---|
| L6b | 2.00 | 1.39 | 1.22 | 1.13 | 1.03 | 0.98 | 0.92 | 0.87 | 0.84 | 1.00 |
| L6−IT−Car3 | 1.42 | 1.02 | 0.94 | 0.86 | 0.82 | 0.78 | 0.75 | 0.71 | 0.70 | 1.00 |
| L6−IT | 1.48 | 1.06 | 0.96 | 0.91 | 0.86 | 0.80 | 0.74 | 0.70 | 0.68 | 1.00 |
| L6−CT | 1.83 | 1.47 | 1.36 | 1.29 | 1.21 | 1.17 | 1.10 | 1.03 | 0.98 | 1.00 |
| L56−NP | 2.14 | 1.63 | 1.45 | 1.36 | 1.27 | 1.19 | 1.09 | 1.00 | 0.93 | 1.00 |
| L5−IT | 1.67 | 1.19 | 1.05 | 1.02 | 0.95 | 0.92 | 0.86 | 0.81 | 0.78 | 1.00 |
| L5−ET | 1.66 | 1.12 | 1.00 | 0.91 | 0.85 | 0.78 | 0.74 | 0.71 | 0.70 | 1.00 |
| L4−IT | 1.04 | 0.79 | 0.74 | 0.69 | 0.66 | 0.64 | 0.60 | 0.58 | 0.59 | 1.00 |
| L23−IT | 1.33 | 0.98 | 0.90 | 0.83 | 0.78 | 0.76 | 0.71 | 0.68 | 0.66 | 1.00 |
| HIP−Misc2 | 1.59 | 1.16 | 1.07 | 1.00 | 0.93 | 0.87 | 0.84 | 0.78 | 0.76 | 1.00 |
| HIP−Misc1 | 2.65 | 2.32 | 2.19 | 2.06 | 1.97 | 1.87 | 1.73 | 1.57 | 1.36 | 1.00 |
| DG | 1.14 | 0.90 | 0.85 | 0.81 | 0.78 | 0.76 | 0.74 | 0.72 | 0.71 | 1.00 |
| CA3 | 1.16 | 1.08 | 1.02 | 1.00 | 0.97 | 0.95 | 0.91 | 0.85 | 0.84 | 1.00 |
| CA1 | 1.18 | 0.82 | 0.75 | 0.71 | 0.66 | 0.64 | 0.60 | 0.60 | 0.60 | 1.00 |
| Amy−Exc | 1.29 | 1.00 | 0.93 | 0.88 | 0.84 | 0.80 | 0.77 | 0.73 | 0.70 | 1.00 |
| Vip | 1.91 | 1.61 | 1.51 | 1.42 | 1.35 | 1.25 | 1.18 | 1.07 | 1.00 | 1.00 |
| THM−MB | 2.22 | 1.68 | 1.48 | 1.39 | 1.25 | 1.17 | 1.07 | 0.99 | 0.92 | 1.00 |
| THM−Inh | 2.61 | 2.12 | 2.01 | 1.94 | 1.83 | 1.74 | 1.67 | 1.54 | 1.35 | 1.00 |
| THM−Exc | 1.56 | 1.44 | 1.37 | 1.32 | 1.27 | 1.22 | 1.17 | 1.12 | 1.04 | 1.00 |
| SubCtx−Cplx | 1.15 | 1.05 | 1.02 | 1.01 | 0.98 | 0.96 | 0.91 | 0.87 | 0.81 | 1.00 |
| Sst | 1.60 | 1.30 | 1.21 | 1.13 | 1.08 | 1.03 | 0.96 | 0.89 | 0.83 | 1.00 |
| Sncg | 1.45 | 1.26 | 1.17 | 1.11 | 1.04 | 0.99 | 0.93 | 0.87 | 0.80 | 1.00 |
| Pvalb−ChC | 1.46 | 1.21 | 1.09 | 1.03 | 0.98 | 0.92 | 0.87 | 0.82 | 0.80 | 1.00 |
| Pvalb | 1.02 | 0.90 | 0.84 | 0.83 | 0.97 | 0.88 | 0.83 | 0.76 | 0.72 | 1.00 |
| PN | 2.13 | 1.34 | 1.14 | 1.02 | 0.95 | 0.88 | 0.82 | 0.79 | 0.74 | 1.00 |
| MSN−D2 | 1.38 | 1.14 | 1.04 | 0.98 | 0.93 | 0.89 | 0.83 | 0.78 | 0.75 | 1.00 |
| MSN−D1 | 1.35 | 1.13 | 1.05 | 0.98 | 0.93 | 0.90 | 0.85 | 0.81 | 0.76 | 1.00 |
| Lamp5−Lhx6 | 1.67 | 1.45 | 1.37 | 1.34 | 1.29 | 1.21 | 1.16 | 1.07 | 0.97 | 1.00 |
| Lamp5 | 1.50 | 1.30 | 1.20 | 1.14 | 1.11 | 1.06 | 0.99 | 0.93 | 0.86 | 1.00 |
| Foxp2 | 1.34 | 1.12 | 1.07 | 1.02 | 0.99 | 0.93 | 0.88 | 0.82 | 0.78 | 1.00 |
| Chd7 | 1.34 | 1.02 | 0.91 | 0.85 | 0.80 | 0.76 | 0.71 | 0.66 | 0.66 | 1.00 |
| CB | 3.20 | 2.75 | 2.60 | 2.49 | 2.34 | 2.14 | 1.97 | 1.73 | 1.49 | 1.00 |

Percent 0.025 0.050 0.075 0.100

### PV_ChCs−PVALB+ chandelier cells

| | 0.0−0.1 | 0.1−0.2 | 0.2−0.3 | 0.3−0.4 | 0.4−0.5 | 0.5−0.6 | 0.6−0.7 | 0.7−0.8 | 0.8−0.9 | 0.9−1.0 |
|---|---|---|---|---|---|---|---|---|---|---|
| L6b | 1.41 | 0.98 | 0.86 | 0.79 | 0.74 | 0.71 | 0.69 | 0.69 | 0.72 | 1.00 |
| L6−IT−Car3 | 0.93 | 0.66 | 0.60 | 0.58 | 0.57 | 0.55 | 0.54 | 0.53 | 0.57 | 1.00 |
| L6−IT | 1.01 | 0.72 | 0.66 | 0.62 | 0.58 | 0.55 | 0.52 | 0.52 | 0.54 | 1.00 |
| L6−CT | 1.28 | 1.02 | 0.97 | 0.93 | 0.89 | 0.87 | 0.87 | 0.85 | 0.84 | 1.00 |
| L56−NP | 1.70 | 1.22 | 1.08 | 1.01 | 0.94 | 0.90 | 0.84 | 0.81 | 0.80 | 1.00 |
| L5−IT | 1.15 | 0.80 | 0.73 | 0.69 | 0.66 | 0.64 | 0.63 | 0.61 | 0.63 | 1.00 |
| L5−ET | 1.08 | 0.72 | 0.67 | 0.61 | 0.57 | 0.55 | 0.55 | 0.53 | 0.57 | 1.00 |
| L4−IT | 0.67 | 0.50 | 0.48 | 0.45 | 0.43 | 0.43 | 0.42 | 0.42 | 0.47 | 1.00 |
| L23−IT | 0.87 | 0.64 | 0.59 | 0.56 | 0.53 | 0.52 | 0.50 | 0.51 | 0.54 | 1.00 |
| HIP−Misc2 | 1.11 | 0.80 | 0.72 | 0.68 | 0.65 | 0.61 | 0.61 | 0.58 | 0.62 | 1.00 |
| HIP−Misc1 | 3.53 | 2.90 | 2.62 | 2.47 | 2.25 | 2.07 | 1.88 | 1.72 | 1.41 | 1.00 |
| DG | 0.78 | 0.60 | 0.56 | 0.54 | 0.52 | 0.54 | 0.54 | 0.54 | 0.57 | 1.00 |
| CA3 | 0.89 | 0.80 | 0.74 | 0.73 | 0.73 | 0.70 | 0.70 | 0.68 | 0.70 | 1.00 |
| CA1 | 0.76 | 0.53 | 0.48 | 0.45 | 0.44 | 0.43 | 0.43 | 0.44 | 0.48 | 1.00 |
| Amy−Exc | 0.86 | 0.67 | 0.62 | 0.58 | 0.57 | 0.55 | 0.54 | 0.55 | 0.56 | 1.00 |
| Vip | 1.49 | 1.19 | 1.10 | 1.03 | 0.99 | 0.92 | 0.90 | 0.84 | 0.84 | 1.00 |
| THM−MB | 1.73 | 1.25 | 1.09 | 1.02 | 0.96 | 0.90 | 0.83 | 0.79 | 0.79 | 1.00 |
| THM−Inh | 3.42 | 2.47 | 2.30 | 2.07 | 1.96 | 1.85 | 1.75 | 1.59 | 1.43 | 1.00 |
| THM−Exc | 1.55 | 1.39 | 1.29 | 1.24 | 1.18 | 1.12 | 1.07 | 1.02 | 0.96 | 1.00 |
| SubCtx−Cplx | 0.82 | 0.74 | 0.71 | 0.69 | 0.68 | 0.66 | 0.65 | 0.65 | 0.66 | 1.00 |
| Sst | 1.15 | 0.91 | 0.83 | 0.79 | 0.75 | 0.73 | 0.68 | 0.66 | 0.68 | 1.00 |
| Sncg | 1.04 | 0.87 | 0.79 | 0.76 | 0.73 | 0.69 | 0.68 | 0.65 | 0.65 | 1.00 |
| Pvalb−ChC | 1.11 | 0.86 | 0.76 | 0.74 | 0.68 | 0.64 | 0.63 | 0.61 | 0.64 | 1.00 |
| Pvalb | 0.74 | 0.64 | 0.61 | 0.65 | 0.70 | 0.60 | 0.59 | 0.56 | 0.55 | 1.00 |
| PN | 1.52 | 0.93 | 0.78 | 0.70 | 0.66 | 0.63 | 0.60 | 0.60 | 0.61 | 1.00 |
| MSN−D2 | 0.91 | 0.74 | 0.69 | 0.66 | 0.62 | 0.59 | 0.59 | 0.58 | 0.58 | 1.00 |
| MSN−D1 | 0.94 | 0.75 | 0.71 | 0.66 | 0.65 | 0.63 | 0.61 | 0.59 | 0.60 | 1.00 |
| Lamp5−Lhx6 | 1.29 | 1.07 | 1.02 | 0.99 | 0.94 | 0.91 | 0.88 | 0.84 | 0.81 | 1.00 |
| Lamp5 | 1.13 | 0.92 | 0.84 | 0.82 | 0.79 | 0.78 | 0.73 | 0.71 | 0.71 | 1.00 |
| Foxp2 | 0.99 | 0.79 | 0.73 | 0.71 | 0.70 | 0.68 | 0.65 | 0.62 | 0.65 | 1.00 |
| Chd7 | 0.91 | 0.66 | 0.60 | 0.55 | 0.52 | 0.51 | 0.48 | 0.48 | 0.52 | 1.00 |
| CB | 3.73 | 3.01 | 2.86 | 2.75 | 2.52 | 2.29 | 2.14 | 1.78 | 1.53 | 1.00 |

Percent 0.02 0.04 0.06

### SEPGA−Dopaminergic neurons from septal nuclei

| | 0.0−0.1 | 0.1−0.2 | 0.2−0.3 | 0.3−0.4 | 0.4−0.5 | 0.5−0.6 | 0.6−0.7 | 0.7−0.8 | 0.8−0.9 | 0.9−1.0 |
|---|---|---|---|---|---|---|---|---|---|---|
| L6b | 0.57 | 0.45 | 0.40 | 0.37 | 0.39 | 0.36 | 0.42 | 0.47 | 0.52 | 1.00 |
| L6−IT−Car3 | 0.30 | 0.23 | 0.21 | 0.23 | 0.24 | 0.23 | 0.28 | 0.29 | 0.40 | 1.00 |
| L6−IT | 0.42 | 0.31 | 0.28 | 0.28 | 0.27 | 0.26 | 0.29 | 0.35 | 0.41 | 1.00 |
| L6−CT | 0.49 | 0.44 | 0.43 | 0.43 | 0.46 | 0.47 | 0.54 | 0.56 | 0.75 | 1.00 |
| L56−NP | 0.84 | 0.59 | 0.51 | 0.46 | 0.50 | 0.41 | 0.46 | 0.54 | 0.61 | 1.00 |
| L5−IT | 0.40 | 0.29 | 0.29 | 0.31 | 0.29 | 0.33 | 0.38 | 0.40 | 0.46 | 1.00 |
| L5−ET | 0.34 | 0.27 | 0.22 | 0.27 | 0.28 | 0.26 | 0.28 | 0.31 | 0.40 | 1.00 |
| L4−IT | 0.20 | 0.16 | 0.15 | 0.15 | 0.16 | 0.18 | 0.21 | 0.26 | 0.33 | 1.00 |
| L23−IT | 0.31 | 0.22 | 0.21 | 0.21 | 0.23 | 0.27 | 0.28 | 0.33 | 0.38 | 1.00 |
| HIP−Misc2 | 0.44 | 0.32 | 0.27 | 0.27 | 0.30 | 0.28 | 0.31 | 0.31 | 0.40 | 1.00 |
| HIP−Misc1 | 11.94 | 8.96 | 6.67 | 6.06 | 5.48 | 4.54 | 4.21 | 3.31 | 2.27 | 1.00 |
| DG | 0.25 | 0.23 | 0.21 | 0.24 | 0.22 | 0.27 | 0.26 | 0.33 | 0.38 | 1.00 |
| CA3 | 0.45 | 0.39 | 0.40 | 0.37 | 0.33 | 0.38 | 0.42 | 0.44 | 0.51 | 1.00 |
| CA1 | 0.23 | 0.17 | 0.17 | 0.18 | 0.20 | 0.18 | 0.24 | 0.28 | 0.34 | 1.00 |
| Amy−Exc | 0.30 | 0.23 | 0.23 | 0.20 | 0.24 | 0.25 | 0.26 | 0.30 | 0.41 | 1.00 |
| Vip | 0.71 | 0.52 | 0.51 | 0.46 | 0.49 | 0.52 | 0.52 | 0.59 | 0.63 | 1.00 |
| THM−MB | 0.72 | 0.55 | 0.47 | 0.55 | 0.53 | 0.50 | 0.51 | 0.52 | 0.61 | 1.00 |
| THM−Inh | 7.53 | 4.36 | 4.21 | 2.96 | 2.56 | 2.49 | 2.29 | 1.87 | 1.45 | 1.00 |
| THM−Exc | 0.98 | 0.89 | 0.85 | 0.81 | 0.77 | 0.71 | 0.71 | 0.73 | 0.69 | 1.00 |
| SubCtx−Cplx | 0.35 | 0.29 | 0.31 | 0.30 | 0.31 | 0.33 | 0.31 | 0.38 | 0.49 | 1.00 |
| Sst | 0.48 | 0.40 | 0.36 | 0.33 | 0.35 | 0.38 | 0.39 | 0.40 | 0.43 | 1.00 |
| Sncg | 0.36 | 0.30 | 0.28 | 0.27 | 0.30 | 0.30 | 0.34 | 0.41 | 0.46 | 1.00 |
| Pvalb−ChC | 0.37 | 0.36 | 0.32 | 0.35 | 0.26 | 0.35 | 0.31 | 0.37 | 0.47 | 1.00 |
| Pvalb | 0.32 | 0.38 | 0.49 | 0.54 | 0.31 | 0.23 | 0.24 | 0.28 | 0.34 | 1.00 |
| PN | 0.57 | 0.36 | 0.31 | 0.29 | 0.27 | 0.28 | 0.32 | 0.35 | 0.43 | 1.00 |
| MSN−D2 | 0.30 | 0.28 | 0.25 | 0.25 | 0.27 | 0.27 | 0.30 | 0.34 | 0.40 | 1.00 |
| MSN−D1 | 0.34 | 0.27 | 0.28 | 0.28 | 0.25 | 0.27 | 0.33 | 0.35 | 0.41 | 1.00 |
| Lamp5−Lhx6 | 0.62 | 0.46 | 0.48 | 0.53 | 0.52 | 0.47 | 0.50 | 0.56 | 0.59 | 1.00 |
| Lamp5 | 0.46 | 0.39 | 0.35 | 0.38 | 0.40 | 0.42 | 0.41 | 0.47 | 0.52 | 1.00 |
| Foxp2 | 0.39 | 0.31 | 0.31 | 0.30 | 0.31 | 0.29 | 0.33 | 0.36 | 0.42 | 1.00 |
| Chd7 | 0.27 | 0.21 | 0.19 | 0.20 | 0.19 | 0.20 | 0.23 | 0.26 | 0.37 | 1.00 |
| CB | 6.37 | 5.30 | 4.66 | 4.39 | 3.77 | 3.50 | 3.17 | 2.67 | 2.00 | 1.00 |

Percent 0.001 0.002

#### SIGA−Dopaminergic neurons from Inferior colliculus and nearby nuclei

| | 0.0−0.1 | 0.1−0.2 | 0.2−0.3 | 0.3−0.4 | 0.4−0.5 | 0.5−0.6 | 0.6−0.7 | 0.7−0.8 | 0.8−0.9 | 0.9−1.0 |
|---|---|---|---|---|---|---|---|---|---|---|
| L6b | 0.66 | 0.46 | 0.43 | 0.39 | 0.35 | 0.41 | 0.42 | 0.44 | 0.54 | 1.00 |
| L6−IT−Car3 | 0.37 | 0.26 | 0.24 | 0.25 | 0.26 | 0.26 | 0.30 | 0.33 | 0.43 | 1.00 |
| L6−IT | 0.43 | 0.31 | 0.28 | 0.27 | 0.28 | 0.29 | 0.29 | 0.32 | 0.39 | 1.00 |
| L6−CT | 0.54 | 0.46 | 0.45 | 0.46 | 0.43 | 0.48 | 0.56 | 0.55 | 0.68 | 1.00 |
| L56−NP | 0.95 | 0.64 | 0.56 | 0.52 | 0.52 | 0.49 | 0.53 | 0.54 | 0.59 | 1.00 |
| L5−IT | 0.49 | 0.33 | 0.33 | 0.32 | 0.35 | 0.35 | 0.38 | 0.41 | 0.49 | 1.00 |
| L5−ET | 0.43 | 0.29 | 0.31 | 0.29 | 0.28 | 0.26 | 0.32 | 0.35 | 0.42 | 1.00 |
| L4−IT | 0.24 | 0.19 | 0.18 | 0.18 | 0.19 | 0.19 | 0.23 | 0.27 | 0.32 | 1.00 |
| L23−IT | 0.38 | 0.25 | 0.25 | 0.23 | 0.26 | 0.27 | 0.31 | 0.33 | 0.40 | 1.00 |
| HIP−Misc2 | 0.53 | 0.37 | 0.33 | 0.31 | 0.32 | 0.32 | 0.33 | 0.37 | 0.47 | 1.00 |
| HIP−Misc1 | 8.47 | 5.87 | 4.59 | 4.18 | 4.18 | 3.56 | 2.66 | 2.25 | 1.87 | 1.00 |
| DG | 0.32 | 0.24 | 0.25 | 0.25 | 0.25 | 0.29 | 0.31 | 0.36 | 0.41 | 1.00 |
| CA3 | 0.49 | 0.41 | 0.40 | 0.39 | 0.41 | 0.41 | 0.45 | 0.45 | 0.55 | 1.00 |
| CA1 | 0.29 | 0.21 | 0.18 | 0.19 | 0.19 | 0.20 | 0.22 | 0.28 | 0.35 | 1.00 |
| Amy−Exc | 0.39 | 0.26 | 0.26 | 0.24 | 0.25 | 0.30 | 0.31 | 0.33 | 0.42 | 1.00 |
| Vip | 0.73 | 0.60 | 0.52 | 0.54 | 0.54 | 0.51 | 0.51 | 0.57 | 0.65 | 1.00 |
| THM−MB | 0.82 | 0.60 | 0.53 | 0.51 | 0.49 | 0.49 | 0.50 | 0.55 | 0.61 | 1.00 |
| THM−Inh | 6.44 | 3.64 | 3.15 | 2.57 | 2.48 | 2.26 | 1.97 | 1.75 | 1.44 | 1.00 |
| THM−Exc | 1.01 | 0.93 | 0.87 | 0.84 | 0.80 | 0.78 | 0.75 | 0.75 | 0.74 | 1.00 |
| SubCtx−Cplx | 0.41 | 0.33 | 0.33 | 0.33 | 0.36 | 0.38 | 0.35 | 0.39 | 0.50 | 1.00 |
| Sst | 0.59 | 0.45 | 0.40 | 0.38 | 0.39 | 0.40 | 0.42 | 0.44 | 0.49 | 1.00 |
| Sncg | 0.45 | 0.32 | 0.32 | 0.34 | 0.32 | 0.32 | 0.38 | 0.40 | 0.46 | 1.00 |
| Pvalb−ChC | 0.44 | 0.38 | 0.34 | 0.37 | 0.35 | 0.38 | 0.39 | 0.41 | 0.49 | 1.00 |
| Pvalb | 0.39 | 0.43 | 0.50 | 0.52 | 0.32 | 0.27 | 0.26 | 0.28 | 0.36 | 1.00 |
| PN | 0.66 | 0.41 | 0.32 | 0.33 | 0.32 | 0.29 | 0.32 | 0.38 | 0.46 | 1.00 |
| MSN−D2 | 0.38 | 0.31 | 0.28 | 0.29 | 0.28 | 0.30 | 0.32 | 0.36 | 0.42 | 1.00 |
| MSN−D1 | 0.45 | 0.33 | 0.31 | 0.29 | 0.31 | 0.31 | 0.34 | 0.34 | 0.43 | 1.00 |
| Lamp5−Lhx6 | 0.74 | 0.58 | 0.56 | 0.58 | 0.56 | 0.54 | 0.59 | 0.62 | 0.67 | 1.00 |
| Lamp5 | 0.56 | 0.44 | 0.41 | 0.43 | 0.41 | 0.44 | 0.45 | 0.49 | 0.53 | 1.00 |
| Foxp2 | 0.51 | 0.35 | 0.34 | 0.34 | 0.35 | 0.35 | 0.35 | 0.40 | 0.45 | 1.00 |
| Chd7 | 0.32 | 0.23 | 0.22 | 0.20 | 0.21 | 0.25 | 0.26 | 0.28 | 0.37 | 1.00 |
| CB | 5.59 | 4.08 | 3.69 | 3.54 | 3.07 | 2.88 | 2.50 | 2.02 | 1.71 | 1.00 |

Percent 0.002 0.004 0.006

#### SMC−Vascular smooth muscle cells

| | 0.0−0.1 | 0.1−0.2 | 0.2−0.3 | 0.3−0.4 | 0.4−0.5 | 0.5−0.6 | 0.6−0.7 | 0.7−0.8 | 0.8−0.9 | 0.9−1.0 |
|---|---|---|---|---|---|---|---|---|---|---|
| L6b | 0.77 | 0.63 | 0.52 | 0.50 | 0.48 | 0.46 | 0.46 | 0.50 | 0.56 | 1.00 |
| L6−IT−Car3 | 0.46 | 0.35 | 0.34 | 0.34 | 0.32 | 0.33 | 0.35 | 0.37 | 0.42 | 1.00 |
| L6−IT | 0.55 | 0.41 | 0.38 | 0.35 | 0.35 | 0.34 | 0.34 | 0.36 | 0.40 | 1.00 |
| L6−CT | 0.71 | 0.61 | 0.58 | 0.57 | 0.58 | 0.60 | 0.61 | 0.63 | 0.72 | 1.00 |
| L56−NP | 1.11 | 0.81 | 0.73 | 0.64 | 0.61 | 0.59 | 0.56 | 0.56 | 0.61 | 1.00 |
| L5−IT | 0.60 | 0.48 | 0.44 | 0.43 | 0.42 | 0.42 | 0.44 | 0.44 | 0.51 | 1.00 |
| L5−ET | 0.50 | 0.41 | 0.36 | 0.35 | 0.34 | 0.32 | 0.35 | 0.37 | 0.44 | 1.00 |
| L4−IT | 0.30 | 0.25 | 0.24 | 0.23 | 0.24 | 0.24 | 0.25 | 0.28 | 0.35 | 1.00 |
| L23−IT | 0.42 | 0.35 | 0.32 | 0.31 | 0.31 | 0.33 | 0.32 | 0.35 | 0.42 | 1.00 |
| HIP−Misc2 | 0.61 | 0.46 | 0.42 | 0.41 | 0.39 | 0.39 | 0.38 | 0.42 | 0.49 | 1.00 |
| HIP−Misc1 | 7.17 | 5.51 | 4.56 | 4.16 | 3.70 | 3.10 | 2.60 | 2.31 | 1.66 | 1.00 |
| DG | 0.37 | 0.32 | 0.32 | 0.31 | 0.30 | 0.33 | 0.33 | 0.37 | 0.44 | 1.00 |
| CA3 | 0.55 | 0.49 | 0.48 | 0.45 | 0.45 | 0.46 | 0.45 | 0.46 | 0.54 | 1.00 |
| CA1 | 0.34 | 0.28 | 0.24 | 0.25 | 0.24 | 0.25 | 0.26 | 0.30 | 0.38 | 1.00 |
| Amy−Exc | 0.43 | 0.36 | 0.35 | 0.34 | 0.34 | 0.34 | 0.35 | 0.36 | 0.42 | 1.00 |
| Vip | 0.89 | 0.72 | 0.66 | 0.64 | 0.62 | 0.59 | 0.58 | 0.61 | 0.65 | 1.00 |
| THM−MB | 0.94 | 0.76 | 0.71 | 0.67 | 0.62 | 0.59 | 0.57 | 0.61 | 0.64 | 1.00 |
| THM−Inh | 5.73 | 3.70 | 3.33 | 2.87 | 2.58 | 2.25 | 1.95 | 1.76 | 1.49 | 1.00 |
| THM−Exc | 1.41 | 1.24 | 1.13 | 1.05 | 0.98 | 0.91 | 0.85 | 0.82 | 0.77 | 1.00 |
| SubCtx−Cplx | 0.49 | 0.43 | 0.40 | 0.41 | 0.41 | 0.39 | 0.40 | 0.43 | 0.49 | 1.00 |
| Sst | 0.65 | 0.52 | 0.49 | 0.46 | 0.46 | 0.45 | 0.46 | 0.47 | 0.52 | 1.00 |
| Sncg | 0.56 | 0.47 | 0.44 | 0.41 | 0.42 | 0.41 | 0.42 | 0.43 | 0.49 | 1.00 |
| Pvalb−ChC | 0.54 | 0.47 | 0.44 | 0.42 | 0.41 | 0.41 | 0.41 | 0.43 | 0.50 | 1.00 |
| Pvalb | 0.44 | 0.37 | 0.40 | 0.45 | 0.39 | 0.33 | 0.31 | 0.33 | 0.38 | 1.00 |
| PN | 0.73 | 0.53 | 0.46 | 0.43 | 0.43 | 0.41 | 0.39 | 0.41 | 0.48 | 1.00 |
| MSN−D2 | 0.46 | 0.39 | 0.36 | 0.35 | 0.34 | 0.35 | 0.37 | 0.39 | 0.44 | 1.00 |
| MSN−D1 | 0.53 | 0.43 | 0.42 | 0.38 | 0.39 | 0.37 | 0.38 | 0.41 | 0.47 | 1.00 |
| Lamp5−Lhx6 | 0.82 | 0.70 | 0.66 | 0.62 | 0.62 | 0.61 | 0.61 | 0.62 | 0.68 | 1.00 |
| Lamp5 | 0.63 | 0.54 | 0.51 | 0.50 | 0.49 | 0.47 | 0.49 | 0.50 | 0.58 | 1.00 |
| Foxp2 | 0.55 | 0.43 | 0.43 | 0.41 | 0.40 | 0.41 | 0.42 | 0.44 | 0.48 | 1.00 |
| Chd7 | 0.40 | 0.33 | 0.30 | 0.29 | 0.29 | 0.30 | 0.30 | 0.32 | 0.38 | 1.00 |
| CB | 5.19 | 3.96 | 3.60 | 3.53 | 3.08 | 2.73 | 2.45 | 2.08 | 1.71 | 1.00 |

Percent 0.005 0.010 0.015 0.020

#### SNCG−SNCG+ GABAergic neurons

| | 0.0−0.1 | 0.1−0.2 | 0.2−0.3 | 0.3−0.4 | 0.4−0.5 | 0.5−0.6 | 0.6−0.7 | 0.7−0.8 | 0.8−0.9 | 0.9−1.0 |
|---|---|---|---|---|---|---|---|---|---|---|
| L6b | 1.46 | 1.03 | 0.92 | 0.85 | 0.80 | 0.76 | 0.77 | 0.77 | 0.79 | 1.00 |
| L6−IT−Car3 | 0.93 | 0.71 | 0.67 | 0.64 | 0.62 | 0.60 | 0.61 | 0.60 | 0.62 | 1.00 |
| L6−IT | 1.06 | 0.77 | 0.70 | 0.67 | 0.64 | 0.62 | 0.59 | 0.58 | 0.61 | 1.00 |
| L6−CT | 1.32 | 1.09 | 1.03 | 0.98 | 0.97 | 0.94 | 0.97 | 0.91 | 0.93 | 1.00 |
| L56−NP | 1.55 | 1.22 | 1.12 | 1.02 | 0.99 | 0.93 | 0.90 | 0.89 | 0.86 | 1.00 |
| L5−IT | 1.14 | 0.84 | 0.79 | 0.76 | 0.73 | 0.72 | 0.71 | 0.69 | 0.70 | 1.00 |
| L5−ET | 1.20 | 0.80 | 0.73 | 0.67 | 0.64 | 0.61 | 0.60 | 0.59 | 0.63 | 1.00 |
| L4−IT | 0.70 | 0.56 | 0.54 | 0.51 | 0.49 | 0.49 | 0.49 | 0.49 | 0.53 | 1.00 |
| L23−IT | 0.94 | 0.69 | 0.65 | 0.61 | 0.59 | 0.59 | 0.57 | 0.58 | 0.61 | 1.00 |
| HIP−Misc2 | 1.11 | 0.85 | 0.77 | 0.75 | 0.71 | 0.68 | 0.67 | 0.66 | 0.67 | 1.00 |
| HIP−Misc1 | 2.84 | 2.34 | 2.15 | 2.03 | 1.91 | 1.82 | 1.66 | 1.53 | 1.32 | 1.00 |
| DG | 0.88 | 0.68 | 0.62 | 0.60 | 0.60 | 0.62 | 0.61 | 0.61 | 0.63 | 1.00 |
| CA3 | 0.93 | 0.83 | 0.82 | 0.80 | 0.80 | 0.76 | 0.76 | 0.75 | 0.77 | 1.00 |
| CA1 | 0.85 | 0.60 | 0.54 | 0.52 | 0.50 | 0.49 | 0.49 | 0.50 | 0.53 | 1.00 |
| Amy−Exc | 0.92 | 0.74 | 0.69 | 0.67 | 0.65 | 0.64 | 0.61 | 0.61 | 0.64 | 1.00 |
| Vip | 1.48 | 1.22 | 1.12 | 1.08 | 1.04 | 1.00 | 0.94 | 0.90 | 0.87 | 1.00 |
| THM−MB | 1.57 | 1.18 | 1.09 | 1.01 | 0.98 | 0.93 | 0.86 | 0.85 | 0.84 | 1.00 |
| THM−Inh | 2.65 | 2.00 | 1.88 | 1.77 | 1.66 | 1.60 | 1.51 | 1.42 | 1.26 | 1.00 |
| THM−Exc | 1.34 | 1.24 | 1.18 | 1.15 | 1.11 | 1.08 | 1.04 | 1.01 | 0.96 | 1.00 |
| SubCtx−Cplx | 0.85 | 0.81 | 0.75 | 0.75 | 0.76 | 0.75 | 0.73 | 0.71 | 0.73 | 1.00 |
| Sst | 1.16 | 0.97 | 0.89 | 0.84 | 0.81 | 0.78 | 0.78 | 0.73 | 0.73 | 1.00 |
| Sncg | 1.11 | 0.93 | 0.86 | 0.83 | 0.80 | 0.75 | 0.75 | 0.71 | 0.70 | 1.00 |
| Pvalb−ChC | 1.05 | 0.87 | 0.81 | 0.76 | 0.75 | 0.72 | 0.71 | 0.69 | 0.70 | 1.00 |
| Pvalb | 0.76 | 0.71 | 0.71 | 0.69 | 0.70 | 0.63 | 0.61 | 0.58 | 0.60 | 1.00 |
| PN | 1.53 | 0.93 | 0.81 | 0.72 | 0.70 | 0.68 | 0.66 | 0.67 | 0.67 | 1.00 |
| MSN−D2 | 0.94 | 0.80 | 0.75 | 0.72 | 0.68 | 0.67 | 0.67 | 0.65 | 0.66 | 1.00 |
| MSN−D1 | 0.95 | 0.79 | 0.75 | 0.72 | 0.71 | 0.70 | 0.68 | 0.68 | 0.67 | 1.00 |
| Lamp5−Lhx6 | 1.40 | 1.15 | 1.09 | 1.05 | 1.02 | 0.97 | 0.96 | 0.90 | 0.91 | 1.00 |
| Lamp5 | 1.26 | 1.01 | 0.92 | 0.90 | 0.88 | 0.85 | 0.81 | 0.79 | 0.79 | 1.00 |
| Foxp2 | 1.02 | 0.84 | 0.81 | 0.76 | 0.76 | 0.75 | 0.72 | 0.70 | 0.71 | 1.00 |
| Chd7 | 1.00 | 0.73 | 0.66 | 0.62 | 0.59 | 0.56 | 0.55 | 0.55 | 0.58 | 1.00 |
| CB | 3.34 | 2.72 | 2.60 | 2.46 | 2.28 | 2.13 | 1.98 | 1.70 | 1.50 | 1.00 |

Percent 0.02 0.04 0.06

SST−SST+ GABAergic neurons

| | 0.0−0.1 | 0.1−0.2 | 0.2−0.3 | 0.3−0.4 | 0.4−0.5 | 0.5−0.6 | 0.6−0.7 | 0.7−0.8 | 0.8−0.9 | 0.9−1.0 |
|---|---|---|---|---|---|---|---|---|---|---|
| L6b | 1.76 | 1.25 | 1.10 | 1.01 | 0.94 | 0.89 | 0.87 | 0.84 | 0.82 | 1.00 |
| L6−IT−Car3 | 1.24 | 0.90 | 0.84 | 0.79 | 0.77 | 0.73 | 0.72 | 0.70 | 0.70 | 1.00 |
| L6−IT | 1.34 | 0.95 | 0.89 | 0.82 | 0.79 | 0.75 | 0.72 | 0.70 | 0.68 | 1.00 |
| L6−CT | 1.65 | 1.34 | 1.26 | 1.20 | 1.17 | 1.12 | 1.11 | 1.03 | 0.99 | 1.00 |
| L56−NP | 1.90 | 1.47 | 1.34 | 1.25 | 1.17 | 1.12 | 1.06 | 0.98 | 0.93 | 1.00 |
| L5−IT | 1.46 | 1.03 | 0.96 | 0.92 | 0.86 | 0.84 | 0.80 | 0.78 | 0.76 | 1.00 |
| L5−ET | 1.49 | 1.00 | 0.90 | 0.84 | 0.79 | 0.72 | 0.70 | 0.67 | 0.69 | 1.00 |
| L4−IT | 0.91 | 0.71 | 0.67 | 0.63 | 0.60 | 0.60 | 0.57 | 0.56 | 0.58 | 1.00 |
| L23−IT | 1.19 | 0.88 | 0.81 | 0.76 | 0.73 | 0.71 | 0.67 | 0.66 | 0.66 | 1.00 |
| HIP−Misc2 | 1.41 | 1.04 | 0.94 | 0.91 | 0.86 | 0.81 | 0.79 | 0.76 | 0.76 | 1.00 |
| HIP−Misc1 | 2.42 | 2.09 | 2.02 | 1.92 | 1.80 | 1.72 | 1.62 | 1.50 | 1.34 | 1.00 |
| DG | 1.04 | 0.82 | 0.77 | 0.75 | 0.74 | 0.72 | 0.71 | 0.70 | 0.70 | 1.00 |
| CA3 | 1.05 | 0.96 | 0.94 | 0.93 | 0.89 | 0.89 | 0.88 | 0.85 | 0.83 | 1.00 |
| CA1 | 1.10 | 0.75 | 0.69 | 0.64 | 0.62 | 0.59 | 0.57 | 0.58 | 0.62 | 1.00 |
| Amy−Exc | 1.14 | 0.90 | 0.85 | 0.80 | 0.77 | 0.75 | 0.74 | 0.70 | 0.70 | 1.00 |
| Vip | 1.63 | 1.41 | 1.34 | 1.29 | 1.22 | 1.15 | 1.10 | 1.04 | 0.98 | 1.00 |
| THM−MB | 1.86 | 1.47 | 1.30 | 1.25 | 1.18 | 1.10 | 1.02 | 0.98 | 0.93 | 1.00 |
| THM−Inh | 2.33 | 1.88 | 1.81 | 1.71 | 1.67 | 1.61 | 1.51 | 1.45 | 1.30 | 1.00 |
| THM−Exc | 1.42 | 1.32 | 1.27 | 1.23 | 1.20 | 1.16 | 1.13 | 1.08 | 1.03 | 1.00 |
| SubCtx−Cplx | 1.00 | 0.95 | 0.93 | 0.92 | 0.91 | 0.89 | 0.86 | 0.85 | 0.82 | 1.00 |
| Sst | 1.46 | 1.19 | 1.09 | 1.04 | 0.99 | 0.97 | 0.92 | 0.86 | 0.83 | 1.00 |
| Sncg | 1.23 | 1.12 | 1.06 | 1.01 | 0.97 | 0.94 | 0.89 | 0.85 | 0.80 | 1.00 |
| Pvalb−ChC | 1.24 | 1.07 | 0.99 | 0.94 | 0.91 | 0.87 | 0.83 | 0.81 | 0.78 | 1.00 |
| Pvalb | 0.91 | 0.84 | 0.79 | 0.78 | 0.84 | 0.80 | 0.77 | 0.72 | 0.70 | 1.00 |
| PN | 1.76 | 1.17 | 1.01 | 0.93 | 0.87 | 0.83 | 0.81 | 0.77 | 0.75 | 1.00 |
| MSN−D2 | 1.19 | 1.02 | 0.95 | 0.92 | 0.86 | 0.83 | 0.80 | 0.77 | 0.75 | 1.00 |
| MSN−D1 | 1.20 | 1.00 | 0.95 | 0.89 | 0.87 | 0.84 | 0.81 | 0.78 | 0.76 | 1.00 |
| Lamp5−Lhx6 | 1.49 | 1.34 | 1.27 | 1.23 | 1.19 | 1.15 | 1.12 | 1.04 | 0.99 | 1.00 |
| Lamp5 | 1.36 | 1.18 | 1.11 | 1.08 | 1.06 | 1.00 | 0.95 | 0.91 | 0.90 | 1.00 |
| Foxp2 | 1.19 | 1.01 | 0.96 | 0.94 | 0.92 | 0.88 | 0.85 | 0.81 | 0.79 | 1.00 |
| Chd7 | 1.13 | 0.89 | 0.84 | 0.79 | 0.74 | 0.73 | 0.69 | 0.65 | 0.66 | 1.00 |
| CB | 2.83 | 2.44 | 2.32 | 2.26 | 2.16 | 1.99 | 1.85 | 1.67 | 1.43 | 1.00 |

Percent 0.025 0.050 0.075 0.100

SST_CHODL−SST+ GABAergic neurons with CHODL+

| | 0.0−0.1 | 0.1−0.2 | 0.2−0.3 | 0.3−0.4 | 0.4−0.5 | 0.5−0.6 | 0.6−0.7 | 0.7−0.8 | 0.8−0.9 | 0.9−1.0 |
|---|---|---|---|---|---|---|---|---|---|---|
| L6b | 1.24 | 0.85 | 0.75 | 0.70 | 0.65 | 0.62 | 0.62 | 0.66 | 0.68 | 1.00 |
| L6−IT−Car3 | 0.76 | 0.52 | 0.50 | 0.48 | 0.49 | 0.46 | 0.47 | 0.49 | 0.53 | 1.00 |
| L6−IT | 0.87 | 0.60 | 0.56 | 0.50 | 0.50 | 0.48 | 0.48 | 0.48 | 0.51 | 1.00 |
| L6−CT | 1.08 | 0.86 | 0.85 | 0.81 | 0.78 | 0.79 | 0.81 | 0.77 | 0.84 | 1.00 |
| L56−NP | 1.46 | 1.05 | 0.93 | 0.85 | 0.82 | 0.79 | 0.77 | 0.76 | 0.74 | 1.00 |
| L5−IT | 0.95 | 0.67 | 0.62 | 0.59 | 0.59 | 0.57 | 0.58 | 0.57 | 0.62 | 1.00 |
| L5−ET | 0.91 | 0.62 | 0.55 | 0.52 | 0.50 | 0.46 | 0.48 | 0.47 | 0.55 | 1.00 |
| L4−IT | 0.53 | 0.41 | 0.39 | 0.36 | 0.36 | 0.36 | 0.36 | 0.38 | 0.44 | 1.00 |
| L23−IT | 0.72 | 0.52 | 0.48 | 0.47 | 0.44 | 0.45 | 0.45 | 0.46 | 0.50 | 1.00 |
| HIP−Misc2 | 0.94 | 0.66 | 0.61 | 0.59 | 0.54 | 0.54 | 0.53 | 0.52 | 0.59 | 1.00 |
| HIP−Misc1 | 3.84 | 3.01 | 2.64 | 2.40 | 2.26 | 2.07 | 1.91 | 1.72 | 1.49 | 1.00 |
| DG | 0.63 | 0.49 | 0.47 | 0.46 | 0.44 | 0.48 | 0.47 | 0.50 | 0.55 | 1.00 |
| CA3 | 0.74 | 0.66 | 0.62 | 0.61 | 0.62 | 0.62 | 0.60 | 0.61 | 0.67 | 1.00 |
| CA1 | 0.62 | 0.44 | 0.39 | 0.37 | 0.36 | 0.36 | 0.37 | 0.39 | 0.45 | 1.00 |
| Amy−Exc | 0.72 | 0.54 | 0.51 | 0.50 | 0.49 | 0.48 | 0.50 | 0.50 | 0.53 | 1.00 |
| Vip | 1.20 | 0.97 | 0.93 | 0.89 | 0.86 | 0.84 | 0.81 | 0.78 | 0.79 | 1.00 |
| THM−MB | 1.35 | 0.99 | 0.92 | 0.85 | 0.81 | 0.78 | 0.75 | 0.75 | 0.76 | 1.00 |
| THM−Inh | 3.51 | 2.37 | 2.20 | 1.97 | 1.91 | 1.78 | 1.63 | 1.51 | 1.34 | 1.00 |
| THM−Exc | 1.30 | 1.17 | 1.11 | 1.07 | 1.04 | 0.99 | 0.96 | 0.93 | 0.89 | 1.00 |
| SubCtx−Cplx | 0.69 | 0.61 | 0.58 | 0.59 | 0.59 | 0.57 | 0.59 | 0.60 | 0.63 | 1.00 |
| Sst | 1.02 | 0.78 | 0.72 | 0.70 | 0.67 | 0.65 | 0.64 | 0.62 | 0.65 | 1.00 |
| Sncg | 0.85 | 0.67 | 0.63 | 0.65 | 0.61 | 0.59 | 0.60 | 0.60 | 0.61 | 1.00 |
| Pvalb−ChC | 0.84 | 0.69 | 0.62 | 0.61 | 0.60 | 0.60 | 0.57 | 0.57 | 0.60 | 1.00 |
| Pvalb | 0.62 | 0.59 | 0.59 | 0.62 | 0.55 | 0.48 | 0.48 | 0.46 | 0.50 | 1.00 |
| PN | 1.21 | 0.75 | 0.64 | 0.60 | 0.57 | 0.56 | 0.55 | 0.55 | 0.60 | 1.00 |
| MSN−D2 | 0.71 | 0.58 | 0.57 | 0.55 | 0.52 | 0.52 | 0.53 | 0.53 | 0.56 | 1.00 |
| MSN−D1 | 0.78 | 0.62 | 0.57 | 0.55 | 0.55 | 0.54 | 0.55 | 0.55 | 0.58 | 1.00 |
| Lamp5−Lhx6 | 1.14 | 0.94 | 0.90 | 0.86 | 0.87 | 0.83 | 0.81 | 0.79 | 0.83 | 1.00 |
| Lamp5 | 0.97 | 0.75 | 0.72 | 0.70 | 0.72 | 0.67 | 0.66 | 0.66 | 0.70 | 1.00 |
| Foxp2 | 0.83 | 0.63 | 0.61 | 0.60 | 0.61 | 0.59 | 0.58 | 0.59 | 0.61 | 1.00 |
| Chd7 | 0.72 | 0.52 | 0.47 | 0.46 | 0.44 | 0.45 | 0.44 | 0.44 | 0.49 | 1.00 |
| CB | 3.80 | 3.09 | 2.87 | 2.74 | 2.57 | 2.34 | 2.13 | 1.91 | 1.59 | 1.00 |

Percent 0.01 0.02 0.03 0.04

SUB−Granule neurons from subicular cortex

| | 0.0−0.1 | 0.1−0.2 | 0.2−0.3 | 0.3−0.4 | 0.4−0.5 | 0.5−0.6 | 0.6−0.7 | 0.7−0.8 | 0.8−0.9 | 0.9−1.0 |
|---|---|---|---|---|---|---|---|---|---|---|
| L6b | 0.73 | 0.52 | 0.49 | 0.47 | 0.40 | 0.41 | 0.42 | 0.47 | 0.54 | 1.00 |
| L6−IT−Car3 | 0.39 | 0.27 | 0.27 | 0.27 | 0.23 | 0.25 | 0.26 | 0.31 | 0.43 | 1.00 |
| L6−IT | 0.52 | 0.36 | 0.30 | 0.27 | 0.27 | 0.25 | 0.31 | 0.33 | 0.40 | 1.00 |
| L6−CT | 0.56 | 0.43 | 0.46 | 0.46 | 0.46 | 0.46 | 0.56 | 0.56 | 0.72 | 1.00 |
| L56−NP | 0.94 | 0.69 | 0.56 | 0.54 | 0.52 | 0.50 | 0.47 | 0.49 | 0.58 | 1.00 |
| L5−IT | 0.51 | 0.39 | 0.37 | 0.35 | 0.36 | 0.35 | 0.37 | 0.46 | 0.53 | 1.00 |
| L5−ET | 0.44 | 0.35 | 0.33 | 0.29 | 0.30 | 0.26 | 0.29 | 0.34 | 0.41 | 1.00 |
| L4−IT | 0.29 | 0.21 | 0.19 | 0.19 | 0.19 | 0.21 | 0.22 | 0.27 | 0.34 | 1.00 |
| L23−IT | 0.42 | 0.29 | 0.27 | 0.26 | 0.26 | 0.28 | 0.31 | 0.35 | 0.45 | 1.00 |
| HIP−Misc2 | 0.54 | 0.41 | 0.32 | 0.32 | 0.31 | 0.31 | 0.34 | 0.38 | 0.45 | 1.00 |
| HIP−Misc1 | 7.96 | 6.02 | 4.61 | 4.06 | 3.76 | 3.01 | 2.68 | 2.27 | 1.51 | 1.00 |
| DG | 0.34 | 0.27 | 0.28 | 0.26 | 0.24 | 0.30 | 0.30 | 0.38 | 0.41 | 1.00 |
| CA3 | 0.53 | 0.47 | 0.46 | 0.45 | 0.43 | 0.43 | 0.45 | 0.52 | 0.53 | 1.00 |
| CA1 | 0.32 | 0.21 | 0.19 | 0.21 | 0.21 | 0.20 | 0.23 | 0.26 | 0.34 | 1.00 |
| Amy−Exc | 0.41 | 0.27 | 0.27 | 0.26 | 0.27 | 0.28 | 0.30 | 0.31 | 0.40 | 1.00 |
| Vip | 0.82 | 0.62 | 0.60 | 0.58 | 0.58 | 0.56 | 0.58 | 0.68 | 0.78 | 1.00 |
| THM−MB | 0.79 | 0.65 | 0.61 | 0.60 | 0.56 | 0.55 | 0.53 | 0.56 | 0.73 | 1.00 |
| THM−Inh | 6.84 | 3.96 | 3.35 | 2.82 | 2.61 | 2.45 | 2.12 | 1.65 | 1.36 | 1.00 |
| THM−Exc | 0.90 | 0.84 | 0.82 | 0.77 | 0.79 | 0.75 | 0.73 | 0.77 | 0.75 | 1.00 |
| SubCtx−Cplx | 0.44 | 0.40 | 0.36 | 0.39 | 0.34 | 0.41 | 0.42 | 0.43 | 0.50 | 1.00 |
| Sst | 0.60 | 0.49 | 0.43 | 0.42 | 0.41 | 0.42 | 0.42 | 0.43 | 0.54 | 1.00 |
| Sncg | 0.45 | 0.36 | 0.36 | 0.36 | 0.35 | 0.38 | 0.41 | 0.44 | 0.51 | 1.00 |
| Pvalb−ChC | 0.45 | 0.41 | 0.43 | 0.38 | 0.37 | 0.44 | 0.44 | 0.45 | 0.49 | 1.00 |
| Pvalb | 0.33 | 0.34 | 0.45 | 0.46 | 0.34 | 0.25 | 0.27 | 0.28 | 0.32 | 1.00 |
| PN | 0.75 | 0.44 | 0.41 | 0.31 | 0.30 | 0.35 | 0.36 | 0.40 | 0.45 | 1.00 |
| MSN−D2 | 0.37 | 0.32 | 0.29 | 0.31 | 0.31 | 0.30 | 0.34 | 0.40 | 0.42 | 1.00 |
| MSN−D1 | 0.42 | 0.33 | 0.32 | 0.34 | 0.30 | 0.33 | 0.35 | 0.39 | 0.43 | 1.00 |
| Lamp5−Lhx6 | 0.81 | 0.69 | 0.56 | 0.62 | 0.64 | 0.66 | 0.59 | 0.64 | 0.68 | 1.00 |
| Lamp5 | 0.62 | 0.51 | 0.43 | 0.48 | 0.42 | 0.51 | 0.46 | 0.56 | 0.55 | 1.00 |
| Foxp2 | 0.53 | 0.42 | 0.33 | 0.36 | 0.39 | 0.38 | 0.44 | 0.42 | 0.50 | 1.00 |
| Chd7 | 0.31 | 0.26 | 0.23 | 0.21 | 0.24 | 0.23 | 0.24 | 0.30 | 0.40 | 1.00 |
| CB | 5.12 | 4.18 | 3.53 | 3.22 | 2.83 | 2.65 | 2.38 | 2.02 | 1.51 | 1.00 |

Percent 0.001 0.002 0.003

#### THMGA−GABAergic neurons from thalamus

| | 0.0−0.1 | 0.1−0.2 | 0.2−0.3 | 0.3−0.4 | 0.4−0.5 | 0.5−0.6 | 0.6−0.7 | 0.7−0.8 | 0.8−0.9 | 0.9−1.0 |
|---|---|---|---|---|---|---|---|---|---|---|
| L6b | 0.85 | 0.67 | 0.60 | 0.57 | 0.54 | 0.54 | 0.56 | 0.57 | 0.63 | 1.00 |
| L6−IT−Car3 | 0.56 | 0.43 | 0.40 | 0.39 | 0.41 | 0.40 | 0.42 | 0.43 | 0.48 | 1.00 |
| L6−IT | 0.63 | 0.48 | 0.45 | 0.41 | 0.42 | 0.43 | 0.41 | 0.43 | 0.48 | 1.00 |
| L6−CT | 0.78 | 0.67 | 0.70 | 0.65 | 0.67 | 0.66 | 0.70 | 0.69 | 0.78 | 1.00 |
| L56−NP | 1.12 | 0.83 | 0.78 | 0.71 | 0.70 | 0.70 | 0.67 | 0.65 | 0.68 | 1.00 |
| L5−IT | 0.71 | 0.54 | 0.53 | 0.50 | 0.50 | 0.50 | 0.52 | 0.52 | 0.58 | 1.00 |
| L5−ET | 0.67 | 0.49 | 0.46 | 0.43 | 0.42 | 0.40 | 0.41 | 0.45 | 0.49 | 1.00 |
| L4−IT | 0.42 | 0.33 | 0.32 | 0.32 | 0.29 | 0.31 | 0.31 | 0.34 | 0.41 | 1.00 |
| L23−IT | 0.56 | 0.42 | 0.40 | 0.39 | 0.39 | 0.40 | 0.39 | 0.43 | 0.47 | 1.00 |
| HIP−Misc2 | 0.69 | 0.55 | 0.50 | 0.49 | 0.47 | 0.45 | 0.48 | 0.45 | 0.55 | 1.00 |
| HIP−Misc1 | 4.71 | 3.67 | 3.22 | 3.04 | 2.81 | 2.53 | 2.12 | 1.96 | 1.52 | 1.00 |
| DG | 0.50 | 0.41 | 0.40 | 0.38 | 0.39 | 0.42 | 0.42 | 0.44 | 0.51 | 1.00 |
| CA3 | 0.68 | 0.61 | 0.55 | 0.59 | 0.58 | 0.58 | 0.57 | 0.59 | 0.64 | 1.00 |
| CA1 | 0.45 | 0.34 | 0.31 | 0.32 | 0.31 | 0.32 | 0.33 | 0.36 | 0.41 | 1.00 |
| Amy−Exc | 0.54 | 0.43 | 0.42 | 0.42 | 0.41 | 0.42 | 0.43 | 0.42 | 0.49 | 1.00 |
| Vip | 0.98 | 0.84 | 0.75 | 0.74 | 0.74 | 0.73 | 0.71 | 0.69 | 0.77 | 1.00 |
| THM−MB | 1.52 | 0.91 | 0.78 | 0.71 | 0.66 | 0.64 | 0.61 | 0.64 | 0.69 | 1.00 |
| THM−Inh | 4.44 | 2.94 | 2.68 | 2.48 | 2.29 | 2.10 | 1.97 | 1.75 | 1.48 | 1.00 |
| THM−Exc | 1.14 | 1.05 | 1.02 | 0.98 | 0.95 | 0.92 | 0.89 | 0.88 | 0.84 | 1.00 |
| SubCtx−Cplx | 0.58 | 0.53 | 0.50 | 0.50 | 0.51 | 0.49 | 0.51 | 0.52 | 0.59 | 1.00 |
| Sst | 0.80 | 0.63 | 0.61 | 0.60 | 0.57 | 0.55 | 0.55 | 0.57 | 0.62 | 1.00 |
| Sncg | 0.64 | 0.54 | 0.50 | 0.50 | 0.51 | 0.50 | 0.50 | 0.51 | 0.55 | 1.00 |
| Pvalb−ChC | 0.64 | 0.55 | 0.51 | 0.53 | 0.52 | 0.51 | 0.50 | 0.49 | 0.55 | 1.00 |
| Pvalb | 0.52 | 0.52 | 0.59 | 0.59 | 0.45 | 0.40 | 0.41 | 0.41 | 0.44 | 1.00 |
| PN | 0.89 | 0.60 | 0.52 | 0.49 | 0.50 | 0.49 | 0.47 | 0.50 | 0.52 | 1.00 |
| MSN−D2 | 0.58 | 0.48 | 0.45 | 0.45 | 0.44 | 0.44 | 0.45 | 0.48 | 0.51 | 1.00 |
| MSN−D1 | 0.60 | 0.51 | 0.47 | 0.45 | 0.46 | 0.47 | 0.46 | 0.47 | 0.52 | 1.00 |
| Lamp5−Lhx6 | 0.95 | 0.80 | 0.78 | 0.78 | 0.76 | 0.73 | 0.75 | 0.74 | 0.77 | 1.00 |
| Lamp5 | 0.80 | 0.64 | 0.62 | 0.63 | 0.59 | 0.60 | 0.61 | 0.61 | 0.66 | 1.00 |
| Foxp2 | 0.67 | 0.54 | 0.52 | 0.51 | 0.51 | 0.50 | 0.50 | 0.53 | 0.57 | 1.00 |
| Chd7 | 0.51 | 0.40 | 0.37 | 0.37 | 0.36 | 0.36 | 0.36 | 0.36 | 0.45 | 1.00 |
| CB | 4.27 | 3.42 | 3.25 | 3.10 | 2.95 | 2.58 | 2.43 | 1.96 | 1.66 | 1.00 |

Percent 0.005 0.010 0.015

#### VIP−VIP+ GABAergic neurons

| | 0.0−0.1 | 0.1−0.2 | 0.2−0.3 | 0.3−0.4 | 0.4−0.5 | 0.5−0.6 | 0.6−0.7 | 0.7−0.8 | 0.8−0.9 | 0.9−1.0 |
|---|---|---|---|---|---|---|---|---|---|---|
| L6b | 1.87 | 1.37 | 1.22 | 1.13 | 1.03 | 0.97 | 0.92 | 0.89 | 0.87 | 1.00 |
| L6−IT−Car3 | 1.32 | 1.02 | 0.94 | 0.88 | 0.85 | 0.80 | 0.77 | 0.73 | 0.71 | 1.00 |
| L6−IT | 1.44 | 1.05 | 0.97 | 0.91 | 0.87 | 0.81 | 0.76 | 0.72 | 0.70 | 1.00 |
| L6−CT | 1.80 | 1.47 | 1.37 | 1.29 | 1.23 | 1.18 | 1.12 | 1.05 | 0.99 | 1.00 |
| L56−NP | 2.05 | 1.61 | 1.44 | 1.37 | 1.29 | 1.20 | 1.10 | 1.01 | 0.96 | 1.00 |
| L5−IT | 1.58 | 1.17 | 1.08 | 1.02 | 0.97 | 0.92 | 0.87 | 0.83 | 0.80 | 1.00 |
| L5−ET | 1.59 | 1.11 | 0.99 | 0.94 | 0.86 | 0.80 | 0.78 | 0.73 | 0.71 | 1.00 |
| L4−IT | 1.00 | 0.80 | 0.75 | 0.70 | 0.67 | 0.65 | 0.61 | 0.59 | 0.60 | 1.00 |
| L23−IT | 1.26 | 0.97 | 0.89 | 0.84 | 0.79 | 0.76 | 0.72 | 0.69 | 0.69 | 1.00 |
| HIP−Misc2 | 1.53 | 1.17 | 1.05 | 1.00 | 0.93 | 0.88 | 0.85 | 0.79 | 0.77 | 1.00 |
| HIP−Misc1 | 2.75 | 2.36 | 2.23 | 2.13 | 2.03 | 1.90 | 1.73 | 1.59 | 1.38 | 1.00 |
| DG | 1.23 | 0.90 | 0.85 | 0.79 | 0.78 | 0.78 | 0.74 | 0.73 | 0.73 | 1.00 |
| CA3 | 1.21 | 1.10 | 1.05 | 1.03 | 0.98 | 0.95 | 0.92 | 0.87 | 0.85 | 1.00 |
| CA1 | 1.18 | 0.86 | 0.77 | 0.71 | 0.68 | 0.65 | 0.62 | 0.60 | 0.61 | 1.00 |
| Amy−Exc | 1.28 | 1.03 | 0.94 | 0.90 | 0.87 | 0.82 | 0.79 | 0.75 | 0.72 | 1.00 |
| Vip | 2.03 | 1.64 | 1.54 | 1.44 | 1.34 | 1.25 | 1.17 | 1.09 | 1.00 | 1.00 |
| THM−MB | 2.13 | 1.65 | 1.46 | 1.35 | 1.26 | 1.19 | 1.10 | 1.00 | 0.94 | 1.00 |
| THM−Inh | 2.66 | 2.19 | 2.08 | 1.94 | 1.88 | 1.77 | 1.68 | 1.56 | 1.36 | 1.00 |
| THM−Exc | 1.54 | 1.42 | 1.36 | 1.31 | 1.26 | 1.21 | 1.16 | 1.10 | 1.04 | 1.00 |
| SubCtx−Cplx | 1.16 | 1.07 | 1.02 | 1.01 | 1.00 | 0.96 | 0.91 | 0.87 | 0.83 | 1.00 |
| Sst | 1.55 | 1.31 | 1.21 | 1.14 | 1.08 | 1.03 | 0.96 | 0.91 | 0.86 | 1.00 |
| Sncg | 1.53 | 1.30 | 1.21 | 1.13 | 1.06 | 0.99 | 0.93 | 0.87 | 0.82 | 1.00 |
| Pvalb−ChC | 1.46 | 1.22 | 1.10 | 1.05 | 1.00 | 0.95 | 0.89 | 0.85 | 0.81 | 1.00 |
| Pvalb | 0.98 | 0.89 | 0.83 | 0.80 | 0.96 | 0.87 | 0.83 | 0.76 | 0.72 | 1.00 |
| PN | 2.21 | 1.35 | 1.15 | 1.02 | 0.93 | 0.89 | 0.84 | 0.80 | 0.77 | 1.00 |
| MSN−D2 | 1.36 | 1.13 | 1.05 | 0.98 | 0.93 | 0.88 | 0.84 | 0.79 | 0.74 | 1.00 |
| MSN−D1 | 1.32 | 1.11 | 1.05 | 0.98 | 0.95 | 0.91 | 0.87 | 0.83 | 0.78 | 1.00 |
| Lamp5−Lhx6 | 1.72 | 1.49 | 1.41 | 1.34 | 1.28 | 1.23 | 1.15 | 1.08 | 1.01 | 1.00 |
| Lamp5 | 1.57 | 1.31 | 1.21 | 1.16 | 1.13 | 1.05 | 0.99 | 0.94 | 0.89 | 1.00 |
| Foxp2 | 1.35 | 1.15 | 1.08 | 1.04 | 1.00 | 0.96 | 0.91 | 0.85 | 0.82 | 1.00 |
| Chd7 | 1.38 | 1.04 | 0.92 | 0.86 | 0.80 | 0.76 | 0.71 | 0.67 | 0.66 | 1.00 |
| CB | 3.39 | 2.84 | 2.75 | 2.56 | 2.40 | 2.22 | 2.03 | 1.75 | 1.49 | 1.00 |

Percent 0.03 0.06 0.09
