## Supplementary figures and images for "Distinct cellular DNA methylation mechanisms underlie common and rare genetic risk for brain disorders"

### Supplementary Figure 1

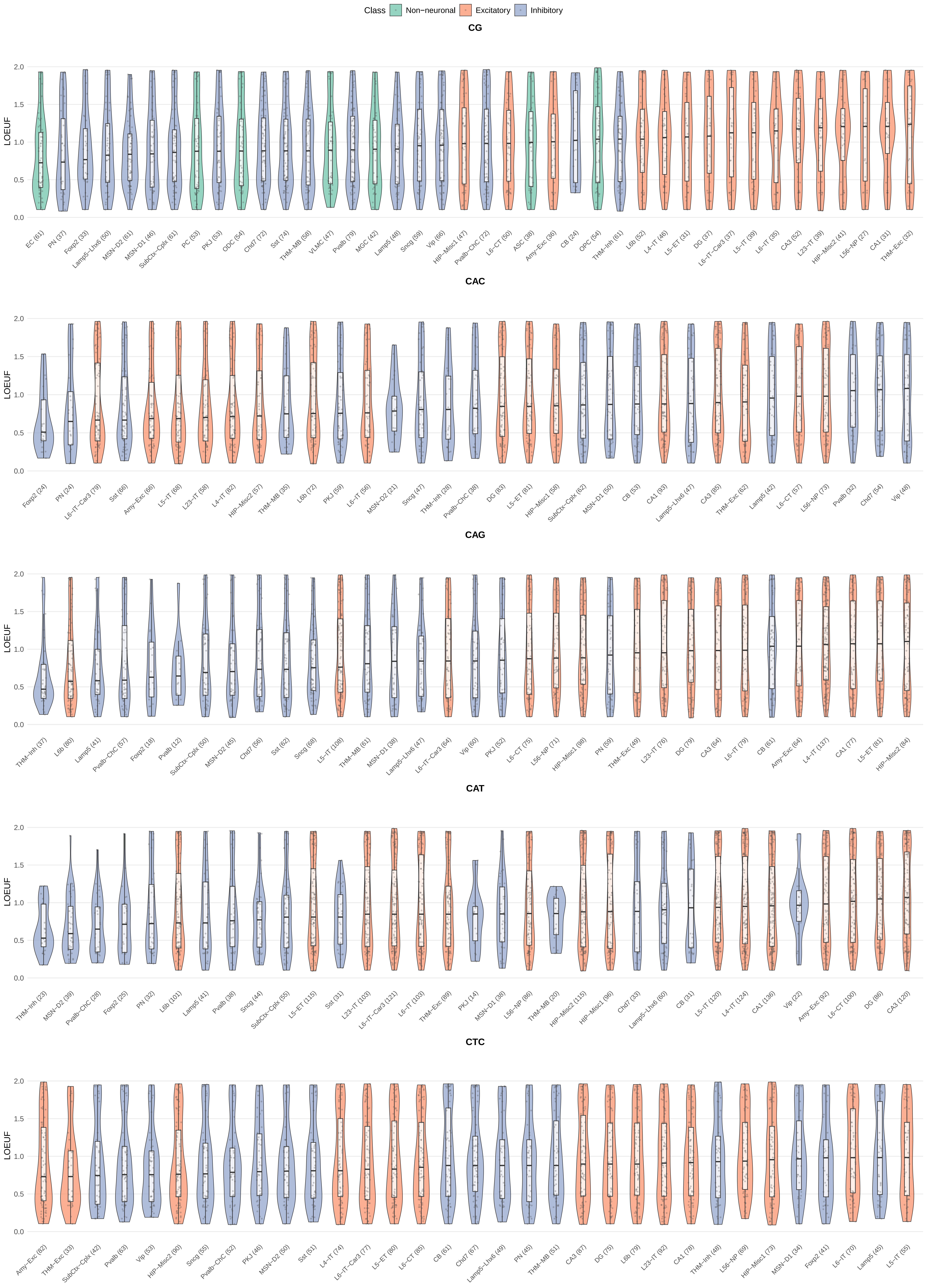

### Supplementary Figure 2

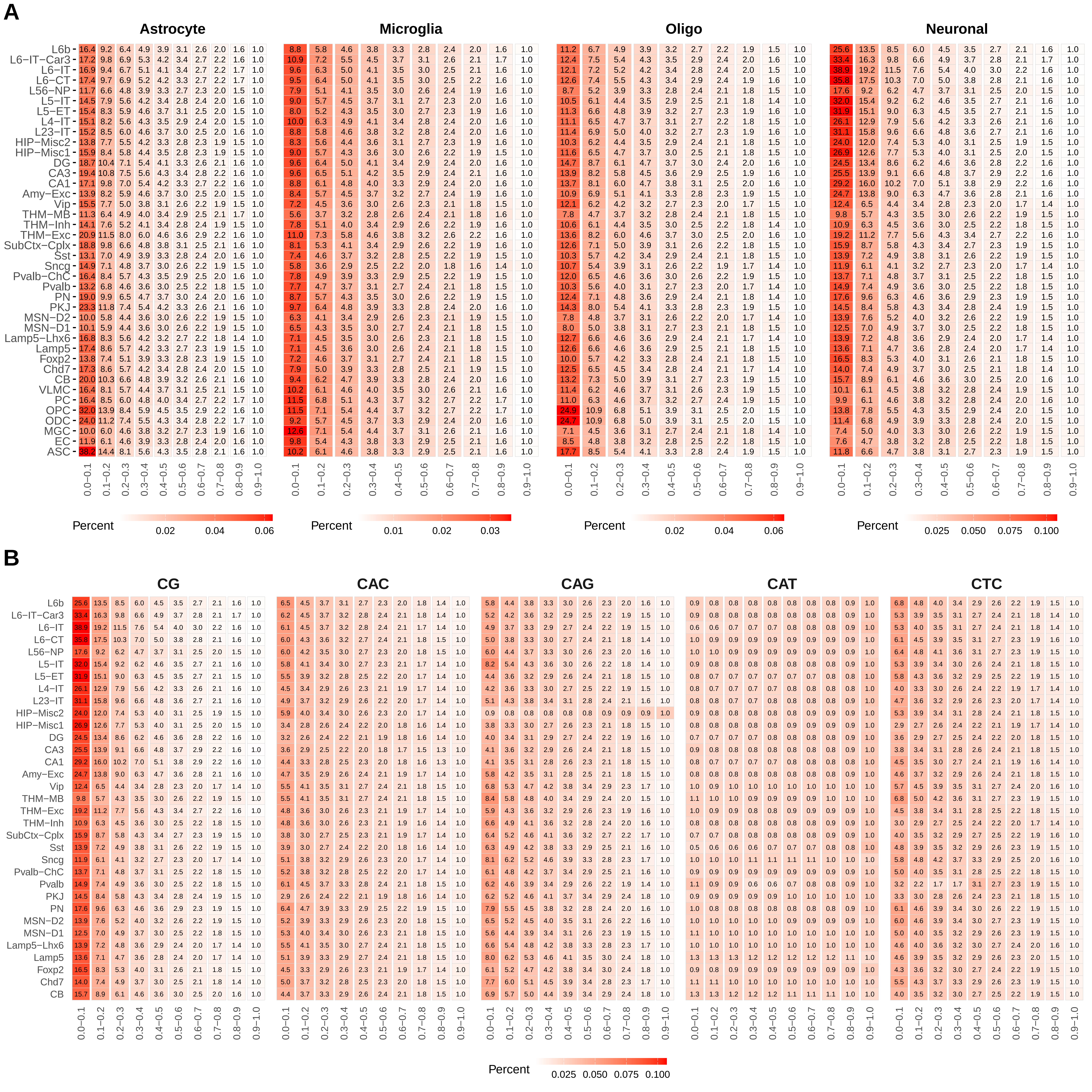

### Supplementary Figure 8

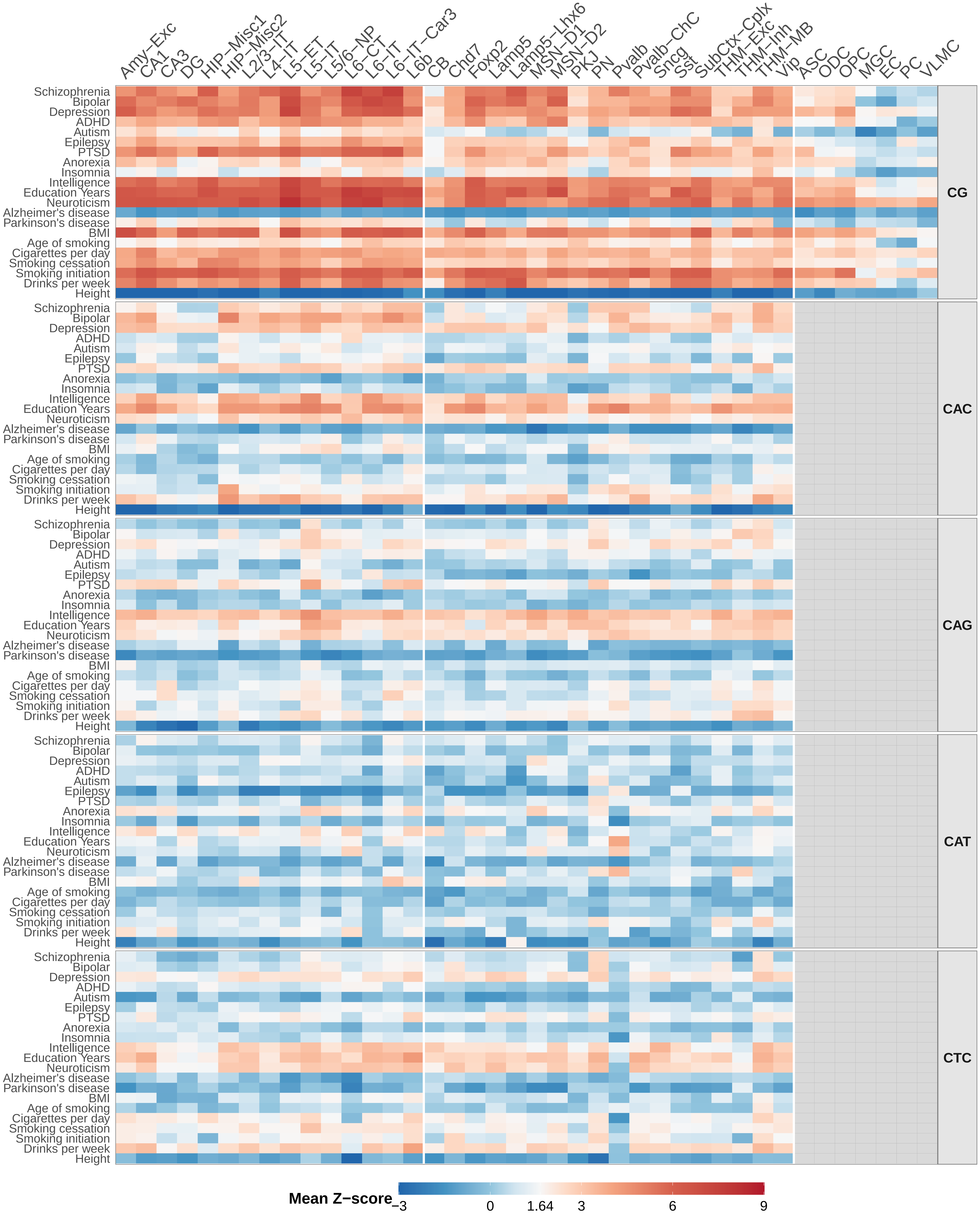
