## Supplementary Figure 9 for "Distinct cellular DNA methylation mechanisms underlie common and rare genetic risk for brain disorders"

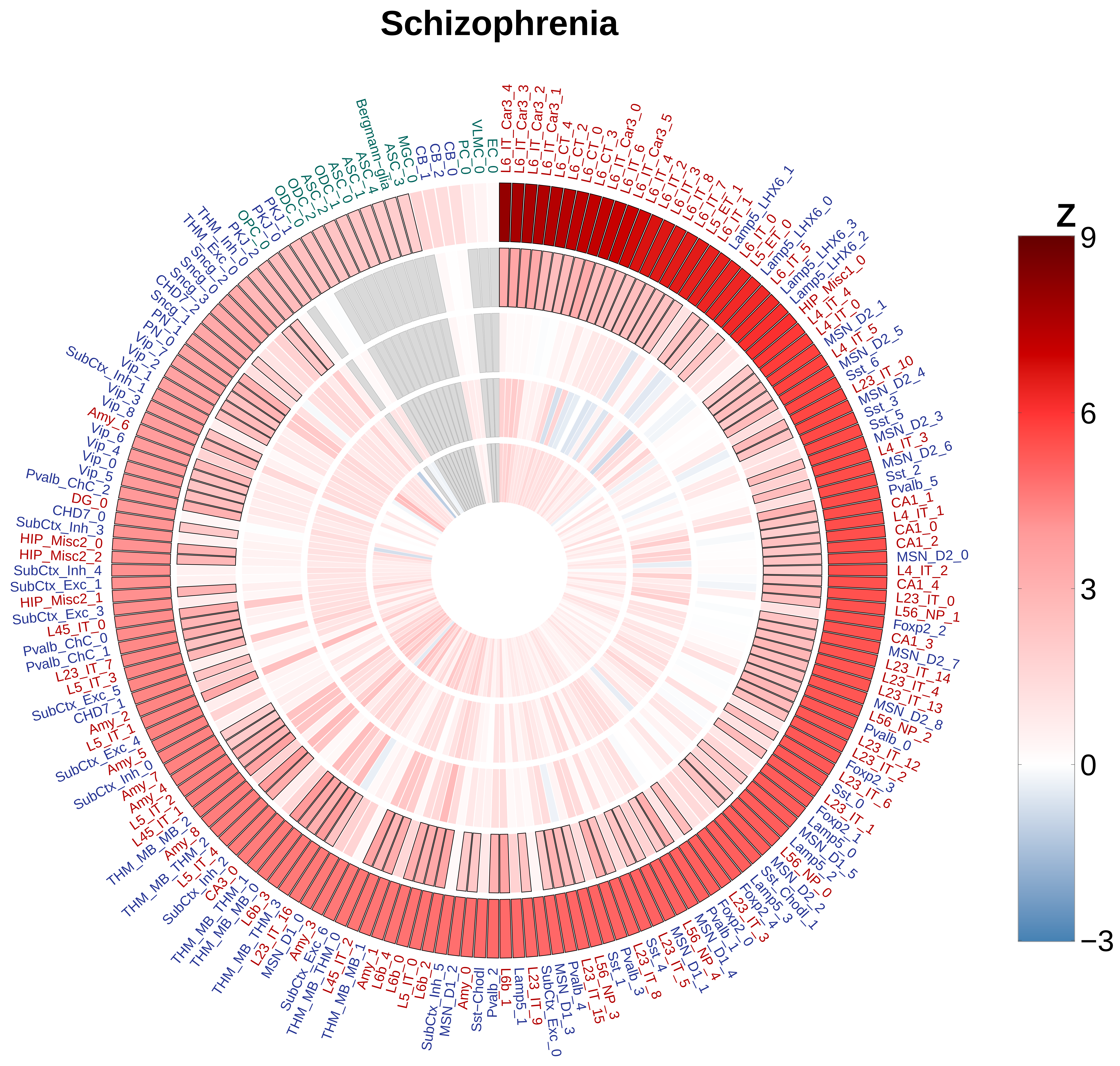
Schizophrenia
Z
9
6
3
0
−3
L6_IT_Car3_4
L6_IT_Car3_3
L6_IT_Car3_2
L6_IT_Car3_1
L6_CT_4
L6_CT_2
L6_CT_0
L6_CT_3
L6_IT_Car3_0
L6_IT_6
L6_IT_Car3_5
L6_IT_4
L6_IT_2
L6_IT_3
L6_IT_8
L6_IT_7
L5_ET_1
L6_IT_1
Lamp5_LHX6_1
L6_IT_0
L5_ET_0
Lamp5_LHX6_0
L6_IT_5
Lamp5_LHX6_3
Lamp5_LHX6_2
HIP_Misc1_0
L4_IT_4
L4_IT_0
MSN_D2_1
L4_IT_5
MSN_D2_5
Sst_6
L23_IT_10
MSN_D2_4
Sst_3
Sst_5
MSN_D2_3
L4_IT_3
MSN_D2_6
Sst_2
Pvalb_5
CA1_1
L4_IT_1
CA1_0
CA1_2
MSN_D2_0
L4_IT_2
CA1_4
L23_IT_0
L56_NP_1
Foxp2_2
CA1_3
MSN_D2_7
L23_IT_14
L23_IT_4
L23_IT_13
MSN_D2_8
L56_NP_2
Pvalb_0
L23_IT_12
L23_IT_2
Foxp2_3
L23_IT_6
Sst_0
L23_IT_1
Foxp2_1
Lamp5_0
MSN_D1_5
Lamp5_2
L56_NP_0
MSN_D2_2
Sst_Chodl_1
Lamp5_3
L23_IT_3
Foxp2_4
Foxp2_0
Pvalb_1
MSN_D1_4
L56_NP_4
MSN_D1_1
L23_IT_5
L23_IT_8
Sst_4
Pvalb_3
Sst_1
L56_NP_3
L23_IT_15
Pvalb_4
MSN_D1_3
SubCtx_Exc_0
L23_IT_9
Lamp5_1
L6b_1
Pvalb_2
Sst-Chodl
Amy_0
MSN_D1_2
SubCtx_Inh_5
L6b_2
L5_IT_0
L6b_0
L6b_4
Amy_1
THM_MB_MB_1
L45_IT_2
THM_MB_THM_0
SubCtx_Exc_6
Amy_3
MSN_D1_0
L23_IT_16
THM_MB_THM_3
L6b_3
THM_MB_MB_0
THM_MB_THM_1
CA3_0
SubCtx_Inh_2
L5_IT_4
THM_MB_THM_2
Amy_8
THM_MB_MB_2
L45_IT_1
L5_IT_2
Amy_4
Amy_7
SubCtx_Inh_0
Amy_5
SubCtx_Exc_4
L5_IT_1
Amy_2
CHD7_1
SubCtx_Exc_5
L5_IT_3
L23_IT_7
Pvalb_ChC_1
Pvalb_ChC_0
L45_IT_0
SubCtx_Exc_3
HIP_Misc2_1
SubCtx_Exc_1
SubCtx_Inh_4
HIP_Misc2_2
HIP_Misc2_0
SubCtx_Inh_3
CHD7_0
DG_0
Pvalb_ChC_2
Vip_5
Vip_0
Vip_4
Vip_6
Amy_6
Vip_8
Vip_3
Vip_1
SubCtx_Inh_1
Vip_2
Vip_7
PN_0
PN_1
Sncg_1
CHD7_2
Sncg_3
Sncg_0
Sncg_2
THM_Exc_0
THM_Inh_0
PKJ_2
OPC_0
PKJ_0
PKJ_1
ODC_0
ODC_2
ODC_1
ASC_2
ASC_0
ASC_1
ASC_4
Bergmann-glia
ASC_3
MGC_0
CB_1
CB_2
CB_0
PC_0
VLMC_0
EC_0

### Bipolar

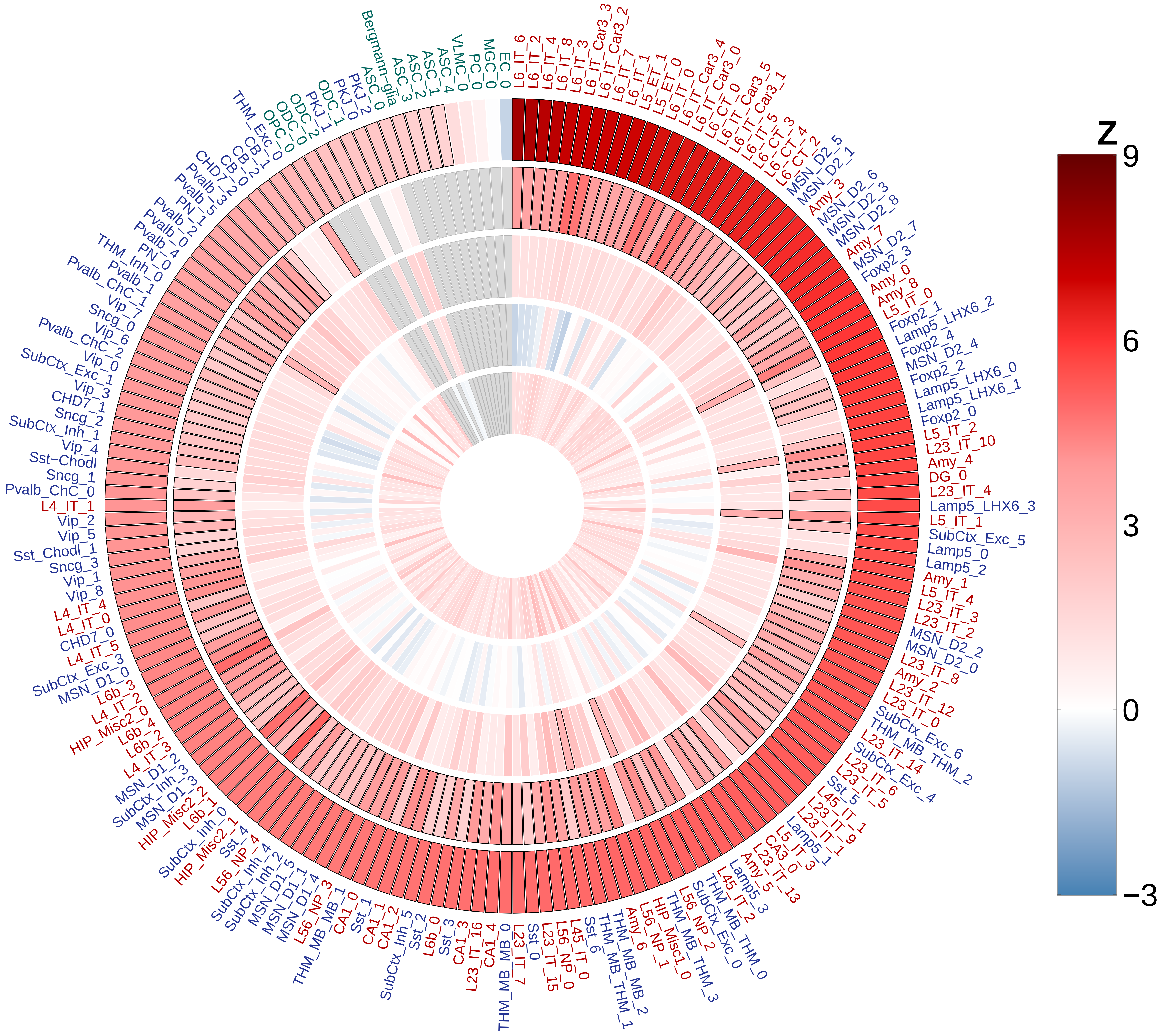

### Depression

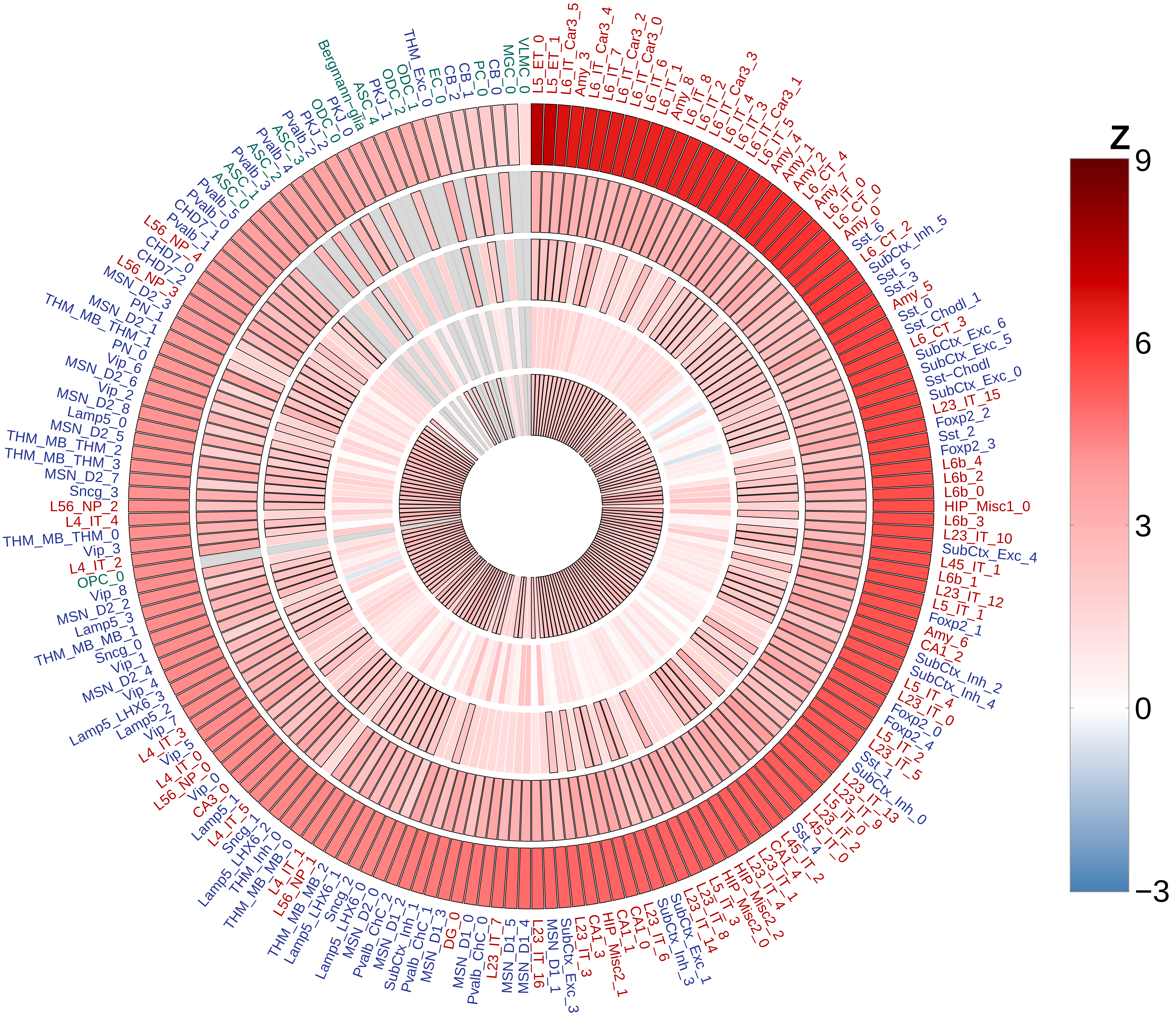

L5_ET_0
L5_ET_1
L6_IT_Car3_5
Amy_3
L6_IT_Car3_4
L6_IT_7
L6_IT_Car3_2
L6_IT_Car3_0
L6_IT_6
L6_IT_1
Amy_8
L6_IT_8
L6_IT_2
L6_IT_Car3_3
L6_IT_4
L6_IT_3
L6_IT_Car3_1
L6_IT_5
Amy_4
Amy_1
Amy_2
L6_CT_4
Amy_7
L6_IT_0
L6_CT_0
Amy_0
Sst_6
L6_CT_2
SubCtx_Inh_5
Sst_5
Sst_3
Amy_5
Sst_0
Sst_Chodl_1
L6_CT_3
SubCtx_Exc_6
SubCtx_Exc_5
Sst−Chodl
SubCtx_Exc_0
L23_IT_15
Foxp2_2
Sst_2
Foxp2_3
L6b_4
L6b_2
L6b_0
HIP_Misc1_0
L6b_3
L23_IT_10
SubCtx_Exc_4
L45_IT_1
L6b_1
L23_IT_12
L5_IT_1
Foxp2_1
Amy_6
CA1_2
SubCtx_Inh_2
SubCtx_Inh_4
L5_IT_4
L23_IT_0
Foxp2_0
Foxp2_4
L5_IT_2
L23_IT_5
Sst_1
SubCtx_Inh_0
L23_IT_13
L23_IT_9
L5_IT_0
L23_IT_2
L45_IT_0
Sst_4
L45_IT_2
CA1_4
L23_IT_1
L23_IT_4
HIP_Misc2_2
HIP_Misc2_0
L5_IT_3
L23_IT_8
L23_IT_14
SubCtx_Exc_1
SubCtx_Inh_3
L23_IT_6
CA1_0
CA1_1
HIP_Misc2_1
CA1_3
L23_IT_3
SubCtx_Exc_3
MSN_D1_1
L23_IT_16
MSN_D1_4
MSN_D1_5
L23_IT_7
Pvalb_ChC_0
MSN_D1_0
DG_0
MSN_D1_3
Pvalb_ChC_1
SubCtx_Inh_1
MSN_D1_2
Pvalb_ChC_2
MSN_D2_0
Lamp5_LHX6_0
Sncg_2
Lamp5_LHX6_1
THM_MB_MB_2
L56_NP_1
L4_IT_1
THM_MB_MB_0
THM_Inh_0
Lamp5_LHX6_2
Sncg_1
L4_IT_5
Lamp5_1
CA3_0
Vip_0
L56_NP_0
L4_IT_0
Vip_5
L4_IT_3
Vip_7
Lamp5_2
Lamp5_LHX6_3
Vip_4
MSN_D2_4
Vip_1
Sncg_0
THM_MB_MB_1
Lamp5_3
MSN_D2_2
Vip_8
OPC_0
L4_IT_2
Vip_3
THM_MB_THM_0
L4_IT_4
L56_NP_2
Sncg_3
MSN_D2_7
THM_MB_THM_3
THM_MB_THM_2
MSN_D2_5
Lamp5_0
MSN_D2_8
Vip_2
MSN_D2_6
Vip_6
PN_0
THM_MB_THM_1
MSN_D2_1
PN_1
MSN_D2_3
L56_NP_3
CHD7_2
CHD7_0
L56_NP_4
Pvalb_1
CHD7_1
Pvalb_0
Pvalb_5
ASC_0
ASC_1
Pvalb_3
ASC_2
Pvalb_4
ASC_3
Pvalb_2
PKJ_2
ODC_0
PKJ_0
Bergmann−glia
ASC_4
PKJ_1
ODC_2
ODC_1
THM_Exc_0
EC_0
CB_2
CB_1
PC_0
CB_0
MGC_0
VLMC_0
Z
9
6
3
0
−3

### ADHD

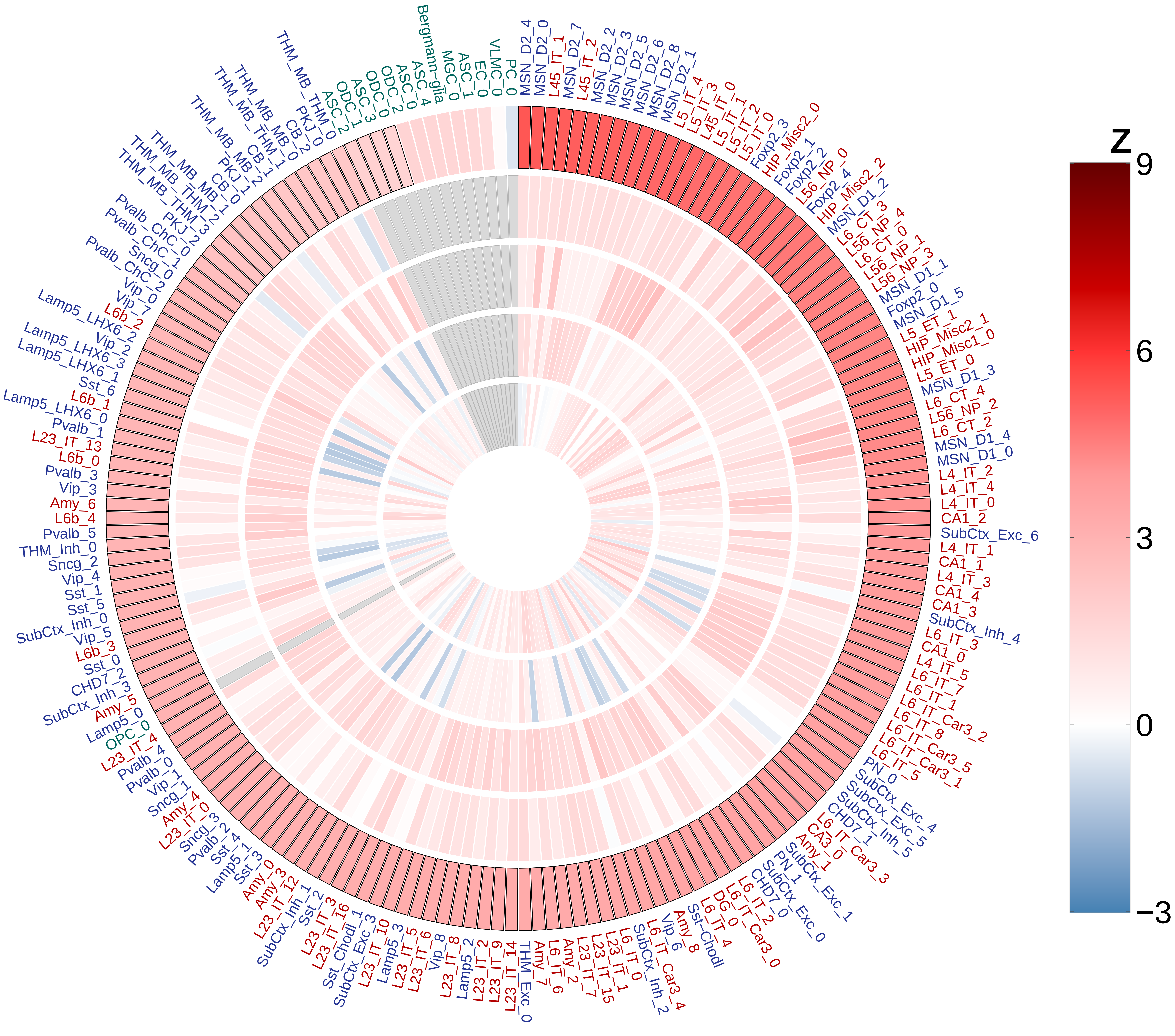

MSN_D2_4
MSN_D2_0
L45_IT_1
MSN_D2_7
L45_IT_2
MSN_D2_2
MSN_D2_3
MSN_D2_5
MSN_D2_6
MSN_D2_8
MSN_D2_1
L5_IT_4
L5_IT_3
L45_IT_0
L5_IT_1
L5_IT_2
L5_IT_0
Foxp2_3
HIP_Misc2_0
Foxp2_1
Foxp2_2
L56_NP_0
Foxp2_4
HIP_Misc2_2
MSN_D1_2
L6_CT_3
L56_NP_4
L6_CT_0
L56_NP_1
L56_NP_3
MSN_D1_1
Foxp2_0
MSN_D1_5
L5_ET_1
HIP_Misc2_1
HIP_Misc1_0
L5_ET_0
MSN_D1_3
L6_CT_4
L56_NP_2
L6_CT_2
MSN_D1_4
MSN_D1_0
L4_IT_2
L4_IT_4
L4_IT_0
CA1_2
SubCtx_Exc_6
L4_IT_1
CA1_1
L4_IT_3
CA1_4
CA1_3
SubCtx_Inh_4
L6_IT_3
CA1_0
L4_IT_5
L6_IT_7
L6_IT_1
L6_IT_Car3_2
L6_IT_8
L6_IT_Car3_5
L6_IT_Car3_1
L6_IT_5
PN_0
SubCtx_Exc_4
SubCtx_Exc_5
SubCtx_Inh_5
CHD7_1
L6_IT_Car3_3
CA3_0
Amy_1
SubCtx_Exc_1
PN_1
SubCtx_Exc_0
CHD7_0
L6_IT_2
L6_IT_Car3_0
DG_0
L6_IT_4
Sst-Chodl
Amy_8
Vip_6
L6_IT_Car3_4
SubCtx_Inh_2
L6_IT_0
L23_IT_1
L23_IT_15
L23_IT_7
Amy_2
L6_IT_6
Amy_7
THM_Exc_0
L23_IT_14
L23_IT_9
L23_IT_2
Lamp5_2
L23_IT_8
Vip_8
L23_IT_6
L23_IT_5
Lamp5_3
L23_IT_10
SubCtx_Exc_3
Sst_Chodl_1
L23_IT_16
L23_IT_3
Sst_2
SubCtx_Inh_1
L23_IT_12
Amy_3
Amy_0
Sst_3
Lamp5_1
Sst_4
Pvalb_2
Sncg_3
L23_IT_0
Amy_4
Sncg_1
Vip_1
Pvalb_0
Pvalb_4
L23_IT_4
OPC_0
Lamp5_0
Amy_5
SubCtx_Inh_3
CHD7_2
Sst_0
L6b_3
Vip_5
SubCtx_Inh_0
Sst_5
Sst_1
Vip_4
Sncg_2
THM_Inh_0
Pvalb_5
L6b_4
Amy_6
Vip_3
Pvalb_3
L6b_0
L23_IT_13
Pvalb_1
Lamp5_LHX6_0
L6b_1
Sst_6
Lamp5_LHX6_1
Lamp5_LHX6_3
Vip_2
Lamp5_LHX6_2
L6b_2
Vip_7
Vip_0
Pvalb_ChC_2
Sncg_0
Pvalb_ChC_1
Pvalb_ChC_0
PKJ_2
THM_MB_THM_3
THM_MB_THM_2
THM_MB_1
CB_0
PKJ_1
THM_MB_2
THM_MB_THM_1
CB_1
THM_MB_0
CB_2
PKJ_0
THM_MB_THM_0
ASC_2
ODC_1
ASC_3
ODC_0
ODC_2
ASC_0
ASC_4
Bergmann-glia
MGC_0
ASC_1
EC_0
VLMC_0
PC_0
Z
9
6
3
0
−3

### Autism

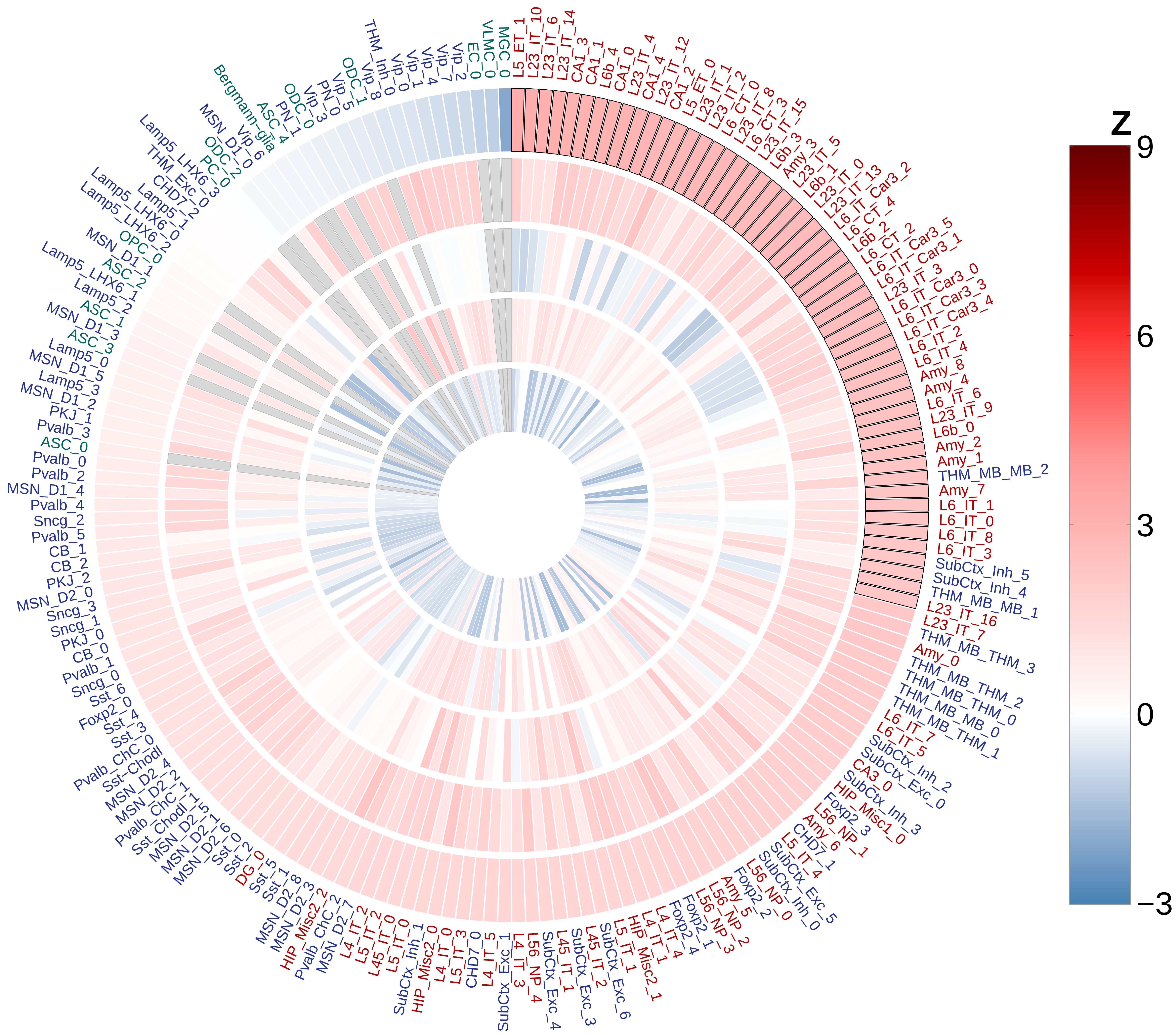

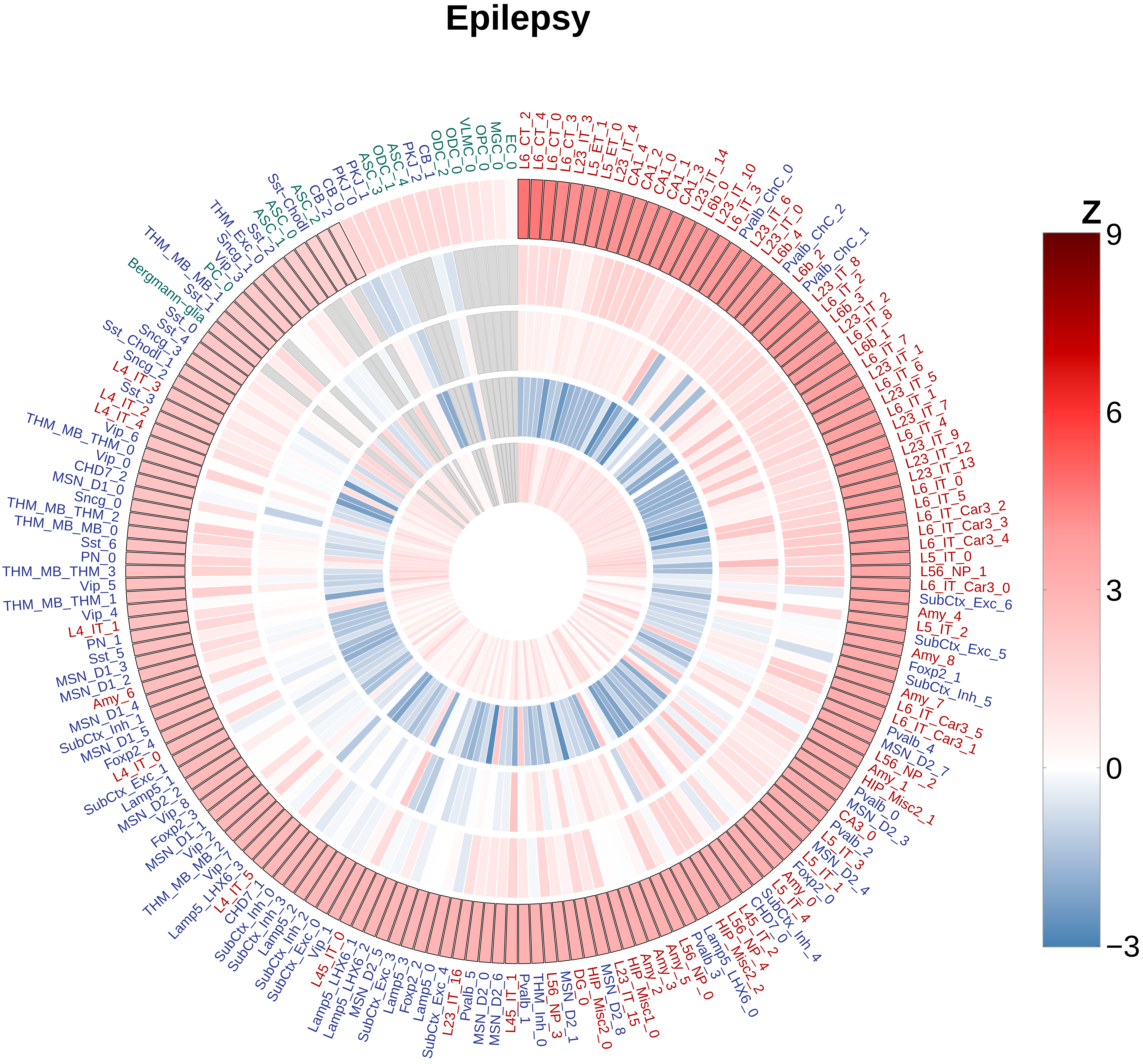

Epilepsy
L6_CT_2
L6_CT_4
L6_CT_0
L6_CT_3
L23_IT_3
L5_ET_1
L5_ET_0
L23_IT_4
CA1_4
CA1_2
CA1_0
CA1_1
CA1_3
L23_IT_14
L6b_0
L23_IT_10
L6_IT_3
Pvalb_ChC_0
L23_IT_6
L23_IT_0
L6b_4
Pvalb_ChC_2
L6b_2
Pvalb_ChC_1
L23_IT_8
L6_IT_2
L6b_3
L23_IT_2
L6_IT_8
L6b_1
L6_IT_7
L23_IT_1
L6_IT_6
L23_IT_5
L6_IT_1
L23_IT_7
L6_IT_4
L23_IT_9
L23_IT_12
L23_IT_13
L6_IT_0
L6_IT_5
L6_IT_Car3_2
L6_IT_Car3_3
L6_IT_Car3_4
L5_IT_0
L56_NP_1
L6_IT_Car3_0
SubCtx_Exc_6
Amy_4
L5_IT_2
SubCtx_Exc_5
Amy_8
Foxp2_1
SubCtx_Inh_5
Amy_7
L6_IT_Car3_5
L6_IT_Car3_1
Pvalb_4
MSN_D2_7
L56_NP_2
Amy_1
HIP_Misc2_1
Pvalb_0
MSN_D2_3
CA3_0
Pvalb_2
L5_IT_3
MSN_D2_4
L5_IT_1
Foxp2_0
Amy_0
L5_IT_4
SubCtx_Inh_4
CHD7_0
L45_IT_2
L56_NP_4
HIP_Misc2_2
Lamp5_LHX6_0
Pvalb_3
L56_NP_0
Amy_5
Amy_3
Amy_2
HIP_Misc1_0
L23_IT_15
MSN_D2_8
HIP_Misc2_0
DG_0
MSN_D2_1
L56_NP_3
THM_Inh_0
Pvalb_1
L45_IT_1
MSN_D2_6
MSN_D2_0
Pvalb_5
L23_IT_16
SubCtx_Exc_4
Lamp5_0
Foxp2_2
Lamp5_3
SubCtx_Exc_3
MSN_D2_5
Lamp5_LHX6_2
Lamp5_LHX6_1
L45_IT_0
Vip_1
SubCtx_Exc_0
SubCtx_Inh_2
Lamp5_2
SubCtx_Inh_3
SubCtx_Inh_0
CHD7_1
L4_IT_5
Lamp5_LHX6_3
THM_MB_MB_2
Vip_3
Vip_7
Vip_2
MSN_D1_1
Foxp2_3
Vip_8
MSN_D2_2
Lamp5_1
SubCtx_Exc_1
L4_IT_0
Foxp2_4
MSN_D1_5
SubCtx_Inh_1
MSN_D1_4
Amy_6
MSN_D1_2
MSN_D1_3
Sst_5
PN_1
L4_IT_1
Vip_4
THM_MB_THM_1
Vip_5
THM_MB_THM_3
PN_0
Sst_6
THM_MB_MB_0
THM_MB_THM_2
Sncg_0
MSN_D1_0
CHD7_2
Vip_0
THM_MB_THM_0
Vip_6
L4_IT_4
L4_IT_2
Sst_3
L4_IT_3
Sncg_2
Sncg_1
Sst_Chodl_3
Sncg_3
Sst_4
Sst_0
Sst_1
Bergmann−glia
THM_MB_MB_1
PC_0
THM_Exc_0
Sst_2
ASC_1
ASC_0
Sst−Chodl
ASC_2
CB_2
CB_0
PKJ_0
PKJ_1
ASC_3
ODC_1
ASC_4
PKJ_2
CB_1
ODC_2
ODC_0
VLMC_0
OPC_0
MGC_0
EC_0
Z
9
6
3
0
−3

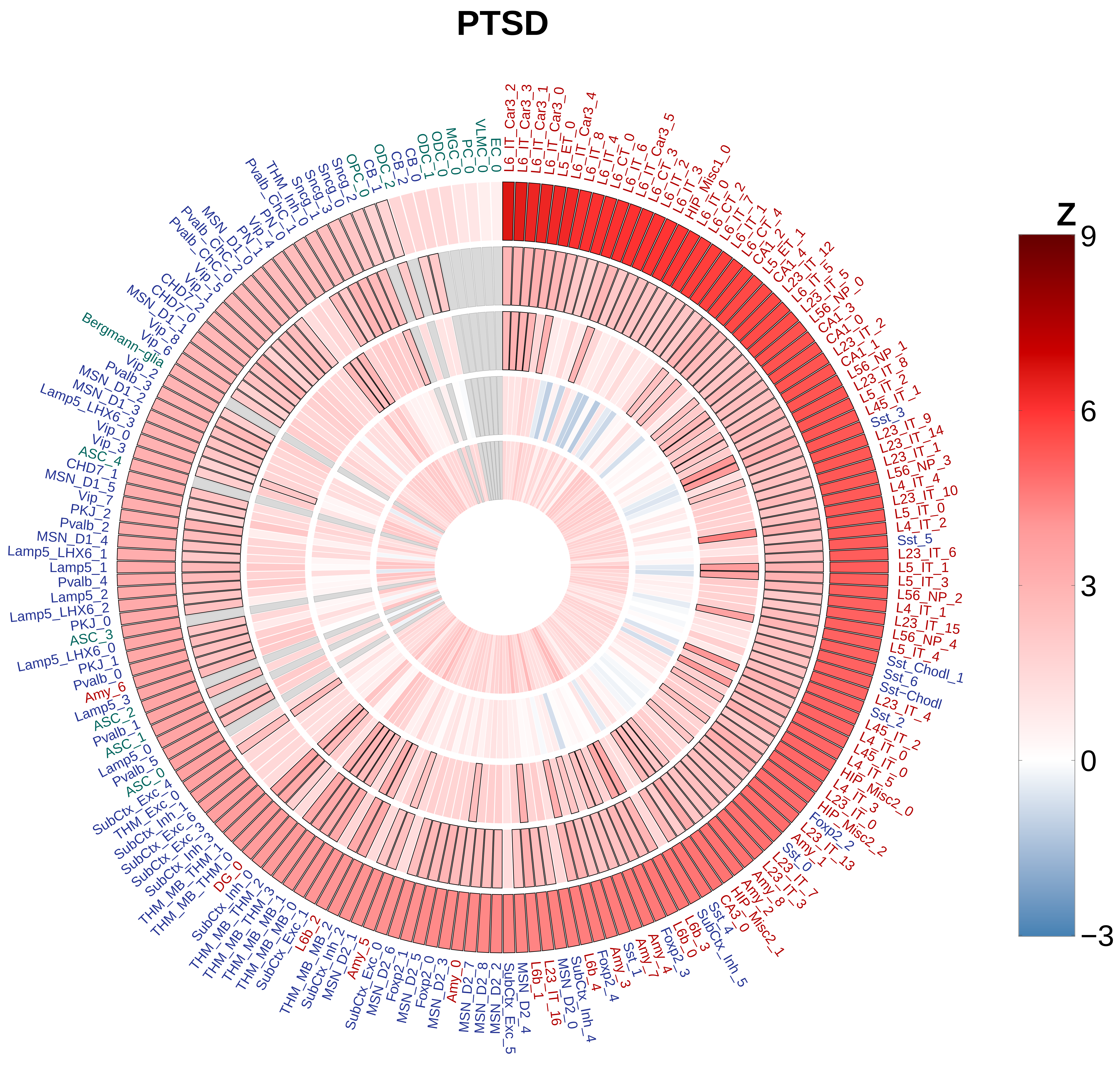
PTSD
Z
9
6
3
0
−3
L6_IT_Car3_2
L6_IT_Car3_3
L6_IT_Car3_1
L6_IT_Car3_0
L5_ET_0
L6_IT_Car3_4
L6_IT_8
L6_IT_4
L6_CT_0
L6_IT_6
L6_IT_Car3_5
L6_CT_3
L6_IT_2
L6_IT_3
HIP_Misc1_0
L6_IT_0
L6_CT_2
L6_IT_7
L6_IT_1
L6_CT_4
CA1_2
L5_ET_1
CA1_4
L23_IT_12
L6_IT_5
L23_IT_5
L56_NP_0
CA1_3
CA1_0
L23_IT_2
CA1_1
L56_NP_1
L23_IT_8
L5_IT_2
L45_IT_1
Sst_3
L23_IT_9
L23_IT_14
L23_IT_1
L56_NP_3
L4_IT_4
L23_IT_10
L5_IT_0
L4_IT_2
Sst_5
L23_IT_6
L5_IT_1
L5_IT_3
L56_NP_2
L4_IT_1
L23_IT_15
L56_NP_4
L5_IT_4
Sst_Chodl_1
Sst_6
Sst-Chodl
L23_IT_4
Sst_2
L45_IT_2
L4_IT_0
L45_IT_0
L4_IT_5
HIP_Misc2_0
L4_IT_3
L23_IT_0
HIP_Misc2_2
Foxp2_2
L23_IT_13
Amy_1
Sst_0
L23_IT_7
L23_IT_3
Amy_8
Amy_2
HIP_Misc2_1
CA3_0
Sst_4
SubCtx_Inh_5
L6b_3
L6b_0
Foxp2_3
Amy_4
Amy_7
Sst_1
Amy_3
Foxp2_4
L6b_4
SubCtx_Inh_4
MSN_D2_0
L23_IT_16
L6b_1
MSN_D2_4
SubCtx_Exc_5
MSN_D2_2
MSN_D2_8
MSN_D2_7
Amy_0
MSN_D2_3
Foxp2_0
MSN_D2_5
Foxp2_1
MSN_D2_6
SubCtx_Exc_0
Amy_5
MSN_D2_1
SubCtx_Inh_2
THM_MB_MB_2
L6b_2
SubCtx_Exc_1
THM_MB_MB_0
THM_MB_MB_1
THM_MB_THM_3
THM_MB_THM_2
SubCtx_Inh_0
DG_0
THM_MB_THM_0
THM_MB_THM_1
SubCtx_Inh_3
SubCtx_Exc_3
SubCtx_Exc_6
SubCtx_Inh_1
THM_Exc_0
SubCtx_Exc_4
ASC_0
Pvalb_5
Lamp5_0
ASC_1
Pvalb_1
ASC_2
Lamp5_3
Amy_6
Pvalb_0
PKJ_1
Lamp5_LHX6_0
ASC_3
PKJ_0
Lamp5_LHX6_2
Lamp5_2
Pvalb_4
Lamp5_1
Lamp5_LHX6_1
MSN_D1_4
Pvalb_2
PKJ_2
Vip_7
MSN_D1_5
CHD7_1
ASC_4
Vip_3
Vip_0
Lamp5_LHX6_3
MSN_D1_3
MSN_D1_2
Pvalb_3
Vip_2
Bergmann-glia
Vip_6
Vip_8
MSN_D1_1
CHD7_0
CHD7_2
Vip_1
Vip_5
Pvalb_ChC_0
Pvalb_ChC_2
MSN_D1_0
PN_1
Vip_4
PN_0
Pvalb_ChC_1
THM_Inh_0
Sncg_1
Sncg_3
Sncg_0
Sncg_2
OPC_0
CB_1
ODC_2
CB_2
CB_0
ODC_1
ODC_0
MGC_0
PC_0
VLMC_0
EC_0

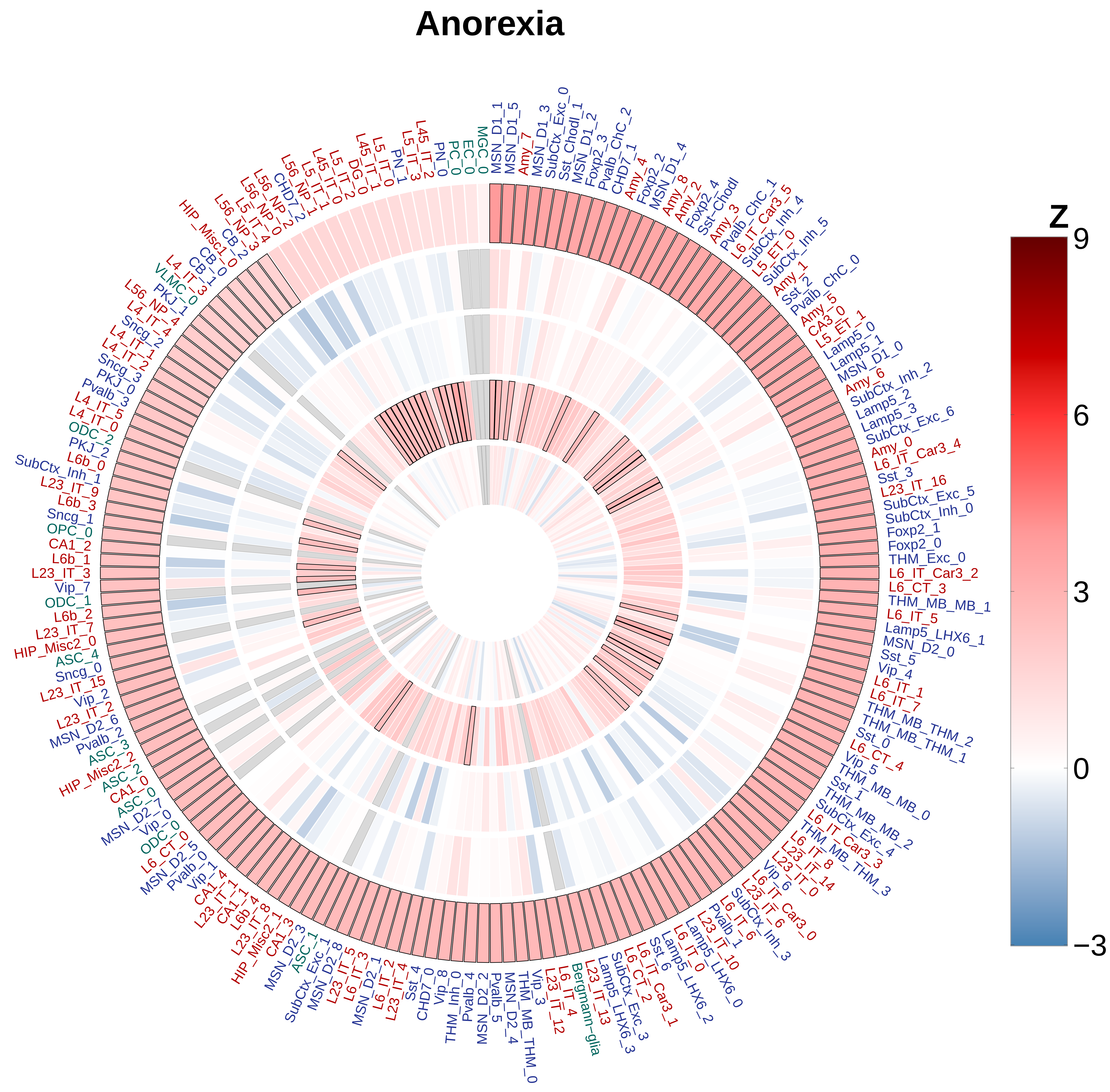
Anorexia
MSN_D1_1
MSN_D1_5
Amy_7
MSN_D1_3
SubCtx_Exc_0
Sst_Chodl_1
MSN_D1_2
Foxp2_3
Pvalb_ChC_2
CHD7_1
Amy_4
Foxp2_2
MSN_D1_4
Amy_8
Amy_2
Foxp2_4
Sst-Chodl
Amy_3
Pvalb_ChC_1
L6_IT_Car3_5
SubCtx_Inh_4
L5_ET_0
SubCtx_Inh_5
Amy_1
Sst_2
Pvalb_ChC_0
Amy_5
CA3_0
L5_ET_1
Lamp5_0
Lamp5_1
MSN_D1_0
Amy_6
SubCtx_Inh_2
Lamp5_2
Lamp5_3
SubCtx_Exc_6
Amy_0
L6_IT_Car3_4
Sst_3
L23_IT_16
SubCtx_Exc_5
SubCtx_Inh_0
Foxp2_1
Foxp2_0
THM_Exc_0
L6_IT_Car3_2
L6_CT_3
THM_MB_MB_1
L6_IT_5
Lamp5_LHX6_1
MSN_D2_0
Sst_5
Vip_4
L6_IT_1
L6_IT_7
THM_MB_THM_2
THM_MB_THM_1
Sst_0
L6_CT_4
Vip_5
THM_MB_MB_0
Sst_1
THM_MB_MB_2
SubCtx_Exc_4
L6_IT_Car3_3
THM_MB_THM_3
L6_IT_8
L23_IT_14
L23_IT_0
Vip_6
L6_IT_Car3_0
L23_IT_6
SubCtx_Inh_3
L6_IT_6
Pvalb_1
L23_IT_10
Lamp5_LHX6_0
L6_IT_0
Lamp5_LHX6_2
Sst_6
L6_IT_Car3_1
L6_CT_2
SubCtx_Exc_3
Lamp5_LHX6_3
L23_IT_13
Bergmann-glia
L6_IT_4
L23_IT_12
Vip_3
THM_MB_THM_0
MSN_D2_4
Pvalb_5
MSN_D2_2
Pvalb_4
THM_Inh_0
Vip_8
CHD7_0
Sst_4
L23_IT_4
L6_IT_2
MSN_D2_1
L6_IT_3
L23_IT_5
MSN_D2_8
SubCtx_Exc_1
ASC_1
MSN_D2_3
CA1_3
HIP_Misc2_1
L23_IT_8
L6b_4
CA1_1
L23_IT_1
CA1_4
Vip_1
Pvalb_0
MSN_D2_5
L6_CT_0
ODC_0
Vip_0
MSN_D2_7
ASC_0
CA1_0
ASC_2
HIP_Misc2_2
ASC_3
Pvalb_2
MSN_D2_6
L23_IT_2
Vip_2
L23_IT_15
Sncg_0
ASC_4
HIP_Misc2_0
L23_IT_7
L6b_2
ODC_1
Vip_7
L23_IT_3
L6b_1
CA1_2
OPC_0
Sncg_1
L6b_3
L23_IT_9
SubCtx_Inh_1
L6b_0
PKJ_2
ODC_2
L4_IT_0
L4_IT_5
Pvalb_3
PKJ_0
Sncg_3
L4_IT_2
L4_IT_1
Sncg_2
L4_IT_4
L56_NP_4
PKJ_1
VLMC_0
L4_IT_3
CB_0
CB_1
HIP_Misc1_0
CB_2
L56_NP_3
L5_IT_4
L56_NP_0
L5_NP_2
CHD7_2
L56_NP_1
L5_IT_1
L45_IT_0
L5_IT_2
DG_0
L45_IT_1
L5_IT_0
PN_1
L5_IT_3
L45_IT_2
PN_0
PC_0
EC_0
MGC_0
Z
9
6
3
0
−3

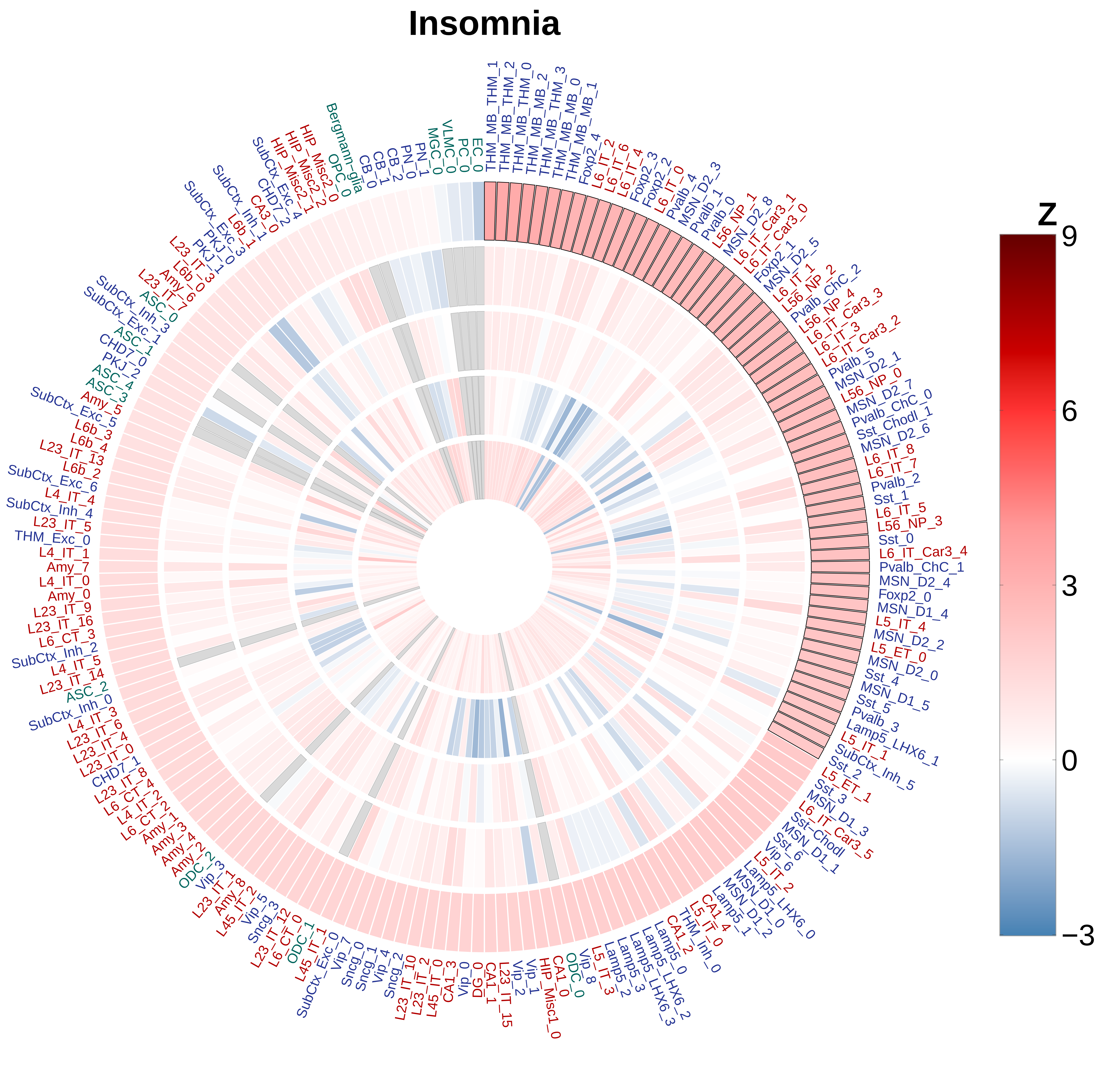

Insomnia
Z
9
6
3
0
−3
THM_MB_THM_1
THM_MB_THM_2
THM_MB_THM_0
THM_MB_MB_2
THM_MB_THM_3
THM_MB_MB_0
THM_MB_MB_1
Foxp2_4
L6_IT_2
L6_IT_6
L6_IT_4
Foxp2_3
Foxp2_2
L6_IT_0
Pvalb_4
MSN_D2_3
Pvalb_1
Pvalb_0
L56_NP_1
MSN_D2_8
L6_IT_Car3_1
L6_IT_Car3_0
Foxp2_1
MSN_D2_5
L6_IT_1
L56_NP_2
Pvalb_ChC_2
L56_NP_4
L6_IT_Car3_3
L6_IT_3
L6_IT_Car3_2
Pvalb_5
MSN_D2_1
L56_NP_0
MSN_D2_7
Pvalb_ChC_0
Sst_Chodl_1
MSN_D2_6
L6_IT_8
L6_IT_7
Pvalb_2
Sst_1
L6_IT_5
L56_NP_3
Sst_0
L6_IT_Car3_4
Pvalb_ChC_1
MSN_D2_4
Foxp2_0
MSN_D1_4
L5_IT_4
MSN_D2_2
L5_ET_0
MSN_D2_0
Sst_4
MSN_D1_5
Sst_5
Pvalb_3
Lamp5_LHX6_1
L5_IT_1
SubCtx_Inh_5
Sst_2
L5_ET_1
Sst_3
MSN_D1_3
L6_IT_Car3_5
Sst_Chodl
MSN_D1_1
Sst_6
Vip_6
L5_IT_2
Lamp5_LHX6_0
MSN_D1_0
MSN_D1_2
Lamp5_1
CA1_4
L5_IT_0
THM_Inh_0
CA1_2
Lamp5_0
Lamp5_LHX6_2
Lamp5_LHX6_3
Lamp5_3
Lamp5_2
L5_IT_3
Vip_8
ODC_0
CA1_0
HIP_Misc1_0
Vip_1
Vip_2
L23_IT_15
CA1_1
DG_0
Vip_0
CA1_3
L45_IT_0
L23_IT_2
L23_IT_10
Sncg_2
Vip_4
Sncg_1
Sncg_0
Vip_7
SubCtx_Exc_0
L45_IT_1
ODC_1
L6_CT_0
L23_IT_12
Sncg_3
Vip_5
L45_IT_2
Amy_8
L23_IT_1
Vip_3
ODC_2
Amy_2
Amy_4
Amy_3
Amy_1
L6_CT_2
L4_IT_2
L6_CT_4
L23_IT_8
CHD7_1
L23_IT_0
L23_IT_4
L23_IT_6
L4_IT_3
SubCtx_Inh_0
ASC_2
L23_IT_14
L4_IT_5
SubCtx_Inh_2
L6_CT_3
L23_IT_16
L23_IT_9
Amy_0
L4_IT_0
Amy_7
L4_IT_1
THM_Exc_0
L23_IT_5
SubCtx_Inh_4
L4_IT_4
SubCtx_Exc_6
L6b_2
L23_IT_13
L6b_4
L6b_3
SubCtx_Exc_5
Amy_5
ASC_3
ASC_4
PKJ_2
CHD7_0
ASC_1
SubCtx_Exc_1
SubCtx_Inh_3
ASC_0
L23_IT_7
Amy_6
L6b_0
L23_IT_3
PKJ_1
PKJ_0
SubCtx_Exc_3
L6b_1
SubCtx_Inh_1
CA3_0
CHD7_2
SubCtx_Exc_4
HIP_Misc2_1
HIP_Misc2_2
HIP_Misc2_0
OPC_0
Bergmann-glia
CB_0
CB_1
CB_2
PN_0
PN_1
MGC_0
VLMC_0
PC_0
EC_0

### Intelligence

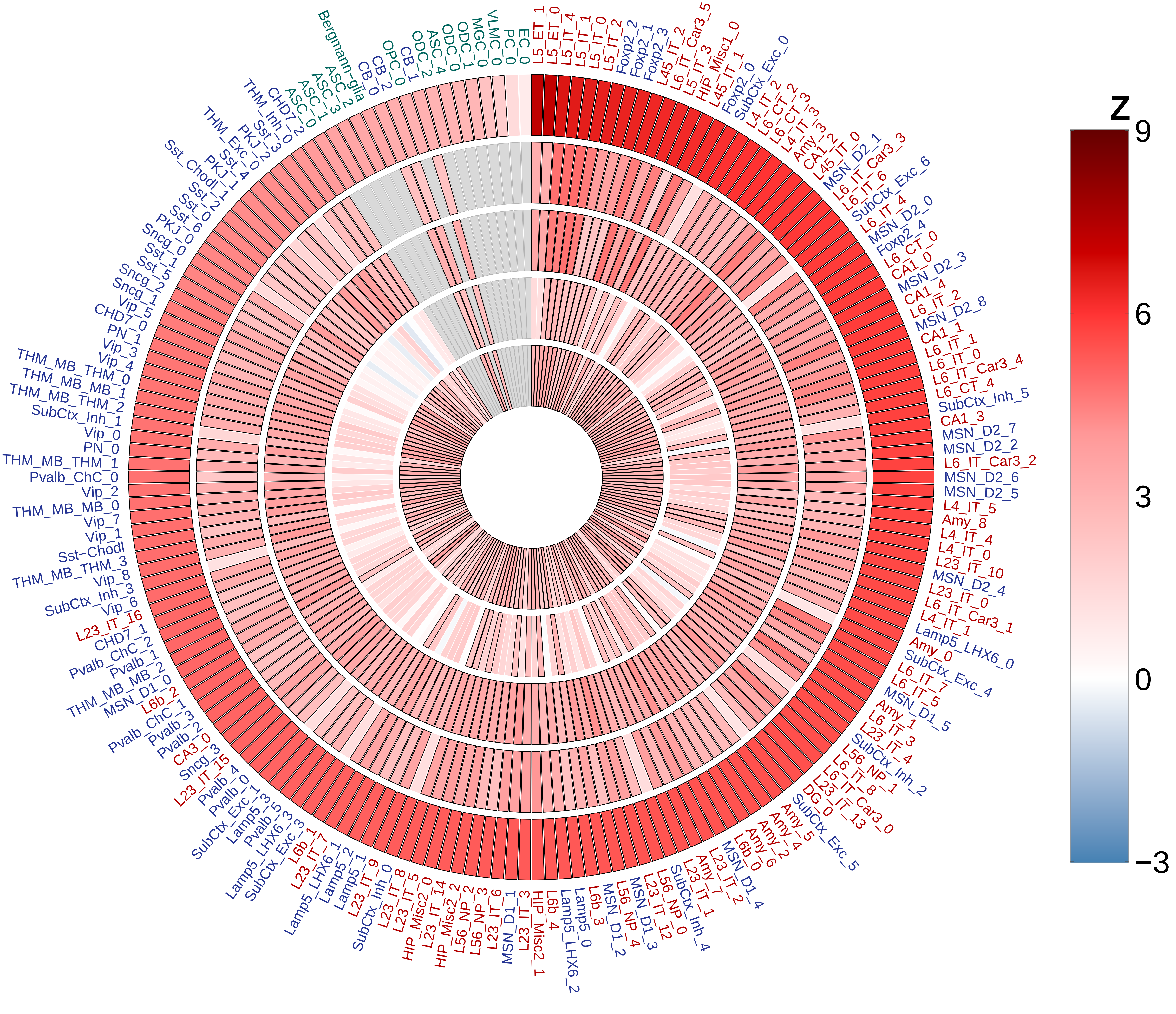

### Education Years

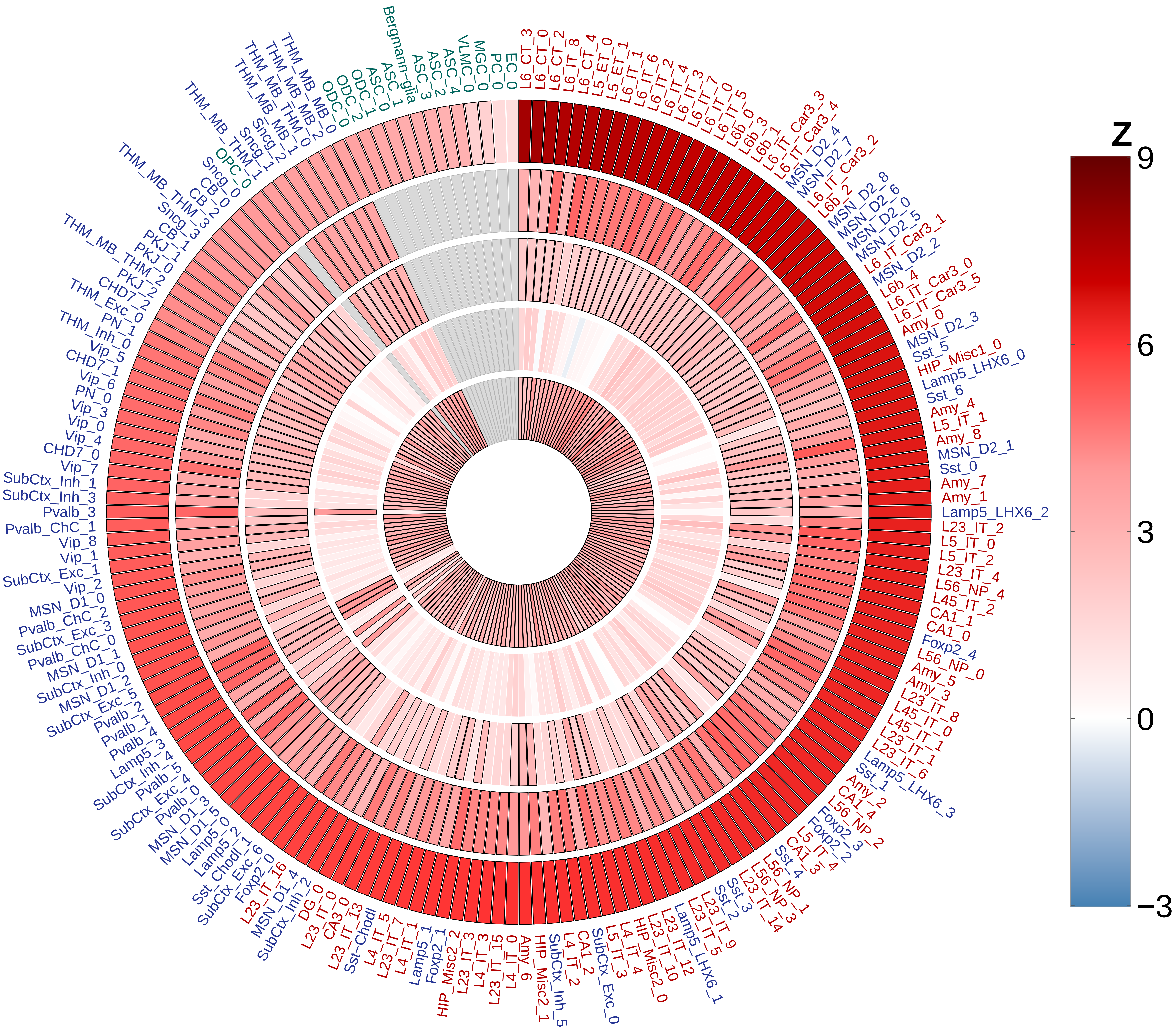

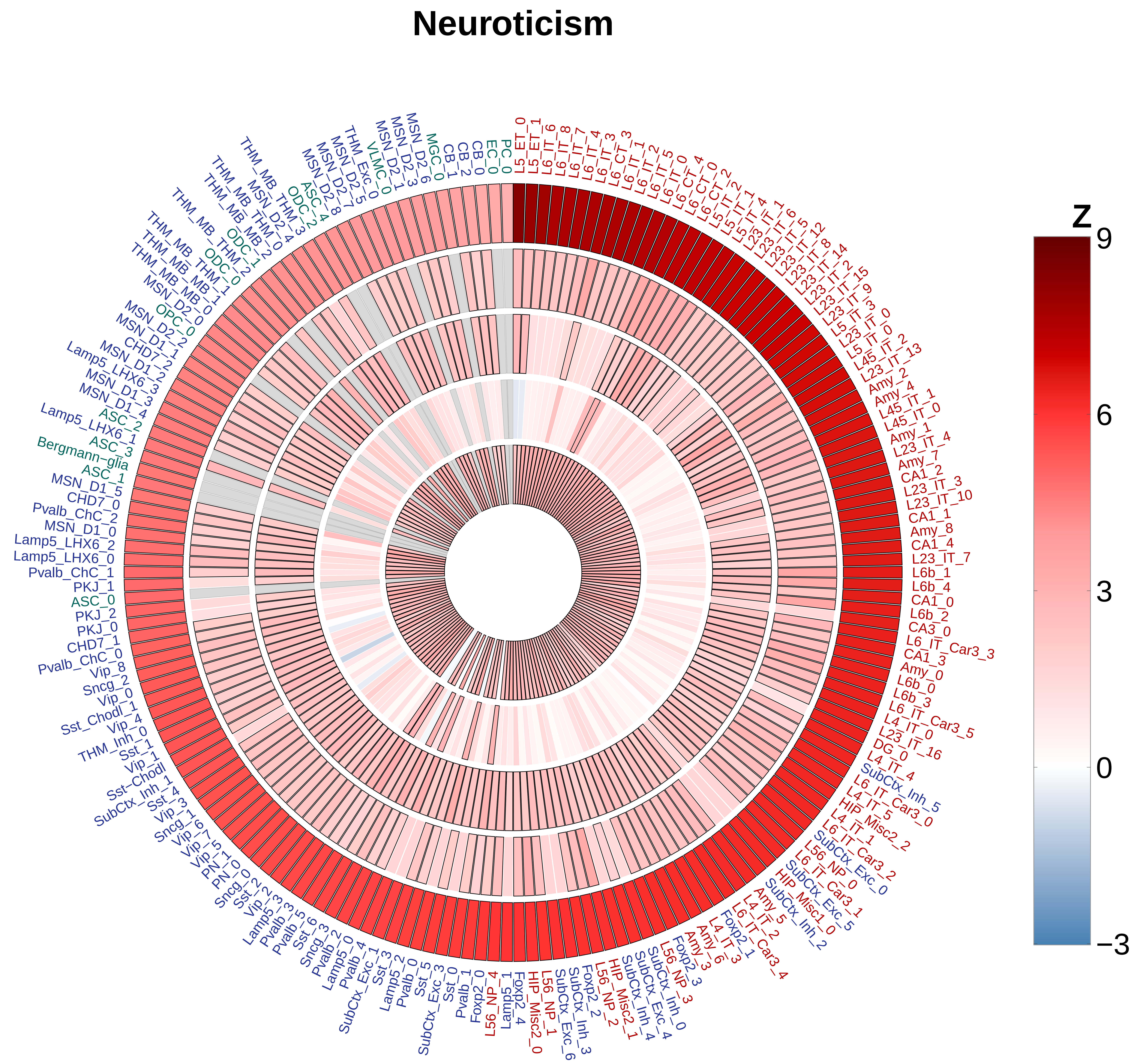
Neuroticism
Z
9
6
3
0
−3
L5_ET_0
L5_ET_1
L6_IT_6
L6_IT_8
L6_IT_7
L6_IT_4
L6_IT_3
L6_CT_3
L6_CT_1
L6_IT_2
L6_IT_5
L6_IT_0
L6_CT_4
L6_CT_0
L6_CT_2
L5_IT_2
L5_IT_1
L5_IT_4
L23_IT_1
L23_IT_6
L23_IT_5
L23_IT_12
L23_IT_8
L23_IT_14
L23_IT_2
L23_IT_15
L23_IT_9
L23_IT_3
L5_IT_3
L23_IT_0
L5_IT_0
L45_IT_2
L23_IT_13
Amy_2
Amy_4
L45_IT_1
L45_IT_0
Amy_1
L23_IT_4
Amy_7
CA1_2
L23_IT_3
L23_IT_10
CA1_1
Amy_8
CA1_4
L23_IT_7
L6b_1
L6b_4
CA1_0
L6b_2
CA3_0
L6_IT_Car3_3
CA1_3
Amy_0
L6b_0
L6b_3
L6_IT_Car3_5
L4_IT_0
L23_IT_16
DG_0
L4_IT_4
SubCtx_Inh_5
L6_IT_Car3_0
L4_IT_5
HIP_Misc2_2
L4_IT_1
L6_IT_Car3_2
SubCtx_Exc_0
L56_NP_0
SubCtx_Exc_5
L6_IT_Car3_1
HIP_Misc1_0
SubCtx_Inh_2
L6_IT_Car3_4
L4_IT_2
Amy_5
Foxp2_1
L4_IT_3
Amy_6
Amy_3
Foxp2_3
L56_NP_3
SubCtx_Inh_0
SubCtx_Exc_4
SubCtx_Inh_4
HIP_Misc2_1
L56_NP_2
Foxp2_2
SubCtx_Inh_3
SubCtx_Exc_6
L56_NP_1
HIP_Misc2_0
Foxp2_4
Lamp5_1
L56_NP_4
Foxp2_0
Pvalb_1
Sst_0
SubCtx_Exc_3
Sst_5
Pvalb_0
Lamp5_2
Sst_3
SubCtx_Exc_1
Pvalb_4
Lamp5_0
Pvalb_2
Sncg_3
Sst_6
Pvalb_5
Pvalb_3
Lamp5_3
Vip_2
Sst_2
Sncg_0
PN_0
PN_1
Vip_5
Vip_7
Vip_6
Sncg_1
Vip_3
Sst_4
SubCtx_Inh_1
Sst-Chodl
Vip_1
Sst_1
THM_Inh_0
Vip_4
Sst_Chodl_1
Vip_0
Sncg_2
Vip_8
Pvalb_ChC_0
CHD7_1
PKJ_0
PKJ_2
ASC_0
PKJ_1
Pvalb_ChC_1
Lamp5_LHX6_0
Lamp5_LHX6_2
MSN_D1_0
Pvalb_ChC_2
CHD7_0
MSN_D1_5
ASC_1
Bergmann-glia
ASC_3
Lamp5_LHX6_1
ASC_2
MSN_D1_4
MSN_D1_3
Lamp5_LHX6_3
MSN_D1_2
CHD7_2
MSN_D1_1
MSN_D2_2
OPC_0
MSN_D2_0
THM_MB_MB_0
THM_MB_MB_1
THM_MB_THM_1
ODC_0
THM_MB_THM_2
ODC_1
THM_MB_MB_2
THM_MB_THM_0
MSN_D2_4
THM_3
ODC_2
ASC_4
MSN_D2_8
MSN_D2_7
MSN_D2_5
THM_Exc_0
VLMC_0
MSN_D2_1
MSN_D2_3
MSN_D2_6
MGC_0
CB_1
CB_2
CB_0
EC_0
PC_0

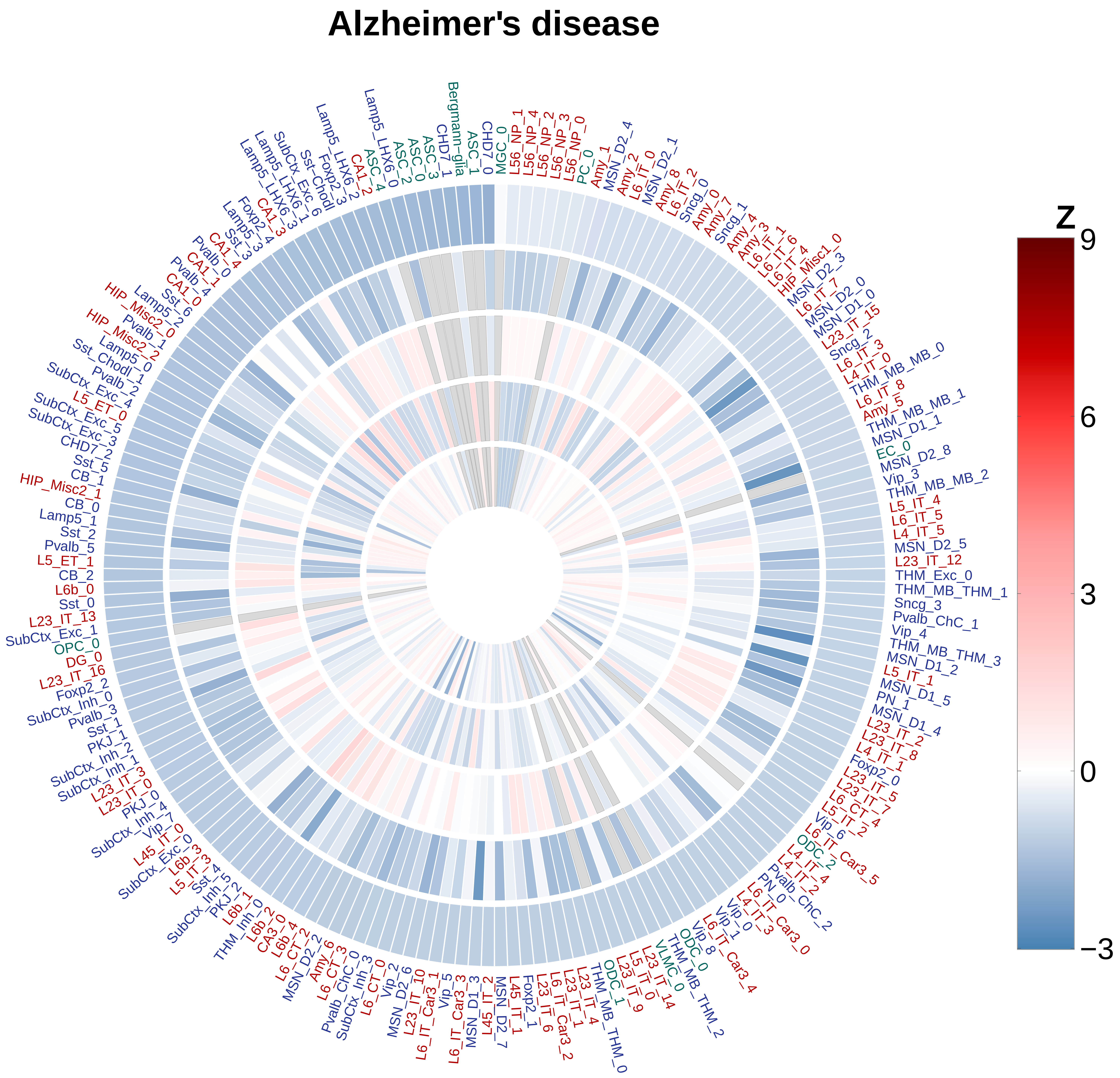

Alzheimer's disease
z
9
6
3
0
−3
CHD7_0
MGC_0
L56_NP_1
L56_NP_4
L56_NP_2
L56_NP_3
L56_NP_0
PC_0
Amy_1
MSN_D2_4
Amy_2
L6_IT_0
MSN_D2_1
Amy_8
L6_IT_2
Sncg_0
Amy_0
Amy_7
Sncg_1
Amy_4
Amy_3
L6_IT_1
L6_IT_6
L6_IT_4
HIP_Misc1_0
MSN_D2_3
L6_IT_7
MSN_D2_0
MSN_D1_0
L23_IT_15
Sncg_2
L6_IT_3
L4_IT_0
THM_MB_MB_0
L6_IT_8
Amy_5
THM_MB_MB_1
MSN_D1_1
EC_0
MSN_D2_8
Vip_3
THM_MB_MB_2
L5_IT_4
L6_IT_5
L4_IT_5
MSN_D2_5
L23_IT_12
THM_Exc_0
THM_MB_THM_1
Sncg_3
Pvalb_ChC_1
Vip_4
THM_MB_THM_3
MSN_D1_2
L5_IT_1
MSN_D1_5
PN_1
MSN_D1_4
L23_IT_2
L23_IT_8
L4_IT_1
Foxp2_0
L23_IT_5
Vip_6
ODC_2
L6_IT_Car3_5
L4_IT_4
L4_IT_2
Pvalb_ChC_2
PN_0
L6_IT_Car3_0
L4_IT_3
Vip_0
Vip_1
L6_IT_Car3_4
Vip_8
ODC_0
THM_MB_THM_2
VLMC_0
L23_IT_14
L5_IT_0
L23_IT_9
ODC_1
THM_MB_THM_0
L23_IT_4
L23_IT_1
L6_IT_Car3_2
L23_IT_6
Foxp2_1
L45_IT_1
MSN_D2_7
L45_IT_2
MSN_D1_3
L6_IT_Car3_3
Vip_5
L6_IT_Car3_1
L23_IT_10
MSN_D2_6
Vip_2
L6_CT_0
SubCtx_Inh_3
Pvalb_ChC_0
L6_CT_3
Amy_6
MSN_D2_2
L6_CT_2
L6b_4
CA3_0
L6b_2
THM_Inh_0
L6b_1
PKJ_2
SubCtx_Inh_5
Sst_4
L5_IT_3
L6b_3
SubCtx_Exc_0
L45_IT_0
Vip_7
SubCtx_Inh_4
PKJ_0
L23_IT_0
L23_IT_3
SubCtx_Inh_1
SubCtx_Inh_2
PKJ_1
Sst_1
Pvalb_3
SubCtx_Inh_0
Foxp2_2
L23_IT_16
DG_0
OPC_0
SubCtx_Exc_1
L23_IT_13
Sst_0
L6b_0
CB_2
L5_ET_1
Pvalb_5
Sst_2
Lamp5_1
CB_0
HIP_Misc2_1
CB_1
Sst_5
CHD7_2
SubCtx_Exc_3
SubCtx_Exc_5
L5_ET_0
SubCtx_Exc_4
Pvalb_2
Sst_Chodl_0
Sst_6
Pvalb_4
CA1_0
CA1_1
Pvalb_0
CA1_4
Sst_3
Lamp5_3
Foxp2_4
CA1_3
Lamp5_LHX6_3
Lamp5_LHX6_1
SubCtx_Exc_6
Sst_Chodl
Foxp2_3
Lamp5_LHX6_2
CA1_2
ASC_4
Lamp5_LHX6_0
ASC_2
ASC_0
ASC_3
CHD7_1
Bergmann-glia
ASC_1

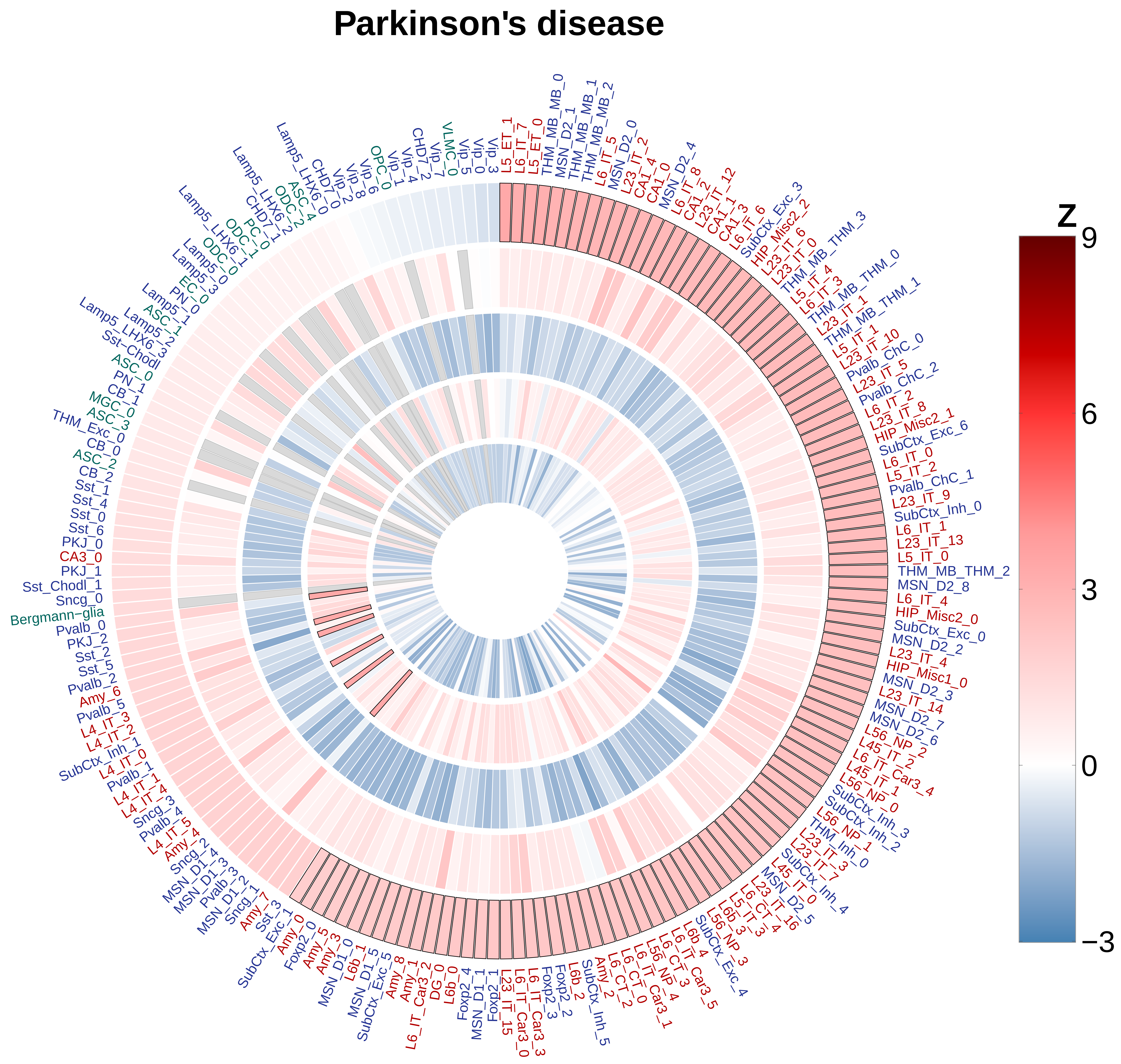
Parkinson's disease
Z
9
6
3
0
−3
L5_ET_1
L6_IT_7
L5_ET_0
THM_MB_MB_0
MSN_D2_1
THM_MB_MB_1
THM_MB_MB_2
L6_IT_5
MSN_D2_0
L23_IT_2
CA1_4
CA1_0
MSN_D2_4
L6_IT_8
CA1_2
L23_IT_12
CA1_1
CA1_3
L6_IT_6
SubCtx_Exc_3
HIP_Misc2_2
L23_IT_6
L23_IT_0
THM_MB_THM_3
L5_IT_4
L6_IT_3
THM_MB_THM_0
L23_IT_1
THM_MB_THM_1
L5_IT_1
L23_IT_10
Pvalb_ChC_0
L23_IT_5
Pvalb_ChC_2
L6_IT_2
L23_IT_8
HIP_Misc2_1
SubCtx_Exc_6
L6_IT_0
L5_IT_2
Pvalb_ChC_1
L23_IT_9
SubCtx_Inh_0
L6_IT_1
L23_IT_13
L5_IT_0
THM_MB_THM_2
MSN_D2_8
L6_IT_4
HIP_Misc2_0
SubCtx_Exc_0
MSN_D2_2
L23_IT_4
HIP_Misc1_0
MSN_D2_3
L23_IT_14
MSN_D2_7
MSN_D2_6
L56_NP_2
L45_IT_2
L6_IT_Car3_4
L45_IT_1
L56_NP_0
SubCtx_Inh_3
SubCtx_Inh_2
L56_NP_1
THM_Inh_0
L23_IT_3
L23_IT_7
SubCtx_Inh_4
L45_IT_0
MSN_D2_5
L23_IT_16
L6_CT_4
L5_IT_3
L6b_3
L56_NP_3
SubCtx_Exc_4
L6_IT_Car3_5
L6b_4
L6_CT_3
L6_IT_Car3_1
L56_NP_4
L6_CT_2
L6_CT_0
Amy_2
SubCtx_Inh_5
L6b_2
Foxp2_2
Foxp2_3
L6_IT_Car3_3
L6_IT_Car3_0
L23_IT_15
Foxp2_1
MSN_D1_1
Foxp2_4
L6b_0
DG_0
L6_IT_Car3_2
Amy_1
Amy_8
SubCtx_Exc_5
MSN_D1_5
L6b_1
MSN_D1_0
Amy_3
Amy_5
Foxp2_0
Amy_0
SubCtx_Exc_1
Sst_3
Amy_7
Sncg_1
MSN_D1_2
Pvalb_3
MSN_D1_3
MSN_D1_4
Sncg_2
Amy_4
L4_IT_5
Pvalb_4
Sncg_3
L4_IT_4
L4_IT_1
Pvalb_1
L4_IT_0
SubCtx_Inh_1
L4_IT_2
L4_IT_3
Pvalb_5
Amy_6
Pvalb_2
Sst_5
Sst_2
PKJ_2
Pvalb_0
Bergmann−glia
Sncg_0
Sst_Chodl_1
PKJ_1
CA3_0
PKJ_0
Sst_6
Sst_0
Sst_4
Sst_1
CB_2
ASC_2
CB_0
THM_Exc_0
ASC_3
MGC_0
CB_1
PN_1
ASC_0
Sst−Chodl
Lamp5_LHX6_3
Lamp5_2
ASC_1
Lamp5_1
PN_0
EC_0
Lamp5_3
Lamp5_0
ODC_0
Lamp5_LHX6_1
ODC_1
PC_0
CHD7_1
Lamp5_LHX6_2
ODC_2
ASC_4
Lamp5_LHX6_0
CHD7_0
Vip_2
Vip_8
Vip_6
OPC_0
Vip_1
Vip_4
CHD7_2
Vip_7
VLMC_0
Vip_5
Vip_0
Vip_3

### BMI

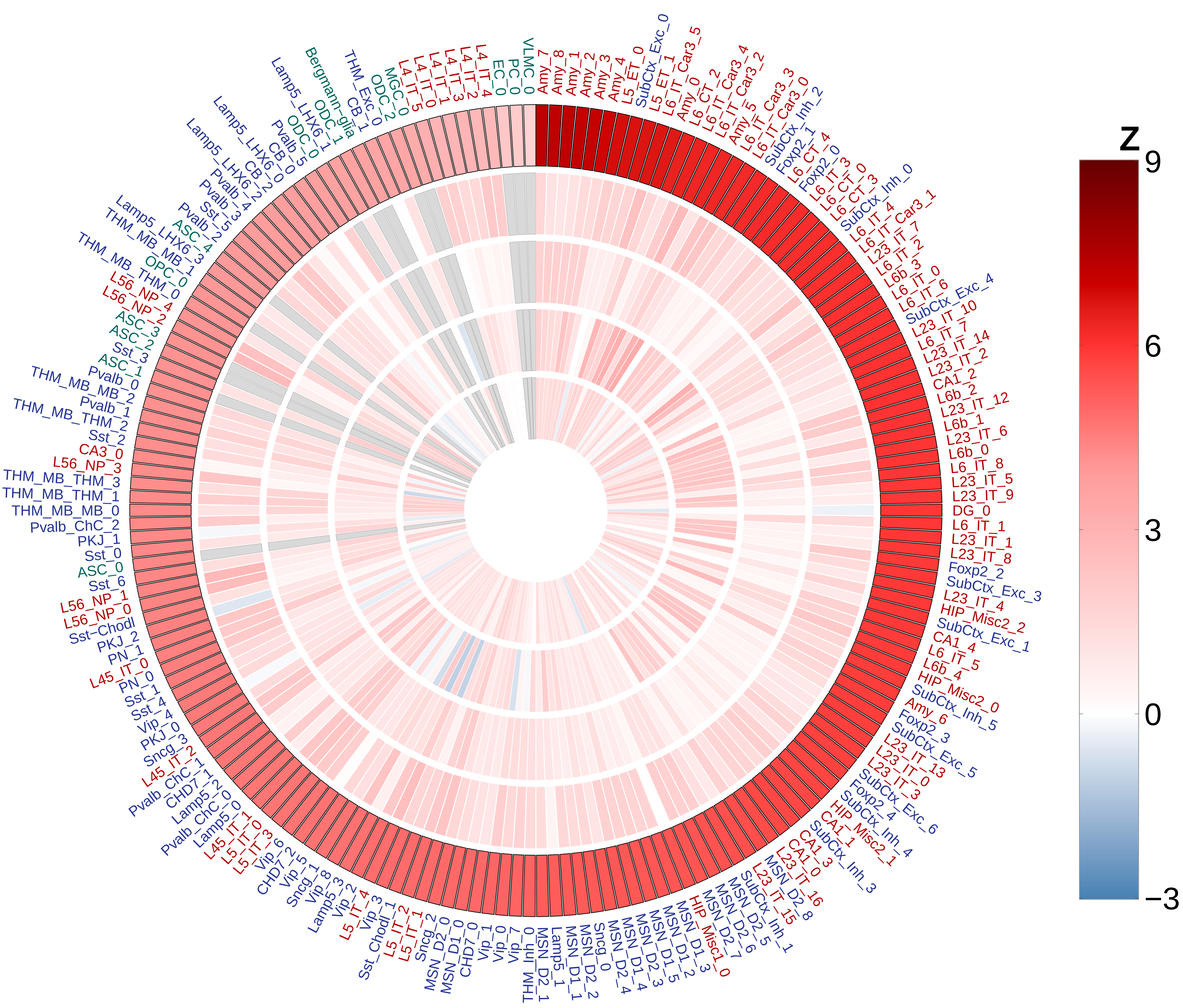
Z
9
6
3
0
−3
VLMC_0
Amy_7
Amy_8
Amy_1
Amy_2
Amy_3
Amy_4
L5_ET_0
SubCtx_Exc_0
L5_ET_1
L6_IT_Car3_5
Amy_0
L6_CT_2
L6_IT_Car3_4
L6_IT_Car3_2
Amy_5
L6_IT_Car3_3
L6_IT_Car3_0
SubCtx_Inh_2
Foxp2_1
L6_CT_4
Foxp2_0
L6_IT_3
L6_CT_0
L6_CT_3
SubCtx_Inh_0
L6_IT_4
L6_IT_Car3_1
L23_IT_7
L6_IT_2
L6b_3
L6_IT_0
L6_IT_6
SubCtx_Exc_4
L23_IT_10
L6_IT_7
L23_IT_14
L23_IT_2
CA1_2
L6b_2
L23_IT_12
L6b_1
L23_IT_6
L6b_0
L6_IT_8
L23_IT_5
L23_IT_9
DG_0
L6_IT_1
L23_IT_1
L23_IT_8
Foxp2_2
SubCtx_Exc_3
L23_IT_4
HIP_Misc2_2
SubCtx_Exc_1
CA1_4
L6_IT_5
L6b_4
HIP_Misc2_0
SubCtx_Inh_5
Amy_6
Foxp2_3
SubCtx_Exc_5
L23_IT_13
L23_IT_0
L23_IT_3
SubCtx_Exc_6
Foxp2_4
SubCtx_Inh_4
HIP_Misc2_1
CA1_1
SubCtx_Inh_3
CA1_3
CA1_0
L23_IT_16
MSN_D2_8
L23_IT_15
SubCtx_Inh_1
MSN_D2_5
MSN_D2_6
MSN_D2_7
HIP_Misc1_0
MSN_D1_3
MSN_D1_2
MSN_D1_5
MSN_D2_3
MSN_D1_4
MSN_D2_4
Sncg_0
MSN_D2_2
MSN_D1_1
Lamp5_1
MSN_D2_1
THM_Inh_0
Vip_7
Vip_0
Vip_1
CHD7_0
MSN_D1_0
MSN_D2_0
Sncg_2
L5_IT_1
L5_IT_2
Sst_Chodl_1
Vip_3
L5_IT_4
Vip_2
Lamp5_3
Vip_8
Sncg_1
Vip_5
CHD7_2
Vip_6
L5_IT_3
L5_IT_0
L45_IT_1
Lamp5_0
Pvalb_ChC_0
Lamp5_2
CHD7_1
Pvalb_ChC_1
L45_IT_2
Sncg_3
PKJ_0
Vip_4
Sst_4
Sst_1
PN_0
L45_IT_0
PN_1
PKJ_2
Sst-Chodl
L56_NP_0
L56_NP_1
Sst_6
ASC_0
Sst_0
PKJ_1
Pvalb_ChC_2
THM_MB_MB_0
THM_MB_THM_1
THM_MB_THM_3
L56_NP_3
CA3_0
Sst_2
THM_MB_THM_2
Pvalb_1
THM_MB_MB_2
Pvalb_0
ASC_1
Sst_3
ASC_2
ASC_3
L56_NP_2
L56_NP_4
THM_MB_THM_0
OPC_1
OPC_0
THM_MB_MB_1
Lamp5_LHX6_3
ASC_4
Pvalb_2
Sst_5
Pvalb_3
Pvalb_4
CB_2
Lamp5_LHX6_2
Lamp5_LHX6_0
CB_0
Pvalb_5
ODC_0
Lamp5_LHX6_1
ODC_1
Bergmann-glia
CB_1
THM_Exc_0
ODC_2
MGC_0
L4_IT_5
L4_IT_0
L4_IT_1
L4_IT_3
L4_IT_2
L4_IT_4
EC_0
PC_0

### Age of smoking

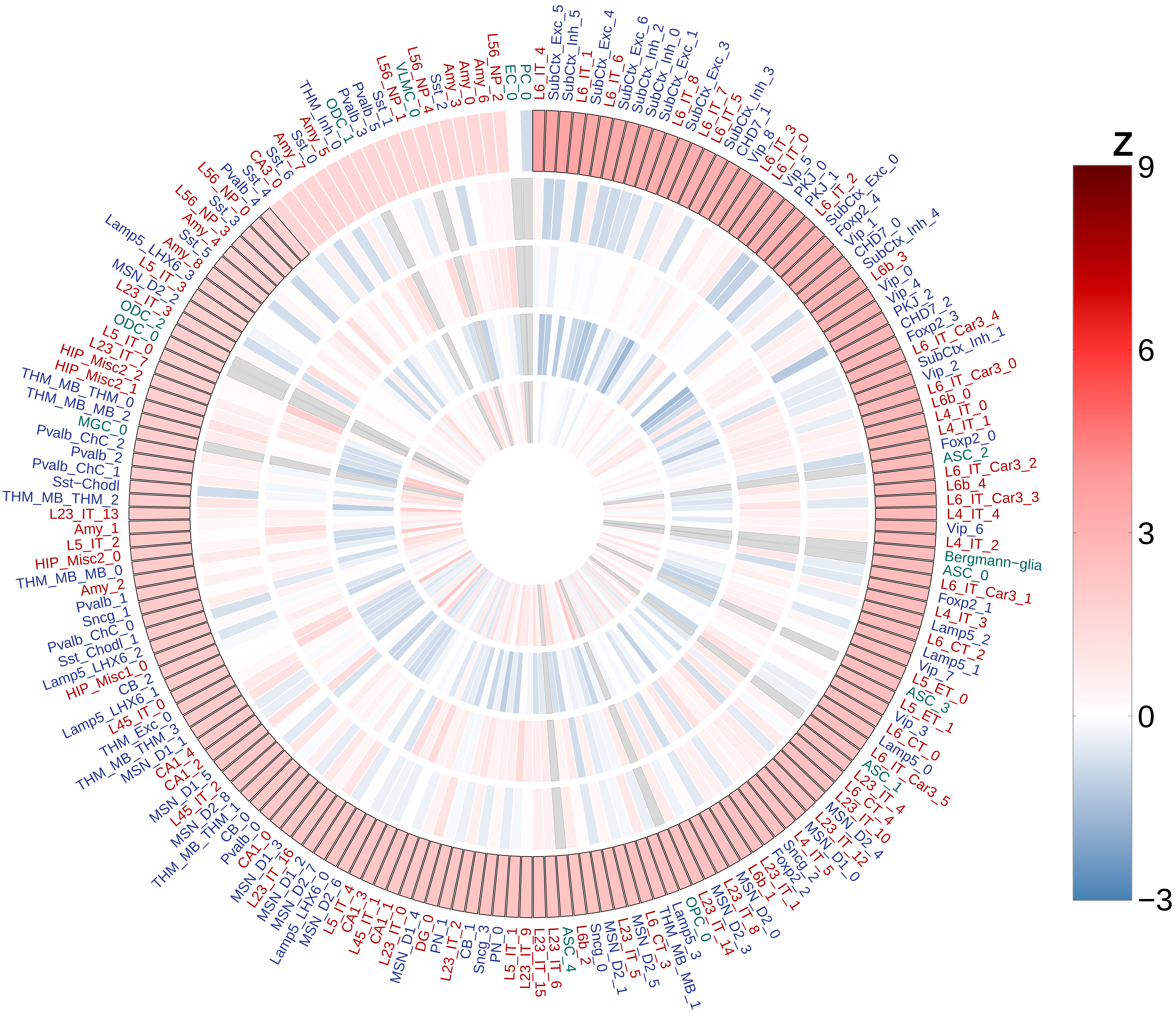

### Cigarettes per day

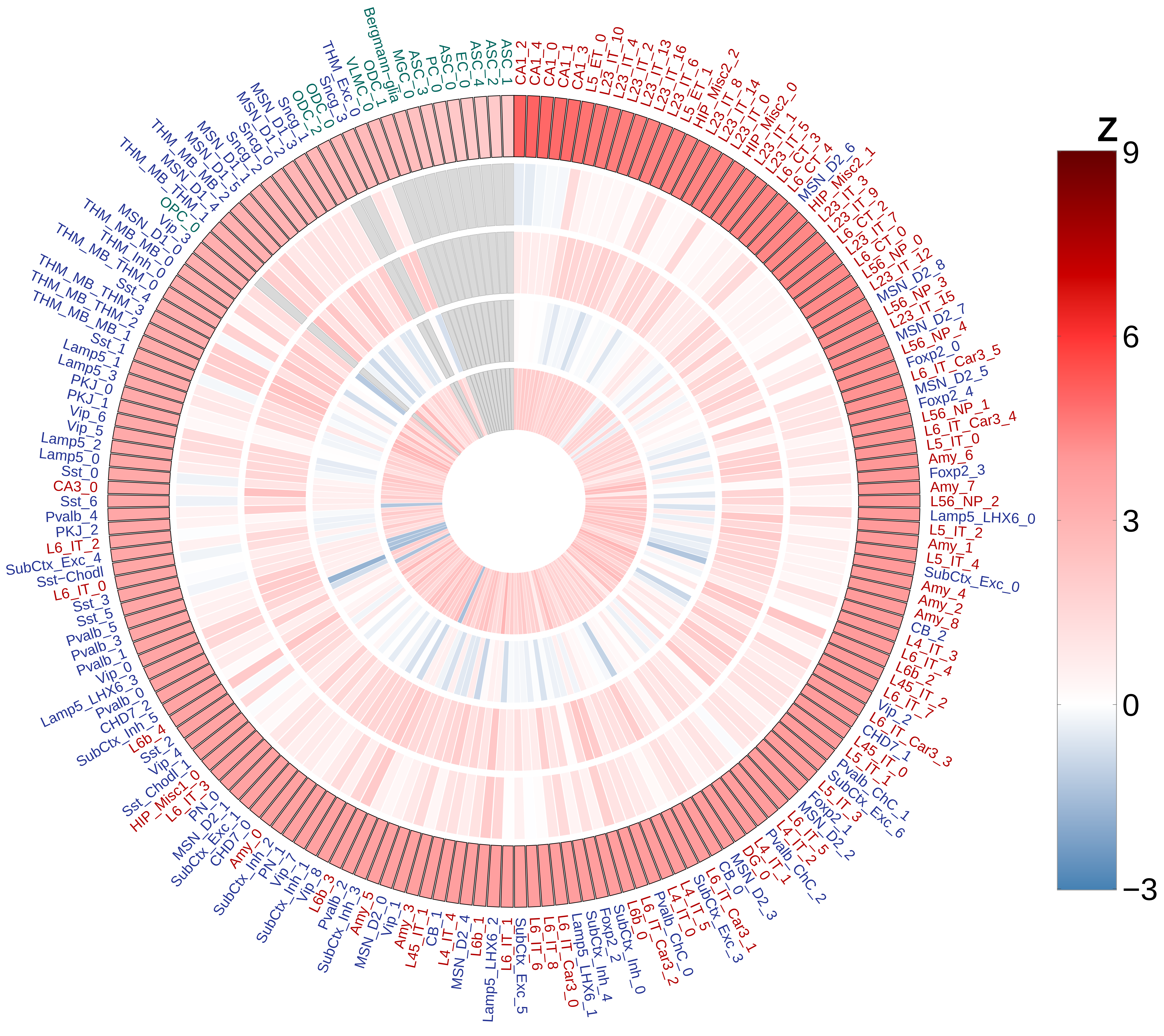

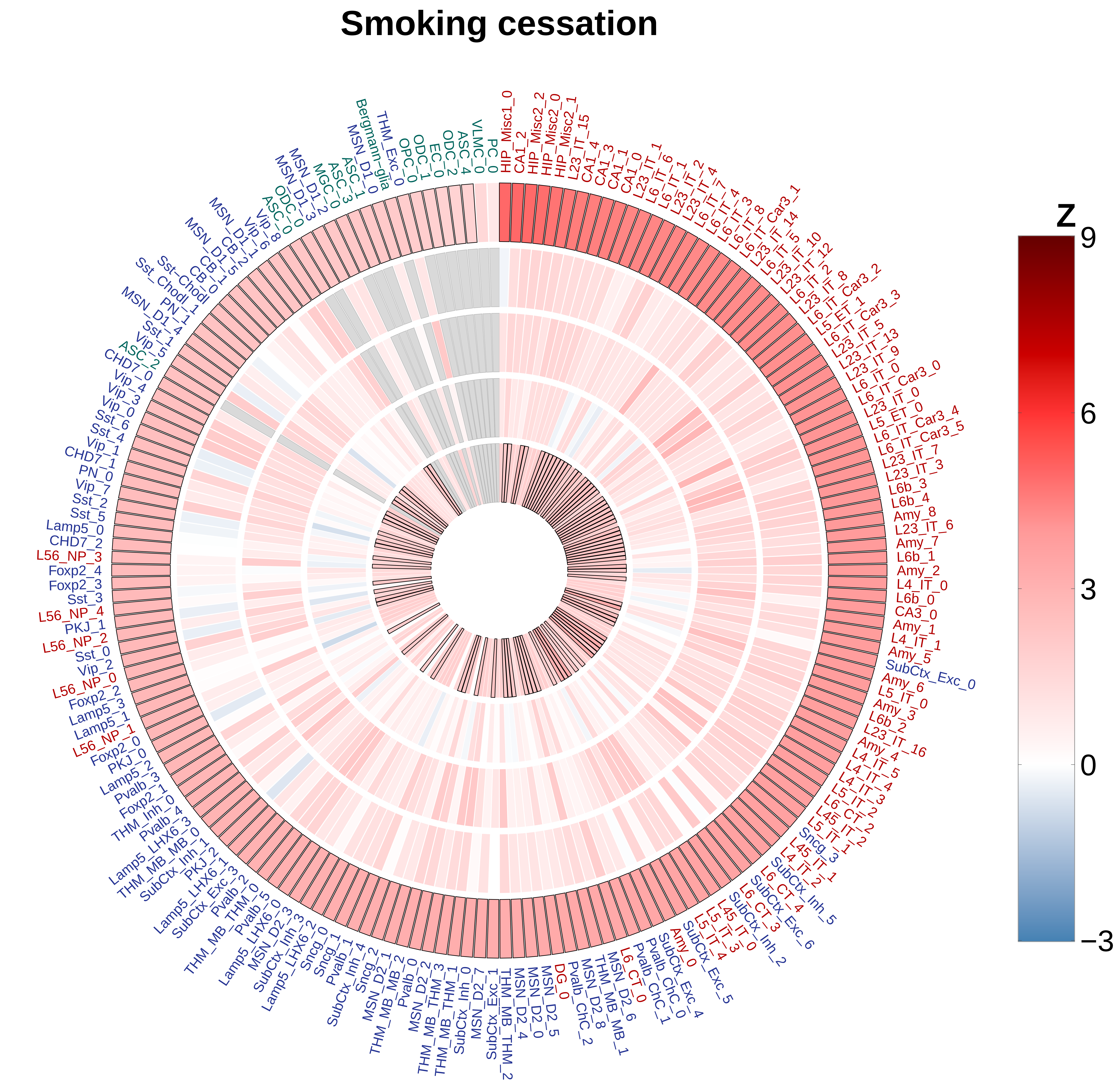
Smoking cessation
HIP_Misc1_0
CA1_2
HIP_Misc2_2
HIP_Misc2_0
HIP_Misc2_1
L23_IT_15
CA1_4
CA1_3
CA1_1
CA1_0
L23_IT_1
L6_IT_6
L6_IT_1
L23_IT_2
L23_IT_4
L6_IT_7
L6_IT_4
L6_IT_3
L6_IT_8
L6_IT_Car3_1
L23_IT_14
L6_IT_5
L23_IT_10
L23_IT_12
L6_IT_2
L23_IT_8
L6_IT_Car3_2
L5_ET_1
L6_IT_Car3_3
L23_IT_5
L23_IT_13
L23_IT_9
L6_IT_0
L6_IT_Car3_0
L23_IT_0
L5_ET_0
L6_IT_Car3_4
L6_IT_Car3_5
L23_IT_7
L23_IT_3
L6b_3
L6b_4
Amy_8
L23_IT_6
Amy_7
L6b_1
Amy_2
L4_IT_0
L6b_0
CA3_0
Amy_1
L4_IT_1
Amy_5
SubCtx_Exc_0
Amy_6
L5_IT_0
Amy_3
L6b_2
L23_IT_16
Amy_4
Sncg_3
SubCtx_Inh_5
L6_CT_4
SubCtx_Exc_6
L6_CT_3
SubCtx_Inh_2
SubCtx_Exc_5
SubCtx_Exc_4
Amy_0
Pvalb_ChC_0
Pvalb_ChC_1
MSN_D2_6
THM_MB_MB_1
MSN_D2_8
Pvalb_ChC_2
DG_0
MSN_D2_5
MSN_D2_0
MSN_D2_4
THM_MB_THM_2
SubCtx_Exc_1
MSN_D2_7
SubCtx_Inh_0
THM_MB_THM_1
THM_MB_THM_3
MSN_D2_2
Pvalb_0
THM_MB_MB_2
MSN_D2_1
Sncg_2
SubCtx_Inh_4
Pvalb_1
Sncg_1
Sncg_0
Lamp5_LHX6_2
SubCtx_Inh_3
MSN_D2_3
Lamp5_LHX6_0
Pvalb_5
THM_MB_THM_0
Pvalb_2
SubCtx_Exc_3
Lamp5_LHX6_1
PKJ_2
SubCtx_Inh_1
THM_MB_MB_0
Lamp5_LHX6_3
Pvalb_4
THM_Inh_0
Foxp2_1
Pvalb_3
Lamp5_2
PKJ_0
Foxp2_0
L56_NP_1
Lamp5_1
Lamp5_3
Foxp2_2
L56_NP_0
Vip_2
Sst_0
L56_NP_2
PKJ_1
L56_NP_4
Sst_3
Foxp2_3
Foxp2_4
L56_NP_3
CHD7_2
Lamp5_0
Sst_5
Sst_2
Vip_7
PN_0
CHD7_1
Vip_1
Sst_4
Sst_6
Vip_0
Vip_3
Vip_4
CHD7_0
ASC_2
Vip_5
Sst_1
PN_1
Sst_Chodl_1
Sst_Chodl_0
CB_1
CB_0
Vip_6
Vip_8
ASC_0
ODC_0
MSN_D1_3
MSN_D1_2
MGC_0
ASC_3
ASC_1
MSN_D1_0
Bergmann-glia
THM_Exc_0
OPC_0
ODC_1
EC_0
ODC_2
ASC_4
VLMC_0
PC_0
Z
9
6
3
0
−3

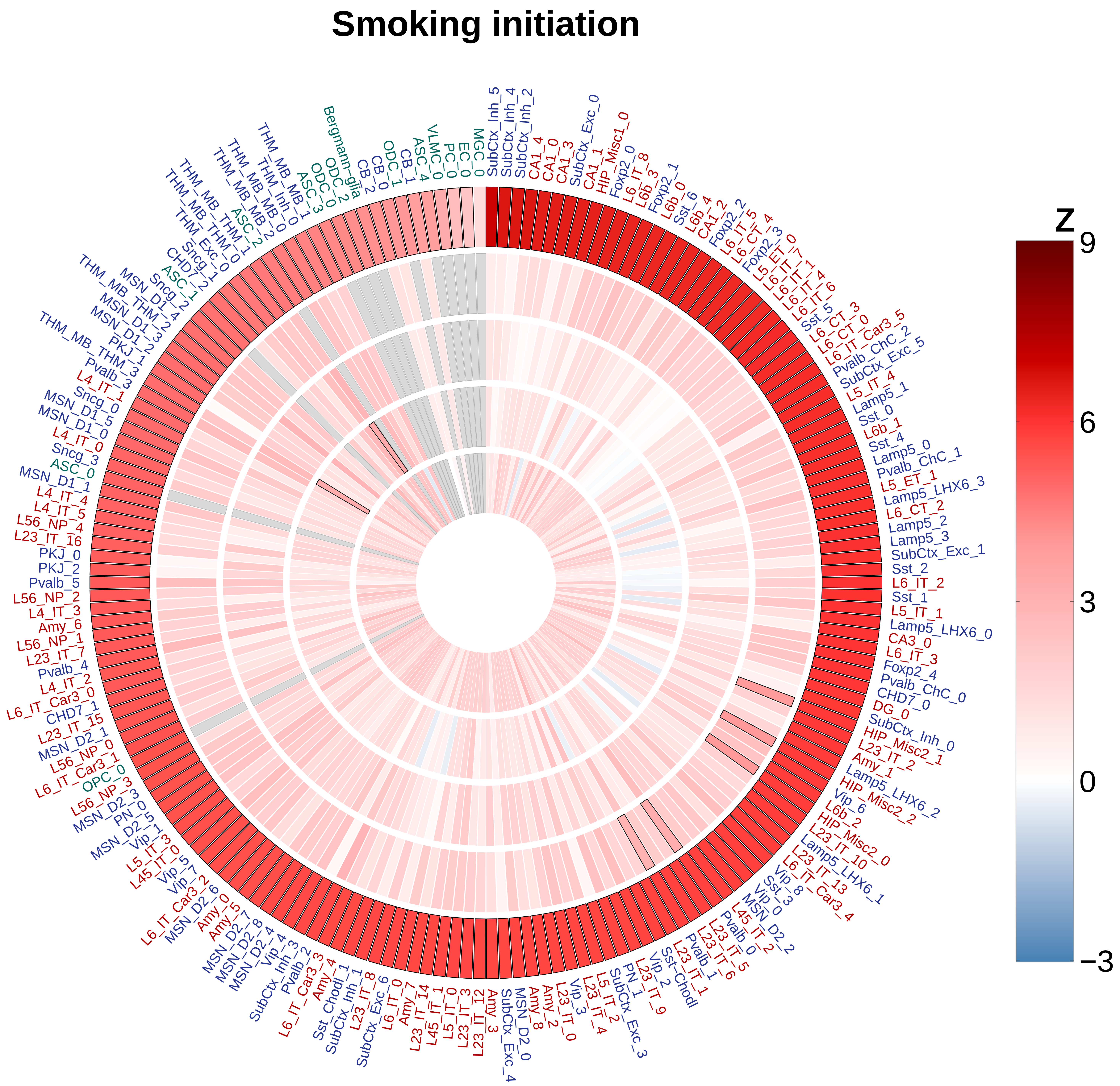

Smoking initiation
Z
9
6
3
0
−3
SubCtx_Inh_5
SubCtx_Inh_4
SubCtx_Inh_2
CA1_4
CA1_0
CA1_3
SubCtx_Exc_0
CA1_1
HIP_Misc1_0
Foxp2_0
L6_IT_8
L6b_3
Foxp2_1
L6b_0
Sst_6
L6b_4
CA1_2
Foxp2_2
L6_IT_5
L6_CT_4
Foxp2_3
L5_ET_0
L6_IT_7
L6_IT_1
L6_IT_4
L6_IT_6
Sst_5
L6_CT_3
L6_CT_0
L6_IT_Car3_5
Pvalb_ChC_2
SubCtx_Exc_5
L5_IT_4
Lamp5_1
Sst_0
L6b_1
Sst_4
Lamp5_0
Pvalb_ChC_1
L5_ET_1
Lamp5_LHX6_3
L6_CT_2
Lamp5_2
Lamp5_3
SubCtx_Exc_1
Sst_2
L6_IT_2
Sst_1
L5_IT_1
Lamp5_LHX6_0
CA3_0
L6_IT_3
Foxp2_4
Pvalb_ChC_0
CHD7_0
DG_0
SubCtx_Inh_0
HIP_Misc2_1
L23_IT_2
Amy_1
Lamp5_LHX6_2
HIP_Misc2_2
Vip_6
L6b_2
HIP_Misc2_0
L23_IT_10
Lamp5_LHX6_1
L23_IT_13
L6_IT_Car3_4
Vip_8
Sst_3
Vip_0
MSN_D2_2
L45_IT_2
Pvalb_0
L23_IT_5
L23_IT_6
Pvalb_1
L23_IT_1
Sst-Chodl
Vip_2
L23_IT_9
PN_1
SubCtx_Exc_3
L5_IT_2
L23_IT_4
Vip_3
L23_IT_0
Amy_2
Amy_8
MSN_D2_0
SubCtx_Exc_4
Amy_3
L23_IT_12
L23_IT_3
L5_IT_0
L45_IT_1
L23_IT_14
Amy_7
L6_IT_0
SubCtx_Exc_6
L23_IT_8
SubCtx_Inh_1
Sst_Chodl_1
Amy_4
L6_IT_Car3_3
Pvalb_2
SubCtx_Inh_3
Vip_4
MSN_D2_4
MSN_D2_8
MSN_D2_7
Amy_5
Amy_0
MSN_D2_6
L6_IT_Car3_2
Vip_7
Vip_5
L45_IT_0
L5_IT_3
Vip_1
MSN_D2_5
PN_0
MSN_D2_3
L56_NP_3
OPC_0
L6_IT_Car3_1
L56_NP_0
MSN_D2_1
L23_IT_15
CHD7_1
L6_IT_Car3_0
L4_IT_2
Pvalb_4
L23_IT_7
L56_NP_1
Amy_6
L4_IT_3
L56_NP_2
Pvalb_5
PKJ_2
PKJ_0
L23_IT_16
L56_NP_4
L4_IT_5
L4_IT_4
MSN_D1_1
ASC_0
Sncg_3
L4_IT_0
MSN_D1_0
MSN_D1_5
Sncg_0
L4_IT_1
Pvalb_3
THM_MB_THM_3
PKJ_1
MSN_D1_2
MSN_D1_3
THM_MB_THM_2
MSN_D1_4
Sncg_2
ASC_1
CHD7_2
Sncg_1
THM_Exc_0
THM_MB_THM_0
THM_MB_THM_1
ASC_2
THM_MB_MB_2
THM_MB_MB_0
THM_Inh_0
THM_MB_MB_1
ASC_3
ODC_0
ODC_2
Bergmann-glia
CB_2
CB_0
ODC_1
CB_1
ASC_4
VLMC_0
PC_0
EC_0
MGC_0

### Drinks per week

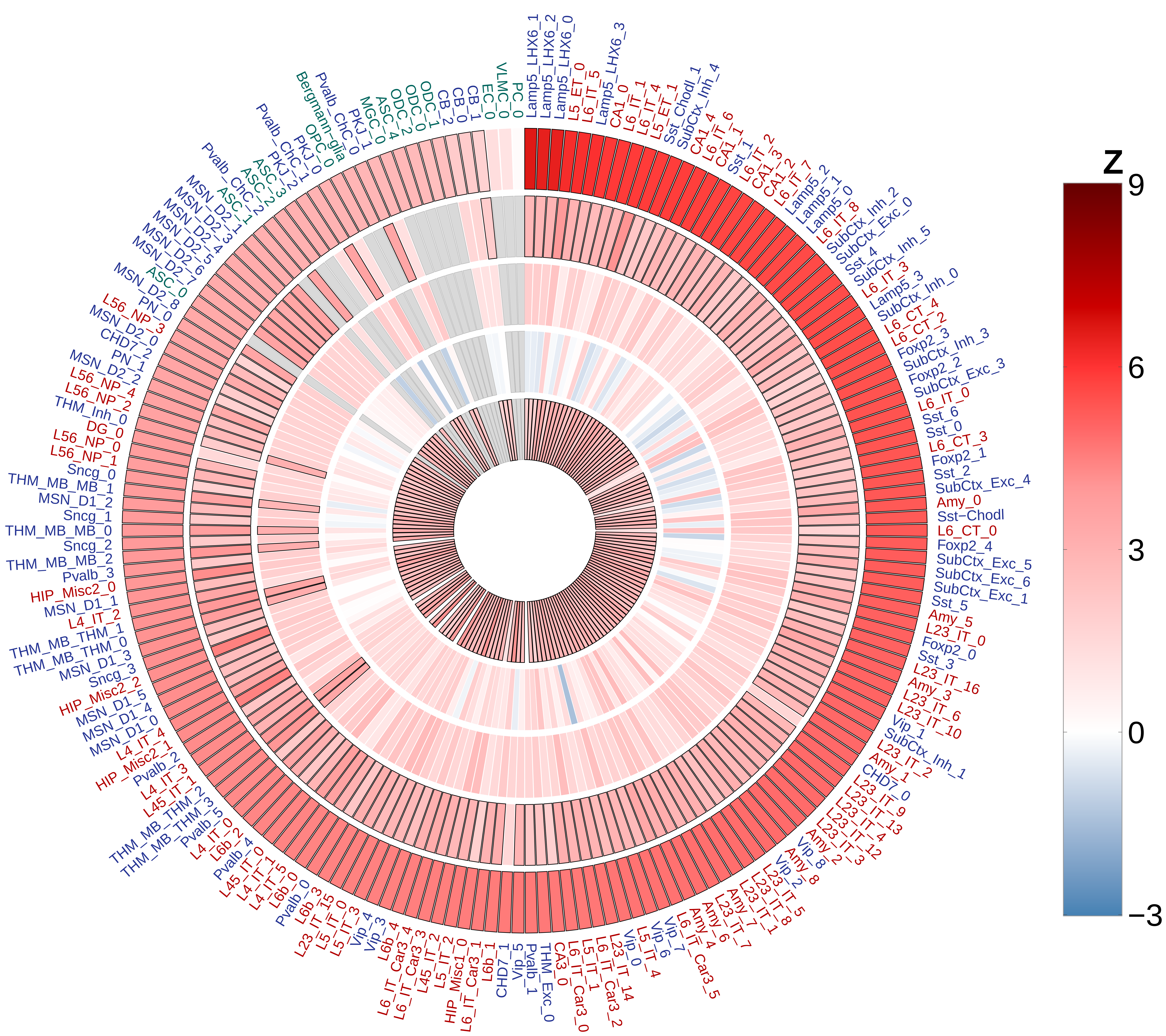

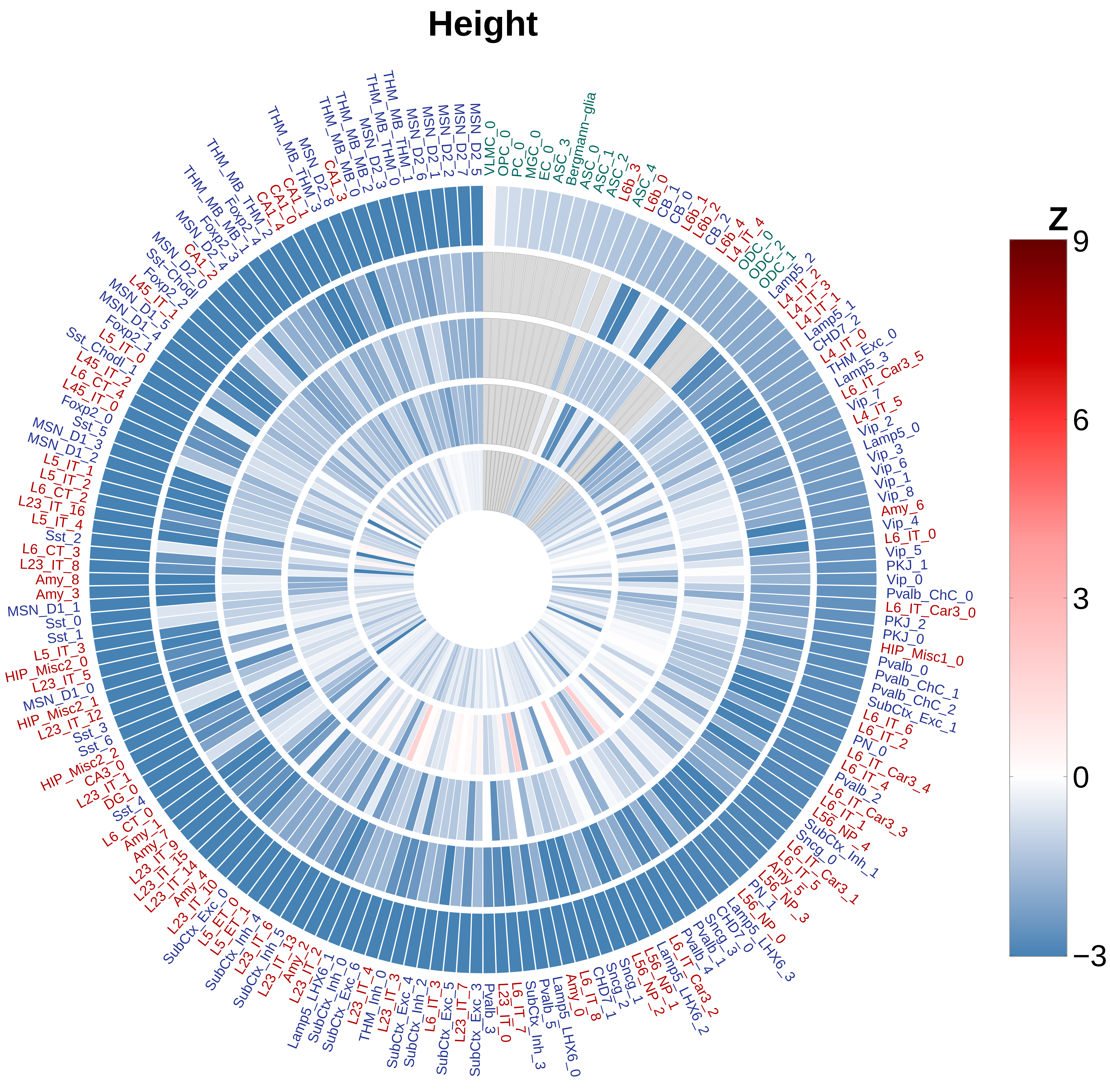
Height
Z
9
6
3
0
−3
VLMC_0
OPC_0
PC_0
MGC_0
EC_0
ASC_3
Bergmann−glia
ASC_0
ASC_1
ASC_2
L6b_3
ASC_4
L6b_0
CB_1
CB_0
L6b_1
L6b_2
CB_2
L6b_4
L4_IT_4
ODC_0
ODC_2
ODC_1
Lamp5_2
L4_IT_2
L4_IT_3
L4_IT_1
Lamp5_1
CHD7_2
L4_IT_0
THM_Exc_0
Lamp5_3
L6_IT_Car3_5
Vip_7
L4_IT_5
Vip_2
Lamp5_0
Vip_3
Vip_6
Vip_1
Vip_8
Amy_6
Vip_4
L6_IT_0
Vip_5
PKJ_1
Vip_0
Pvalb_ChC_0
L6_IT_Car3_0
PKJ_2
PKJ_0
HIP_Misc1_0
Pvalb_0
Pvalb_ChC_1
Pvalb_ChC_2
SubCtx_Exc_1
L6_IT_6
L6_IT_2
PN_0
L6_IT_Car3_4
L6_IT_4
Pvalb_2
L6_IT_Car3_3
L6_IT_1
L56_NP_4
SubCtx_Inh_1
Sncg_0
L6_IT_Car3_1
L6_IT_5
Amy_5
L56_NP_3
PN_1
L56_NP_0
Lamp5_LHX6_3
CHD7_0
Sncg_3
Pvalb_1
Pvalb_4
L6_IT_Car3_2
Lamp5_LHX6_2
L56_NP_1
L56_NP_2
Sncg_1
Sncg_2
CHD7_1
L6_IT_8
Amy_0
Lamp5_LHX6_0
Pvalb_5
SubCtx_Inh_3
L6_IT_7
L23_IT_0
Pvalb_3
SubCtx_Exc_3
L23_IT_7
SubCtx_Exc_5
L6_IT_3
SubCtx_Inh_2
SubCtx_Exc_4
L23_IT_3
THM_Inh_0
L23_IT_4
SubCtx_Exc_6
SubCtx_Inh_0
Lamp5_LHX6_1
L23_IT_2
Amy_2
L23_IT_13
SubCtx_Inh_5
L23_IT_6
SubCtx_Inh_4
L5_ET_1
L5_ET_0
SubCtx_Exc_0
L23_IT_10
Amy_4
L23_IT_14
L23_IT_15
L23_IT_9
Amy_7
Amy_1
L6_CT_0
Sst_4
DG_0
L23_IT_1
CA3_0
HIP_Misc2_2
Sst_6
Sst_3
L23_IT_12
HIP_Misc2_1
MSN_D1_0
L23_IT_5
HIP_Misc2_0
L5_IT_3
Sst_1
Sst_0
MSN_D1_1
Amy_3
Amy_8
L23_IT_8
L6_CT_3
Sst_2
L5_IT_4
L23_IT_16
L6_CT_2
L5_IT_2
L5_IT_1
MSN_D1_2
MSN_D1_3
Sst_5
Foxp2_0
L45_IT_0
L6_CT_4
L45_IT_2
L6_CT_1
Sst_Chodl_1
L5_IT_0
Foxp2_1
MSN_D1_4
MSN_D1_5
L45_IT_1
Foxp2_2
Sst−Chodl
MSN_D2_0
CA1_2
MSN_D2_4
Foxp2_3
THM_MB_MB_1
Foxp2_4
THM_MB_THM_2
CA1_4
CA1_0
CA1_1
THM_MB_THM_3
MSN_D2_8
CA1_3
THM_MB_MB_0
THM_MB_MB_2
MSN_D2_3
THM_MB_THM_0
THM_MB_THM_1
MSN_D2_6
MSN_D2_1
MSN_D2_2
MSN_D2_7
MSN_D2_5
